# Field dependent deep learning enables high-throughput whole-cell 3D super-resolution imaging

**DOI:** 10.1101/2022.10.14.512179

**Authors:** Shuang Fu, Wei Shi, Tingdan Luo, Yingchuan He, Lulu Zhou, Jie Yang, Zhichao Yang, Jiadong Liu, Xiaotian Liu, Zhiyong Guo, Chengyu Yang, Chao Liu, Zhen-li Huang, Jonas Ries, Mingjie Zhang, Peng Xi, Dayong Jin, Yiming Li

**Affiliations:** Guangdong Provincial Key Laboratory of Advanced Biomaterials, Department of Biomedical Engineering, Southern University of Science and Technology, Shenzhen 518055, China; School of Life Sciences, Southern University of Science and Technology, Shenzhen 518055, China; Key Laboratory of Biomedical Engineering of Hainan Province, School of Biomedical Engineering, Hainan University, Haikou 570228, China; European Molecular Biology Laboratory, Cell Biology and Biophysics, Heidelberg 69117, Germany; Department of Biomedical Engineering, College of Future Technology, Peking University, Beijing 100871, China; Institute for Biomedical Materials and Devices (IBMD), Faculty of Science, University of Technology Sydney, NSW 2007, Australia

**Author notes:** These authors contributed equally: Shuang Fu, Wei Shi. Correspondence should be addressed to Y.L.

## Abstract

Single-molecule localization microscopy (SMLM) in a typical wide-field setup has been widely used for investigating sub-cellular structures with super resolution. However, field-dependent aberrations restrict the field of view (FOV) to only few tens of micrometers. Here, we present a deep learning method for precise localization of spatially variant point emitters (FD-DeepLoc) over a large FOV covering the full chip of a modern sCMOS camera. Using a graphic processing unit (GPU) based vectorial PSF fitter, we can fast and accurately model the spatially variant point spread function (PSF) of a high numerical aperture (NA) objective in the entire FOV. Combined with deformable mirror based optimal PSF engineering, we demonstrate high-accuracy 3D SMLM over a volume of ~180 × 180 × 5 μm^3^, allowing us to image mitochondria and nuclear pore complex in the entire cells in a single imaging cycle without hardware scanning - a 100-fold increase in throughput compared to the state-of-the-art.

## Introduction

While fluorescence microscopy with high contrast and super resolution has revolutionized structural cell biology studies^1^, high-throughput imaging with rich information content has enabled quantitative biology research^2^. The trend has been pointed to developing high-throughput super-resolution imaging techniques for high content screening^3–5^. The typical strategy is for automated microscope to acquire small field of view (FOV) images one by one^6^ and generate a mosaic image by post processing^7^. However, hardware automation is often not available in the typical microscopes, and some biological samples are not suitable to be volumetrically scanned. Moreover, it is not easy to stitch the super resolution images with accuracy comparable to its high spatial resolution^7^. As most biological samples contain rich structural information in three dimensions, it becomes particularly challenging to obtain the whole-cell-scale 3D single-molecule-resolution image at high throughput.

As a wide-field super-resolution imaging technique, single molecule localization microscopy (SMLM) has the potential to increase the imaging throughput by just using a modern sCMOS camera with ultra-high number of pixels on a large chip without compromising the imaging speed or spatial resolution. Through homogeneous illumination, FOVs as large as 221 μm × 221 μm ^8^ can be achieved using waveguides^9^, multimode fibers^8^, microlens arrays^10^ or widefield illumination scanning^11^. However, 3D super-resolution imaging across large FOV to utilize the full camera chip remains difficult. We believe the main reason is that optical aberrations become more pronounced at the margins of the large FOV, leading to the degraded 3D imaging quality.

Conventional 3D SMLM analysis requires spatially invariant 3D point spread function (PSF) to achieve accurate fitting of single molecules. However, it is difficult to correct the aberrations if the emitters are far away from the central optical axis^12^, which is particularly true for the high NA objective lenses^13,14^. Efforts include the use of a regularly spaced nanohole array to build an approximated PSF model on the spatially variant of localization bias^13^, and the recent introduction of vectorial PSF theory by rendering the field-dependent PSFs with a group of Zernike coefficients^14^. The later has the potential to increase the accuracy, but it requires computational resources, *e.g*. costing ~2 min per localization for data analysis^14^. Therefore, such methods are difficult to use in practice, especially for high throughput imaging with large datasets.

In recent years, deep learning has proved to be a powerful tool in SMLM^15–18^, especially for conditions with highly overlapping molecules. Examples include high density emitters^16,17^, classifying colors^18^, optimizing engineered PSFs^16,19^, which are difficult to solve using conventional model fitting methods. As the image formation of single molecule signals by a microscope is well understood, a realistic PSF model incorporating aberrations can be simulated using established mathematical models with high accuracy^15^. This allows generation of virtually unlimited training data and helps the localization accuracy of the neural network achieve Cramér -Rao lower bound (CRLB), the theoretical minimum uncertainty^17,20^. However, conventional convolution neural networks (CNNs) are spatially invariant and can only poorly learn spatial information^21^. They perform well for object recognition when the objects are decoupled from their spatial coordinates. Optical aberrations are often static and highly correlated to their positions. As a result, the pattern of point emitters is field dependent. For large FOV imaging, a spatially sensitive neural network is needed.

To overcome the field-dependent aberrations, we present FD-DeepLoc (**Fig. 1**), which combines both fast spatially variant PSF modeling and deep learning based single molecule localization to achieve 3D super resolution over a large FOV and depth of field (DOF). Based on a GPU-accelerated vectorial PSF fitter, it can quickly model the spatially variant PSF by Zernike based aberration maps over a large FOV (**Fig. 1a**). The spatially variant PSF model is then used as a training-data generator for deep learning-based 3D single molecule localization (**Fig. 1c**). To enable CNNs to learn the spatially variant single molecule patterns, FD-DeepLoc incorporates CoordConv^22^ channels to the network architecture to encode the spatial context to the conventional CNNs which is otherwise spatially invariant. Moreover, we introduce a small aberration variation to the trained PSF models, which makes the network more robust when the theoretical and experimental PSF models are mismatched. With the accelerated PSF modeling and optimized CNNs, we demonstrate on various biological structures that FD-DeepLoc enables 3D whole cell super-resolution imaging across a large FOV with high fidelity, allowing quantitative analysis of super resolution images using larger sample set.

**Fig. 1.**
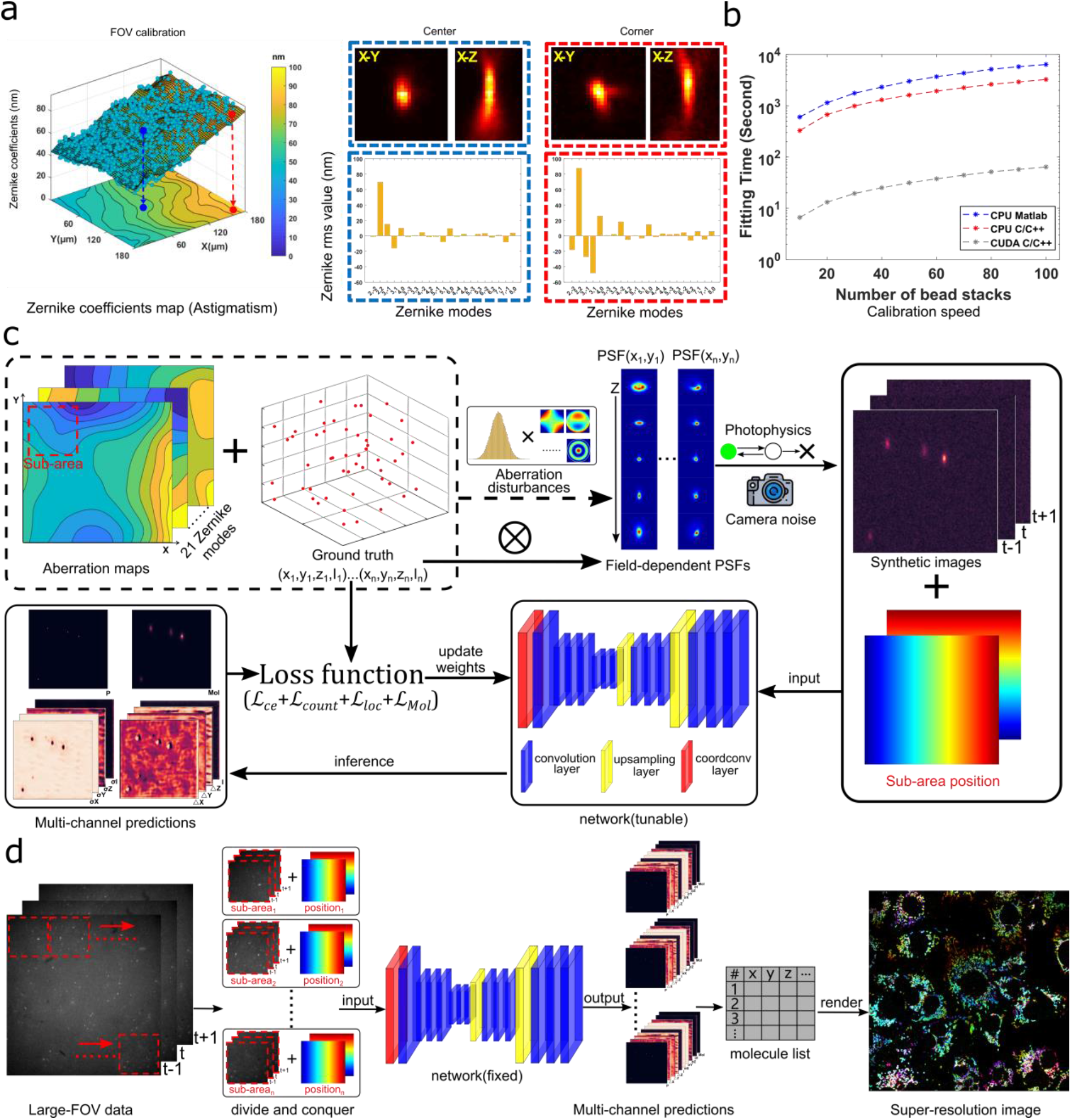
Schematic of FD-DeepLoc. **a**, Field-dependent aberration calibration. Thousands of through-focus bead stacks sampling the entire FOV are fitted to a vectorial PSF model with a GPU based fitter to retrieve the field-dependent Zernike aberrations. The left panel shows an aberration map for astigmatism. The two right panels are the fitted Zernike coefficients for two bead stacks at the center and corner positions, respectively. **b**, Fitting speed of vectorial PSF fitter implemented in different programming languages running with CPU and GPU individually. The bead stack used for evaluation has a ROI size of 27×27×41 pixels with a PC equipped with an Intel Core i9-9900 processor of 128GB RAM clocked at 3.50GHz and an NVIDIA GeForce GTX 3090 graphics card with 24.0 GB memory. **c**, FD-DeepLoc training process. The training dataset is generated by the spatially variant vectorial PSF model using the calibrated aberration maps with a small variant to compensate for the models mismatch between theoretical and experimental PSF models. For each single molecule, the coordinate is randomly placed in the FOV. Three consecutive images of a sub areas (usually 128×128 pixels) with corresponding coordinate channels are used in each training unit. **d**, FD-DeepLoc inference process. Full frame images are divided into multiple sub-areas which are sequentially fed into the network with the corresponding global position. The output of the network are ten prediction channels with the same size as the input image, each corresponds to an estimated parameter (probability 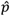, positions 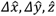, intensity 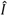, and uncertainties *σ_x_, σ_y_, σ_z_, σ_I_* and single molecule image estimation 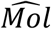). A molecule list with uncertainty estimation of parameters is generated from these channels. We finally used SMAP^31^ to render the super-resolution images and ViSP^41^ to generate the reconstructed 3D super-resolution movies from the molecule list.

## Results

### GPU based fast and accurate spatially variant vectorial PSF modeling

To accurately model field dependent aberrations, we employed a GPU based vectorial PSF fitter to fit thousands of through-focus bead stacks evenly illuminated by a multimode fiber and randomly distributed in the entire FOV using maximum likelihood estimation (MLE, **Extended Data Fig. 1**). The pupil function of each bead was decomposed into Zernike polynomials whose coefficients were then interpolated to generate spatially variant aberration maps across the entire FOV (**Fig. 2 and Supplementary Fig. 1**). Although the vectorial PSF could model the experimental PSF of a high NA objective with high accuracy, it is still not widely used for single molecule localization^23,24^. A key limitation for direct vectorial PSF fit is its great computational burden due to the complex functions involved in the Debye integral^25^ (**Supplementary Note 1**). Here, we overcome the computational challenge of vectorial PSF model fit in three ways. First, we implement the vectorial PSF fitter in CUDA C/C++ and execute it on the GPU, so that a fast modeling of the spatially variant PSF models over a large FOV is achieved. Running on an NVIDIA GeForce RTX™ 3090 graphics card, our implementation amounts to ~50 fold speed advantage over a similar implementation running on an Intel^®^ Core™ i9-9900 processor (**Fig. 1b**). Second, we integrated globLoc with flexible parameter sharing^26^ functionality to our vectorial PSF fitter which could link parameters (e.g. aberrations, *xyz* positions, photons, background) among different single molecule images. Therefore, the fitter could account for the photon bleaching and system drift during the acquisition of the bead stacks (**Methods**). The fitted PSF model agrees well with the experimental PSF model across the entire FOV (~180 μm × 180 μm) at different *z* positions (**Extended Data Fig. 2, Supplementary Fig. 2**). Third, we developed a GPU based vectorial PSF simulator for the neural network so that online simulation and training of the neural network could be achieved (**Supplementary Fig. 3**). The whole GPU based vectorial PSF calibration, fitting and simulation pipeline enabled fast modeling and training of the neural network for analyzing the single molecule data in a large FOV.

**Fig. 2.**
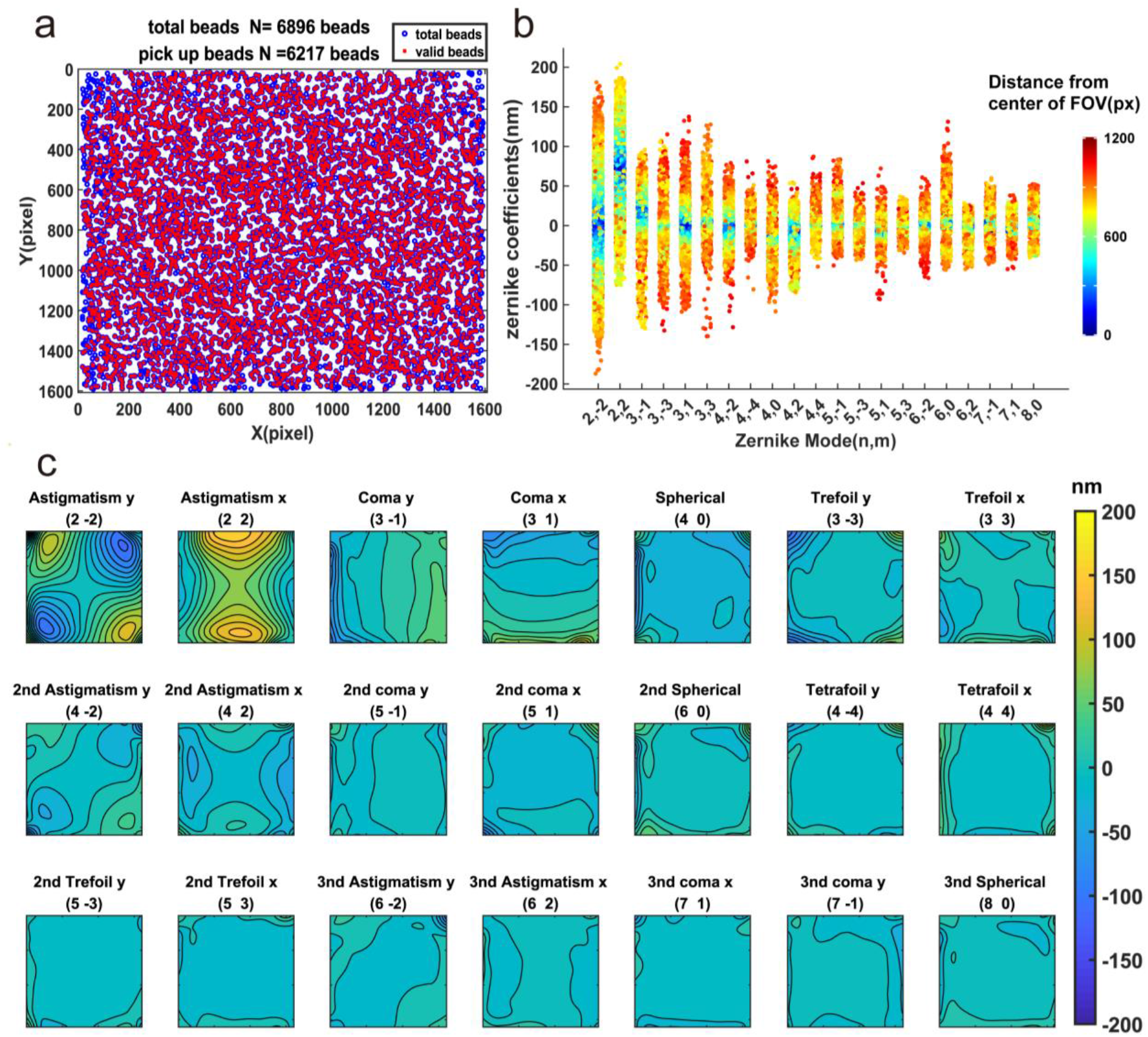
Field-dependent aberration maps for an astigmatic PSF induced by a cylindrical lens. **a**, The distribution of the beads used for aberration map calibration (summed from 50 sets of beads stacks). Blue circles denote all beads collected and red crosses denote the beads after filtering. **b**, The relationship between aberration magnitude and the distance to center of FOV. Color denotes the distance to the FOV center. **c**, The interpolated 21 Zernike aberration maps of our microscope after inserting a cylindrical lens to the system. The contour interval is 16 nm.

### Features of the FD-DeepLoc network

CNNs use convolution kernels to detect features within an image and have been successfully applied in SMLM to localize molecules with precision as high as CRLB^17,20^. In conventional CNNs, such as our previously published DECODE network^17^, all feature patterns in the images share the same convolution kernels. As a result, the detected features are spatially invariant to their local positions. To enable the CNN to precisely localize the field dependent single molecule patterns, we introduced two additional coordinate channels (*x* and *y*) to the DECODE input. The essences of the coordinate channels are the pixel-wise *x*/*y* positions (**Fig. 1c**). Therefore, the position is explicitly modelled and can have an influence on the training and inference performance (CoordConv^22^, **Fig. 1c and d** and **Extended Data Fig. 3**). We then combined CoordConv with a conventional spatially invariant CNNs (DECODE^17^) and enabled FD-DeepLoc to precisely determine both what feature it reads and where it is positioned in the FOV. To adapt to the large FOV, which is memory consuming during training, we sample a sub-area (usually 128×128 pixels) of the entire single molecule image to the network. The global position of each sub-area was encoded in the coordinate channels. Therefore, the network could retrieve the field dependent feature pattern of each single molecule within the sub-area using the local coordinate information and corresponding simulated PSFs based on the calibrated aberration map of entire FOV.

Normally, deep learning methods perform well when the training data accurately resembles the experimental data. However, experimental PSFs are often slightly different between different samples. Such a model mismatch between training and experimental data results in localization errors. To illustrate this problem, we simulated three datasets with different aberrations of various amplitudes: 70 nm astigmatism; 70 nm astigmatism plus 20 nm spherical aberration; 70 nm astigmatism plus 20 nm spherical aberration plus 20 nm coma (**Extended Data Fig. 4**). The network was trained using only 70 nm astigmatism. As shown in **Extended Data Fig. 4a**, DECODE performs well when the PSF models between training and test datasets are matched. However, if a little spherical aberration is added to the test dataset, the network predicts large artifacts, especially at imaging planes away from focus (**Extended Data Fig. 4d and e**). The artifacts became more severe when we further added the coma aberration (**Extended Data Fig. 4g and h**). We suspect that this is due to the PSF model mismatch. We then gave more freedom to the training data generator by adding small random disturbances (normal distribution with zero mean and 0.06 rad standard deviation) to all Zernike coefficients (**Methods**). The additional aberrations account for the measurement error of the PSF calibration and sample induced aberrations. It makes the network more robust to different situations while maintaining relatively high accuracy. As shown in **Extended Data Fig. 4c, f and i**, the modified loss function (**Methods**) and robust training strategy greatly reduced the artifacts. Additionally, the root mean square error (RMSE) only deteriorates slightly compared to that when the training model is totally matched with the test data. We also compared the reconstruction of nuclear pore complex (NPC) protein Nup96 by the network trained with/without robust training. In **Supplementary Fig. 4**, it is clearly shown that robust training reduces the artifacts at defocus area significantly, where model mismatch happens most frequently.

Moreover, we found that training using a non-uniform background can effectively improve the network’s detection accuracy on experimental images. As shown in **Supplementary Fig. 5**, the network trained with uniform background is prone to predict many false positives in the experimental data. This is because biological images often contain structured background^27^ which are recognized as dim, large-defocus emitters by the network^17^. Although some of these artifacts can be filtered in the postprocessing step, training with data under non-uniform background greatly reduces these artifacts at the first place and avoids the reconstruction being contaminated by these false positives.

### Performance of FD-DeepLoc on simulated large-FOV data with field-dependent aberrations

We first quantify the accuracy of FD-DeepLoc using single molecules with field-dependent aberrations. To this end, we trained FD-DeepLoc and DECODE^17^ using the same synthetic data with simulated field-dependent aberrations with an FOV of 204.8 × 204.8 μm^2^ based on our customized microscope system (**Supplementary Fig. 6**). We then chose single molecules locate at five different positions in the FOV: left top, right top, left bottom, right bottom, and middle positions for localization accuracy evaluation. The PSFs are visually different at these positions (**Extended Data Fig. 5**). The localization results of FD-DeepLoc and DECODE on these datasets are shown in **Extended Data Fig. 5**. The theoretical localization precision limit, CRLB, are also shown as a reference. As indicated by **Extended Data Fig. 5**, FD-DeepLoc was able to simultaneously reach CRLB at different locations, while DECODE, trained with multiple PSF models simultaneously corresponding to the entire FOV, but without spatial context, was difficult to achieve CRLB in all locations. As the PSF patterns are not fixed with respect to their global positions during the training of DECODE, different PSF models probably confuse the network to make a proper prediction.

To quantify the performance of FD-Deeploc on SMLM images over the entire FOV with field dependent aberrations, we simulated six evaluation datasets which consisted of randomly distributed hollow rods pattern in a 204.8×204.8×1. 4 μm^3^ volume (**Fig. 3a**). The structure coordinates were generated using TestSTORM^28^. For each single molecule, we employed the same photophysics model as the SMLM challenge^29^. For each dataset, three levels of signal to background (low, medium, high) and two levels of aberrations (normal, strong) were used to generate the SMLM images (**Fig. 3b, Supplementary Fig. 6, Supplementary Fig. 7 and Supplementary Note 2**). The density is set as 1.5 emitters/μm^2^. As before, we compared FD-DeepLoc with DECODE, trained with all PSF models simultaneously, and with the cubic spline fitter^30^ from SMAP^31^ using an average PSF model.

**Fig. 3.**
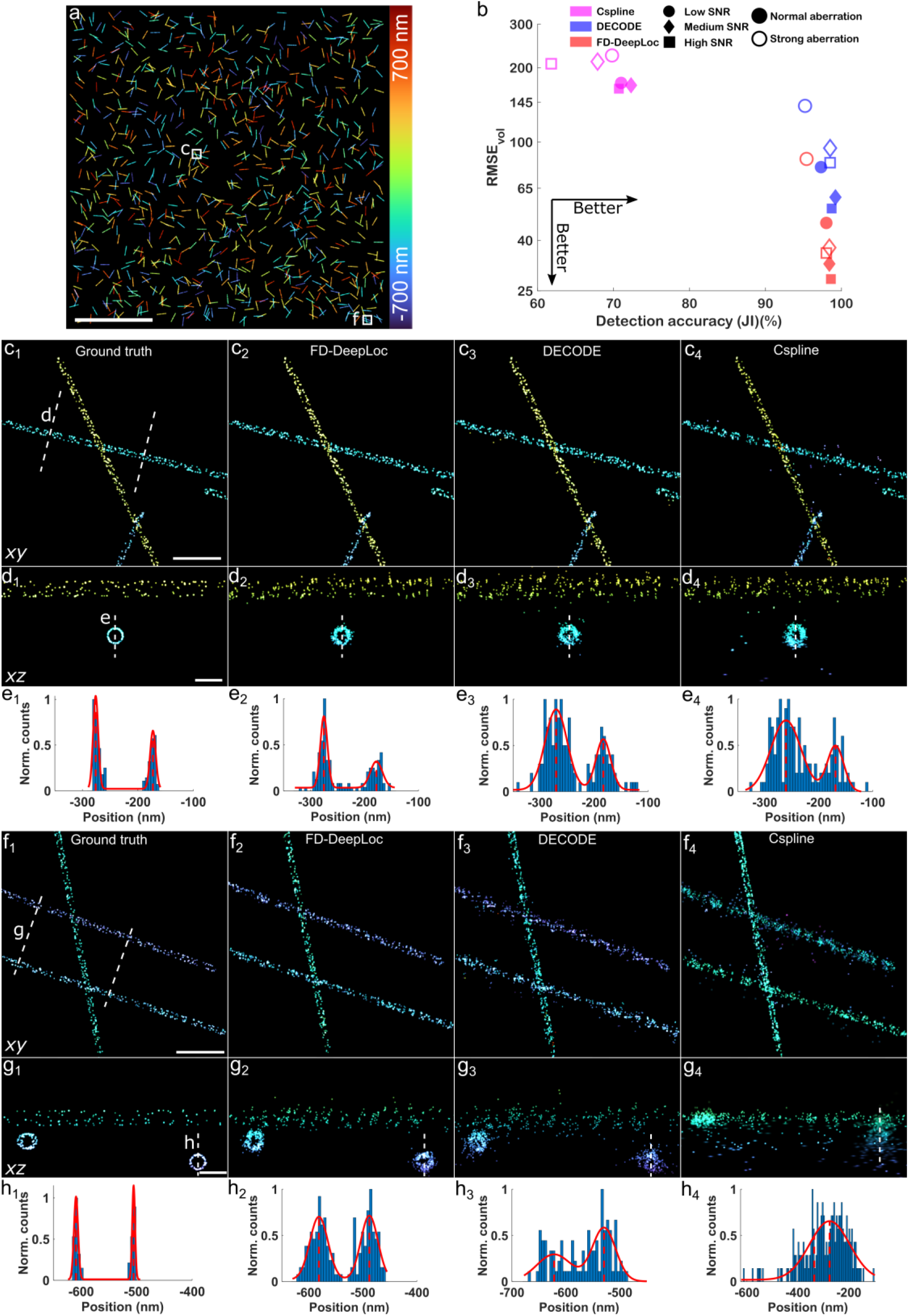
Performance of FD-DeepLoc on simulated datasets with field-dependent aberrations. **a**, Simulated hollow rods are randomly distributed in a 204.8×204.8×1. 4 μm^3^ volume. **b**, Performance evaluation on simulated datasets with different magnitudes of aberrations and SNRs using Jaccard index and RMSE. Aberration maps with two different magnitudes were used (Normal and Strong, **Methods**). Low, medium, and high SNRs are corresponding to average 1000/5000/10000 photons per emitter and 10/50/100 background photons per pixel, respectively. The fitting model for Cspline is an averaged PSF model. The training data for DECODE and FD-DeepLoc is the same. **c1-4** are zoomed views of the region indicated by the box **c** in **a**, reconstructed using ground truth and predictions of three different algorithms,respectively. **d1-4**, side-view cross-sections along the dashed lines in **c1-4**. **e1-4**, axial intensity profiles along the dashed lines in **d1-4. f**,**g**,**h** have the same meaning as **c**,**d**,**e** respectively,but for the region indicated by the box **f** in **a**. The visual comparison **c** and **f** are based on the dataset with Normal aberration and medium SNR. Scale bars, 50 μm (**a**), 1 μm (**c, f**), 200 nm (**d**, **g**).

To quantitatively compare the algorithms, we employed two metrics: Jaccard Index (JI) and volumetric Root Mean Squared Error (RMSE_vol_), which represent detection efficiency and 3D localization accuracy, respectively (**Supplementary Note 3**). Similar to previous findings^17^, deep learning-based algorithms could achieve higher JI score compared to the model fitting-based algorithms (**Fig. 3b**). Returned localizations close to the ground truth positions (< 250 nm for lateral and < 500 nm for axial) were used for calculation of RMSE_vol_. **Fig. 3b** shows the RMSE_vol_ results of these algorithms. Both deep learning methods behaved much better than the model fitting-based method as the fitter only utilizes an averaged PSF model. By additionally encoding the global position information into the data channel using CoordConv^22^, FD-DeepLoc improved the RMSE_vol_ by almost a factor of 2 compared to that of DECODE. **Fig. 3c and f** are the reconstructed 3D images of central and peripheral regions using different algorithms, respectively. As the aberrations in the central area are usually small, the hollow structure of the simulated rod can be resolved in all reconstructions (**Fig. 3d and e**). By contrast, only FD-DeepLoc can clearly resolve the hollow structure of the simulated rod in the marginal area (**Fig. 3g and h**) where the field-dependent aberrations are quite large.

### FD-DeepLoc enables large FOV 3D super-resolution imaging with high fidelity

We then applied FD-DeepLoc to experimental data from a biological sample. We imaged the nucleoporin Nup96 in U2OS cells (**Fig. 4**, **Extended Data Fig. 6** and **Supplementary Movie 1**), which is widely used as a quantitative reference structure^32^, and mitochondria TOM20 in COS7 cells (**Extended Data Fig. 7** and **Supplementary Movie 2**). Thanks to the multi-mode fiber illumination, we were able to homogeneously illuminate an FOV (~180 μm × 180 μm) that covered the full chip of our sCMOS camera, thus fully utilizing the imaging capability of the modern sCMOS technologies with ultra-high pixel numbers. We used conventional astigmatism-based 3D SMLM by inserting a cylindrical lens to the imaging path, which introduced an astigmatism of ~70 nm rms wavefront aberration to the system (**Fig. 2**).

**Fig. 4.**
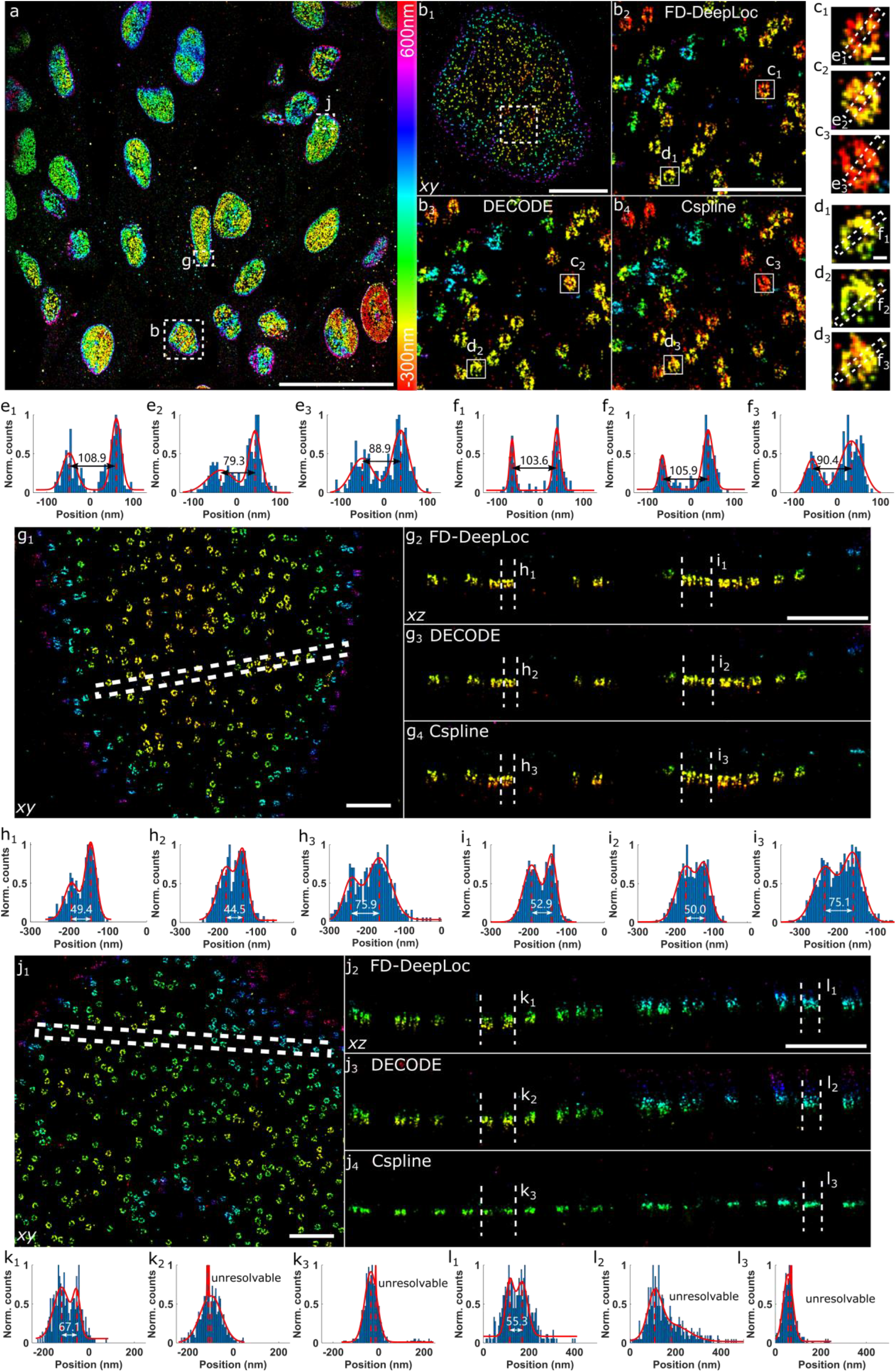
Performance of FD-DeepLoc on experimental astigmatic 3D data of NPCs in a large FOV. **a**, Overview of the panoramic 3D super-resolution image of Nup96-SNAP-AF647 reconstructed by FD-DeepLoc. **b**, Top views of the NPCs reconstructed with FD-DeepLoc, Cspline and DECODE. **b1**, Zoomed view of the region indicated by the dashed box b in **a**. **b2**, **b3** and **b4** are the zoomed images of the rectangle region indicated in **b1** reconstructed by FD-DeepLoc, Cspline and DECODE, respectively. **c** and **d**, Zoomed views of the regions indicated by boxes c and d in **b**. **e** and **f**, Intensity profiles of the dashed lines denoted in **c** and **d**, respectively. **g**, Zoomed view of the region indicated by the dashed box g in **a**. **g2-4**, Side-view cross-sections of the region as indicated by the dashed lines in **g1** reconstructed by: **g2**, FD-DeepLoc; **g3**, Cspline; **g4**, DECODE. **h** and **i**, Intensity profiles of the dashed lines denoted in **g**. **j**- **l** are the same as **g**- **i**, but for regions denoted by the dashed box j in **a**. Representative results are shown from 3 experiments. Scale bars, 50μm (**a**), 5μm (**b1**), 1μm (**b2**, **g1**, **g2**, **j1**, **j2**) and 50nm (**c1**, **d1**).

We then compared the reconstructed images analyzed by FD-DeepLoc, DECODE and Cspline. For model fitting based Cspline method, an averaged PSF model in the central area (center 512 ×512 pixels) was used. In the central region, where the aberrations are relatively small, all algorithms can reconstruct the double ring structure of the nucleoporin Nup96 as indicated by the axial intensity profiles (**Fig. 4g-i**). In contrast, the returned localizations by Cspline tend to converge to the same z-positions in the peripheral areas (**Fig. 4j**, **Extended Data Fig. 6** and **7**) where the model mismatch happens. For the deep learning-based methods, all PSF features across the entire FOV were included in the training dataset. However, since the spatially variant PSF patterns are trained without spatial context in DECODE, DECODE returned ghost images (**Fig. 4j**, **Extended Data Fig. 6**), which agrees with the previous simulated data. In contrast, FD-DeepLoc could nicely reconstruct the double ring structure of Nup96 at different locations with different aberrations (**Fig. 4h, i, k, l**), showing superior Fourier ring correlation (FRC) resolution (**Supplementary Fig. 8**).

### FD-DeepLoc combined with DM based optimal PSF engineering enables whole cell 3D super-resolution imaging over large FOV

PSF engineering encodes the *z* information to the shape of the PSF and could enable 3D super-resolution imaging without the need to scan the sample^33,34^. In order to perform volumetric super-resolution imaging of the whole cell at high throughput, we optimized deformable mirror (DM) based PSF^35^ that has best 3D CRLB in a 6 μm axial detection range which could cover the entire cell (**Methods**). As the DM has limited number of actuators, Zernike polynomial or pixelwise based pupil function optimization methods could lead to approximation error using DM. Here, we directly employed the influence functions of the DM as the basis function for the pupil function optimized and called the PSF optimized in this way as DM based optimal PSF (DMO PSF)^35^. We then applied the 6 μm DMO PSF to image the mitochondria and NPC in the whole cell (**Fig. 5**, **Extended Data Fig. 8** and **Supplementary Fig. 9**).

**Fig. 5.**
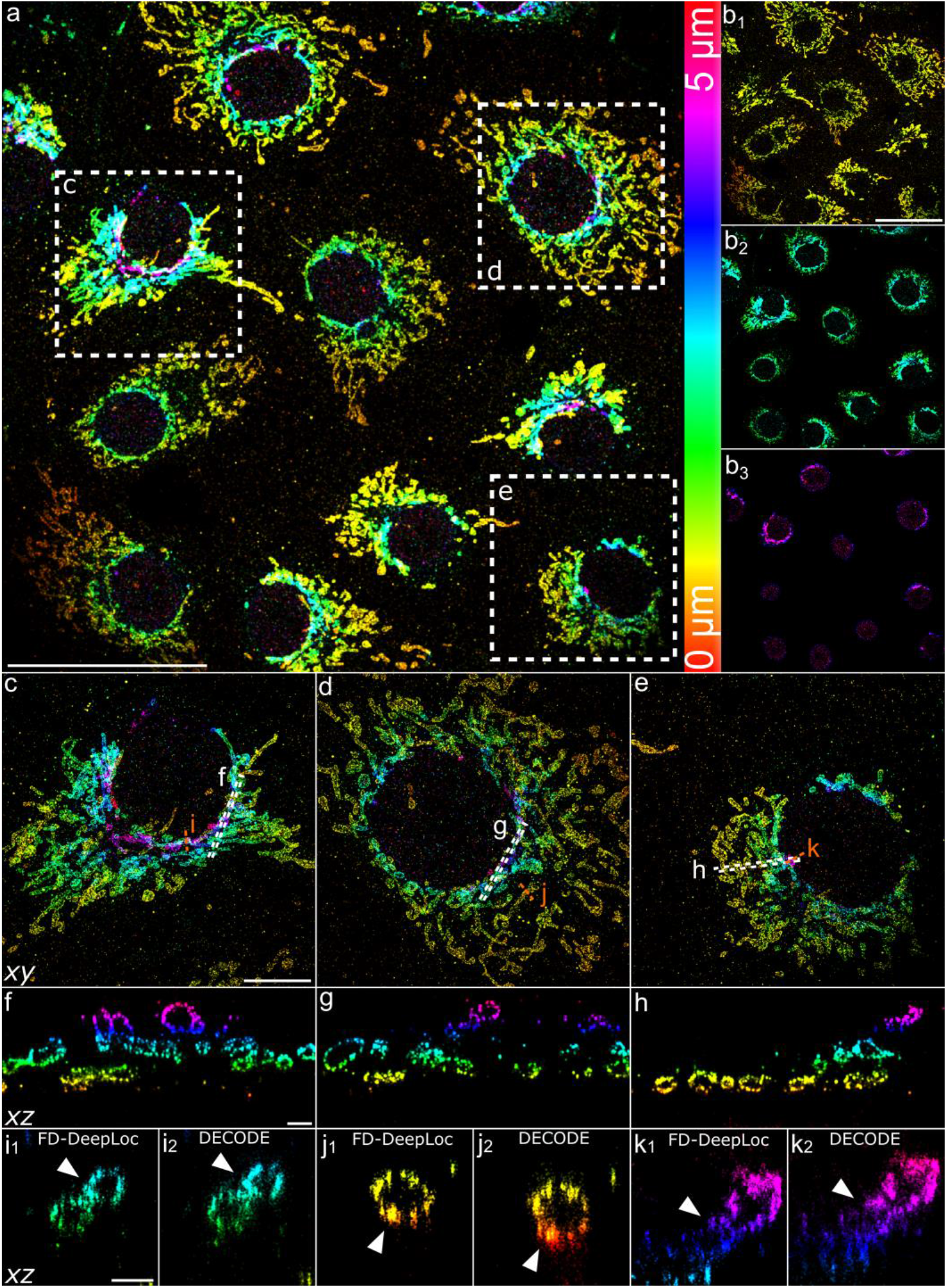
FD-DeepLoc enables 3D super-resolution imaging of mitochondria within a large FOV (180 × 180 μm^2^) and DOF (**5 μm**). **a**, Overview of the panoramic 3D super-resolution image of mitochondria, where *z* position is indicated by color. **b**, Different axial sections of **a**. **b1-3** are top views of axial range of (0 μm - 1.7 μm), (1.7 μm - 3.4 μm) and (3.4 μm – 5 μm), respectively. **c**,**d**,**e**, Magnified views of areas denoted by the dashed white boxes in **a**. **f**, **g** and **h** are side-view cross-sections of the region denoted by the white dashed lines in **c**, **d** and **e**, respectively. **i**, **j** and **k** are side-view cross-sections of areas denoted by the orange dashed lines in **c**, **d** and **e**, respectively. Raw beads and single molecule data for DMO Tetropad PSF can be found in **Supplementary Fig. 9**. Representative results are shown from 7 experiments. Scale bars, 50 μm (**a**, **b1**), 10 μm (**c**), 1 μm (**f**) and 500 nm (**i1**).

With the help of FD-DeepLoc, we were able to reconstruct high quality 3D super resolution image over the entire FOV (**Fig. 5a** and **Supplementary Movie 3**). The axial imaging range spanned over ~ 5 μm (**Fig. 5b** and **Supplementary Movie 4**). We then chose three cells locates at different corners of the FOV. The hollow structure of the mitochondria in these cells could be nicely resolved in the side-view cross section images. Compared with DECODE, FD-DeepLoc reconstructed sharper mitochondria images as shown in **Fig. 5i - k**. The improved resolution is also confirmed by the FRC analysis of the reconstructed images by these two methods. The overall FRC resolution of these cells is about ~50 - 60 nm for FD-DeepLoc while it is ~80 - 100 nm for DECODE (**Supplementary Fig. 10**). We also performed 3D whole cell imaging of NPCs. As shown in **Extended Data Fig. 8, Supplementary Movie 5 and 6**, FD-DeepLoc could resolve the structure of the whole nuclear envelope. In the zoomed image as shown in **Extended Data Fig. 8c and d**, we were able to resolve the nuclear pores as rings both in the top and bottom of the nuclear envelope. The consistency of the radii of NPCs is also verified at the top and bottom of the cells in both central and marginal regions (**Extended Data Fig. 8e and f**, **Supplementary Note 4**).

As FD-DeepLoc enables super-resolution imaging with both large FOV and DOF, quantitative analysis of groups of super-resolved whole cells becomes feasible. To this end, we used an ImageJ plugin, Mitochondria Analyzer^36^, to analyze the reconstructed super-resolution images of whole cell mitochondria. Thanks to the high throughput imaging capabilities, 121 whole cell 3D super-resolution mitochondria images can be acquired in only 16 ROIs within few hours (**Extended Data Fig. 9**). We then extract their morphology parameters and network connectivity (*e.g*., volume, sphericity, branch length, *etc*.) using Mitochondria Analyzer plugin (**Supplementary Note 4**). The features (eight-dimensional feature vectors) were finally classified using k-means++ algorithm^37^. To visualize the classified results, principal component analysis was performed on the same feature vectors with the first 3 principal components shown in **Extended Data Fig. 9**. 3 types of cells with different mitochondria features were observed: Type1 cells contain more small round mitochondria and fewer branches; Type2 cells contain outspreading mitochondria and more complex networks; Type3 cells contain a mixture of spherical and tubular mitochondria. These results demonstrate that FD-DeepLoc is compatible with many downstream biological image analysis tools. The high throughput data with super-resolution would hopefully give new insights to many biological applications.

### FD-DeepLoc combined with DMO PSF with flexible DOF enables high resolution imaging of neuronal processes across large FOV

Neuronal cells grow neurites over hundreds of microns in culture. Conventional astigmatic 3D SMLM imaging of neurites is normally limited to a DOF < 1.2 μm and FOV < 50 × 50 μm ^2^, impeding the visualization of neurites with large diameter and their 3D organization in large scale. Here, we imaged neuronal cells which were induced from mouse embryonic stem cells (**Methods**). The induced neuronal cells were grown for about 21 days and labeled by β2-spectrin which forms a periodic submembrane scaffold along axons^38^. We first applied the widely used astigmatic PSF for the 3D imaging. The periodic organization of spectrin along the neurites is clearly resolved (**Fig. 6a - c**). However, due to the limited axial range, the neurites showed discontinuous structure where their axial spread is larger than the DOF of astigmatic PSF (**Fig. 6d and e**). We therefore engineered a 3 μm DMO PSF using a deformable mirror as described before^35^. Compared to the astigmatic PSF, the 3μm DMO PSF improved the axial range by more than 2 times, while the averaged 3D CRLB is almost the same (**Supplementary Fig. 11**). We then applied the 3μm DMO PSF to image the thick neurite network across large FOV. The 3D distribution of β2-spectrin can also be nicely reconstructed even in the neurites with large diameter using the 3μm DMO PSF (**Fig. 6f - j, Supplementary Movie 7**). The periodic structure can be observed in both the top and bottom surfaces of the neurites (**Fig. 6j**). The demonstrated high resolution over large FOV and DOF would benefit many applications with large sample size.

**Fig. 6.**
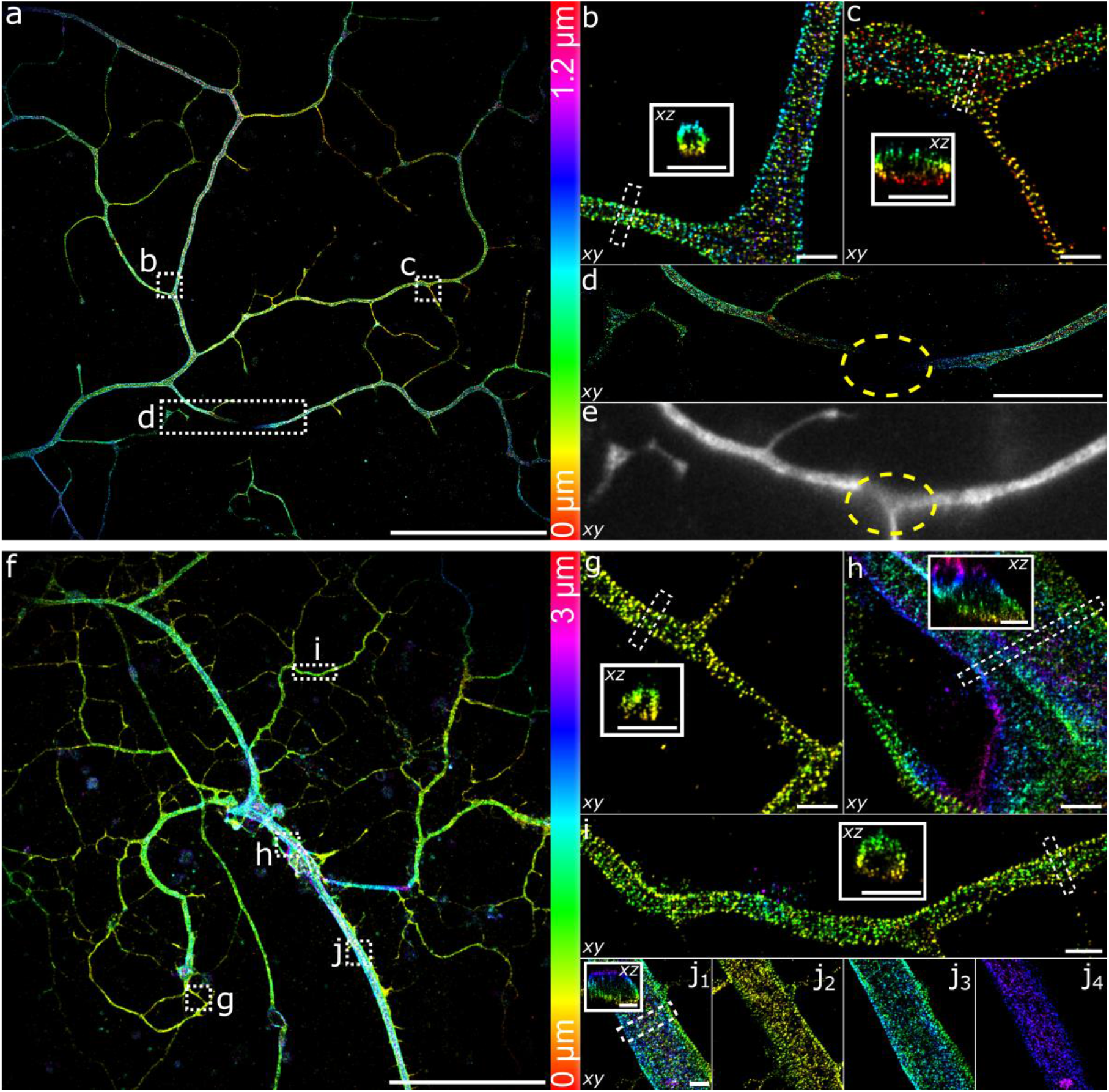
FD-DeepLoc with flexible DMO PSF engineering enables high quality 3D super-resolution imaging of thick neurites over a large FOV. **a**, Overview of the panoramic 3D super-resolution image of neurites using astigmatic PSF. **b**, **c**, **d**, Magnified views of areas denoted by the dashed white boxes in **a**, respectively. Insets in **b** and **c** are the corresponding side-view cross-section of the areas denoted by the dashed lines in **b** and **c**, respectively. **e**, Wide-field diffraction limited image of the area **d**. **f**, Overview of the panoramic 3D super-resolution image of neurites using 3 μm DMO PSF. **g**- **j**, Magnified views of areas denoted by the dashed white boxes in **f**, respectively. Insets in **g - j** are the corresponding side-view cross-section of the areas denoted by the dashed lines in **g - j**, respectively. **e**, Wide-field diffraction limited image of the area **d**. **j2 - 3** are top views of axial range of (0 μm – 1 μm), (1 μm - 2 μm) and (2μm – 3 μm), of **j1** respectively. Representative results are shown from 7 experiments. Scale bars, 50 μm (**a**, **f**), 10 μm (**d**) and 1 μm (**b, c, g, h, i, j** and **insets**).

## Discussion

To summarize, we presented FD-DeepLoc, a special deep learning-based method for SMLM with spatial awareness that enables high throughput whole cell 3D super-resolution imaging over a large FOV and DOF. We attribute the broad applicability of high throughput super-resolution imaging to three advances made in this work: (1) a GPU based vectorial PSF fitter for fast and accurately modeling spatially variant PSFs over large FOV under high NA objectives; (2) full frame large FOV imaging using the modern sCMOS technology and large DOF imaging using an PSF optimized for DM with large axial range; (3) a robust spatially variant CNN, FD-DeepLoc, that is able to learn spatially variant features to correct the field dependent aberrations. As a result, both in the popular astigmatism and DMO Tetrapod PSF based 3D super-resolution imaging, FD-DeepLoc achieved high accuracy reconstruction of biological structures with high fidelity across the entire camera frame.

An accurate and comprehensive PSF model is the key to extract multi-dimensional information from single molecules, and vectorial PSF model is superior for a high NA objective system as both polarization effects and refractive index variations are considered. Our GPU based vectorial PSF fitter takes full advantage of the accurate vectorial PSF model incorporating high-dimensional optical parameters, while maintaining a fast computational speed (<1 s for bead stacks size of 27×27×41). Furthermore, our vectorial PSF fitter is very flexible in terms of parameter sharing between different single molecule images, which could be used to extract the rich information (*i.e*., aberration, dipole orientation, wavelength, *etc*.) within single molecules by global optimization of different single molecule images.

Aberrations in the optical system are often static and field dependent. Correcting the aberrations with a shift-varying PSF models is normally slow and computationally intensive^39^. FD-DeepLoc with CoordConv can successfully encode the position information into the neural networks, without significantly increasing the size of the network. Therefore, the CoordConv strategy is very suitable to many position sensitive imaging applications, such as field dependent PSF deconvolution^39^ and multi-channel PSF modeling with shift-invariant transformation^19^, with only minor modification to the conventional CNNs.

Correction of the field dependent aberrations, as well as with the optimized large DOF PSF engineering, volumetric 3D super-resolution imaging of samples with large size, *e.g*., organoid, tissue sections, mammalian embryos *etc*., becomes accessible to many labs without the need of complicated hardware automation. For thick samples with more prominent sample induced aberrations, a more accurate in-situ PSF model (*e.g*., INSPR^40^) could be used to account for the sample-induced aberrations. Correctly training FD-DeepLoc with such models would make FD-DeepLoc robust to sample-induced aberrations. Future development could include 3D CoordConv for correction of sample induced depth dependent aberrations. Finally, as an open-source code with detailed examples and tutorials, FD-DeepLoc will transform 3D SMLM from a low throughput technique into a high throughput imaging technology even under a microscope without much hardware modification.

## Supporting information

Supplementary Software

## Methods

### GPU based vectorial PSF fitter

The field dependent aberrations are retrieved from through focus imaging stacks of fluorescent beads using vectorial PSF model (**Supplementary Note 1 and Supplementary Fig. 12**). MLE fitting routine was employed to analyze the bead stacks with Zernike coefficients as the fitting parameters. We then employed globLoc^26^ for the parameter estimation. Each image in the bead stack can be regarded as an individual channel, whose fitting parameters can be shared or fit independently. Since the aberration term *ψ_aber_* in each bead stack is the same at different *z* positions, the Zernike coefficients are treated as global parameters across the *z* stack. The other parameters can either be treated as global or local parameters depending on different imaging conditions. Since the system drift is very small during the acquisition of beads stack in our microscope, we linked *x*, *y*, and *z* parameters while the photons and background parameters are unlinked to account for the photon bleaching effect during imaging acquisition.

Since vectorial PSF fitter is computational challenging, we implemented the fitting pipeline using CUDA C/C++ in NVIDIA CUDA® enabled graphic cards to enable fast modeling of the spatially variant PSFs in the large FOV. The framework of the MLE fitting method follows the previous work^26^ which employed a modified Levenberg-Marquardt iterative schemes for non-linear optimization. To accurately retrieve the aberrations from point emitter, single molecule images at different *z* positions are normally needed. To make full use of GPU multithreaded parallel operation, pupil function of single molecules at different *z* positions are calculated parallelly in different blocks before fast Fourier transform (FFT). As calculation of the derivatives for each parameter (21 Zernike coefficients plus x, y and z) in each iteration requires 24 FFTs (including forwards and inverse FFT) and FFT is very time consuming, we parallelly executed multiple FFTs with GPU. We set the maximum parallelism for 64 molecules to avoid the memory overflow. 512 threads were used in each block. Our GPU-based vectorial PSF fitter is about 50 times faster than CPU based C/C++ code. The GPU-based CUDA code is compiled in Microsoft Visual Studio 2019 for both Matlab and Python individually. A PC equipped with an Intel Core i9-9900 processor of 128GB RAM clocked at 3.50GHz and an NVIDIA GeForce GTX 3090 graphics card with 24.0 GB memory was used for speed benchmarking.

### Calibration of aberration maps over a large field of view

The field-dependent aberrations are retrieved from thousands of through-focus bead stacks randomly distributed in the whole FOV on the cover glass (**Fig. 2**). For astigmatism PSF calibration, 41 single emitter images were recorded with a *z* step of 50 nm over an axial range of 1 μm above and below the focus. For Tetrapod PSF calibration, 61 single emitter images were recorded with a *z* step of 100 nm over an axial range of 3 μm above and below the focus. All bead stacks were acquired with an exposure time of 100 ms and illumination intensities of ~400 mW (over 180 ×180 μm^2^ FOV) on the sample. We removed bead stacks whose center position are less than 25 and 50 pixels from their nearest neighbor beads for astigmatism and Tetrapod beads calibration, respectively. Each bead stack was segmented in a maximum-intensity projected image by maximum finding and thresholding. We then used the GPU based vectorial PSF fitter to fit each bead stack by MLE to retrieve the Zernike based aberration coefficients. In this work, the coefficients of all the 21 tertiary Zernike polynomials were retrieved for each bead stack. The retrieved coefficients represent the local aberration on the position where the bead stack locates. To exclude some outlier bead stacks (*e.g*., aggregated beads, background contaminated beads, *etc*.), we filtered the beads whose Zernike coefficients with a difference to the local mean value (256**×** 256 pixels around the bead for astigmatism PSF and 1024**×** 1024 pixels for Tetrapod PSF) are more than 2.5 times of the standard deviation of the local coefficient values. Bead stacks whose relative root square error (RRSE)^42^ between the raw data and fitted model was more than 15% for astigmatism PSF and 20% for Tetrapod PSF were also discarded (**Supplementary Fig. 2**). The RRSE is defined by

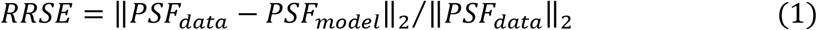

where *PSF*_*data*_ is the bead stack raw data and *PSF*_*model*_ is the fitted PSF model. After filtering, we interpolated each aberration coefficient across the whole FOV using natural interpolation. Gaussian smoothing with a sigma of 100 and 200 pixels were applied to the aberration maps for astigmatism and Tetrapod beads, respectively.

### PSF engineering

A deformable mirror based optimal PSF (DMO PSF) was employed in this work. Similar to our previous work^35^, we optimized the pupil function of the engineered PSF by minimizing its 3D CRLB. Instead of using Zernike polynomials as the solution space, we used the influence function of each DM actuator as the basis function of the pupil function optimized. It offers more accurate and flexible wavefront design and avoids the approximation error which often happens during the Zernike based and pixelwise wavefront control. The objective function for pupil function optimization is given by:

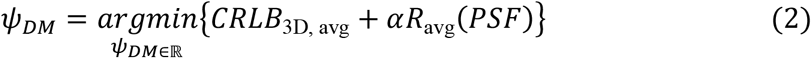

where 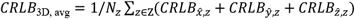 is the averaged 3D CRLB over *N*_*z*_ discrete *z* positions in a predefined axial range Ζ. *R*_avg_(*PSF*) term is used to confine the spatial extent of the PSF to reduce overlap:

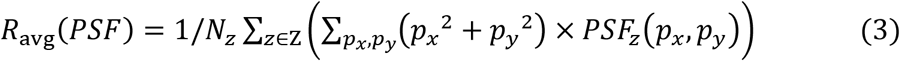

where (*p_x_, p_y_*) represents pixel coordinates with the center of PSF model as zero. *α* is a hyper-parameter. In this work, we set *α* as 0 and 30 for the 1.2 μm DMO saddle point PSF and 6 μm DMO Tetrapod PSF optimization, respectively. The *z* steps were set as 100 nm and 200 nm for DMO saddle point and Tetrapod PSF respectively.

### FD-DeepLoc network

To enable the network to analyze SMLM images with field-dependent features, we employed the CoordConv strategy^22^. Two extra coordinate channels were added to the input, one for the *x* position and the other one for the *y* position in normalized coordinates. In CoordConv layer, there are extra convolution kernels with size of *n* × 2 × 1 × 1 working on the coordinate channels (**Extended Data Fig. 3**), where *n* is the number of output channels of the original convolutional layer, 2 refers to the input *xy* channels, 1 × 1 is the size of the convolutional kernel. The result of the coordinate convolution is then added on the result of image convolution to form the output of CoordConv layer. With only 2*n* additional weights for each CoordConv layer, the network is sensitive to the spatially variant PSFs.

Similar to DECODE, the main network architecture of FD-DeepLoc consists of two stacked U-nets^43^: frame analysis module and temporal context module (**Extended Data Fig. 3**). We incorporated the first convolutional layer of each module with CoordConv^22^. Each module contains 16 layers, including twelve 3×3 convolutional layers, two 2×2 convolutional layers and two up-sampling layers. At each 2×2 convolutional layer and up-sampling layer, the resolution will be halved and doubled, respectively. The frame analysis module receives 3 consecutive frames and outputs 48-channel features for each frame. These features are then analyzed by the temporal context module, whose outputs are used to form the final prediction. Exponential linear unit (ELU) activation is used after each hidden layer.

For each 3 consecutive frames’ input unit, FD-DeepLoc outputs ten-channel maps (**Supplementary Note 5**): the first two channels are pixel-wise probability 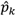 of an emitter exists in the pixel *k* and its corresponding photons 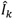. To achieve sub-pixel precision, another three channels output two lateral continuous-valued offset 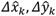 relative to the center of the pixel *k* and axial distance 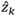 relative to the focal plane. Four additional channels represent the uncertainty *σ_xk_, σ_yk_, σ_zk_, σ_Ik_* for *x*, *y*, *z* and photon parameters estimation, respectively. The last channel 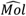 is optional, which predicts the theoretical single molecule image of the inputs. This can also be used to predict background by subtracting it from the raw image. The Tanh activation function were used for localization offset 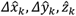 channels, while the Sigmoid function were used for all other channels.

### Loss function

The loss function of FD-DeepLoc is a modified version of that of DECODE and consists of four parts: cross-entropy term, count loss term, localization loss term and PSF loss term:

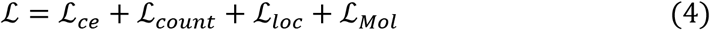

Here, 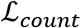 and 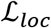 are the same as the ones in the DECODE loss function. 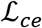 is the cross-entropy (CE) between the predicted probability map and its corresponding ground truth map:

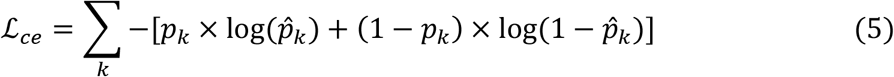

where 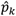 is the predicted probability that an emitter exists in the pixel *k*, *p_k_* is the binary ground truth probability. As the network outputs continuous-valued localizations formed by pixel-wise probability 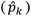 and subpixel offsets 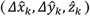. The overall localization problem can be treated as a combination of classification (whether an emitter exists in a pixel) and position regression tasks. Here, the 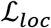 term is the negative log-likelihood of the position regression task (Gaussian-mixture model) and the 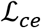 term is the negative log-likelihood of the classification task (Bernoulli distribution). The CE term could be regarded as a complement for 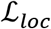 as it provides a clearer objective for the probability channel. In our simulated dataset, it could help the probability output channel of the network converge better and leads to better RMSE (**Extended Data Fig. 4b, Supplementary Fig. 13**). Although the two categories are highly imbalanced (most pixels are empty even for dense localization), the 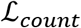 term could effectively avoid predicting an empty map.

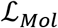 measures the error between the predicted single molecule image 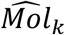 and the ground truth single molecule image *Mol_k_* without adding background and camera noise:

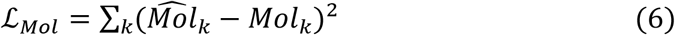

Here, 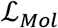 is used to enforce the network to learn the spatially variant features of PSFs. The predicted theoretical single molecule images can also be utilized to generate the background image by subtracting it from the raw data.

### Training data simulation

FD-DeepLoc follows the training procedure as shown in **Fig. 1c**. The training data are generated online using the spatially variant PSF model calibrated before (**Supplementary Note 5**). To generate temporal image sequences for training, we simulated the data in the units of three consecutive images. The network’s prediction is for the molecules in the middle frame of each unit. Different from the sophisticated photoactivation model^44^, we simply let the emitters with predefined density be randomly activated, and the probability of on-state emitters appear in the next frame follows a simple binomial distribution since only three consecutive images are used in each unit. The emitter positions are randomly placed in the entire FOV. The photon of each emitter is evenly drawn from a predefined range, which could be adjusted with different dyes and excitation intensity. The aberration coefficients of each emitter are indexed according to its lateral position.

In addition, considering the potential measurement error of PSF calibration and the sample induced aberrations, the network may be overfitted to a biased model different from the real experimental data, leading to artifacts in the reconstruction. Therefore, we gave more freedom to the PSF simulator by adding small random Zernike aberrations to each single molecule in the training data. To this end, we added each field dependent Zernike coefficient with an additional zero mean normal distributed value with a standard deviation of λ/100. We found this helped the network perform much better when models were mismatched (**Extended Data Fig. 4**).

In real biological specimens, the background is spatially variant due to nonspecific labeling or autofluorescence from different structures of samples^27^. Training the network with constant background is prone to misrecognize the structured background as dim, large-defocus emitters (**Supplementary Fig. 5**). To solve this problem, we added random non-uniform background to the training data and found this improved detection accuracy a lot in experiments. We choose Perlin noise with a principal frequency of 64 image pixels and an octave number of 1 to mimic the non-uniform background^45^. The frequency and octave number are empirically chosen as we found it ran well on our experimental images.

Both EMCCD and sCMOS based training data were simulated. To simulate the EMCCD camera noise, we incorporated the same noise model as SMLM challenge^29^. The noise model mainly consists of the shot noise, electron multiplication noise and camera readout noise. For the sCMOS camera, there is no electron multiplication process. Therefore, only the Poisson shot noise and Gaussian readout noise were considered. Although it should be noted that the gain, offset and readout noise of sCMOS camera are pixel-dependent, we found using simple constants is enough as the pixel variant noise is quite small and did not affect the result too much (**Supplementary Fig. 14**).

### Optical setups for large FOV

In this work, high-throughput SMLM imaging was performed at room temperature (24 °C) on a custom-built microscope (**Extended Data Fig. 10**). We combined a laser box with a multi-mode fiber (WFRCT200×200/230×230/440/620/1100N, NA=0.22, CeramOptec) to deliver homogeneous illumination to the sample^8^. Single-mode fiber (P3-405BPM-FC-2, Thorlabs) excitation for total internal reflection fluorescence (TIRF) imaging can also be achieved with a flip mirror (KSHM40/90, Owis). The multi-mode fiber was bound with a vibrator to reduce the laser speckle^46^. The lasers were triggered by a field-programmable gate array (Mojo, Embedded Micro), allowing microsecond pulsing control of lasers. The illumination beam was filtered by a laser clean-up filter (ZET405/488/561/640xv2, Chroma) to remove fiber-induced fluorescence. The excitation laser was then reflected by a main dichroic (ZT405/488/561/640rpcxt-UF2, Chroma) before entering the objective for sample illumination. Sample fluorescence was collected by a high NA objective (NA 1.5, UPLAPO 100XOHR or NA 1.35, UPLSAPO 100XS, Olympus) and then filtered by a quad-band emission filter (ZET405/488/561/640mv2, Chroma). In this work, NA 1.5 objective was used for astigmatism 3D imaging, and NA 1.35 objective was used for DMO Tetrapod PSF based 3D imaging. After the tube lens (TTL-180-A, Thorlabs), the back focal plane of the objective was imaged onto a deformable mirror (DM140A-35-P01, Boston Micromachines) for PSF engineering. Before entering the sCMOS camera (PRIME 95B, Teledyne Photometrics), a band-pass filter (ET680/40m, Chroma) was inserted to the beam path to reject residual laser light. A 785 nm near infrared laser (iBEAM-SMART-785-S, Toptica Photonics) was introduced to the system by a dichroic mirror (FF750-SDi02, Semrock) for sample focus stabilization. The reflected laser from coverslip was detected by a quadrant photodiode (SD197-23-21-041, Advanced Photonix Inc) whose position dependent output voltage was used as feedback to the objective *z* stage (P-726.1CD, Physik Instrumente). The software control of the microscope was integrated in Micro-Manager with EMU^47^. Typically, we acquired 50,000-100,000 frames with a 15 ms exposure time.

### Sample preparation for imaging of the nuclear pore complex and mitochondria

#### Cell culture

COS-7 cells (catalog no. 100040, BNCC) were grown in DMEM (catalog no. 11995, Gibco) containing 10% (v/v) fetal bovine serum (FBS; catalog no. 10270-106, Gibco), 100 U/ml penicillin and 100 μg/ml streptomycin (PS; catalog no. 15140-122, Gibco). U2OS cells (Nup96-SNAP no. 300444, Cell Line Services) were grown in DMEM (catalog no. 10569, Gibco) containing 10% (v/v) FBS, 1× PS and 1×MEM NEAA (catalog no. 11140-050, Gibco). Cells were cultured in a humidified atmosphere with 5% CO_2_ at 37 °C and passaged every two or three days. Prior to cell plating, high-precision 25-mm-round glass coverslips (no. 1.5H, catalog no. CG15XH, Thorlabs) were cleaned by sequentially sonicating in 1 M potassium hydroxide (KOH), Milli-Q water and ethanol, and finally irradiated under ultraviolet light for 30 min. For super-resolution imaging, COS-7 and U2OS cells were cultured on the clean coverslips for 2d with a confluency of 50-70%.

Mouse embryonic stem cells (mESCs) were a subclone of an established ESC line originally named E14. mESCs were cultured in feeder-free conditions, using serum/LIF medium containing high-glucose DMEM (HyClone, SH30022.01), 15% fetal bovine serum (Gibco, 30044-333), 1× sodium pyruvate (Gibco, 11360070), 1× penicillin-streptomycin (Gibco, 15070063), 1× nonessential amino acids (NEAA, Gibco, 11140050), 1× GlutaMAX (Gibco, 35050061), 1 mM 2-mercaptoethanol (Sigma, M3148) and leukemia inhibitory factor (LIF). All the cells were cultured at 37°C in a humidified atmosphere containing 5% CO_2_. Mycoplasma detection were conducted routinely to ensure mycoplasma-free conditions throughout the study.

### Generation of induced neuronal cells from mESCs

Mouse ESCs were induced into neuronal cells as described previously^48,49^ with modifications. In brief, constitutive expressed rtTA and hygromycin resistance gene and Ngn2-P2A-puro driven by Tet-on promoter were transfected into mESCs using Piggybac system. The cells were selected with 200 μg/ml hygromycin (Sigma, V900372) for 3-5 days. On day 0, the cells were digested and plated (6-well plates, 1× 10^5^ cells/well) on poly-D-lysine/laminin coated coverslips in serum/LIF medium. Twelve hours later, the culture medium was replaced with N2B27 (1:1 mixture of DMEM/F12 (Gibco, 11320033) and Neurobasal (Gibco, 21103049), 1% N2 (Gibco, 17502048), 2% B27 (Gibco, 17504044), 1× sodium pyruvate, 1× penicillin - streptomycin, 1× NEAA, 1× GlutaMAX, 1 mM 2-mercaptoethanol) containing 5 μM retinoic acid (Sigma, R2625) and 2 mg/ml doxycycline (TargetMol, T1687). On day 1, a 24 hours puromycin selection (1 mg/ml) period was started. On day 6, Ara-C (2 mM, TargetMol, T1272) was added to the medium to inhibit neural stem cell proliferation. After day 2, 50% of the medium in each well was exchanged every 2 days, and induced neuronal cells were assayed on day 21 in most experiments.

#### SNAP-tag labeling of Nup96

To label Nup96, U2OS-Nup96-SNAP cells were prepared as previously reported^32^. Briefly, cells were prefixed in 2.4% paraformaldehyde (PFA) for 30s, permeabilized in 0.4% Triton X-100 for 3 min and subsequently fixed in 2.4% PFA for 30 min. Then, cells were quenched in 0.1 M NH_4_Cl for 5 min and washed twice with PBS. To decrease unspecific binding, cells were blocked for 30 min with Image-iT FX Signal Enhancer (catalog no. I36933, Invitrogen). For labeling, cells were incubated in dye solution (1 μM BG-AF647 (catalog no. S9136S, New England Biolabs), 1 mM DTT (catalog no. 1111GR005, BioFroxx) and 0.5% bovine serum albumin (BSA) in PBS) for 2 h and washed 3 times in PBS for 5 min each to remove excess dyes. Lastly, cells were postfixed with 4% PFA for 10 min, washed with PBS 3 times for 3 minutes each and stored at 4 °C until imaged.

#### Mitochondria labeling

Mitochondria sample were prepared as previously described^50^. Briefly, COS-7 cells were fixed with 4% PFA (preheat to 37 °C before using) in PBS for 12 min, incubated in permeabilization buffer (0.3% CA-630 (catalog no. I8896, Sigma), 0.05% TX-100, 0.1% BSA and 1× PBS) for 3 min, and then quenched in 0.1 M NH_4_Cl for 5 min. After washed 3 times for 5 minutes each with PBS, cells were blocked in 3% BSA for 60 min. For labeling, cells were stained by rabbit anti-Tom20 (catalog no. ab78547, Abcam, 1 mg/ml) with 1:1,000 dilution in 3% BSA and incubated overnight at 4 °C and washed 3 times for 5 minutes each with PBS. Cells were then stained with the corresponding secondary antibodies conjugated with AF647 (catalog no. A21245, Invitrogen, 2 mg/ml) with 1:2,000 dilution in 3% BSA for 2 h and washed with PBS. Finally, cells were postfixed with 4% PFA for 10 min, washed 3 times for 5 minutes each with PBS and stored in PBS at 4 °C.

#### Neuronal cells labeling

Neuronal sample were labeled according to Ref. ^51^. Briefly, cultured mESCs induced neuronal cells were fixed between 21 - 25 days in vitro using 4% PFA (preheat to 37 °C before using) in PBS for 30 min and washed three times in PBS. The cells were then permeabilized with 0.15% Triton X-100 in PBS for 10 min. After washed 3 times for 5 minutes each with PBS, cells were blocked in 3% BSA for 60 min. For labeling, cells were incubated in mouse anti-βII spectrin (catalog no. 612563, BD Biosiences, 250 μg/ml) with 1:100 dilution in 3% BSA, incubated overnight at 4 °C and washed 3 times for 5 minutes each with PBS. Cells were then stained with secondary antibodies goat anti-mouse IgG conjugated AF647 (catalog no. A21235, Invitrogen, 2 mg/ml) with 1:800 dilution in 3% BSA for 2 h and washed with PBS 3 times for 5 minutes each. Finally, cells were postfixed with 4% PFA for 20 min, washed 3 times for 5 minutes each with PBS and stored in PBS at 4 °C before imaging.

#### Imaging Buffer

Samples were imaged in refractive index matching buffer including 50 mM Tris-HCl (pH 8.0), 10 mM NaCl, 10% (w/v) glucose, 0.5 mg/ml glucose oxidase (G7141, Sigma), 40 μg/ml catalase (C100, Sigma), 35 mM cysteamine and 28.5% (v/v) 2,2’-thiodiethanol (166782, Sigma). The refractive index of the final imaging buffer is 1.406.

### Statistics and reproducibility

Figures show representative data from 3 (Fig. 4, Extended Data Fig. 7, Supplementary Figure 3, Supplementary Figure 4, Supplementary Figure 7) or 7 (Fig. 5, Extended Data Fig. 9, Supplementary Figure 8) or 2 (Extended Data Fig. 8) independent experiments.

## Acknowledgements

This work was supported by the Key Technology Research and Development Program of Shandong (2021CXGC010212), Guangdong Natural Science Foundation Joint Fund (2020A1515110380), Shenzhen Science and Technology Innovation Commission (Grant No. KQTD20200820113012029, KQTD20210811090115021), Guangdong Provincial Key Laboratory of Advanced Biomaterials (2022B1212010003), China National Postdoctoral Program for Innovative Talents (BX20220141) and the Startup grant from Southern University of Science and Technology.

## Author contributions

Y.L. conceived the concept and supervised the entire project. S. F. and W. S. developed the deep learning-based algorithms. Y.L. and T.L. developed the GPU based vectorial PSF fitter. W.S. developed the field dependent PSF calibration software. J.Y., Y.L., Y.H., Z.H., Z.Y. and Z. G. designed the optical setup. W.S., L.Z., C.Y. and C. L. prepared the U2OS and COS7 cell samples. J. L., X. L., and M. Z. contributed the induced neuronal cells from mESCs. W.S. and L. Z. labeled the cells and acquired the data. Y.H. and S.W. set up the microscope hardware. S.F., S.W. and T.L. analyzed the data. Y.L., S.F., W.S., J.R., P.X. and D.J. wrote the manuscript with the input from all other authors.

## Data availability

All data are available upon reasonable request from the corresponding authors.

## Code availability

Source code for the software used in this manuscript is contained in Supplementary Software 1 and updated versions can be freely downloaded at: https://github.com/Li-Lab-SUSTech/FD-DeepLoc

## Extended Data figure

**Extended Data Fig. 1.**
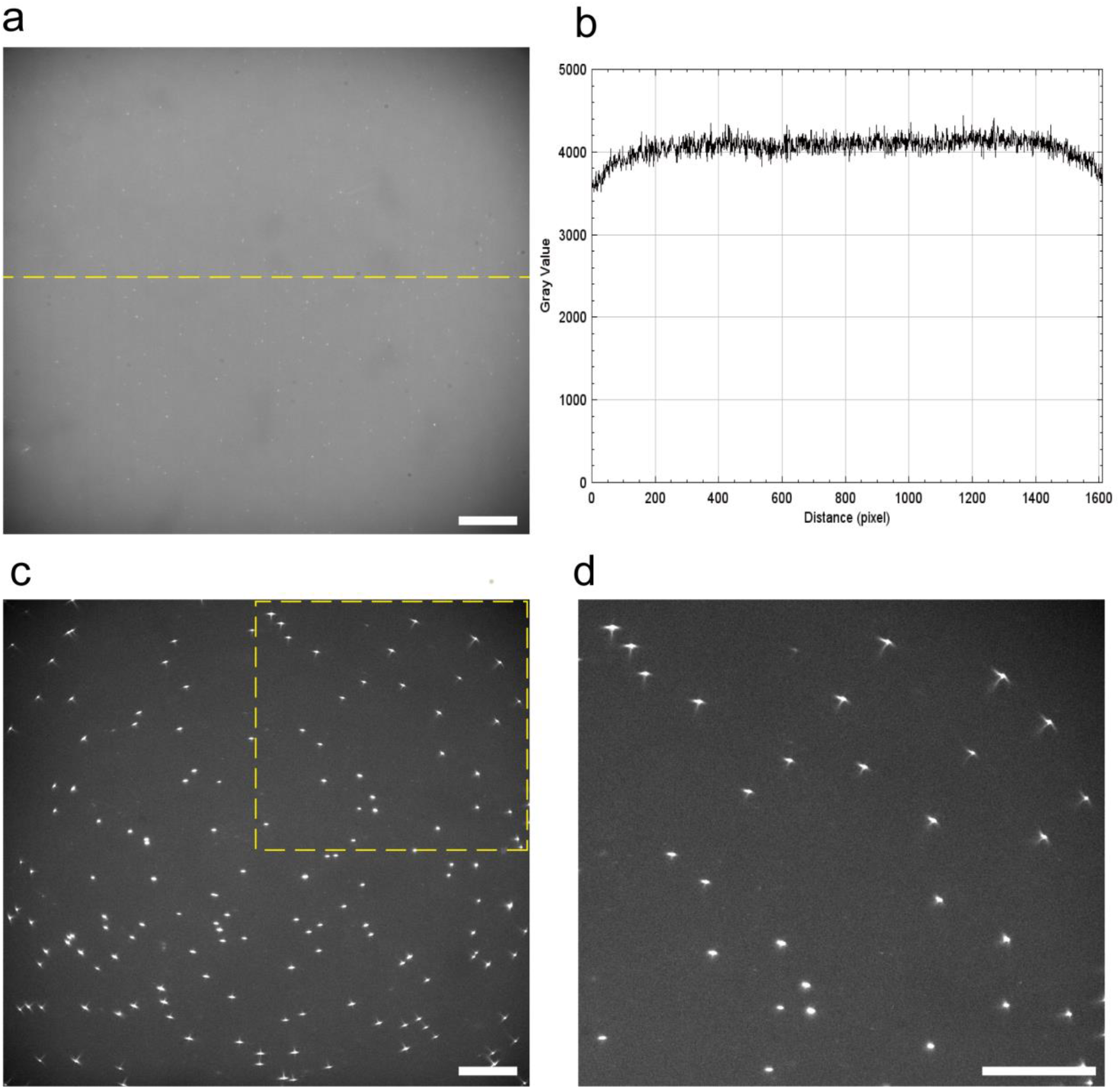
Uniform illumination of large FOV. **a**, Full frame (1608 ×1608 pixels) imaging of BG-AF647 dye solution ( S9136S, New England Biolabs, 1:1000 dilution from stock in Milli-Q water)on coverslip. **b**, Intensity profile of the yellow dashed line in **a**. **c**, Example image of beads (T7279, Invitrogen, TetraSpeck) on coverslip (defocused by 300 nm) under uniform illumination. **d**, Zoomed image of the yellow rectangle indicated in **c**. Scale bars, 20μm.

**Extended Data Fig. 2.**
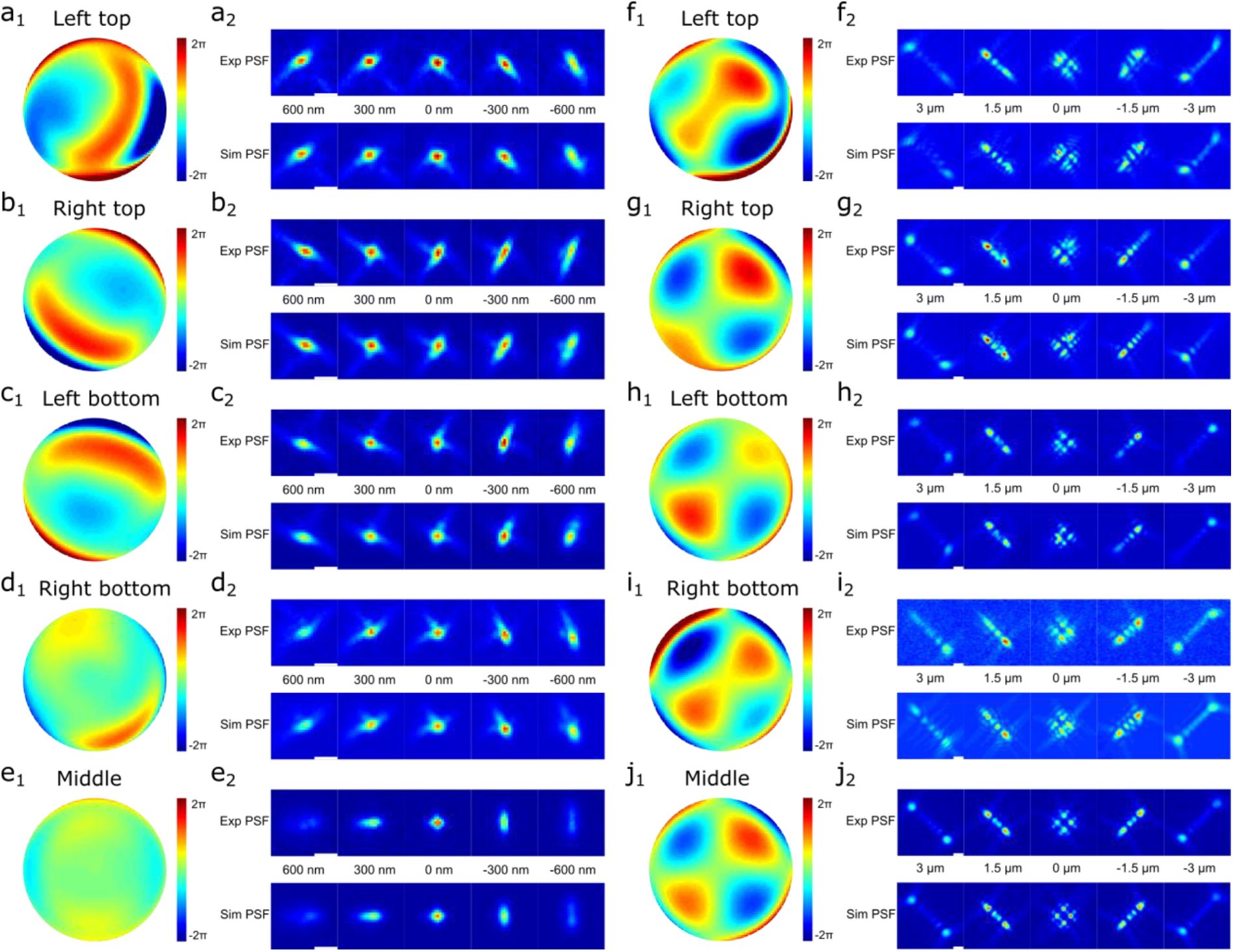
Comparison of experimental and fitted PSF models across the FOV. Experiment and fitted astigmatism PSF models (**a-e**) and DMO Tetrapod PSF models (**f-j**) acquired from different positions in the full frame image (1608×1608 pixels). **a1**, Fitted pupil function of bead stacks locate at the left top position (79, 136). **a2**, Comparison of the experimental and fitted PSF model within ± 600 nm axial range. **b-e** are the same as **a**, but with different locations: **b** right top (1417, 22); **c** left bottom (246, 1588); **d** right bottom (1439, 1503); **e** middle (812, 805). **f-j** are experiment and fitted DMO Tetrapod PSF models within ± 3 μm axial range acquired from different positions: **f** left top (139, 75); **g** right top (1329, 206); **h** left bottom (177, 1422); **i** right bottom (1444, 1577); **j** middle (855, 810). Scale bars, 1 μm.

**Extended Data Fig. 3.**
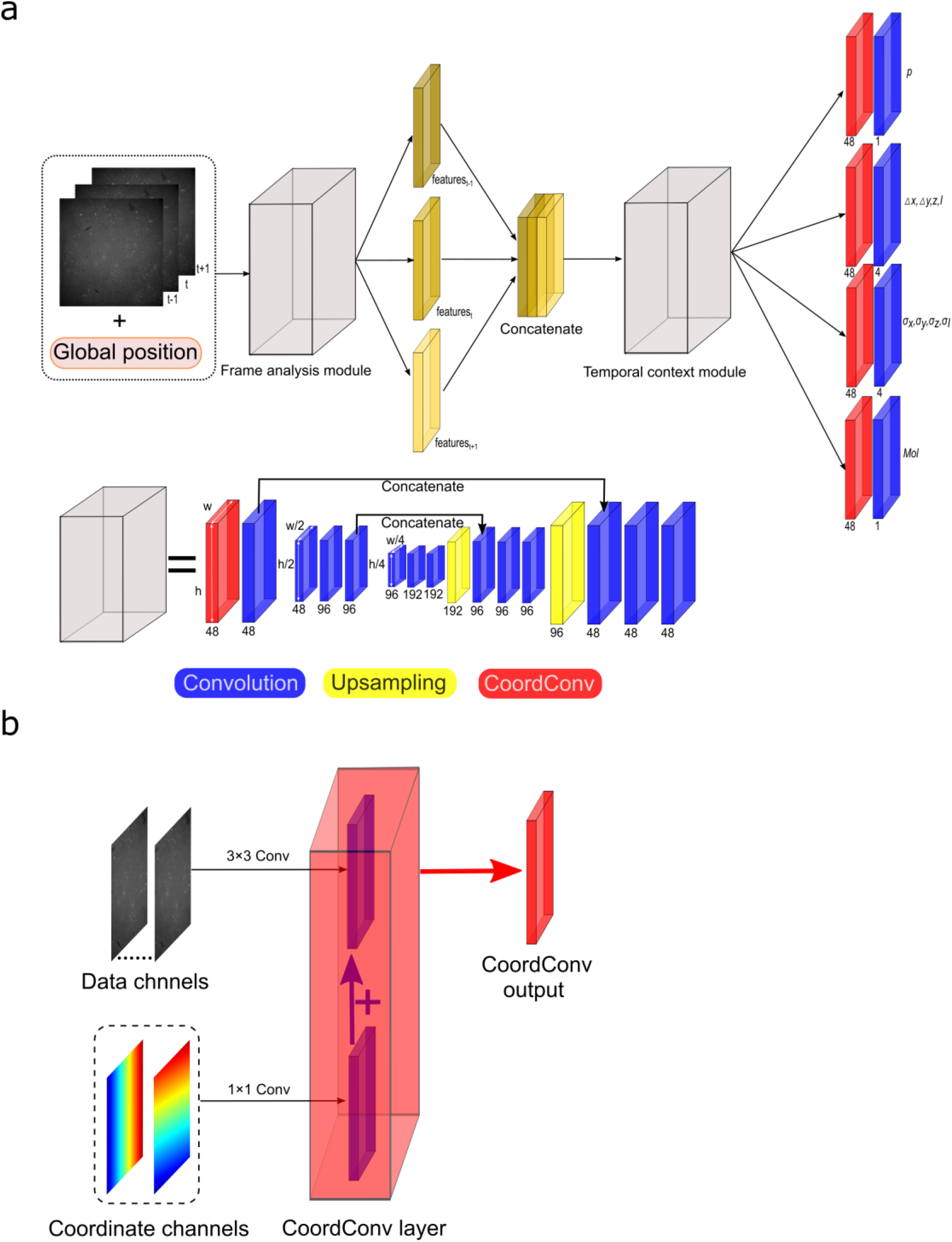
FD-DeepLoc architecture. **a**, Overall network architecture of FD-DeepLoc. Similar to the DECODE network, FD-DeepLoc contains two U-Nets as indicated by the gray box. For each input frame, features will be extracted by the frame analysis module and then concatenated together to the temporal context module. Finally, a ten-channel map will be output to form the emitter predictions of each frame. The output of each layer is depicted with colorful blocks, where *h* and *w* represent the height and width of the channels. **b**, CoordConv channels. The CoordConv layer is placed in the first convolution layer of each module and has two convolution operations. One convolution (3 × 3) is operated on the input image. The other one (1 × 1) is operated on two coordinate channels. The results are then added as the output of the Coordconv layer.

**Extended Data Fig. 4.**
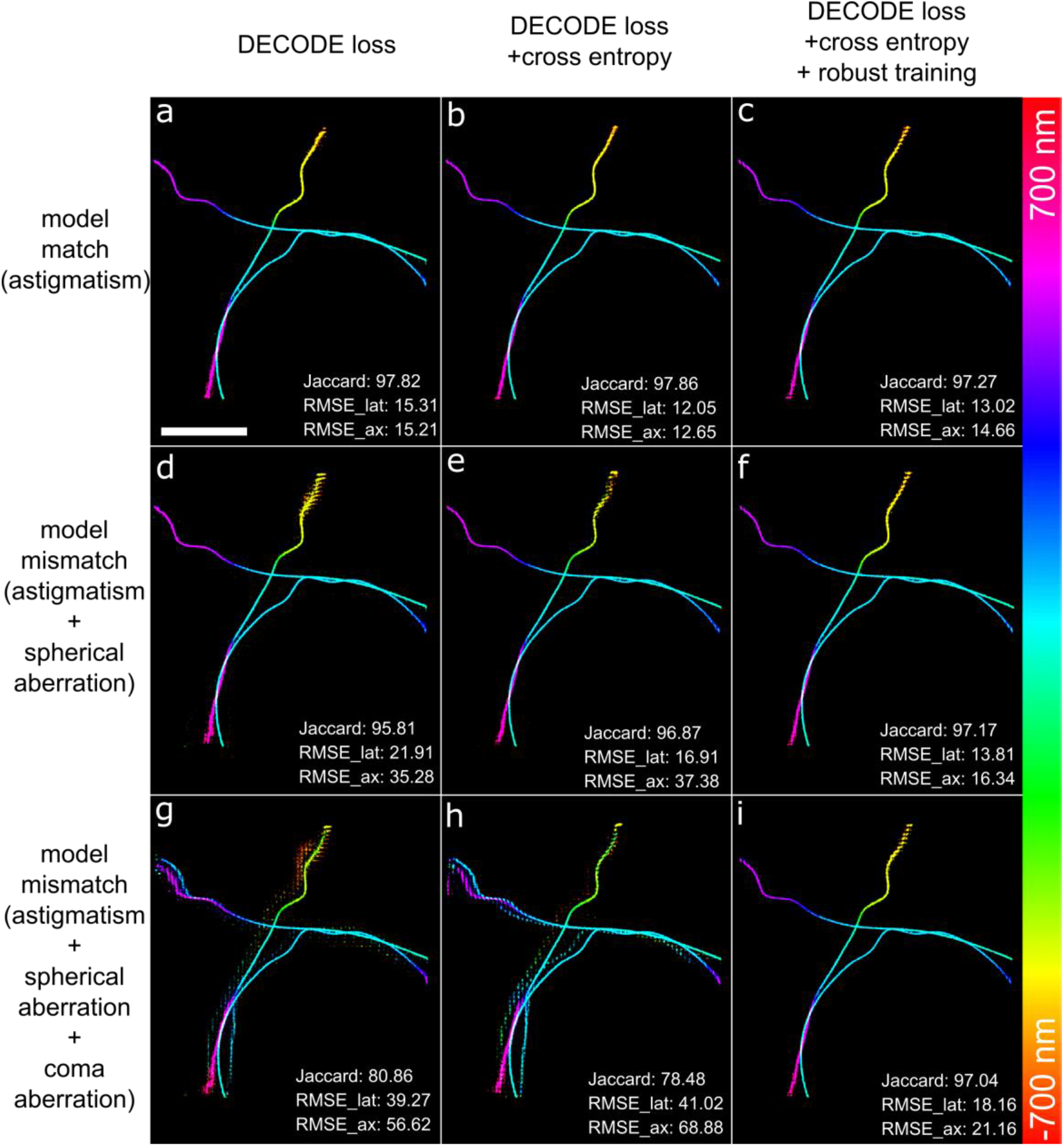
Effects of PSF model mismatch between training and test dataset. The training dataset is generated by PSF model with astigmatism aberration. Reconstructed images for test data using PSF models with astigmatism aberration (**a, b, c**), astigmatism plus spherical aberration (**d, e, f**) and astigmatism plus spherical and coma aberration (**g, h, i**) are shown. The performance of network with different loss function are compared: original DECODE loss function (**a, d, g**), original DECODE loss function plus cross entropy term (**b, e, h**) and original DECODE loss function plus cross entropy term and robust training (**c, f, i**). For robust training, small random aberrations were added (**Methods**). The signal to background ratio for test data is set as 5500 average photons per emitter (sampled from a uniform distribution within [1000, 10000]) and 50 background photons per pixel. The rms wavefront error of spherical aberration and coma aberration are both set as 20 nm (λ = 660 nm). The ground truth coordinates are taken from the SMLM challenge training dataset MT0^29^. Scale bar, 2 μm.

**Extended Data Fig. 5.**
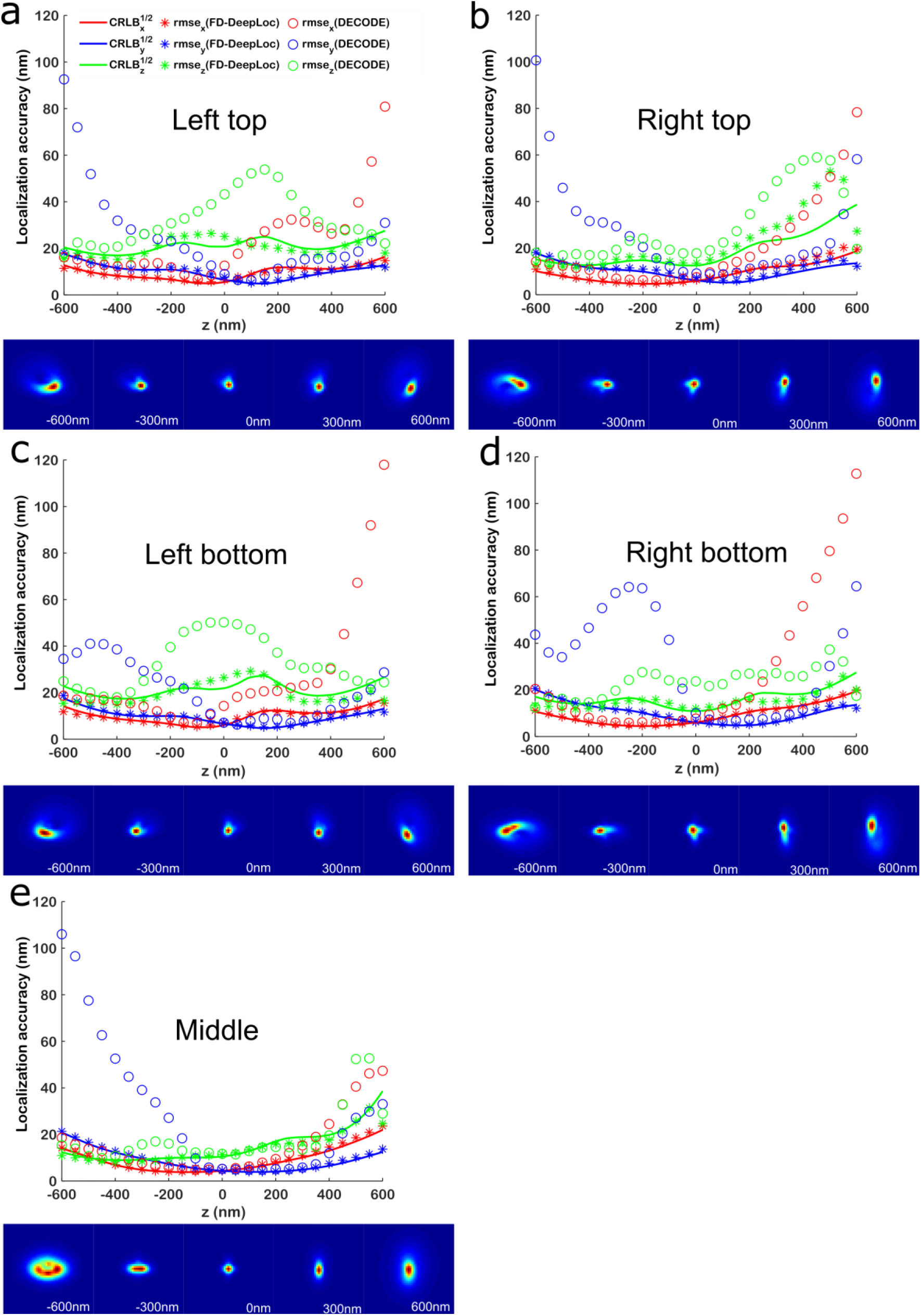
Comparison of localization accuracy for single molecules with field dependent aberrations analyzed by FD-DeepLoc and DECODE. **a**, Localization accuracy of a left top single molecule at different axial positions analyzed by FD-DeepLoc and DECODE. Both networks were trained using the same training data containing all spatially variant PSFs. The bottom panel of **a** shows the corresponding 3D PSF. **b-e** are the same as a, but at different locations: **b** right top; **c** left bottom; **d** right bottom; **e** middle. The data is simulated using the aberration maps in **Supplementary Fig. 6**. 5,000 photons and 50 background were used for each single molecule. 3,000 single-emitter images are generated with random x, y positions at each axial position (x, y is random within a pixel, z is random within a 50 nm step, r.m.s.e is averaged over 50 nm bins).

**Extended Data Fig. 6.**
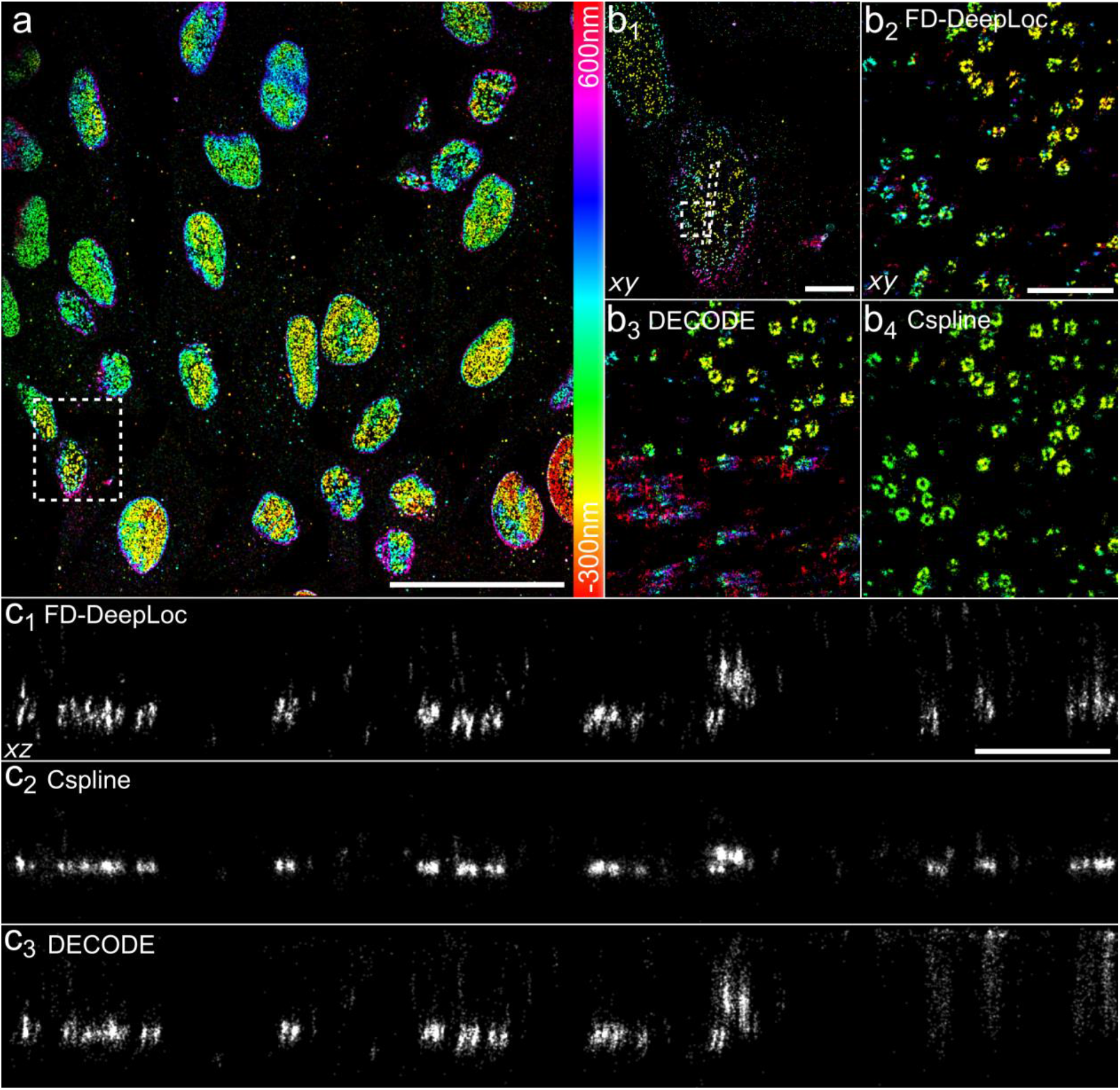
FD-DeepLoc enables large FOV 3D super-resolution imaging of nuclear pores with high fidelity. **a**, Overview of the panoramic 3D super-resolution image of Nup96-SNAP-Alxea Fluor 647. **b**, Top views of the NPCs reconstructed with FD-DeepLoc, DECODE and Cspline. **b1**, Zoomed view of the region indicated by the white dashed box in **a**. **b2**, **b3** and **b4** are the zoomed images of the rectangle region indicated in **b1** reconstructed by FD-DeepLoc, DECODE and Cspline separately. **c**, Side view images of the region bounded by the 500 nm dashed lines in **b1** reconstructed by FD-DeepLoc, Cspline and DECODE separately. Representative results are shown from 3 experiments. Scale bars, 50 μm (**a**), 5 μm (**b1**,) and 1 μm (**b2**, **c1**).

**Extended Data Fig. 7.**
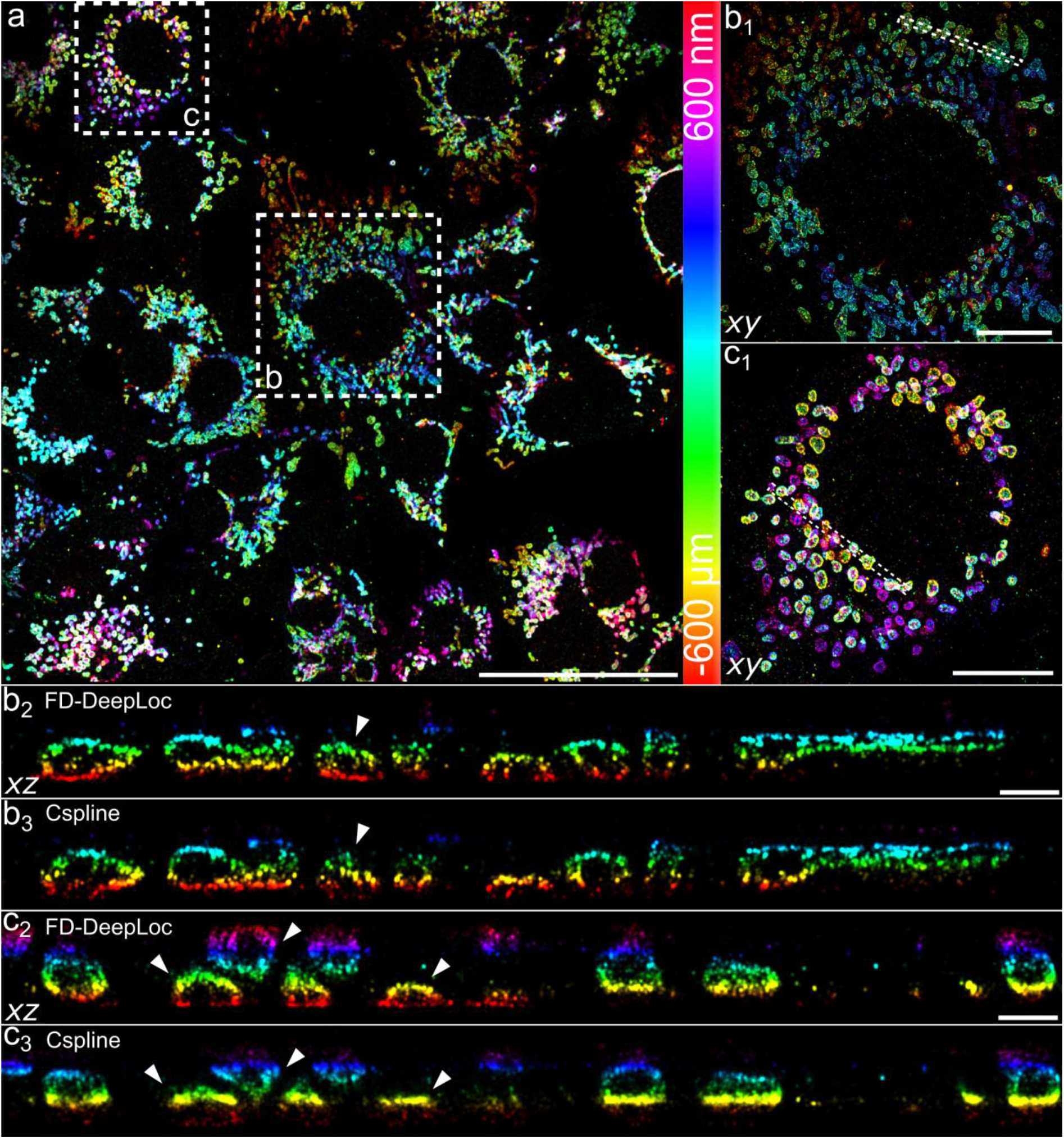
Performance of FD-DeepLoc on 3D astigmatism imaging of mitochondria in a large FOV. **a**, Overview of the panoramic 3D super-resolution image of immunolabeled TOM 20 in mitochondria. **b1, c1**, Zoomed images of the center and marginal areas as indicated by the dashed boxes **b, c** in **a**. **b2, b3**, Side-view cross-section of the region bounded by the dashed lines in **b1** reconstructed by FD-DeepLoc and Cspline, separately. **c2, c3** the same as **b2, b3**, but for the region bounded by the dashed lines in **c1**. Representative results are shown from 2 experiments. Scale bars, 50μm (**a**), 10μm (**b1**, **c1**) and 1μm (**b2**, **c2**).

**Extended Data Fig. 8.**
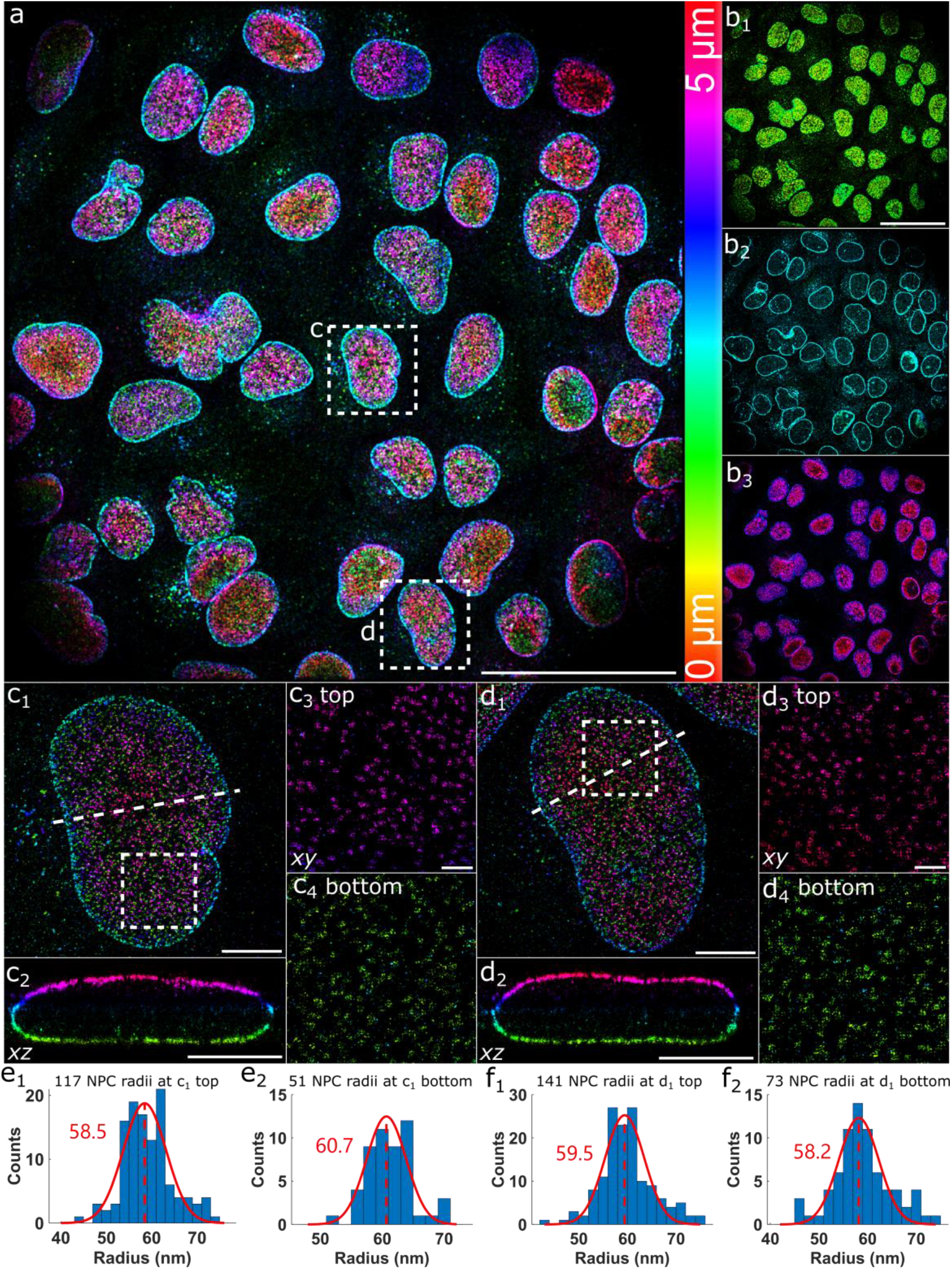
FD-DeepLoc enables large FOV whole cell 3D super-resolution imaging of NPC. **a**, Overview of the panoramic whole cell 3D super-resolution image of NPC. **b**, Different axial sections of **a**. **b1-3** are top views of axial range of (0 μm - 2 μm), (2 μm - 3 μm) and (3 μm – 5 μm), respectively. **c**, Magnified views of areas denoted by box c in **a**. **c1**, Top view of the box c area denoted in **a**. **c2**, Side-view cross-section of region denoted by the dashed line in **c1**. **c3-4** are zoomed views of the top and bottom surface of the boxed area denoted in **c1**. **d**, Same as **c** for magnified views of region denoted by box d in **a**. **e**, The distribution of the fitted radii of the NPCs for the cell in **c**. **e1** and **e2** are for the top and bottom nucleus surface, respectively. **f** is the same as **e**, but for the cell in **d**. Representative results are shown from 7 experiments. Scale bars, 50 μm (**a**, **b1**), 5 μm (**c1**, **c2**, **d1**, **d2**) and 1 μm (**c3**, **d3**).

**Extended Data Fig. 9.**
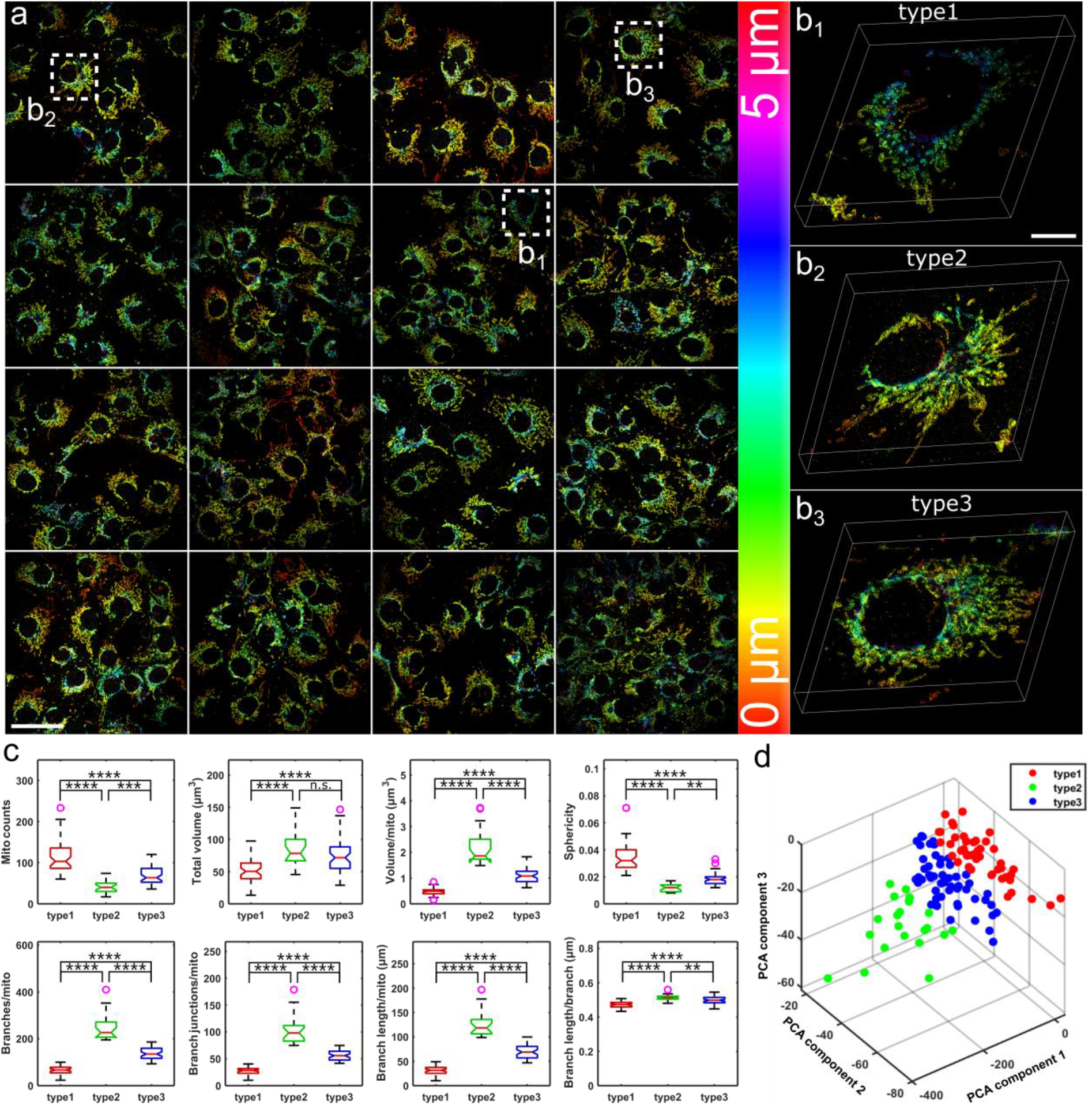
Quantitative analysis of whole cell 3D super-resolved mitochondria images. **a**, Overview of 16 full frame images of whole cell 3D super-resolved mitochondria. **b**, Magnified angled views of 3 representative cells denoted by the boxes in **a**. **c**, Quantitative analysis of mitochondria morphology and network connectivity using Mitochondria Analyzer^36^. The boxplots illustrate the median (horizontal red lines in each box), interquartile range (extent of each box), adjacent values (vertical extending lines denote the most extreme values within 1.5 interquartile range), notch (the variability of the median) and outlier values (purple circle) for each group of shape parameters. n.s., not significant. \*\**P*<0.01, *** *P*<0.001, and \*\*\*\**P*<0.0001 (one-way ANOVA with Sidak post hoc test, one-sided). Statistical significance was set at a threshold of *P*<0.05. **d**, Principle component analysis was performed on the features used in **c** with the first 3 principle components plotted, color indicates the categorization results of 121 cells. The numbers of cells in type1/2/3 are 47/22/52, respectively. Scale bars, 50 μm (**a**) and 10 μm (**b1**).

**Extended Data Fig. 10.**
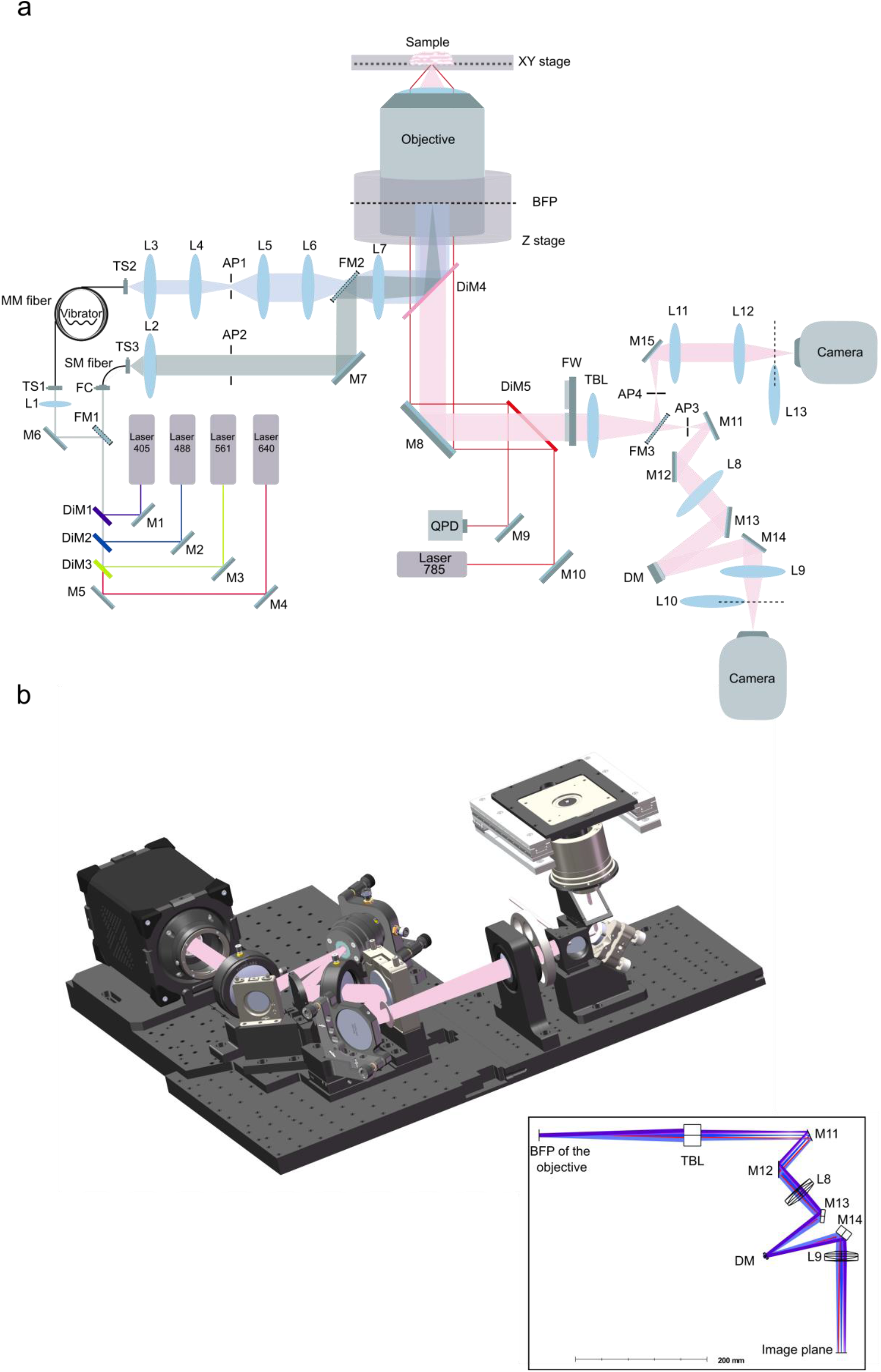
Layout of custom-built microscope used for this study. M: mirror, DiM: dichroic mirror, FM: flip mirror, L: lens, TS: translation stage, FC: fiber coupler, MM fiber: multi-mode fiber, SM fiber: single-mode fiber, AP: aperture, BFP: back focal plane, FW: filter-wheel, TBL: tube lens, QPD: quadrant photodiode, DM: deformable mirror. **a**, Optical path of the microscope. Our setup has two excitation options: single-mode (SM) excitation and multi-mode (MM) excitation. In this work, we mainly use the MM mode for large FOV excitation. Excitation lasers are firstly reflected by dichroic mirrors DiM1-DiM3 (DMLP 425/505/605, Thorlabs) and then coupled by lens L1 into a multi-mode fiber (WFRCT200×200- 230×230-440-620-1100N, NA = 0.22, CeramOptec). The fiber is bound with a vibrator to generate a homogeneous illumination ^8,46^. There are two imaging paths for the fluorescence detection which is separated by a flip mirror FM3. In the reflection path of FM3 (KSHM90, Owis), cylindrical lens (LJ1516L1-A, Thorlabs) based astigmatism 3D super-resolution imaging was performed. For the other imaging path without FM3, DM engineered PSFs were used for 3D super-resolution imaging. The focal length for each lens: L1 (f = 19 mm, Ø0.5 inch), L2 (f = 75 mm, Ø1 inch), L3 (PLN 10X/NA0.25, Olympus), L4(f = 400 mm, Ø1 inch), L5 (f = 35 mm, Ø1 inch), L6 (f = 400 mm, Ø75 mm), L7 (f = 400 mm, Ø75 mm), TBL (f = 180 mm, TTL180-A, Thorlabs), L8 (f = 150 mm, Ø2 inch), L9 (f = 150 mm, Ø2 inch), L10 (f = 75 mm, Ø1 inch), L11 (f = 150 mm, Ø2 inch), L12 (f = 150 mm, Ø2 inch), L13 (f = 1000 mm, LJ1516L1-A, Thorlabs). **b**, The rendered mechanical design using SolidWorks (Dassault Systèmes). Inset is the ray-tracing diagram of the imaging path rendered in OpticStudio (Zemax).

