## Supplementary Information for "Field dependent deep learning enables high-throughput whole-cell 3D super-resolution imaging"

### Supplementary materials

| | |
|---|---|
| **Supplementary Figure 1** | Field-dependent aberration maps for DMO Tetrapod PSF induced by DM |
| **Supplementary Figure 2** | The relative root square error of the fitted PSF model |
| **Supplementary Figure 3** | Comparison of the speed of vectorial PSF simulation using CPU and GPU |
| **Supplementary Figure 4** | Effects of robust training on experimental data of 3D super-resolution imaging by DMO saddle point PSF |
| **Supplementary Figure 5** | Effect of non-uniform background simulation |
| **Supplementary Figure 6** | Aberration maps for simulated datasets |
| **Supplementary Figure 7** | Schematic of the artificial structures and example images of the evaluation set |
| **Supplementary Figure 8** | Fourier ring correlation (FRC) analysis of the large FOV 3D super-resolution image of multiple NPCs reconstructed using different algorithms |
| **Supplementary Figure 9** | Raw beads and single molecule images for DMO Tetrapod PSF |
| **Supplementary Figure 10** | Fourier ring correlation (FRC) analysis of whole cell 3D super-resolution image of mitochondria within a large FOV (180 × 180 μm$^2$) and DOF (5 μm) |
| **Supplementary Figure 11** | 3D CRLB of different experimental PSFs |
| **Supplementary Figure 12** | Schematic of the imaging formation in our vectorial PSF model. |
| **Supplementary Figure 13** | Contribution of cross entropy and CoordConv to the overall performance of FD-DeepLoc |
| **Supplementary Figure 14** | Impact of the performance of FD-DeepLoc based on training with/without per-pixel readout noise (RN) simulation |
| **Supplementary Note 1** | Vectorial PSF model |
| **Supplementary Note 2** | Evaluation set generation |
| **Supplementary Note 3** | Evaluation metrics |
| **Supplementary Note 4** | NPC radii analysis and mitochondria morphology analysis |
| **Supplementary Note 5** | FD-DeepLoc details, including training process and inference process |
| **Supplementary Note 6** | Data postprocessing and rendering |
| **Supplementary Movie 1** | A tour view animation of the large-FOV 3D super-resolution image of NPC in Fig. 4 |
| **Supplementary Movie 2** | A tour view animation of the large-FOV 3D super-resolution image of mitochondria in Extended Data Fig. 7 |
| **Supplementary Movie 3** | Panoramic view animation of the large-FOV and DOF 3D super-resolution image of mitochondria in Fig. 5 |

| | |
|---|---|
| **Supplementary Movie 4** | Animation of a single cell in the large-FOV and DOF 3D super-resolution image of mitochondria in Fig. 5 |
| **Supplementary Movie 5** | Panoramic view animation of the large-FOV and DOF 3D super-resolution image of NPC in Extended Data Fig. 8 |
| **Supplementary Movie 6** | Animation of a single cell in the large-FOV and DOF 3D super-resolution image of NPC in Extended Data Fig. 8 |
| **Supplementary Movie 7** | A tour view animation of the large-FOV and DOF 3D super-resolution image of neurites in Fig. 6 |
| **Supplementary Code** | Example and source code for FD-DeepLoc |

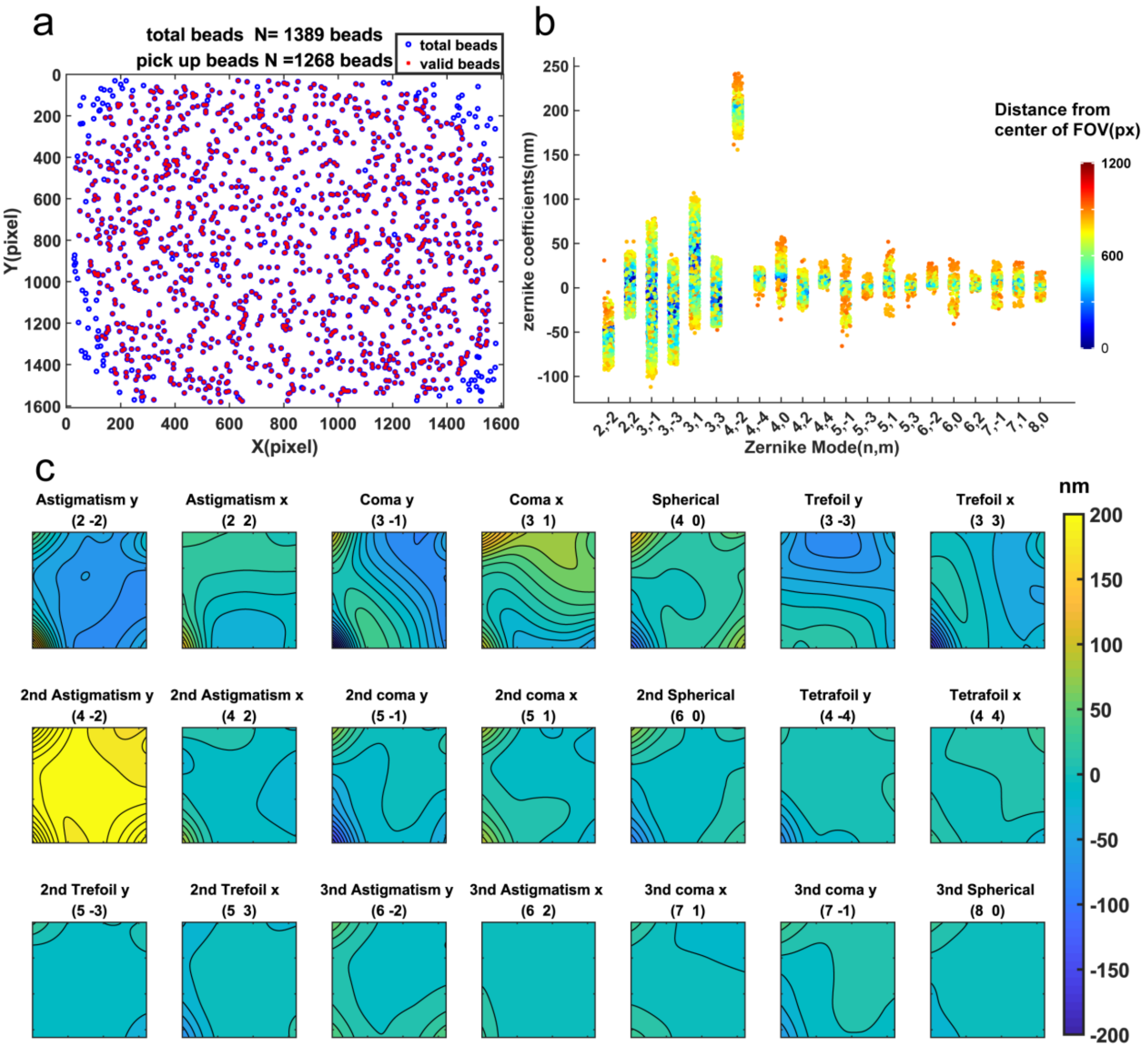


**Supplementary Figure 1. Field-dependent aberration maps for DMO Tetrapod PSF induced by DM. a,** The distribution of the beads used for aberration map calibration (summed from 80 sets of beads stacks). Blue circles denote all beads collected and red crosses denote the beads after filtering. **b,** The relationship between aberration magnitude and the distance to the center of the FOV. Color denotes the distance to the center of the FOV. **c,** The interpolated 21 Zernike aberration maps of our microscope after load the control matrix of DMO Tetrapod PSF to the DM. The contour interval is 16 nm.

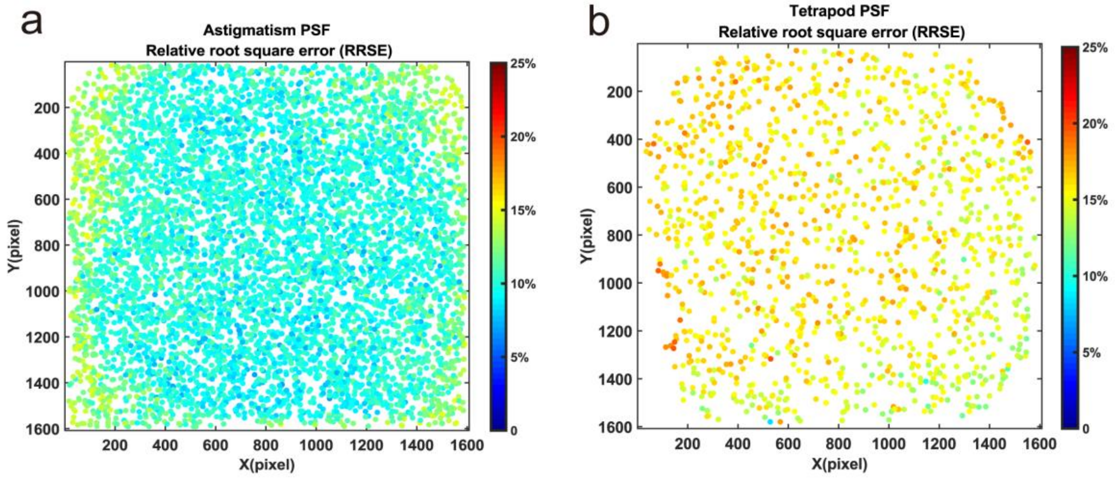


**Supplementary Figure 2. The relative root square error of the fitted PSF model.** The relative root square error (RRSE) between data and the fitted model across the full frame image (1608×1608 pixels). **a,** RRSE of astigmatism PSF across the full frame image. **b,** RRSE of Tetrapod PSF across the full frame image.

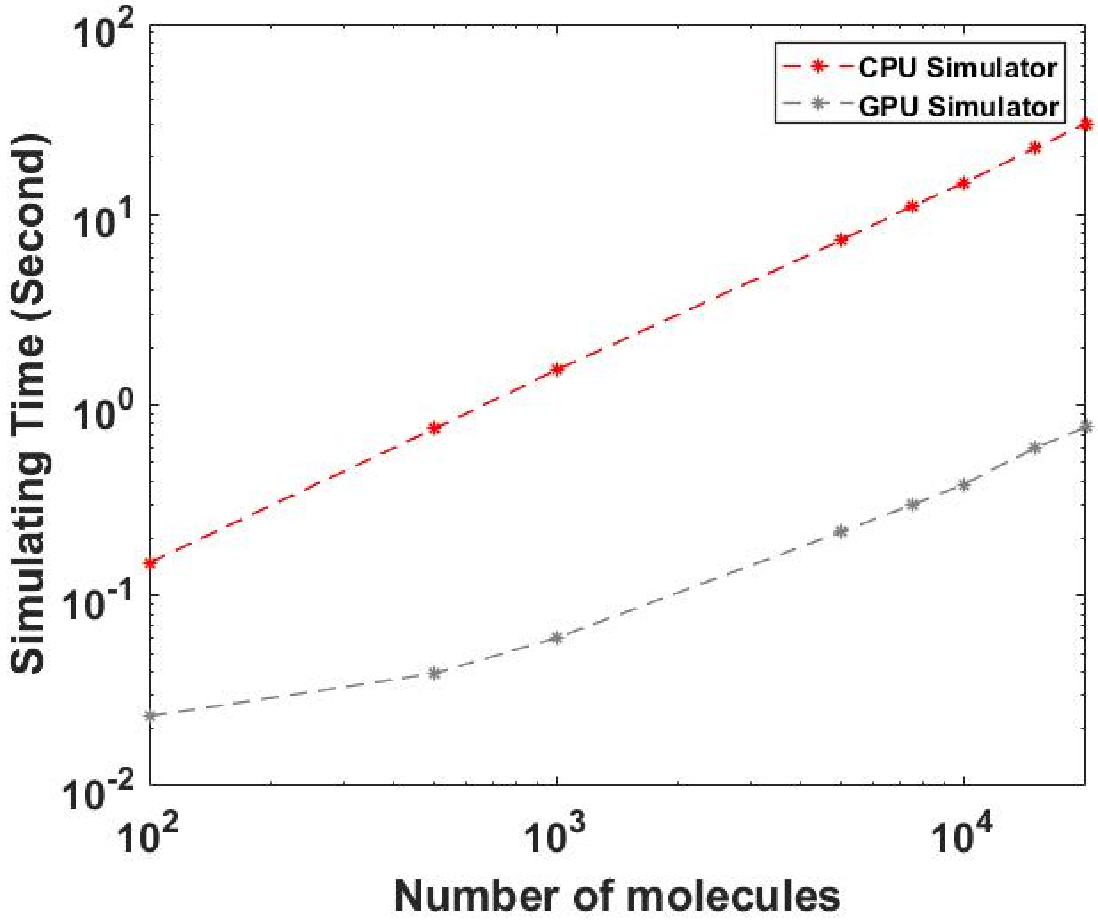


**Supplementary Figure 3. Comparison of the speed of vectorial PSF simulation using CPU and GPU.** Molecules with ROI size of 27×27 pixels were simulated. The speed evaluation was performed on an i9-9900 CPU and a RTX3090 consumer graphics card running CPU and GPU, separately.

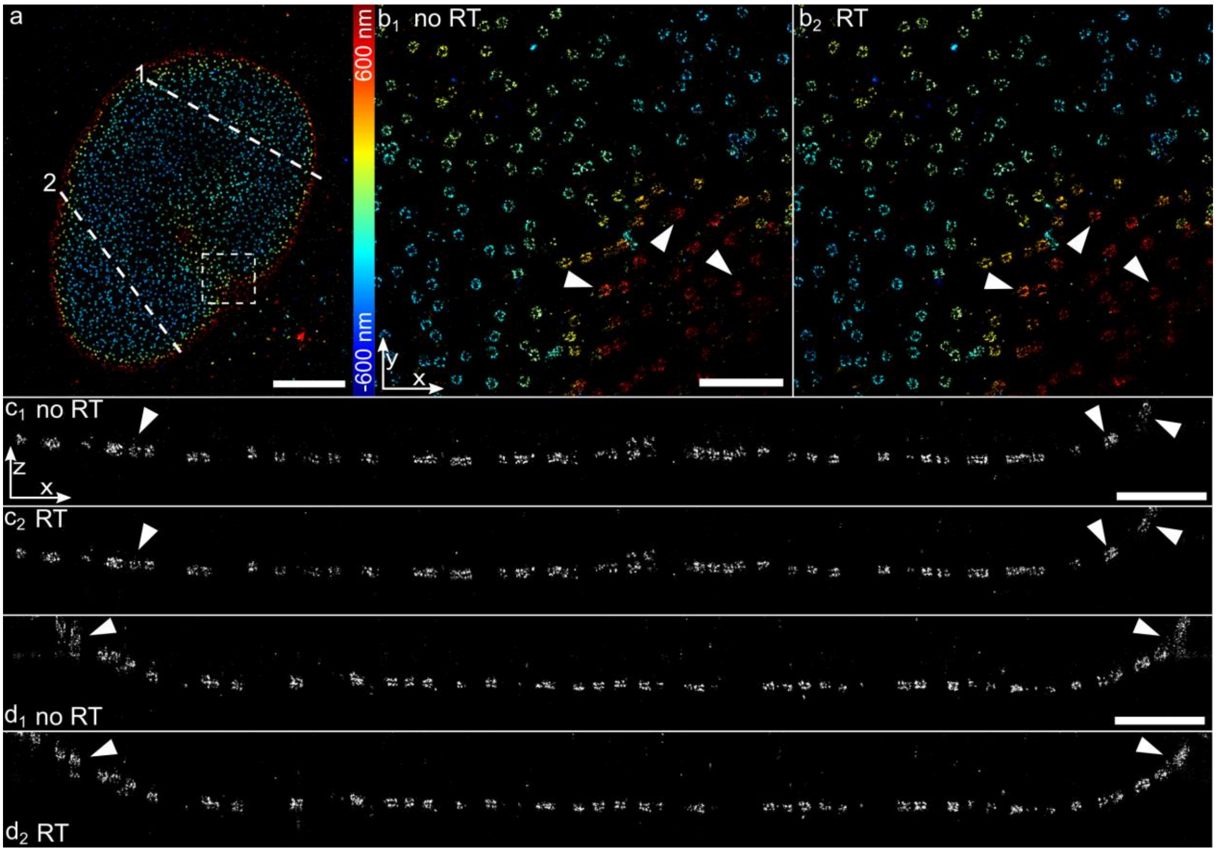


**Supplementary Figure 4. Effects of robust training on experimental data of 3D super-resolution imaging by DMO saddle point PSF. a,** 3D super-resolution imaging of Nup96-SNAP-AF647 reconstructed by FD-DeepLoc. **b**, Magnified images of the region as indicated in **a** reconstructed by the FD-DeepLoc without (**b1**) and with (**b2**) applying the robust training strategy (**Methods**). **c,** Comparison of the side view of the regions denoted by line 1 (500 nm width) in **a**. Images are reconstructed without robust training (**c1**) and with robust training (**c2**) separately. **d**, the same as **c** but along line 2 (500 nm width) denoted in **a**. Representative results are shown from 3 experiments. Scale bars, 5 μm (**a**) and 1 μm (**b, c, d**).

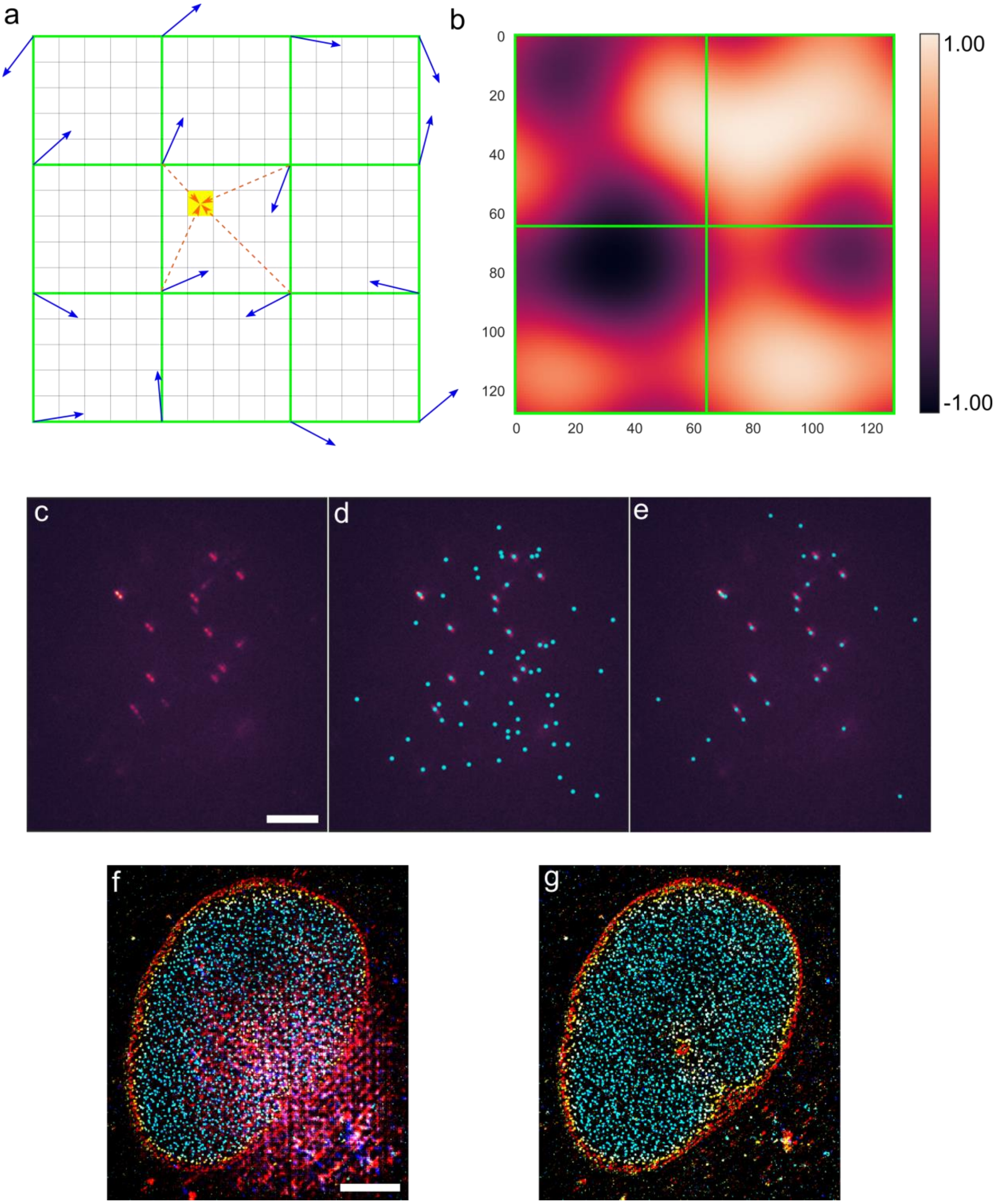


**Supplementary Figure 5. Effect of non-uniform background simulation. a,** Principle of Perlin noise. First, several image pixels (black) are used to form a grid (green). For each grid corner, a unit gradient vector with random orientation is assigned (blue arrows). To determine the value of a specific pixel (yellow), the distance vectors from four nearest grid corners are calculated (orange arrows). Then the dot products of the distance and gradient vectors are computed and assigned to each grid corner. To make the final background noise map smooth and continuous, the coordinates of the yellow pixel in a grid (every grid pixel size is normalized to 1) are converted to a new faded coordinates by a fade function: $f(t) = 6t^5 - 15t^4 + 10t^3$, where $t$ is the yellow pixel coordinates. The yellow pixel value is defined as the bilinear interpolation of four grid corner values on the faded coordinates. The number of image pixels in each grid pixel is the frequency of the noise. **b,** An example Perlin noise map with a frequency of 64 image pixels and an octave number of 1, the value is normalized to $[-1,1]$. **c,** An example 3D SMLM raw frame (1.2 μm DMO PSF). **d** and **e** are the localization results of **c**, predicted by the network trained with constant background and non-uniform background, respectively. The probability threshold is set as 0.7. **f** and **g** are the final reconstructed images corresponding to **d** and **e**. Representative results are shown from 3 experiments. Scale bar, 5 μm (**c**, **f**).

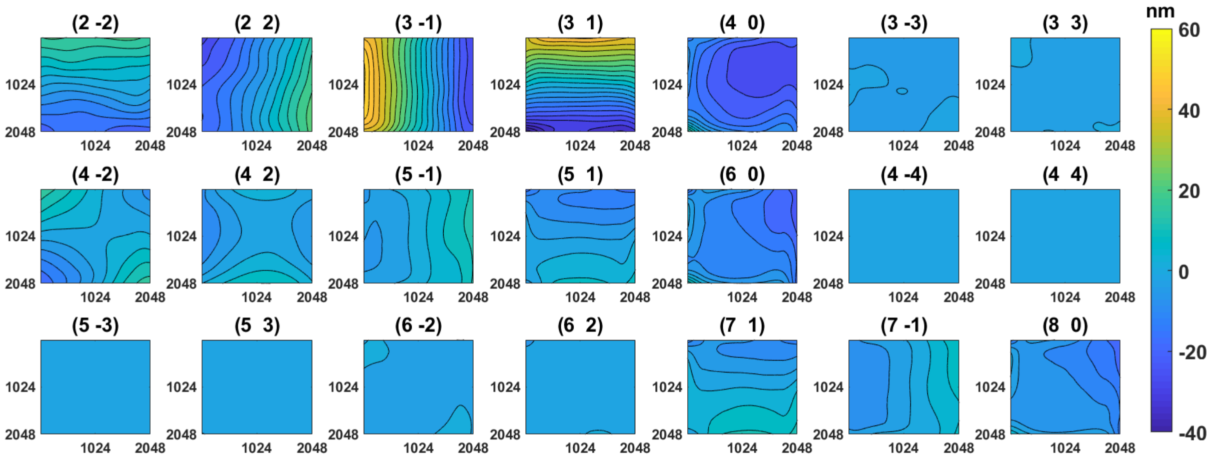


**Supplementary Figure 6. Aberration maps for simulated datasets.** The aberration maps were calibrated in a microscope equipped with a deformable mirror (DM140A-35-P01, Boston Micromachines). DM was set as flat using the flat control matrix provided by the manufacture during aberration calibration. The aberration maps were interpolated to 2048×2048 pixels and defined as Normal aberration map for simulation test in **Fig. 3**. Strong aberration maps were equal to twice magnitude of the Normal aberration maps. For astigmatism PSF simulation, an additional constant astigmatism with rms of 70 nm was added to the (2 ,2) Zernike aberration map. The contour interval is 4 nm.

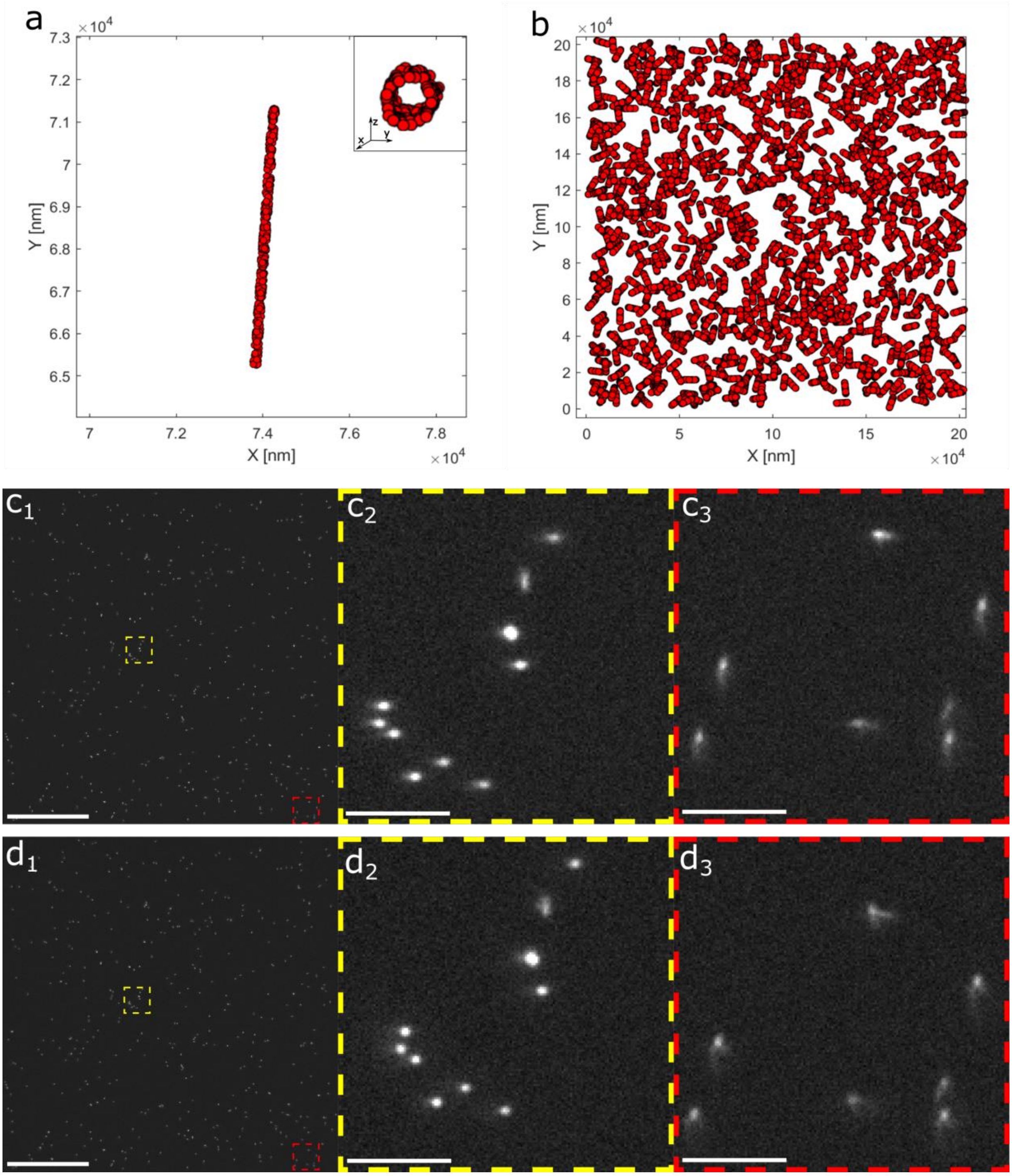


**Supplementary Figure 7. Schematic of the artificial structures and example images of the evaluation set. a,** The schematic of the simulated hollow rod with a radius of 50 nm and length of 6,000 nm. **b,** 1,000 hollow rods were generated in a 204.8×204.8×1.4 μm$^3$ space as the ground truth structures. **c,** An example image of the evaluation set (10,000 photons per emitter, and 100 constant background per pixel) generated using the Gradient 1 aberration maps, **c2** and **c3** are zoomed views of the region indicated by the yellow and red dashed boxes in **c1** respectively. **d,** the same as **c**, but generated using the Gradient 2 aberration maps, which change more quickly across the FOV. Scale bars 50 μm (**c1, d1**), 5μm (**c2, c3, d2, d3**).

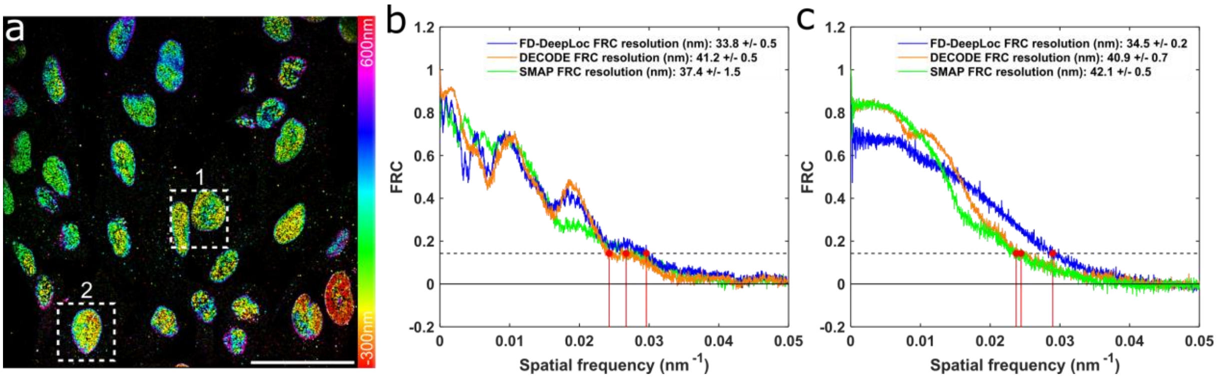


**Supplementary Figure 8. Fourier ring correlation (FRC) analysis of the large FOV 3D super-resolution image of multiple NPCs reconstructed using different algorithms. a,** Large FOV 3D super-resolution image of NPCs. **b** and **c** are FRC curves correspond to areas denoted by dashed box **1** and box **2** in **a** reconstructed by different algorithms, respectively. Representative results are shown from 3 experiments. Scale bar, 50 µm (**a**).

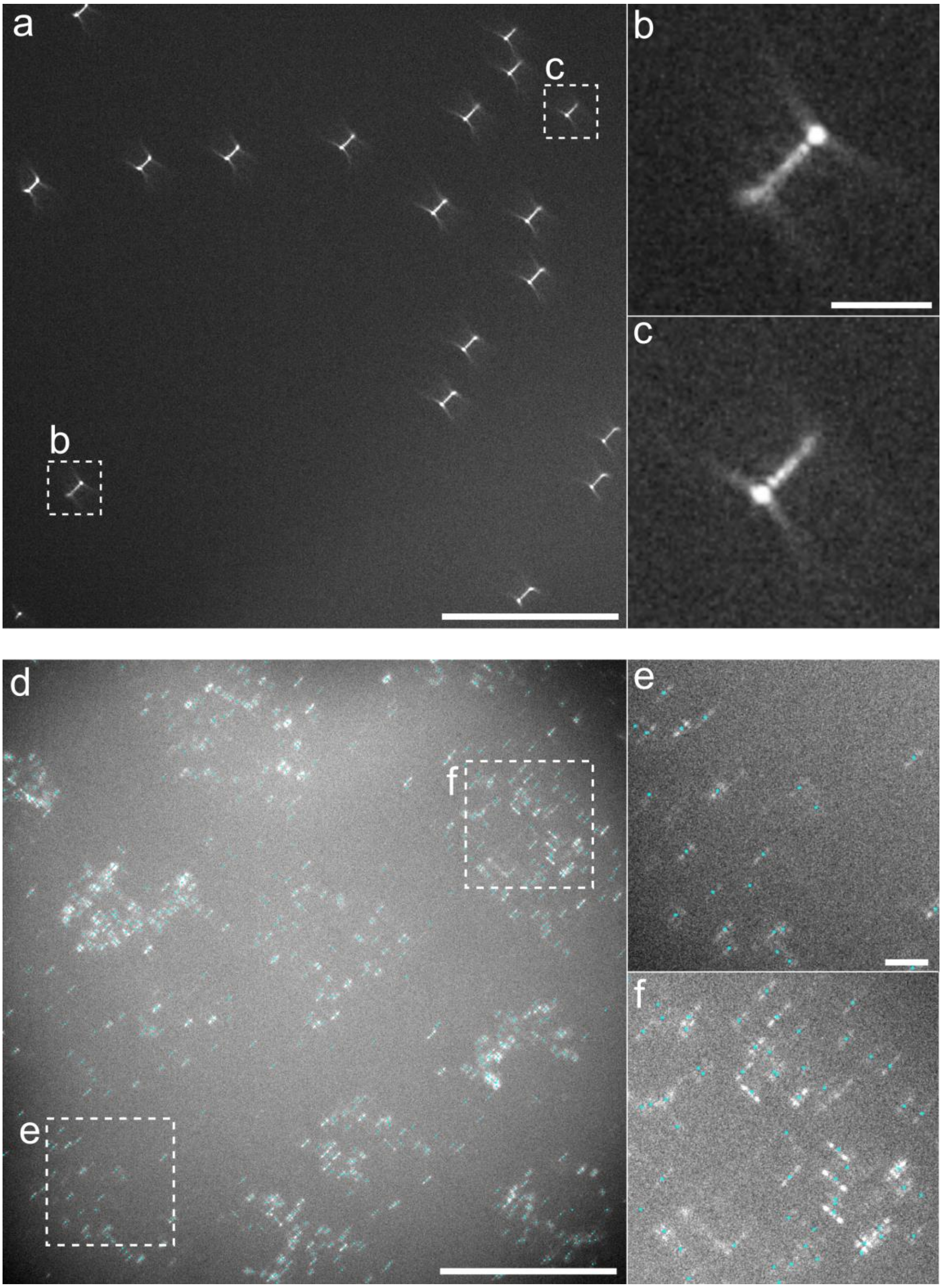


**Supplementary Figure 9. Raw beads and single molecule images for DMO Tetrapod PSF (6 µm).** **a,** Beads images for Tetrapod PSF. **b** and **c** are the magnified images of corresponding boxed area in **a**. **d**, Single molecule image of mitochondria labeled by anti-Tom 20 antibodies. **e** and **f** are the magnified images of corresponding boxed area in **d**. Cyan filled circles are the pixel-level localizations predicted by FD-DeepLoc. Scale bars 50 µm (**a, d**), 5 µm (**b, e**).

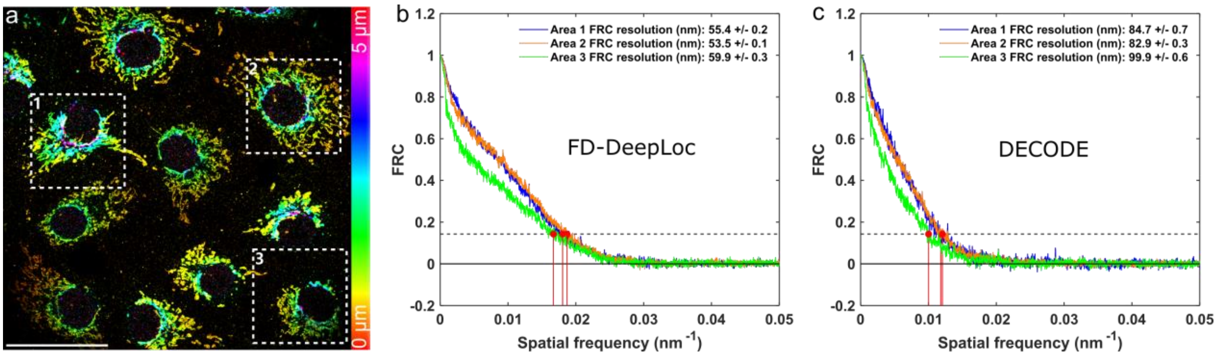


**Supplementary Figure 10. Fourier ring correlation (FRC) analysis of whole cell 3D super-resolution image of mitochondria within a large FOV ($\mathbf{180 \times 180\ \mu m^2}$) and DOF ($\mathbf{5\ \mu m}$).** **a,** Whole cell 3D super-resolution image of mitochondria over large FOV. **b** and **c** are the FRC analysis of the box regions **1, 2** and **3** in **a** reconstructed by FD-DeepLoc and DECODE, respectively. Representative results are shown from 7 experiments. Scale bar, 50 µm (**a**).

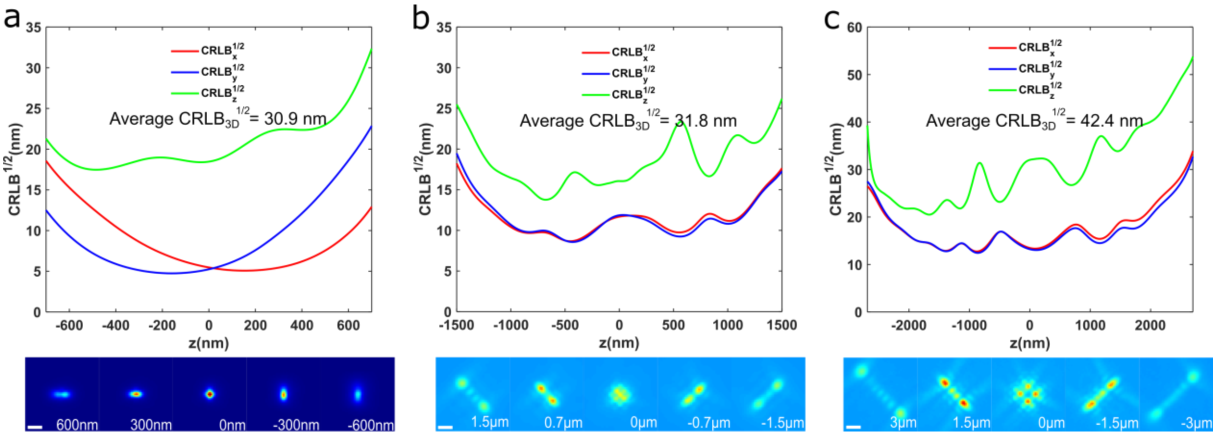


**Supplementary Figure 11. 3D CRLB of different experimental PSFs. a**, astigmatic PSF; **b**, 3 μm DMO PSF; **c**, 6 μm DMO PSF. We used 5000 photons and 50 background photons for CRLB calculation. Scale bars, 1 μm.

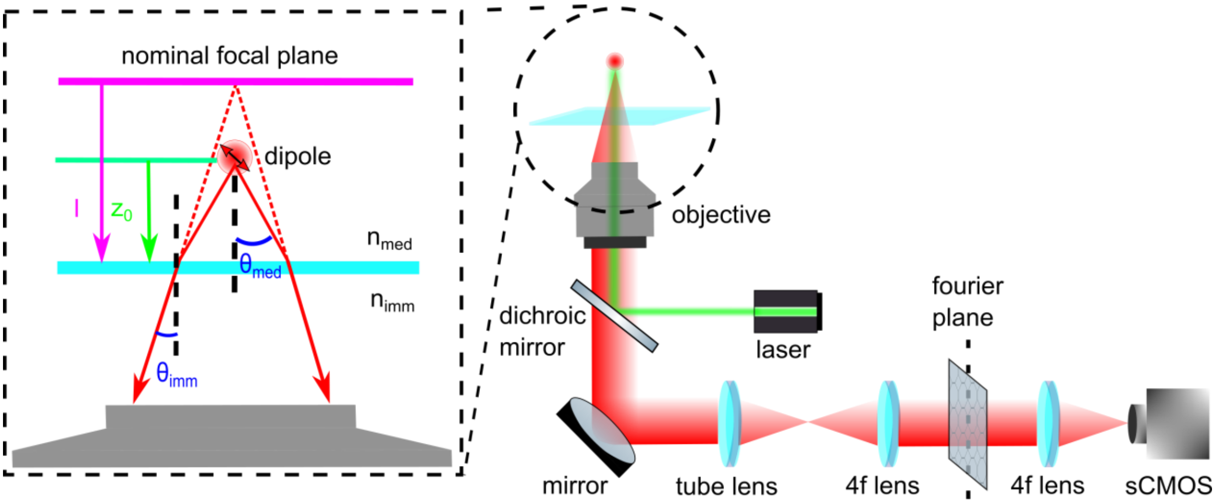


**Supplementary Figure 12. Schematic of the imaging formation in our vectorial PSF model.**

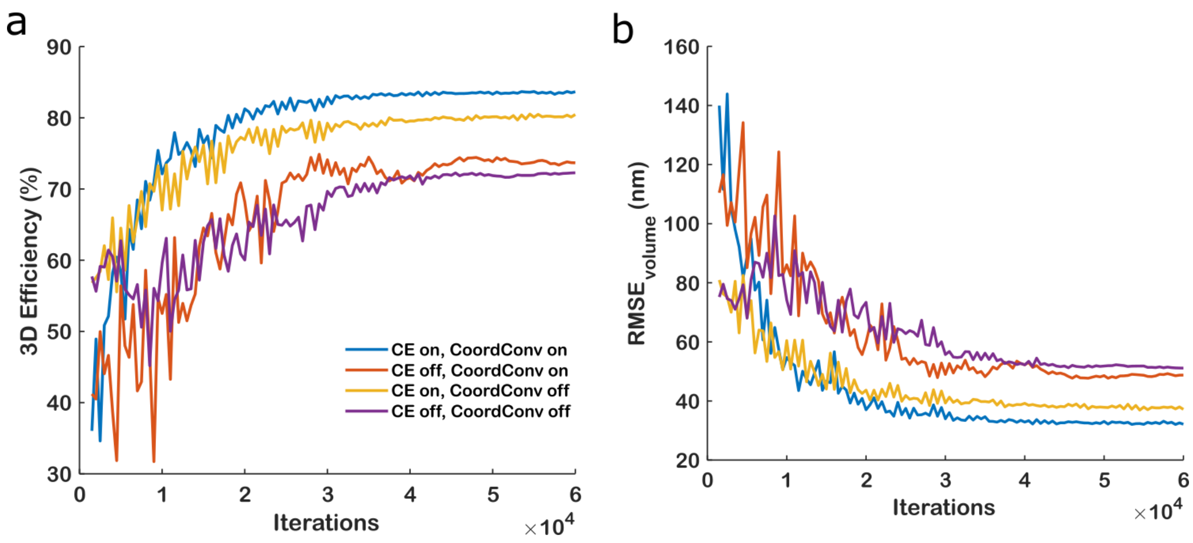


**Supplementary Figure 13. Contribution of cross entropy and CoordConv to the overall performance of FD-DeepLoc.** The network was trained on the dataset with Normal field-dependent aberration and medium SNR (**Supplementary Note 2**). The performance was recorded every 500 iterations.

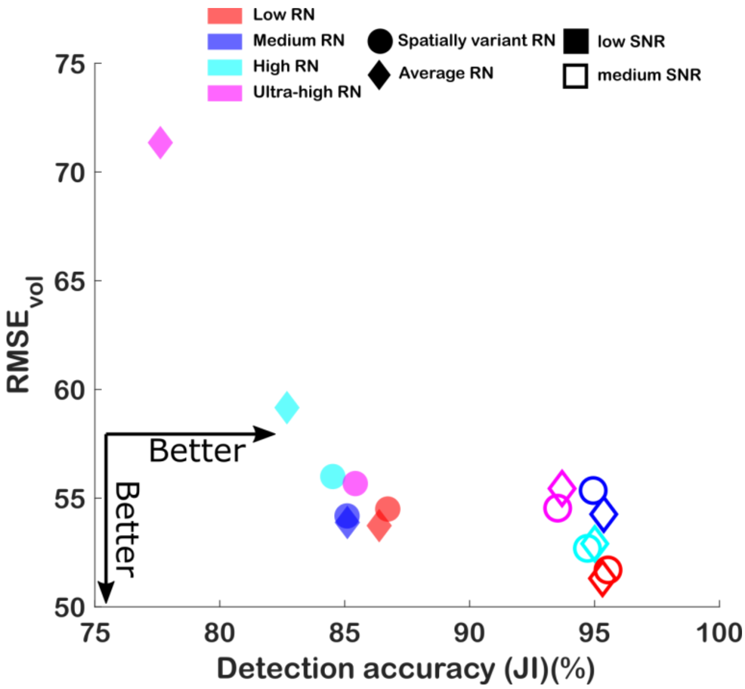


**Supplementary Figure 14. Impact of the performance of FD-DeepLoc based on training with/without per-pixel readout noise (RN) simulation.** 4 types of readout noise (low, medium, high, ultra-high) and 2 types of SNRs (low, medium) were simulated for evaluation. For each simulated dataset, 2 networks were trained. One was trained with averaged RN and the other one was trained with pixel-dependent RN. Here, we assumed that the distribution of the pixel-dependent RN for the low RN case follows Gaussian distribution with a mean of 2.3e- and a standard deviation of 0.53e-, which corresponds to a commercial sCMOS camera (Prime 95B, Photometrics, USA). To generate RN map with higher readout noise, we add hot pixels with high readout noise to the map. For the medium/high/ultra-high RN dataset, 5%/10%/10% pixels of the low RN map was added with an extra readout noise which is a Gaussian distribution with a mean of 5e-/10e-/20e- and a shared sigma of 1e-, respectively.

**Supplementary Note 1 Vectorial PSF model**

Since polarization exhibit non-negligible effects for optical system with a high numerical aperture (NA) objective, a vectorial PSF model accommodating for refractive index mismatch between medium-cover slip interface and cover slip-immersion medium interface was used[1]. Here, we modeled the PSF of an isotropic emitter as the fluorescent probes are normally attached to the molecules of interest flexibly and can rotate or wobble freely. Therefore, the PSF are treated as the summed image of three orthogonal dipoles. To further consider the field dependent aberrations, the PSF model can be written as:

$$PSF \propto \sum_{p=x,y} \sum_{d=x,y,z} \left| \mathcal{F}_{2D}\left\{ A(\rho,\varphi) \cdot e^{i\psi_{aber}(\rho,\varphi;x_0,y_0)} \cdot e^{i\psi_{pos}(\rho,\varphi;x_0,y_0,z_0)} \cdot E_{p,d}^{BFP} \right\} \right|^2 \otimes Gauss_{2D}(\sigma_x,\sigma_y) \quad \text{(S1)}$$

where $\mathcal{F}_{2D}$ denotes 2D Fourier transform, ($\rho$, $\varphi$) is the normalized polar coordinate in the back focal plane (BFP) whereas $\rho_{max}$ corresponds to limiting aperture angle $NA/n_{imm}$. Additional 2D Gaussian blurring is also applied to account for the high-spatial-frequency attenuation[2], and the standard deviations of the Gaussian kernel are determined experimentally by fitting to a real beads data. $A(\rho,\varphi)$ is the amplitude function with aplanatic correction factor. ($x_0$, $y_0$, $z_0$) is the 3D position of the point source in the sample plane. $\psi_{aber}$ represents the field-dependent aberrations and is expressed as a liner sum of variance normalized Zernike polynomials:

$$\psi_{aber} = \sum_{n+|m|\le 8} C_n^m(x_0,y_0) \cdot Z_n^m(\rho,\varphi) \quad \text{(S2)}$$

where $C_n^m(x_0,y_0)$ is the aberration coefficients at $(x_0,y_0)$, $Z_n^m$ is the normalized Zernike polynomial, $n$ is the radial order and $m$ is the angular frequency. The phase shift correlated with the emitter position and objective position $\psi_{pos}$ is defined as:

$$\psi_{pos} = \frac{2\pi}{\lambda}\left( NAx_0\rho \cos\varphi + NAy_0\rho \sin\varphi + n_{med} z_0 \sqrt{1-(\frac{\rho NA}{n_{med}})^2} - n_{imm} l \sqrt{1-(\frac{\rho NA}{n_{imm}})^2} \right) \quad \text{(S3)}$$

Here, $n_{med}$, $n_{cov}$ and $n_{imm}$ are the refractive indexes of sample medium, cover glass and objective immersion fluid, respectively. $z_0$ represents the emitter's height above cover glass. $l$ is the distance between the nominal focal plane and the cover glass. $E_{p,d}^{BFP}$ represents the polarization vector with components $p = x, y$ in the image plane and each dipole components $d = x, y, z$ in the sample contributes to components $p$:

$$E_{x,d}^{BFP} = T_p \vec{P}_d \cos\varphi - T_s \vec{S}_d \sin\varphi \quad \text{(S4a)}$$

$$E_{y,d}^{BFP} = T_p \vec{P}_d \sin\varphi + T_s \vec{S}_d \cos\varphi \quad \text{(S4b)}$$

$$with \left( \vec{P} = \begin{bmatrix} \cos\theta_1 \cos\varphi \\ \cos\theta_1 \sin\varphi \\ -\sin\theta_1 \end{bmatrix}, \vec{S} = \begin{bmatrix} -\sin\varphi \\ \cos\varphi \\ 0 \end{bmatrix} \right). \quad \text{(S4c)}$$

Here $\vec{P}$ and $\vec{S}$ are the basis polarization vectors. $T_p$ and $T_s$ are the Fresnel transmission coefficients for p and s polarization. The transmission loss is computed for each refractive index boundary:

$$T_p = T_{p,med-cov} \times T_{p,cov-imm} \qquad \text{(S5a)}$$

$$T_s = T_{s,med-cov} \times T_{s,cov-imm} \qquad \text{(S5b)}$$

With

$$T_{p,\mathrm{med-cov}} = \frac{2n_{\mathrm{med}}\, cos\, \theta_{\mathrm{med}}}{n_{\mathrm{med}}\, cos\, \theta_{\mathrm{cov}} + n_{cov}\, cos\, \theta_{\mathrm{med}}} \qquad \text{(S6a)}$$

$$T_{p,\mathrm{cov-imm}} = \frac{2n_{\mathrm{cov}}\, cos\, \theta_{\mathrm{cov}}}{n_{\mathrm{cov}}\, cos\, \theta_{\mathrm{imm}} + n_{imm}\, cos\, \theta_{\mathrm{cov}}} \qquad \text{(S6b)}$$

$$T_{s,\mathrm{med-cov}} = \frac{2n_{\mathrm{med}}\, cos\, \theta_{\mathrm{med}}}{n_{\mathrm{med}}\, cos\, \theta_{\mathrm{med}} + n_{\mathrm{cov}}\, cos\, \theta_{\mathrm{cov}}} \qquad \text{(S6c)}$$

$$T_{s,\mathrm{cov-imm}} = \frac{2n_{\mathrm{cov}}\, cos\, \theta_{\mathrm{cov}}}{n_{\mathrm{cov}}\, cos\, \theta_{\mathrm{cov}} + n_{\mathrm{imm}}\, cos\, \theta_{\mathrm{imm}}} \qquad \text{(S6d)}$$

For the numerical implementation of **Equation (s1)**, chirp *z*-transform was used to replace the 2D Fourier transform which allows breaking the relationship of the sampling points between imaging space and Fourier space.

**Supplementary Note 2 Evaluation set generation**

To quantitatively evaluate how well FD-DeepLoc could perform on SMLM data with field dependent aberrations, we utilized TestSTORM[3] to simulate 1,000 hollow rods randomly distributed in a 204.8*204.8*1.4 $\mu m^3$ three-dimensional space, with each rod has a radius of 50 nm and length of 6,000 nm. For each rod, it has 320 epitopes to link dye labels. The length of the linker is 7 nm. The bonding angle of dye labels follows a normal distribution with a mean of 0° and a standard deviation of 30°. After getting the molecular positions, we incorporated the photo physics activation model from the SMLM challenge[4] to determine which molecule and when the molecules will blink. The average on-time and dark-time are set as 3 frames and 2.5 frames, respectively. The evaluation sets contain three types of SNR: low, medium and high SNR, each with a mean of 1,000/5,000/10,000 photons per emitter, and 10/50/100 constant background per pixel, respectively. The spatially variant aberration maps in **Supplementary Fig. 5** were used, including Normal and Strong field-dependent aberrations. An additional astigmatism aberration $Z_2^2$ with an amplitude of 70 nm rms (λ=660nm) was added to the aberration maps to engineer the 3D PSF.

After convolving the molecular positions with the PSFs, we incorporated the noise model from SMLM challenge[4]. For the EMCCD camera data, the noise model mainly contained the shot noise, excess noise and camera readout noise. The final value of pixel $k$ ($x_k$) in simulated training image is given by:

$$x_k = \frac{\mu_{2,k}}{e_{ADU}} + BL \quad (S7)$$

where,

$$\mu_{2,k} = Gamma\left(\mu_{1,k}, EM_{gain}\right) + Gauss(0, \sigma_R) \quad (S8)$$

and

$$\mu_{1,k} = Poisson\left(QE \cdot \left(\mu_{0,k} + BG\right) + c\right) \quad (S9)$$

Here, $\mu_{0,k}$ is the expected number of photons in the $k_{th}$ pixel, given by the spatially variant vectorial PSF model described before. *QE* is the quantum efficiency. *c* is the spurious charge, and *BG* is either a constant value or a nonuniform background with Perlin noise. Shot noise is added by the function of $Poisson$ to form $\mu_{1,k}$. Then $\mu_{1,k}$ signal goes through an electron multiplication process, where the excess noise is described by a Gamma distribution with scale parameter $EM_{gain}$. The camera readout noise $Gauss(0, \sigma_R)$ follows a zero-mean Gaussian distribution with standard deviation $\sigma_R$ . The final pixel value $x_k$ is calculated by **Equation s7**, where $e_{ADU}$ and *BL* are analog-to-digital conversion factor and camera baseline, respectively. For the sCMOS camera data, there is no electron multiplication process. Only shot noise and Gaussian readout noise were considered. Although the gain and readout noise of the sCMOS camera are pixel-dependent, we found using constant $e_{ADU}$ and $\sigma_R$ (1.61 $e^-$ was used in this work) does not affect the result too much.

**Supplementary Note 3 Evaluation metrics**

To quantify the performance of FD-DeepLoc on the simulated large-FOV SMLM data, Jaccard Index(JI) and Root Mean Squared Error(RMSE) were used. JI measures the degree that an algorithm could accurately recognize the blinking emitters and avoid taking non-emitter spots for predictions:

$$JI = \frac{TP}{TP + FP + FN} \quad (S10)$$

where TP (True Positive) is the number of predicted emitters that can be matched to the ground truth emitters, FP (False Positive) is the number of predicted emitters that can not be matched to the ground truth emitters, FN (False Negative) is the number of ground truth emitters that are not matched by the predictions. The matching threshold for lateral displacement is 250 nm, while the axial matching threshold is 500 nm. To further quantify how close the TP predictions are to the ground truth positions, RMSE is used to measure the error between the predicted emitter positions and the ground truth positions:

$$RMSE = \sqrt{\frac{1}{TP}\sum_{i}^{TP}(\hat{x}_i - x_i^{GT})^2 + (\hat{y}_i - y_i^{GT})^2 + (\hat{z}_i - z_i^{GT})^2} \quad (S11)$$

where $\hat{x}_i, \hat{y}_i, \hat{z}_i$ are the coordinates of the predicted $TP$ emitter $i$. $x_i^{GT}, y_i^{GT}, z_i^{GT}$ are the corresponding ground truth.

**Supplementary Note 4 NPC radii analysis and mitochondria morphology analysis**

We used the workflow provided in Ref. 9 to automatically segment and analyze the NPCs in **Extended Data Fig. 8**. The rendered super-resolution image was first convolved with a Gaussian ring kernel, with the diameter range set as 90 nm – 130 nm and cutoff value set as 0.04. Then the local maxima are treated as candidates and passed through a clean-up process. All localizations of each candidate were fitted to a circle and discarded if the fitted radius was too small (< 40 nm) or too large (> 70 nm). Next, the rest candidates were re-fitted with a circle with fixed radius (55 nm), and rejected if more than 30% of localizations were within 45 nm or more than 75% of localizations were more than 65 nm away from the NPC center. The radii of these filtered NPC candidates were finally fitted using a circular model with central coordinates and radius as free fitting parameters.

To extract the morphology parameters of mitochondria, we combined the pipeline of the ImageJ plugin Mitochondria Analyzer[10] with FD-DeepLoc. It can be summarized as three simple steps: 1) Analyze the raw data and export the molecule list by FD-DeepLoc; 2) Import the molecule list into the SMAP for rendering; 3) Export 3D super-resolution image stacks by SMAP and process these image stacks using the batch command of Mitochondria Analyzer in ImageJ.

In this work, we rendered the images using the “constant Gaussian” mode with a Gaussian size of 20 nm in SMAP. The intensity mode was set as “photons”. The contrast mode was set as “quantile = -2”, which means $10^{-2}$ proportion of pixels are saturated. Considering the FRC resolution of the 3D whole-cell mitochondria reconstruction is around 50 nm (**Supplementary Figure 10**), the image stacks were rendered with a lateral pixel-size of 50 nm and a z-step of 100 nm. During the mitochondria analysis, we skipped the pre-process step of “Subtract Background” and “Sigma Filter Plus” as the input localization-based image is very sharp and has little fluorescence background. We used an “Enhance Local Contrast (CLAHE)” value of 10 and “Adjust Gamma” value of 0.1 instead of the default value (1.5 and 0.9 respectively). This is because localization-based image has many localization clusters that should belong to continuous structure which is different from that for wide-field image. Although the present segmentation method is enough for the current analysis, a better pre-process method could be designed specifically for SMLM images for more precise feature extraction in the future. For the SMLM image thresholding, we set the “block size” as 1.25 and “C-value” as 1. The post-processing commands “Remove Outliers” and “Fill 3D Holes” were used in our analysis. The thresholded images were then analyzed by the Mitochondria Analyzer with default settings. The final output was a series of feature parameters describing the mitochondria morphology (counts, volume, sphericity, *etc.*) and network connectivity (branches, branch junctions, branch length, etc.). We further applied the built-in k-means++ clustering algorithm in MATLAB to classify these feature vectors into three groups. A k-value of 3 was set empirically and the clustering was repeated for 200 times to find a lowest sum distance.

**Supplementary Note 5 FD-DeepLoc details, including training process and inference process**

For the training process, FD-DeepLoc employs an online training strategy where training data is generated randomly for each iteration. This assures that the network would not be overfitted to specific SMLM images. At the beginning of each training, an extra evaluation dataset is built to check how well the network performs on the training data. As described in the main text, the data generator could simulate spatially variant PSFs at different positions using the calibrated aberration maps. Once the aberration maps of the FOV are fixed, the network will be trained to localize emitters under these specified field-dependent aberrations. As most sCMOS cameras have a large chip size (more than 1,000 ×1,000 pixels), it is intractable to train the network directly with images of the same size as the full frame images since traditional graphics cards do not have enough memory. Therefore, the data generator only simulates smaller SMLM images which are sub-areas of the whole FOV. During training, the aberration maps are traversed with a sliding window that has the same size as the simulated images to generate the local spatially variant PSFs. It is worth noting that some overlap between sliding windows of two adjacent iterations is necessary as it ensures that the PSFs locating in the marginal region of the training images are fully learned by the network. For the coordinate channel of FD-DeepLoc, the global position is translated into coordinate channels normalized by the size of sliding window and aberration maps. Depending on the size of GPU's available memory, our training is normally performed on a window size of 128×128 pixels. We used a batch size of 10 which we found is enough to ensure the converge of optimization. AdamW[5] optimizer is used with an initial learning rate of $6\times10^{-4}$. The learning rate is then multiplied by 0.9 every 1,000 iterations. The norm of the gradient of all trainable parameters is clipped with a maximum of 0.03 for training stabilization. It is worth noting that training data with relatively high SNR is preferred to enhance the network to learn the field-dependent features, with a cost of ignoring some dim molecules in the inference process.

For the inference process, the real experimental SMLM images and its global position relative to the entire FOV are needed. If the input images are too large to be processed with the graphic card, the input images will be divided into multiple sub-areas with 20 pixels' overlap between each area. As the network could successfully link the local PSF features to their global positions in the FOV, it is very important to provide correct global positions. The network prediction consists of multiple channels: the pixel-wise probability $\hat{p}_k$of containing a molecule in the *kth* pixel, photons of the molecule $\hat{I}_k$, the lateral continuous-valued offset ($\Delta\hat{x}_k$ and $\Delta\hat{y}_k$), axial position $\hat{z}_k$, and the uncertainty of the estimated parameters ($\sigma_{xk}$, $\sigma_{yk}$, $\sigma_{zk}$, $\sigma_{Ik}$ for *x*, *y*, *z* and $I$ respectively). Local maximum detection (3×3 pixels) is applied to $\hat{p}_k$. Local maxima with $\hat{p}_k > 0.3$ and non-maximum pixels with $\hat{p}_k > 0.6$ are identified as molecule candidates. This ensures that two emitters in adjacent pixels can still be detected when the $\hat{p}_k$ is not a local maximum. The probability values of four nearest neighbor pixels around the candidate pixels are then added to the candidate pixels, forming a new probability map, $\hat{p}_k^{new}$. The new probability map was then thresholded (normally 0.7) to produce a deterministic binary map, which indicates whether a pixel contains a molecule or not. After combining the deterministic binary map with offset $\Delta\hat{x}_k$, $\Delta\hat{y}_k$, axial position $\hat{z}_k$, and uncertainties $\sigma_{xk}$, $\sigma_{yk}$, $\sigma_{zk}$, $\sigma_{Ik}$, the final molecule list with precision estimation could be obtained.

The large-FOV PSF calibration costs about 3.5 hours for Tetrapod beads (1389 beads stacks with size of 51×51×61 pixels) and 2 hours for astigmatism beads (6987 beads stacks with size of 27×27×41 pixels) on our workstation equipped with an Intel Core i9-9900 processor of 128GB RAM clocked at 3.50GHz and an NVIDIA GeForce GTX 3090 graphics card with 24.0 GB memory. The training process costs about 4 hours (30,000 iterations of network update). Once these are established, we can normally use them for weeks if the imaging path was not modified. The inference process takes about 20 hours (100,000 frames with size of 1608×1608 pixels, ~500 GBs).

**Supplementary Note 6 Data postprocessing and rendering**

For all localizations returned by FD-DeepLoc, we used SMAP[6] for the post-processing. To filter bad localizations, we normally set the *xy* localization precision threshold as 30 nm and 50 nm for astigmatic PSF and DMO PSF based 3D imaging, respectively. The threshold for the probability value $\hat{p}$ was normally set as ~0.7 in both cases and can be adjusted to reach a balance between high confidence and enough localizations. To group consecutive blinking events that are likely to originate from the same molecule, localizations close than 35nm/100nm within 3 adjacent frames were grouped as one molecule for astigmatic PSF /DMO PSF based 3D imaging, respectively. To avoid the artifacts caused by the sample drift during the acquisition, redundant cross-correlation algorithm[7] was used for drift correction in this work, with a spatial bin size of 10 nm and 10 time windows. To render the final pseudo-colored super-resolution image, we used SMAP and Vutara SRX software (Bruker) for all static super-resolution images. ViSP software[8] was used to generate the 3D super-resolution movies (**Supplementary Movie 1-6)**.
