## Supplementary Software for "Field dependent deep learning enables high-throughput whole-cell 3D super-resolution imaging": FD-DeepLoc tutorial.pdf

### Getting started

In this document, we show how to use FD-DeepLoc step by step, including large-FOV beads calibration, network training and inferring.

### System requirements

FD-DeepLoc was tested on a workstation equipped with 128 GB of memory, an Intel(R) Core(TM) i9-11900K, 3.50GHz CPU, and an NVidia GeForce RTX 3080 GPU with 10 GB of video memory. To use FD-DeepLoc yourself, a computer with CPU memory ≥ 32GB and GPU memory ≥ 8GB is recommended since FD-DeepLoc would process large-FOV SMLM images (usually > 500GB). CUDA Driver (>=11.3) is required for fast training PSF simulation, PSF fitting and PyTorch.

For field-dependent aberration map calibration, we used Matlab 2021b with CUDA 11.3 on a Windows 10 system. The deep learning part of FD-DeepLoc is based on Python and Pytorch. We recommend *conda* (https://anaconda.org) to manage the environment.

### Installation in Terminal

To manage the deep learning environment, we provide ***fd_deeploc_env.yaml*** file under the folder ***FD-DeepLoc/Field Dependent PSF Learning*** to build the conda environment. After the installation of Anaconda from ([https://anaconda.org](https://anaconda.org)), open the Anaconda Prompt on Windows. Change the current work directory to ***FD-DeepLoc/Field Dependent PSF Learning***. Enter the following commands in terminal to open the Jupyter Notebook and check the demo pipelines.

```
# CUDA capable GPU
conda env create -f fd_deeploc_env.yaml

# after previous command (all platforms)
conda activate fd_deeploc

# open the notebook
jupyter notebook
```

Several example notebooks can be found under the folder ***demo_notebooks***. The examples include training, testing and inference pipelines with commonly used functions. Users need to manually download the demo datasets using the links below and uncompress them under the folder ***demo_datasets***.

### To download demo datasets

Demo1:

The test dataset can be downloaded from: FD-DeepLoc datasets: demo1 dataset | Zenodo.

Demo2:

The test dataset can be downloaded from FD-DeepLoc datasets: demo2 dataset | Zenodo. The raw bead stacks files can be downloaded from FD-DeepLoc datasets: beads stacks for demo2 | Zenodo.

Demo3:

The test dataset can be downloaded from FD-DeepLoc datasets: demo3 dataset | Zenodo. The raw bead stacks files can be downloaded from FD-DeepLoc datasets: beads stacks for demo3 | Zenodo.

Demo4:

The test dataset can be downloaded from FD-DeepLoc datasets: demo4 dataset | Zenodo.

### Field-dependent aberration map calibration

1. Start Matlab
2. Run the file ***FD-DeepLoc\Field Dependent PSF Modeling\fitFDaberration\calibrate_abr_map_GUI.m*** to open the GUI (Figure 1).
3. Select camera files to open a dialog box to select files.
   a) Click add to add bead stacks files in the same directory, or click add dir to add multiple directories and the software will automatically find bead stacks files in these directories.
   b) Press Done to finish adding the bead stacks files to the GUI listbox.
3. The output file is set automatically, but you can change it manually with Select output file.
4. In the general parameters setting:
   a) Select the 3D modality: choose the Astigmatic or DMO Tetrapod and automatically change the corresponding fitting parameters.
   b) Enter the distance between the frames of bead stacks you used for acquiring the z-stacks. At the same time, you can check set frames to set the frame interval of z-stacks to improve the fitting speed, but may reduce the fitting accuracy due to fewer data usage.
   c) PSF Rescale: gaussian smoothing for the PSF model, also known as OTF rescale.

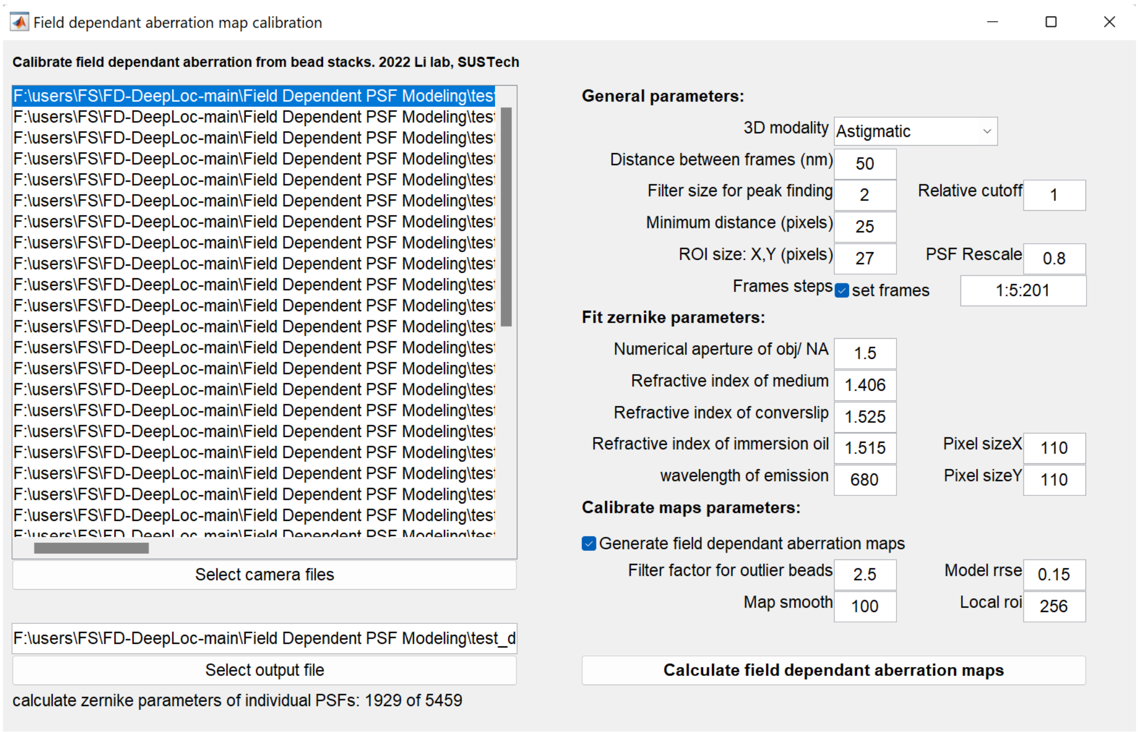

Figure 1 The GUI of field dependent aberration map calibration software

5. In the Fit zernike parameters setting:
   Set these parameters according to your microscope, and remember these parameters to set the training parameters of the network.
6. In the calibrate maps parameters setting
   a) Filter factor for outlier beads excludes some outlier beads, set Local roi to filtered outlier bead around a fixed range of pixels, usually keep the default value.

b) Model rrse discards the beads with bad fitting precision, usually keep 0.15 for astigmatism PSF and 0.2 for DMO Tetrapod PSF.

c) Map smooth sets gaussian smoothing parameter applied to the aberration maps, usually use 100 for astigmatism PSF and 200 for Tetrapod PSF.

7. Calculate field dependent aberration maps is to calculate and save aberration maps. You can monitor the progress in the status bar of the GUI. After calculation, you can check the ZernikeMap, localization precision, beads distribution and other information (Figure 2).
8. The aberration maps for network training are finally stored in a file named as ~***aber_map.mat***.

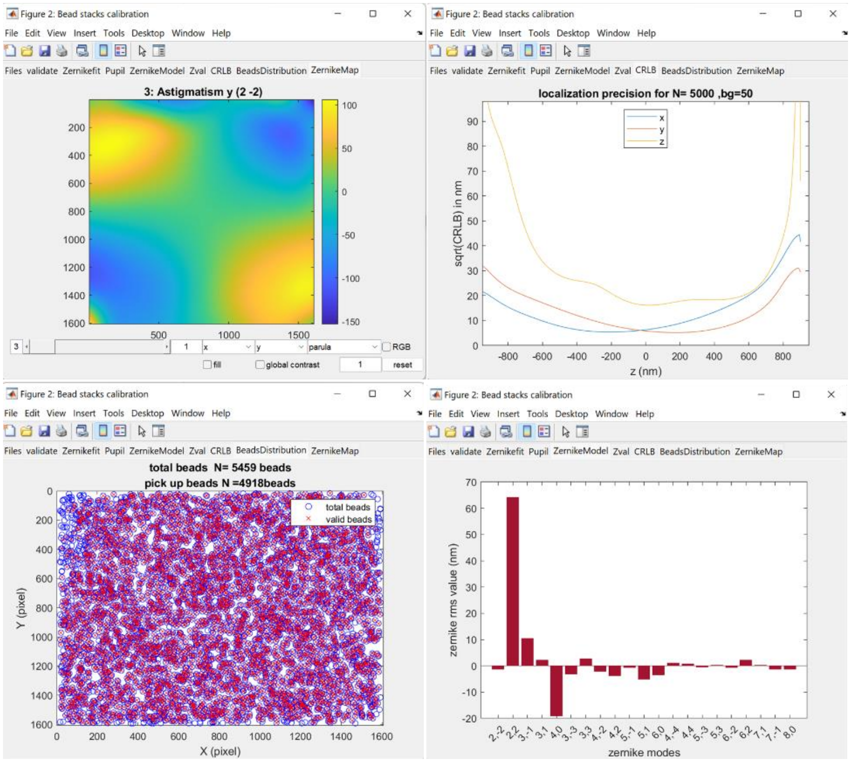

Figure 2 Example results of aberration map calibration

### Field-dependent deep-learning localization network

#### Network training

After getting the calibrated Zernike aberration map, now we can switch to the deep learning part. The following example training pipeline is based on demo2 jupyter notebook file ***Field Dependent PSF Learning\demo_notebooks\demo2_FD_astig_NPC\demo2_train.ipynb*** with detailed instruction. The training process generally includes 5 main steps as shown in Figure 3.

#### FD-DeepLoc training example

The training process of the FD-DeepLoc includes the following steps:

1. Load the experimental SMLM images and calibrated aberration map.
2. Set the parameters for the training.
3. Visually check the aberration maps, PSFs, and training frames.
4. Init an evaluation dataset for testing network performance during training.
5. Start training.

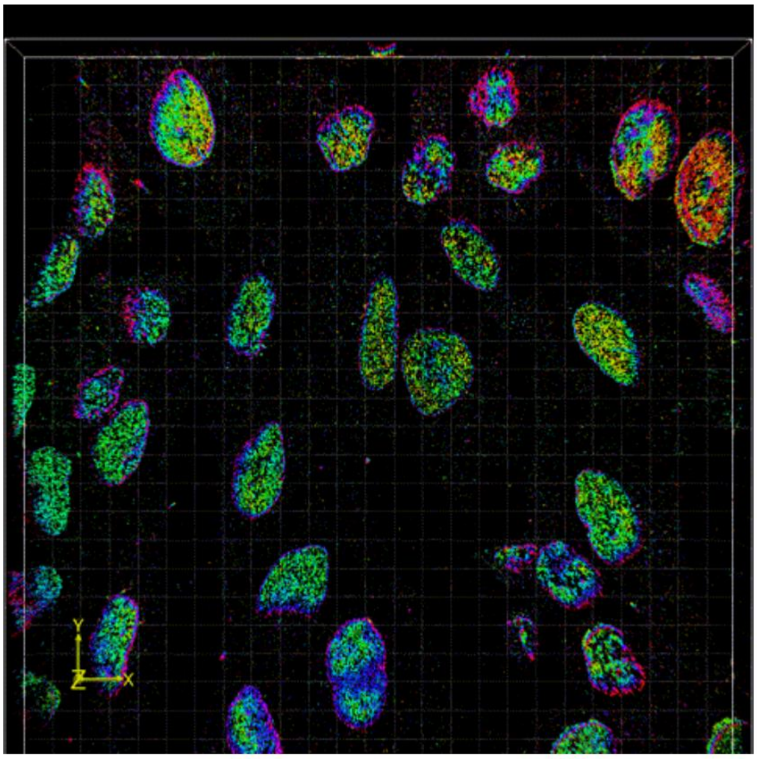

Figure 3 Main steps to train the network model.

All parameters for network building, PSF model setting, training data simulation, evaluation data simulation and model training are well explained in the ***demo2_train.ipynb***, as shown in Figure 4.

#### 2. Set the parameters for the training.

1. `net_params` :

- `local_context` means whether use three consecutive frames as input;
- `sig_pred` means whether output uncertainty about the *x,y,z,I* prediction;
- `psf_pred` means whether predict the noisefree molecule image of the input;
- `use_coordconv` means whether use the CoordConv technique to built the relationship between the PSF model and global position in the entire FOV;
- We recommend to set the remaining paprameters as default.

2. `psf_params` :

- These parameters should be set the same as when calibrating aberration maps.
- `ph_scale` is the maximum possible photon number that could be assigned to each single molecule during training;
- `initial_obj_stage` is the nominal focal plane with respect to the coverslip, it should be set carefully when there is a refractive index (RI) mismatch between `refmed` and `refimm` . If there is no RI mismatch, `initial_obj_stage` becomes meaningless;

3. `simulation_params` :

- `train_size` is the size of simulated training images, set it small when GPU memory is limited, recommend 64,128,256...;
- `surv_p` is the probability of on-state emitters appear in the next frame follows a simple binomial distribution since only three consecutive images are used in each unit;
- `min_ph` is the lower bound of the uniform distribution where photon number is sampled from, the final photon distribution is $U(minph, 1) * phscale$;
- `density` is the average number of molecules simulated on each training frame, if use `local_context` , the real average number of the middle frame will be increased by a factor of `surv_p` ;
- `z_prior` means the *z* position is sampled from $U(zprior) * zscale$;
- `margin_empty` means molecules will not be simulated at the XX% marginal area of training images, this avoids the network to learn too many incomplete PSFs;
- `camera` could be set as 'EMCCD' or 'sCMOS', if 'sCMOS', set `em_gain=1` ;
- `qe` is quantum efficiency;
- `sig_read` and `e_per_adu` are Gaussian read out noise and analog-to-digital conversion factor, respectively. Although for 'sCMOS'case they are theoretically pixel-dependent, but it does not matter a lot and here we assume them to be constant across the whole FOV;
- `baseline` is the final offset added to the image;
- `robust_training` means add small random Zernike aberration disturbance to the training PSF model at each iteration, this helps network more robust when analyzing experimental images. If in simulation where the PSF model is accurate, turn it off;
- `perlin_noise` means whether add the perlin noise to the uniform background `backg` to simulate non-uniform background; `pn_factor` should be in the range of [0,1], which implies PV degree of the added Perlin nonuniform background, set it small when experimental background looks uniform. The range of extra Perlin noise is $pnfactor * [-1, 1] * (backg - baseline) * eperadu/emgain/qe$; `pn_res` is the resolution(or frequency) of the Perlin noise, we have tested on 64/128 and found it works well;

4. `evaluation_params` :

- `eval_imgs_number` is the number of images in evaluation dataset, these images have the same size as the whole FOV (in pixels);
- `mols_per_img` is the average number of molecules on each evaluation image. If use `local_context` , the real average number of molecules will be increased by a factor of `surv_p` ;
- `batch_size` means evaluation images are processed in batches of given number. When the images are large, `batch_size` has to be lowered to save GPU memory. But if `divide_and_conquer` on, the evaluation images (as large as whole FOV) will be split into sub-areas and be processed sequentially, the `batch_size` can be larger then.

5. `train_params` :

- `lr` is the learning rate for the optimizer; `lr_decay` means learning rate will be reduced by a given factor every 1000 iterations; `clip_g_n` means gradient norm clipping; `w_decay` is the weight decay coefficient for AdamW optimizer. We don't recommend to change these training parameters as they have been tested under variant situations.
- `ph_filt` means whether ignore molecules with photon number lower than `ph_filt_thre` , thse molecules will be excluded from the ground-truth.
- `P_locs_cse` means whether include the cross entropy term in the loss function.

After the `DeepLocModel` is instantiated, it will print all training sliding windows, which indicates the order that the sub-area training images of the large FOV are simulated in cycle. The printed `field_xy` means the sub-area image's position in the whole FOV (*xy* starts from the upper left, which is opposite to [row,column]). The form is: [x_start,x_end,y_start,y_end], where (x_start,y_start) is the position of the upper left pixel of the sub-area image in the entire FOV.

Figure 4 Example snapshot of the parameters explanation in ***demo2_train.ipynb***.

After setting all parameters, one can visually check the aberration maps, PSFs, and training images by run several code blocks (Figure 5). When you make sure everything is set up properly, you can simply run the last two code blocks to create an evaluation dataset and start training. We usually trained with 30,000 iterations in 4 hours using the NVidia GeForce RTX 3080 GPU.

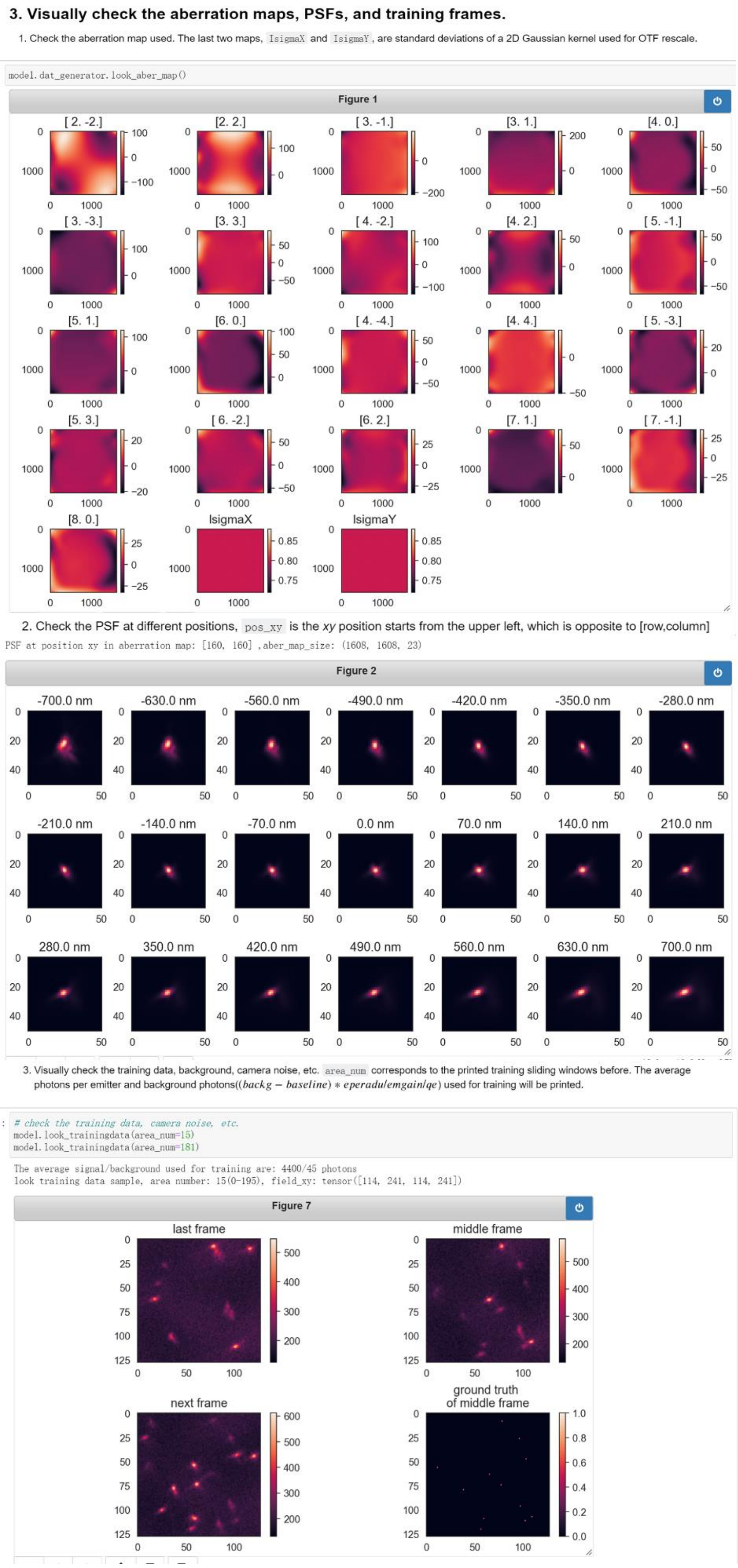

Figure 5 Example snapshots of the plotted aberration maps, PSFs and training images that need to be checked.

#### Network inference

After training, we will get a network file ***FD-DeepLoc.pkl*** to analyze the experimental data. The example inference code is provided as a jupyter notebook file ***demo2_inference.ipynb*** with detailed instruction. The inference process includes 5 main steps as shown in Figure 6.

### FD-DeepLoc inference example

The inference process includes the following steps:

1. Set the path for the trained network model and experimental images.
2. Set necessary parameters.
3. Load the network and plot the training process.
4. Check a specific experiment frame and corresponding network's multi-channel predictions.
5. Start inferring.
6. Load the ground-truth and assess(optional).

Figure 6 Main steps for using the network to analyze the experiment data.

After setting some necessary parameters, you can check the predictions of the network about a specific image frame (Figure 7). This will indicate that whether your inference parameters are set correctly or whether the network is well trained.

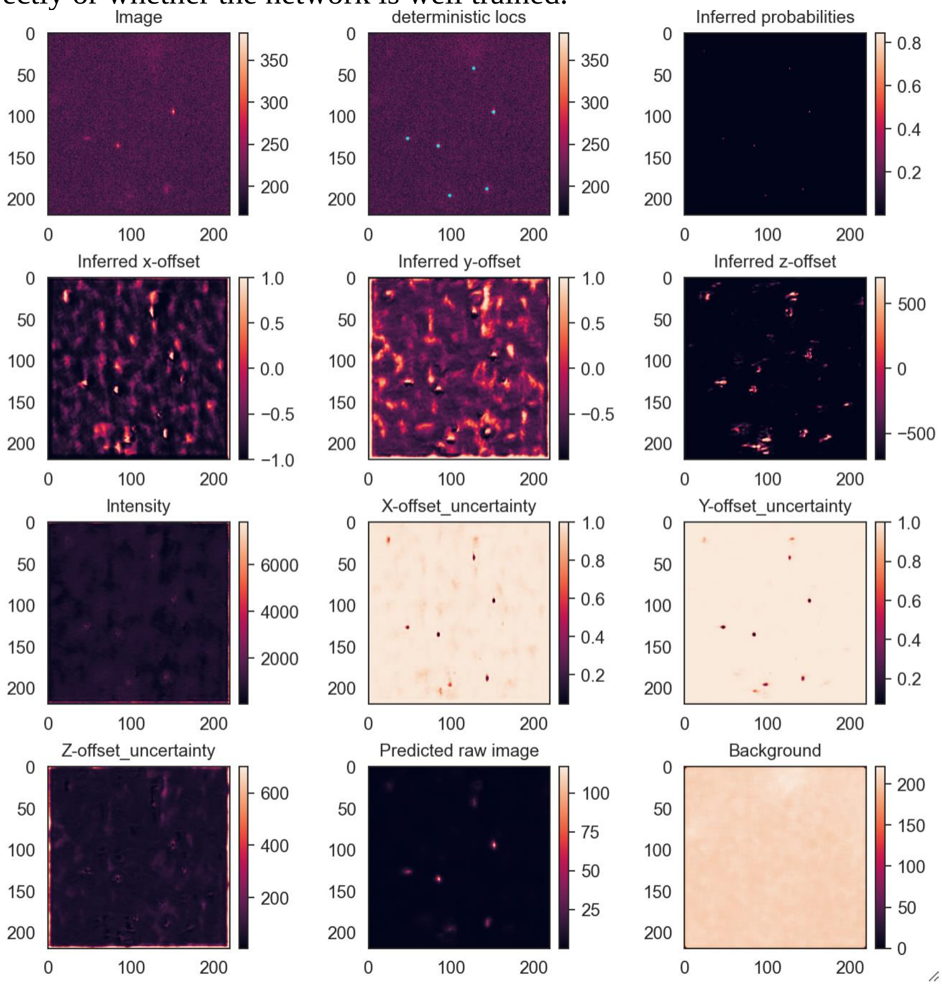

Figure 7 Example network predictions about a specific frame.

Finally, you can run the code block (Figure 8) to analyze the large-FOV SMLM data, which is usually larger than 500 GBs (100,000 frames with 1608×1608 pixels). All predictions will be saved in a ***.csv*** file in the form of molecule list.

##### 5. Start inferring.

Read big tiff and predict, save the predictions every finish processing `stack_giga` -sized SMLM images, even some accidents happen you will not lose all results. If there is already a prediction file with the same name as `save_path` , inference will start from the last saved frame number in the `save_path` file. We recommend using SMAP to postprocess the prediction list (drift correction, grouping, etc.) and render the super-resolution image (Ries, J. SMAP: a modular super-resolution microscopy analysis platform for SMLM data. Nat Methods 17, 870–872 (2020). https://doi.org/10.1038/s41592-020-0938-1).

- The parameters haven been explained before.

```
# read big tiff and predict, save the predictions every finish processing stack_giga-sized SMLM images,
# even some accidents happen you will not lose all results
total_shape, fov_size = read_bigtiff_and_predict(model, image_path_roi1, stack_giga=stack_giga, batch_size=10,
                                                 use_tqdm=True, nms=True, candi_thre=0.3, nms_thre=0.3,
                                                 rescale_xy=False, pixel_size=pixel_size, start_field_pos=start_field_pos_roi1,
                                                 divide_and_conquer=True, win_size=256, padding=True, save_path=save_path_roi1)
```

```
the file to save the predictions is:  ./demo2_FD-DeepLoc_roi_startpos_1280_1250.csv
stack: 1/22, contain imgs: 4583, already analyzed:0/100826

processing area:1/1, input field_xy:[1280 1499 1250 1505], use_coordconv:True, retain locs in area:[1280, 1499, 1250, 1505], aber_map siz
e:(1608, 1608, 23)

100%|████████████████████████████████████████████████████████████████| 459/459 [00:33<00:00, 13.63it/s]
```

Figure 8 Snapshot of the code block for data analysis in ***demo2_inference.ipynb***.
