## Supplementary Software for "Field dependent deep learning enables high-throughput whole-cell 3D super-resolution imaging": readme.txt

For each demo, we provide an individual Zenodo link to download the dataset. 
The user should download the zip file and uncompress it under this folder.


The downloading links can be found in the FD-DeepLoc tutorial.pdf or 
the 'Demo examples' section in the README.md on our GitHub "https://github.com/Li-Lab-SUSTech/FD-DeepLoc".
