## Supplementary Software for "Field dependent deep learning enables high-throughput whole-cell 3D super-resolution imaging": demo1_inference.ipynb

```
# %% [markdown]
# CRLB comparison example
The CRLB comparison process of the FD-DeepLoc includes the following steps:
1. Set the path for the trained network model
2. Load the network and plot the training process
3. Set necessary parameters
4. Check the CRLB of a local PSF at given position
5. Compare network's performance with local single-emitter CRLB

# %% [code]
import pickle
import sys
sys.path.append('../..')

import matplotlib.pyplot as plt
import numpy as np
import seaborn as sns
import torch

%matplotlib notebook
sns.set_style("white")
sys.setrecursionlimit(10000)

from fd_deeploc_core import *
from local_utils import *

# %% [markdown]
## 1. Set the path for the trained network model.

# %% [code]
fd_deeploc_path = "../../demo_datasets/demo1_FD_astig_simu_comparison_and_CRLBtest/demo1_FD-DeepLoc.pkl"
decode_path = "../../demo_datasets/demo1_FD_astig_simu_comparison_and_CRLBtest/demo1_DECODE.pkl"
image_path = "../../demo_datasets/demo1_FD_astig_simu_comparison_and_CRLBtest/sim_data_normal_aberration_medium_SNR.tif"

fd_deeploc_save_path = './'+os.path.split(fd_deeploc_path)[-1].split('.')[0]+'_'+os.path.split(image_path)[-1].split('.')[0]+'.csv'
print(fd_deeploc_save_path)

decode_save_path = './'+os.path.split(decode_path)[-1].split('.')[0]+'_'+os.path.split(image_path)[-1].split('.')[0]+'.csv'
print(decode_save_path)

# %% [markdown]
## 2. Load the network and plot the training process.


# %% [code]
# load the completely trained FD_DECODE model to test or predict
with open(fd_deeploc_path, 'rb') as f:
    fd_deeploc_model = pickle.load(f)
with open(decode_path, 'rb') as f:
    decode_model = pickle.load(f)

# plot performance during the training
print('------------------------------------------FD-DeepLoc model-------------------------------------------')
plot_train_record(fd_deeploc_model)
print('------------------------------------------DECODE model-------------------------------------------')
plot_train_record(decode_model)

# %% [markdown]
## 3. Set necessary parameters.
* `mol_photons` is the average number of signal photons used in training.
* `bg_photons` is the average number of background photons used in training.
* `test_pos` is the global position of the PSF whose CRLB will be computed.
* `Nmol` is the number of sampled z positions, which are uniformly distributed in the training z range.
* `test_num` is the number of simulated random PSFs at each sampled z position for RMSE evaluation, the total number of test set is $testnum*Nmol$.The xy positions for each single emitter are random in the center two pixel, the z position is random in a step range $(zrange/(Nmol-1))$ around each z position.

# %% [code]
# set necessary parameters
print(fd_deeploc_model.dat_generator.psf_pars['aber_map'].shape)
bg_photons = (fd_deeploc_model.dat_generator.simulation_pars['backg'] - fd_deeploc_model.dat_generator.simulation_pars['baseline']) \
                     / fd_deeploc_model.dat_generator.simulation_pars['em_gain'] * fd_deeploc_model.dat_generator.simulation_pars['e_per_adu'] \
                     / fd_deeploc_model.dat_generator.simulation_pars['qe']
mol_photons = (fd_deeploc_model.dat_generator.simulation_pars['min_ph'] + 1) / 2 * fd_deeploc_model.dat_generator.psf_pars['ph_scale']
print('{}{}{}{}{}'.format('The average signal/background used for training are: ', int(mol_photons), '/', int(bg_photons), ' photons'))
aber_w = fd_deeploc_model.dat_generator.psf_pars['aber_map'].shape[1]
aber_h = fd_deeploc_model.dat_generator.psf_pars['aber_map'].shape[0]
test_pos_list = [[int(aber_w*0.5), int(aber_h*0.5)],
                 [int(aber_w*0.95), int(aber_h*0.95)]]
print(test_pos_list)

Nmol = 25
test_num = 2000

# %% [markdown]
## 4. Check the CRLB of a local PSF at given position.
* `model` is the FD-DeepLoc model.
* `test_pos` is the global position where the PSF CRLB will be computed and checked.
* `test_photons` is the number of signal photons used to compute CRLB.
* `test_bg` is the number of background photons used to compute CRLB.
* `Nmol` is the number of sampled z positions, which are uniformly distributed in the training z range.

# %% [code]
print('------------------------------------------FD-DeepLoc model-------------------------------------------')
check_local_CRLB(model=fd_deeploc_model, test_pos=test_pos_list[0], test_photons=mol_photons, test_bg=bg_photons, Nmol=Nmol)
# print('------------------------------------------DECODE model-------------------------------------------')
# check_local_CRLB(model=decode_model, test_pos=test_pos_list[0], test_photons=mol_photons, test_bg=bg_photons, Nmol=Nmol)

# %% [markdown]
## 5. Compare network's performance with local single-emitter CRLB.
* `use_train_cam`: If true, the single-emitter test set will be simulated using the camera noise of training (e.g., qe, readout noise, analog-to-digital conversion factor). Otherwise only poisson noise will be added.
* `test_num` is the number of simulated random PSFs at each sampled z position for RMSE evaluation, the total number of test set is $testnum*Nmol$.The xy positions for each single emitter are random in the center two pixel, the z position is random in a step range $(zrange/(Nmol-1))$ around each z position.
* The remaining is the same as above.

# %% [code]
print('------------------------------------------FD-DeepLoc model-------------------------------------------')
zpos, x_crlb, y_crlb, z_crlb, rmse_xyz_fd_deeploc = test_local_CRLB(model=fd_deeploc_model, test_pos=test_pos_list[0], test_photons=mol_photons, test_bg=bg_photons, Nmol=Nmol,
                use_train_cam=True, test_num=test_num, show_res=True)
print('------------------------------------------DECODE model-------------------------------------------')
zpos, x_crlb, y_crlb, z_crlb, rmse_xyz_decode = test_local_CRLB(model=decode_model, test_pos=test_pos_list[0], test_photons=mol_photons, test_bg=bg_photons, Nmol=Nmol,
                use_train_cam=True, test_num=test_num, show_res=True)

# compare directly
plt.figure(constrained_layout=True)
plt.plot(zpos, x_crlb,'b', zpos, y_crlb,'g', zpos, z_crlb,'r')
plt.scatter(zpos, rmse_xyz_fd_deeploc[0, :],c="b",marker="*")
plt.scatter(zpos, rmse_xyz_fd_deeploc[1, :],c="g",marker="*")
plt.scatter(zpos, rmse_xyz_fd_deeploc[2, :],c="r",marker="*")
plt.scatter(zpos, rmse_xyz_decode[0, :],c="b",marker="o")
plt.scatter(zpos, rmse_xyz_decode[1, :],c="g",marker="o")
plt.scatter(zpos, rmse_xyz_decode[2, :],c="r",marker="o")
plt.legend(('$CRLB_x^{1/2}$', '$CRLB_y^{1/2}$', '$CRLB_z^{1/2}$', 
            '$FD-DeepLoc RMSE_x$', '$FD-DeepLoc RMSE_y$', '$FD-DeepLoc RMSE_z$', 
            '$DECODE RMSE_x$', '$DECODE RMSE_y$', '$DECODE RMSE_z$'), ncol=3,
           loc='upper center')
plt.xlim([np.min(zpos), np.max(zpos)])
plt.show()

# %% [code]
print('------------------------------------------FD-DeepLoc model-------------------------------------------')
zpos, x_crlb, y_crlb, z_crlb, rmse_xyz_fd_deeploc = test_local_CRLB(model=fd_deeploc_model, test_pos=test_pos_list[1], test_photons=mol_photons, test_bg=bg_photons, Nmol=Nmol,
                use_train_cam=True, test_num=test_num, show_res=True)
print('------------------------------------------DECODE model-------------------------------------------')
zpos, x_crlb, y_crlb, z_crlb, rmse_xyz_decode = test_local_CRLB(model=decode_model, test_pos=test_pos_list[1], test_photons=mol_photons, test_bg=bg_photons, Nmol=Nmol,
                use_train_cam=True, test_num=test_num, show_res=True)

# compare directly
plt.figure(constrained_layout=True)
plt.plot(zpos, x_crlb,'b', zpos, y_crlb,'g', zpos, z_crlb,'r')
plt.scatter(zpos, rmse_xyz_fd_deeploc[0, :],c="b",marker="*")
plt.scatter(zpos, rmse_xyz_fd_deeploc[1, :],c="g",marker="*")
plt.scatter(zpos, rmse_xyz_fd_deeploc[2, :],c="r",marker="*")
plt.scatter(zpos, rmse_xyz_decode[0, :],c="b",marker="o")
plt.scatter(zpos, rmse_xyz_decode[1, :],c="g",marker="o")
plt.scatter(zpos, rmse_xyz_decode[2, :],c="r",marker="o")
plt.legend(('$CRLB_x^{1/2}$', '$CRLB_y^{1/2}$', '$CRLB_z^{1/2}$', 
            '$FD-DeepLoc RMSE_x$', '$FD-DeepLoc RMSE_y$', '$FD-DeepLoc RMSE_z$', 
            '$DECODE RMSE_x$', '$DECODE RMSE_y$', '$DECODE RMSE_z$'), ncol=3,
           loc='upper center')
plt.xlim([np.min(zpos), np.max(zpos)])
plt.show()

# %% [markdown]
## 6. Start inferring on dataset
* `stack_giga` is the size of sequentially processed images (in gigabyte), it is only an approximate value, set it small when you have limited RAM.
* `pixel_size` is the physical size of each camera pixel (_xy_ in nm).
* `start_field_pos`: **Important**, it is the **_xy_** position of the upper left pixel of the input images in the entire FOV. For example, `start_field_pos` [102,41] means the upper left pixel (namely local position [0,0]) of the input images is located at [102,41] of the whole FOV. Thus CoordConv can get the global position of the input images.

Read big tiff and predict, save the predictions every finish processing `stack_giga`-sized SMLM images, even some accidents happen you will not lose all results. If there is already a prediction file with the same name as `save_path`, inference will start from the last saved frame number in the `save_path` file. We recommend using SMAP to postprocess the prediction list (drift correction, grouping, etc.) and render the super-resolution image (Ries, J. SMAP: a modular super-resolution microscopy analysis platform for SMLM data. Nat Methods 17, 870–872 (2020). https://doi.org/10.1038/s41592-020-0938-1). 
* The parameters haven been explained before.

# %% [code]
# set the size of file to be processed sequentially, unit: gigabyte
stack_giga = 0.5
pixel_size = [100, 100]
start_field_pos = [0, 0]

print('------------------------------------------FD-DeepLoc model-------------------------------------------')
# read big tiff and predict, save the predictions every finish processing stack_giga-sized SMLM images,
# even some accidents happen you will not lose all results
total_shape,fov_size = read_bigtiff_and_predict(fd_deeploc_model, image_path, stack_giga=stack_giga, batch_size=10,
                                                use_tqdm=True, nms=True, candi_thre=0.3, nms_thre=0.3,
                                                rescale_xy=False, pixel_size=pixel_size, start_field_pos=start_field_pos, 
                                                divide_and_conquer=True, win_size=256, padding=True, save_path=fd_deeploc_save_path)

print('------------------------------------------DECODE model-------------------------------------------')
total_shape,fov_size = read_bigtiff_and_predict(decode_model, image_path, stack_giga=stack_giga, batch_size=10,
                                                use_tqdm=True, nms=True, candi_thre=0.3, nms_thre=0.3,
                                                rescale_xy=False, pixel_size=pixel_size, start_field_pos=start_field_pos, 
                                                divide_and_conquer=True, win_size=256, padding=True, save_path=decode_save_path)

# %% [markdown]
## 7. Load the ground-truth and assess.
If in simulation case, we can compare the network's predictions with the ground truth and calculate performance metrics.

# %% [code]
eval_csv = read_csv("../../demo_datasets/demo1_FD_astig_simu_comparison_and_CRLBtest/ground_truth.csv")

print('------------------------------------------FD-DeepLoc model-------------------------------------------')
# if in simulation, read the predictions and ground truth, then evaluate
fd_deeploc_preds = read_csv(fd_deeploc_save_path).tolist()
# compare the raw predictions and the ground truth
match_dict_raw, matches_raw = assess(test_frame_nbr=total_shape[0], test_csv=eval_csv, pred_inp=fd_deeploc_preds,
                                     size_xy=fov_size, tolerance=250, border=450,
                                     print_res=True, min_int=False, tolerance_ax=500, segmented=False)

print('------------------------------------------DECODE model-------------------------------------------')
decode_preds = read_csv(decode_save_path).tolist()
# compare the raw predictions and the ground truth
match_dict_raw, matches_raw = assess(test_frame_nbr=total_shape[0], test_csv=eval_csv, pred_inp=decode_preds,
                                     size_xy=fov_size, tolerance=250, border=450,
                                     print_res=True, min_int=False, tolerance_ax=500, segmented=False)

# %% [code]
# We recommend using SMAP to postprocess the prediction list (drift correction, grouping, etc.) 
# and render the super-resolution image 
fd_deeploc_preds = read_csv(fd_deeploc_save_path).tolist()

decode_preds = read_csv(decode_save_path).tolist()

hist_img_fd_deeploc = create_3d_hist(fd_deeploc_preds, z_clip=[-700, 700], pix_size=50, sigma=5, contrast_fac=20)

hist_img_decode = create_3d_hist(decode_preds, z_clip=[-700, 700], pix_size=50, sigma=5, contrast_fac=20)

print('------------------------------------------compare the reconstructed image-------------------------------------------')
plt.figure()
plt.subplot(1,2,1)
plt.imshow(hist_img_fd_deeploc)
plt.subplot(1,2,2)
plt.imshow(hist_img_decode)

# %% [code]
```
