## Supplementary Software for "Field dependent deep learning enables high-throughput whole-cell 3D super-resolution imaging": demo1_train.ipynb

```
# %% [markdown]
# FD-DeepLoc training example
The training process of the FD-DeepLoc includes the following steps:
1. Load the experimental SMLM images and calibrated aberration map.
2. Set the parameters for the training.
3. Visually check the aberration maps, PSFs, and training frames.
4. Init an evaluation dataset for testing network performance during training.
5. Start training.
![traverse_large_FOV_NPC_ast2%2000_00_00-00_00_30.gif](attachment:traverse_large_FOV_NPC_ast2%2000_00_00-00_00_30.gif)

# %% [code]
import datetime
import sys
sys.path.append('../..')

import scipy.io
import seaborn as sns

sns.set_style("white")
%matplotlib notebook

from fd_deeploc_core import *
from local_utils import *

sys.setrecursionlimit(10000)
setup_seed(25)

# %% [markdown]
## 1. Load the experimental SMLM images and calibrated aberration map. 
Estimate the background of experimental images from the pixel values. One can also manually set a background value by directly assigning the variable `bg_exp`.

# %% [code]
# loading experimental images and aberration map
exp_dat = "../../demo_datasets/demo1_FD_astig_simu_comparison_and_CRLBtest/sim_data_normal_aberration_medium_SNR.tif"
aber_map = scipy.io.loadmat("../../demo_datasets/demo1_FD_astig_simu_comparison_and_CRLBtest/simulated_2048ast_normal_aberration.mat")
aber_map = aber_map['aber_map']

# estimate the experimental background (since the bg of marginal area is much lower, we choose the center area)
# exp_imgs = imread(exp_dat)
exp_imgs = read_first_size_gb_tiff(exp_dat, size_gb=8)
exp_imgs=exp_imgs[:,int(0.1*exp_imgs.shape[1]):-int(0.1*exp_imgs.shape[1]),int(0.1*exp_imgs.shape[2]):-int(0.1*exp_imgs.shape[2])]
print('size of images used to estiamte bg(adu):',exp_imgs.shape)
bg_exp, _ = get_bg_stats(exp_imgs, percentile=50, plot=False)
factor = exp_imgs.mean(0).max().astype('float32')
offset = exp_imgs.mean().astype('float32')
del exp_imgs
print('background adu est(normally contain offset):%.2f, factor(adu):%.2f, offset(adu):%.2f'%(bg_exp,factor,offset))

# %% [markdown]
## 2. Set the parameters for the training.
1. `net_params`: 
* `local_context` means whether use three consecutive frames as input; 
* `sig_pred` means whether output uncertainty about the _x,y,z,I_ prediction; 
* `psf_pred` means whether predict the noisefree molecule image of the input; 
* `use_coordconv` means whether use the CoordConv technique to built the relationship between the PSF model and global position in the entire FOV;
* We recommend to set the remaining paprameters as default.

2. `psf_params`: 
* These parameters should be set the same as when calibrating aberration maps. 
* `ph_scale` is the maximum possible photon number that could be assigned to each single molecule during training; 
* `initial_obj_stage` is the nominal focal plane with respect to the coverslip, it should be set carefully when there is a refractive index (RI) mismatch between `refmed` and `refimm`. If there is no RI mismatch, `initial_obj_stage` becomes meaningless;

3. `simulation_params`: 
* `train_size` is the size of simulated training images, set it small when GPU memory is limited, recommend 64,128,256...;
* `surv_p` is the probability of on-state emitters appear in the next frame follows a simple binomial distribution since only three consecutive images are used in each unit; 
* `min_ph` is the lower bound of the uniform distribution where photon number is sampled from, the final photon distribution is $ U(minph,1)*phscale$; 
* `density` is the average number of molecules simulated on each training frame, if use `local_context`, the real average number of the middle frame will be increased by a factor of `surv_p`; 
* `z_prior` means the _z_ position is sampled from $U(zprior)*zscale$; 
* `margin_empty` means molecules will not be simulated at the XX% marginal area of training images, this avoids the network to learn too many incomplete PSFs; 
* `camera` could be set as 'EMCCD' or 'sCMOS', if 'sCMOS', set `em_gain=1`; 
* `qe` is quantum efficiency; 
* `sig_read` and `e_per_adu` are Gaussian read out noise and analog-to-digital conversion factor, respectively. Although for 'sCMOS'case they are theoretically pixel-dependent, but it does not matter a lot and here we assume them to be constant across the whole FOV; 
* `baseline` is the final offset added to the image;
* `robust_training` means add small random Zernike aberration disturbance to the training PSF model at each iteration, this helps network more robust when analyzing experimental images. If in simulation where the PSF model is accurate, turn it off;
* `perlin_noise` means whether add the perlin noise to the uniform background `backg` to simulate non-uniform background; `pn_factor` should be in the range of [0,1], which implies PV degree of the added Perlin nonuniform background, set it small when experimental background looks uniform. The range of extra Perlin noise is $pnfactor*[-1,1]*(backg-baseline)*eperadu/emgain/qe$; `pn_res` is the resolution(or frequency) of the Perlin noise, we have tested on 64/128 and found it works well;

4. `evaluation_params`: 
* `eval_imgs_number` is the number of images in evaluation dataset, these images have the same size as the whole FOV (in pixels); 
* `mols_per_img` is the average number of molecules on each evaluation image. If use `local_context`, the real average number of molecules will be increased by a factor of `surv_p`; 
* `batch_size` means evaluation images are processed in batches of given number. When the images are large, `batch_size` has to be lowered to save GPU memory. But if `divide_and_conquer` on, the evaluation images (as large as whole FOV) will be split into sub-areas and be processed sequentially, the `batch_size` can be larger then.

5. `train_params`: 
* `lr` is the learning rate for the optimizer; `lr_decay` means learning rate will be reduced by a given factor every 1000 iterations; `clip_g_n` means gradient norm clipping; `w_decay` is the weight decay coefficient for AdamW optimizer. We don't recommend to change these training parameters as they have been tested under variant situations.
* `ph_filt` means whether ignore molecules with photon number lower than `ph_filt_thre`, thse molecules will be excluded from the ground-truth. 
* `P_locs_cse` means whether include the cross entropy term in the loss function.

After the `DeepLocModel` is instantiated, it will print all training sliding windows, which indicates the order that the sub-area training images of the large FOV are simulated in cycle. The printed `field_xy` means the sub-area image's position in the whole FOV (_xy_ starts from the upper left, which is opposite to [row,column]). The form is: [x_start,x_end,y_start,y_end], where (x_start,y_start) is the position of the upper left pixel of the sub-area image in the entire FOV.

# %% [code]
# set parameters
# network architecture parameters
net_params = {'n_filters': 48, 'padding': 1, 'kernel_size': 3, 'factor': factor, 'offset': offset,
              'local_context': False, 'sig_pred': True, 'psf_pred': True, 'use_coordconv': True}
# psf simulation parameters
psf_params = {'aber_map': aber_map, 'ph_scale': 7000, 'psf_size': 51, 'z_scale': 700,
              'pixel_size_xy': [100, 100], 'NA': 1.49, 'refmed': 1.518, 'refcov': 1.518, 'refimm': 1.518,
              'lambda': 660, 'initial_obj_stage': -800}
# training data simulation parameters
simulation_params = {'train_size': 128, 'backg': bg_exp, 'surv_p': 0.5, 'min_ph': 0.428,
                     'density': 10, 'z_prior': [-1, 1], 'margin_empty': 0.05, 'camera': 'sCMOS', 'em_gain': 1.0,
                     'qe': 1, 'spurious_c': 0.000, 'sig_read': 0.0001, 'e_per_adu': 1, 'baseline': 0,
                     'robust_training': False, 'perlin_noise': False, 'pn_factor': 0.2, 'pn_res':64}
# evaluation data simulation parameters, and recognition parameters when test on evaluation set
evaluation_params = {'eval_imgs_number': 30, 'mols_per_img': 500, 'batch_size': 4, 'divide_and_conquer': True}
# training parameters
train_params = {'lr': 6e-4, 'lr_decay': 0.9, 'clip_g_n': 0.03, 'w_decay': 0.1, 'ph_filt': False,
                'ph_filt_thre': 500, 'P_locs_cse': True}

# initialize model
fd_deeploc_model = DeepLocModel(net_pars=net_params, psf_pars=psf_params, simulation_pars=simulation_params,
                     evaluation_pars=evaluation_params, train_pars=train_params)


# %% [markdown]
## 3. Visually check the aberration maps, PSFs, and training frames.
1. Check the aberration map used. The last two maps, `IsigmaX` and `IsigmaY`, are standard deviations of a 2D Gaussian kernel used for OTF rescale.

# %% [code]
fd_deeploc_model.dat_generator.look_aber_map()

# %% [markdown]
2. Check the PSF at different positions, `pos_xy` is the _xy_ position starts from the upper left, which is opposite to [row,column]

# %% [code]
aber_w = fd_deeploc_model.dat_generator.psf_pars['aber_map'].shape[1]
aber_h = fd_deeploc_model.dat_generator.psf_pars['aber_map'].shape[0]
test_pos_list = [[int(aber_w*0.1), int(aber_h*0.1)],
                [int(aber_w*0.9), int(aber_h*0.1)],
                [int(aber_w*0.1), int(aber_h*0.9)],
                [int(aber_w*0.9), int(aber_h*0.9)],
                [int(aber_w*0.5), int(aber_h*0.5)]]
fd_deeploc_model.dat_generator.look_psf(pos_xy=test_pos_list[0], z_scale=fd_deeploc_model.dat_generator.psf_pars['z_scale'])
fd_deeploc_model.dat_generator.look_psf(pos_xy=test_pos_list[1], z_scale=fd_deeploc_model.dat_generator.psf_pars['z_scale'])
fd_deeploc_model.dat_generator.look_psf(pos_xy=test_pos_list[2], z_scale=fd_deeploc_model.dat_generator.psf_pars['z_scale'])
fd_deeploc_model.dat_generator.look_psf(pos_xy=test_pos_list[3], z_scale=fd_deeploc_model.dat_generator.psf_pars['z_scale'])
fd_deeploc_model.dat_generator.look_psf(pos_xy=test_pos_list[4], z_scale=fd_deeploc_model.dat_generator.psf_pars['z_scale'])

# %% [markdown]
3. Visually check the training data, background, camera noise, etc. `area_num` corresponds to the printed training sliding windows before. The average photons per emitter and background photons($(backg-baseline)*eperadu/emgain/qe$) used for training will be printed.

# %% [code]
# check the training data, camera noise, etc.
fd_deeploc_model.look_trainingdata(area_num=183)
fd_deeploc_model.look_trainingdata(area_num=112)

# %% [markdown]
## 4. Init an evaluation dataset for testing network performance during training.

# %% [code]
# init the eval_dataset for track performance during training (it may take tens of minutes),
# performance indicates how well the network analyzes the evaluation data 
fd_deeploc_model.init_eval_data()
# init the record dict, including loss, efficiency, etc.
fd_deeploc_model.init_recorder()

# %% [markdown]
## 5. Start training.
* `batch_size` is the number of training images for each iteration, normally the bigger the better, it depends on your available GPU memory. We found 10 is enough to ensure the convergence.
* `max_iters` is the number of training iterations, we usually set it as 3,0000. The network's performance (_Efficiency3D, Jaccard, RMSE, etc._) on evlauation dataset will be printed every `print_freq` iterations (except first 1000 iterations) while `print_output` is on.

# %% [code]
# train the FD_DeepLoc model
# del aber_map; 
# torch.autograd.set_detect_anomaly(True)
fd_deeploc_model.filename = datetime.datetime.now().strftime('%Y-%m-%d-%H') + 'FD-DeepLoc'
fd_deeploc_model.fit(batch_size=10, max_iters=60000, print_freq=500, print_output=True)

# %% [code]
del fd_deeploc_model

# %% [markdown]
## Train another network without using CoordConv and Cross Entropy

# %% [code]
# set parameters
# network architecture parameters
net_params = {'n_filters': 48, 'padding': 1, 'kernel_size': 3, 'factor': factor, 'offset': offset,
              'local_context': False, 'sig_pred': True, 'psf_pred': False, 'use_coordconv': False}
# psf simulation parameters
psf_params = {'aber_map': aber_map, 'ph_scale': 7000, 'psf_size': 51, 'z_scale': 700,
              'pixel_size_xy': [100, 100], 'NA': 1.49, 'refmed': 1.518, 'refcov': 1.518, 'refimm': 1.518,
              'lambda': 660, 'initial_obj_stage': -800}
# training data simulation parameters
simulation_params = {'train_size': 128, 'backg': bg_exp, 'surv_p': 0.5, 'min_ph': 0.428,
                     'density': 10, 'z_prior': [-1, 1], 'margin_empty': 0.05, 'camera': 'sCMOS', 'em_gain': 1.0,
                     'qe': 1, 'spurious_c': 0.000, 'sig_read': 0.0001, 'e_per_adu': 1, 'baseline': 0,
                     'robust_training': False, 'perlin_noise': False, 'pn_factor': 0.2, 'pn_res':64}
# evaluation data simulation parameters, and recognition parameters when test on evaluation set
evaluation_params = {'eval_imgs_number': 30, 'mols_per_img': 500, 'batch_size': 4, 'divide_and_conquer': True}
# training parameters
train_params = {'lr': 6e-4, 'lr_decay': 0.9, 'clip_g_n': 0.03, 'w_decay': 0.1, 'ph_filt': False,
                'ph_filt_thre': 500, 'P_locs_cse': False}

# initialize model
decode_model = DeepLocModel(net_pars=net_params, psf_pars=psf_params, simulation_pars=simulation_params,
                     evaluation_pars=evaluation_params, train_pars=train_params)

# %% [code]
decode_model.dat_generator.look_aber_map()

aber_w = decode_model.dat_generator.psf_pars['aber_map'].shape[1]
aber_h = decode_model.dat_generator.psf_pars['aber_map'].shape[0]
test_pos_list = [[int(aber_w*0.1), int(aber_h*0.1)],
                [int(aber_w*0.9), int(aber_h*0.1)],
                [int(aber_w*0.1), int(aber_h*0.9)],
                [int(aber_w*0.9), int(aber_h*0.9)],
                [int(aber_w*0.5), int(aber_h*0.5)]]
decode_model.dat_generator.look_psf(pos_xy=test_pos_list[0], z_scale=decode_model.dat_generator.psf_pars['z_scale'])
decode_model.dat_generator.look_psf(pos_xy=test_pos_list[1], z_scale=decode_model.dat_generator.psf_pars['z_scale'])
decode_model.dat_generator.look_psf(pos_xy=test_pos_list[2], z_scale=decode_model.dat_generator.psf_pars['z_scale'])
decode_model.dat_generator.look_psf(pos_xy=test_pos_list[3], z_scale=decode_model.dat_generator.psf_pars['z_scale'])
decode_model.dat_generator.look_psf(pos_xy=test_pos_list[4], z_scale=decode_model.dat_generator.psf_pars['z_scale'])

# check the training data, camera noise, etc.
decode_model.look_trainingdata(area_num=183)
decode_model.look_trainingdata(area_num=112)

# init the eval_dataset for track performance during training (it may take tens of minutes),
# performance indicates how well the network analyzes the evaluation data 
decode_model.init_eval_data()
# init the record dict, including loss, efficiency, etc.
decode_model.init_recorder()

# %% [code]
decode_model.filename = datetime.datetime.now().strftime('%Y-%m-%d-%H') + 'DECODE'
decode_model.fit(batch_size=10, max_iters=60000, print_freq=500, print_output=True)

# %% [code]
```
