## Supplementary Software for "Field dependent deep learning enables high-throughput whole-cell 3D super-resolution imaging": demo2_inference.ipynb

```
# %% [markdown]
# FD-DeepLoc inference example
The inference process includes the following steps:
1. Set the path for the trained network model and experimental images.
2. Set necessary parameters.
3. Load the network and plot the training process.
4. Check a specific experiment frame and corresponding network's multi-channel predictions.
5. Start inferring.
6. Load the ground-truth and assess(optional). 

# %% [code]
import pickle
import sys
sys.path.append('../..')

import seaborn as sns

sns.set_style("white")
sys.setrecursionlimit(10000)
%matplotlib notebook

from fd_deeploc_core import *
from local_utils import *

# %% [markdown]
## 1. Set the path for the trained network model and experimental images.
The printed `save_path` is where to save the network's predicted molecule list.

# %% [code]
# set the trained model path and the image path that need to be analysed
network_path = "../../demo_datasets/demo2_FD_astig_NPC/demo2_FD-DeepLoc.pkl"

image_path_roi1 = "../../demo_datasets/demo2_FD_astig_NPC/roi_startpos_1280_1250.tif"
save_path_roi1 = './'+os.path.split(network_path)[-1].split('.')[0]+'_'+os.path.split(image_path_roi1)[-1].split('.')[0]+'.csv'
print(save_path_roi1)

image_path_roi2 = "../../demo_datasets/demo2_FD_astig_NPC/roi_startpos_810_790.tif"
save_path_roi2 = './'+os.path.split(network_path)[-1].split('.')[0]+'_'+os.path.split(image_path_roi2)[-1].split('.')[0]+'.csv'
print(save_path_roi2)

# %% [markdown]
## 2. Set necessary parameters.
* `stack_giga` is the size of sequentially processed images (in gigabyte), it is only an approximate value, set it small when you have limited RAM.
* `pixel_size` is the physical size of each camera pixel (_xy_ in nm).
* `start_field_pos`: **Important**, it is the **_xy_** position of the upper left pixel of the input images in the entire FOV. For example, `start_field_pos` [102,41] means the upper left pixel (namely local position [0,0]) of the input images is located at [102,41] of the whole FOV. Thus CoordConv can get the global position of the input images.

# %% [code]
# set the size of file to be processed sequentially, unit: gigabyte
stack_giga = 0.5
pixel_size = [110, 110]

# make sure the field position of sub-region is correct!
start_field_pos_roi1 = [1280, 1250]

start_field_pos_roi2 = [810, 790]

# %% [markdown]
## 3. Load the network and plot the training process.

# %% [code]
# load the completely trained FD_DECODE model
with open(network_path, 'rb') as f:
    model = pickle.load(f)

# plot performance during the training
plot_train_record(model)

# %% [markdown]
## 4. Check a specific experiment frame and corresponding network's multi-channel predictions.
* `plot_num`: The number of specific image that will be checked, better $\geq$ 5 (considering temporal context and background estimation)
* `nms`: If False, only use a simple threshold `nms_thre` to filter probability channel to get the deterministic pixels. If True, probability channel pixels with value>`candi_thre` will be treated as candidates for local maximum searching. Then add the values from the 4 adjacent pixels to local maximums and filter the modified probability channel with the threshold `nms_thre` to get deterministic pixel-level localizations.
* `rescale_xy`: Rescales _x_ and _y_ offsets (inplace) so that they are distributed uniformly within [-0.5, 0.5] to correct for biased outputs (usually False).
* `pixel_size`,`start_field_pos` are explained before.
* `divide_and_conquer`: Divide the large images into small sub-area images. This is necessary as large input images may lead to GPU memory problem, this enables the network process the large images by sub-areas.
* `win_size`: If `divide_and_conquer`=True, set the size of sub-areas to be analyzed, must be a multiple of 4 (avoid error when down-sampling or up-sampling).
* `padding`: If `divide_and_conquer`=True and `padding`=True, this will segment a larger area (20 pixels) than `win_size` and traverse with overlap to avoid error from incomplete PSFs at margin.

# %% [code]
# check the specified frame and corresponding multi-channel output
check_specific_frame_output(plot_num=18, model=model, image_path=image_path_roi1, eval_csv=None,
                            nms=True, candi_thre=0.3, nms_thre=0.3, rescale_xy=False, pixel_size=pixel_size,
                            start_field_pos=start_field_pos_roi1, divide_and_conquer=True, win_size=256, padding=True)


check_specific_frame_output(plot_num=18, model=model, image_path=image_path_roi2, eval_csv=None,
                            nms=True, candi_thre=0.3, nms_thre=0.3, rescale_xy=False, pixel_size=pixel_size,
                            start_field_pos=start_field_pos_roi2, divide_and_conquer=True, win_size=256, padding=True)

# %% [markdown]
## 5. Start inferring.
Read big tiff and predict, save the predictions every finish processing `stack_giga`-sized SMLM images, even some accidents happen you will not lose all results. If there is already a prediction file with the same name as `save_path`, inference will start from the last saved frame number in the `save_path` file. We recommend using SMAP to postprocess the prediction list (drift correction, grouping, etc.) and render the super-resolution image (Ries, J. SMAP: a modular super-resolution microscopy analysis platform for SMLM data. Nat Methods 17, 870–872 (2020). https://doi.org/10.1038/s41592-020-0938-1). 
* The parameters haven been explained before.

# %% [code]
# read big tiff and predict, save the predictions every finish processing stack_giga-sized SMLM images,
# even some accidents happen you will not lose all results
total_shape,fov_size = read_bigtiff_and_predict(model, image_path_roi1, stack_giga=stack_giga, batch_size=10,
                                                use_tqdm=True, nms=True, candi_thre=0.3, nms_thre=0.3,
                                                rescale_xy=False, pixel_size=pixel_size, start_field_pos=start_field_pos_roi1, 
                                                divide_and_conquer=True, win_size=256, padding=True, save_path=save_path_roi1)

# %% [code]
total_shape,fov_size = read_bigtiff_and_predict(model, image_path_roi2, stack_giga=stack_giga, batch_size=10,
                                                use_tqdm=True, nms=True, candi_thre=0.3, nms_thre=0.3,
                                                rescale_xy=False, pixel_size=pixel_size, start_field_pos=start_field_pos_roi2, 
                                                divide_and_conquer=True, win_size=256, padding=True, save_path=save_path_roi2)

# %% [markdown]
### filter and render (recommend to be done in SMAP)

# %% [code]
# read the prediction list
# render, if the FOV is very big, this might be slow and cost large RAM, 
# you may need a postprocessing software to do drift correction and grouping for better image rendering.    
preds_roi1 = read_csv(save_path_roi1).tolist()
hist_img_roi1 = create_3d_hist(preds_roi1, z_clip=[-300, 600], pix_size=50, sigma=10, contrast_fac=100)

preds_roi2 = read_csv(save_path_roi2).tolist()
hist_img_roi2 = create_3d_hist(preds_roi2, z_clip=[-300, 600], pix_size=50, sigma=10, contrast_fac=100)

plt.figure()
plt.subplot(1,2,1)
plt.imshow(hist_img_roi1)
plt.subplot(1,2,2)
plt.imshow(hist_img_roi2)

# %% [code]
# filter the raw predictions by probability channel and uncertainty channel (can also be done in SMAP)
preds_roi1_filtered = filt_preds(preds_roi1, nms_p_thre=0.9, sig_perc=95, is_3d=True)
fig, axes = plot_preds_distribution(preds_roi1,preds_roi1_filtered)
hist_img_roi1_filtered = create_3d_hist(preds_roi1_filtered, z_clip=[-300, 600], pix_size=50, sigma=10, contrast_fac=100)

preds_roi2_filtered = filt_preds(preds_roi2, nms_p_thre=0.9, sig_perc=95, is_3d=True)
fig, axes = plot_preds_distribution(preds_roi2,preds_roi2_filtered)
hist_img_roi2_filtered = create_3d_hist(preds_roi2_filtered, z_clip=[-300, 600], pix_size=50, sigma=10, contrast_fac=100)

# render, if the FOV is very big, this might be slow and cost large RAM
plt.figure()
plt.subplot(1,2,1)
plt.imshow(hist_img_roi1_filtered)
plt.subplot(1,2,2)
plt.imshow(hist_img_roi2_filtered)

# %% [code]
```
