## Supplementary Software for "Field dependent deep learning enables high-throughput whole-cell 3D super-resolution imaging": demo3_checkpoint.ipynb

```
# %% [markdown]
# FD-DeepLoc training from a chekpoint
Load the saved `.pkl` file and run this script with parameters explained in the `demo1_train.ipynb`

# %% [code]
import datetime
import sys
sys.path.append('../..')

import scipy.io
import seaborn as sns

sns.set_style("white")

from fd_deeploc_core import *
from local_utils import *

sys.setrecursionlimit(10000)
setup_seed(20)


# %% [code]
if __name__ == '__main__':
    with open("../../demo_datasets/demo3_FD_dmo3tetrapod_neuron/demo3_FD-DeepLoc.pkl", 'rb') as f:
        model = pickle.load(f)

    # set the training parameters and retrain
    model.fit(batch_size=10, max_iters=30500, print_freq=500, print_output=True)

# %% [code]
```
