## Supplementary Software for "Field dependent deep learning enables high-throughput whole-cell 3D super-resolution imaging": __init__.py

```
# import all functions of all modules in this package
from fd_deeploc_core.network import *
from fd_deeploc_core.data_simulator import *
from fd_deeploc_core.model import *
from fd_deeploc_core.train_loss_rec import *
from fd_deeploc_core.anlz_eval import *
```
