## Supplementary Software for "Field dependent deep learning enables high-throughput whole-cell 3D super-resolution imaging": anlz_eval.py

```
import numpy as np
import torch.nn.functional as func
from torch.cuda.amp import autocast as autocast
from tqdm import tqdm
from scipy.spatial.distance import cdist
import os

from local_utils import *

def decode_func(model, images, field_xy, batch_size=100, z_scale=10, int_scale=10, use_tqdm=False):
    """Performs inference for a given set of images.
    
    Parameters
    ----------
    model: Model
        FD-DeepLoc model
    images: numpy array
        Three dimensional array of SMLM images
    batch_size: int
        Images are proccessed in batches of the given size. 
        When the images are large, the batch size has to be lowered to save GPU memory. 
    z_scale: float
        The model outputs z values between -1 and 1 that are rescaled.
    int_scale: float
        The model outputs photon values between 0 and 1 that are rescaled.
        
    Returns
    -------
    infs: dict
        Dictionary of arrays with the rescaled network outputs
    """

    with torch.no_grad():
        N = len(images)
        if N != 1:
            images = np.concatenate([images[1:2], images, images[-2:-1]], 0).astype('float32')
        else:
            images = np.concatenate([images, images, images], 0).astype('float32')

        if use_tqdm:
            tqdm_func = tqdm
        else:
            def tqdm_func(x):
                return x

        infs = {'Probs': [], 'XO': [], 'YO': [], 'ZO': [], 'Int': []}
        if model.psf_pred:
            infs['BG'] = []
        if model.sig_pred:
            infs['XO_sig'] = []
            infs['YO_sig'] = []
            infs['ZO_sig'] = []
            infs['Int_sig'] = []

        for i in tqdm_func(range(int(np.ceil(N / batch_size)))):
            p, xyzi, xyzi_sig, bg = model.inferring(X=gpu(images[i * batch_size:(i + 1) * batch_size + 2]),
                                                    field_xy=field_xy,
                                                    aber_map_size=[model.dat_generator.aber_map.shape[1],
                                                                   model.dat_generator.aber_map.shape[0]])

            infs['Probs'].append(p[1:-1].cpu())  # index[1：-1] because the input stack was concatenated with two images
            infs['XO'].append(xyzi[1:-1, 0].cpu())
            infs['YO'].append(xyzi[1:-1, 1].cpu())
            infs['ZO'].append(xyzi[1:-1, 2].cpu())
            infs['Int'].append(xyzi[1:-1, 3].cpu())
            if model.sig_pred:
                infs['XO_sig'].append(xyzi_sig[1:-1, 0].cpu())
                infs['YO_sig'].append(xyzi_sig[1:-1, 1].cpu())
                infs['ZO_sig'].append(xyzi_sig[1:-1, 2].cpu())
                infs['Int_sig'].append(xyzi_sig[1:-1, 3].cpu())
            if model.psf_pred:  # rescale the psf estimation
                infs['BG'].append(bg[1:-1].cpu() * model.dat_generator.psf_pars['ph_scale'] / 10
                                  * model.dat_generator.simulation_pars['qe'] *
                                  model.dat_generator.simulation_pars['em_gain']
                                  / model.dat_generator.simulation_pars['e_per_adu'])

        for k in infs.keys():
            infs[k] = np.vstack(infs[k])

        # scale the predictions
        infs['ZO'] = z_scale * infs['ZO']
        infs['Int'] = int_scale * infs['Int']
        if model.sig_pred:
            infs['Int_sig'] = int_scale * infs['Int_sig']
            infs['ZO_sig'] = z_scale * infs['ZO_sig']
        return infs


def nms_sampling(res_dict, threshold=0.3, candi_thre=0.3, batch_size=500, nms=True, nms_cont=False):
    """Performs Non-maximum Suppression to obtain deterministic samples from the probabilities provided by the decode function. 
    
    Parameters
    ----------
    res_dict: dict
        Dictionary of arrays created with decode_func
    threshold: float
        Processed probabilities above this threshold are considered as final detections
    candi_thre:float
        Probabilities above this threshold are treated as candidates
    batch_size: int
        Outputs are proccessed in batches of the given size. 
        When the arrays are large, the batch size has to be lowered to save GPU memory. 
    nms: bool
        If False don't perform Non-maximum Suppression and simply applies a theshold to the probablities to obtain detections
    nms_cont: bool
        If true also averages the offset variables according to the probabilties that count towards a given detection
        
    Returns
    -------
    res_dict: dict
        Dictionary of arrays where 'Samples_ps' contains the final detections
    """

    res_dict['Probs_ps'] = res_dict['Probs'] + 0  # after nms, this is the final-probability
    res_dict['XO_ps'] = res_dict['XO'] + 0
    res_dict['YO_ps'] = res_dict['YO'] + 0
    res_dict['ZO_ps'] = res_dict['ZO'] + 0

    if nms:

        N = len(res_dict['Probs'])
        for i in range(int(np.ceil(N / batch_size))):
            sl = np.index_exp[i * batch_size:(i + 1) * batch_size]
            if nms_cont:
                res_dict['Probs_ps'][sl], res_dict['XO_ps'][sl], res_dict['YO_ps'][sl], res_dict['ZO_ps'][sl] \
                    = nms_func(res_dict['Probs'][sl], candi_thre,
                               res_dict['XO'][sl], res_dict['YO'][sl], res_dict['ZO'][sl])
            else:
                res_dict['Probs_ps'][sl] = nms_func(res_dict['Probs'][sl], candi_thre=candi_thre)

    res_dict['Samples_ps'] = np.where(res_dict['Probs_ps'] > threshold, 1, 0)  # deterministic locs


def rescale(arr_infs, rescale_bins=50, sig_3d=False):
    """Rescales x and y offsets (inplace) so that they are distributed uniformly within [-0.5, 0.5] to correct for biased outputs. 
    
    Parameters
    ----------
    arr_infs: dict
        Dictionary of arrays created with decode_func and nms_sampling
    rescale_bins: int
        The bias scales with the uncertainty of the localization. Therefore all detections are binned according to their predicted uncertainty.
        Detections within different bins are then rescaled seperately. This specifies the number of bins. 
    sig_3d: bool
        If true also the uncertainty in z when performing the binning
    """
    if arr_infs['Samples_ps'].sum() > 0:

        s_inds = arr_infs['Samples_ps'].nonzero()

        x_sig_var = np.var(arr_infs['XO_sig'][s_inds])
        y_sig_var = np.var(arr_infs['YO_sig'][s_inds])
        z_sig_var = np.var(arr_infs['ZO_sig'][s_inds])

        tot_sig = arr_infs['XO_sig'] ** 2 + (np.sqrt(x_sig_var / y_sig_var) * arr_infs['YO_sig']) ** 2
        if sig_3d:
            tot_sig += (np.sqrt(x_sig_var / z_sig_var) * arr_infs['ZO_sig']) ** 2

        arr = np.where(arr_infs['Samples_ps'], tot_sig, 0)
        bins = histedges_equalN(arr[s_inds], rescale_bins)
        for i in range(rescale_bins):
            inds = np.where((arr > bins[i]) & (arr < bins[i + 1]) & (arr != 0))
            arr_infs['XO_ps'][inds] = uniformize(arr_infs['XO_ps'][inds]) + np.mean(arr_infs['XO_ps'][inds])
            arr_infs['YO_ps'][inds] = uniformize(arr_infs['YO_ps'][inds]) + np.mean(arr_infs['YO_ps'][inds])


def array_to_list(infs, wobble=[0, 0], pix_nm=[100, 100], drifts=None, start_img=0, start_n=0):
    """Transform the the output of the model (dictionary of outputs at imaging resolution) into a list of predictions.
    
    Parameters
    ----------
    infs: dict
        Dictionary of arrays created with decode_func
    wobble: list of floats
        When working with challenge data two constant offsets can be substracted from the x,y variables to
        account for shifts introduced in the PSF fitting.
    pix_nm: list of floats
        x, y pixel size (nano meter)
    drifts:
        If drifts is not None, add the drifts to the xyz.
    start_img: int
        When processing data in multiple batches this variable should be set to the last image count of the
        previous batch to get continuous counting
    start_n: int
        When processing data in multiple batches this variable should be set to the last localization count
        of the previous batch to get continuous counting
        
    Returns
    -------
    res_dict: pred_list
        List of localizations with columns: 'localization', 'frame', 'x', 'y', 'z', 'intensity',
        'x_sig', 'y_sig', 'z_sig'
    """
    samples = infs['Samples_ps']  # determine which pixel has a molecule
    # probs = infs['Probs_ps']

    if drifts is None:
        drifts = np.zeros([len(samples), 4])

    pred_list = []
    count = 1 + start_n

    num_areas = len(infs)
    n = np.sqrt(num_areas)

    for i in range(len(samples)):
        pos = np.nonzero(infs['Samples_ps'][i])  # get the deterministic pixel position
        xo = infs['XO_ps'][i] - drifts[i, 1]
        yo = infs['YO_ps'][i] - drifts[i, 2]
        zo = infs['ZO_ps'][i] - drifts[i, 3]
        ints = infs['Int'][i]
        p_nms = infs['Probs_ps'][i]

        if 'XO_sig' in infs:
            xos = infs['XO_sig'][i]
            yos = infs['YO_sig'][i]
            zos = infs['ZO_sig'][i]
            int_sig = infs['Int_sig'][i]

        for j in range(len(pos[0])):
            pred_list.append([count, i + 1 + start_img,
                              (0.5 + pos[1][j] + xo[pos[0][j], pos[1][j]]) * pix_nm[0] + wobble[0],
                              (0.5 + pos[0][j] + yo[pos[0][j], pos[1][j]]) * pix_nm[1] + wobble[1],
                              zo[pos[0][j], pos[1][j]], ints[pos[0][j], pos[1][j]],
                              p_nms[pos[0][j], pos[1][j]]])
            if 'XO_sig' in infs:
                pred_list[-1] += [xos[pos[0][j], pos[1][j]] * pix_nm[0], yos[pos[0][j], pos[1][j]] * pix_nm[1],
                                  zos[pos[0][j], pos[1][j]], int_sig[pos[0][j], pos[1][j]]]
            else:
                pred_list[-1] += [None, None, None, None]
            count += 1

    return pred_list


def filt_preds(preds, nms_p_thre=0.7, sig_perc=100, is_3d=True):
    """Removes the localizations with the highest uncertainty estimate
    
    Parameters
    ----------
    preds: list
        List of localizations
    sig_perc: float between 0 and 100
        Percentage of localizations that remain
    is_3d: int
        If false only uses x and y uncertainty to filter
        
    Returns
    -------
    preds: list
        List of remaining localizations
    """
    if len(preds):
        # filter by nms_p
        preds = np.array(preds, dtype=np.float32)
        nmsp = preds[:, 6]
        filt_nmsp_index = np.where(nmsp >= nms_p_thre)
        preds = preds[filt_nmsp_index]

        if len(preds):
            if preds[0][-1] is not None:
                x_sig_var = np.var(preds[:, -4])
                y_sig_var = np.var(preds[:, -3])
                z_sig_var = np.var(preds[:, -2])

                tot_var = preds[:, -4] ** 2 + (np.sqrt(x_sig_var / y_sig_var) * preds[:, -3]) ** 2
                if is_3d:
                    tot_var += (np.sqrt(x_sig_var / z_sig_var) * preds[:, -2]) ** 2

                max_s = np.percentile(tot_var, sig_perc)
                filt_sig = np.where(tot_var <= max_s)

                preds = list(preds[filt_sig])

    return list(preds)


def filt_preds_xyz(preds, nms_p_thre=0.7, sigma_x=100, sigma_y=100, sigma_z=100, is_3d=True):
    """Removes the localizations < nms_p threshold and > sigma_xyz

    Parameters
    ----------
    preds: list
        List of localizations
    sig_perc: float between 0 and 100
        Percentage of localizations that remain
    is_3d: int
        If false only uses x and y uncertainty to filter

    Returns
    -------
    preds: list
        List of remaining localizations
    """
    if len(preds):
        # filter by nms_p
        preds = np.array(preds, dtype=np.float32)
        nmsp = preds[:, 6]
        filt_nmsp_index = np.where(nmsp >= nms_p_thre)
        preds = preds[filt_nmsp_index]

        if len(preds):
            if preds[0][-1] is not None:
                # filter by sigma_xyz
                filt_sigmaxyz_index = np.where(
                    (preds[:, -4] <= sigma_x) & (preds[:, -3] <= sigma_y) & (preds[:, -2] <= sigma_z))
                preds = preds[filt_sigmaxyz_index]

    return list(preds)


def nms_func(p, candi_thre=0.3, xo=None, yo=None, zo=None):
    with torch.no_grad():
        diag = 0  # 1/np.sqrt(2)

        p = gpu(p)

        p_copy = p + 0

        # probability values > 0.3 are regarded as possible locations

        # p_clip = torch.where(p > 0.3, p, torch.zeros_like(p))[:, None]
        p_clip = torch.where(p > candi_thre, p, torch.zeros_like(p))[:, None]  # fushuang

        # localize maximum values within a 3x3 patch

        pool = func.max_pool2d(p_clip, 3, 1, padding=1)
        max_mask1 = torch.eq(p[:, None], pool).float()

        # Add probability values from the 4 adjacent pixels

        filt = np.array([[diag, 1, diag], [1, 1, 1], [diag, 1, diag]], ndmin=4)
        conv = func.conv2d(p[:, None], gpu(filt), padding=1)
        p_ps1 = max_mask1 * conv

        # In order do be able to identify two fluorophores in adjacent pixels we look for probablity values > 0.6 that are not part of the first mask

        p_copy *= (1 - max_mask1[:, 0])
        p_clip = torch.where(p_copy > 0.6, p_copy, torch.zeros_like(p_copy))[:, None]  # fushuang
        max_mask2 = torch.where(p_copy > 0.6, torch.ones_like(p_copy), torch.zeros_like(p_copy))[:, None]  # fushuang
        p_ps2 = max_mask2 * conv

        # This is our final clustered probablity which we then threshold (normally > 0.7) to get our final discrete locations 
        p_ps = p_ps1 + p_ps2

        if xo is None:
            return p_ps[:, 0].cpu()

        xo = gpu(xo)
        yo = gpu(yo)
        zo = gpu(zo)

        max_mask = torch.clamp(max_mask1 + max_mask2, 0, 1)

        mult_1 = max_mask1 / p_ps1
        mult_1[torch.isnan(mult_1)] = 0
        mult_2 = max_mask2 / p_ps2
        mult_2[torch.isnan(mult_2)] = 0

        # The rest is weighting the offset variables by the probabilities

        z_mid = zo * p
        z_conv1 = func.conv2d((z_mid * (1 - max_mask2[:, 0]))[:, None], gpu(filt), padding=1)
        z_conv2 = func.conv2d((z_mid * (1 - max_mask1[:, 0]))[:, None], gpu(filt), padding=1)

        zo_ps = z_conv1 * mult_1 + z_conv2 * mult_2
        zo_ps[torch.isnan(zo_ps)] = 0

        x_mid = xo * p
        x_mid_filt = np.array([[0, 1, 0], [0, 1, 0], [0, 1, 0]], ndmin=4)
        xm_conv1 = func.conv2d((x_mid * (1 - max_mask2[:, 0]))[:, None], gpu(x_mid_filt), padding=1)
        xm_conv2 = func.conv2d((x_mid * (1 - max_mask1[:, 0]))[:, None], gpu(x_mid_filt), padding=1)

        x_left = (xo + 1) * p
        x_left_filt = flip_filt(np.array([[diag, 0, 0], [1, 0, 0], [diag, 0, 0]], ndmin=4))
        xl_conv1 = func.conv2d((x_left * (1 - max_mask2[:, 0]))[:, None], gpu(x_left_filt), padding=1)
        xl_conv2 = func.conv2d((x_left * (1 - max_mask1[:, 0]))[:, None], gpu(x_left_filt), padding=1)

        x_right = (xo - 1) * p
        x_right_filt = flip_filt(np.array([[0, 0, diag], [0, 0, 1], [0, 0, diag]], ndmin=4))
        xr_conv1 = func.conv2d((x_right * (1 - max_mask2[:, 0]))[:, None], gpu(x_right_filt), padding=1)
        xr_conv2 = func.conv2d((x_right * (1 - max_mask1[:, 0]))[:, None], gpu(x_right_filt), padding=1)

        xo_ps = (xm_conv1 + xl_conv1 + xr_conv1) * mult_1 + (xm_conv2 + xl_conv2 + xr_conv2) * mult_2

        y_mid = yo * p
        y_mid_filt = np.array([[0, 0, 0], [1, 1, 1], [0, 0, 0]], ndmin=4)
        ym_conv1 = func.conv2d((y_mid * (1 - max_mask2[:, 0]))[:, None], gpu(y_mid_filt), padding=1)
        ym_conv2 = func.conv2d((y_mid * (1 - max_mask1[:, 0]))[:, None], gpu(y_mid_filt), padding=1)

        y_up = (yo + 1) * p
        y_up_filt = flip_filt(np.array([[diag, 1, diag], [0, 0, 0], [0, 0, 0]], ndmin=4))
        yu_conv1 = func.conv2d((y_up * (1 - max_mask2[:, 0]))[:, None], gpu(y_up_filt), padding=1)
        yu_conv2 = func.conv2d((y_up * (1 - max_mask1[:, 0]))[:, None], gpu(y_up_filt), padding=1)

        y_down = (yo - 1) * p
        y_down_filt = flip_filt(np.array([[0, 0, 0], [0, 0, 0], [diag, 1, diag]], ndmin=4))
        yd_conv1 = func.conv2d((y_down * (1 - max_mask2[:, 0]))[:, None], gpu(y_down_filt), padding=1)
        yd_conv2 = func.conv2d((y_down * (1 - max_mask1[:, 0]))[:, None], gpu(y_down_filt), padding=1)

        yo_ps = (ym_conv1 + yu_conv1 + yd_conv1) * mult_1 + (ym_conv2 + yu_conv2 + yd_conv2) * mult_2

        return p_ps[:, 0].cpu(), xo_ps[:, 0].cpu(), yo_ps[:, 0].cpu(), zo_ps[:, 0].cpu()


def cdf_get(cdf, val):
    ind = (val + 1) / 2 * 200 - 1.
    dec = ind - np.floor(ind)

    return dec * cdf[[int(i) + 1 for i in ind]] + (1 - dec) * cdf[[int(i) for i in ind]]


def uniformize(x):
    x = np.clip(x, -0.99, 0.99)
    x_cdf = np.histogram(x, bins=np.linspace(-1, 1, 201))
    x_re = cdf_get(np.cumsum(x_cdf[0]) / sum(x_cdf[0]), x)

    return x_re - 0.5


def histedges_equalN(x, nbin):
    npt = len(x)
    return np.interp(np.linspace(0, npt, nbin + 1),
                     np.arange(npt),
                     np.sort(x))


def list_to_arr(list_or_path, img_size=64, pix_xy=[100, 100], wobble=[0, 0]):
    if isinstance(list_or_path, str):
        pos = []
        reader = csv.reader(open(list_or_path, 'r'))
        for row in reader:
            gt, f, x, y, z, i = row
            if f != 'frame':
                pos.append([float(gt), float(f), float(x) - wobble[0], float(y) - wobble[1], float(z), float(i)])

        pos = np.array(pos).T.astype('float32')
    else:
        pos = np.array(list_or_path).T

    N = int(pos[1].max())

    locs = {'Samples': np.zeros([N, img_size, img_size]), 'XO': np.zeros([N, img_size, img_size]),
            'YO': np.zeros([N, img_size, img_size]), 'ZO': np.zeros([N, img_size, img_size]),
            'Int': np.zeros([N, img_size, img_size])}

    for i in range(N):
        curr = pos[:, np.where(pos[1] - 1 == i)][:, 0]
        for p in curr.T:
            x_ind = np.min([int(p[2] / pix_xy[0]), img_size - 1])
            y_ind = np.min([int(p[3] / pix_xy[0]), img_size - 1])
            if locs['Samples'][i, y_ind, x_ind] == 0:
                locs['Samples'][i, y_ind, x_ind] = 1
                locs['XO'][i, y_ind, x_ind] -= (x_ind * 100 - p[2]) / 100 + 0.5
                locs['YO'][i, y_ind, x_ind] -= (y_ind * 100 - p[3]) / 100 + 0.5
                locs['ZO'][i, y_ind, x_ind] = p[4]
                locs['Int'][i, y_ind, x_ind] = p[5]

    return locs


def assess(test_frame_nbr, test_csv, pred_inp, size_xy=[204800, 204800], tolerance=250, border=450,
                print_res=False, min_int=False, tolerance_ax=np.inf, segmented=False):
    """
    Matches localizations to ground truth positions and provides assessment metrics used in the
    SMLM2016 challenge.

    Parameters
    ----------
    test_frame_nbr:
        number of frames that be analyzed
    test_csv:
        Ground truth positions with columns: 'localization', 'frame', 'x', 'y', 'z', 'photons'
        Either list or str with locations of csv file.
    pred_inp:
        List of predicted localizations
    size_xy:
        Size of the FOV, which contains localizations need to be assessed (nano meter)
    tolerance:
        Localizations are matched when they are within a circle of the given radius.
    border:
        Localizations that are close to the edge of the recording are excluded because they often suffer from artifacts.
    print_res:
        If true prints a list of assessment metrics.
    min_int:
         If true only uses the brightest 75% of ground truth locations.
        This is the setting used in the leaderboard of the challenge.
        However this implementation does not exactly match the method used in the localization tool.
    tolerance_ax:
        Localizations are matched when they are closer than this value in z direction.
        Should be infinity for 2D recordings. 500nm is used for 3D recordings in the challenge.
    segmented:
        If true outputs localization evaluations of different regions of image.(not completed, unnecessary)
    Returns
    -------
    perf_dict, matches: dict, list
        Dictionary of perfomance metrics.
        List of all matches localizations for further evaluation in format: [x_gt, y_gt, z_gt, intensity_gt,
        x_pred, y_pred, z_pred,	intensity_pred,	nms_p, x_sig, y_sig, z_sig]
    """

    perf_dict = None
    matches = []

    test_list = []
    if isinstance(test_csv, str):
        with open(test_csv, 'r') as csvfile:
            reader = csv.reader(csvfile, delimiter=',')
            for row in reader:
                if 'truth' not in row[0]:
                    test_list.append([float(r) for r in row])
    else:
        for r in test_csv:
            test_list.append([i for i in r])

    test_list = sorted(test_list, key=itemgetter(1))  # csv文件按frame升序来排列
    test_list = test_list[:find_frame(test_list, test_frame_nbr)]  # 只要和eval_images对应帧数上的molecule list
    print('{}{}{}{}{}'.format('\nevaluation on ', test_frame_nbr,
                              ' images, ', 'contain ground truth: ', len(test_list)), end='')

    # If true only uses the brightest 75% of ground truth locations.
    if min_int:
        min_int = np.percentile(np.array(test_list)[:, -1], 25)
    else:
        min_int = 0

    if isinstance(pred_inp, str):
        pred_list = []
        with open(pred_inp, 'r') as csvfile:
            reader = csv.reader(csvfile, delimiter=',')
            for row in reader:
                if 'truth' not in row[0]:
                    pred_list.append([float(r) for r in row])

    pred_list = copy.deepcopy(pred_inp)
    print('{}{}'.format(', preds:', len(pred_list)))

    if len(pred_list) == 0:
        perf_dict = {'recall': np.nan, 'precision': np.nan, 'jaccard': np.nan, 'f_score': np.nan, 'rmse_lat': np.nan,
                     'rmse_ax': np.nan,
                     'rmse_x': np.nan, 'rmse_y': np.nan, 'jor': np.nan, 'eff_lat': np.nan, 'eff_ax': np.nan,
                     'eff_3d': np.nan}
        print('original pred_list is empty!')
        return perf_dict, matches

    perf_dict, matches = limited_matching(test_list, pred_list, min_int, limited_x=[0, size_xy[0]],
                                          limited_y=[0, size_xy[1]], border=border,
                                          print_res=print_res, tolerance=tolerance, tolerance_ax=tolerance_ax)

    if segmented:
        _, _1 = limited_matching(test_list, pred_list, min_int, limited_x=[0, 12800],
                                 limited_y=[0, 12800], border=border,
                                 print_res=print_res, tolerance=tolerance, tolerance_ax=tolerance_ax)

        _, _1 = limited_matching(test_list, pred_list, min_int, limited_x=[38400, 51200],
                                 limited_y=[0, 12800], border=border,
                                 print_res=print_res, tolerance=tolerance, tolerance_ax=tolerance_ax)

        _, _1 = limited_matching(test_list, pred_list, min_int, limited_x=[12800, 25600],
                                 limited_y=[12800, 25600], border=border,
                                 print_res=print_res, tolerance=tolerance, tolerance_ax=tolerance_ax)

        _, _1 = limited_matching(test_list, pred_list, min_int, limited_x=[0, 12800],
                                 limited_y=[38400, 51200], border=border,
                                 print_res=print_res, tolerance=tolerance, tolerance_ax=tolerance_ax)

        _, _1 = limited_matching(test_list, pred_list, min_int, limited_x=[38400, 51200],
                                 limited_y=[38400, 51200], border=border,
                                 print_res=print_res, tolerance=tolerance, tolerance_ax=tolerance_ax)

    return perf_dict, matches


def limited_matching(truth_origin, pred_list_origin, min_int, limited_x=[0, 204800], limited_y=[0, 204800],
                     border=450, print_res=True, tolerance=250, tolerance_ax=np.inf):
    print('{}{}{}{}'.format('FOV: x=', limited_x, ' y=', limited_y))

    matches = []

    truth = copy.deepcopy(truth_origin)
    pred_list = copy.deepcopy(pred_list_origin)

    truth_array = np.array(truth)
    pred_array = np.array(pred_list)

    # filter prediction and gt according to limited_x;y
    t_inds = np.where(
        (truth_array[:, 2] < limited_x[0]) | (truth_array[:, 2] > limited_x[1]) |
        (truth_array[:, 3] < limited_y[0]) | (truth_array[:, 3] > limited_y[1]))
    p_inds = np.where(
        (pred_array[:, 2] < limited_x[0]) | (pred_array[:, 2] > limited_x[1]) |
        (pred_array[:, 3] < limited_y[0]) | (pred_array[:, 3] > limited_y[1]))
    for t in reversed(t_inds[0]):
        del (truth[t])
    for p in reversed(p_inds[0]):
        del (pred_list[p])

    if len(pred_list) == 0:
        perf_dict = {'recall': np.nan, 'precision': np.nan, 'jaccard': np.nan, 'f_score': np.nan, 'rmse_lat': np.nan,
                     'rmse_ax': np.nan,
                     'rmse_x': np.nan, 'rmse_y': np.nan, 'jor': np.nan, 'eff_lat': np.nan, 'eff_ax': np.nan,
                     'eff_3d': np.nan}
        print('after FOV segmentation, pred_list is empty!')
        return perf_dict, matches

    # delete molecules of ground truth/estimation in the margin area
    if border:
        test_arr = np.array(truth)
        pred_arr = np.array(pred_list)

        t_inds = np.where(
            (test_arr[:, 2] < limited_x[0] + border) | (test_arr[:, 2] > (limited_x[1] - border)) |
            (test_arr[:, 3] < limited_y[0] + border) | (test_arr[:, 3] > (limited_y[1] - border)))
        p_inds = np.where(
            (pred_arr[:, 2] < limited_x[0] + border) | (pred_arr[:, 2] > (limited_x[1] - border)) |
            (pred_arr[:, 3] < limited_y[0] + border) | (pred_arr[:, 3] > (limited_y[1] - border)))
        for t in reversed(t_inds[0]):
            del (truth[t])
        for p in reversed(p_inds[0]):
            del (pred_list[p])

    if len(pred_list) == 0:
        perf_dict = {'recall': np.nan, 'precision': np.nan, 'jaccard': np.nan, 'f_score': np.nan, 'rmse_lat': np.nan,
                     'rmse_ax': np.nan,
                     'rmse_x': np.nan, 'rmse_y': np.nan, 'jor': np.nan, 'eff_lat': np.nan, 'eff_ax': np.nan,
                     'eff_3d': np.nan}
        print('after border, pred_list is empty!')
        return perf_dict, matches

    print('{}{}{}{}{}'.format('after FOV and border segmentation,'
                              , 'truth: ', len(truth), ' ,preds: ', len(pred_list)))

    TP = 0
    FP = 0.0001
    FN = 0.0001
    MSE_lat = 0
    MSE_ax = 0
    MSE_vol = 0

    if len(pred_list):
        for i in range(1, int(truth_origin[-1][1]) + 1):  # traverse all gt frames

            tests = []  # gt in each frame
            preds = []  # prediction in each frame

            if len(truth) > 0:  # after border filtering and area segmentation, truth could be empty
                while truth[0][1] == i:
                    tests.append(truth.pop(0))  # put all gt in the tests
                    if len(truth) < 1:
                        break
            if len(pred_list) > 0:
                while pred_list[0][1] == i:
                    preds.append(pred_list.pop(0))  # put all predictions in the preds
                    if len(pred_list) < 1:
                        break

            # if preds is empty, it means no detection on the frame, all tests are FN
            if len(preds) == 0:
                FN += len(tests)
                continue  # no need to calculate metric
            # if the gt of this frame is empty, all preds on this frame are FP
            if len(tests) == 0:
                FP += len(preds)
                continue  # no need to calculate metric

            # calculate the Euclidean distance between all gt and preds, get a matrix [number of gt, number of preds]
            dist_arr = cdist(np.array(tests)[:, 2:4], np.array(preds)[:, 2:4])
            ax_arr = cdist(np.array(tests)[:, 4:5], np.array(preds)[:, 4:5])
            tot_arr = np.sqrt(dist_arr ** 2 + ax_arr ** 2)

            if tolerance_ax == np.inf:
                tot_arr = dist_arr

            match_tests = copy.deepcopy(tests)
            match_preds = copy.deepcopy(preds)

            if dist_arr.size > 0:
                while dist_arr.min() < tolerance:
                    r, c = np.where(tot_arr == tot_arr.min())  # select the positions pair with shortest distance
                    r = r[0]
                    c = c[0]
                    if ax_arr[r, c] < tolerance_ax and dist_arr[r, c] < tolerance:  # compare the distance and tolerance
                        if match_tests[r][-1] > min_int:  # photons should be larger than min_int

                            MSE_lat += dist_arr[r, c] ** 2
                            MSE_ax += ax_arr[r, c] ** 2
                            MSE_vol += dist_arr[r, c] ** 2 + ax_arr[r, c] ** 2
                            TP += 1
                            matches.append([match_tests[r][2], match_tests[r][3], match_tests[r][4], match_tests[r][5],
                                            match_preds[c][2], match_preds[c][3], match_preds[c][4], match_preds[c][5],
                                            match_preds[c][6], match_preds[c][-4], match_preds[c][-3],
                                            match_preds[c][-2]])

                        dist_arr[r, :] = np.inf
                        dist_arr[:, c] = np.inf
                        tot_arr[r, :] = np.inf
                        tot_arr[:, c] = np.inf

                        tests[r][-1] = -100  # photon cannot be negative, work as a flag
                        preds.pop()

                    dist_arr[r, c] = np.inf
                    tot_arr[r, c] = np.inf

            for i in reversed(range(len(tests))):
                if tests[i][-1] < min_int:  # delete matched gt
                    del (tests[i])

            FP += len(preds)  # all remaining preds are FP
            FN += len(tests)  # all remaining gt are FN

    else:
        print('after border and FOV segmentation, pred list is empty!')

    precision = TP / (TP + FP)
    recall = TP / (TP + FN)
    jaccard = TP / (TP + FP + FN)
    rmse_lat = np.sqrt(MSE_lat / (TP + 0.00001))
    rmse_ax = np.sqrt(MSE_ax / (TP + 0.00001))
    rmse_vol = np.sqrt(MSE_vol / (TP + 0.00001))
    jor = 100 * jaccard / rmse_lat

    eff_lat = 100 - np.sqrt((100 - 100 * jaccard) ** 2 + 1 ** 2 * rmse_lat ** 2)
    eff_ax = 100 - np.sqrt((100 - 100 * jaccard) ** 2 + 0.5 ** 2 * rmse_ax ** 2)
    eff_3d = (eff_lat + eff_ax) / 2

    matches = np.array(matches)
    rmse_x = np.nan
    rmse_y = np.nan
    rmse_z = np.nan
    rmse_i = np.nan
    if len(matches):
        rmse_x = np.sqrt(((matches[:, 0] - matches[:, 4]) ** 2).mean())
        rmse_y = np.sqrt(((matches[:, 1] - matches[:, 5]) ** 2).mean())
        rmse_z = np.sqrt(((matches[:, 2] - matches[:, 6]) ** 2).mean())
        rmse_i = np.sqrt(((matches[:, 3] - matches[:, 7]) ** 2).mean())
    else:
        print('matches is empty!')

    if print_res:
        print('{}{:0.3f}'.format('Recall: ', recall))
        print('{}{:0.3f}'.format('Precision: ', precision))
        print('{}{:0.3f}'.format('Jaccard: ', 100 * jaccard))
        print('{}{:0.3f}'.format('RMSE_lat: ', rmse_lat))
        print('{}{:0.3f}'.format('RMSE_ax: ', rmse_ax))
        print('{}{:0.3f}'.format('RMSE_vol: ', rmse_vol))
        print('{}{:0.3f}'.format('Jaccard/RMSE: ', jor))
        print('{}{:0.3f}'.format('Eff_lat: ', eff_lat))
        print('{}{:0.3f}'.format('Eff_ax: ', eff_ax))
        print('{}{:0.3f}'.format('Eff_3d: ', eff_3d))
        print('FN: ' + str(np.round(FN)) + ' FP: ' + str(np.round(FP)))

    perf_dict = {'recall': recall, 'precision': precision, 'jaccard': jaccard, 'rmse_lat': rmse_lat,
                 'rmse_ax': rmse_ax, 'rmse_vol': rmse_vol, 'rmse_x': rmse_x, 'rmse_y': rmse_y,
                 'rmse_z': rmse_z, 'rmse_i': rmse_i, 'jor': jor, 'eff_lat': eff_lat, 'eff_ax': eff_ax,
                 'eff_3d': eff_3d}

    return perf_dict, matches


def recognition(model, eval_imgs_all, batch_size, use_tqdm, nms, rescale_xy, pix_nm, plot_num,
                wobble=[0, 0], start_field_pos=[0, 0], stack_interval=100000, divide_and_conquer=True, win_size=128,
                padding=True, candi_thre=0.3, nms_thre=0.3):
    """ Analyze the SMLM images using FD-DeepLoc model

    Parameters
    ----------
    model:
        FD-DeepLoc model
    eval_imgs_all:
        Input images that need to be analyzed
    batch_size: int
        Number of images in each batch
    use_tqdm: bool
        Progress bar
    nms: bool
        If False, only use a simple threshold to filter P channel to get the deterministic pixels. If True, add the
        values from the 4 adjacent pixels to local maximums and then filter with the threshold.
    rescale_xy: bool
        Rescales x and y offsets (inplace) so that they are distributed uniformly within [-0.5, 0.5] to
        correct for biased outputs.
    wobble:
        Constant x y offset added to the final molecule list
    pix_nm:
        Pixel size, [X_pixelsize Y_pixelsize]
    plot_num:
        Choose a specific frame to return the output (int or None)
    start_field_pos:
        The global position [x, y] (not [row, column]) of the input images in the whole training aberration map.
        Start from the top left. For example, start_field_pos [102, 41] means the top left pixel (namely local position
        [0, 0]) of the input images is located at [102, 41] of the whole FOV. Thus CoordConv can get the
        global position of the input images.
    stack_interval: int
        Process the long-time frames by time interval. If RAM is out of memory, set it smaller.
    divide_and_conquer: bool
        Divide the large images into small sub-area images. This is necessary as large input images lead to GPU memory
        error, this enables the network process the large images by sub-areas.
    win_size:
        If divide_and_conquer=True, set the size of segmented areas, must be a multiple of 4 to avoid error when down-
        sampling or up-sampling.
    padding: bool
        If divide_and_conquer=True, padding=True, this will cut a larger area (20 pixels) than win_size and traverse
        with overlap to avoid error from incomplete PSFs at margin.
    candi_thre:
        In the probability channel, only pixels with value>candi_thre will be treated as candidates for local maximum
        searching.
    nms_thre:
        This works as a threshold to filter P channel to get the deterministic pixels
    Returns
    -------
    pred_list:
        List of localizations with columns: 'localization', 'frame', 'x', 'y', 'z', 'intensity',
        'x_sig', 'y_sig', 'z_sig'
    """
    N = len(eval_imgs_all)
    plot_data = []
    preds_frames = []
    for stack_num in range(int(np.ceil(N / stack_interval))):
        eval_imgs_tmp = eval_imgs_all[stack_num * stack_interval: (stack_num + 1) * stack_interval if
                        (stack_num + 1) * stack_interval < N else N]

        start_img = stack_num * stack_interval
        start_field = copy.deepcopy(start_field_pos)
        h, w = eval_imgs_tmp.shape[1], eval_imgs_tmp.shape[2]

        # enforce the image size to be multiple of 4, pad with estimated background. start_field_pos should be modified
        # according to the padding, and field_xy for sub-area images should be modified too.
        pad_h = 0
        pad_w = 0
        bg_test_whole_field = None
        if (h % 4 != 0) or (w % 4 != 0):
            bg_test_whole_field, _ = get_bg_stats(eval_imgs_tmp, percentile=50)
            if h % 4 != 0:
                new_h = (h // 4 + 1) * 4
                pad_h = new_h - h
                eval_imgs_tmp = np.pad(eval_imgs_tmp, [[0, 0], [pad_h, 0], [0, 0]],
                                       mode='constant', constant_values=bg_test_whole_field)
                start_field[1] -= pad_h
                h += pad_h
            if w % 4 != 0:
                new_w = (w // 4 + 1) * 4
                pad_w = new_w - w
                eval_imgs_tmp = np.pad(eval_imgs_tmp, [[0, 0], [0, 0], [pad_w, 0]],
                                       mode='constant', constant_values=bg_test_whole_field)
                start_field[0] -= pad_w
                w += pad_w

        n_per_img = 0

        if divide_and_conquer:
            area_rows = int(np.ceil(h / win_size))
            area_columns = int(np.ceil(w / win_size))

            images_areas = []
            origin_areas_list = []
            areas_list = []
            if padding:
                for i in range(area_rows):
                    for j in range(area_columns):
                        x_origin = j * win_size
                        y_origin = i * win_size
                        x_origin_end = w if x_origin + win_size > w else x_origin + win_size
                        y_origin_end = h if y_origin + win_size > h else y_origin + win_size

                        x_start = j * win_size if j * win_size - 20 < 0 else j * win_size - 20
                        y_start = i * win_size if i * win_size - 20 < 0 else i * win_size - 20
                        x_end = w if x_origin + win_size + 20 > w else x_origin + win_size + 20
                        y_end = h if y_origin + win_size + 20 > h else y_origin + win_size + 20

                        sub_imgs_tmp = eval_imgs_tmp[:, y_start:y_end, x_start:x_end]
                        # set the background of eval_images the same as training set
                        bg_test, _ = get_bg_stats(sub_imgs_tmp,percentile=50)
                        sub_imgs_tmp = sub_imgs_tmp - bg_test + model.dat_generator.simulation_pars['backg']
                        images_areas.append(sub_imgs_tmp)

                        areas_list.append([x_start + start_field[0], x_end - 1 + start_field[0],
                                           y_start + start_field[1], y_end - 1 + start_field[1]])

                        origin_areas_list.append([x_origin + start_field[0], x_origin_end - 1 + start_field[0],
                                                  y_origin + start_field[1], y_origin_end - 1 + start_field[1]])
            else:
                for i in range(area_rows):
                    for j in range(area_columns):
                        x_start = j * win_size
                        y_start = i * win_size
                        x_end = w if x_start + win_size > w else x_start + win_size
                        y_end = h if y_start + win_size > h else y_start + win_size

                        sub_imgs_tmp = eval_imgs_tmp[:, y_start:y_end, x_start:x_end]
                        # set the background of eval_images the same as training set
                        bg_test, _ = get_bg_stats(sub_imgs_tmp,percentile=50)
                        sub_imgs_tmp = sub_imgs_tmp - bg_test + model.dat_generator.simulation_pars['backg']
                        images_areas.append(sub_imgs_tmp)

                        areas_list.append([x_start + start_field[0], x_end - 1 + start_field[0],
                                           y_start + start_field[1], y_end - 1 + start_field[1]])

                        origin_areas_list.append([x_start + start_field[0], x_end - 1 + start_field[0],
                                                  y_start + start_field[1], y_end - 1 + start_field[1]])

            if plot_num:
                if 0 <= plot_num - 1 - start_img < N:
                    plot_areas = [{} for i in range(area_rows * area_columns + 1)]
                    plot_areas[0]['raw_img'] = eval_imgs_tmp[plot_num - 1 - start_img]
                    plot_areas[0]['rows'] = area_rows
                    plot_areas[0]['columns'] = area_columns
                    plot_areas[0]['win_size'] = win_size
                else:
                    plot_areas = []
            else:
                plot_areas = []

            del eval_imgs_tmp

            preds_areas = []

            for i in range(area_rows * area_columns):
                field_xy = torch.tensor(areas_list[i])
                # field_xy = torch.tensor(areas_list[area_rows*area_columns-1-i])+\
                #            torch.tensor([100,100,100,100])  # test wrong global position

                if use_tqdm:
                    # print('\nprocessing area:', str(i + 1), '/', str(area_rows * area_columns),
                    #       ', input field_xy:', str(cpu(field_xy)),
                    #       ', use_coordconv:', str(model.net_pars['use_coordconv']),
                    #       ', retain locs in area:', str(origin_areas_list[i]),
                    #       ', aber_map size:', str(model.dat_generator.psf_pars['aber_map'].shape) )
                    print('{}{}{}{}{}{}{}{}{}{}{}{}'.format('\nprocessing area:', i + 1, '/', area_rows * area_columns,
                                                            ', input field_xy:', cpu(field_xy), ', use_coordconv:',
                                                            model.net_pars['use_coordconv'], ', retain locs in area:',
                                                            origin_areas_list[i], ', aber_map size:',
                                                            model.dat_generator.psf_pars['aber_map'].shape))
                else:
                    # print('\rprocessing area:', str(i + 1), '/', str(area_rows * area_columns),
                    #       ', input field_xy:', str(cpu(field_xy)),
                    #       ', use_coordconv:', str(model.net_pars['use_coordconv']),
                    #       ', retain locs in area:', str(origin_areas_list[i]),
                    #       ', aber_map size:', str(model.dat_generator.psf_pars['aber_map'].shape), end='')
                    print('{}{}{}{}{}{}{}{}{}{}{}{}'.format('\rprocessing area:',i+1, '/', area_rows * area_columns,
                                                            ', input field_xy:', cpu(field_xy), ', use_coordconv:',
                                                            model.net_pars['use_coordconv'],', retain locs in area:',
                                                            origin_areas_list[i],', aber_map size:',
                                                            model.dat_generator.psf_pars['aber_map'].shape), end='')

                arr_infs = decode_func(model, images_areas[0],
                                       field_xy=field_xy,
                                       batch_size=batch_size, use_tqdm=use_tqdm,
                                       z_scale=model.dat_generator.psf_pars['z_scale'],
                                       int_scale=model.dat_generator.psf_pars['ph_scale'])

                nms_sampling(arr_infs, threshold=nms_thre, candi_thre=candi_thre, batch_size=batch_size, nms=nms, nms_cont=False)

                if rescale_xy:
                    rescale(arr_infs, 20, sig_3d=False)  # 根据sigma来rescale xyoffset

                preds_list = array_to_list(arr_infs, wobble=wobble, pix_nm=pix_nm, start_img=start_img)

                if padding:
                    # drop the molecules in the overlap between sub-areas (padding), shift the remaining molecules
                    # back to correct positions.
                    preds_list = padding_shift(origin_areas_list[i], areas_list[i], preds_list, pix_nm=pix_nm)

                preds_areas.append(preds_list)

                # calculate the n_per_image, need to take account for the padding
                x_index = int((torch.tensor(origin_areas_list[i]) - torch.tensor(areas_list[i]))[0])
                y_index = int((torch.tensor(origin_areas_list[i]) - torch.tensor(areas_list[i]))[2])
                n_per_img += arr_infs['Probs'][:, y_index: y_index + win_size,
                             x_index: x_index + win_size].sum(-1).sum(-1).mean()

                if plot_num:
                    if 0 <= plot_num - 1 - start_img < N:
                        for k in arr_infs.keys():
                            x_index = int((torch.tensor(origin_areas_list[i]) - torch.tensor(areas_list[i]))[0])
                            y_index = int((torch.tensor(origin_areas_list[i]) - torch.tensor(areas_list[i]))[2])
                            plot_areas[i + 1][k] = copy.deepcopy(arr_infs[k][plot_num - 1 - start_img,
                                                                 y_index: y_index + win_size,
                                                                 x_index: x_index + win_size])

                del arr_infs
                del images_areas[0]

            print('')

            plot_data += plot_areas
            preds_frames += post_process(preds_areas, h, w, pixel_size=pix_nm, win_size=win_size)

        # divide_and_conquer = False
        else:
            if bg_test_whole_field is None:
                bg_test_whole_field, _ = get_bg_stats(eval_imgs_tmp,percentile=50)
            eval_imgs_tmp = eval_imgs_tmp - bg_test_whole_field + model.dat_generator.simulation_pars['backg']

            if plot_num:
                if 0 <= plot_num - 1 - start_img < N:
                    plot_areas = [{} for i in range(2)]
                    plot_areas[0]['raw_img'] = eval_imgs_tmp[plot_num - 1 - start_img]
                    plot_areas[0]['rows'] = 1
                    plot_areas[0]['columns'] = 1
                    plot_areas[0]['win_size'] = max(eval_imgs_tmp.shape[1], eval_imgs_tmp.shape[2])
                else:
                    plot_areas = []
            else:
                plot_areas = []

            field_xy = torch.tensor([start_field[0], start_field[0] + w - 1, start_field[1], start_field[1] + h - 1])

            print('field_xy:', cpu(field_xy), ', use_coordconv:', model.net_pars['use_coordconv'],
                  ', aber_map_size:', str(model.dat_generator.psf_pars['aber_map'].shape) )

            arr_infs = decode_func(model, eval_imgs_tmp,
                                   field_xy=field_xy,
                                   batch_size=batch_size, use_tqdm=use_tqdm,
                                   z_scale=model.dat_generator.psf_pars['z_scale'],
                                   int_scale=model.dat_generator.psf_pars['ph_scale'])

            nms_sampling(arr_infs, threshold=nms_thre, candi_thre=candi_thre, batch_size=batch_size, nms=nms, nms_cont=False)

            if rescale_xy:
                rescale(arr_infs, 20, sig_3d=False)  # rescale xyoffset using uncertainty

            preds_list = array_to_list(arr_infs, wobble=wobble, pix_nm=pix_nm, start_img=start_img)

            n_per_img += arr_infs['Probs'].sum(-1).sum(-1).mean()

            if plot_num:
                if 0 <= plot_num - 1 - start_img < N:
                    for k in arr_infs.keys():
                        plot_areas[1][k] = copy.deepcopy(arr_infs[k][plot_num - 1 - start_img])

            del eval_imgs_tmp
            del arr_infs

            plot_data += plot_areas
            preds_frames += preds_list

    # if modification was made to the raw images to have a size of multiple of 4, this resulted in offsets.
    preds_frames = np.array(preds_frames, dtype=np.float32)
    if len(preds_frames):
        preds_frames[:, 2] -= pad_w * pix_nm[0]
        preds_frames[:, 3] -= pad_h * pix_nm[1]
    preds_raw = sorted(preds_frames.tolist(), key=itemgetter(1))

    return preds_raw, n_per_img, plot_data


def post_process(preds_areas, height, width, pixel_size=[100, 100], win_size=128):
    """transform the sub-area coordinate to whole-filed coordinate"""
    rows = int(np.ceil(height / win_size))
    columns = int(np.ceil(width / win_size))
    preds = []
    for i in range(rows * columns):
        field_xy = [i % columns * win_size, i % columns * win_size + win_size - 1, i // columns * win_size,
                    i // columns * win_size + win_size - 1]
        tmp = preds_areas[i]
        for j in range(len(tmp)):
            tmp[j][2] += field_xy[0] * pixel_size[0]
            tmp[j][3] += field_xy[2] * pixel_size[1]
        preds = preds + tmp

    preds = sorted(preds, key=itemgetter(1))

    return preds


def padding_shift(origin_areas, padded_areas, preds_list, pix_nm):
    """the sub-image cropped is larger than win_size, we need to select the predictions in win_size"""
    if origin_areas[0] == padded_areas[0]:
        x_offset = 0
    else:
        x_offset = 20
    if origin_areas[2] == padded_areas[2]:
        y_offset = 0
    else:
        y_offset = 20

    for i in range(len(preds_list)):
        preds_list[i][2] -= x_offset * pix_nm[0]
        preds_list[i][3] -= y_offset * pix_nm[1]

    preds_shift = []
    for j in range(len(preds_list)):
        if preds_list[j][2] < 0 or preds_list[j][3] < 0 or \
                preds_list[j][2] > (origin_areas[1] - origin_areas[0] + 1) * pix_nm[0] or \
                preds_list[j][3] > (origin_areas[3] - origin_areas[2] + 1) * pix_nm[1]:
            continue

        preds_shift.append(preds_list[j])

    return preds_shift


def read_bigtiff_and_predict(model, image_path, stack_giga=0.5, batch_size=10, use_tqdm=True,
                             nms=True, candi_thre=0.3, nms_thre=0.3,
                             rescale_xy=False, wobble=[0, 0], pixel_size=[100,100], plot_num=None,
                             start_field_pos=[0, 0], stack_interval=100000, divide_and_conquer=True,
                             win_size=256, padding=True, save_path='./pred_list.csv'):
    """ Analyze the SMLM images using FD-DeepLoc model

    Parameters
    ----------
    model:
        FD-DeepLoc model
    image_path:
        Path of images (tiff) that need to be analyzed
    stack_giga:
        As large-FOV image analysis needs a lot of RAM, we load and analyze SMLM images by stack_giga (Gb) sequentially.
    batch_size: int
        Number of images in each batch
    use_tqdm: bool
        Progress bar
    nms: bool
        It False, only use a simple threshold to filter P channel to get the deterministic pixels. If True, add the
        values from the 4 adjacent pixels to local maximums and then filter with the threshold.
    rescale_xy: bool
        Rescales x and y offsets (inplace) so that they are distributed uniformly within [-0.5, 0.5] to
        correct for biased outputs.
    wobble:
        Constant x y offset added to the final molecule list
    pixel_size:
        Pixel size, [X_pixelsize Y_pixelsize]
    plot_num:
        Choose a specific frame to return the output (int or None)
    start_field_pos:
        The global position [x, y] (not [row, column]) of the input images in the whole training aberration map.
        Start from the top left. For example, start_field_pos [102, 41] means the upper left pixel (namely local position
        [0, 0]) of the input images is located at [102, 41] of the whole FOV. Thus CoordConv can get the
        global position of the input images.
    stack_interval: int
        Process the long-time frames by time interval. If RAM is out of memory, set it smaller.
    divide_and_conquer: bool
        Divide the large images into small sub-area images. This is necessary as large input images lead to GPU memory
        error, this enables the model process the large images by sub-areas.
    win_size:
        If divide_and_conquer=True, set the size of segmented areas, must be a multiple of 4 to avoid error when down-
        sampling or up-sampling.
    padding: bool
        If divide_and_conquer=True, padding=True, this will cut a larger area (20 pixels) than win_size and traverse
        with overlap to avoid error from incomplete PSFs at margin.
    candi_thre:
        In the probability channel, only pixels with value>candi_thre will be treated as candidates for local maximum
        searching.
    nms_thre:
        This works as a threshold to filter P channel to get the deterministic pixels
    save_path:
        Directory to save the prediction, after processing stack_giga-sized SMLM images, save the pred_list.csv once.

    Returns
    -------
    pred_list:
        List of localizations with columns: ['Localization number', 'frame', 'xnano', 'ynano', 'znano',
        'intensity','nms_p', 'x_sig', 'y_sig', 'z_sig','I_sig']
    """

    # can not load too big file directly to RAM, so we need to use this way
    print('the file to save the predictions is: ', save_path)
    if os.path.exists(save_path):
        last_preds = read_csv(save_path)
        last_frame_num = int(last_preds[-1, 1])
        del last_preds
        print('append the pred list to existed csv, the last analyzed frame is:', last_frame_num)
    else:
        last_frame_num = 0
        preds_empty = []
        write_csv(preds_empty, name=save_path, write_gt=False, append=False)

    with TiffFile(image_path, is_ome=True) as tif:
        total_shape = tif.series[0].shape
        occu_mem = total_shape[0] * total_shape[1] * total_shape[2] * 16 / (1024 ** 3) / 8
        stack_num = int(np.ceil(occu_mem / stack_giga))
        frames_per_stack = int(total_shape[0] // stack_num)

        fov_size = [total_shape[2] * pixel_size[0], total_shape[1] * pixel_size[1]]

        frames_ind = []
        i = 0
        while (i + 1) * frames_per_stack + last_frame_num <= total_shape[0]:
            frames_ind.append(range(i * frames_per_stack + last_frame_num, (i + 1) * frames_per_stack + last_frame_num))
            i += 1
        if (i * frames_per_stack + last_frame_num < total_shape[0]) &\
                ((i + 1) * frames_per_stack + last_frame_num > total_shape[0]):
            frames_ind.append(range(i * frames_per_stack + last_frame_num, total_shape[0]))

        for tif_ind in range(len(frames_ind)):
            eval_images = tif.asarray(key=frames_ind[tif_ind], series=0)
            if len(eval_images.shape) == 2:
                eval_images = eval_images[np.newaxis, :, :]
            print('{}{}{}{}{}{}{}{}{}{}'.format('stack: ', tif_ind + 1, '/', len(frames_ind), ', contain imgs: ',
                                                eval_images.shape[0], ', already analyzed:', frames_ind[tif_ind].start,
                                                '/', total_shape[0]))

            # print('stack:', tif_ind + 1, '/', len(frames_ind), ', contain imgs:', eval_images.shape[0],
            #       'finished:', frames_ind[tif_ind].start,'/',total_shape[0])

            # # only process specific subarea of the image
            # eval_images = eval_images[:, 48:48+256, 436:436+256]

            # fov_size = [eval_images.shape[2] * pixel_size[0], eval_images.shape[1] * pixel_size[1]]

            # the recognition process
            with autocast():  # speed up, but with precision loss
                preds_tmp, n_per_img, plot_data = recognition(model=model, eval_imgs_all=eval_images,
                                                              batch_size=batch_size,
                                                              use_tqdm=use_tqdm,
                                                              nms=nms, rescale_xy=rescale_xy,
                                                              wobble=wobble,
                                                              pix_nm=pixel_size,
                                                              plot_num=plot_num,
                                                              start_field_pos=start_field_pos,
                                                              stack_interval=stack_interval,
                                                              divide_and_conquer=divide_and_conquer,
                                                              win_size=win_size, padding=padding,
                                                              candi_thre=candi_thre, nms_thre=nms_thre)

            preds_tmp = np.array(preds_tmp)
            preds_tmp[:, 1] += frames_ind[tif_ind].start
            write_csv(preds_tmp.tolist(), name=save_path, write_gt=False, append=True)

            # preds.append(preds_tmp)
    # preds_raw = []
    # for i in range(len(frames_ind)):
    #     tmp = np.array(preds[i])
    #     tmp[:, 1] = tmp[:, 1] + last_tmp[-1, 1] if i > 0 else tmp[:, 1]
    #     preds_raw = preds_raw + tmp.tolist()
    #     last_tmp = tmp
    print('analysis finished ! the file containing results is:', save_path)

    return total_shape, fov_size


def check_specific_frame_output(plot_num, model, image_path, eval_csv=None,
                                nms=True, candi_thre=0.3, nms_thre=0.3, rescale_xy=False, pixel_size=[100,100],
                                start_field_pos=[0, 0], divide_and_conquer=True,
                                win_size=256, padding=True):
    """ Check the network output of a specified single frame.

    Parameters
    ----------
    plot_num:
        The number of the frame that want to check, better be greater than 5 considering temporal context and
        background estimation.
    model:
        FD-DeepLoc model
    image_path:
        Path of images (tiff) that need to be analyzed
    eval_csv:
        List of ground truth localizations with columns: 'localization', 'frame', 'x', 'y', 'z', 'intensity/photons'
    nms: bool
        It False, only use a simple threshold to filter P channel to get the deterministic pixels. If True, add the
        values from the 4 adjacent pixels to local maximums and then filter with the threshold.
    candi_thre:
        In the probability channel, only pixels with value>candi_thre will be treated as candidates for local maximum
        searching.
    nms_thre:
        This works as a threshold to filter P channel to get the deterministic pixels
    rescale_xy: bool
        Rescales x and y offsets (inplace) so that they are distributed uniformly within [-0.5, 0.5] to
        correct for biased outputs.
    pixel_size:
        Pixel size, [X_pixelsize Y_pixelsize]
    start_field_pos:
        The global position [x, y] (not [row, column]) of the input images in the whole training aberration map.
        Start from the top left. For example, start_field_pos [102, 41] means the top left pixel (namely local position
        [0, 0]) of the input images is located at [102, 41] of the whole FOV. Thus CoordConv can get the
        global position of the input images.
    divide_and_conquer: bool
        Divide the large images into small sub-area images. This is necessary as large input images lead to GPU memory
        error, this enables the model process the large images by sub-areas.
    win_size:
        If divide_and_conquer=True, set the size of segmented areas, must be a multiple of 4 to avoid error when down-
        sampling or up-sampling.
    padding: bool
        If divide_and_conquer=True, padding=True, this will cut a larger area (20 pixels) than win_size and traverse
        with overlap to avoid error from incomplete PSFs at margin.
    Returns
    -------

    """
    with TiffFile(image_path, is_ome=True) as tif:
        total_shape = tif.series[0].shape
        frames_ind = range(plot_num - 5, plot_num + 4) if 5 <= plot_num < total_shape[0] \
            else [plot_num-1,plot_num-1,plot_num-1,plot_num-1,plot_num-1,
                  plot_num-1,plot_num-1,plot_num-1,plot_num-1]

        eval_images = tif.asarray(key=frames_ind, series=0)

        # # only process specific subarea of the image
        # eval_images = eval_images[:, 48:48+256, 436:436+256]

        fov_size = [eval_images.shape[2] * pixel_size[0], eval_images.shape[1] * pixel_size[1]]

        # the recognition process
        with autocast():  # speed up, but with precision loss
            preds_tmp, n_per_img, plot_data = recognition(model=model, eval_imgs_all=eval_images,
                                                          batch_size=1,
                                                          use_tqdm=False,
                                                          nms=nms, candi_thre=candi_thre, nms_thre=nms_thre,
                                                          rescale_xy=rescale_xy,
                                                          wobble=[0,0],
                                                          pix_nm=pixel_size,
                                                          plot_num=5,
                                                          start_field_pos=start_field_pos,
                                                          stack_interval=20000,
                                                          divide_and_conquer=divide_and_conquer,
                                                          win_size=win_size, padding=padding)
    plot_sample_predictions(model, plot_infs=plot_data, eval_csv=eval_csv, plot_num=plot_num,
                            fov_size=fov_size, pixel_size=pixel_size)


def check_local_CRLB(model, test_pos, test_photons, test_bg, Nmol=25):
    """
    Check the CRLB of the local PSF at given positions
    Parameters
    ----------
    model:
        FD-DeepLoc model
    test_pos:
        Position (x,y) where the PSF CRLB will be computed and checked
    test_photons:
        Number of signal photons used to compute CRLB
    test_bg:
        Number of background photons used to compute CRLB
    Nmol:
        Number of sampled z positions, which are uniformly distributed in the training z range
    Returns
    -------

    """
    # set PSF parameters
    NA = model.dat_generator.psf_pars['NA']
    refmed = model.dat_generator.psf_pars['refmed']
    refcov = model.dat_generator.psf_pars['refcov']
    refimm = model.dat_generator.psf_pars['refimm']
    lambda_fs = model.dat_generator.psf_pars['lambda']
    objstage0 = model.dat_generator.psf_pars['initial_obj_stage']
    zemit0 = -1 * refmed / refimm * objstage0
    pixel_size_xy = list(reversed(model.dat_generator.psf_pars['pixel_size_xy']))
    Npupil = 64
    zernike_aber = np.array([2, -2, 0, 2, 2, 0, 3, -1, 0, 3, 1, 0, 4, 0, 0, 3, -3, 0, 3, 3, 0,
                             4, -2, 0, 4, 2, 0, 5, -1, 0, 5, 1, 0, 6, 0, 0, 4, -4, 0, 4, 4, 0,
                             5, -3, 0, 5, 3, 0, 6, -2, 0, 6, 2, 0, 7, 1, 0, 7, -1, 0, 8, 0, 0],
                            dtype=np.float32).reshape([21, 3])
    zernike_aber[:, 2] = model.dat_generator.aber_map[test_pos[1], test_pos[0], 0:21] * model.dat_generator.psf_pars[
        'lambda']
    Npixels = model.dat_generator.psf_pars['psf_size']
    Nphotons = test_photons * np.ones(Nmol)
    bg = test_bg * np.ones(Nmol)
    xemit = 0 * np.ones(Nmol)
    yemit = 0 * np.ones(Nmol)
    zemit = 1 * np.linspace(-model.dat_generator.psf_pars['z_scale'], model.dat_generator.psf_pars['z_scale'], Nmol)
    objstage = 0 * np.linspace(-1000, 1000, Nmol)
    otf_rescale = [model.dat_generator.aber_map[test_pos[1], test_pos[0], 21],
                   model.dat_generator.aber_map[test_pos[1], test_pos[0], 22]]

    psf_pars = {'NA': NA, 'refmed': refmed, 'refcov': refcov, 'refimm': refimm, 'lambda': lambda_fs,
                'pixel_size_xy': pixel_size_xy, 'Npupil': Npupil, 'zernike_aber': zernike_aber, 'Npixels': Npixels,
                'Nphotons': Nphotons, 'bg': bg, 'objstage0': objstage0, 'zemit0': zemit0, 'objstage': objstage,
                'xemit': xemit, 'yemit': yemit, 'zemit': zemit, 'otf_rescale': otf_rescale}

    # instantiate the PSF model
    PSF_torch = FreeDipolePSF_torch(psf_pars, req_grad=False)
    data_torch = PSF_torch.gen_psf()

    # show PSFs
    print('look PSF model at this position', test_pos)
    plt.figure(constrained_layout=True)
    for j in range(Nmol):
        plt.subplot(int(np.sqrt(Nmol)), int(np.sqrt(Nmol)), j + 1)
        plt.imshow(cpu(data_torch)[j])
        plt.title(str(np.round(zemit[j])) + ' nm')
    plt.show()

    # calculate CRLB
    y_crlb, x_crlb, z_crlb, model_torch = PSF_torch.cal_crlb()
    x_crlb_np = x_crlb.detach().cpu().numpy()
    y_crlb_np = y_crlb.detach().cpu().numpy()
    z_crlb_np = z_crlb.detach().cpu().numpy()
    print('average 3D CRLB is:', np.sum(x_crlb_np ** 2 + y_crlb_np ** 2 + z_crlb_np ** 2) / PSF_torch.Nmol)
    plt.figure(constrained_layout=True)
    plt.plot(zemit, x_crlb_np, 'b', zemit, y_crlb_np,'g', zemit, z_crlb_np,'r')
    plt.legend(('$CRLB_x^{1/2}$', '$CRLB_y^{1/2}$', '$CRLB_z^{1/2}$'))
    plt.xlim([np.min(zemit), np.max(zemit)])
    plt.show()


def test_local_CRLB(model, test_pos, test_photons, test_bg, Nmol=25, use_train_cam=False, test_num=3000, show_res=True):
    """

    Compare the CRLB of the local PSF at given positions with the RMSE of the network prediction
    Parameters
    ----------
    model:
        FD-DeepLoc model
    test_pos:
        Position (x,y) where the PSF CRLB will be computed and checked
    test_photons:
        Number of signal photons used to compute CRLB
    test_bg:
        Number of background photons used to compute CRLB
    Nmol:
        Number of sampled z positions, which are uniformly distributed in the training z range
    use_train_cam:
        If true, the single-emitter test set will be simulated using the camera noise of training
        (e.g., qe, readout noise, analog-to-digital conversion factor). Otherwise only poisson noise will be added.
    test_num:
        Number of random PSFs at each sampled z position, the total number of test set is test_num*Nmol.
        The xy positions for each single emitter are random in the center two pixel, the z position is random in
        a step range (z_range/(Nmol-1)) around each z position.
    Returns
    -------

    """
    print('compare with PSF CRLB at this position', test_pos)

    # set PSF parameters
    NA = model.dat_generator.psf_pars['NA']
    refmed = model.dat_generator.psf_pars['refmed']
    refcov = model.dat_generator.psf_pars['refcov']
    refimm = model.dat_generator.psf_pars['refimm']
    lambda_fs = model.dat_generator.psf_pars['lambda']
    objstage0 = model.dat_generator.psf_pars['initial_obj_stage']
    zemit0 = -1 * refmed / refimm * objstage0
    pixel_size_xy = list(reversed(model.dat_generator.psf_pars['pixel_size_xy']))
    Npupil = 64
    zernike_aber = np.array([2, -2, 0, 2, 2, 0, 3, -1, 0, 3, 1, 0, 4, 0, 0, 3, -3, 0, 3, 3, 0,
                             4, -2, 0, 4, 2, 0, 5, -1, 0, 5, 1, 0, 6, 0, 0, 4, -4, 0, 4, 4, 0,
                             5, -3, 0, 5, 3, 0, 6, -2, 0, 6, 2, 0, 7, 1, 0, 7, -1, 0, 8, 0, 0],
                            dtype=np.float32).reshape([21, 3])
    zernike_aber[:, 2] = model.dat_generator.aber_map[test_pos[1], test_pos[0], 0:21] * model.dat_generator.psf_pars[
        'lambda']
    Npixels = model.dat_generator.psf_pars['psf_size']
    Nphotons = test_photons * np.ones(Nmol)
    bg = test_bg * np.ones(Nmol)
    xemit = 0 * np.ones(Nmol)
    yemit = 0 * np.ones(Nmol)
    zemit = 1 * np.linspace(-model.dat_generator.psf_pars['z_scale'], model.dat_generator.psf_pars['z_scale'], Nmol)
    objstage = 0 * np.linspace(-1000, 1000, Nmol)
    otf_rescale = [model.dat_generator.aber_map[test_pos[1], test_pos[0], 21],
                   model.dat_generator.aber_map[test_pos[1], test_pos[0], 22]]

    psf_pars = {'NA': NA, 'refmed': refmed, 'refcov': refcov, 'refimm': refimm, 'lambda': lambda_fs,
                'pixel_size_xy': pixel_size_xy, 'Npupil': Npupil, 'zernike_aber': zernike_aber, 'Npixels': Npixels,
                'Nphotons': Nphotons, 'bg': bg, 'objstage0': objstage0, 'zemit0': zemit0, 'objstage': objstage,
                'xemit': xemit, 'yemit': yemit, 'zemit': zemit, 'otf_rescale': otf_rescale}

    # instantiate the PSF model
    PSF_torch = FreeDipolePSF_torch(psf_pars, req_grad=False)
    # data_torch = PSF_torch.gen_psf()
    #
    # # show PSFs
    # print('look PSF model at this position')
    # plt.figure(constrained_layout=True)
    # for j in range(Nmol):
    #     plt.subplot(int(np.sqrt(Nmol)), int(np.sqrt(Nmol)), j + 1)
    #     plt.imshow(cpu(data_torch)[j])
    #     plt.title(str(np.round(zemit[j])) + ' nm')
    # plt.show()

    # calculate CRLB
    y_crlb, x_crlb, z_crlb, model_torch = PSF_torch.cal_crlb()
    x_crlb_np = x_crlb.detach().cpu().numpy()
    y_crlb_np = y_crlb.detach().cpu().numpy()
    z_crlb_np = z_crlb.detach().cpu().numpy()
    print('average 3D CRLB is:', np.sum(x_crlb_np ** 2 + y_crlb_np ** 2 + z_crlb_np ** 2) / PSF_torch.Nmol)
    # plt.figure(constrained_layout=True)
    # plt.plot(zemit, x_crlb_np, zemit, y_crlb_np, zemit, z_crlb_np)
    # plt.legend(('$CRLB_y^{1/2}$', '$CRLB_x^{1/2}$', '$CRLB_z^{1/2}$'))
    # plt.xlim([np.min(zemit), np.max(zemit)])
    # plt.show()


    # generate test single emitter set for CRLB comparison
    oldzemit = zemit
    dz = np.abs(oldzemit[2] - oldzemit[1])
    data_buffer = np.zeros([1, Npixels, Npixels])
    ground_truth = []
    frame_count = 1
    print('simulating single-emitter images for CRLB test')
    for i in range(test_num):
        PSF_torch.xemit = gpu((np.ones(Nmol) - 2 * np.random.rand(1)) * pixel_size_xy[0])  # nm
        PSF_torch.yemit = gpu((np.ones(Nmol) - 2 * np.random.rand(1)) * pixel_size_xy[1])  # nm
        PSF_torch.zemit = gpu(oldzemit + (np.ones(Nmol) - 2 * np.random.rand(1)) * (dz / 2 - 1))  # nm

        for j in range(Nmol):
            ground_truth.append([frame_count, frame_count,
                                 cpu(PSF_torch.yemit[j]) + Npixels / 2 * pixel_size_xy[1],
                                 cpu(PSF_torch.xemit[j]) + Npixels / 2 * pixel_size_xy[0],
                                 cpu(PSF_torch.zemit[j]) + 0, Nphotons[j]])
            frame_count += 1

        if use_train_cam:
            field_xy = [test_pos[0] - np.floor(Npixels / 2), test_pos[0] - np.floor(Npixels / 2) + Npixels - 1,
                        test_pos[1] - np.floor(Npixels / 2), test_pos[1] - np.floor(Npixels / 2) + Npixels - 1]
            data_model = PSF_torch.gen_psf() * PSF_torch.Nphotons
            data_tmp = cpu(model.dat_generator.sim_noise(data_model, field_xy))
        else:
            data_model = PSF_torch.gen_psf() * PSF_torch.Nphotons + PSF_torch.bg
            data_tmp = np.random.poisson(cpu(data_model))

        data_buffer = np.concatenate((data_buffer, data_tmp), axis=0)

        print('{}{}{}{}{}{}'.format('\r', 'simulated ', (i + 1) * Nmol, '/', test_num * Nmol, ' images'), end='')

    data_buffer = data_buffer[1:]
    print('\n')

    # show test data
    if show_res:
        print('example single-emitter data for CRLB test')
        plt.figure(constrained_layout=True)
        for j in range(Nmol):
            plt.subplot(int(np.sqrt(Nmol)), int(np.sqrt(Nmol)), j + 1)
            plt.imshow(data_buffer[j])
            plt.title(str(np.round(ground_truth[j][4])) + ' nm')
        plt.show()

    print('start inferring')
    # compare network's prediction with CRLB
    preds_raw, n_per_img, _ = recognition(model=model, eval_imgs_all=data_buffer,
                                          batch_size=model.evaluation_pars['batch_size'], use_tqdm=True,
                                          nms=True, candi_thre=0.3, nms_thre=0.7, rescale_xy=False,
                                          pix_nm=model.dat_generator.psf_pars['pixel_size_xy'],
                                          plot_num=None,
                                          start_field_pos=[test_pos[0] - np.floor(Npixels / 2),
                                                           test_pos[1] - np.floor(Npixels / 2)],
                                          stack_interval=200000, divide_and_conquer=False,
                                          win_size=model.dat_generator.simulation_pars['train_size'],
                                          padding=True)

    print('start computing evaluation metrics')
    match_dict, matches = assess(test_frame_nbr=data_buffer.shape[0], test_csv=ground_truth, pred_inp=preds_raw,
                                 tolerance=250,
                                 border=450, print_res=True, min_int=False, tolerance_ax=500, segmented=False)

    rmse_xyz = np.zeros([3, Nmol])
    for i in range(Nmol):
        z = zemit[i]
        ind = np.where(((z - dz / 2) < matches[:, 2]) & (matches[:, 2] < (z + dz / 2)))
        tmp = np.squeeze(matches[ind, :])
        if tmp.shape[0]:
            rmse_xyz[0, i] = np.sqrt(np.mean(np.square(tmp[:, 0] - tmp[:, 4])))
            rmse_xyz[1, i] = np.sqrt(np.mean(np.square(tmp[:, 1] - tmp[:, 5])))
            rmse_xyz[2, i] = np.sqrt(np.mean(np.square(tmp[:, 2] - tmp[:, 6])))

    if show_res:
        print('plot the RMSE of network prediction vs CRLB')
        plt.figure(constrained_layout=True)
        plt.plot(zemit, x_crlb_np, 'b', zemit, y_crlb_np, 'g', zemit, z_crlb_np, 'r')
        plt.scatter(zemit, rmse_xyz[0, :],c='b')
        plt.scatter(zemit, rmse_xyz[1, :],c='g')
        plt.scatter(zemit, rmse_xyz[2, :],c='r')
        plt.legend(('$CRLB_x^{1/2}$', '$CRLB_y^{1/2}$', '$CRLB_z^{1/2}$', '$RMSE_x$', '$RMSE_y$', '$RMSE_z$'), ncol=2,
                   loc='upper center')
        plt.xlim([np.min(zemit), np.max(zemit)])
        plt.show()

    return zemit, x_crlb_np, y_crlb_np, z_crlb_np, rmse_xyz
```
