## Supplementary Software for "Field dependent deep learning enables high-throughput whole-cell 3D super-resolution imaging": data_simulator.py

```
import matplotlib.pyplot as plt
import torch
# from PSF_vector_c import *
from PSF_vector_gpu import *
from local_utils import *


# class DataSimulator(PSF_VECTOR_C):
class DataSimulator(PSF_VECTOR_GPU):

    def __init__(self, psf_params, simulation_params):

        self.psf_pars = psf_params
        self.psf_size = psf_params['psf_size']
        self.aber_map = psf_params['aber_map']
        self.simulation_pars = simulation_params

    def transform_offsets(self, S, XYZI):

        n_samples = S.shape[0] // XYZI.shape[0]
        XYZI_rep = XYZI.repeat_interleave(n_samples, 0)

        s_inds = tuple(S.nonzero().transpose(1, 0))
        x_os_vals = (XYZI_rep[:, 0][s_inds])[:, None, None]
        y_os_vals = (XYZI_rep[:, 1][s_inds])[:, None, None]
        z_vals = self.psf_pars['z_scale'] * XYZI_rep[:, 2][s_inds][:, None, None]
        i_vals = (XYZI_rep[:, 3][s_inds])[:, None, None]

        return x_os_vals, y_os_vals, z_vals, i_vals

    def psf_vector(self, S, X_os, Y_os, Z, I, field_xy, robust_training):
        W = self.sim_psf_vector(cpu(S.nonzero()), cpu(X_os), cpu(Y_os), cpu(Z), self.aber_map,
                                field_xy, robust_training)

        W /= W.sum(-1).sum(-1)[:, None, None]  # photon normalization
        W *= I

        return W

    def look_aber_map(self):
        zernikeModes = np.array([2, -2, 0, 2, 2, 0, 3, -1, 0, 3, 1, 0, 4, 0, 0, 3, -3, 0, 3, 3, 0,
                                 4, -2, 0, 4, 2, 0, 5, -1, 0, 5, 1, 0, 6, 0, 0, 4, -4, 0, 4, 4, 0,
                                 5, -3, 0, 5, 3, 0, 6, -2, 0, 6, 2, 0, 7, 1, 0, 7, -1, 0, 8, 0, 0],
                                dtype=np.float32).reshape(21, 3).T
        plt.figure(constrained_layout=True)
        # plt.figure()
        for i in range(zernikeModes.shape[1]):
            ax_tmp = plt.subplot(5, 5, i + 1)
            img_tmp = plt.imshow(self.aber_map[:, :, i] * self.psf_pars['lambda'])
            plt.colorbar(mappable=img_tmp,ax=ax_tmp,fraction=0.046, pad=0.04)
            plt.title(str(zernikeModes[0:2, i]))
        ax_tmp=plt.subplot(5, 5, i + 2)
        img_tmp=plt.imshow(self.aber_map[:, :, 21])
        plt.colorbar(mappable=img_tmp,ax=ax_tmp,fraction=0.046, pad=0.04)
        plt.title('IsigmaX')
        ax_tmp=plt.subplot(5, 5, i + 3)
        img_tmp=plt.imshow(self.aber_map[:, :, 22])
        plt.colorbar(mappable=img_tmp,ax=ax_tmp,fraction=0.046, pad=0.04)
        plt.title('IsigmaY')
        # plt.tight_layout()
        plt.show()

    def look_psf(self, pos_xy, z_scale=1000):
        print('PSF at position xy in aberration map:', pos_xy,
              ',aber_map_size:', self.psf_pars['aber_map'].shape)

        # [x,y] position corresponds to [column, row]
        pos_rc = [pos_xy[1], pos_xy[0]]
        pos_rc = np.array(pos_rc).reshape(1, 2)

        locs_in_map = np.repeat(np.insert(pos_rc, 0, 0, axis=1), repeats=21, axis=0)
        x_offset = np.zeros([21, 1, 1])
        y_offset = np.zeros([21, 1, 1])
        z = np.linspace(-z_scale, z_scale, 21).reshape(21, 1, 1)

        field_xy = [0, self.aber_map.shape[1] - 1, 0, self.aber_map.shape[0] - 1]

        psf_samples = self.sim_psf_vector(locs_in_map, x_offset, y_offset, z, self.aber_map, field_xy)
        psf_samples /= psf_samples.sum(-1).sum(-1)[:, None, None]
        psf_samples = cpu(psf_samples)

        plt.figure(constrained_layout=True)
        for j in range(21):
            plt.subplot(3, 7, j + 1)
            plt.imshow(psf_samples[j])
            plt.title(str(z[j, 0, 0]) + ' nm')
        plt.show()

    def place_psfs(self, W, S):

        recs = torch.zeros_like(S)
        h, w = S.shape[1], S.shape[2]

        s_inds = tuple(S.nonzero().transpose(1, 0))
        relu = nn.ReLU()

        r_inds = S.nonzero()[:, 1:]
        uni_inds = S.sum(0).nonzero()

        x_rl = relu(uni_inds[:, 0] - self.psf_size // 2)
        y_rl = relu(uni_inds[:, 1] - self.psf_size // 2)

        x_wl = relu(self.psf_size // 2 - uni_inds[:, 0])
        x_wh = self.psf_size - (uni_inds[:, 0] + self.psf_size // 2 - h) - 1

        y_wl = relu(self.psf_size // 2 - uni_inds[:, 1])
        y_wh = self.psf_size - (uni_inds[:, 1] + self.psf_size // 2 - w) - 1

        r_inds_r = h * r_inds[:, 0] + r_inds[:, 1]
        uni_inds_r = h * uni_inds[:, 0] + uni_inds[:, 1]

        for i in range(len(uni_inds)):
            curr_inds = torch.nonzero(r_inds_r == uni_inds_r[i])[:, 0]
            w_cut = W[curr_inds, x_wl[i]: x_wh[i], y_wl[i]: y_wh[i]]

            recs[s_inds[0][curr_inds], x_rl[i]:x_rl[i] + w_cut.shape[1], y_rl[i]:y_rl[i] + w_cut.shape[2]] += w_cut

        return recs

    def genfunc(self, S, XYZI, field_xy, robust_training):
        X_os, Y_os, Z, I = self.transform_offsets(S, XYZI)
        psf_patches = self.psf_vector(S, X_os, Y_os, Z, I, field_xy, robust_training)
        return self.psf_pars['ph_scale'] * self.place_psfs(psf_patches, S)

    def datagen_func(self, S, X_os, Y_os, Z, I, field_xy, robust_training):
        batch_size, n_inp, h, w = S.shape[0], S.shape[1], S.shape[2], S.shape[3]
        xyzi = torch.cat([X_os.reshape([-1, 1, h, w]), Y_os.reshape([-1, 1, h, w]), Z.reshape([-1, 1, h, w]),
                          I.reshape([-1, 1, h, w])], 1)
        recs = self.genfunc(S.reshape([-1, h, w]), xyzi, field_xy, robust_training)
        torch.clamp_min_(recs, 0)
        imgs_sim = recs.reshape([batch_size, n_inp, h, w])

        return imgs_sim

    def sim_noise(self, imgs_sim, field_xy, add_noise=True):
        if self.simulation_pars['camera'] == 'EMCCD':
            bg_photons = (self.simulation_pars['backg'] - self.simulation_pars['baseline']) \
                         / self.simulation_pars['em_gain'] * self.simulation_pars['e_per_adu'] \
                         / self.simulation_pars['qe']
            if bg_photons<0:
                print('converted bg_photons is less than 0, please check the parameters setting!')

            if self.simulation_pars['perlin_noise']:
                size_x, size_y = imgs_sim.shape[-2], imgs_sim.shape[-1]
                self.PN_res = self.simulation_pars['pn_res']
                self.PN_octaves_num = 1
                space_range_x = size_x / self.PN_res
                space_range_y = size_y / self.PN_res
                self.PN = PerlinNoiseFactory(dimension=2, octaves=self.PN_octaves_num,
                                             tile=(space_range_x, space_range_y),
                                             unbias=True)
                PN_tmp_map = np.zeros([size_x, size_y])
                for x in range(size_x):
                    for y in range(size_y):
                        cal_PN_tmp = self.PN(x / self.PN_res, y / self.PN_res)
                        PN_tmp_map[x, y] = cal_PN_tmp
                PN_noise = PN_tmp_map * bg_photons * self.simulation_pars['pn_factor']
                bg_photons += PN_noise
                bg_photons = gpu(bg_photons)

            imgs_sim += bg_photons

            if add_noise:
                imgs_sim = torch.distributions.Poisson(
                    imgs_sim * self.simulation_pars['qe'] + self.simulation_pars['spurious_c']).sample()

                imgs_sim = torch.distributions.Gamma(imgs_sim, 1 / self.simulation_pars['em_gain']).sample()

                if type(self.simulation_pars['sig_read']) == np.ndarray:
                    RN = gpu(self.simulation_pars['sig_read'][field_xy[2]:field_xy[3] + 1, field_xy[0]:field_xy[1] + 1]+1e-6)
                else:
                    RN = self.simulation_pars['sig_read']
                zeros = torch.zeros_like(imgs_sim)
                read_out_noise = torch.distributions.Normal(zeros, zeros + RN).sample()

                imgs_sim = imgs_sim + read_out_noise
                imgs_sim = torch.clamp(imgs_sim / self.simulation_pars['e_per_adu'] + self.simulation_pars['baseline'], min=0)

        elif self.simulation_pars['camera'] == 'sCMOS':
            bg_photons = (self.simulation_pars['backg'] - self.simulation_pars['baseline']) \
                         * self.simulation_pars['e_per_adu'] / self.simulation_pars['qe']
            if bg_photons<0:
                print('converted bg_photons is less than 0, please check the parameters setting!')

            if self.simulation_pars['perlin_noise']:
                size_x, size_y = imgs_sim.shape[-2], imgs_sim.shape[-1]
                self.PN_res = self.simulation_pars['pn_res']
                self.PN_octaves_num = 1
                space_range_x = size_x / self.PN_res
                space_range_y = size_y / self.PN_res
                self.PN = PerlinNoiseFactory(dimension=2, octaves=self.PN_octaves_num,
                                             tile=(space_range_x, space_range_y),
                                             unbias=True)
                PN_tmp_map = np.zeros([size_x, size_y])
                for x in range(size_x):
                    for y in range(size_y):
                        cal_PN_tmp = self.PN(x / self.PN_res, y / self.PN_res)
                        PN_tmp_map[x, y] = cal_PN_tmp
                PN_noise = PN_tmp_map * bg_photons * self.simulation_pars['pn_factor']
                bg_photons += PN_noise
                bg_photons = gpu(bg_photons)

            imgs_sim += bg_photons

            if add_noise:
                imgs_sim = torch.distributions.Poisson(
                    imgs_sim * self.simulation_pars['qe'] + self.simulation_pars['spurious_c']).sample()

                if type(self.simulation_pars['sig_read']) == np.ndarray:
                    RN = gpu(self.simulation_pars['sig_read'][field_xy[2]:field_xy[3] + 1, field_xy[0]:field_xy[1] + 1]+1e-6)
                else:
                    RN = self.simulation_pars['sig_read']
                zeros = torch.zeros_like(imgs_sim)
                read_out_noise = torch.distributions.Normal(zeros, zeros + RN).sample()

                imgs_sim = imgs_sim + read_out_noise
                imgs_sim = torch.clamp(imgs_sim / self.simulation_pars['e_per_adu'] + self.simulation_pars['baseline'], min=0)
        else:
            print('wrong camera types! please choose EMCCD or sCMOS!')

        return imgs_sim

    def sampling(self, M, batch_size=1, local_context=False, iter_num=None, train_size=128):
        """ randomly generate the molecule localizations (discrete pixel+continuous offset), photons, apply a simple
            photo-physics model
        """

        blink_p = M
        blink_p = blink_p.reshape(1, 1, blink_p.shape[-2], blink_p.shape[-1]).repeat_interleave(batch_size, 0)

        # every pixel has a probability blink_p of existing a molecule, following binomial distribution
        locs1 = torch.distributions.Binomial(1, blink_p).sample().to('cuda')
        zeros = torch.zeros_like(locs1).to('cuda')
        # z position follows a uniform distribution with predefined range
        z = torch.distributions.Uniform(zeros + self.simulation_pars['z_prior'][0],
                                        zeros + self.simulation_pars['z_prior'][1]).sample().to('cuda')
        # xy offset follow uniform distribution
        x_os = torch.distributions.Uniform(zeros - 0.5, zeros + 0.5).sample().to('cuda')
        y_os = torch.distributions.Uniform(zeros - 0.5, zeros + 0.5).sample().to('cuda')

        if local_context:
            surv_p = self.simulation_pars['surv_p']
            a11 = 1 - (1 - blink_p) * (1 - surv_p)
            locs2 = torch.distributions.Binomial(1, (1 - locs1) * blink_p + locs1 * a11).sample().to('cuda')
            locs3 = torch.distributions.Binomial(1, (1 - locs2) * blink_p + locs2 * a11).sample().to('cuda')
            locs = torch.cat([locs1, locs2, locs3], 1)
            x_os = x_os.repeat_interleave(3, 1)
            y_os = y_os.repeat_interleave(3, 1)
            z = z.repeat_interleave(3, 1)
        else:
            locs = locs1
        #  photon number is sampled from a uniform distribution
        ints = torch.distributions.Uniform(torch.zeros_like(locs) + self.simulation_pars['min_ph'],
                                           torch.ones_like(locs)).sample().to('cuda')
        x_os *= locs
        y_os *= locs
        z *= locs
        ints *= locs
        # when generating the training data, width==height==train_size
        if M.shape[1] == M.shape[2] == train_size:
            index = iter_num % len(self.sliding_win)
            # global xy position in the whole aberration map. range is [0，aber_map.size-1]
            field_xy = torch.tensor(self.sliding_win[index])
        # when generating evaluation set, field_xy corresponds to the full size of the aber_map
        else:
            field_xy = torch.tensor([0, self.aber_map.shape[1] - 1, 0, self.aber_map.shape[0] - 1])

        return locs, x_os, y_os, z, ints, field_xy

    def sim_func(self, M, batch_size=1, local_context=False, photon_filt=False, photon_filt_thre=500,
                 P_locs_cse=False, iter_num=None, train_size=128, robust_training=False):
        """Main function for simulating SMLM images
        
        Parameters
        ----------
        M: 
            A map with pixel values indicating the probability of a molecule existing in that pixel
        batch_size: 
            Number of images simulated for each iteration
        local_context: 
            Generate 1(False) or 3 consecutive(True) frames
        photon_filt:
            Enforce the network to ignore too dim molecules
        photon_filt_thre:
            Threshold used to decide which molecules should be ignored
        P_locs_cse:
            If True, the loss function will add a cross-entropy term
        iter_num:
            Number of simulating iterations, used to determine the global position of the sub-area image
        train_size:
            Size of training data
        robust_training:
            If True, add small zernike disturbance to the simulated PSFs
        Returns
        -------

        """
        imgs_sim = torch.zeros([batch_size, 3, M.shape[1], M.shape[2]]).type(torch.cuda.FloatTensor) \
            if local_context else torch.zeros([batch_size, 1, M.shape[1], M.shape[2]]).type(torch.cuda.FloatTensor)

        xyzi_gt = torch.zeros([batch_size, 0, 4]).type(torch.cuda.FloatTensor)
        s_mask = torch.zeros([batch_size, 0]).type(torch.cuda.FloatTensor)
        pix_cor = torch.zeros([batch_size, 0, 2]).type(torch.cuda.FloatTensor)

        S, X_os, Y_os, Z, I, field_xy = self.sampling(batch_size=batch_size, M=M, local_context=local_context,
                                                      iter_num=iter_num, train_size=train_size)

        if S.sum():

            imgs_sim += self.datagen_func(S, X_os, Y_os, Z, I, field_xy, robust_training)
            xyzi = torch.cat([X_os[:, :, None], Y_os[:, :, None], Z[:, :, None], I[:, :, None]], 2)

            S = S[:, 1] if local_context else S[:, 0]

            if S.sum():
                # if simulate the local context, take the middle frame otherwise the first one
                xyzi = xyzi[:, 1] if local_context else xyzi[:, 0]
                # get all molecules' discrete pixel positions [number_in_batch, row, column]
                s_inds = tuple(S.nonzero().transpose(1, 0))
                # get these molecules' sub-pixel xy offsets, z positions and photons
                xyzi_true = xyzi[s_inds[0], :, s_inds[1], s_inds[2]]
                # get the xy continuous pixel positions
                xyzi_true[:, 0] += s_inds[2].type(torch.cuda.FloatTensor) + 0.5
                xyzi_true[:, 1] += s_inds[1].type(torch.cuda.FloatTensor) + 0.5
                # return the gt numbers of molecules on each training images of this batch
                # (if local_context, return the number of molecules on the middle frame)
                s_counts = torch.unique_consecutive(s_inds[0], return_counts=True)[1]
                s_max = s_counts.max()
                # for each training images of this batch, build a molecule list with length=s_max
                xyzi_gt_curr = torch.cuda.FloatTensor(batch_size, s_max, 4).fill_(0)
                s_mask_curr = torch.cuda.FloatTensor(batch_size, s_max).fill_(0)
                pix_cor_curr = torch.cuda.LongTensor(batch_size, s_max, 2).fill_(0)
                s_arr = torch.cat([torch.arange(c) for c in s_counts], dim=0)
                # put the gt in the molecule list, with remaining=0
                xyzi_gt_curr[s_inds[0], s_arr] = xyzi_true
                s_mask_curr[s_inds[0], s_arr] = 1
                pix_cor_curr[s_inds[0], s_arr, 0] = s_inds[1].clone().detach()
                pix_cor_curr[s_inds[0], s_arr, 1] = s_inds[2].clone().detach()

                xyzi_gt = torch.cat([xyzi_gt, xyzi_gt_curr], 1)
                s_mask = torch.cat([s_mask, s_mask_curr], 1)
                pix_cor = torch.cat([pix_cor, pix_cor_curr], 1)

        # add_noise, bg is actually the normalized un-noised PSF image
        psf_imgs_gt = imgs_sim.clone() / self.psf_pars['ph_scale'] * 10
        psf_imgs_gt = psf_imgs_gt[:, 1] if local_context else psf_imgs_gt[:, 0]
        imgs_sim = self.sim_noise(imgs_sim, field_xy)

        # only return the ground truth with photon>threshold,
        if photon_filt:
            for i in range(xyzi_gt.shape[0]):
                for j in range(xyzi_gt.shape[1]):
                    if xyzi_gt[i, j, 3] * self.psf_pars['ph_scale'] < photon_filt_thre:
                        xyzi_gt[i, j] = torch.tensor([0, 0, 0, 0])
                        s_mask[i, j] = 0
                        S[i, int(pix_cor[i, j][0]), int(pix_cor[i, j][1])] = 0

        locs = S if P_locs_cse else None

        return imgs_sim, xyzi_gt, s_mask, psf_imgs_gt, locs, field_xy  # return images and ground truth
```
