## Supplementary Software for "Field dependent deep learning enables high-throughput whole-cell 3D super-resolution imaging": model.py

```
import collections
import pickle
import time

from fd_deeploc_core.train_loss_rec import *
from fd_deeploc_core.network import *
from fd_deeploc_core.data_simulator import *
from fd_deeploc_core.anlz_eval import *


class DeepLocModel(TrainFuncs, LossFuncs, RecFuncs):

    def __init__(self, net_pars, psf_pars, simulation_pars, train_pars, evaluation_pars):
        """FD-DeepLoc Model
        
        Parameters
        ----------
        net_pars : dict
            Parameters for the network
        psf_pars : dict
            Parameters for the Point Spread Function
        simulation_pars : dict
            Parameters for the data simulator
        train_pars: dict
            Parameters for the training process and loss function
        """
        self.train_pars = train_pars
        self.evaluation_pars = evaluation_pars
        self.net_pars = net_pars

        self.local_context = net_pars['local_context']
        self.sig_pred = net_pars['sig_pred']
        self.psf_pred = net_pars['psf_pred']
        self.n_filters = net_pars['n_filters']

        self.n_inp = 3 if self.local_context else 1
        n_features = self.n_filters * self.n_inp
        self.frame_module = UnetCoordConv(n_inp=1, n_filters=self.n_filters, n_stages=2,
                                          use_coordconv=self.net_pars['use_coordconv']).to(torch.device('cuda'))
        self.context_module = UnetCoordConv(n_inp=n_features, n_filters=self.n_filters, n_stages=2,
                                            use_coordconv=self.net_pars['use_coordconv']).to(torch.device('cuda'))
        self.out_module = OutnetCoordConv(self.n_filters, self.sig_pred, self.psf_pred, pad=self.net_pars['padding'],
                                          ker_size=self.net_pars['kernel_size'],
                                          use_coordconv=self.net_pars['use_coordconv']).to(torch.device('cuda'))

        self.dat_generator = DataSimulator(psf_pars, simulation_pars)

        self.filename = None

        self.recorder = {}
        self._iter_count = 0

        # network structure
        self.net_weights = list(self.frame_module.parameters()) + list(self.context_module.parameters()) + list(self.out_module.parameters())
        self.optimizer_rec = torch.optim.AdamW(self.net_weights, lr=self.train_pars['lr'],
                                               weight_decay=self.train_pars['w_decay'])
        self.scheduler_rec = torch.optim.lr_scheduler.StepLR(self.optimizer_rec, step_size=1000,
                                                             gamma=self.train_pars['lr_decay'])

        # init the sliding windows' positions in the whole FOV, namely the field_xy
        self.init_sliding_win()

    def init_recorder(self):

        self.recorder['cost_hist'] = collections.OrderedDict([])
        self.recorder['update_time'] = collections.OrderedDict([])
        self.recorder['n_per_img'] = collections.OrderedDict([])
        self.recorder['recall'] = collections.OrderedDict([])
        self.recorder['precision'] = collections.OrderedDict([])
        self.recorder['jaccard'] = collections.OrderedDict([])
        self.recorder['rmse_lat'] = collections.OrderedDict([])
        self.recorder['rmse_ax'] = collections.OrderedDict([])
        self.recorder['rmse_vol'] = collections.OrderedDict([])
        self.recorder['jor'] = collections.OrderedDict([])
        self.recorder['eff_lat'] = collections.OrderedDict([])
        self.recorder['eff_ax'] = collections.OrderedDict([])
        self.recorder['eff_3d'] = collections.OrderedDict([])

    def init_eval_data(self):
        train_size_x = self.dat_generator.psf_pars['aber_map'].shape[1]
        train_size_y = self.dat_generator.psf_pars['aber_map'].shape[0]
        density = self.evaluation_pars['mols_per_img']

        # the probability of a molecule in central area is larger
        M = np.ones([1, train_size_y, train_size_x])
        M[0, int(0.05 * train_size_y):int(0.95 * train_size_y), int(0.05 * train_size_x):int(0.95 * train_size_x)] += 9
        M = M / M.sum() * density

        ground_truth = []
        eval_imgs = np.zeros([1, train_size_y, train_size_x])
        for j in range(self.evaluation_pars['eval_imgs_number']):
            imgs_sim, xyzi_mat, s_mask, psf_est, locs, field_xy = self.dat_generator.sim_func(
                M=gpu(M), batch_size=1,
                local_context=self.local_context,
                photon_filt=False,
                photon_filt_thre=0,
                P_locs_cse=False,
                iter_num=self._iter_count, train_size=self.dat_generator.simulation_pars['train_size'],
                robust_training=False)

            imgs_tmp = cpu(imgs_sim)[:, 1] if self.local_context else cpu(imgs_sim)[:, 0]
            eval_imgs = np.concatenate((eval_imgs, imgs_tmp), axis=0)

            for i in range(xyzi_mat.shape[1]):
                ground_truth.append(
                    [i + 1, j + 1, cpu(xyzi_mat[0, i, 0]) * self.dat_generator.psf_pars['pixel_size_xy'][0],
                     cpu(xyzi_mat[0, i, 1]) * self.dat_generator.psf_pars['pixel_size_xy'][1],
                     cpu(xyzi_mat[0, i, 2]) * self.dat_generator.psf_pars['z_scale'],
                     cpu(xyzi_mat[0, i, 3]) * self.dat_generator.psf_pars['ph_scale']])

            print('{}{}{}'.format('\rAlready simulated ', j+1, ' evaluation images'), end='')

        self.evaluation_pars['eval_imgs'] = eval_imgs[1:]
        self.evaluation_pars['ground_truth'] = ground_truth
        self.evaluation_pars['fov_size'] = [train_size_x * self.dat_generator.psf_pars['pixel_size_xy'][0],
                                            train_size_y * self.dat_generator.psf_pars['pixel_size_xy'][1]]

        print('\neval images shape:', self.evaluation_pars['eval_imgs'].shape, 'contain', len(ground_truth), 'molecules,',
              'field_xy:', field_xy)

        plt.figure(constrained_layout=True)
        ax_tmp = plt.subplot(1,1,1)
        img_tmp = plt.imshow(self.evaluation_pars['eval_imgs'][0])
        plt.colorbar(mappable=img_tmp,ax=ax_tmp, fraction=0.046, pad=0.04)
        plt.title('the first image of eval set,check the background')
        # plt.tight_layout()
        plt.show()

    def eval_func(self, candi_thre=0.3, nms_thre=0.7, print_res=False):
        if self.evaluation_pars['ground_truth'] is not None:
            preds_raw, n_per_img, _ = recognition(model=self, eval_imgs_all=self.evaluation_pars['eval_imgs'],
                                                  batch_size=self.evaluation_pars['batch_size'], use_tqdm=False,
                                                  nms=True, candi_thre=candi_thre, nms_thre=nms_thre,
                                                  rescale_xy=False, pix_nm=self.dat_generator.psf_pars['pixel_size_xy'],
                                                  plot_num=None, start_field_pos=[0, 0], stack_interval=20000,
                                                  divide_and_conquer=self.evaluation_pars['divide_and_conquer'],
                                                  win_size=self.dat_generator.simulation_pars['train_size'],
                                                  padding=True)
            match_dict, _ = assess(test_frame_nbr=self.evaluation_pars['eval_imgs_number'],
                                   test_csv=self.evaluation_pars['ground_truth'], pred_inp=preds_raw,
                                   size_xy=self.evaluation_pars['fov_size'], tolerance=250, border=450,
                                   print_res=print_res, min_int=False, tolerance_ax=500, segmented=False)

            for k in self.recorder.keys():
                if k in match_dict:
                    self.recorder[k][self._iter_count] = match_dict[k]

            self.recorder['n_per_img'][self._iter_count] = n_per_img

    def fit(self, batch_size=16, max_iters=50000, print_output=True, print_freq=100):
        """Train the FD-DeepLoc model
        
        Parameters
        ----------
        batch_size: int
            The amount of training data used per iteration
        max_iters: int
            Number of training iterations
        print_output: bool
            If True, the model performance on the evaluation dataset will be printed
        print_freq:
            Number of iterations between evaluations of the training progress
        """
        self.batch_size = batch_size
        self.train_size = self.dat_generator.simulation_pars['train_size']
        self.print_freq = print_freq

        last_iter = self._iter_count
        tot_t = 0
        best_record = -1e5
        iter_best = 0

        print('start training!')  # fushuang

        if self._iter_count > 1000:
            iter_best = self._iter_count
            print('train from checkpoint! the last print is:')
            if self.evaluation_pars['ground_truth'] is not None:
                best_record = self.recorder['eff_3d'][self._iter_count]
                # Jor = 100*jaccard/rmse_lat 越大越好,Factor是所有测试图像的全图概率p的和的平均
                print('{}{:0.3f}'.format('JoR: ', float(self.recorder['jor'][self._iter_count])), end='')
                # print('{}{}{:0.3f}'.format(' || ', 'Eff_lat: ', self.recorder['eff_lat'][self._iter_count]),end='')
                print('{}{}{:0.3f}'.format(' || ', 'Eff_3d: ', self.recorder['eff_3d'][self._iter_count]), end='')
                print('{}{}{:0.3f}'.format(' || ', 'Jaccard: ', self.recorder['jaccard'][self._iter_count]), end='')
                print('{}{}{:0.3f}'.format(' || ', 'Factor: ', self.recorder['n_per_img'][self._iter_count]), end='')
                print('{}{}{:0.3f}'.format(' || ', 'RMSE_lat: ', self.recorder['rmse_lat'][self._iter_count]), end='')
                print('{}{}{:0.3f}'.format(' || ', 'RMSE_ax: ', self.recorder['rmse_ax'][self._iter_count]), end='')
                print('{}{}{:0.3f}'.format(' || ', 'Cost: ', self.recorder['cost_hist'][self._iter_count]), end='')
                print('{}{}{:0.3f}'.format(' || ', 'Recall: ', self.recorder['recall'][self._iter_count]), end='')
                print('{}{}{:0.3f}'.format(' || ', 'Precision: ', self.recorder['precision'][self._iter_count]), end='')
                print('{}{}{}'.format(' || ', 'BatchNr.: ', self._iter_count))
            else:
                # print('{}{:0.3f}'.format('Factor: ', self.recorder['n_per_img'][self._iter_count]), end='')
                print('{}{}{:0.3f}'.format(' || ', 'Cost: ', self.recorder['cost_hist'][self._iter_count]), end='')
                print('{}{}{}'.format(' || ', 'BatchNr.: ', self._iter_count))

        # main loop for training
        while self._iter_count < max_iters:

            t0 = time.time()
            tot_cost = []

            # evaluate the performance and save model every print_freq iterations
            for _ in range(self.print_freq):
                loss = self.training(self.train_size, self.dat_generator.simulation_pars['density'])
                tot_cost.append(cpu(loss))
            tot_t += (time.time() - t0)
            self.recorder['cost_hist'][self._iter_count] = np.mean(tot_cost)
            updatetime = 1000 * tot_t / (self._iter_count - last_iter)
            last_iter = self._iter_count
            tot_t = 0
            self.recorder['update_time'][self._iter_count] = updatetime

            if print_output:
                if self._iter_count > 1000 and self.evaluation_pars['ground_truth'] is not None:
                    self.eval_func()
                    # Jor = 100*jaccard/rmse_lat. the larger, the better, Factor is the average sum of the probability
                    # channel
                    print('{}{:0.3f}'.format('JoR: ', float(self.recorder['jor'][self._iter_count])), end='')
                    # print('{}{}{:0.3f}'.format(' || ', 'Eff_lat: ', self.recorder['eff_lat'][self._iter_count]),end='')
                    print('{}{}{:0.3f}'.format(' || ', 'Eff_3d: ', self.recorder['eff_3d'][self._iter_count]), end='')
                    print('{}{}{:0.3f}'.format(' || ', 'Jaccard: ', self.recorder['jaccard'][self._iter_count]), end='')
                    print('{}{}{:0.3f}'.format(' || ', 'Factor: ', self.recorder['n_per_img'][self._iter_count]),end='')
                    print('{}{}{:0.3f}'.format(' || ', 'RMSE_lat: ', self.recorder['rmse_lat'][self._iter_count]),end='')
                    print('{}{}{:0.3f}'.format(' || ', 'RMSE_ax: ', self.recorder['rmse_ax'][self._iter_count]), end='')
                    print('{}{}{:0.3f}'.format(' || ', 'Cost: ', self.recorder['cost_hist'][self._iter_count]), end='')
                    print('{}{}{:0.3f}'.format(' || ', 'Recall: ', self.recorder['recall'][self._iter_count]), end='')
                    print('{}{}{:0.3f}'.format(' || ', 'Precision: ', self.recorder['precision'][self._iter_count]),end='')
                    print('{}{}{}'.format(' || ', 'BatchNr.: ', self._iter_count), end='')
                    print('{}{}{:0.1f}{}'.format(' || ', 'Time Upd.: ', float(updatetime), ' ms '))
                else:
                    # print('{}{:0.3f}'.format('Factor: ', self.recorder['n_per_img'][self._iter_count]), end='')
                    print('{}{}{:0.3f}'.format(' || ', 'Cost: ', self.recorder['cost_hist'][self._iter_count]), end='')
                    print('{}{}{:0.1f}{}'.format(' || ', 'Time Upd.: ', float(updatetime), ' ms '), end='')
                    print('{}{}{}'.format(' || ', 'BatchNr.: ', self._iter_count))

            #  save the model
            if self.filename:
                if self._iter_count > 1000 and self.evaluation_pars['ground_truth'] is not None:
                    best_record = self.recorder['eff_3d'][self._iter_count]
                    rmse_lat_best = self.recorder['rmse_lat'][self._iter_count]
                    rmse_ax_best = self.recorder['rmse_ax'][self._iter_count]
                    iter_best = self._iter_count
                    print('{}{:0.3f}{}{:0.3f}{}{:0.3f}{}{}'.format(
                        'saving this model, eff_3d, rmse_lat, rmse_ax and BatchNr are : ',
                        best_record, ' || ', rmse_lat_best, ' || ', rmse_ax_best, ' || ', iter_best))
                    print('\n')
                    with open(self.filename + '.pkl', 'wb') as f:
                        pickle.dump(self, f)
                else:
                    with open(self.filename + '.pkl', 'wb') as f:
                        pickle.dump(self, f)
        print('training finished!')

    def init_sliding_win(self):
        """ Init the sliding windows (sub-area training data) on the big aberration map (FOV), ensure PSF at everywhere
        is properly learned
        """
        vacuum_size = int(np.ceil(
            self.dat_generator.simulation_pars['train_size'] * self.dat_generator.simulation_pars['margin_empty']))
        over_lap = 2 * vacuum_size
        # traverse the aberration map, the area field_xy denotes should have the same size as train_size, ensure the
        # field_xy is in the range of aberration map, when field_xy exceeds the boundary, index win_size backward from
        # the boundary
        row_num = int(np.ceil(
            (self.dat_generator.aber_map.shape[0]-over_lap) / (self.dat_generator.simulation_pars['train_size'] - over_lap)))
        column_num = int(np.ceil(
            (self.dat_generator.aber_map.shape[1]-over_lap) / (self.dat_generator.simulation_pars['train_size'] - over_lap)))

        sliding_win = []
        for iter_num in range(0, row_num * column_num):
            x_field = iter_num % column_num * (self.dat_generator.simulation_pars['train_size'] - over_lap) \
                if iter_num % column_num * (self.dat_generator.simulation_pars['train_size'] - over_lap) + \
                   self.dat_generator.simulation_pars['train_size'] <= self.dat_generator.aber_map.shape[1] \
                else self.dat_generator.aber_map.shape[1] - self.dat_generator.simulation_pars['train_size']

            y_field = iter_num // column_num % row_num * (self.dat_generator.simulation_pars['train_size'] - over_lap) \
                if iter_num // column_num % row_num * (self.dat_generator.simulation_pars['train_size'] - over_lap) + \
                   self.dat_generator.simulation_pars['train_size'] <= self.dat_generator.aber_map.shape[0] \
                else self.dat_generator.aber_map.shape[0] - self.dat_generator.simulation_pars['train_size']

            # it represents the [x, y] position, not [row, column], numerical range is [0, aber_map.shape-1]
            sliding_win.append([x_field, x_field + self.dat_generator.simulation_pars['train_size'] - 1,
                                y_field, y_field + self.dat_generator.simulation_pars['train_size'] - 1])

        self.dat_generator.sliding_win = sliding_win
        print('training sliding windows on aber_map: ')
        for i in range(0, len(sliding_win)):
            print('area_num:', i, 'field_xy:', sliding_win[i])
```
