## Supplementary Software for "Field dependent deep learning enables high-throughput whole-cell 3D super-resolution imaging": network.py

```
import torch
import torch.nn as nn
import torch.nn.functional as F
import math
from torch.optim.optimizer import Optimizer


def add_coord(input, field_xy, aber_map_size):
    """ concatenate global coordinate channels to the input data

    Parameters
    ----------
    input:
         tensors with shape [batchsize, channel, height, width]
    field_xy:
        [xstart, xend, ystart, yend], the global position of the input sub-area image in the big aberration map.
        should satisfies this relationship: xstart - xend + 1 = input.width
    aber_map_size:
        [sizex sizey], the size of the aberration map, sizex corresponds to column, sizey corresponds to row.
    """
    x_start = field_xy[0].float()
    y_start = field_xy[2].float()
    x_end = field_xy[1].float()
    y_end = field_xy[3].float()

    batch_size = input.size()[0]
    x_dim = input.size()[3]
    y_dim = input.size()[2]

    x_step = 1 / (aber_map_size[0] - 1)
    y_step = 1 / (aber_map_size[1] - 1)

    xx_range = torch.arange(x_start / (aber_map_size[0] - 1), x_end / (aber_map_size[0] - 1) + 1e-6, step=x_step,
                            dtype=torch.float32).repeat([y_dim, 1]).reshape([1, y_dim, x_dim])

    xx_range = xx_range.repeat_interleave(repeats=batch_size, dim=0).reshape([batch_size, 1, y_dim, x_dim])

    yy_range = torch.arange(y_start / (aber_map_size[1] - 1), y_end / (aber_map_size[1] - 1) + 1e-6, step=y_step,
                            dtype=torch.float32).repeat([x_dim, 1]).transpose(1, 0).reshape([1, y_dim, x_dim])

    yy_range = yy_range.repeat_interleave(repeats=batch_size, dim=0).reshape([batch_size, 1, y_dim, x_dim])

    xx_range = xx_range.cuda()
    yy_range = yy_range.cuda()

    ret = torch.cat([input, xx_range, yy_range], dim=1)

    return ret


class CoordConv(nn.Module):
    """ CoordConv class, add coordinate channels to the data,
    apply extra 2D convolution on the coordinate channels and add the result"""
    def __init__(self, in_channels, out_channels, kernel_size, padding):
        super(CoordConv, self).__init__()
        self.conv2d_im = nn.Conv2d(in_channels=in_channels, out_channels=out_channels, kernel_size=kernel_size,
                                   padding=padding)
        self.conv2d_coord = nn.Conv2d(in_channels=2, out_channels=out_channels, kernel_size=1, padding=0)

    def forward(self, input, field_xy, aber_map_size):
        y = add_coord(input, field_xy, aber_map_size)
        ret_1 = self.conv2d_im(y[:, 0:-2])
        ret_2 = self.conv2d_coord(y[:, -2:])
        return ret_1 + ret_2


class OutnetCoordConv(nn.Module):
    """output module"""
    def __init__(self, n_filters, pred_sig=False, pred_bg=False, pad=1, ker_size=3, use_coordconv=True):
        super(OutnetCoordConv, self).__init__()

        self.pred_bg = pred_bg
        self.pred_sig = pred_sig
        self.use_coordconv = use_coordconv

        if self.use_coordconv:
            self.p_out1 = CoordConv(in_channels=n_filters, out_channels=n_filters, kernel_size=ker_size,
                                    padding=pad).cuda()
            self.p_out2 = nn.Conv2d(in_channels=n_filters, out_channels=1, kernel_size=1, padding=0).cuda()  # fu
            self.xyzi_out1 = CoordConv(in_channels=n_filters, out_channels=n_filters, kernel_size=ker_size,
                                       padding=pad).cuda()
            self.xyzi_out2 = nn.Conv2d(in_channels=n_filters, out_channels=4, kernel_size=1, padding=0).cuda()  # fu

            nn.init.kaiming_normal_(self.p_out1.conv2d_im.weight, mode='fan_in', nonlinearity='relu')
            nn.init.kaiming_normal_(self.p_out1.conv2d_coord.weight, mode='fan_in', nonlinearity='relu')
            nn.init.kaiming_normal_(self.p_out2.weight, mode='fan_in', nonlinearity='sigmoid')
            nn.init.constant_(self.p_out2.bias, -6.)  # -6

            nn.init.kaiming_normal_(self.xyzi_out1.conv2d_im.weight, mode='fan_in', nonlinearity='relu')
            nn.init.kaiming_normal_(self.xyzi_out1.conv2d_coord.weight, mode='fan_in', nonlinearity='relu')
            nn.init.kaiming_normal_(self.xyzi_out2.weight, mode='fan_in', nonlinearity='tanh')
            nn.init.zeros_(self.xyzi_out2.bias)

            if self.pred_sig:
                self.xyzis_out1 = CoordConv(in_channels=n_filters, out_channels=n_filters, kernel_size=ker_size,
                                            padding=pad).cuda()
                self.xyzis_out2 = nn.Conv2d(in_channels=n_filters, out_channels=4, kernel_size=1, padding=0).cuda()

                nn.init.kaiming_normal_(self.xyzis_out1.conv2d_im.weight, mode='fan_in', nonlinearity='relu')
                nn.init.kaiming_normal_(self.xyzis_out1.conv2d_coord.weight, mode='fan_in', nonlinearity='relu')
                nn.init.kaiming_normal_(self.xyzis_out2.weight, mode='fan_in', nonlinearity='sigmoid')
                nn.init.zeros_(self.xyzis_out2.bias)

            if self.pred_bg:
                self.bg_out1 = CoordConv(in_channels=n_filters, out_channels=n_filters, kernel_size=ker_size,
                                         padding=pad).cuda()
                self.bg_out2 = nn.Conv2d(in_channels=n_filters, out_channels=1, kernel_size=1, padding=0).cuda()

                nn.init.kaiming_normal_(self.bg_out1.conv2d_im.weight, mode='fan_in', nonlinearity='relu')
                nn.init.kaiming_normal_(self.bg_out1.conv2d_coord.weight, mode='fan_in', nonlinearity='relu')
                nn.init.kaiming_normal_(self.bg_out2.weight, mode='fan_in', nonlinearity='sigmoid')
                nn.init.zeros_(self.bg_out2.bias)
        else:
            self.p_out1 = nn.Conv2d(in_channels=n_filters, out_channels=n_filters, kernel_size=ker_size,
                                    padding=pad).cuda()
            self.p_out2 = nn.Conv2d(in_channels=n_filters, out_channels=1, kernel_size=1, padding=0).cuda()  # fu
            self.xyzi_out1 = nn.Conv2d(in_channels=n_filters, out_channels=n_filters, kernel_size=ker_size,
                                       padding=pad).cuda()
            self.xyzi_out2 = nn.Conv2d(in_channels=n_filters, out_channels=4, kernel_size=1, padding=0).cuda()  # fu

            nn.init.kaiming_normal_(self.p_out1.weight, mode='fan_in', nonlinearity='relu')
            nn.init.kaiming_normal_(self.p_out2.weight, mode='fan_in', nonlinearity='sigmoid')
            nn.init.constant_(self.p_out2.bias, -6.)  # -6

            nn.init.kaiming_normal_(self.xyzi_out1.weight, mode='fan_in', nonlinearity='relu')
            nn.init.kaiming_normal_(self.xyzi_out2.weight, mode='fan_in', nonlinearity='tanh')
            nn.init.zeros_(self.xyzi_out2.bias)

            if self.pred_sig:
                self.xyzis_out1 = nn.Conv2d(in_channels=n_filters, out_channels=n_filters, kernel_size=ker_size,
                                            padding=pad).cuda()
                self.xyzis_out2 = nn.Conv2d(in_channels=n_filters, out_channels=4, kernel_size=1, padding=0).cuda()

                nn.init.kaiming_normal_(self.xyzis_out1.weight, mode='fan_in', nonlinearity='relu')
                nn.init.kaiming_normal_(self.xyzis_out2.weight, mode='fan_in', nonlinearity='sigmoid')
                nn.init.zeros_(self.xyzis_out2.bias)

            if self.pred_bg:
                self.bg_out1 = nn.Conv2d(in_channels=n_filters, out_channels=n_filters, kernel_size=ker_size,
                                         padding=pad).cuda()
                self.bg_out2 = nn.Conv2d(in_channels=n_filters, out_channels=1, kernel_size=1, padding=0).cuda()

                nn.init.kaiming_normal_(self.bg_out1.weight, mode='fan_in', nonlinearity='relu')
                nn.init.kaiming_normal_(self.bg_out2.weight, mode='fan_in', nonlinearity='sigmoid')
                nn.init.zeros_(self.bg_out2.bias)

    def forward(self, x, field_xy, aber_map_size):

        outputs = {}

        if self.use_coordconv:
            p = F.elu(self.p_out1(x, field_xy, aber_map_size))
            outputs['p'] = self.p_out2(p)

            xyzi = F.elu(self.xyzi_out1(x, field_xy, aber_map_size))
            outputs['xyzi'] = self.xyzi_out2(xyzi)

            if self.pred_sig:
                xyzis = F.elu(self.xyzis_out1(x, field_xy, aber_map_size))
                outputs['xyzi_sig'] = self.xyzis_out2(xyzis)

            if self.pred_bg:
                bg = F.elu(self.bg_out1(x, field_xy, aber_map_size))
                outputs['bg'] = self.bg_out2(bg)
        else:
            p = F.elu(self.p_out1(x))
            outputs['p'] = self.p_out2(p)

            xyzi = F.elu(self.xyzi_out1(x))
            outputs['xyzi'] = self.xyzi_out2(xyzi)

            if self.pred_sig:
                xyzis = F.elu(self.xyzis_out1(x))
                outputs['xyzi_sig'] = self.xyzis_out2(xyzis)

            if self.pred_bg:
                bg = F.elu(self.bg_out1(x))
                outputs['bg'] = self.bg_out2(bg)

        return outputs


class UnetCoordConv(nn.Module):
    """used for frame analysis module and temporal context module"""
    def __init__(self, n_inp, n_filters=64, n_stages=5, pad=1, ker_size=3, use_coordconv=True):
        super(UnetCoordConv, self).__init__()
        curr_N = n_filters
        self.n_stages = n_stages
        self.layer_path = nn.ModuleList()
        self.use_coordconv = use_coordconv

        if self.use_coordconv:
            self.layer_path.append(
                CoordConv(in_channels=n_inp, out_channels=curr_N, kernel_size=ker_size, padding=pad).cuda())
        else:
            self.layer_path.append(
                nn.Conv2d(in_channels=n_inp, out_channels=curr_N, kernel_size=ker_size, padding=pad).cuda())

        self.layer_path.append(
            nn.Conv2d(in_channels=curr_N, out_channels=curr_N, kernel_size=ker_size, padding=pad).cuda())

        for i in range(n_stages):
            self.layer_path.append(
                nn.Conv2d(in_channels=curr_N, out_channels=curr_N, kernel_size=2, stride=2, padding=0).cuda())
            self.layer_path.append(
                nn.Conv2d(in_channels=curr_N, out_channels=curr_N * 2, kernel_size=ker_size, padding=pad).cuda())
            curr_N *= 2
            self.layer_path.append(
                nn.Conv2d(in_channels=curr_N, out_channels=curr_N, kernel_size=ker_size, padding=pad).cuda())

        for i in range(n_stages):
            self.layer_path.append(nn.UpsamplingNearest2d(scale_factor=2).cuda())
            self.layer_path.append(
                nn.Conv2d(in_channels=curr_N, out_channels=curr_N // 2, kernel_size=ker_size, padding=pad).cuda())

            curr_N = curr_N // 2

            self.layer_path.append(
                nn.Conv2d(in_channels=curr_N * 2, out_channels=curr_N, kernel_size=ker_size, padding=pad).cuda())
            self.layer_path.append(
                nn.Conv2d(in_channels=curr_N, out_channels=curr_N, kernel_size=ker_size, padding=pad).cuda())

        for m in self.layer_path:
            if isinstance(m, CoordConv):
                nn.init.kaiming_normal_(m.conv2d_im.weight, mode='fan_in', nonlinearity='relu')  # 初始化卷积层权重
                nn.init.kaiming_normal_(m.conv2d_coord.weight, mode='fan_in', nonlinearity='relu')
            if isinstance(m, nn.Conv2d):
                nn.init.kaiming_normal_(m.weight, mode='fan_in', nonlinearity='relu')

    def forward(self, x, xy_field, aber_map_size):

        n_l = 0
        x_bridged = []

        if self.use_coordconv:
            x = F.elu(list(self.layer_path)[n_l](x, xy_field, aber_map_size))
        else:
            x = F.elu(list(self.layer_path)[n_l](x))
        n_l += 1
        x = F.elu(list(self.layer_path)[n_l](x))
        n_l += 1

        x_bridged.append(x)

        for i in range(self.n_stages):
            for n in range(3):
                if isinstance(list(self.layer_path)[n_l], CoordConv):
                    x = F.elu(list(self.layer_path)[n_l](x, xy_field, aber_map_size))
                else:
                    x = F.elu(list(self.layer_path)[n_l](x))
                n_l += 1
                if n == 2 and i < self.n_stages - 1:
                    x_bridged.append(x)

        for i in range(self.n_stages):
            for n in range(4):
                if isinstance(list(self.layer_path)[n_l], CoordConv):
                    x = F.elu(list(self.layer_path)[n_l](x, xy_field, aber_map_size))
                else:
                    x = F.elu(list(self.layer_path)[n_l](x))
                n_l += 1
                if n == 1:
                    x = torch.cat([x, x_bridged.pop()], 1)  # concatenate

        return x
```
