## Supplementary Software for "Field dependent deep learning enables high-throughput whole-cell 3D super-resolution imaging": train_loss_rec.py

```
import torch.nn as nn
import numpy as np
import torch
from local_utils import *


class TrainFuncs:

    def training(self, train_size, density):
        """
        main function for training process

        Parameters
        ----------
        train_size:
            size of training image
        density:
            number of molecules per image
        Returns
        -------
        """
        map = np.zeros([1, train_size, train_size])
        map[0, int(self.dat_generator.simulation_pars['margin_empty'] * train_size):
               int((1 - self.dat_generator.simulation_pars['margin_empty']) * train_size),
        int(self.dat_generator.simulation_pars['margin_empty'] * train_size):
        int((1 - self.dat_generator.simulation_pars['margin_empty']) * train_size)] += 1
        map = map / map.sum() * density

        imgs_sim, xyzi_gt, s_mask, psf_imgs_gt, locs, field_xy = self.dat_generator.sim_func(
            M=gpu(map), batch_size=self.batch_size,
            local_context=self.local_context,
            photon_filt=self.train_pars['ph_filt'],
            photon_filt_thre=self.train_pars['ph_filt_thre'],
            P_locs_cse=self.train_pars['P_locs_cse'],
            iter_num=self._iter_count, train_size=train_size,
            robust_training=self.dat_generator.simulation_pars['robust_training'])

        P, xyzi_est, xyzi_sig, psf_imgs_est = self.inferring(imgs_sim, field_xy,
                                                             aber_map_size=[self.dat_generator.aber_map.shape[1],
                                                                            self.dat_generator.aber_map.shape[0]])

        # loss
        loss_total = self.final_loss(P, xyzi_est, xyzi_sig, xyzi_gt, s_mask, psf_imgs_est, psf_imgs_gt, locs)

        self.optimizer_rec.zero_grad()
        loss_total.backward()

        # avoid too large gradient
        torch.nn.utils.clip_grad_norm_(self.net_weights, max_norm=self.train_pars['clip_g_n'], norm_type=2)

        # update the network and the optimizer state
        self.optimizer_rec.step()
        self.scheduler_rec.step()

        self._iter_count += 1

        return loss_total.detach()

    def look_trainingdata(self, area_num=0):
        train_size = self.dat_generator.simulation_pars['train_size']
        density = self.dat_generator.simulation_pars['density']

        map = np.zeros([1, train_size, train_size])
        map[0, int(self.dat_generator.simulation_pars['margin_empty'] * train_size):
               int((1 - self.dat_generator.simulation_pars['margin_empty']) * train_size),
        int(self.dat_generator.simulation_pars['margin_empty'] * train_size):
        int((1 - self.dat_generator.simulation_pars['margin_empty']) * train_size)] += 1
        map = map / map.sum() * density

        imgs_sim, xyzi_gt, s_mask, bg, locs, field_xy = self.dat_generator.sim_func(
            M=gpu(map), batch_size=1,
            local_context=self.local_context,
            photon_filt=False,
            photon_filt_thre=0,
            P_locs_cse=True,
            iter_num=area_num, train_size=train_size,
            robust_training=self.dat_generator.simulation_pars['robust_training'])

        bg_photons = (self.dat_generator.simulation_pars['backg'] - self.dat_generator.simulation_pars['baseline']) \
                     / self.dat_generator.simulation_pars['em_gain'] * self.dat_generator.simulation_pars['e_per_adu'] \
                     / self.dat_generator.simulation_pars['qe']

        mol_photons = (self.dat_generator.simulation_pars['min_ph'] + 1) / 2 * self.dat_generator.psf_pars['ph_scale']

        print('{}{}{}{}{}'.format('The average signal/background used for training are: ', int(mol_photons), '/',
                                  int(bg_photons), ' photons'))

        print('{}{}{}{}{}{}{}'.format('look training data sample, area number: ', area_num, '(0-',
                                      len(self.dat_generator.sliding_win) - 1, '), ', 'field_xy: ', field_xy))

        if self.local_context:
            plt.figure(constrained_layout=True)
            ax_tmp = plt.subplot(2, 2, 1)
            img_tmp = plt.imshow(cpu(imgs_sim)[0, 0])
            plt.colorbar(mappable=img_tmp,ax=ax_tmp,fraction=0.046, pad=0.04)
            plt.title('last frame')

            ax_tmp = plt.subplot(2, 2, 2)
            img_tmp = plt.imshow(cpu(imgs_sim)[0, 1])
            plt.colorbar(mappable=img_tmp,ax=ax_tmp,fraction=0.046, pad=0.04)
            plt.title('middle frame')

            ax_tmp = plt.subplot(2, 2, 3)
            img_tmp = plt.imshow(cpu(imgs_sim)[0, 2])
            plt.colorbar(mappable=img_tmp,ax=ax_tmp,fraction=0.046, pad=0.04)
            plt.title('next frame')

            ax_tmp = plt.subplot(2, 2, 4)
            img_tmp = plt.imshow(cpu(locs)[0])
            plt.colorbar(mappable=img_tmp,ax=ax_tmp,fraction=0.046, pad=0.04)
            plt.title('ground truth \nof middle frame')
            # plt.tight_layout()
            plt.show()
        else:
            plt.figure(constrained_layout=True)
            ax_tmp = plt.subplot(1, 2, 1)
            img_tmp = plt.imshow(cpu(imgs_sim)[0, 0])
            plt.colorbar(mappable=img_tmp,ax=ax_tmp,fraction=0.046, pad=0.04)
            plt.title('frame')

            ax_tmp = plt.subplot(1, 2, 2)
            img_tmp = plt.imshow(cpu(locs)[0])
            plt.colorbar(mappable=img_tmp,ax=ax_tmp,fraction=0.046, pad=0.04)
            plt.title('ground truth of the frame')
            # plt.tight_layout()
            plt.show()


class LossFuncs:

    def eval_bg_sq_loss(self, psf_imgs_est, psf_imgs_gt):
        loss = nn.MSELoss(reduction='none')
        cost = loss(psf_imgs_est, psf_imgs_gt)
        cost = cost.sum(-1).sum(-1)
        return cost

    def eval_P_locs_loss(self, P, locs):
        loss_cse = -(locs * torch.log(P) + (1 - locs) * torch.log(1 - P))
        loss_cse = loss_cse.sum(-1).sum(-1)
        return loss_cse

    def count_loss_analytical(self, P, s_mask):
        log_prob = 0
        prob_mean = P.sum(-1).sum(-1)
        prob_var = (P - P ** 2).sum(-1).sum(-1)
        X = s_mask.sum(-1)
        log_prob += 1 / 2 * ((X - prob_mean) ** 2) / prob_var + 1 / 2 * torch.log(2 * np.pi * prob_var)
        return log_prob

    def loc_loss_analytical(self, P, xyzi_est, xyzi_sig, xyzi_gt, s_mask):
        # each pixel is a component of Gaussian Mixture Model, with weights prob_normed
        prob_normed = P / (P.sum(-1).sum(-1)[:, None, None])

        p_inds = tuple((P + 1).nonzero().transpose(1, 0))

        xyzi_mu = xyzi_est[p_inds[0], :, p_inds[1], p_inds[2]]
        xyzi_mu[:, 0] += p_inds[2].type(torch.cuda.FloatTensor) + 0.5
        xyzi_mu[:, 1] += p_inds[1].type(torch.cuda.FloatTensor) + 0.5

        xyzi_mu = xyzi_mu.reshape(self.batch_size, 1, -1, 4)
        xyzi_sig = xyzi_sig[p_inds[0], :, p_inds[1], p_inds[2]].reshape(self.batch_size, 1, -1, 4)  # >=0.01
        # xyzi_lnsig2 = xyzi_sig[p_inds[0], :, p_inds[1], p_inds[2]].reshape(self.batch_size, 1, -1, 4)  # >=0.01
        XYZI = xyzi_gt.reshape(self.batch_size, -1, 1, 4).repeat_interleave(self.train_size * self.train_size, 2)

        numerator = -1 / 2 * ((XYZI - xyzi_mu) ** 2)
        denominator = (xyzi_sig ** 2)  # >0
        # denominator = torch.exp(xyzi_lnsig2)
        log_p_gauss_4d = (numerator / denominator).sum(3) - 1 / 2 * (torch.log(2 * np.pi * denominator[:, :, :, 0]) +
                                                                     torch.log(2 * np.pi * denominator[:, :, :, 1]) +
                                                                     torch.log(2 * np.pi * denominator[:, :, :, 2]) +
                                                                     torch.log(2 * np.pi * denominator[:, :, :, 3]))

        gauss_coef = prob_normed.reshape(self.batch_size, 1, self.train_size * self.train_size)
        gauss_coef_logits = torch.log(gauss_coef)
        gauss_coef_logmax = torch.log_softmax(gauss_coef_logits, dim=2)

        gmm_log = torch.logsumexp(log_p_gauss_4d + gauss_coef_logmax, dim=2)
        # c = torch.sum(p_gauss_4d * gauss_coef,-1)
        # gmm_log = (torch.log(c) * s_mask).sum(-1)
        return (gmm_log * s_mask).sum(-1)

    def final_loss(self, P, xyzi_est, xyzi_sig, xyzi_gt, s_mask, psf_imgs_est, psf_imgs_gt, locs):
        count_loss = torch.mean(self.count_loss_analytical(P, s_mask) * s_mask.sum(-1))
        loc_loss = -torch.mean(self.loc_loss_analytical(P, xyzi_est, xyzi_sig, xyzi_gt, s_mask))
        bg_loss = torch.mean(self.eval_bg_sq_loss(psf_imgs_est, psf_imgs_gt)) if psf_imgs_est is not None else 0
        P_locs_error = torch.mean(self.eval_P_locs_loss(P, locs)) if locs is not None else 0

        loss_total = count_loss + loc_loss + bg_loss + P_locs_error

        return loss_total


class RecFuncs:

    def inferring(self, X, field_xy, aber_map_size):
        """
        main function for inferring process

        Parameters
        ----------
        X:
            input images
        field_xy:
            the global position of images in the aberration map
        aber_map_size:
            size of aberration map, used to calculate the normalized global position
        Returns
        -------

        """
        img_h, img_w = X.shape[-2], X.shape[-1]

        # simple normalization
        scaled_x = (X - self.net_pars['offset']) / self.net_pars['factor']

        if X.ndimension() == 3:  # when test, X.ndimension = 3
            scaled_x = scaled_x[:, None]
            fm_out = self.frame_module(scaled_x, field_xy, aber_map_size)
            if self.local_context:
                zeros = torch.zeros_like(fm_out[:1])
                h_t0 = fm_out
                h_tm1 = torch.cat([zeros, fm_out], 0)[:-1]
                h_tp1 = torch.cat([fm_out, zeros], 0)[1:]
                fm_out = torch.cat([h_tm1, h_t0, h_tp1], 1)
        else:  # when train, X.ndimension = 4
            fm_out = self.frame_module(scaled_x.reshape([-1, 1, img_h, img_w]), field_xy, aber_map_size) \
                .reshape(-1, self.n_filters * self.n_inp, img_h, img_w)

        cm_in = fm_out

        cm_out = self.context_module(cm_in, field_xy, aber_map_size)
        outputs = self.out_module.forward(cm_out, field_xy, aber_map_size)

        if self.sig_pred:
            xyzi_sig = torch.sigmoid(outputs['xyzi_sig']) + 0.001
        else:
            xyzi_sig = 0.2 * torch.ones_like(outputs['xyzi'])

        probs = torch.sigmoid(torch.clamp(outputs['p'], -16., 16.))

        xyzi_est = outputs['xyzi']
        xyzi_est[:, :2] = torch.tanh(xyzi_est[:, :2])  # xy
        xyzi_est[:, 2] = torch.tanh(xyzi_est[:, 2])  # z
        xyzi_est[:, 3] = torch.sigmoid(xyzi_est[:, 3])  # ph
        psf_est = torch.sigmoid(outputs['bg'])[:, 0] if self.psf_pred else None

        return probs[:, 0], xyzi_est, xyzi_sig, psf_est
```
