## Supplementary Software for "Field dependent deep learning enables high-throughput whole-cell 3D super-resolution imaging": fd_deeploc_env.yaml

```
name: fd_deeploc
channels:
  - pytorch
  - conda-forge
  - defaults
dependencies:
  - python=3.8.13
  - seaborn
  - jupyter
  - opencv
  - imagecodecs
  - tqdm
  - scipy=1.9.1
  - pytorch=1.10.0
  - torchvision=0.11.0
  - torchaudio=0.10.0
  - cudatoolkit=11.3
```
