## Supplementary Software for "Field dependent deep learning enables high-throughput whole-cell 3D super-resolution imaging": __init__.py

```
from local_utils.local_tifffile import *
from local_utils.csv_utils import *
from local_utils.utils import *
from local_utils.perlin_noise import *
from local_utils.plot_funcs import *
from local_utils.freedipolepsf_torch import *
```
