## Supplementary Software for "Field dependent deep learning enables high-throughput whole-cell 3D super-resolution imaging": csv_utils.py

```
from numpy import cross, eye, dot
import numpy as np
import csv
# from theano import config
import pandas as pd


def read_csv(path, flip_z=False, z_fac=1, pix_scale=[1, 1], drift_txt=None):
    """Reads a csv_file with leading columns: 'localization', 'frame', 'x', 'y', 'z'
    
    Parameters
    ----------
    flip_z : bool
        If True flips the z variable
    z_fac: float
        Multiplies z variable with the given factor to correct for eventual aberrations
    pix_scale: list of two ints
        Multiplies x and y locations with the given factors
    drift_txt : str
        Reads a drift corredtion txt file and applies drift correction for x and y locations
        
    Returns
    -------
    preds :list
        List of localizations with x,y,z locations given in nano meter
    """
    preds = pd.read_csv(path, header=None, skiprows=[0]).values

    if drift_txt is not None:
        drift_data = pd.read_csv(drift_txt, sep='	', header=None, skiprows=0).values

        for p in preds:
            p[2] = float(p[2]) - 100 * drift_data[np.clip(int(p[1]) - 1, 0, len(drift_data) - 1), 1]
            p[3] = float(p[3]) - 100 * drift_data[np.clip(int(p[1]) - 1, 0, len(drift_data) - 1), 2]

    preds[:, 2] = preds[:, 2] * pix_scale[0] + 0
    preds[:, 3] = preds[:, 3] * pix_scale[1] + 0
    preds[:, 4] = preds[:, 4] * z_fac + 0
    if flip_z:
        preds[:, 4] = -preds[:, 4] + 0

    return preds


def write_csv(pred_list, name, write_gt=False, append=False):
    """Writes a csv_file with columns: 'localizatioh', 'frame', 'x', 'y', 'z', 'intensity','x_sig','y_sig','z_sig'
    
    Parameters
    ----------
    pred_list : list
        List of localizations
    name: str
        File name
    """
    if write_gt:
        with open(name, 'w', newline='') as csvfile:
            csvwriter = csv.writer(csvfile, delimiter=',', quoting=csv.QUOTE_MINIMAL)
            csvwriter.writerow(['x_gt', 'y_gt', 'z_gt', 'intensity_gt','x_pred','y_pred','z_pred',
                                'intensity_pred','nms_p', 'x_sig', 'y_sig', 'z_sig'])
            for p in pred_list:
                csvwriter.writerow([repr(f) for f in p])
    else:
        if not append:
            with open(name, 'w', newline='') as csvfile:
                csvwriter = csv.writer(csvfile, delimiter=',', quoting=csv.QUOTE_MINIMAL)
                csvwriter.writerow(['Ground-truth', 'frame', 'xnano', 'ynano', 'znano', 'intensity','nms_p', 'x_sig',
                                    'y_sig', 'z_sig','I_sig'])
                for p in pred_list:
                    csvwriter.writerow([repr(f) for f in p])
        else:
            with open(name, 'a', newline='') as csvfile:
                csvwriter = csv.writer(csvfile, delimiter=',', quoting=csv.QUOTE_MINIMAL)
                for p in pred_list:
                    csvwriter.writerow([repr(f) for f in p])


def crop_preds(preds, sl=None, lims=None, z_lim=[-np.inf, np.inf], pix_nm=[100, 100]):
    """Crops a list of predictions to the given x,y,z limits
    
    Parameters
    ----------
    sl : numpy index_exp
        Specifies a slice of the recording.
    pix_nm: list of floats
        Specifies the size of a pixel in nano meters to translate a slice expression into nano meter
    lims: list of four floats
        Lower x, upper x, lower y, upper y limits in nano meter
    z_lim : list of two floats
        Lower z, upper z limits in nano meter
        
    Returns
    -------
    preds :list
        List of localizations with localizations outside the specified limits filteres out.
        x and y positions are shifted to start at zero
    """
    i_l = 0
    i_h = np.array(preds)[:, 1].max()

    if sl[0].start:
        i_l = sl[0].start
    if sl[0].stop:
        i_h = sl[0].stop

    if sl:
        y_l, y_h = pix_nm[1] * sl[1].start, pix_nm[1] * sl[1].stop
        x_l, x_h = pix_nm[0] * sl[2].start, pix_nm[0] * sl[2].stop

    if lims:
        x_l, x_h, y_l, y_h = lims[0], lims[1], lims[2], lims[3]

    preds = np.array(preds)
    inds = np.argwhere(
        (preds[:, 1] > i_l) & (preds[:, 1] < i_h) & (preds[:, 3] > y_l) & (preds[:, 3] < y_h) & (preds[:, 2] > x_l) & (
                    preds[:, 2] < x_h) & (preds[:, 4] > z_lim[0]) & (preds[:, 4] < z_lim[1]))[:, 0]
    preds = preds[inds]
    preds[:, 2] -= x_l
    preds[:, 3] -= y_l

    return list(preds)
```
