## Supplementary Software for "Field dependent deep learning enables high-throughput whole-cell 3D super-resolution imaging": freedipolepsf_torch.py

```
import time

import torch
import numpy as np
import matplotlib.pyplot as plt
import torch.nn as nn
from copy import deepcopy
from local_utils import *


class FreeDipolePSF_torch:
    def __init__(self, psf_pars, req_grad=False):
        self.psf_pars = deepcopy(psf_pars)
        self.NA = torch.tensor(self.psf_pars['NA'], device='cuda', dtype=torch.float64)
        self.refmed = torch.tensor(self.psf_pars['refmed'], device='cuda', dtype=torch.float64, requires_grad=req_grad)
        self.refcov = torch.tensor(self.psf_pars['refcov'], device='cuda', dtype=torch.float64, requires_grad=req_grad)
        self.refimm = torch.tensor(self.psf_pars['refimm'], device='cuda', dtype=torch.float64, requires_grad=req_grad)
        self.wavelength = torch.tensor(self.psf_pars['lambda'], device='cuda', dtype=torch.float64)
        self.objstage0 = torch.tensor(self.psf_pars['objstage0'], device='cuda', dtype=torch.float64, requires_grad=req_grad)
        self.zemit0 = torch.tensor(self.psf_pars['zemit0'], device='cuda', dtype=torch.float64, requires_grad=req_grad)
        self.pixel_size_x = torch.tensor(self.psf_pars['pixel_size_xy'][0], device='cuda', dtype=torch.float64)
        self.pixel_size_y = torch.tensor(self.psf_pars['pixel_size_xy'][1], device='cuda', dtype=torch.float64)
        self.Npupil = self.psf_pars['Npupil']
        self.zernike_aber = self.psf_pars['zernike_aber']
        self.orders = torch.tensor(self.zernike_aber[:, 0:2], device='cuda', dtype=torch.float64)
        self.zernikecoefs = torch.tensor(self.zernike_aber[:, 2], device='cuda', dtype=torch.float64,
                                         requires_grad=req_grad)
        self.Npixels = self.psf_pars['Npixels']
        self.Nphotons = torch.tensor(np.expand_dims(self.psf_pars['Nphotons'], axis=[1,2])
                                     , device='cuda', dtype=torch.float64, requires_grad=req_grad)
        self.bg = torch.tensor(np.expand_dims(self.psf_pars['bg'],axis=[1,2])
                               , device='cuda', dtype=torch.float64, requires_grad=req_grad)
        self.xemit = torch.tensor(self.psf_pars['xemit'], device='cuda', dtype=torch.float64, requires_grad=req_grad)
        self.yemit = torch.tensor(self.psf_pars['yemit'], device='cuda', dtype=torch.float64, requires_grad=req_grad)
        self.zemit = torch.tensor(self.psf_pars['zemit'], device='cuda', dtype=torch.float64, requires_grad=req_grad)
        self.objstage = torch.tensor(self.psf_pars['objstage'], device='cuda', dtype=torch.float64, requires_grad=req_grad)
        self.otf_rescale = psf_pars['otf_rescale']

    @staticmethod
    def get_zernikefunc(orders, xpupil, ypupil):
        xpupil = torch.real(xpupil)
        ypupil = torch.real(ypupil)
        zersize = orders.shape
        Nzer = zersize[0]
        radormax = int(max(orders[:, 0]))
        azormax = int(max(abs(orders[:, 1])))
        [Nx, Ny] = xpupil.shape

        # zerpol = np.zeros( [radormax+1,azormax+1,Nx,Ny] )
        zerpol = torch.zeros([21, 6, Nx, Ny], device='cuda')
        rhosq = xpupil ** 2 + ypupil ** 2
        rho = torch.sqrt(rhosq)
        zerpol[0, 0, :, :] = torch.ones_like(xpupil)

        for jm in range(1, azormax + 2 + 1):
            m = jm - 1
            if m > 0:
                zerpol[jm - 1, jm - 1, :, :] = rho * torch.squeeze(zerpol[jm - 1 - 1, jm - 1 - 1, :, :])

            zerpol[jm + 2 - 1, jm - 1, :, :] = ((m + 2) * rhosq - m - 1) * torch.squeeze(zerpol[jm - 1, jm - 1, :, :])
            for p in range(2, radormax - m + 2 + 1):
                n = m + 2 * p
                jn = n + 1
                zerpol[jn - 1, jm - 1, :, :] = (2 * (n - 1) * (n * (n - 2) * (2 * rhosq - 1) - m ** 2) * torch.squeeze(zerpol[jn - 2 - 1, jm - 1, :, :]) -
                                                n * (n + m - 2) * (n - m - 2) * torch.squeeze(zerpol[jn - 4 - 1, jm - 1, :, :])) / ((n - 2) * (n + m) * (n - m))

        phi = torch.atan2(ypupil, xpupil)
        allzernikes = torch.zeros([Nzer, Nx, Ny], device='cuda')
        for j in range(1, Nzer + 1):
            n = int(orders[j - 1, 0])
            m = int(orders[j - 1, 1])
            if m >= 0:
                allzernikes[j - 1, :, :] = torch.squeeze(zerpol[n + 1 - 1, m + 1 - 1, :, :]) * torch.cos(m * phi)
            else:
                allzernikes[j - 1, :, :] = torch.squeeze(zerpol[n + 1 - 1, -m + 1 - 1, :, :]) * torch.sin(-m * phi)

        # plt.figure(constrained_layout=True)
        # for i in range(21):
        #     plt.subplot(3, 7, i + 1)
        #     plt.imshow(cpu(allzernikes[i]))
        # plt.show()

        return allzernikes

    def prechirpz(self, xsize, qsize, N, M):
        L = N + M - 1
        sigma = 2 * np.pi * xsize * qsize / N / M
        Afac = torch.exp(2 * 1j * sigma * (1 - M))
        Bfac = torch.exp(2 * 1j * sigma * (1 - N))
        sqW = torch.exp(2 * 1j * sigma)
        W = sqW ** 2

        # fixed phase factor and amplitude factor
        Gfac = (2 * xsize / N) * torch.exp(1j * sigma * (1 - N) * (1 - M))

        # integration about n
        Utmp = torch.zeros([1, N], dtype=torch.complex64, device='cuda')
        A = torch.zeros([1, N], dtype=torch.complex64, device='cuda')
        Utmp[0, 0] = sqW * Afac
        A[0, 0] = 1.0
        for i in range(1, N):
            A[0, i] = Utmp[0, i - 1] * A[0, i - 1]
            Utmp[0, i] = Utmp[0, i - 1] * W

        #  the factor before the summation
        Utmp = torch.zeros([1, M], dtype=torch.complex64, device='cuda')
        B = torch.ones([1, M], dtype=torch.complex64, device='cuda')
        Utmp[0, 0] = sqW * Bfac
        B[0, 0] = Gfac
        for i in range(1, M):
            B[0, i] = Utmp[0, i - 1] * B[0, i - 1]
            Utmp[0, i] = Utmp[0, i - 1] * W

        # for circular convolution
        Utmp = torch.zeros([1, max(N, M) + 1], dtype=torch.complex64, device='cuda')
        Vtmp = torch.zeros([1, max(N, M) + 1], dtype=torch.complex64, device='cuda')
        Utmp[0, 0] = sqW
        Vtmp[0, 0] = 1.0
        # Utmp_cp = Utmp.clone()
        # Vtmp_cp = Vtmp.clone()
        for i in range(1, max(N, M) + 1):
            Vtmp[0, i] = Utmp[0, i - 1] * Vtmp[0, i - 1]
            Utmp[0, i] = Utmp[0, i - 1] * W
            # Vtmp[0, i] = Utmp_cp[0, i - 1] * Vtmp_cp[0, i - 1]
            # Utmp[0, i] = Utmp_cp[0, i - 1] * W
            # Vtmp_cp[0, i] = Vtmp[0, i].clone()
            # Utmp_cp[0, i] = Utmp[0, i].clone()

        D = torch.ones([1, L], dtype=torch.complex64, device='cuda')
        for i in range(0, M):
            D[0, i] = torch.conj(Vtmp[0, i])
        for i in range(0, N):
            D[0, L - 1 - i] = torch.conj(Vtmp[0, i + 1])

        D = torch.fft.fft(D, axis=1)

        return A, B, D

    def czt(self, datain, A, B, D):
        N = A.shape[1]
        M = B.shape[1]
        L = D.shape[1]
        K = datain.shape[0]

        # torch.repeat_interleave is too slow
        # t0 = time.time()
        # Amt = torch.repeat_interleave(A, K, 0)
        # Bmt = torch.repeat_interleave(B, K, 0)
        # Dmt = torch.repeat_interleave(D, K, 0)
        # print('torch czt: ', time.time() - t0)
        Amt = A.expand(K, N)
        Bmt = B.expand(K, M)
        Dmt = D.expand(K, L)

        cztin = torch.zeros([K, L], dtype=torch.complex64, device='cuda')
        cztin[:, 0:N] = Amt * datain
        tmp = Dmt * torch.fft.fft(cztin)
        cztout = torch.fft.ifft(tmp)

        dataout = Bmt * cztout[:, 0:M]

        return dataout

    def gen_psf(self):

        Nmol = self.xemit.shape[0]

        # pupil radius (in diffraction units) and pupil coordinate sampling
        pupil_size = torch.tensor(1.0, device='cuda', dtype=torch.float64)
        dxypupil = 2 * pupil_size / self.Npupil
        xypupil = torch.arange(-pupil_size + dxypupil / 2, pupil_size, dxypupil, device='cuda', dtype=torch.float64)
        [xpupil, ypupil] = torch.meshgrid(xypupil, xypupil)
        ypupil = torch.complex(ypupil, torch.zeros_like(ypupil))
        xpupil = torch.complex(xpupil, torch.zeros_like(xpupil))

        # calculation of relevant Fresnel-coefficients for the interfaces
        costhetamed = torch.sqrt(1.0 - (xpupil ** 2 + ypupil ** 2) * (self.NA ** 2) / (self.refmed ** 2))
        costhetacov = torch.sqrt(1.0 - (xpupil ** 2 + ypupil ** 2) * (self.NA ** 2) / (self.refcov ** 2))
        costhetaimm = torch.sqrt(1.0 - (xpupil ** 2 + ypupil ** 2) * (self.NA ** 2) / (self.refimm ** 2))
        fresnelpmedcov = 2 * self.refmed * costhetamed / (self.refmed * costhetacov + self.refcov * costhetamed)
        fresnelsmedcov = 2 * self.refmed * costhetamed / (self.refmed * costhetamed + self.refcov * costhetacov)
        fresnelpcovimm = 2 * self.refcov * costhetacov / (self.refcov * costhetaimm + self.refimm * costhetacov)
        fresnelscovimm = 2 * self.refcov * costhetacov / (self.refcov * costhetacov + self.refimm * costhetaimm)
        fresnelp = fresnelpmedcov * fresnelpcovimm
        fresnels = fresnelsmedcov * fresnelscovimm

        # apodization
        apod = 1 / torch.sqrt(costhetaimm)
        # define aperture
        aperturemask = torch.where((xpupil ** 2 + ypupil ** 2).real < 1.0, 1.0, 0.0)
        amplitude = aperturemask * apod

        # setting of vectorial functions
        phi = torch.atan2(torch.real(ypupil), torch.real(xpupil))
        cosphi = torch.cos(phi)
        sinphi = torch.sin(phi)
        costheta = costhetamed
        sintheta = torch.sqrt(1 - costheta ** 2)

        pvec = torch.empty([3,self.Npupil,self.Npupil], dtype=torch.complex64, device='cuda')
        pvec[0] = fresnelp * costheta * cosphi
        pvec[1] = fresnelp * costheta * sinphi
        pvec[2] = -fresnelp * sintheta
        svec = torch.empty([3,self.Npupil,self.Npupil], dtype=torch.complex64, device='cuda')
        svec[0] = -fresnels * sinphi
        svec[1] = fresnels * cosphi
        svec[2] = 0 * cosphi

        polarizationvector = torch.empty([2, 3,self.Npupil,self.Npupil], dtype=torch.complex64, device='cuda')
        for ipol in range(3):
            polarizationvector[0, ipol] = cosphi * pvec[ipol] - sinphi * svec[ipol]
            polarizationvector[1, ipol] = sinphi * pvec[ipol] + cosphi * svec[ipol]

        wavevector = torch.empty([2,self.Npupil,self.Npupil], dtype=torch.complex64,device='cuda')
        wavevector[0] = 2 * np.pi * self.NA / self.wavelength * xpupil
        wavevector[1] = 2 * np.pi * self.NA / self.wavelength * ypupil
        wavevectorzimm = 2 * np.pi * self.refimm / self.wavelength * costhetaimm
        wavevectorzmed = 2 * np.pi * self.refmed / self.wavelength * costhetamed

        # calculate aberration function
        waberration = torch.zeros_like(xpupil,dtype=torch.complex64,device='cuda')
        normfac = torch.sqrt(2 * (self.orders[:, 0] + 1) / (1 + torch.where(self.orders[:, 1] == 0,1.0,0.0)))
        zernikecoefs_norm = self.zernikecoefs * normfac
        allzernikes = self.get_zernikefunc(self.orders, xpupil, ypupil)

        for izer in range(self.orders.shape[0]):
            waberration += zernikecoefs_norm[izer] * allzernikes[izer]
        waberration *= aperturemask
        phasefactor = torch.exp(1j * 2 * np.pi * waberration / self.wavelength)

        pupilmatrix = torch.empty([2, 3,self.Npupil,self.Npupil], dtype=torch.complex64, device='cuda')
        for imat in range(2):
            for jmat in range(3):
                pupilmatrix[imat, jmat] = amplitude * phasefactor * polarizationvector[imat, jmat]

        # czt transform(fft the pupil)
        xrange = self.pixel_size_x * self.Npixels / 2
        yrange = self.pixel_size_y * self.Npixels / 2
        imagesizex = xrange * self.NA / self.wavelength
        imagesizey = yrange * self.NA / self.wavelength

        # calculate the auxiliary vectors for chirpz
        ax, bx, dx = self.prechirpz(pupil_size, imagesizex, self.Npupil, self.Npixels)
        ay, by, dy = self.prechirpz(pupil_size, imagesizey, self.Npupil, self.Npixels)

        FieldMatrix = torch.empty([2, 3, Nmol,self.Npixels,self.Npixels], dtype=torch.complex64, device='cuda')
        for jz in range(Nmol):
            # xyz induced phase
            if self.zemit[jz] + self.zemit0 >= 0:
                Wxyz = (-1 * self.xemit[jz]) * wavevector[0] + (-1 * self.yemit[jz]) * wavevector[1] + (
                            self.zemit[jz] + self.zemit0) * wavevectorzmed
                PositionPhase = torch.exp(1j * (Wxyz + (self.objstage[jz] + self.objstage0) * wavevectorzimm))
            else:
                # print("warning! the emitter's position may not have physical meaning")
                Wxyz = (-1 * self.xemit[jz]) * wavevector[0] + (-1 * self.yemit[jz]) * wavevector[1]
                PositionPhase = torch.exp(1j * (Wxyz + (self.objstage[jz] + self.objstage0 + self.zemit[jz] + self.zemit0) * wavevectorzimm))
            # t0 = time.time()
            for itel in range(2):
                for jtel in range(3):
                    Pupilfunction = PositionPhase * pupilmatrix[itel, jtel]
                    inter_image = torch.transpose(self.czt(Pupilfunction, ay, by, dy),1,0)
                    FieldMatrix[itel, jtel, jz] = torch.transpose(self.czt(inter_image, ax, bx, dx),1,0)
            # print('torch z loop: ', time.time() - t0)

        PSFs = torch.zeros([Nmol, self.Npixels, self.Npixels],device='cuda',dtype=torch.float64)
        for jz in range(Nmol):
            for jtel in range(3):
                for itel in range(2):
                    PSFs[jz, :, :] += 1 / 3 * (torch.abs(FieldMatrix[itel, jtel, jz])) ** 2

        # calculate intensity normalization function using the PSFs at focus
        FieldMatrix_norm = torch.empty([2, 3, self.Npixels, self.Npixels], dtype=torch.complex64,device='cuda')
        for itel in range(2):
            for jtel in range(3):
                Pupilfunction_norm = amplitude * polarizationvector[itel, jtel]
                inter_image_norm = torch.transpose(self.czt(Pupilfunction_norm, ay, by, dy),1,0)
                FieldMatrix_norm[itel, jtel] = torch.transpose(self.czt(inter_image_norm, ax, bx, dx),1,0)

        intFocus = torch.zeros([self.Npixels, self.Npixels], dtype=torch.float64,device='cuda')
        for jtel in range(3):
            for itel in range(2):
                intFocus += 1 / 3 * (torch.abs(FieldMatrix_norm[itel, jtel])) ** 2
        normIntensity = torch.sum(intFocus)
        PSFs /= normIntensity

        # otf rescale
        if len(self.otf_rescale):
            I_sigmax = self.otf_rescale[0]
            I_sigmay = self.otf_rescale[1]
            if I_sigmax == 0 or I_sigmay == 0:
                print('sigma for otf rescale is 0')
            else:
                h = torch.cuda.DoubleTensor(otf_gauss2D(shape=[5, 5], Isigmax=I_sigmax, Isigmay=I_sigmay)) \
                                            .reshape([1, 1, 5, 5])
                PSFs_tmp = PSFs.view(Nmol, 1, self.Npixels, self.Npixels)
                tmp = nn.functional.conv2d(PSFs_tmp, h, padding=2, stride=1)
                PSFs = tmp.view(Nmol, self.Npixels, self.Npixels)

        # PSFs = PSFs * self.Nphotons + self.bg

        # plt.figure(constrained_layout=True)
        # for i in range(Nmol):
        #     plt.subplot(5,5,i+1)
        #     plt.imshow(cpu(PSFs[i]))
        # plt.show()

        return PSFs

    def cal_crlb(self):
        self.Nmol = self.xemit.shape[0]

        # pupil radius (in diffraction units) and pupil coordinate sampling
        pupil_size = torch.tensor(1.0, device='cuda', dtype=torch.float64)
        dxypupil = 2 * pupil_size / self.Npupil
        xypupil = torch.arange(-pupil_size + dxypupil / 2, pupil_size, dxypupil, device='cuda', dtype=torch.float64)
        [xpupil, ypupil] = torch.meshgrid(xypupil, xypupil)
        ypupil = torch.complex(ypupil,torch.zeros_like(ypupil))
        xpupil = torch.complex(xpupil, torch.zeros_like(xpupil))

        # calculation of relevant Fresnel-coefficients for the interfaces
        costhetamed = torch.sqrt(1.0 - (xpupil ** 2 + ypupil ** 2) * (self.NA ** 2) / (self.refmed ** 2))
        costhetacov = torch.sqrt(1.0 - (xpupil ** 2 + ypupil ** 2) * (self.NA ** 2) / (self.refcov ** 2))
        costhetaimm = torch.sqrt(1.0 - (xpupil ** 2 + ypupil ** 2) * (self.NA ** 2) / (self.refimm ** 2))
        fresnelpmedcov = 2 * self.refmed * costhetamed / (self.refmed * costhetacov + self.refcov * costhetamed)
        fresnelsmedcov = 2 * self.refmed * costhetamed / (self.refmed * costhetamed + self.refcov * costhetacov)
        fresnelpcovimm = 2 * self.refcov * costhetacov / (self.refcov * costhetaimm + self.refimm * costhetacov)
        fresnelscovimm = 2 * self.refcov * costhetacov / (self.refcov * costhetacov + self.refimm * costhetaimm)
        fresnelp = fresnelpmedcov * fresnelpcovimm
        fresnels = fresnelsmedcov * fresnelscovimm

        # apodization
        apod = 1 / torch.sqrt(costhetaimm)
        # define aperture
        aperturemask = torch.where((xpupil ** 2 + ypupil ** 2).real < 1.0, 1.0, 0.0)
        self.amplitude = aperturemask * apod

        # setting of vectorial functions
        phi = torch.atan2(torch.real(ypupil), torch.real(xpupil))
        cosphi = torch.cos(phi)
        sinphi = torch.sin(phi)
        costheta = costhetamed
        sintheta = torch.sqrt(1 - costheta ** 2)

        pvec = torch.empty([3,self.Npupil,self.Npupil], dtype=torch.complex64, device='cuda')
        pvec[0] = fresnelp * costheta * cosphi
        pvec[1] = fresnelp * costheta * sinphi
        pvec[2] = -fresnelp * sintheta
        svec = torch.empty([3,self.Npupil,self.Npupil], dtype=torch.complex64, device='cuda')
        svec[0] = -fresnels * sinphi
        svec[1] = fresnels * cosphi
        svec[2] = 0 * cosphi

        self.polarizationvector = torch.empty([2, 3,self.Npupil,self.Npupil], dtype=torch.complex64, device='cuda')
        for ipol in range(3):
            self.polarizationvector[0, ipol] = cosphi * pvec[ipol] - sinphi * svec[ipol]
            self.polarizationvector[1, ipol] = sinphi * pvec[ipol] + cosphi * svec[ipol]

        self.wavevector = torch.empty([2,self.Npupil,self.Npupil], dtype=torch.complex64, device='cuda')
        self.wavevector[0] = 2 * np.pi * self.NA / self.wavelength * xpupil
        self.wavevector[1] = 2 * np.pi * self.NA / self.wavelength * ypupil
        self.wavevectorzimm = 2 * np.pi * self.refimm / self.wavelength * costhetaimm
        self.wavevectorzmed = 2 * np.pi * self.refmed / self.wavelength * costhetamed

        # calculate aberration function
        waberration = torch.zeros_like(xpupil,dtype=torch.complex64,device='cuda')
        normfac = torch.sqrt(2 * (self.orders[:, 0] + 1) / (1 + torch.where(self.orders[:, 1] == 0,1.0,0.0)))
        zernikecoefs_norm = self.zernikecoefs * normfac
        allzernikes = self.get_zernikefunc(self.orders, xpupil, ypupil)

        for izer in range(self.orders.shape[0]):
            waberration += zernikecoefs_norm[izer] * allzernikes[izer]
        waberration *= aperturemask
        phasefactor = torch.exp(1j * 2 * np.pi * waberration / self.wavelength)

        self.pupilmatrix = torch.empty([2, 3,self.Npupil,self.Npupil], dtype=torch.complex64, device='cuda')
        for imat in range(2):
            for jmat in range(3):
                self.pupilmatrix[imat, jmat] = self.amplitude * phasefactor * self.polarizationvector[imat, jmat]

        # czt transform(fft the pupil)
        xrange = self.pixel_size_x * self.Npixels / 2
        yrange = self.pixel_size_y * self.Npixels / 2
        imagesizex = xrange * self.NA / self.wavelength
        imagesizey = yrange * self.NA / self.wavelength

        # calculate the auxiliary vectors for chirpz
        self.ax, self.bx, self.dx = self.prechirpz(pupil_size, imagesizex, self.Npupil, self.Npixels)
        self.ay, self.by, self.dy = self.prechirpz(pupil_size, imagesizey, self.Npupil, self.Npixels)

        # # calculate PSFs
        # FieldMatrix = torch.empty([2, 3, Nmol,self.Npixels,self.Npixels], dtype=torch.complex64, device='cuda')
        # for jz in range(Nmol):
        #     # xyz induced phase
        #     if self.zemit[jz] + self.zemit0 >= 0:
        #         Wxyz = (-1 * self.xemit[jz]) * wavevector[0] + (-1 * self.yemit[jz]) * wavevector[1] + (
        #                     self.zemit[jz] + self.zemit0) * wavevectorzmed
        #         PositionPhase = torch.exp(1j * (Wxyz + (self.objstage[jz] + self.objstage0) * wavevectorzimm))
        #     else:
        #         Wxyz = (-1 * self.xemit[jz]) * wavevector[0] + (-1 * self.yemit[jz]) * wavevector[1]
        #         PositionPhase = torch.exp(1j * (Wxyz + (self.objstage[jz] + self.objstage0 + self.zemit[jz] + self.zemit0) * wavevectorzimm))
        #     # t0 = time.time()
        #     for itel in range(2):
        #         for jtel in range(3):
        #             Pupilfunction = PositionPhase * pupilmatrix[itel, jtel]
        #             inter_image = torch.transpose(self.czt(Pupilfunction, ay, by, dy),1,0)
        #             FieldMatrix[itel, jtel, jz] = torch.transpose(self.czt(inter_image, ax, bx, dx),1,0)
        #     # print('torch z loop: ', time.time() - t0)
        #
        # PSFs = torch.zeros([Nmol, self.Npixels, self.Npixels],device='cuda',dtype=torch.float64)
        # for jz in range(Nmol):
        #     for jtel in range(3):
        #         for itel in range(2):
        #             PSFs[jz, :, :] += 1 / 3 * (torch.abs(FieldMatrix[itel, jtel, jz])) ** 2

        # calculate intensity normalization function using the PSFs at focus
        FieldMatrix_norm = torch.empty([2, 3, self.Npixels, self.Npixels], dtype=torch.complex64, device='cuda')
        for itel in range(2):
            for jtel in range(3):
                Pupilfunction_norm = self.amplitude * self.polarizationvector[itel, jtel]
                inter_image_norm = torch.transpose(self.czt(Pupilfunction_norm, self.ay, self.by, self.dy),1,0)
                FieldMatrix_norm[itel, jtel] = torch.transpose(self.czt(inter_image_norm, self.ax, self.bx, self.dx),1,0)

        intFocus = torch.zeros([self.Npixels, self.Npixels], dtype=torch.float64,device='cuda')
        for jtel in range(3):
            for itel in range(2):
                intFocus += 1 / 3 * (torch.abs(FieldMatrix_norm[itel, jtel])) ** 2
        self.normIntensity = torch.sum(intFocus)

        # build theta_GT
        self.num_zernike = len(self.zernikecoefs)
        self.shared = np.concatenate((np.ones(self.num_zernike),np.array([0, 0, 0, 0, 0])),axis=0)
        sum_shared = np.sum(self.shared)
        self.num_pars = int((self.num_zernike+5)*self.Nmol-sum_shared*(self.Nmol-1))


        bg_GT = torch.zeros(self.Nmol)
        Nph_GT = torch.zeros(self.Nmol)
        x_GT = torch.zeros(self.Nmol)
        y_GT = torch.zeros(self.Nmol)
        z_GT = torch.zeros(self.Nmol)

        for i in range(self.Nmol):
            bg_GT[i] = torch.squeeze(self.bg[i])
            Nph_GT[i] = torch.squeeze(self.Nphotons[i])
        x_GT = torch.squeeze(self.xemit)
        y_GT = torch.squeeze(self.yemit)
        z_GT = torch.squeeze(self.zemit)

        alltheta_GT = torch.zeros((self.num_zernike+5,self.Nmol))
        alltheta_GT[self.num_zernike, :] = x_GT
        alltheta_GT[self.num_zernike+1, :] = y_GT
        alltheta_GT[self.num_zernike+2, :] = z_GT
        alltheta_GT[self.num_zernike+3, :] = Nph_GT
        alltheta_GT[self.num_zernike+4, :] = bg_GT
        alltheta_GT[0:self.num_zernike,:] = self.zernikecoefs.unflatten(0, (self.num_zernike, 1)).expand(self.num_zernike,self.Nmol)

        # shared map
        shared_map = np.zeros((self.num_pars, 3),dtype=int)
        n = 0
        for i in range(self.num_zernike+5):
            if self.shared[i] == 1:
                shared_map[n, 0] = 1
                shared_map[n, 1] = i+1
                shared_map[n, 2] = 0
                n += 1
            elif self.shared[i] == 0:
                for j in range(self.Nmol):
                    shared_map[n, 0] = 0
                    shared_map[n, 1] = i+1
                    shared_map[n, 2] = j+1
                    n += 1
        self.shared_map = shared_map

        thetainit_GT = torch.zeros(self.num_pars)
        for i in range(self.num_pars):
            if shared_map[i,0]==1:
                thetainit_GT[i] = torch.mean(alltheta_GT[shared_map[i,1]-1,:])
            elif shared_map[i,0]==0:
                thetainit_GT[i] = alltheta_GT[shared_map[i, 1]-1, shared_map[i, 2]-1]

        # calculate dudt using analytical functions
        [dudt, model] = self._cal_dudt(thetainit_GT, alltheta_GT)

        # calculate dudt using pytorch
        # self.optimizer_pars = torch.optim.AdamW([self.refmed, self.refcov, self.refimm, self.objstage0, self.zemit0,
        #                                    self.zernikecoefs, self.Nphotons, self.bg, self.xemit, self.yemit,self.zemit,
        #                                    self.objstage],lr=1e-3)
        # dudt = np.zeros([self.Npixels,self.Npixels,Nmol,5])
        # for jz in range(Nmol):
        #     print(jz)
        #     for jx in range(self.Npixels):
        #         for jy in range(self.Npixels):
        #             self.optimizer_pars.zero_grad()
        #             model[jz, jx, jy].backward(retain_graph=True)
        #             dudt[jx, jy, jz, 0] = cpu(self.xemit.grad[jz])
        #             dudt[jx, jy, jz, 1] = cpu(self.yemit.grad[jz])
        #             dudt[jx, jy, jz, 2] = cpu(self.zemit.grad[jz])
        #             dudt[jx, jy, jz, 3] = cpu(self.Nphotons.grad[jz, 0, 0])
        #             dudt[jx, jy, jz, 4] = cpu(self.bg.grad[jz, 0, 0])


        # calculate hessian matrix
        Npars = self.Nmol*5
        dudt = dudt
        t2 = 1/model
        hessian = torch.zeros([Npars,Npars],device='cuda')
        for p1 in range(Npars):
            temp1_zind = int(np.floor(p1 / 5))
            temp1_pind = int(p1%5)
            temp1 = torch.squeeze(dudt[:,:,:,temp1_pind])

            for p2 in range(p1,Npars):
                temp2_zind = int(np.floor(p2 / 5))
                temp2_pind = int(p2%5)
                temp2 = torch.squeeze(dudt[:, :, :, temp2_pind])
                if 1:
                    if temp1_zind==temp2_zind:
                        temp = t2[temp1_zind,:,:]*temp1[temp1_zind,:,:]*temp2[temp2_zind,:,:]
                        hessian[p1,p2]=torch.sum(temp)
                        hessian[p2,p1]=hessian[p1,p2]

        # calculate local fisher matrix and crlb
        x_crlb = torch.zeros([self.Nmol,1],device='cuda')
        y_crlb = torch.zeros([self.Nmol,1],device='cuda')
        z_crlb = torch.zeros([self.Nmol,1],device='cuda')
        for j in range(self.Nmol):
            fisher_tmp = hessian[j*5:j*5+5,j*5:j*5+5]
            sqrt_crlb_tmp = torch.sqrt(torch.diag(torch.inverse(fisher_tmp)))
            x_crlb[j] = sqrt_crlb_tmp[0]
            y_crlb[j] = sqrt_crlb_tmp[1]
            z_crlb[j] = sqrt_crlb_tmp[2]

        return x_crlb,y_crlb,z_crlb, model

    def _cal_dudt(self, thetainit_GT, alltheta_GT):
        # calculate PSFs
        num_ders = 5
        FieldMatrix = torch.empty([2, 3, self.Nmol,self.Npixels,self.Npixels], dtype=torch.complex64, device='cuda')
        FieldMatrix_ders = torch.empty([2, 3, self.Nmol, num_ders,self.Npixels, self.Npixels], dtype=torch.complex64,
                                       device='cuda')
        for jz in range(self.Nmol):
            # xyz induced phase
            if self.zemit[jz] + self.zemit0 >= 0:
                Wxyz = (-1 * self.xemit[jz]) * self.wavevector[0] + (-1 * self.yemit[jz]) * self.wavevector[1] + (
                            self.zemit[jz] + self.zemit0) * self.wavevectorzmed
                PositionPhase = torch.exp(1j * (Wxyz + (self.objstage[jz] + self.objstage0) * self.wavevectorzimm))
            else:
                print("warning! the emitter's position may not have physical meaning, CRLB is not guaranteed accurate if refmed is not equal to refimm")
                Wxyz = (-1 * self.xemit[jz]) * self.wavevector[0] + (-1 * self.yemit[jz]) * self.wavevector[1]
                PositionPhase = torch.exp(1j * (Wxyz + (self.objstage[jz] + self.objstage0 + self.zemit[jz] + self.zemit0)
                                                * self.wavevectorzimm))
            # t0 = time.time()
            for itel in range(2):
                for jtel in range(3):
                    Pupilfunction = PositionPhase * self.pupilmatrix[itel, jtel]
                    inter_image = torch.transpose(self.czt(Pupilfunction, self.ay, self.by, self.dy),1,0)
                    FieldMatrix[itel, jtel, jz] = torch.transpose(self.czt(inter_image, self.ax, self.bx, self.dx),1,0)

                    # xy derivatives
                    Pupilfunction_x = -1j * self.wavevector[0] * PositionPhase * self.pupilmatrix[itel, jtel]
                    inter_image_x = torch.transpose(self.czt(Pupilfunction_x, self.ay, self.by, self.dy), 1, 0)
                    FieldMatrix_ders[itel,jtel,jz,0] = torch.transpose(
                        self.czt(inter_image_x, self.ax, self.bx, self.dx), 1, 0)

                    Pupilfunction_y = -1j * self.wavevector[1] * PositionPhase * self.pupilmatrix[itel, jtel]
                    inter_image_y = torch.transpose(self.czt(Pupilfunction_y, self.ay, self.by, self.dy), 1, 0)
                    FieldMatrix_ders[itel, jtel, jz, 1] = torch.transpose(
                        self.czt(inter_image_y, self.ax, self.bx, self.dx), 1, 0)

                    # z derivatives
                    Pupilfunction_z = 1j * self.wavevectorzmed * PositionPhase * self.pupilmatrix[itel, jtel]
                    inter_image_z = torch.transpose(self.czt(Pupilfunction_z, self.ay, self.by, self.dy), 1, 0)
                    FieldMatrix_ders[itel, jtel, jz, 2] = torch.transpose(
                        self.czt(inter_image_z, self.ax, self.bx, self.dx), 1, 0)

        PSFs = torch.zeros([self.Nmol, self.Npixels, self.Npixels],device='cuda',dtype=torch.float64)
        PSFs_ders = torch.zeros([self.Nmol, self.Npixels, self.Npixels, 3], device='cuda', dtype=torch.float64)
        for jz in range(self.Nmol):
            for jtel in range(3):
                for itel in range(2):
                    PSFs[jz, :, :] += 1 / 3 * (torch.abs(FieldMatrix[itel, jtel, jz])) ** 2
                    for jder in range(3):
                        PSFs_ders[jz,:,:,jder] = PSFs_ders[jz,:,:,jder]+2/3*torch.real(torch.conj(FieldMatrix[itel,jtel,jz])*
                                                                                       FieldMatrix_ders[itel,jtel,jz,jder])

        PSFs /= self.normIntensity
        PSFs_ders /= self.normIntensity

        # otf rescale
        if len(self.otf_rescale):
            I_sigmax = self.otf_rescale[0]
            I_sigmay = self.otf_rescale[1]
            if I_sigmax == 0 or I_sigmay == 0:
                print('sigma for otf rescale is 0')
            else:
                h = torch.cuda.DoubleTensor(otf_gauss2D(shape=[5, 5], Isigmax=I_sigmax, Isigmay=I_sigmay)) \
                                            .reshape([1, 1, 5, 5])
                PSFs_tmp = PSFs.view(self.Nmol, 1, self.Npixels, self.Npixels)
                tmp = nn.functional.conv2d(PSFs_tmp, h, padding=2, stride=1)
                PSFs = tmp.view(self.Nmol, self.Npixels, self.Npixels)

                PSFs_ders_tmp0 = PSFs_ders[:, :, :, 0].view(self.Nmol, 1, self.Npixels, self.Npixels)
                tmp0 = nn.functional.conv2d(PSFs_ders_tmp0, h, padding=2, stride=1)
                PSFs_ders_tmp1 = PSFs_ders[:, :, :, 1].view(self.Nmol, 1, self.Npixels, self.Npixels)
                tmp1 = nn.functional.conv2d(PSFs_ders_tmp1, h, padding=2, stride=1)
                PSFs_ders_tmp2 = PSFs_ders[:, :, :, 2].view(self.Nmol, 1, self.Npixels, self.Npixels)
                tmp2 = nn.functional.conv2d(PSFs_ders_tmp2, h, padding=2, stride=1)

                PSFs_ders[:, :, :, 0] = tmp0.view(self.Nmol, self.Npixels, self.Npixels)
                PSFs_ders[:, :, :, 1] = tmp1.view(self.Nmol, self.Npixels, self.Npixels)
                PSFs_ders[:, :, :, 2] = tmp2.view(self.Nmol, self.Npixels, self.Npixels)

        model = torch.zeros_like(PSFs)
        for i in range(self.Nmol):
            model[i,:,:] = PSFs[i,:,:]*self.Nphotons[i]+self.bg[i]

        dudt = torch.zeros([self.Nmol, self.Npixels, self.Npixels, 5], device='cuda', dtype=torch.float64)
        for i in range(self.Nmol):
            dudt[i, :, :, 0] = self.Nphotons[i] * PSFs_ders[i, :, :, 0]
            dudt[i, :, :, 1] = self.Nphotons[i] * PSFs_ders[i, :, :, 1]
            dudt[i, :, :, 2] = self.Nphotons[i] * PSFs_ders[i, :, :, 2]

        dudt[:, :, :, 3] = PSFs
        dudt[:, :, :, 4] = torch.ones_like(PSFs)
        return dudt, model
```
