## Supplementary Software for "Field dependent deep learning enables high-throughput whole-cell 3D super-resolution imaging": local_tifffile.py

```
# -*- coding: utf-8 -*-
# local_tifffile.py

# Copyright (c) 2008-2019, Christoph Gohlke
# Copyright (c) 2008-2019, The Regents of the University of California
# Produced at the Laboratory for Fluorescence Dynamics
# All rights reserved.
#
# Redistribution and use in source and binary forms, with or without
# modification, are permitted provided that the following conditions are met:
#
# * Redistributions of source code must retain the above copyright notice,
#   this list of conditions and the following disclaimer.
#
# * Redistributions in binary form must reproduce the above copyright notice,
#   this list of conditions and the following disclaimer in the documentation
#   and/or other materials provided with the distribution.
#
# * Neither the name of the copyright holder nor the names of its
#   contributors may be used to endorse or promote products derived from
#   this software without specific prior written permission.
#
# THIS SOFTWARE IS PROVIDED BY THE COPYRIGHT HOLDERS AND CONTRIBUTORS "AS IS"
# AND ANY EXPRESS OR IMPLIED WARRANTIES, INCLUDING, BUT NOT LIMITED TO, THE
# IMPLIED WARRANTIES OF MERCHANTABILITY AND FITNESS FOR A PARTICULAR PURPOSE
# ARE DISCLAIMED. IN NO EVENT SHALL THE COPYRIGHT HOLDER OR CONTRIBUTORS BE
# LIABLE FOR ANY DIRECT, INDIRECT, INCIDENTAL, SPECIAL, EXEMPLARY, OR
# CONSEQUENTIAL DAMAGES (INCLUDING, BUT NOT LIMITED TO, PROCUREMENT OF
# SUBSTITUTE GOODS OR SERVICES; LOSS OF USE, DATA, OR PROFITS; OR BUSINESS
# INTERRUPTION) HOWEVER CAUSED AND ON ANY THEORY OF LIABILITY, WHETHER IN
# CONTRACT, STRICT LIABILITY, OR TORT (INCLUDING NEGLIGENCE OR OTHERWISE)
# ARISING IN ANY WAY OUT OF THE USE OF THIS SOFTWARE, EVEN IF ADVISED OF THE
# POSSIBILITY OF SUCH DAMAGE.

"""Read and write TIFF(r) files.

Tifffile is a Python library to

(1) store numpy arrays in TIFF (Tagged Image File Format) files, and
(2) read image and metadata from TIFF-like files used in bioimaging.

Image and metadata can be read from TIFF, BigTIFF, OME-TIFF, STK, LSM, SGI,
NIHImage, ImageJ, MicroManager, FluoView, ScanImage, SEQ, GEL, SVS, SCN, SIS,
ZIF, QPI, NDPI, and GeoTIFF files.

Numpy arrays can be written to TIFF, BigTIFF, and ImageJ hyperstack compatible
files in multi-page, memory-mappable, tiled, predicted, or compressed form.

Only a subset of the TIFF specification is supported, mainly uncompressed and
losslessly compressed 1, 8, 16, 32 and 64-bit integer, 16, 32 and 64-bit float,
grayscale and RGB(A) images.
Specifically, reading slices of image data, CCITT and OJPEG compression,
chroma subsampling without JPEG compression, or IPTC and XMP metadata are not
implemented.

TIFF(r), the Tagged Image File Format, is a trademark and under control of
Adobe Systems Incorporated. BigTIFF allows for files greater than 4 GB.
STK, LSM, FluoView, SGI, SEQ, GEL, and OME-TIFF, are custom extensions
defined by Molecular Devices (Universal Imaging Corporation), Carl Zeiss
MicroImaging, Olympus, Silicon Graphics International, Media Cybernetics,
Molecular Dynamics, and the Open Microscopy Environment consortium
respectively.

For command line usage run ``python -m tifffile --help``

:Author:
  `Christoph Gohlke <https://www.lfd.uci.edu/~gohlke/>`_

:Organization:
  Laboratory for Fluorescence Dynamics, University of California, Irvine

:License: 3-clause BSD

:Version: 2019.3.18

Requirements
------------
This release has been tested with the following requirements and dependencies
(other versions may work):

* `CPython 2.7.16, 3.5.4, 3.6.8, 3.7.2, 64-bit <https://www.python.org>`_
* `Numpy 1.15.4 <https://www.numpy.org>`_
* `Imagecodecs 2019.2.22 <https://pypi.org/project/imagecodecs/>`_
  (optional; used for decoding LZW, JPEG, etc.)
* `Matplotlib 2.2 <https://www.matplotlib.org>`_ (optional; used for plotting)
* Python 2.7 requires 'futures', 'enum34', and 'pathlib'.

Revisions
---------
2019.3.18
    Pass 2756 tests.
    Fix regression decoding JPEG with RGB photometrics.
    Fix reading OME-TIFF files with corrupted but unused pages.
    Allow to load TiffFrame without specifying keyframe.
    Calculate virtual TiffFrames for non-BigTIFF ScanImage files > 2GB.
    Rename property is_chroma_subsampled to is_subsampled.
    Make more attributes and methods private (WIP).
2019.3.8
    Fix MemoryError when RowsPerStrip > ImageLength.
    Fix SyntaxWarning on Python 3.8.
    Fail to decode JPEG to planar RGB for now.
    Separate public from private test files (WIP).
    Allow testing without data files or imagecodecs.
2019.2.22
    Use imagecodecs-lite as a fallback for imagecodecs.
    Simplify reading numpy arrays from file.
    Use TiffFrames when reading arrays from page sequences.
    Support slices and iterators in TiffPageSeries sequence interface.
    Auto-detect uniform series.
    Use page hash to determine generic series.
    Turn off page cache (tentative).
    Pass through more parameters in imread.
    Discontinue movie parameter in imread and TiffFile (breaking).
    Discontinue bigsize parameter in imwrite (breaking).
    Raise TiffFileError in case of issues with TIFF structure.
    Return TiffFile.ome_metadata as XML (breaking).
    Ignore OME series when last dimensions are not stored in TIFF pages.
2019.2.10
    Assemble IFDs in memory to speed-up writing on some slow media.
    Handle discontinued arguments fastij, multifile_close, and pages.
2019.1.30
    Use black background in imshow.
    Do not write datetime tag by default (breaking).
    Fix OME-TIFF with SamplesPerPixel > 1.
    Allow 64-bit IFD offsets for NDPI (files > 4GB still not supported).
2019.1.4
    Fix decoding deflate without imagecodecs.
2019.1.1
    Update copyright year.
    Require imagecodecs >= 2018.12.16.
    Do not use JPEG tables from keyframe.
    Enable decoding large JPEG in NDPI.
    Decode some old-style JPEG.
    Reorder OME channel axis to match PlanarConfiguration storage.
    Return tiled images as contiguous arrays.
    Add decode_lzw proxy function for compatibility with old czifile module.
    Use dedicated logger.
2018.11.28
    Make SubIFDs accessible as TiffPage.pages.
    Make parsing of TiffSequence axes pattern optional (breaking).
    Limit parsing of TiffSequence axes pattern to file names, not path names.
    Do not interpolate in imshow if image dimensions <= 512, else use bilinear.
    Use logging.warning instead of warnings.warn in many cases.
    Fix numpy FutureWarning for out == 'memmap'.
    Adjust ZSTD and WebP compression to libtiff-4.0.10 (WIP).
    Decode old-style LZW with imagecodecs >= 2018.11.8.
    Remove TiffFile.qptiff_metadata (QPI metadata are per page).
    Do not use keyword arguments before variable positional arguments.
    Make either all or none return statements in a function return expression.
    Use pytest parametrize to generate tests.
    Replace test classes with functions.
2018.11.6
    Rename imsave function to imwrite.
    Readd Python implementations of packints, delta, and bitorder codecs.
    Fix TiffFrame.compression AttributeError.
2018.10.18
    Rename tiffile package to tifffile.
2018.10.10
    Read ZIF, the Zoomable Image Format (WIP).
    Decode YCbCr JPEG as RGB (tentative).
    Improve restoration of incomplete tiles.
    Allow to write grayscale with extrasamples without specifying planarconfig.
    Enable decoding of PNG and JXR via imagecodecs.
    Deprecate 32-bit platforms (too many memory errors during tests).
2018.9.27
    Read Olympus SIS (WIP).
    Allow to write non-BigTIFF files up to ~4 GB (fix).
    Fix parsing date and time fields in SEM metadata.
    Detect some circular IFD references.
    Enable WebP codecs via imagecodecs.
    Add option to read TiffSequence from ZIP containers.
    Remove TiffFile.isnative.
    Move TIFF struct format constants out of TiffFile namespace.
2018.8.31
    Fix wrong TiffTag.valueoffset.
    Towards reading Hamamatsu NDPI (WIP).
    Enable PackBits compression of byte and bool arrays.
    Fix parsing NULL terminated CZ_SEM strings.
2018.8.24
    Move local_tifffile.py and related modules into tiffile package.
    Move usage examples to module docstring.
    Enable multi-threading for compressed tiles and pages by default.
    Add option to concurrently decode image tiles using threads.
    Do not skip empty tiles (fix).
    Read JPEG and J2K compressed strips and tiles.
    Allow floating-point predictor on write.
    Add option to specify subfiletype on write.
    Depend on imagecodecs package instead of _tifffile, lzma, etc modules.
    Remove reverse_bitorder, unpack_ints, and decode functions.
    Use pytest instead of unittest.
2018.6.20
    Save RGBA with unassociated extrasample by default (breaking).
    Add option to specify ExtraSamples values.
2018.6.17 (included with 0.15.1)
    Towards reading JPEG and other compressions via imagecodecs package (WIP).
    Read SampleFormat VOID as UINT.
    Add function to validate TIFF using 'jhove -m TIFF-hul'.
    Save bool arrays as bilevel TIFF.
    Accept pathlib.Path as filenames.
    Move 'software' argument from TiffWriter __init__ to save.
    Raise DOS limit to 16 TB.
    Lazy load lzma and zstd compressors and decompressors.
    Add option to save IJMetadata tags.
    Return correct number of pages for truncated series (fix).
    Move EXIF tags to TIFF.TAG as per TIFF/EP standard.
2018.2.18
    Always save RowsPerStrip and Resolution tags as required by TIFF standard.
    Do not use badly typed ImageDescription.
    Coherce bad ASCII string tags to bytes.
    Tuning of __str__ functions.
    Fix reading 'undefined' tag values.
    Read and write ZSTD compressed data.
    Use hexdump to print byte strings.
    Determine TIFF byte order from data dtype in imsave.
    Add option to specify RowsPerStrip for compressed strips.
    Allow memory-map of arrays with non-native byte order.
    Attempt to handle ScanImage <= 5.1 files.
    Restore TiffPageSeries.pages sequence interface.
    Use numpy.frombuffer instead of fromstring to read from binary data.
    Parse GeoTIFF metadata.
    Add option to apply horizontal differencing before compression.
    Towards reading PerkinElmer QPI (QPTIFF, no test files).
    Do not index out of bounds data in tifffile.c unpackbits and decodelzw.
2017.9.29
    Many backward incompatible changes improving speed and resource usage:
    Add detail argument to __str__ function. Remove info functions.
    Fix potential issue correcting offsets of large LSM files with positions.
    Remove TiffFile sequence interface; use TiffFile.pages instead.
    Do not make tag values available as TiffPage attributes.
    Use str (not bytes) type for tag and metadata strings (WIP).
    Use documented standard tag and value names (WIP).
    Use enums for some documented TIFF tag values.
    Remove 'memmap' and 'tmpfile' options; use out='memmap' instead.
    Add option to specify output in asarray functions.
    Add option to concurrently decode pages using threads.
    Add TiffPage.asrgb function (WIP).
    Do not apply colormap in asarray.
    Remove 'colormapped', 'rgbonly', and 'scale_mdgel' options from asarray.
    Consolidate metadata in TiffFile _metadata functions.
    Remove non-tag metadata properties from TiffPage.
    Add function to convert LSM to tiled BIN files.
    Align image data in file.
    Make TiffPage.dtype a numpy.dtype.
    Add 'ndim' and 'size' properties to TiffPage and TiffPageSeries.
    Allow imsave to write non-BigTIFF files up to ~4 GB.
    Only read one page for shaped series if possible.
    Add memmap function to create memory-mapped array stored in TIFF file.
    Add option to save empty arrays to TIFF files.
    Add option to save truncated TIFF files.
    Allow single tile images to be saved contiguously.
    Add optional movie mode for files with uniform pages.
    Lazy load pages.
    Use lightweight TiffFrame for IFDs sharing properties with key TiffPage.
    Move module constants to 'TIFF' namespace (speed up module import).
    Remove 'fastij' option from TiffFile.
    Remove 'pages' parameter from TiffFile.
    Remove TIFFfile alias.
    Deprecate Python 2.
    Require enum34 and futures packages on Python 2.7.
    Remove Record class and return all metadata as dict instead.
    Add functions to parse STK, MetaSeries, ScanImage, SVS, Pilatus metadata.
    Read tags from EXIF and GPS IFDs.
    Use pformat for tag and metadata values.
    Fix reading some UIC tags.
    Do not modify input array in imshow (fix).
    Fix Python implementation of unpack_ints.
2017.5.23
    Write correct number of SampleFormat values (fix).
    Use Adobe deflate code to write ZIP compressed files.
    Add option to pass tag values as packed binary data for writing.
    Defer tag validation to attribute access.
    Use property instead of lazyattr decorator for simple expressions.
2017.3.17
    Write IFDs and tag values on word boundaries.
    Read ScanImage metadata.
    Remove is_rgb and is_indexed attributes from TiffFile.
    Create files used by doctests.
2017.1.12 (included with scikit-image 0.14.x)
    Read Zeiss SEM metadata.
    Read OME-TIFF with invalid references to external files.
    Rewrite C LZW decoder (5x faster).
    Read corrupted LSM files missing EOI code in LZW stream.
2017.1.1
    ...

Refer to the CHANGES file for older revisions.

Notes
-----
The API is not stable yet and might change between revisions.

Tested on little-endian platforms only.

Python 2.7 and 32-bit versions are deprecated.

Tifffile relies on the `imagecodecs <https://pypi.org/project/imagecodecs/>`_
package for decoding LZW, JPEG, and other compressed images. Alternatively,
the `imagecodecs-lite <https://pypi.org/project/imagecodecs-lite/>`_ package
can be used for decoding LZW compressed images.

There are several TIFF-like formats (not adhering to the TIFF6 specification)
that allow files to exceed the 4 GB limit:

* *BigTIFF* is identified by version number 43 and uses different file
  header, IFD, and tag structures with 64-bit offsets. It adds more data types.
  Tifffile can read and write BigTIFF files.
* *ImageJ* hyperstacks store all image data, which may exceed 4 GB,
  contiguously after the first IFD. Files > 4 GB contain one IFD only.
  The size (shape and dtype) of the image data can be determined from the
  ImageDescription of the first IFD. Tifffile can read and write ImageJ
  hyperstacks.
* *LSM* stores all IFDs below 4 GB but wraps around 32-bit StripOffsets.
  The StripOffsets of each series and position require separate unwrapping.
  The StripByteCounts tag contains the number of bytes for the uncompressed
  data. Tifffile can read large LSM files.
* *NDPI* uses some 64-bit offsets in the file header, IFD, and tag structures
  and might require correcting 32-bit offsets found in tags.
  JPEG compressed tiles with dimensions > 65536 are not readable with libjpeg.
  Tifffile can read NDPI files < 4 GB and decompress large JPEG tiles using
  the imagecodecs library on Windows.
* *ScanImage* optionally writes corrupt non-BigTIFF files > 2 GB. The values
  of StripOffsets and StripByteCounts can be recovered using the constant
  differences of the offsets of IFD and tag values throughout the file.
  TiffFile can read such files on Python 3 if the image data is stored
  contiguously in each page.

Other libraries for reading scientific TIFF files from Python:

*  `Python-bioformats <https://github.com/CellProfiler/python-bioformats>`_
*  `Imread <https://github.com/luispedro/imread>`_
*  `GDAL <https://github.com/OSGeo/gdal/tree/master/gdal/swig/python>`_
*  `OpenSlide-python <https://github.com/openslide/openslide-python>`_
*  `PyLibTiff <https://github.com/pearu/pylibtiff>`_
*  `SimpleITK <https://github.com/SimpleITK/SimpleITK>`_
*  `PyLSM <https://launchpad.net/pylsm>`_
*  `PyMca.TiffIO.py <https://github.com/vasole/pymca>`_ (same as fabio.TiffIO)
*  `BioImageXD.Readers <http://www.bioimagexd.net/>`_
*  `CellCognition <https://cellcognition-project.org/>`_
*  `pymimage <https://github.com/ardoi/pymimage>`_
*  `pytiff <https://github.com/FZJ-INM1-BDA/pytiff>`_
*  `ScanImageTiffReaderPython
   <https://gitlab.com/vidriotech/scanimagetiffreader-python>`_

Acknowledgements
----------------
*   Egor Zindy, University of Manchester, for lsm_scan_info specifics.
*   Wim Lewis for a bug fix and some LSM functions.
*   Hadrien Mary for help on reading MicroManager files.
*   Christian Kliche for help writing tiled and color-mapped files.

References
----------
1)  TIFF 6.0 Specification and Supplements. Adobe Systems Incorporated.
    https://www.adobe.io/open/standards/TIFF.html
2)  TIFF File Format FAQ. https://www.awaresystems.be/imaging/tiff/faq.html
3)  MetaMorph Stack (STK) Image File Format.
    http://mdc.custhelp.com/app/answers/detail/a_id/18862
4)  Image File Format Description LSM 5/7 Release 6.0 (ZEN 2010).
    Carl Zeiss MicroImaging GmbH. BioSciences. May 10, 2011
5)  The OME-TIFF format.
    https://docs.openmicroscopy.org/ome-model/5.6.4/ome-tiff/
6)  UltraQuant(r) Version 6.0 for Windows Start-Up Guide.
    http://www.ultralum.com/images%20ultralum/pdf/UQStart%20Up%20Guide.pdf
7)  Micro-Manager File Formats.
    https://micro-manager.org/wiki/Micro-Manager_File_Formats
8)  Tags for TIFF and Related Specifications. Digital Preservation.
    https://www.loc.gov/preservation/digital/formats/content/tiff_tags.shtml
9)  ScanImage BigTiff Specification - ScanImage 2016.
    http://scanimage.vidriotechnologies.com/display/SI2016/
    ScanImage+BigTiff+Specification
10) CIPA DC-008-2016: Exchangeable image file format for digital still cameras:
    Exif Version 2.31.
    http://www.cipa.jp/std/documents/e/DC-008-Translation-2016-E.pdf
11) ZIF, the Zoomable Image File format. http://zif.photo/

Examples
--------
Save a 3D numpy array to a multi-page, 16-bit grayscale TIFF file:

>>> data = numpy.random.randint(0, 2**12, (4, 301, 219), 'uint16')
>>> imwrite('temp.tif', data, photometric='minisblack')

Read the whole image stack from the TIFF file as numpy array:

>>> image_stack = imread('temp.tif')
>>> image_stack.shape
(4, 301, 219)
>>> image_stack.dtype
dtype('uint16')

Read the image from first page in the TIFF file as numpy array:

>>> image = imread('temp.tif', key=0)
>>> image.shape
(301, 219)

Read images from a sequence of TIFF files as numpy array:

>>> image_sequence = imread(['temp.tif', 'temp.tif'])
>>> image_sequence.shape
(2, 4, 301, 219)

Save a numpy array to a single-page RGB TIFF file:

>>> data = numpy.random.randint(0, 255, (256, 256, 3), 'uint8')
>>> imwrite('temp.tif', data, photometric='rgb')

Save a floating-point array and metadata, using zlib compression:

>>> data = numpy.random.rand(2, 5, 3, 301, 219).astype('float32')
>>> imwrite('temp.tif', data, compress=6, metadata={'axes': 'TZCYX'})

Save a volume with xyz voxel size 2.6755x2.6755x3.9474 µm^3 to an ImageJ file:

>>> volume = numpy.random.randn(57*256*256).astype('float32')
>>> volume.shape = 1, 57, 1, 256, 256, 1  # dimensions in TZCYXS order
>>> imwrite('temp.tif', volume, imagej=True, resolution=(1./2.6755, 1./2.6755),
...         metadata={'spacing': 3.947368, 'unit': 'um'})

Read hyperstack and metadata from the ImageJ file:

>>> with TiffFile('temp.tif') as tif:
...     imagej_hyperstack = tif.asarray()
...     imagej_metadata = tif.imagej_metadata
>>> imagej_hyperstack.shape
(57, 256, 256)
>>> imagej_metadata['slices']
57

Read the "XResolution" tag from the first page in the TIFF file:

>>> with TiffFile('temp.tif') as tif:
...     tag = tif.pages[0].tags['XResolution']
>>> tag.value
(2000, 5351)
>>> tag.name
'XResolution'
>>> tag.code
282
>>> tag.count
1
>>> tag.dtype
'2I'
>>> tag.valueoffset
360

Read images from a selected range of pages:

>>> image = imread('temp.tif', key=range(4, 40, 2))
>>> image.shape
(18, 256, 256)

Create an empty TIFF file and write to the memory-mapped numpy array:

>>> memmap_image = memmap('temp.tif', shape=(256, 256), dtype='float32')
>>> memmap_image[255, 255] = 1.0
>>> memmap_image.flush()
>>> memmap_image.shape, memmap_image.dtype
((256, 256), dtype('float32'))
>>> del memmap_image

Memory-map image data of the first page in the TIFF file:

>>> memmap_image = memmap('temp.tif', page=0)
>>> memmap_image[255, 255]
1.0
>>> del memmap_image

Successively append images to a BigTIFF file:

>>> data = numpy.random.randint(0, 255, (5, 2, 3, 301, 219), 'uint8')
>>> with TiffWriter('temp.tif', bigtiff=True) as tif:
...     for i in range(data.shape[0]):
...         tif.save(data[i], compress=6, photometric='minisblack')

Iterate over pages and tags in the TIFF file and successively read images:

>>> with TiffFile('temp.tif') as tif:
...     image_stack = tif.asarray()
...     for page in tif.pages:
...         for tag in page.tags.values():
...             tag_name, tag_value = tag.name, tag.value
...         image = page.asarray()

Save two image series to a TIFF file:

>>> data0 = numpy.random.randint(0, 255, (301, 219, 3), 'uint8')
>>> data1 = numpy.random.randint(0, 255, (5, 301, 219), 'uint16')
>>> with TiffWriter('temp.tif') as tif:
...     tif.save(data0, compress=6, photometric='rgb')
...     tif.save(data1, compress=6, photometric='minisblack', contiguous=False)

Read the second image series from the TIFF file:

>>> series1 = imread('temp.tif', series=1)
>>> series1.shape
(5, 301, 219)

Read an image stack from a sequence of TIFF files with a file name pattern:

>>> imwrite('temp_C001T001.tif', numpy.random.rand(64, 64))
>>> imwrite('temp_C001T002.tif', numpy.random.rand(64, 64))
>>> image_sequence = TiffSequence('temp_C001*.tif', pattern='axes')
>>> image_sequence.shape
(1, 2)
>>> image_sequence.axes
'CT'
>>> data = image_sequence.asarray()
>>> data.shape
(1, 2, 64, 64)

"""

from __future__ import division, print_function

__version__ = '2019.3.18'
__docformat__ = 'restructuredtext en'
__all__ = ('imwrite', 'imsave', 'imread', 'imshow', 'memmap', 'lsm2bin',
           'TiffFile', 'TiffFileError', 'TiffWriter', 'TiffSequence',
           'FileHandle', 'TiffPage', 'TiffFrame', 'TiffTag', 'TIFF',
           # utility functions used by oiffile, czifile, etc
           'lazyattr', 'natural_sorted', 'stripnull', 'transpose_axes',
           'squeeze_axes', 'create_output', 'repeat_nd', 'format_size',
           'product', 'xml2dict', 'pformat', 'str2bytes', '_app_show',
           'decode_lzw')

import sys
import os
import io
import re
import glob
import math
import time
import json
import enum
import struct
import pathlib
import logging
import warnings
import binascii
import datetime
import threading
import collections

try:
    from collections.abc import Iterable
except ImportError:
    from collections import Iterable

from concurrent.futures import ThreadPoolExecutor

import numpy

try:
    import imagecodecs
except ImportError:
    import zlib
    try:
        import imagecodecs_lite as imagecodecs
    except ImportError:
        imagecodecs = None

# delay import of mmap, pprint, fractions, xml, tkinter, lxml, matplotlib,
#   subprocess, multiprocessing, tempfile, zipfile, fnmatch

log = logging.getLogger(__name__)  # .addHandler(logging.NullHandler())


def imread(files, **kwargs):
    """Return image data from TIFF file(s) as numpy array.

    Refer to the TiffFile and  TiffSequence classes and their asarray
    functions for documentation.

    Parameters
    ----------
    files : str, binary stream, or sequence
        File name, seekable binary stream, glob pattern, or sequence of
        file names.
    kwargs : dict
        Parameters 'name', 'offset', 'size', 'multifile', and 'is_ome'
        are passed to the TiffFile constructor.
        The 'pattern' parameter is passed to the TiffSequence constructor.
        Other parameters are passed to the asarray functions.
        The first image series in the file is returned if no arguments are
        provided.

    """
    kwargs_file = parse_kwargs(kwargs, 'is_ome', 'multifile', '_useframes',
                               'name', 'offset', 'size',
                               'multifile_close', 'fastij', 'movie')  # legacy
    kwargs_seq = parse_kwargs(kwargs, 'pattern')

    if kwargs.get('pages', None) is not None:
        if kwargs.get('key', None) is not None:
            raise TypeError(
                "the 'pages' and 'key' arguments cannot be used together")
        log.warning("imread: the 'pages' argument is deprecated")
        kwargs['key'] = kwargs.pop('pages')

    if isinstance(files, basestring) and any(i in files for i in '?*'):
        files = glob.glob(files)
    if not files:
        raise ValueError('no files found')
    if not hasattr(files, 'seek') and len(files) == 1:
        files = files[0]

    if isinstance(files, basestring) or hasattr(files, 'seek'):
        with TiffFile(files, **kwargs_file) as tif:
            return tif.asarray(**kwargs)
    else:
        with TiffSequence(files, **kwargs_seq) as imseq:
            return imseq.asarray(**kwargs)


def imwrite(file, data=None, shape=None, dtype=None, **kwargs):
    """Write numpy array to TIFF file.

    Refer to the TiffWriter class and its asarray function for documentation.

    A BigTIFF file is created if the data size in bytes is larger than 4 GB
    minus 32 MB (for metadata), and 'bigtiff' is not specified, and 'imagej'
    or 'truncate' are not enabled.

    Parameters
    ----------
    file : str or binary stream
        File name or writable binary stream, such as an open file or BytesIO.
    data : array_like
        Input image. The last dimensions are assumed to be image depth,
        height, width, and samples.
        If None, an empty array of the specified shape and dtype is
        saved to file.
        Unless 'byteorder' is specified in 'kwargs', the TIFF file byte order
        is determined from the data's dtype or the dtype argument.
    shape : tuple
        If 'data' is None, shape of an empty array to save to the file.
    dtype : numpy.dtype
        If 'data' is None, data-type of an empty array to save to the file.
    kwargs : dict
        Parameters 'append', 'byteorder', 'bigtiff', and 'imagej', are passed
        to the TiffWriter constructor. Other parameters are passed to the
        TiffWriter.save function.

    Returns
    -------
    offset, bytecount : tuple or None
        If the image data are written contiguously, return offset and bytecount
        of image data in the file.

    """
    tifargs = parse_kwargs(kwargs, 'append', 'bigtiff', 'byteorder', 'imagej')
    if data is None:
        dtype = numpy.dtype(dtype)
        size = product(shape) * dtype.itemsize
        byteorder = dtype.byteorder
    else:
        try:
            size = data.nbytes
            byteorder = data.dtype.byteorder
        except Exception:
            size = 0
            byteorder = None
    bigsize = kwargs.pop('bigsize', 2**32-2**25)
    if 'bigtiff' not in tifargs and size > bigsize and not (
            tifargs.get('imagej', False) or tifargs.get('truncate', False)):
        tifargs['bigtiff'] = True
    if 'byteorder' not in tifargs:
        tifargs['byteorder'] = byteorder

    with TiffWriter(file, **tifargs) as tif:
        return tif.save(data, shape, dtype, **kwargs)


imsave = imwrite


def memmap(filename, shape=None, dtype=None, page=None, series=0, mode='r+',
           **kwargs):
    """Return memory-mapped numpy array stored in TIFF file.

    Memory-mapping requires data stored in native byte order, without tiling,
    compression, predictors, etc.
    If 'shape' and 'dtype' are provided, existing files will be overwritten or
    appended to depending on the 'append' parameter.
    Otherwise the image data of a specified page or series in an existing
    file will be memory-mapped. By default, the image data of the first page
    series is memory-mapped.
    Call flush() to write any changes in the array to the file.
    Raise ValueError if the image data in the file is not memory-mappable.

    Parameters
    ----------
    filename : str
        Name of the TIFF file which stores the array.
    shape : tuple
        Shape of the empty array.
    dtype : numpy.dtype
        Data-type of the empty array.
    page : int
        Index of the page which image data to memory-map.
    series : int
        Index of the page series which image data to memory-map.
    mode : {'r+', 'r', 'c'}
        The file open mode. Default is to open existing file for reading and
        writing ('r+').
    kwargs : dict
        Additional parameters passed to imwrite() or TiffFile().

    """
    if shape is not None and dtype is not None:
        # create a new, empty array
        kwargs.update(data=None, shape=shape, dtype=dtype, returnoffset=True,
                      align=TIFF.ALLOCATIONGRANULARITY)
        result = imwrite(filename, **kwargs)
        if result is None:
            # TODO: fail before creating file or writing data
            raise ValueError('image data are not memory-mappable')
        offset = result[0]
    else:
        # use existing file
        with TiffFile(filename, **kwargs) as tif:
            if page is not None:
                page = tif.pages[page]
                if not page.is_memmappable:
                    raise ValueError('image data are not memory-mappable')
                offset, _ = page.is_contiguous
                shape = page.shape
                dtype = page.dtype
            else:
                series = tif.series[series]
                if series.offset is None:
                    raise ValueError('image data are not memory-mappable')
                shape = series.shape
                dtype = series.dtype
                offset = series.offset
            dtype = tif.byteorder + dtype.char
    return numpy.memmap(filename, dtype, mode, offset, shape, 'C')


class lazyattr(object):
    """Attribute whose value is computed on first access."""
    # TODO: help() doesn't work
    __slots__ = ('func',)

    def __init__(self, func):
        self.func = func
        # self.__name__ = func.__name__
        # self.__doc__ = func.__doc__
        # self.lock = threading.RLock()

    def __get__(self, instance, owner):
        # with self.lock:
        if instance is None:
            return self
        try:
            value = self.func(instance)
        except AttributeError as exc:
            raise RuntimeError(exc)
        if value is NotImplemented:
            return getattr(super(owner, instance), self.func.__name__)
        setattr(instance, self.func.__name__, value)
        return value


class TiffFileError(Exception):
    """Exception to indicate invalid TIFF structure."""


class TiffWriter(object):
    """Write numpy arrays to TIFF file.

    TiffWriter instances must be closed using the 'close' method, which is
    automatically called when using the 'with' context manager.

    TiffWriter's main purpose is saving nD numpy array's as TIFF,
    not to create any possible TIFF format. Specifically, JPEG compression,
    SubIFDs, ExifIFD, or GPSIFD tags are not supported.

    """

    def __init__(self, file, bigtiff=False, byteorder=None, append=False,
                 imagej=False):
        """Open a TIFF file for writing.

        An empty TIFF file is created if the file does not exist, else the
        file is overwritten with an empty TIFF file unless 'append'
        is true. Use 'bigtiff=True' when creating files larger than 4 GB.

        Parameters
        ----------
        file : str, binary stream, or FileHandle
            File name or writable binary stream, such as an open file
            or BytesIO.
        bigtiff : bool
            If True, the BigTIFF format is used.
        byteorder : {'<', '>', '=', '|'}
            The endianness of the data in the file.
            By default, this is the system's native byte order.
        append : bool
            If True and 'file' is an existing standard TIFF file, image data
            and tags are appended to the file.
            Appending data may corrupt specifically formatted TIFF files
            such as LSM, STK, ImageJ, or FluoView.
        imagej : bool
            If True, write an ImageJ hyperstack compatible file.
            This format can handle data types uint8, uint16, or float32 and
            data shapes up to 6 dimensions in TZCYXS order.
            RGB images (S=3 or S=4) must be uint8.
            ImageJ's default byte order is big-endian but this implementation
            uses the system's native byte order by default.
            ImageJ hyperstacks do not support BigTIFF or compression.
            The ImageJ file format is undocumented.
            When using compression, use ImageJ's Bio-Formats import function.

        """
        if append:
            # determine if file is an existing TIFF file that can be extended
            try:
                with FileHandle(file, mode='rb', size=0) as fh:
                    pos = fh.tell()
                    try:
                        with TiffFile(fh) as tif:
                            if append != 'force' and not tif.is_appendable:
                                raise TiffFileError('cannot append to file'
                                                    ' containing metadata')
                            byteorder = tif.byteorder
                            bigtiff = tif.is_bigtiff
                            self._ifdoffset = tif.pages.next_page_offset
                    finally:
                        fh.seek(pos)
            except (IOError, FileNotFoundError):
                append = False

        if byteorder in (None, '=', '|'):
            byteorder = '<' if sys.byteorder == 'little' else '>'
        elif byteorder not in ('<', '>'):
            raise ValueError('invalid byteorder %s' % byteorder)
        if imagej and bigtiff:
            warnings.warn('writing incompatible BigTIFF ImageJ')

        self._byteorder = byteorder
        self._imagej = bool(imagej)
        self._truncate = False
        self._metadata = None
        self._colormap = None

        self._descriptionoffset = 0
        self._descriptionlen = 0
        self._descriptionlenoffset = 0
        self._tags = None
        self._shape = None  # normalized shape of data in consecutive pages
        self._datashape = None  # shape of data in consecutive pages
        self._datadtype = None  # data type
        self._dataoffset = None  # offset to data
        self._databytecounts = None  # byte counts per plane
        self._tagoffsets = None  # strip or tile offset tag code

        if bigtiff:
            self._bigtiff = True
            self._offsetsize = 8
            self._tagsize = 20
            self._tagnoformat = 'Q'
            self._offsetformat = 'Q'
            self._valueformat = '8s'
        else:
            self._bigtiff = False
            self._offsetsize = 4
            self._tagsize = 12
            self._tagnoformat = 'H'
            self._offsetformat = 'I'
            self._valueformat = '4s'

        if append:
            self._fh = FileHandle(file, mode='r+b', size=0)
            self._fh.seek(0, 2)
        else:
            self._fh = FileHandle(file, mode='wb', size=0)
            self._fh.write({'<': b'II', '>': b'MM'}[byteorder])
            if bigtiff:
                self._fh.write(struct.pack(byteorder+'HHH', 43, 8, 0))
            else:
                self._fh.write(struct.pack(byteorder+'H', 42))
            # first IFD
            self._ifdoffset = self._fh.tell()
            self._fh.write(struct.pack(byteorder+self._offsetformat, 0))

    def save(self, data=None, shape=None, dtype=None, returnoffset=False,
             photometric=None, planarconfig=None, extrasamples=None, tile=None,
             contiguous=True, align=16, truncate=False, compress=0,
             rowsperstrip=None, predictor=False, colormap=None,
             description=None, datetime=None, resolution=None, subfiletype=0,
             software='local_tifffile.py', metadata={}, ijmetadata=None,
             extratags=()):
        """Write numpy array and tags to TIFF file.

        The data shape's last dimensions are assumed to be image depth,
        height (length), width, and samples.
        If a colormap is provided, the data's dtype must be uint8 or uint16
        and the data values are indices into the last dimension of the
        colormap.
        If 'shape' and 'dtype' are specified, an empty array is saved.
        This option cannot be used with compression or multiple tiles.
        Image data are written uncompressed in one strip per plane by default.
        Dimensions larger than 2 to 4 (depending on photometric mode, planar
        configuration, and SGI mode) are flattened and saved as separate pages.
        The SampleFormat and BitsPerSample tags are derived from the data type.

        Parameters
        ----------
        data : numpy.ndarray or None
            Input image array.
        shape : tuple or None
            Shape of the empty array to save. Used only if 'data' is None.
        dtype : numpy.dtype or None
            Data-type of the empty array to save. Used only if 'data' is None.
        returnoffset : bool
            If True and the image data in the file is memory-mappable, return
            the offset and number of bytes of the image data in the file.
        photometric : {'MINISBLACK', 'MINISWHITE', 'RGB', 'PALETTE', 'CFA'}
            The color space of the image data.
            By default, this setting is inferred from the data shape and the
            value of colormap.
            For CFA images, DNG tags must be specified in 'extratags'.
        planarconfig : {'CONTIG', 'SEPARATE'}
            Specifies if samples are stored interleaved or in separate planes.
            By default, this setting is inferred from the data shape.
            If this parameter is set, extra samples are used to store grayscale
            images.
            'CONTIG': last dimension contains samples.
            'SEPARATE': third last dimension contains samples.
        extrasamples : tuple of {'UNSPECIFIED', 'ASSOCALPHA', 'UNASSALPHA'}
            Defines the interpretation of extra components in pixels.
            'UNSPECIFIED': no transparency information (default).
            'ASSOCALPHA': single, true transparency with pre-multiplied color.
            'UNASSALPHA': independent transparency masks.
        tile : tuple of int
            The shape (depth, length, width) of image tiles to write.
            If None (default), image data are written in strips.
            The tile length and width must be a multiple of 16.
            If the tile depth is provided, the SGI ImageDepth and TileDepth
            tags are used to save volume data.
            Unless a single tile is used, tiles cannot be used to write
            contiguous files.
            Few software can read the SGI format, e.g. MeVisLab.
        contiguous : bool
            If True (default) and the data and parameters are compatible with
            previous ones, if any, the image data are stored contiguously after
            the previous one. In that case, 'photometric', 'planarconfig',
            'rowsperstrip', are ignored. Metadata such as 'description',
            'metadata', 'datetime', and 'extratags' are written to the first
            page of a contiguous series only.
        align : int
            Byte boundary on which to align the image data in the file.
            Default 16. Use mmap.ALLOCATIONGRANULARITY for memory-mapped data.
            Following contiguous writes are not aligned.
        truncate : bool
            If True, only write the first page including shape metadata if
            possible (uncompressed, contiguous, not tiled).
            Other TIFF readers will only be able to read part of the data.
        compress : int or str or (str, int)
            If 0 (default), data are written uncompressed.
            If 0-9, the level of ADOBE_DEFLATE compression.
            If a str, one of TIFF.COMPRESSION, e.g. 'LZMA' or 'ZSTD'.
            If a tuple, first item is one of TIFF.COMPRESSION and second item
            is compression level.
            Compression cannot be used to write contiguous files.
        rowsperstrip : int
            The number of rows per strip. By default strips will be ~64 KB
            if compression is enabled, else rowsperstrip is set to the image
            length. Bilevel images are always stored in one strip per plane.
        predictor : bool
            If True, apply horizontal differencing or floating-point predictor
            before compression.
        colormap : numpy.ndarray
            RGB color values for the corresponding data value.
            Must be of shape (3, 2**(data.itemsize*8)) and dtype uint16.
        description : str
            The subject of the image. Must be 7-bit ASCII. Cannot be used with
            the ImageJ format. Saved with the first page only.
        datetime : datetime, str, or bool
            Date and time of image creation in '%Y:%m:%d %H:%M:%S' format or
            datetime object. Else if True, the current date and time is used.
            Saved with the first page only.
        resolution : (float, float[, str]) or ((int, int), (int, int)[, str])
            X and Y resolutions in pixels per resolution unit as float or
            rational numbers. A third, optional parameter specifies the
            resolution unit, which must be None (default for ImageJ),
            'INCH' (default), or 'CENTIMETER'.
        subfiletype : int
            Bitfield to indicate the kind of data. Set bit 0 if the image
            is a reduced-resolution version of another image. Set bit 1 if
            the image is part of a multi-page image. Set bit 2 if the image
            is transparency mask for another image (photometric must be
            MASK, SamplesPerPixel and BitsPerSample must be 1).
        software : str
            Name of the software used to create the file. Must be 7-bit ASCII.
            Saved with the first page only.
        metadata : dict
            Additional metadata to be saved along with shape information
            in JSON or ImageJ formats in an ImageDescription tag.
            If None, do not write a second ImageDescription tag.
            Strings must be 7-bit ASCII. Saved with the first page only.
        ijmetadata : dict
            Additional metadata to be saved in application specific
            IJMetadata and IJMetadataByteCounts tags. Refer to the
            imagej_metadata_tag function for valid keys and values.
            Saved with the first page only.
        extratags : sequence of tuples
            Additional tags as [(code, dtype, count, value, writeonce)].

            code : int
                The TIFF tag Id.
            dtype : str
                Data type of items in 'value' in Python struct format.
                One of B, s, H, I, 2I, b, h, i, 2i, f, d, Q, or q.
            count : int
                Number of data values. Not used for string or byte string
                values.
            value : sequence
                'Count' values compatible with 'dtype'.
                Byte strings must contain count values of dtype packed as
                binary data.
            writeonce : bool
                If True, the tag is written to the first page only.

        """
        # TODO: refactor this function
        fh = self._fh
        byteorder = self._byteorder

        if data is None:
            if compress:
                raise ValueError('cannot save compressed empty file')
            datashape = shape
            datadtype = numpy.dtype(dtype).newbyteorder(byteorder)
            datadtypechar = datadtype.char
        else:
            data = numpy.asarray(data, byteorder+data.dtype.char, 'C')
            if data.size == 0:
                raise ValueError('cannot save empty array')
            datashape = data.shape
            datadtype = data.dtype
            datadtypechar = data.dtype.char

        returnoffset = returnoffset and datadtype.isnative
        bilevel = datadtypechar == '?'
        if bilevel:
            index = -1 if datashape[-1] > 1 else -2
            datasize = product(datashape[:index])
            if datashape[index] % 8:
                datasize *= datashape[index] // 8 + 1
            else:
                datasize *= datashape[index] // 8
        else:
            datasize = product(datashape) * datadtype.itemsize

        # just append contiguous data if possible
        self._truncate = bool(truncate)
        if self._datashape:
            if (not contiguous
                    or self._datashape[1:] != datashape
                    or self._datadtype != datadtype
                    or (compress and self._tags)
                    or tile
                    or not numpy.array_equal(colormap, self._colormap)):
                # incompatible shape, dtype, compression mode, or colormap
                self._write_remaining_pages()
                self._write_image_description()
                self._truncate = False
                self._descriptionoffset = 0
                self._descriptionlenoffset = 0
                self._datashape = None
                self._colormap = None
                if self._imagej:
                    raise ValueError(
                        'ImageJ does not support non-contiguous data')
            else:
                # consecutive mode
                self._datashape = (self._datashape[0] + 1,) + datashape
                if not compress:
                    # write contiguous data, write IFDs/tags later
                    offset = fh.tell()
                    if data is None:
                        fh.write_empty(datasize)
                    else:
                        fh.write_array(data)
                    if returnoffset:
                        return offset, datasize
                    return None

        input_shape = datashape
        tagnoformat = self._tagnoformat
        valueformat = self._valueformat
        offsetformat = self._offsetformat
        offsetsize = self._offsetsize
        tagsize = self._tagsize

        MINISBLACK = TIFF.PHOTOMETRIC.MINISBLACK
        MINISWHITE = TIFF.PHOTOMETRIC.MINISWHITE
        RGB = TIFF.PHOTOMETRIC.RGB
        CFA = TIFF.PHOTOMETRIC.CFA
        PALETTE = TIFF.PHOTOMETRIC.PALETTE
        CONTIG = TIFF.PLANARCONFIG.CONTIG
        SEPARATE = TIFF.PLANARCONFIG.SEPARATE

        # parse input
        if photometric is not None:
            photometric = enumarg(TIFF.PHOTOMETRIC, photometric)
        if planarconfig:
            planarconfig = enumarg(TIFF.PLANARCONFIG, planarconfig)
        if extrasamples is None:
            extrasamples_ = None
        else:
            extrasamples_ = tuple(enumarg(TIFF.EXTRASAMPLE, es)
                                  for es in sequence(extrasamples))
        if not compress:
            compress = False
            compresstag = 1
            # TODO: support predictors without compression
            predictor = False
            predictortag = 1
        else:
            if isinstance(compress, (tuple, list)):
                compress, compresslevel = compress
            elif isinstance(compress, int):
                compress, compresslevel = 'ADOBE_DEFLATE', int(compress)
                if not 0 <= compresslevel <= 9:
                    raise ValueError('invalid compression level %s' % compress)
            else:
                compresslevel = None
            compress = compress.upper()
            compresstag = enumarg(TIFF.COMPRESSION, compress)

        if predictor:
            if datadtype.kind in 'iu':
                predictortag = 2
                predictor = TIFF.PREDICTORS[2]
            elif datadtype.kind == 'f':
                predictortag = 3
                predictor = TIFF.PREDICTORS[3]
            else:
                raise ValueError('cannot apply predictor to %s' % datadtype)

        # prepare ImageJ format
        if self._imagej:
            # if predictor or compress:
            #     warnings.warn(
            #         'ImageJ cannot handle predictors or compression')
            if description:
                warnings.warn('not writing description to ImageJ file')
                description = None
            volume = False
            if datadtypechar not in 'BHhf':
                raise ValueError(
                    'ImageJ does not support data type %s' % datadtypechar)
            ijrgb = photometric == RGB if photometric else None
            if datadtypechar not in 'B':
                ijrgb = False
            ijshape = imagej_shape(datashape, ijrgb)
            if ijshape[-1] in (3, 4):
                photometric = RGB
                if datadtypechar not in 'B':
                    raise ValueError('ImageJ does not support data type %s '
                                     'for RGB' % datadtypechar)
            elif photometric is None:
                photometric = MINISBLACK
                planarconfig = None
            if planarconfig == SEPARATE:
                raise ValueError('ImageJ does not support planar images')
            planarconfig = CONTIG if ijrgb else None

        # define compress function
        if compress:
            compressor = TIFF.COMPESSORS[compresstag]
            if predictor:
                def compress(data, level=compresslevel):
                    data = predictor(data, axis=-2)
                    return compressor(data, level)
            else:
                def compress(data, level=compresslevel):
                    return compressor(data, level)

        # verify colormap and indices
        if colormap is not None:
            if datadtypechar not in 'BH':
                raise ValueError('invalid data dtype for palette mode')
            colormap = numpy.asarray(colormap, dtype=byteorder+'H')
            if colormap.shape != (3, 2**(datadtype.itemsize * 8)):
                raise ValueError('invalid color map shape')
            self._colormap = colormap

        # verify tile shape
        if tile:
            tile = tuple(int(i) for i in tile[:3])
            volume = len(tile) == 3
            if (len(tile) < 2 or tile[-1] % 16 or tile[-2] % 16 or
                    any(i < 1 for i in tile)):
                raise ValueError('invalid tile shape')
        else:
            tile = ()
            volume = False

        # normalize data shape to 5D or 6D, depending on volume:
        #   (pages, planar_samples, [depth,] height, width, contig_samples)
        datashape = reshape_nd(datashape, 3 if photometric == RGB else 2)
        shape = datashape
        ndim = len(datashape)

        samplesperpixel = 1
        extrasamples = 0
        if volume and ndim < 3:
            volume = False
        if colormap is not None:
            photometric = PALETTE
            planarconfig = None
        if photometric is None:
            photometric = MINISBLACK
            if bilevel:
                photometric = MINISWHITE
            elif planarconfig == CONTIG:
                if ndim > 2 and shape[-1] in (3, 4):
                    photometric = RGB
            elif planarconfig == SEPARATE:
                if volume and ndim > 3 and shape[-4] in (3, 4):
                    photometric = RGB
                elif ndim > 2 and shape[-3] in (3, 4):
                    photometric = RGB
            elif ndim > 2 and shape[-1] in (3, 4):
                photometric = RGB
            elif self._imagej:
                photometric = MINISBLACK
            elif volume and ndim > 3 and shape[-4] in (3, 4):
                photometric = RGB
            elif ndim > 2 and shape[-3] in (3, 4):
                photometric = RGB
        if planarconfig and len(shape) <= (3 if volume else 2):
            planarconfig = None
            if photometric not in (0, 1, 3, 4):
                photometric = MINISBLACK
        if photometric == RGB:
            if len(shape) < 3:
                raise ValueError('not a RGB(A) image')
            if len(shape) < 4:
                volume = False
            if planarconfig is None:
                if shape[-1] in (3, 4):
                    planarconfig = CONTIG
                elif shape[-4 if volume else -3] in (3, 4):
                    planarconfig = SEPARATE
                elif shape[-1] > shape[-4 if volume else -3]:
                    planarconfig = SEPARATE
                else:
                    planarconfig = CONTIG
            if planarconfig == CONTIG:
                datashape = (-1, 1) + shape[(-4 if volume else -3):]
                samplesperpixel = datashape[-1]
            else:
                datashape = (-1,) + shape[(-4 if volume else -3):] + (1,)
                samplesperpixel = datashape[1]
            if samplesperpixel > 3:
                extrasamples = samplesperpixel - 3
        elif photometric == CFA:
            if len(shape) != 2:
                raise ValueError('invalid CFA image')
            volume = False
            planarconfig = None
            datashape = (-1, 1) + shape[-2:] + (1,)
            if 50706 not in (et[0] for et in extratags):
                raise ValueError('must specify DNG tags for CFA image')
        elif planarconfig and len(shape) > (3 if volume else 2):
            if planarconfig == CONTIG:
                datashape = (-1, 1) + shape[(-4 if volume else -3):]
                samplesperpixel = datashape[-1]
            else:
                datashape = (-1,) + shape[(-4 if volume else -3):] + (1,)
                samplesperpixel = datashape[1]
            extrasamples = samplesperpixel - 1
        else:
            planarconfig = None
            while len(shape) > 2 and shape[-1] == 1:
                shape = shape[:-1]  # remove trailing 1s
            if len(shape) < 3:
                volume = False
            if extrasamples_ is None:
                datashape = (-1, 1) + shape[(-3 if volume else -2):] + (1,)
            else:
                datashape = (-1, 1) + shape[(-4 if volume else -3):]
                samplesperpixel = datashape[-1]
                extrasamples = samplesperpixel - 1

        if subfiletype & 0b100:
            # FILETYPE_MASK
            if not (bilevel and samplesperpixel == 1 and
                    photometric in (0, 1, 4)):
                raise ValueError('invalid SubfileType MASK')
            photometric = TIFF.PHOTOMETRIC.MASK

        # normalize shape to 6D
        assert len(datashape) in (5, 6)
        if len(datashape) == 5:
            datashape = datashape[:2] + (1,) + datashape[2:]
        if datashape[0] == -1:
            s0 = product(input_shape) // product(datashape[1:])
            datashape = (s0,) + datashape[1:]
        shape = datashape
        if data is not None:
            data = data.reshape(shape)

        if tile and not volume:
            tile = (1, tile[-2], tile[-1])

        if photometric == PALETTE:
            if (samplesperpixel != 1 or extrasamples or
                    shape[1] != 1 or shape[-1] != 1):
                raise ValueError('invalid data shape for palette mode')

        if photometric == RGB and samplesperpixel == 2:
            raise ValueError('not a RGB image (samplesperpixel=2)')

        if bilevel:
            if compresstag not in (1, 32773):
                raise ValueError('cannot compress bilevel image')
            if tile:
                raise ValueError('cannot save tiled bilevel image')
            if photometric not in (0, 1, 4):
                raise ValueError('cannot save bilevel image as %s' %
                                 str(photometric))
            datashape = list(datashape)
            if datashape[-2] % 8:
                datashape[-2] = datashape[-2] // 8 + 1
            else:
                datashape[-2] = datashape[-2] // 8
            datashape = tuple(datashape)
            assert datasize == product(datashape)
            if data is not None:
                data = numpy.packbits(data, axis=-2)
                assert datashape[-2] == data.shape[-2]

        bytestr = bytes if sys.version[0] == '2' else (
            lambda x: bytes(x, 'ascii') if isinstance(x, str) else x)
        tags = []  # list of (code, ifdentry, ifdvalue, writeonce)

        strip_or_tile = 'Tile' if tile else 'Strip'
        tagbytecounts = TIFF.TAG_NAMES[strip_or_tile + 'ByteCounts']
        tagoffsets = TIFF.TAG_NAMES[strip_or_tile + 'Offsets']
        self._tagoffsets = tagoffsets

        def pack(fmt, *val):
            return struct.pack(byteorder+fmt, *val)

        def addtag(code, dtype, count, value, writeonce=False):
            # Compute ifdentry & ifdvalue bytes from code, dtype, count, value
            # Append (code, ifdentry, ifdvalue, writeonce) to tags list
            code = int(TIFF.TAG_NAMES.get(code, code))
            try:
                tifftype = TIFF.DATA_DTYPES[dtype]
            except KeyError:
                raise ValueError('unknown dtype %s' % dtype)
            rawcount = count

            if dtype == 's':
                # strings
                value = bytestr(value) + b'\0'
                count = rawcount = len(value)
                rawcount = value.find(b'\0\0')
                if rawcount < 0:
                    rawcount = count
                else:
                    rawcount += 1  # length of string without buffer
                value = (value,)
            elif isinstance(value, bytes):
                # packed binary data
                dtsize = struct.calcsize(dtype)
                if len(value) % dtsize:
                    raise ValueError('invalid packed binary data')
                count = len(value) // dtsize
            if len(dtype) > 1:
                count *= int(dtype[:-1])
                dtype = dtype[-1]
            ifdentry = [pack('HH', code, tifftype),
                        pack(offsetformat, rawcount)]
            ifdvalue = None
            if struct.calcsize(dtype) * count <= offsetsize:
                # value(s) can be written directly
                if isinstance(value, bytes):
                    ifdentry.append(pack(valueformat, value))
                elif count == 1:
                    if isinstance(value, (tuple, list, numpy.ndarray)):
                        value = value[0]
                    ifdentry.append(pack(valueformat, pack(dtype, value)))
                else:
                    ifdentry.append(pack(valueformat,
                                         pack(str(count)+dtype, *value)))
            else:
                # use offset to value(s)
                ifdentry.append(pack(offsetformat, 0))
                if isinstance(value, bytes):
                    ifdvalue = value
                elif isinstance(value, numpy.ndarray):
                    assert value.size == count
                    assert value.dtype.char == dtype
                    ifdvalue = value.tostring()
                elif isinstance(value, (tuple, list)):
                    ifdvalue = pack(str(count)+dtype, *value)
                else:
                    ifdvalue = pack(dtype, value)
            tags.append((code, b''.join(ifdentry), ifdvalue, writeonce))

        def rational(arg, max_denominator=1000000):
            """"Return nominator and denominator from float or two integers."""
            from fractions import Fraction  # delayed import
            try:
                f = Fraction.from_float(arg)
            except TypeError:
                f = Fraction(arg[0], arg[1])
            f = f.limit_denominator(max_denominator)
            return f.numerator, f.denominator

        if description:
            # user provided description
            addtag('ImageDescription', 's', 0, description, writeonce=True)

        # write shape and metadata to ImageDescription
        self._metadata = {} if not metadata else metadata.copy()
        if self._imagej:
            description = imagej_description(
                input_shape, shape[-1] in (3, 4), self._colormap is not None,
                **self._metadata)
        elif metadata or metadata == {}:
            if self._truncate:
                self._metadata.update(truncated=True)
            description = json_description(input_shape, **self._metadata)
        # elif metadata is None and self._truncate:
        #     raise ValueError('cannot truncate without writing metadata')
        else:
            description = None
        if description:
            # add 64 bytes buffer
            # the image description might be updated later with the final shape
            description = str2bytes(description, 'ascii')
            description += b'\0' * 64
            self._descriptionlen = len(description)
            addtag('ImageDescription', 's', 0, description, writeonce=True)

        if software:
            addtag('Software', 's', 0, software, writeonce=True)
        if datetime:
            if isinstance(datetime, str):
                if len(datetime) != 19 or datetime[16] != ':':
                    raise ValueError('invalid datetime string')
            else:
                try:
                    datetime = datetime.strftime('%Y:%m:%d %H:%M:%S')
                except AttributeError:
                    datetime = self._now().strftime('%Y:%m:%d %H:%M:%S')
            addtag('DateTime', 's', 0, datetime, writeonce=True)
        addtag('Compression', 'H', 1, compresstag)
        if predictor:
            addtag('Predictor', 'H', 1, predictortag)
        addtag('ImageWidth', 'I', 1, shape[-2])
        addtag('ImageLength', 'I', 1, shape[-3])
        if tile:
            addtag('TileWidth', 'I', 1, tile[-1])
            addtag('TileLength', 'I', 1, tile[-2])
            if tile[0] > 1:
                addtag('ImageDepth', 'I', 1, shape[-4])
                addtag('TileDepth', 'I', 1, tile[0])
        addtag('NewSubfileType', 'I', 1, subfiletype)
        if not bilevel:
            sampleformat = {'u': 1, 'i': 2, 'f': 3, 'c': 6}[datadtype.kind]
            addtag('SampleFormat', 'H', samplesperpixel,
                   (sampleformat,) * samplesperpixel)
        addtag('PhotometricInterpretation', 'H', 1, photometric.value)
        if colormap is not None:
            addtag('ColorMap', 'H', colormap.size, colormap)
        addtag('SamplesPerPixel', 'H', 1, samplesperpixel)
        if bilevel:
            pass
        elif planarconfig and samplesperpixel > 1:
            addtag('PlanarConfiguration', 'H', 1, planarconfig.value)
            addtag('BitsPerSample', 'H', samplesperpixel,
                   (datadtype.itemsize * 8,) * samplesperpixel)
        else:
            addtag('BitsPerSample', 'H', 1, datadtype.itemsize * 8)
        if extrasamples:
            if extrasamples_ is not None:
                if extrasamples != len(extrasamples_):
                    raise ValueError('wrong number of extrasamples specified')
                addtag('ExtraSamples', 'H', extrasamples, extrasamples_)
            elif photometric == RGB and extrasamples == 1:
                # Unassociated alpha channel
                addtag('ExtraSamples', 'H', 1, 2)
            else:
                # Unspecified alpha channel
                addtag('ExtraSamples', 'H', extrasamples, (0,) * extrasamples)
        if resolution is not None:
            addtag('XResolution', '2I', 1, rational(resolution[0]))
            addtag('YResolution', '2I', 1, rational(resolution[1]))
            if len(resolution) > 2:
                unit = resolution[2]
                unit = 1 if unit is None else enumarg(TIFF.RESUNIT, unit)
            elif self._imagej:
                unit = 1
            else:
                unit = 2
            addtag('ResolutionUnit', 'H', 1, unit)
        elif not self._imagej:
            addtag('XResolution', '2I', 1, (1, 1))
            addtag('YResolution', '2I', 1, (1, 1))
            addtag('ResolutionUnit', 'H', 1, 1)
        if ijmetadata:
            for t in imagej_metadata_tag(ijmetadata, byteorder):
                addtag(*t)

        contiguous = not compress
        if tile:
            # one chunk per tile per plane
            tiles = ((shape[2] + tile[0] - 1) // tile[0],
                     (shape[3] + tile[1] - 1) // tile[1],
                     (shape[4] + tile[2] - 1) // tile[2])
            numtiles = product(tiles) * shape[1]
            databytecounts = [
                product(tile) * shape[-1] * datadtype.itemsize] * numtiles
            addtag(tagbytecounts, offsetformat, numtiles, databytecounts)
            addtag(tagoffsets, offsetformat, numtiles, [0] * numtiles)
            contiguous = contiguous and product(tiles) == 1
            if not contiguous:
                # allocate tile buffer
                chunk = numpy.empty(tile + (shape[-1],), dtype=datadtype)
        elif contiguous and (bilevel or rowsperstrip is None):
            # one strip per plane
            if bilevel:
                databytecounts = [product(datashape[2:])] * shape[1]
            else:
                databytecounts = [
                    product(datashape[2:]) * datadtype.itemsize] * shape[1]
            addtag(tagbytecounts, offsetformat, shape[1], databytecounts)
            addtag(tagoffsets, offsetformat, shape[1], [0] * shape[1])
            addtag('RowsPerStrip', 'I', 1, shape[-3])
        else:
            # use rowsperstrip
            rowsize = product(shape[-2:]) * datadtype.itemsize
            if rowsperstrip is None:
                # compress ~64 KB chunks by default
                rowsperstrip = 65536 // rowsize if compress else shape[-3]
            if rowsperstrip < 1:
                rowsperstrip = 1
            elif rowsperstrip > shape[-3]:
                rowsperstrip = shape[-3]
            addtag('RowsPerStrip', 'I', 1, rowsperstrip)

            numstrips1 = (shape[-3] + rowsperstrip - 1) // rowsperstrip
            numstrips = numstrips1 * shape[1]
            if compress:
                databytecounts = [0] * numstrips
            else:
                # TODO: save bilevel data with rowsperstrip
                stripsize = rowsperstrip * rowsize
                databytecounts = [stripsize] * numstrips
                stripsize -= rowsize * (numstrips1 * rowsperstrip - shape[-3])
                for i in range(numstrips1-1, numstrips, numstrips1):
                    databytecounts[i] = stripsize
            addtag(tagbytecounts, offsetformat, numstrips, databytecounts)
            addtag(tagoffsets, offsetformat, numstrips, [0] * numstrips)

        if data is None and not contiguous:
            raise ValueError('cannot write non-contiguous empty file')

        # add extra tags from user
        for t in extratags:
            addtag(*t)

        # TODO: check TIFFReadDirectoryCheckOrder warning in files containing
        #   multiple tags of same code
        # the entries in an IFD must be sorted in ascending order by tag code
        tags = sorted(tags, key=lambda x: x[0])

        fhpos = fh.tell()
        if not (self._bigtiff or self._imagej) and fhpos + datasize > 2**32-1:
            raise ValueError('data too large for standard TIFF file')

        # if not compressed or multi-tiled, write the first IFD and then
        # all data contiguously; else, write all IFDs and data interleaved
        for pageindex in range(1 if contiguous else shape[0]):

            ifdpos = fhpos
            if ifdpos % 2:
                # location of IFD must begin on a word boundary
                fh.write(b'\0')
                ifdpos += 1

            # update pointer at ifdoffset
            fh.seek(self._ifdoffset)
            fh.write(pack(offsetformat, ifdpos))
            fh.seek(ifdpos)

            # create IFD in memory
            if pageindex < 2:
                ifd = io.BytesIO()
                ifd.write(pack(tagnoformat, len(tags)))
                tagoffset = ifd.tell()
                ifd.write(b''.join(t[1] for t in tags))
                ifdoffset = ifd.tell()
                ifd.write(pack(offsetformat, 0))  # offset to next IFD
                # write tag values and patch offsets in ifdentries
                for tagindex, tag in enumerate(tags):
                    offset = tagoffset + tagindex * tagsize + offsetsize + 4
                    code = tag[0]
                    value = tag[2]
                    if value:
                        pos = ifd.tell()
                        if pos % 2:
                            # tag value is expected to begin on word boundary
                            ifd.write(b'\0')
                            pos += 1
                        ifd.seek(offset)
                        ifd.write(pack(offsetformat, ifdpos + pos))
                        ifd.seek(pos)
                        ifd.write(value)
                        if code == tagoffsets:
                            dataoffsetsoffset = offset, pos
                        elif code == tagbytecounts:
                            databytecountsoffset = offset, pos
                        elif code == 270 and value.endswith(b'\0\0\0\0'):
                            # image description buffer
                            self._descriptionoffset = ifdpos + pos
                            self._descriptionlenoffset = (
                                ifdpos + tagoffset + tagindex*tagsize + 4)
                    elif code == tagoffsets:
                        dataoffsetsoffset = offset, None
                    elif code == tagbytecounts:
                        databytecountsoffset = offset, None
                ifdsize = ifd.tell()
                if ifdsize % 2:
                    ifd.write(b'\0')
                    ifdsize += 1

            # write IFD later when strip/tile bytecounts and offsets are known
            fh.seek(ifdsize, 1)

            # write image data
            dataoffset = fh.tell()
            skip = align - dataoffset % align
            fh.seek(skip, 1)
            dataoffset += skip
            if contiguous:
                if data is None:
                    fh.write_empty(datasize)
                else:
                    fh.write_array(data)
            elif tile:
                if data is None:
                    fh.write_empty(numtiles * databytecounts[0])
                else:
                    stripindex = 0
                    for plane in data[pageindex]:
                        for tz in range(tiles[0]):
                            for ty in range(tiles[1]):
                                for tx in range(tiles[2]):
                                    c0 = min(tile[0], shape[2] - tz*tile[0])
                                    c1 = min(tile[1], shape[3] - ty*tile[1])
                                    c2 = min(tile[2], shape[4] - tx*tile[2])
                                    chunk[c0:, c1:, c2:] = 0
                                    chunk[:c0, :c1, :c2] = plane[
                                        tz*tile[0]:tz*tile[0]+c0,
                                        ty*tile[1]:ty*tile[1]+c1,
                                        tx*tile[2]:tx*tile[2]+c2]
                                    if compress:
                                        t = compress(chunk)
                                        fh.write(t)
                                        databytecounts[stripindex] = len(t)
                                        stripindex += 1
                                    else:
                                        fh.write_array(chunk)
                                        # fh.flush()
            elif compress:
                # write one strip per rowsperstrip
                assert data.shape[2] == 1  # not handling depth
                numstrips = (shape[-3] + rowsperstrip - 1) // rowsperstrip
                stripindex = 0
                for plane in data[pageindex]:
                    for i in range(numstrips):
                        strip = plane[0, i*rowsperstrip: (i+1)*rowsperstrip]
                        strip = compress(strip)
                        fh.write(strip)
                        databytecounts[stripindex] = len(strip)
                        stripindex += 1
            else:
                fh.write_array(data[pageindex])

            # update strip/tile offsets
            offset, pos = dataoffsetsoffset
            ifd.seek(offset)
            if pos:
                ifd.write(pack(offsetformat, ifdpos + pos))
                ifd.seek(pos)
                offset = dataoffset
                for size in databytecounts:
                    ifd.write(pack(offsetformat, offset))
                    offset += size
            else:
                ifd.write(pack(offsetformat, dataoffset))

            if compress:
                # update strip/tile bytecounts
                offset, pos = databytecountsoffset
                ifd.seek(offset)
                if pos:
                    ifd.write(pack(offsetformat, ifdpos + pos))
                    ifd.seek(pos)
                    for size in databytecounts:
                        ifd.write(pack(offsetformat, size))
                else:
                    ifd.write(pack(offsetformat, databytecounts[0]))

            fhpos = fh.tell()
            fh.seek(ifdpos)
            fh.write(iogetbuffer(ifd))
            fh.flush()
            fh.seek(fhpos)

            self._ifdoffset = ifdpos + ifdoffset

            # remove tags that should be written only once
            if pageindex == 0:
                tags = [tag for tag in tags if not tag[-1]]

        self._shape = shape
        self._datashape = (1,) + input_shape
        self._datadtype = datadtype
        self._dataoffset = dataoffset
        self._databytecounts = databytecounts

        if contiguous:
            # write remaining IFDs/tags later
            self._tags = tags
            # return offset and size of image data
            if returnoffset:
                return dataoffset, sum(databytecounts)
        return None

    def _write_remaining_pages(self):
        """Write outstanding IFDs and tags to file."""
        if not self._tags or self._truncate:
            return

        pageno = self._shape[0] * self._datashape[0] - 1
        if pageno < 1:
            self._tags = None
            self._datadtype = None
            self._dataoffset = None
            self._databytecounts = None
            return

        fh = self._fh
        fhpos = fh.tell()
        if fhpos % 2:
            fh.write(b'\0')
            fhpos += 1

        pack = struct.pack
        offsetformat = self._byteorder + self._offsetformat
        offsetsize = self._offsetsize
        tagnoformat = self._byteorder + self._tagnoformat
        tagsize = self._tagsize
        dataoffset = self._dataoffset
        pagedatasize = sum(self._databytecounts)

        # construct template IFD in memory
        # need to patch offsets to next IFD and data before writing to file
        ifd = io.BytesIO()
        ifd.write(pack(tagnoformat, len(self._tags)))
        tagoffset = ifd.tell()
        ifd.write(b''.join(t[1] for t in self._tags))
        ifdoffset = ifd.tell()
        ifd.write(pack(offsetformat, 0))  # offset to next IFD
        # tag values
        for tagindex, tag in enumerate(self._tags):
            offset = tagoffset + tagindex * tagsize + offsetsize + 4
            code = tag[0]
            value = tag[2]
            if value:
                pos = ifd.tell()
                if pos % 2:
                    # tag value is expected to begin on word boundary
                    ifd.write(b'\0')
                    pos += 1
                ifd.seek(offset)
                try:
                    ifd.write(pack(offsetformat, fhpos + pos))
                except Exception:  # struct.error
                    if self._imagej:
                        warnings.warn('truncating ImageJ file')
                        self._truncate = True
                        return
                    raise ValueError('data too large for non-BigTIFF file')
                ifd.seek(pos)
                ifd.write(value)
                if code == self._tagoffsets:
                    # save strip/tile offsets for later updates
                    dataoffsetsoffset = offset, pos
            elif code == self._tagoffsets:
                dataoffsetsoffset = offset, None

        ifdsize = ifd.tell()
        if ifdsize % 2:
            ifd.write(b'\0')
            ifdsize += 1

        # check if all IFDs fit in file
        if not self._bigtiff and fhpos + ifdsize * pageno > 2**32 - 32:
            if self._imagej:
                warnings.warn('truncating ImageJ file')
                self._truncate = True
                return
            raise ValueError('data too large for non-BigTIFF file')

        # assemble IFD chain in memory from IFD template
        ifds = io.BytesIO(bytes(ifdsize * pageno))
        ifdpos = fhpos
        for _ in range(pageno):
            # update strip/tile offsets in IFD
            dataoffset += pagedatasize  # offset to image data
            offset, pos = dataoffsetsoffset
            ifd.seek(offset)
            if pos:
                ifd.write(pack(offsetformat, ifdpos + pos))
                ifd.seek(pos)
                offset = dataoffset
                for size in self._databytecounts:
                    ifd.write(pack(offsetformat, offset))
                    offset += size
            else:
                ifd.write(pack(offsetformat, dataoffset))
            # update pointer at ifdoffset to point to next IFD in file
            ifdpos += ifdsize
            ifd.seek(ifdoffset)
            ifd.write(pack(offsetformat, ifdpos))
            # write IFD entry
            ifds.write(iogetbuffer(ifd))

        # terminate IFD chain
        ifdoffset += ifdsize * (pageno - 1)
        ifds.seek(ifdoffset)
        ifds.write(pack(offsetformat, 0))
        # write IFD chain to file
        fh.write(iogetbuffer(ifds))
        # update file to point to new IFD chain
        pos = fh.tell()
        fh.seek(self._ifdoffset)
        fh.write(pack(offsetformat, fhpos))
        fh.flush()
        fh.seek(pos)

        self._ifdoffset = fhpos + ifdoffset
        self._tags = None
        self._datadtype = None
        self._dataoffset = None
        self._databytecounts = None
        # do not reset _shape or _datashape

    def _write_image_description(self):
        """Write metadata to ImageDescription tag."""
        if (not self._datashape or self._datashape[0] == 1 or
                self._descriptionoffset <= 0):
            return

        colormapped = self._colormap is not None
        if self._imagej:
            isrgb = self._shape[-1] in (3, 4)
            description = imagej_description(
                self._datashape, isrgb, colormapped, **self._metadata)
        else:
            description = json_description(self._datashape, **self._metadata)

        # rewrite description and its length to file
        description = description.encode('utf-8')
        description = description[:self._descriptionlen-1]
        pos = self._fh.tell()
        self._fh.seek(self._descriptionoffset)
        self._fh.write(description)
        self._fh.seek(self._descriptionlenoffset)
        self._fh.write(struct.pack(self._byteorder+self._offsetformat,
                                   len(description)+1))
        self._fh.seek(pos)

        self._descriptionoffset = 0
        self._descriptionlenoffset = 0
        self._descriptionlen = 0

    def _now(self):
        """Return current date and time."""
        return datetime.datetime.now()

    def close(self):
        """Write remaining pages and close file handle."""
        if not self._truncate:
            self._write_remaining_pages()
        self._write_image_description()
        self._fh.close()

    def __enter__(self):
        return self

    def __exit__(self, exc_type, exc_value, traceback):
        self.close()


class TiffFile(object):
    """Read image and metadata from TIFF file.

    TiffFile instances must be closed using the 'close' method, which is
    automatically called when using the 'with' context manager.

    Attributes
    ----------
    pages : TiffPages
        Sequence of TIFF pages in file.
    series : list of TiffPageSeries
        Sequences of closely related TIFF pages. These are computed
        from OME, LSM, ImageJ, etc. metadata or based on similarity
        of page properties such as shape, dtype, and compression.
    is_flag : bool
        If True, file is of a certain format.
        Flags are: bigtiff, uniform, shaped, ome, imagej, stk, lsm, fluoview,
        nih, vista, micromanager, metaseries, mdgel, mediacy, tvips, fei,
        sem, scn, svs, scanimage, andor, epics, ndpi, pilatus, qpi.

    All attributes are read-only.

    """
    def __init__(self, arg, name=None, offset=None, size=None,
                 multifile=True, _useframes=None, **kwargs):
        """Initialize instance from file.

        Parameters
        ----------
        arg : str or open file
            Name of file or open file object.
            The file objects are closed in TiffFile.close().
        name : str
            Optional name of file in case 'arg' is a file handle.
        offset : int
            Optional start position of embedded file. By default, this is
            the current file position.
        size : int
            Optional size of embedded file. By default, this is the number
            of bytes from the 'offset' to the end of the file.
        multifile : bool
            If True (default), series may include pages from multiple files.
            Currently applies to OME-TIFF only.
        kwargs : bool
            'is_ome': If False, disable processing of OME-XML metadata.

        """
        if kwargs:
            for key in ('movie', 'fastij', 'multifile_close'):
                if key in kwargs:
                    del kwargs[key]
                    log.warning("TiffFile: the '%s' argument is ignored" % key)
            if 'pages' in kwargs:
                raise TypeError(
                    "the TiffFile 'pages' argument is no longer supported.\n\n"
                    "Use TiffFile.asarray(keys=[...]) to read image data "
                    "from specific pages.\n")

            for key, value in kwargs.items():
                if key[:3] == 'is_' and key[3:] in TIFF.FILE_FLAGS:
                    if value is not None and not value:
                        setattr(self, key, bool(value))
                else:
                    raise TypeError('unexpected keyword argument: %s' % key)

        fh = FileHandle(arg, mode='rb', name=name, offset=offset, size=size)
        self._fh = fh
        self._multifile = bool(multifile)
        self._files = {fh.name: self}  # cache of TiffFiles
        try:
            fh.seek(0)
            header = fh.read(4)
            try:
                byteorder = {b'II': '<', b'MM': '>'}[header[:2]]
            except KeyError:
                raise TiffFileError('not a TIFF file')

            version = struct.unpack(byteorder+'H', header[2:4])[0]
            if version == 43:
                # BigTiff
                offsetsize, zero = struct.unpack(byteorder+'HH', fh.read(4))
                if zero != 0 or offsetsize != 8:
                    raise TiffFileError('invalid BigTIFF file')
                if byteorder == '>':
                    self.tiff = TIFF.BIG_BE
                else:
                    self.tiff = TIFF.BIG_LE
            elif version == 42:
                # Classic TIFF
                if byteorder == '>':
                    self.tiff = TIFF.CLASSIC_BE
                elif kwargs.get('is_ndpi', False):
                    # NDPI uses 64 bit IFD offsets
                    # TODO: fix offsets in NDPI tags if file size > 4 GB
                    self.tiff = TIFF.NDPI_LE
                else:
                    self.tiff = TIFF.CLASSIC_LE
            else:
                raise TiffFileError('invalid TIFF file')

            # file handle is at offset to offset to first page
            self.pages = TiffPages(self)

            if self.is_lsm and (self.filehandle.size >= 2**32 or
                                self.pages[0].compression != 1 or
                                self.pages[1].compression != 1):
                self._lsm_load_pages()
            elif self.is_scanimage and (not self.is_bigtiff and
                                        self.filehandle.size >= 2**31):
                self.pages._load_virtual_frames()
            elif _useframes:
                self.pages.useframes = True

        except Exception:
            fh.close()
            raise

    @property
    def byteorder(self):
        return self.tiff.byteorder

    @property
    def is_bigtiff(self):
        return self.tiff.version == 43

    @property
    def filehandle(self):
        """Return file handle."""
        return self._fh

    @property
    def filename(self):
        """Return name of file handle."""
        return self._fh.name

    @lazyattr
    def fstat(self):
        """Return status of file handle as stat_result object."""
        try:
            return os.fstat(self._fh.fileno())
        except Exception:  # io.UnsupportedOperation
            return None

    def close(self):
        """Close open file handle(s)."""
        for tif in self._files.values():
            tif.filehandle.close()
        self._files = {}

    def asarray(self, key=None, series=None, out=None, validate=True,
                maxworkers=None):
        """Return image data from selected TIFF page(s) as numpy array.

        By default, the data from the first series is returned.

        Parameters
        ----------
        key : int, slice, or sequence of indices
            Defines which pages to return as array.
            If None (default), data from a series (default 0) is returned.
            If not None, data from the specified pages in the whole file
            (if 'series' is None) or a specified series are returned as a
            stacked array.
            Requesting an array from multiple pages that are not compatible
            wrt. shape, dtype, compression etc is undefined, i.e. may crash
            or return incorrect values.
        series : int or TiffPageSeries
            Defines which series of pages to return as array.
        out : numpy.ndarray, str, or file-like object
            Buffer where image data will be saved.
            If None (default), a new array will be created.
            If numpy.ndarray, a writable array of compatible dtype and shape.
            If 'memmap', directly memory-map the image data in the TIFF file
            if possible; else create a memory-mapped array in a temporary file.
            If str or open file, the file name or file object used to
            create a memory-map to an array stored in a binary file on disk.
        validate : bool
            If True (default), validate various tags.
            Passed to TiffPage.asarray().
        maxworkers : int or None
            Maximum number of threads to concurrently get data from pages
            or tiles. If None (default), mutli-threading is enabled if data
            are compressed. If 0, up to half the CPU cores are used.
            If 1, mutli-threading is disabled.
            Reading data from file is limited to a single thread.
            Using multiple threads can significantly speed up this function
            if the bottleneck is decoding compressed data, e.g. in case of
            large LZW compressed LSM files or JPEG compressed tiled slides.
            If the bottleneck is I/O or pure Python code, using multiple
            threads might be detrimental.

        Returns
        -------
        numpy.ndarray
            Image data from the specified pages.
            See the TiffPage.asarray function for operations that are
            applied (or not) to the raw data stored in the file.

        """
        if not self.pages:
            return numpy.array([])
        if key is None and series is None:
            series = 0
        if series is None:
            pages = self.pages
        else:
            try:
                series = self.series[series]
            except (KeyError, TypeError):
                pass
            pages = series.pages

        if key is None:
            pass
        elif series is None:
            pages = self.pages._getlist(key)
        elif isinstance(key, inttypes):
            pages = [pages[key]]
        elif isinstance(key, slice):
            pages = pages[key]
        elif isinstance(key, Iterable):
            pages = [pages[k] for k in key]
        else:
            raise TypeError('key must be an int, slice, or sequence')

        if not pages:
            raise ValueError('no pages selected')

        if key is None and series and series.offset:
            typecode = self.byteorder + series.dtype.char
            if pages[0].is_memmappable and (isinstance(out, str) and
                                            out == 'memmap'):
                # direct mapping
                result = self.filehandle.memmap_array(
                    typecode, series.shape, series.offset)
            else:
                # read into output
                if out is not None:
                    out = create_output(out, series.shape, series.dtype)
                self.filehandle.seek(series.offset)
                result = self.filehandle.read_array(
                    typecode, product(series.shape), out=out)
        elif len(pages) == 1:
            result = pages[0].asarray(out=out, validate=validate,
                                      maxworkers=maxworkers)
        else:
            result = stack_pages(pages, out=out, maxworkers=maxworkers)

        if result is None:
            return None

        if key is None:
            try:
                result.shape = series.shape
            except ValueError:
                try:
                    log.warning('TiffFile.asarray: failed to reshape %s to %s',
                                result.shape, series.shape)
                    # try series of expected shapes
                    result.shape = (-1,) + series.shape
                except ValueError:
                    # revert to generic shape
                    result.shape = (-1,) + pages[0].shape
        elif len(pages) == 1:
            result.shape = pages[0].shape
        else:
            result.shape = (-1,) + pages[0].shape
        return result

    @lazyattr
    def series(self):
        """Return related pages as TiffPageSeries.

        Side effect: after calling this function, TiffFile.pages might contain
        TiffPage and TiffFrame instances.

        """
        if not self.pages:
            return []

        useframes = self.pages.useframes
        keyframe = self.pages.keyframe.index
        series = []
        for name in ('lsm', 'ome', 'imagej', 'shaped', 'fluoview', 'sis',
                     'uniform', 'mdgel'):
            if getattr(self, 'is_' + name, False):
                series = getattr(self, '_series_' + name)()
                break
        self.pages.useframes = useframes
        self.pages.keyframe = keyframe
        if not series:
            series = self._series_generic()

        # remove empty series, e.g. in MD Gel files
        series = [s for s in series if product(s.shape) > 0]

        for i, s in enumerate(series):
            s.index = i
        return series

    def _series_generic(self):
        """Return image series in file.

        A series is a sequence of TiffPages with the same hash.

        """
        pages = self.pages
        pages._clear(False)
        pages.useframes = False
        if pages.cache:
            pages._load()

        result = []
        keys = []
        series = {}
        for page in pages:
            if not page.shape or product(page.shape) == 0:
                continue
            key = page.hash
            if key in series:
                series[key].append(page)
            else:
                keys.append(key)
                series[key] = [page]

        for key in keys:
            pages = series[key]
            page = pages[0]
            shape = page.shape
            axes = page.axes
            if len(pages) > 1:
                shape = (len(pages),) + shape
                axes = 'I' + axes
            result.append(TiffPageSeries(pages, shape, page.dtype, axes,
                                         kind='Generic'))

        self.is_uniform = len(result) == 1
        return result

    def _series_uniform(self):
        """Return all images in file as single series."""
        page = self.pages[0]
        shape = page.shape
        axes = page.axes
        dtype = page.dtype
        validate = not (page.is_scanimage or page.is_nih)
        pages = self.pages._getlist(validate=validate)
        lenpages = len(pages)
        if lenpages > 1:
            shape = (lenpages,) + shape
            axes = 'I' + axes
        if page.is_scanimage:
            kind = 'ScanImage'
        elif page.is_nih:
            kind = 'NIHImage'
        else:
            kind = 'Uniform'
        return [TiffPageSeries(pages, shape, dtype, axes, kind=kind)]

    def _series_shaped(self):
        """Return image series in "shaped" file."""
        pages = self.pages
        pages.useframes = True
        lenpages = len(pages)

        def append_series(series, pages, axes, shape, reshape, name,
                          truncated):
            page = pages[0]
            if not axes:
                shape = page.shape
                axes = page.axes
                if len(pages) > 1:
                    shape = (len(pages),) + shape
                    axes = 'Q' + axes
            size = product(shape)
            resize = product(reshape)
            if page.is_contiguous and resize > size and resize % size == 0:
                if truncated is None:
                    truncated = True
                axes = 'Q' + axes
                shape = (resize // size,) + shape
            try:
                axes = reshape_axes(axes, shape, reshape)
                shape = reshape
            except ValueError as exc:
                log.warning('Shaped series: %s: %s',
                            exc.__class__.__name__, exc)
            series.append(
                TiffPageSeries(pages, shape, page.dtype, axes, name=name,
                               kind='Shaped', truncated=truncated))

        keyframe = axes = shape = reshape = name = None
        series = []
        index = 0
        while True:
            if index >= lenpages:
                break
            # new keyframe; start of new series
            pages.keyframe = index
            keyframe = pages.keyframe
            if not keyframe.is_shaped:
                log.warning(
                    'Shaped series: invalid metadata or corrupted file')
                return None
            # read metadata
            axes = None
            shape = None
            metadata = json_description_metadata(keyframe.is_shaped)
            name = metadata.get('name', '')
            reshape = metadata['shape']
            truncated = metadata.get('truncated', None)
            if 'axes' in metadata:
                axes = metadata['axes']
                if len(axes) == len(reshape):
                    shape = reshape
                else:
                    axes = ''
                    log.warning('Shaped series: axes do not match shape')
            # skip pages if possible
            spages = [keyframe]
            size = product(reshape)
            npages, mod = divmod(size, product(keyframe.shape))
            if mod:
                log.warning(
                    'Shaped series: series shape does not match page shape')
                return None
            if 1 < npages <= lenpages - index:
                size *= keyframe._dtype.itemsize
                if truncated:
                    npages = 1
                elif (keyframe.is_final and
                      keyframe.offset + size < pages[index+1].offset):
                    truncated = False
                else:
                    # need to read all pages for series
                    truncated = False
                    for j in range(index+1, index+npages):
                        page = pages[j]
                        page.keyframe = keyframe
                        spages.append(page)
            append_series(series, spages, axes, shape, reshape, name,
                          truncated)
            index += npages

        self.is_uniform = len(series) == 1

        return series

    def _series_imagej(self):
        """Return image series in ImageJ file."""
        # ImageJ's dimension order is always TZCYXS
        # TODO: fix loading of color, composite, or palette images
        pages = self.pages
        pages.useframes = True
        pages.keyframe = 0
        page = pages[0]
        ij = self.imagej_metadata

        def is_hyperstack():
            # ImageJ hyperstack store all image metadata in the first page and
            # image data are stored contiguously before the second page, if any
            if not page.is_final:
                return False
            images = ij.get('images', 0)
            if images <= 1:
                return False
            offset, count = page.is_contiguous
            if (count != product(page.shape) * page.bitspersample // 8
                    or offset + count*images > self.filehandle.size):
                raise ValueError()
            # check that next page is stored after data
            if len(pages) > 1 and offset + count*images > pages[1].offset:
                return False
            return True

        try:
            hyperstack = is_hyperstack()
        except ValueError:
            log.warning('ImageJ series: invalid metadata or corrupted file')
            return None
        if hyperstack:
            # no need to read other pages
            pages = [page]
        else:
            pages = pages[:]

        shape = []
        axes = []
        if 'frames' in ij:
            shape.append(ij['frames'])
            axes.append('T')
        if 'slices' in ij:
            shape.append(ij['slices'])
            axes.append('Z')
        if 'channels' in ij and not (page.photometric == 2 and not
                                     ij.get('hyperstack', False)):
            shape.append(ij['channels'])
            axes.append('C')
        remain = ij.get('images', len(pages))//(product(shape) if shape else 1)
        if remain > 1:
            shape.append(remain)
            axes.append('I')
        if page.axes[0] == 'I':
            # contiguous multiple images
            shape.extend(page.shape[1:])
            axes.extend(page.axes[1:])
        elif page.axes[:2] == 'SI':
            # color-mapped contiguous multiple images
            shape = page.shape[0:1] + tuple(shape) + page.shape[2:]
            axes = list(page.axes[0]) + axes + list(page.axes[2:])
        else:
            shape.extend(page.shape)
            axes.extend(page.axes)

        truncated = (
            hyperstack and len(self.pages) == 1 and
            page.is_contiguous[1] != product(shape) * page.bitspersample // 8)

        self.is_uniform = True

        return [TiffPageSeries(pages, shape, page.dtype, axes, kind='ImageJ',
                               truncated=truncated)]

    def _series_fluoview(self):
        """Return image series in FluoView file."""
        pages = self.pages._getlist(validate=False)

        mm = self.fluoview_metadata
        mmhd = list(reversed(mm['Dimensions']))
        axes = ''.join(TIFF.MM_DIMENSIONS.get(i[0].upper(), 'Q')
                       for i in mmhd if i[1] > 1)
        shape = tuple(int(i[1]) for i in mmhd if i[1] > 1)
        self.is_uniform = True
        return [TiffPageSeries(pages, shape, pages[0].dtype, axes,
                               name=mm['ImageName'], kind='FluoView')]

    def _series_mdgel(self):
        """Return image series in MD Gel file."""
        # only a single page, scaled according to metadata in second page
        self.pages.useframes = False
        self.pages.keyframe = 0
        md = self.mdgel_metadata
        if md['FileTag'] in (2, 128):
            dtype = numpy.dtype('float32')
            scale = md['ScalePixel']
            scale = scale[0] / scale[1]  # rational
            if md['FileTag'] == 2:
                # squary root data format
                def transform(a):
                    return a.astype('float32')**2 * scale
            else:
                def transform(a):
                    return a.astype('float32') * scale
        else:
            transform = None
        page = self.pages[0]
        self.is_uniform = False
        return [TiffPageSeries([page], page.shape, dtype, page.axes,
                               transform=transform, kind='MDGel')]

    def _series_sis(self):
        """Return image series in Olympus SIS file."""
        pages = self.pages._getlist(validate=False)
        page = pages[0]
        lenpages = len(pages)
        md = self.sis_metadata
        if 'shape' in md and 'axes' in md:
            shape = md['shape'] + page.shape
            axes = md['axes'] + page.axes
        elif lenpages == 1:
            shape = page.shape
            axes = page.axes
        else:
            shape = (lenpages,) + page.shape
            axes = 'I' + page.axes
        self.is_uniform = True
        return [TiffPageSeries(pages, shape, page.dtype, axes, kind='SIS')]

    def _series_ome(self):
        """Return image series in OME-TIFF file(s)."""
        from xml.etree import cElementTree as etree  # delayed import
        omexml = self.pages[0].description
        try:
            root = etree.fromstring(omexml)
        except etree.ParseError as exc:
            # TODO: test badly encoded OME-XML
            log.warning('OME series: %s: %s', exc.__class__.__name__, exc)
            try:
                # might work on Python 2
                omexml = omexml.decode('utf-8', 'ignore').encode('utf-8')
                root = etree.fromstring(omexml)
            except Exception:
                return None

        self.pages.cache = True
        self.pages.useframes = True
        self.pages.keyframe = 0
        self.pages._load(keyframe=None)

        uuid = root.attrib.get('UUID', None)
        self._files = {uuid: self}
        dirname = self._fh.dirname
        modulo = {}
        series = []
        for element in root:
            if element.tag.endswith('BinaryOnly'):
                # TODO: load OME-XML from master or companion file
                log.warning('OME series: not an ome-tiff master file')
                break
            if element.tag.endswith('StructuredAnnotations'):
                for annot in element:
                    if not annot.attrib.get('Namespace',
                                            '').endswith('modulo'):
                        continue
                    for value in annot:
                        for modul in value:
                            for along in modul:
                                if not along.tag[:-1].endswith('Along'):
                                    continue
                                axis = along.tag[-1]
                                newaxis = along.attrib.get('Type', 'other')
                                newaxis = TIFF.AXES_LABELS[newaxis]
                                if 'Start' in along.attrib:
                                    step = float(along.attrib.get('Step', 1))
                                    start = float(along.attrib['Start'])
                                    stop = float(along.attrib['End']) + step
                                    labels = numpy.arange(start, stop, step)
                                else:
                                    labels = [label.text for label in along
                                              if label.tag.endswith('Label')]
                                modulo[axis] = (newaxis, labels)

            if not element.tag.endswith('Image'):
                continue

            attr = element.attrib
            name = attr.get('Name', None)

            for pixels in element:
                if not pixels.tag.endswith('Pixels'):
                    continue
                attr = pixels.attrib
                # dtype = attr.get('PixelType', None)
                axes = ''.join(reversed(attr['DimensionOrder']))
                shape = idxshape = list(int(attr['Size'+ax]) for ax in axes)
                size = product(shape[:-2])
                ifds = None
                spp = 1  # samples per pixel
                for data in pixels:
                    if data.tag.endswith('Channel'):
                        attr = data.attrib
                        if ifds is None:
                            spp = int(attr.get('SamplesPerPixel', spp))
                            ifds = [None] * (size // spp)
                            if spp > 1:
                                # correct channel dimension for spp
                                idxshape = list((shape[i] // spp if ax == 'C'
                                                 else shape[i])
                                                for i, ax in enumerate(axes))
                        elif int(attr.get('SamplesPerPixel', 1)) != spp:
                            raise ValueError(
                                'cannot handle differing SamplesPerPixel')
                        continue
                    if ifds is None:
                        ifds = [None] * (size // spp)
                    if not data.tag.endswith('TiffData'):
                        continue
                    attr = data.attrib
                    ifd = int(attr.get('IFD', 0))
                    num = int(attr.get('NumPlanes', 1 if 'IFD' in attr else 0))
                    num = int(attr.get('PlaneCount', num))
                    idx = [int(attr.get('First'+ax, 0)) for ax in axes[:-2]]
                    try:
                        idx = numpy.ravel_multi_index(idx, idxshape[:-2])
                    except ValueError:
                        # ImageJ produces invalid ome-xml when cropping
                        log.warning('OME series: invalid TiffData index')
                        continue
                    for uuid in data:
                        if not uuid.tag.endswith('UUID'):
                            continue
                        if uuid.text not in self._files:
                            if not self._multifile:
                                # abort reading multifile OME series
                                # and fall back to generic series
                                return []
                            fname = uuid.attrib['FileName']
                            try:
                                tif = TiffFile(os.path.join(dirname, fname))
                                tif.pages.cache = True
                                tif.pages.useframes = True
                                tif.pages.keyframe = 0
                                tif.pages._load(keyframe=None)
                            except (IOError, FileNotFoundError, ValueError):
                                log.warning("OME series: failed to read '%s'",
                                            fname)
                                break
                            self._files[uuid.text] = tif
                            tif.close()
                        pages = self._files[uuid.text].pages
                        try:
                            for i in range(num if num else len(pages)):
                                ifds[idx + i] = pages[ifd + i]
                        except IndexError:
                            log.warning('OME series: index out of range')
                        # only process first UUID
                        break
                    else:
                        pages = self.pages
                        try:
                            for i in range(num if num else
                                           min(len(pages), len(ifds))):
                                ifds[idx + i] = pages[ifd + i]
                        except IndexError:
                            log.warning('OME series: index out of range')

                if all(i is None for i in ifds):
                    # skip images without data
                    continue

                # find a keyframe
                keyframe = None
                for i in ifds:
                    # try find a TiffPage
                    if i and i == i.keyframe:
                        keyframe = i
                        break
                if keyframe is None:
                    # reload a TiffPage from file
                    for i, keyframe in enumerate(ifds):
                        if keyframe:
                            keyframe.parent.pages.keyframe = keyframe.index
                            keyframe = keyframe.parent.pages[keyframe.index]
                            ifds[i] = keyframe
                            break

                # move channel axis to match PlanarConfiguration storage
                # TODO: is this a bug or a inconsistency in the OME spec?
                if spp > 1:
                    if keyframe.planarconfig == 1 and axes[-1] != 'C':
                        i = axes.index('C')
                        axes = axes[:i] + axes[i+1:] + axes[i:i+1]
                        shape = shape[:i] + shape[i+1:] + shape[i:i+1]

                # FIXME: this implementation assumes the last dimensions are
                # stored in TIFF pages. Apparently that is not always the case.
                # For now, verify that shapes of keyframe and series match
                # If not, skip series.
                if keyframe.shape != tuple(shape[-len(keyframe.shape):]):
                    log.warning('OME series: incompatible page shape %s; '
                                'expected %s', keyframe.shape,
                                tuple(shape[-len(keyframe.shape):]))
                    del ifds
                    continue

                # set a keyframe on all IFDs
                for i in ifds:
                    if i is not None:
                        i.keyframe = keyframe

                series.append(
                    TiffPageSeries(ifds, shape, keyframe.dtype, axes,
                                   parent=self, name=name, kind='OME'))
                del ifds

        for serie in series:
            shape = list(serie.shape)
            for axis, (newaxis, labels) in modulo.items():
                i = serie.axes.index(axis)
                size = len(labels)
                if shape[i] == size:
                    serie.axes = serie.axes.replace(axis, newaxis, 1)
                else:
                    shape[i] //= size
                    shape.insert(i+1, size)
                    serie.axes = serie.axes.replace(axis, axis+newaxis, 1)
            serie.shape = tuple(shape)

        # squeeze dimensions
        for serie in series:
            serie.shape, serie.axes = squeeze_axes(serie.shape, serie.axes)
        self.is_uniform = len(series) == 1
        return series

    def _series_lsm(self):
        """Return main and thumbnail series in LSM file."""
        lsmi = self.lsm_metadata
        axes = TIFF.CZ_LSMINFO_SCANTYPE[lsmi['ScanType']]
        if self.pages[0].photometric == 2:  # RGB; more than one channel
            axes = axes.replace('C', '').replace('XY', 'XYC')
        if lsmi.get('DimensionP', 0) > 1:
            axes += 'P'
        if lsmi.get('DimensionM', 0) > 1:
            axes += 'M'
        axes = axes[::-1]
        shape = tuple(int(lsmi[TIFF.CZ_LSMINFO_DIMENSIONS[i]]) for i in axes)
        name = lsmi.get('Name', '')
        pages = self.pages._getlist(slice(0, None, 2), validate=False)
        dtype = pages[0].dtype
        series = [TiffPageSeries(pages, shape, dtype, axes, name=name,
                                 kind='LSM')]

        if self.pages[1].is_reduced:
            pages = self.pages._getlist(slice(1, None, 2), validate=False)
            dtype = pages[0].dtype
            cp = 1
            i = 0
            while cp < len(pages) and i < len(shape)-2:
                cp *= shape[i]
                i += 1
            shape = shape[:i] + pages[0].shape
            axes = axes[:i] + 'CYX'
            series.append(TiffPageSeries(pages, shape, dtype, axes, name=name,
                                         kind='LSMreduced'))

        self.is_uniform = False
        return series

    def _lsm_load_pages(self):
        """Load and fix all pages from LSM file."""
        # cache all pages to preserve corrected values
        pages = self.pages
        pages.cache = True
        pages.useframes = True
        # use first and second page as keyframes
        pages.keyframe = 1
        pages.keyframe = 0
        # load remaining pages as frames
        pages._load(keyframe=None)
        # fix offsets and bytecounts first
        self._lsm_fix_strip_offsets()
        self._lsm_fix_strip_bytecounts()
        # assign keyframes for data and thumbnail series
        keyframe = pages[0]
        for page in pages[::2]:
            page.keyframe = keyframe
        keyframe = pages[1]
        for page in pages[1::2]:
            page.keyframe = keyframe

    def _lsm_fix_strip_offsets(self):
        """Unwrap strip offsets for LSM files greater than 4 GB.

        Each series and position require separate unwrapping (undocumented).

        """
        if self.filehandle.size < 2**32:
            return

        pages = self.pages
        npages = len(pages)
        series = self.series[0]
        axes = series.axes

        # find positions
        positions = 1
        for i in 0, 1:
            if series.axes[i] in 'PM':
                positions *= series.shape[i]

        # make time axis first
        if positions > 1:
            ntimes = 0
            for i in 1, 2:
                if axes[i] == 'T':
                    ntimes = series.shape[i]
                    break
            if ntimes:
                div, mod = divmod(npages, 2*positions*ntimes)
                assert mod == 0
                shape = (positions, ntimes, div, 2)
                indices = numpy.arange(product(shape)).reshape(shape)
                indices = numpy.moveaxis(indices, 1, 0)
        else:
            indices = numpy.arange(npages).reshape(-1, 2)

        # images of reduced page might be stored first
        if pages[0]._offsetscounts[0][0] > pages[1]._offsetscounts[0][0]:
            indices = indices[..., ::-1]

        # unwrap offsets
        wrap = 0
        previousoffset = 0
        for i in indices.flat:
            page = pages[int(i)]
            dataoffsets = []
            for currentoffset in page._offsetscounts[0]:
                if currentoffset < previousoffset:
                    wrap += 2**32
                dataoffsets.append(currentoffset + wrap)
                previousoffset = currentoffset
            page._offsetscounts = dataoffsets, page._offsetscounts[1]

    def _lsm_fix_strip_bytecounts(self):
        """Set databytecounts to size of compressed data.

        The StripByteCounts tag in LSM files contains the number of bytes
        for the uncompressed data.

        """
        pages = self.pages
        if pages[0].compression == 1:
            return
        # sort pages by first strip offset
        pages = sorted(pages, key=lambda p: p._offsetscounts[0][0])
        npages = len(pages) - 1
        for i, page in enumerate(pages):
            if page.index % 2:
                continue
            offsets, bytecounts = page._offsetscounts
            if i < npages:
                lastoffset = pages[i+1]._offsetscounts[0][0]
            else:
                # LZW compressed strips might be longer than uncompressed
                lastoffset = min(offsets[-1] + 2*bytecounts[-1], self._fh.size)
            for j in range(len(bytecounts) - 1):
                bytecounts[j] = offsets[j+1] - offsets[j]
            bytecounts[-1] = lastoffset - offsets[-1]

    def __getattr__(self, name):
        """Return 'is_flag' attributes from first page."""
        if name[3:] in TIFF.FILE_FLAGS:
            if not self.pages:
                return False
            value = bool(getattr(self.pages[0], name))
            setattr(self, name, value)
            return value
        raise AttributeError("'%s' object has no attribute '%s'" %
                             (self.__class__.__name__, name))

    def __enter__(self):
        return self

    def __exit__(self, exc_type, exc_value, traceback):
        self.close()

    def __str__(self, detail=0, width=79):
        """Return string containing information about file.

        The detail parameter specifies the level of detail returned:

        0: file only.
        1: all series, first page of series and its tags.
        2: large tag values and file metadata.
        3: all pages.

        """
        info = [
            "TiffFile '%s'",
            format_size(self._fh.size),
            '' if byteorder_isnative(self.tiff.byteorder) else {
                '<': 'little-endian', '>': 'big-endian'}[self.tiff.byteorder]]
        if self.is_bigtiff:
            info.append('BigTiff')
        info.append(' '.join(f.lower() for f in self.flags))
        if len(self.pages) > 1:
            info.append('%i Pages' % len(self.pages))
        if len(self.series) > 1:
            info.append('%i Series' % len(self.series))
        if len(self._files) > 1:
            info.append('%i Files' % (len(self._files)))
        info = '  '.join(info)
        info = info.replace('    ', '  ').replace('   ', '  ')
        info = info % snipstr(self._fh.name, max(12, width+2-len(info)))
        if detail <= 0:
            return info
        info = [info]
        info.append('\n'.join(str(s) for s in self.series))
        if detail >= 3:
            info.extend((
                TiffPage.__str__(p, detail=detail, width=width)
                for p in self.pages
                if p is not None))
        elif self.series:
            info.extend((
                TiffPage.__str__(s.pages[0], detail=detail, width=width)
                for s in self.series
                if s.pages[0] is not None))
        elif self.pages and self.pages[0]:
            info.append(
                TiffPage.__str__(self.pages[0], detail=detail, width=width))
        if detail >= 2:
            for name in sorted(self.flags):
                if hasattr(self, name + '_metadata'):
                    m = getattr(self, name + '_metadata')
                    if m:
                        info.append(
                            '%s_METADATA\n%s' % (name.upper(),
                                                 pformat(m, width=width,
                                                         height=detail*12)))
        return '\n\n'.join(info).replace('\n\n\n', '\n\n')

    @lazyattr
    def flags(self):
        """Return set of file flags."""
        return set(name.lower() for name in sorted(TIFF.FILE_FLAGS)
                   if getattr(self, 'is_' + name))

    @lazyattr
    def is_mdgel(self):
        """File has MD Gel format."""
        # TODO: this likely reads the second page from file
        try:
            ismdgel = self.pages[0].is_mdgel or self.pages[1].is_mdgel
            if ismdgel:
                self.is_uniform = False
            return ismdgel
        except IndexError:
            return False

    @lazyattr
    def is_uniform(self):
        """Return if file contains a uniform series of pages."""
        # the hashes of IFDs 0, 7, and -1 are the same
        pages = self.pages
        page = pages[0]
        if page.is_scanimage or page.is_nih:
            return True
        try:
            useframes = pages.useframes
            pages.useframes = False
            h = page.hash
            for i in (1, 7, -1):
                if pages[i].aspage().hash != h:
                    return False
        except IndexError:
            return False
        finally:
            pages.useframes = useframes
        return True

    @property
    def is_appendable(self):
        """Return if pages can be appended to file without corrupting."""
        # TODO: check other formats
        return not (self.is_lsm or self.is_stk or self.is_imagej or
                    self.is_fluoview or self.is_micromanager)

    @lazyattr
    def shaped_metadata(self):
        """Return tifffile metadata from JSON descriptions as dicts."""
        if not self.is_shaped:
            return None
        return tuple(json_description_metadata(s.pages[0].is_shaped)
                     for s in self.series if s.kind.lower() == 'shaped')

    @property
    def ome_metadata(self):
        """Return OME XML."""
        if not self.is_ome:
            return None
        # return xml2dict(self.pages[0].description)['OME']
        return self.pages[0].description

    @property
    def lsm_metadata(self):
        """Return LSM metadata from CZ_LSMINFO tag as dict."""
        if not self.is_lsm:
            return None
        return self.pages[0].tags['CZ_LSMINFO'].value

    @lazyattr
    def stk_metadata(self):
        """Return STK metadata from UIC tags as dict."""
        if not self.is_stk:
            return None
        page = self.pages[0]
        tags = page.tags
        result = {}
        result['NumberPlanes'] = tags['UIC2tag'].count
        if page.description:
            result['PlaneDescriptions'] = page.description.split('\0')
            # result['plane_descriptions'] = stk_description_metadata(
            #    page.image_description)
        if 'UIC1tag' in tags:
            result.update(tags['UIC1tag'].value)
        if 'UIC3tag' in tags:
            result.update(tags['UIC3tag'].value)  # wavelengths
        if 'UIC4tag' in tags:
            result.update(tags['UIC4tag'].value)  # override uic1 tags
        uic2tag = tags['UIC2tag'].value
        result['ZDistance'] = uic2tag['ZDistance']
        result['TimeCreated'] = uic2tag['TimeCreated']
        result['TimeModified'] = uic2tag['TimeModified']
        try:
            result['DatetimeCreated'] = numpy.array(
                [julian_datetime(*dt) for dt in
                 zip(uic2tag['DateCreated'], uic2tag['TimeCreated'])],
                dtype='datetime64[ns]')
            result['DatetimeModified'] = numpy.array(
                [julian_datetime(*dt) for dt in
                 zip(uic2tag['DateModified'], uic2tag['TimeModified'])],
                dtype='datetime64[ns]')
        except ValueError as exc:
            log.warning('STK metadata: %s: %s', exc.__class__.__name__, exc)
        return result

    @lazyattr
    def imagej_metadata(self):
        """Return consolidated ImageJ metadata as dict."""
        if not self.is_imagej:
            return None
        page = self.pages[0]
        result = imagej_description_metadata(page.is_imagej)
        if 'IJMetadata' in page.tags:
            try:
                result.update(page.tags['IJMetadata'].value)
            except Exception:
                pass
        return result

    @lazyattr
    def fluoview_metadata(self):
        """Return consolidated FluoView metadata as dict."""
        if not self.is_fluoview:
            return None
        result = {}
        page = self.pages[0]
        result.update(page.tags['MM_Header'].value)
        # TODO: read stamps from all pages
        result['Stamp'] = page.tags['MM_Stamp'].value
        # skip parsing image description; not reliable
        # try:
        #     t = fluoview_description_metadata(page.image_description)
        #     if t is not None:
        #         result['ImageDescription'] = t
        # except Exception as exc:
        #     log.warning('FluoView metadata: '
        #                 'failed to parse image description (%s)', str(exc))
        return result

    @lazyattr
    def nih_metadata(self):
        """Return NIH Image metadata from NIHImageHeader tag as dict."""
        if not self.is_nih:
            return None
        return self.pages[0].tags['NIHImageHeader'].value

    @lazyattr
    def fei_metadata(self):
        """Return FEI metadata from SFEG or HELIOS tags as dict."""
        if not self.is_fei:
            return None
        tags = self.pages[0].tags
        if 'FEI_SFEG' in tags:
            return tags['FEI_SFEG'].value
        if 'FEI_HELIOS' in tags:
            return tags['FEI_HELIOS'].value
        return None

    @property
    def sem_metadata(self):
        """Return SEM metadata from CZ_SEM tag as dict."""
        if not self.is_sem:
            return None
        return self.pages[0].tags['CZ_SEM'].value

    @lazyattr
    def sis_metadata(self):
        """Return Olympus SIS metadata from SIS and INI tags as dict."""
        if not self.is_sis:
            return None
        tags = self.pages[0].tags
        result = {}
        try:
            result.update(tags['OlympusINI'].value)
        except Exception:
            pass
        try:
            result.update(tags['OlympusSIS'].value)
        except Exception:
            pass
        return result

    @lazyattr
    def mdgel_metadata(self):
        """Return consolidated metadata from MD GEL tags as dict."""
        for page in self.pages[:2]:
            if 'MDFileTag' in page.tags:
                tags = page.tags
                break
        else:
            return None
        result = {}
        for code in range(33445, 33453):
            name = TIFF.TAGS[code]
            if name not in tags:
                continue
            result[name[2:]] = tags[name].value
        return result

    @property
    def andor_metadata(self):
        """Return Andor tags as dict."""
        return self.pages[0].andor_tags

    @property
    def epics_metadata(self):
        """Return EPICS areaDetector tags as dict."""
        return self.pages[0].epics_tags

    @property
    def tvips_metadata(self):
        """Return TVIPS tag as dict."""
        if not self.is_tvips:
            return None
        return self.pages[0].tags['TVIPS'].value

    @lazyattr
    def metaseries_metadata(self):
        """Return MetaSeries metadata from image description as dict."""
        if not self.is_metaseries:
            return None
        return metaseries_description_metadata(self.pages[0].description)

    @lazyattr
    def pilatus_metadata(self):
        """Return Pilatus metadata from image description as dict."""
        if not self.is_pilatus:
            return None
        return pilatus_description_metadata(self.pages[0].description)

    @lazyattr
    def micromanager_metadata(self):
        """Return consolidated MicroManager metadata as dict."""
        if not self.is_micromanager:
            return None
        # from file header
        result = read_micromanager_metadata(self._fh)
        # from tag
        result.update(self.pages[0].tags['MicroManagerMetadata'].value)
        return result

    @lazyattr
    def scanimage_metadata(self):
        """Return ScanImage non-varying frame and ROI metadata as dict."""
        if not self.is_scanimage:
            return None
        result = {}
        try:
            framedata, roidata = read_scanimage_metadata(self._fh)
            result['FrameData'] = framedata
            result.update(roidata)
        except ValueError:
            pass
        # TODO: scanimage_artist_metadata
        try:
            result['Description'] = scanimage_description_metadata(
                self.pages[0].description)
        except Exception as exc:
            log.warning('ScanImage metadata: %s: %s',
                        exc.__class__.__name__, exc)
        return result

    @property
    def geotiff_metadata(self):
        """Return GeoTIFF metadata from first page as dict."""
        if not self.is_geotiff:
            return None
        return self.pages[0].geotiff_tags


class TiffPages(object):
    """Sequence of TIFF image file directories (IFD chain).

    Instances of TiffPages have a state (cache, keyframe, etc.) and are not
    thread-safe.

    """
    def __init__(self, parent):
        """Initialize instance and read first TiffPage from file.

        If parent is a TiffFile, the file position must be at an offset to an
        offset to a TiffPage. If parent is a TiffPage, page offsets are read
        from the SubIFDs tag.

        """
        self.parent = None
        self.pages = []  # cache of TiffPages, TiffFrames, or their offsets
        self._indexed = False  # True if offsets to all pages were read
        self._cached = False  # True if all pages were read into cache
        self._tiffpage = TiffPage  # class used for reading pages
        self._keyframe = None  # current page that is used as keyframe
        self._cache = False  # do not cache frames or pages (if not keyframe)
        self._nextpageoffset = None

        if isinstance(parent, TiffFile):
            # read offset to first page from current file position
            self.parent = parent
            fh = parent.filehandle
            self._nextpageoffset = fh.tell()
            offset = struct.unpack(parent.tiff.ifdoffsetformat,
                                   fh.read(parent.tiff.ifdoffsetsize))[0]
        elif 'SubIFDs' not in parent.tags:
            self._indexed = True
            return
        else:
            # use offsets from SubIFDs tag
            self.parent = parent.parent
            fh = self.parent.filehandle
            offsets = parent.tags['SubIFDs'].value
            offset = offsets[0]

        if offset == 0:
            log.warning('TiffPages: file contains no pages')
            self._indexed = True
            return
        if offset >= fh.size:
            log.warning('TiffPages: invalid page offset (%i)', offset)
            self._indexed = True
            return

        # read and cache first page
        fh.seek(offset)
        page = TiffPage(self.parent, index=0)
        self.pages.append(page)
        self._keyframe = page
        if self._nextpageoffset is None:
            # offsets from SubIFDs tag
            self.pages.extend(offsets[1:])
            self._indexed = True
            self._cached = True

    @property
    def cache(self):
        """Return if pages/frames are currently being cached."""
        return self._cache

    @cache.setter
    def cache(self, value):
        """Enable or disable caching of pages/frames. Clear cache if False."""
        value = bool(value)
        if self._cache and not value:
            self._clear()
        self._cache = value

    @property
    def useframes(self):
        """Return if currently using TiffFrame (True) or TiffPage (False)."""
        return self._tiffpage == TiffFrame and TiffFrame is not TiffPage

    @useframes.setter
    def useframes(self, value):
        """Set to use TiffFrame (True) or TiffPage (False)."""
        self._tiffpage = TiffFrame if value else TiffPage

    @property
    def keyframe(self):
        """Return current keyframe."""
        return self._keyframe

    @keyframe.setter
    def keyframe(self, index):
        """Set current keyframe. Load TiffPage from file if necessary."""
        index = int(index)
        if index < 0:
            index %= len(self)
        if self._keyframe.index == index:
            return
        if index == 0:
            self._keyframe = self.pages[0]
            return
        if self._indexed or index < len(self.pages):
            page = self.pages[index]
            if isinstance(page, TiffPage):
                self._keyframe = page
                return
            if isinstance(page, TiffFrame):
                # remove existing TiffFrame
                self.pages[index] = page.offset
        # load TiffPage from file
        tiffpage = self._tiffpage
        self._tiffpage = TiffPage
        try:
            self._keyframe = self._getitem(index)
        finally:
            self._tiffpage = tiffpage
        # always cache keyframes
        self.pages[index] = self._keyframe

    @property
    def next_page_offset(self):
        """Return offset where offset to a new page can be stored."""
        if not self._indexed:
            self._seek(-1)
        return self._nextpageoffset

    def _load(self, keyframe=True):
        """Read all remaining pages from file."""
        if self._cached:
            return
        pages = self.pages
        if not pages:
            return
        if not self._indexed:
            self._seek(-1)
        if not self._cache:
            return
        fh = self.parent.filehandle
        if keyframe is not None:
            keyframe = self._keyframe
        for i, page in enumerate(pages):
            if isinstance(page, inttypes):
                fh.seek(page)
                page = self._tiffpage(self.parent, index=i, keyframe=keyframe)
                pages[i] = page
        self._cached = True

    def _load_virtual_frames(self):
        """Calculate virtual TiffFrames."""
        pages = self.pages
        try:
            if sys.version_info[0] == 2:
                raise ValueError('not supported on Python 2')
            if len(pages) > 1:
                raise ValueError('pages already loaded')
            page = pages[0]
            bytecounts = page._offsetscounts[1]
            if len(bytecounts) != 1:
                raise ValueError('data not contiguous')
            self._seek(4)
            delta = pages[2] - pages[1]
            if pages[3] - pages[2] != delta or pages[4] - pages[3] != delta:
                raise ValueError('page offsets not equidistant')
            page1 = self._getitem(1, validate=page.hash)
            offsetoffset = page1._offsetscounts[0][0] - page1.offset
            if offsetoffset < 0 or offsetoffset > delta:
                raise ValueError('page offsets not equidistant')
            pages = [page, page1]
            filesize = self.parent.filehandle.size - delta
            for index, offset in enumerate(range(page1.offset+delta,
                                                 filesize, delta)):
                offsets = [offset + offsetoffset]
                offset = offset if offset < 2**31 else None
                pages.append(
                    TiffFrame(parent=page.parent, index=index+2, offset=None,
                              offsets=offsets, bytecounts=bytecounts,
                              keyframe=page))
        except Exception as exc:
            log.warning(
                'TiffPages: failed to load virtual frames: %s', str(exc))
        assert pages[1]
        self.pages = pages
        self._cache = True
        self._cached = True
        self._indexed = True

    def _clear(self, fully=True):
        """Delete all but first page from cache. Set keyframe to first page."""
        pages = self.pages
        if not pages:
            return
        self._keyframe = pages[0]
        if fully:
            # delete all but first TiffPage/TiffFrame
            for i, page in enumerate(pages[1:]):
                if not isinstance(page, inttypes) and page.offset is not None:
                    pages[i+1] = page.offset
        elif TiffFrame is not TiffPage:
            # delete only TiffFrames
            for i, page in enumerate(pages):
                if isinstance(page, TiffFrame) and page.offset is not None:
                    pages[i] = page.offset
        self._cached = False

    def _seek(self, index, maxpages=None):
        """Seek file to offset of page specified by index."""
        pages = self.pages
        lenpages = len(pages)
        if lenpages == 0:
            raise IndexError('index out of range')

        fh = self.parent.filehandle
        if fh.closed:
            raise ValueError('seek of closed file')

        if self._indexed or 0 <= index < lenpages:
            page = pages[index]
            offset = page if isinstance(page, inttypes) else page.offset
            fh.seek(offset)
            return

        tiff = self.parent.tiff
        offsetformat = tiff.ifdoffsetformat
        offsetsize = tiff.ifdoffsetsize
        tagnoformat = tiff.tagnoformat
        tagnosize = tiff.tagnosize
        tagsize = tiff.tagsize
        unpack = struct.unpack

        page = pages[-1]
        offset = page if isinstance(page, inttypes) else page.offset

        if maxpages is None:
            maxpages = 2**22
        while lenpages < maxpages:
            # read offsets to pages from file until index is reached
            fh.seek(offset)
            # skip tags
            try:
                tagno = unpack(tagnoformat, fh.read(tagnosize))[0]
                if tagno > 4096:
                    raise TiffFileError(
                        'suspicious number of tags: %i' % tagno)
            except Exception:
                log.warning('TiffPages: corrupted tag list of page %i @ %i',
                            lenpages, offset)
                del pages[-1]
                lenpages -= 1
                self._indexed = True
                break
            self._nextpageoffset = offset + tagnosize + tagno * tagsize
            fh.seek(self._nextpageoffset)

            # read offset to next page
            offset = unpack(offsetformat, fh.read(offsetsize))[0]
            if offset == 0:
                self._indexed = True
                break
            if offset >= fh.size:
                log.warning('TiffPages: invalid page offset (%i)', offset)
                self._indexed = True
                break

            pages.append(offset)
            lenpages += 1
            if 0 <= index < lenpages:
                break

            # detect some circular references
            if lenpages == 100:
                for p in pages[:-1]:
                    if offset == (p if isinstance(p, inttypes) else p.offset):
                        raise TiffFileError('invalid circular IFD reference')

        if index >= lenpages:
            raise IndexError('index out of range')

        page = pages[index]
        fh.seek(page if isinstance(page, inttypes) else page.offset)

    def _getlist(self, key=None, useframes=True, validate=True):
        """Return specified pages as list of TiffPages or TiffFrames.

        The first item is a TiffPage, and is used as a keyframe for
        following TiffFrames.

        """
        getitem = self._getitem
        _useframes = self.useframes

        if key is None:
            key = iter(range(len(self)))
        elif isinstance(key, Iterable):
            key = iter(key)
        elif isinstance(key, slice):
            start, stop, _ = key.indices(2**31-1)
            if not self._indexed and max(stop, start) > len(self.pages):
                self._seek(-1)
            key = iter(range(*key.indices(len(self.pages))))
        elif isinstance(key, inttypes):
            # return single TiffPage
            self.useframes = False
            if key == 0:
                return [self.pages[key]]
            try:
                return [getitem(key)]
            finally:
                self.useframes = _useframes
        else:
            raise TypeError('key must be an integer, slice, or iterable')

        # use first page as keyframe
        keyframe = self._keyframe
        self.keyframe = next(key)
        if validate:
            validate = self._keyframe.hash
        if useframes:
            self.useframes = True
        try:
            pages = [getitem(i, validate) for i in key]
            pages.insert(0, self._keyframe)
        finally:
            # restore state
            self._keyframe = keyframe
            if useframes:
                self.useframes = _useframes

        return pages

    def _getitem(self, key, validate=False):
        """Return specified page from cache or file."""
        key = int(key)
        pages = self.pages

        if key < 0:
            key %= len(self)
        elif self._indexed and key >= len(pages):
            raise IndexError('index out of range')

        if key < len(pages):
            page = pages[key]
            if self._cache:
                if not isinstance(page, inttypes):
                    if validate and validate != page.hash:
                        raise RuntimeError('page hash mismatch')
                    return page
            elif isinstance(page, (TiffPage, self._tiffpage)):
                if validate and validate != page.hash:
                    raise RuntimeError('page hash mismatch')
                return page

        self._seek(key)
        page = self._tiffpage(self.parent, index=key, keyframe=self._keyframe)
        if validate and validate != page.hash:
            raise RuntimeError('page hash mismatch')
        if self._cache:
            pages[key] = page
        return page

    def __getitem__(self, key):
        """Return specified page(s)."""
        pages = self.pages
        getitem = self._getitem

        if isinstance(key, inttypes):
            if key == 0:
                return pages[key]
            return getitem(key)

        if isinstance(key, slice):
            start, stop, _ = key.indices(2**31-1)
            if not self._indexed and max(stop, start) > len(pages):
                self._seek(-1)
            return [getitem(i) for i in range(*key.indices(len(pages)))]

        if isinstance(key, Iterable):
            return [getitem(k) for k in key]

        raise TypeError('key must be an integer, slice, or iterable')

    def __iter__(self):
        """Return iterator over all pages."""
        i = 0
        while True:
            try:
                yield self._getitem(i)
                i += 1
            except IndexError:
                break
        if self._cache:
            self._cached = True

    def __bool__(self):
        """Return True if file contains any pages."""
        return len(self.pages) > 0

    def __len__(self):
        """Return number of pages in file."""
        if not self._indexed:
            self._seek(-1)
        return len(self.pages)


class TiffPage(object):
    """TIFF image file directory (IFD).

    Attributes
    ----------
    index : int
        Index of page in file.
    dtype : numpy.dtype or None
        Data type (native byte order) of the image in IFD.
    shape : tuple
        Dimensions of the image in IFD.
    axes : str
        Axes label codes:
        'X' width, 'Y' height, 'S' sample, 'I' image series|page|plane,
        'Z' depth, 'C' color|em-wavelength|channel, 'E' ex-wavelength|lambda,
        'T' time, 'R' region|tile, 'A' angle, 'P' phase, 'H' lifetime,
        'L' exposure, 'V' event, 'Q' unknown, '_' missing
    tags : dict
        Dictionary of tags in IFD. {tag.name: TiffTag}
    colormap : numpy.ndarray
        Color look up table, if exists.

    All attributes are read-only.

    Notes
    -----
    The internal, normalized '_shape' attribute is 6 dimensional:

    0 : number planes/images  (stk, ij).
    1 : planar samplesperpixel.
    2 : imagedepth Z  (sgi).
    3 : imagelength Y.
    4 : imagewidth X.
    5 : contig samplesperpixel.

    """
    # default properties; will be updated from tags
    subfiletype = 0
    imagewidth = 0
    imagelength = 0
    imagedepth = 1
    tilewidth = 0
    tilelength = 0
    tiledepth = 1
    bitspersample = 1
    samplesperpixel = 1
    sampleformat = 1
    rowsperstrip = 2**32-1
    compression = 1
    planarconfig = 1
    fillorder = 1
    photometric = 0
    predictor = 1
    extrasamples = 1
    colormap = None
    software = ''
    description = ''
    description1 = ''

    def __init__(self, parent, index, keyframe=None):
        """Initialize instance from file.

        The file handle position must be at offset to a valid IFD.

        """
        self.parent = parent
        self.index = index
        self.shape = ()
        self._shape = ()
        self.dtype = None
        self._dtype = None
        self.axes = ''
        self.tags = tags = {}
        self.dataoffsets = ()
        self.databytecounts = ()

        tiff = parent.tiff

        # read TIFF IFD structure and its tags from file
        fh = parent.filehandle
        self.offset = fh.tell()  # offset to this IFD
        try:
            tagno = struct.unpack(
                tiff.tagnoformat, fh.read(tiff.tagnosize))[0]
            if tagno > 4096:
                raise TiffFileError('suspicious number of tags')
        except Exception:
            raise TiffFileError(
                'corrupted tag list at offset %i' % self.offset)

        tagoffset = self.offset + tiff.tagnosize  # fh.tell()
        tagsize = tiff.tagsize
        tagindex = -tagsize

        data = fh.read(tagsize * tagno)

        for _ in range(tagno):
            tagindex += tagsize
            try:
                tag = TiffTag(parent, data[tagindex:tagindex+tagsize],
                              tagoffset+tagindex)
            except TiffFileError as exc:
                log.warning('%s: %s', exc.__class__.__name__, exc)
                continue
            tagname = tag.name
            if tagname not in tags:
                name = tagname
                tags[name] = tag
            else:
                # some files contain multiple tags with same code
                # e.g. MicroManager files contain two ImageDescription tags
                i = 1
                while True:
                    name = '%s%i' % (tagname, i)
                    if name not in tags:
                        tags[name] = tag
                        break
            name = TIFF.TAG_ATTRIBUTES.get(name, '')
            if name:
                if name[:3] in 'sof des' and not isinstance(tag.value, str):
                    pass  # wrong string type for software, description
                else:
                    setattr(self, name, tag.value)

        if not tags:
            return  # found in FIBICS

        if 'SubfileType' in tags and self.subfiletype == 0:
            sft = tags['SubfileType'].value
            if sft == 2:
                self.subfiletype = 0b1  # reduced image
            elif sft == 3:
                self.subfiletype = 0b10  # multi-page

        # consolidate private tags; remove them from self.tags
        if self.is_andor:
            self.andor_tags
        elif self.is_epics:
            self.epics_tags
        # elif self.is_ndpi:
        #     self.ndpi_tags

        if self.is_sis and 'GPSTag' in tags:
            # TODO: can't change tag.name
            tags['OlympusSIS2'] = tags['GPSTag']
            del tags['GPSTag']

        if self.is_lsm or (self.index and self.parent.is_lsm):
            # correct non standard LSM bitspersample tags
            tags['BitsPerSample']._fix_lsm_bitspersample(self)
            if self.compression == 1 and self.predictor != 1:
                # work around bug in LSM510 software
                self.predictor = 1

        if self.is_vista or (self.index and self.parent.is_vista):
            # ISS Vista writes wrong ImageDepth tag
            self.imagedepth = 1

        if self.is_stk and 'UIC1tag' in tags and not tags['UIC1tag'].value:
            # read UIC1tag now that plane count is known
            uic1tag = tags['UIC1tag']
            fh.seek(uic1tag.valueoffset)
            tags['UIC1tag'].value = read_uic1tag(
                fh, tiff.byteorder, uic1tag.dtype,
                uic1tag.count, None, tags['UIC2tag'].count)

        if 'IJMetadata' in tags:
            # decode IJMetadata tag
            try:
                tags['IJMetadata'].value = imagej_metadata(
                    tags['IJMetadata'].value,
                    tags['IJMetadataByteCounts'].value,
                    tiff.byteorder)
            except Exception as exc:
                log.warning('TiffPage: %s: %s', exc.__class__.__name__, exc)

        if 'BitsPerSample' in tags:
            tag = tags['BitsPerSample']
            if tag.count == 1:
                self.bitspersample = tag.value
            else:
                # LSM might list more items than samplesperpixel
                value = tag.value[:self.samplesperpixel]
                if any((v-value[0] for v in value)):
                    self.bitspersample = value
                else:
                    self.bitspersample = value[0]

        if 'SampleFormat' in tags:
            tag = tags['SampleFormat']
            if tag.count == 1:
                self.sampleformat = tag.value
            else:
                value = tag.value[:self.samplesperpixel]
                if any((v-value[0] for v in value)):
                    self.sampleformat = value
                else:
                    self.sampleformat = value[0]

        if 'TileWidth' in tags:
            self.rowsperstrip = None
        elif 'ImageLength' in tags:
            if 'RowsPerStrip' not in tags or tags['RowsPerStrip'].count > 1:
                self.rowsperstrip = self.imagelength
            self.rowsperstrip = min(self.rowsperstrip, self.imagelength)
            # self.stripsperimage = int(math.floor(
            #    float(self.imagelength + self.rowsperstrip - 1) /
            #    self.rowsperstrip))

        # determine dtype
        dtype = self.sampleformat, self.bitspersample
        dtype = TIFF.SAMPLE_DTYPES.get(dtype, None)
        if dtype is not None:
            dtype = numpy.dtype(dtype)
        self.dtype = self._dtype = dtype

        # determine shape of data
        imagelength = self.imagelength
        imagewidth = self.imagewidth
        imagedepth = self.imagedepth
        samplesperpixel = self.samplesperpixel

        if self.is_stk:
            assert self.imagedepth == 1
            uictag = tags['UIC2tag'].value
            planes = tags['UIC2tag'].count
            if self.planarconfig == 1:
                self._shape = (
                    planes, 1, 1, imagelength, imagewidth, samplesperpixel)
                if samplesperpixel == 1:
                    self.shape = (planes, imagelength, imagewidth)
                    self.axes = 'YX'
                else:
                    self.shape = (
                        planes, imagelength, imagewidth, samplesperpixel)
                    self.axes = 'YXS'
            else:
                self._shape = (
                    planes, samplesperpixel, 1, imagelength, imagewidth, 1)
                if samplesperpixel == 1:
                    self.shape = (planes, imagelength, imagewidth)
                    self.axes = 'YX'
                else:
                    self.shape = (
                        planes, samplesperpixel, imagelength, imagewidth)
                    self.axes = 'SYX'
            # detect type of series
            if planes == 1:
                self.shape = self.shape[1:]
            elif numpy.all(uictag['ZDistance'] != 0):
                self.axes = 'Z' + self.axes
            elif numpy.all(numpy.diff(uictag['TimeCreated']) != 0):
                self.axes = 'T' + self.axes
            else:
                self.axes = 'I' + self.axes
        elif self.photometric == 2 or samplesperpixel > 1:  # PHOTOMETRIC.RGB
            if self.planarconfig == 1:
                self._shape = (
                    1, 1, imagedepth, imagelength, imagewidth, samplesperpixel)
                if imagedepth == 1:
                    self.shape = (imagelength, imagewidth, samplesperpixel)
                    self.axes = 'YXS'
                else:
                    self.shape = (
                        imagedepth, imagelength, imagewidth, samplesperpixel)
                    self.axes = 'ZYXS'
            else:
                self._shape = (1, samplesperpixel, imagedepth,
                               imagelength, imagewidth, 1)
                if imagedepth == 1:
                    self.shape = (samplesperpixel, imagelength, imagewidth)
                    self.axes = 'SYX'
                else:
                    self.shape = (
                        samplesperpixel, imagedepth, imagelength, imagewidth)
                    self.axes = 'SZYX'
        else:
            self._shape = (1, 1, imagedepth, imagelength, imagewidth, 1)
            if imagedepth == 1:
                self.shape = (imagelength, imagewidth)
                self.axes = 'YX'
            else:
                self.shape = (imagedepth, imagelength, imagewidth)
                self.axes = 'ZYX'

        # dataoffsets and databytecounts
        if 'TileOffsets' in tags:
            self.dataoffsets = tags['TileOffsets'].value
        elif 'StripOffsets' in tags:
            self.dataoffsets = tags['StripOffsets'].value

        if 'TileByteCounts' in tags:
            self.databytecounts = tags['TileByteCounts'].value
        elif 'StripByteCounts' in tags:
            self.databytecounts = tags['StripByteCounts'].value
        else:
            self.databytecounts = (
                product(self.shape) * (self.bitspersample // 8),)
            if self.compression != 1:
                log.warning('TiffPage: ByteCounts tag is missing')

        # assert len(self.shape) == len(self.axes)

    def asarray(self, out=None, squeeze=True, lock=None, reopen=True,
                maxsize=None, maxworkers=None, validate=True):
        """Read image data from file and return as numpy array.

        Raise ValueError if format is unsupported.

        Parameters
        ----------
        out : numpy.ndarray, str, or file-like object
            Buffer where image data will be saved.
            If None (default), a new array will be created.
            If numpy.ndarray, a writable array of compatible dtype and shape.
            If 'memmap', directly memory-map the image data in the TIFF file
            if possible; else create a memory-mapped array in a temporary file.
            If str or open file, the file name or file object used to
            create a memory-map to an array stored in a binary file on disk.
        squeeze : bool
            If True (default), all length-1 dimensions (except X and Y) are
            squeezed out from the array.
            If False, the shape of the returned array might be different from
            the page.shape.
        lock : {RLock, NullContext}
            A reentrant lock used to syncronize reads from file.
            If None (default), the lock of the parent's filehandle is used.
        reopen : bool
            If True (default) and the parent file handle is closed, the file
            is temporarily re-opened and closed if no exception occurs.
        maxsize: int
            Maximum size of data before a ValueError is raised.
            Can be used to catch DOS. Default: 16 TB.
        maxworkers : int or None
            Maximum number of threads to concurrently decode tile data.
            If None (default), up to half the CPU cores are used for
            compressed tiles.
            See remarks in TiffFile.asarray.
        validate : bool
            If True (default), validate various parameters.
            If None, only validate parameters and return None.

        Returns
        -------
        numpy.ndarray
            Numpy array of decompressed, depredicted, and unpacked image data
            read from Strip/Tile Offsets/ByteCounts, formatted according to
            shape and dtype metadata found in tags and parameters.
            Photometric conversion, pre-multiplied alpha, orientation, and
            colorimetry corrections are not applied. Specifically, CMYK images
            are not converted to RGB, MinIsWhite images are not inverted,
            and color palettes are not applied.

        """
        # properties from TiffPage or TiffFrame
        fh = self.parent.filehandle
        byteorder = self.parent.tiff.byteorder
        offsets, bytecounts = self._offsetscounts
        self_ = self
        self = self.keyframe  # self or keyframe

        if not self._shape or product(self._shape) == 0:
            return None

        tags = self.tags

        if validate or validate is None:
            if maxsize is None:
                maxsize = 2**44
            if maxsize and product(self._shape) > maxsize:
                raise ValueError('data are too large %s' % str(self._shape))
            if self.dtype is None:
                raise ValueError('data type not supported: %s%i' % (
                    self.sampleformat, self.bitspersample))
            if self.compression not in TIFF.DECOMPESSORS:
                raise ValueError(
                    'cannot decompress %s' % self.compression.name)
            if 'SampleFormat' in tags:
                tag = tags['SampleFormat']
                if tag.count != 1 and any((i-tag.value[0] for i in tag.value)):
                    raise ValueError(
                        'sample formats do not match %s' % tag.value)
            if self.is_subsampled and (self.compression not in (6, 7) or
                                       self.planarconfig == 2):
                raise NotImplementedError('chroma subsampling not supported')
            if validate is None:
                return None

        lock = fh.lock if lock is None else lock
        with lock:
            closed = fh.closed
            if closed:
                if reopen:
                    fh.open()
                else:
                    raise IOError('file handle is closed')

        dtype = self._dtype
        shape = self._shape
        imagewidth = self.imagewidth
        imagelength = self.imagelength
        imagedepth = self.imagedepth
        bitspersample = self.bitspersample
        typecode = byteorder + dtype.char
        lsb2msb = self.fillorder == 2
        istiled = self.is_tiled

        if istiled:
            tilewidth = self.tilewidth
            tilelength = self.tilelength
            tiledepth = self.tiledepth
            tw = (imagewidth + tilewidth - 1) // tilewidth
            tl = (imagelength + tilelength - 1) // tilelength
            td = (imagedepth + tiledepth - 1) // tiledepth
            tiledshape = (td, tl, tw)
            tileshape = (tiledepth, tilelength, tilewidth, shape[-1])
            runlen = tilewidth
        else:
            runlen = imagewidth

        if self.planarconfig == 1:
            runlen *= self.samplesperpixel

        if isinstance(out, str) and out == 'memmap' and self.is_memmappable:
            # direct memory map array in file
            with lock:
                result = fh.memmap_array(typecode, shape, offset=offsets[0])
        elif self.is_contiguous:
            # read contiguous bytes to array
            if out is not None:
                out = create_output(out, shape, dtype)
            with lock:
                fh.seek(offsets[0])
                result = fh.read_array(typecode, product(shape), out=out)
            if lsb2msb:
                bitorder_decode(result, out=result)
        else:
            # decompress, unpack,... individual strips or tiles
            result = create_output(out, shape, dtype)

            decompress = TIFF.DECOMPESSORS[self.compression]

            if self.compression in (6, 7):  # COMPRESSION.JPEG
                colorspace = None
                outcolorspace = None
                jpegtables = None
                if lsb2msb:
                    log.warning('TiffPage.asarray: disabling LSB2MSB for JPEG')
                    lsb2msb = False
                if 'JPEGTables' in tags:
                    # load JPEGTables from TiffFrame
                    jpegtables = self_._gettags({347}, lock=lock)[0][1].value
                # TODO: obtain table from OJPEG tags
                # elif ('JPEGInterchangeFormat' in tags and
                #       'JPEGInterchangeFormatLength' in tags and
                #       tags['JPEGInterchangeFormat'].value != offsets[0]):
                #     fh.seek(tags['JPEGInterchangeFormat'].value)
                #     fh.read(tags['JPEGInterchangeFormatLength'].value)
                if 'ExtraSamples' in tags:
                    pass
                elif self.photometric == 6:
                    # YCBCR -> RGB
                    outcolorspace = 'RGB'
                elif self.photometric == 2:
                    if self.planarconfig == 2:
                        # TODO: decode JPEG to planar RGB
                        raise NotImplementedError(
                            'cannot decode JPEG to planar RGB')
                    colorspace = outcolorspace = 'RGB'
                else:
                    outcolorspace = TIFF.PHOTOMETRIC(self.photometric).name
                if istiled:
                    heightwidth = tilelength, tilewidth
                else:
                    heightwidth = imagelength, imagewidth

                def decompress(data, bitspersample=bitspersample,
                               jpegtables=jpegtables, colorspace=colorspace,
                               outcolorspace=outcolorspace, shape=heightwidth,
                               out=None, _decompress=decompress):
                    return _decompress(data, bitspersample, jpegtables,
                                       colorspace, outcolorspace, shape, out)

                def unpack(data):
                    return data.reshape(-1)

            elif bitspersample in (8, 16, 32, 64, 128):
                if (bitspersample * runlen) % 8:
                    raise ValueError('data and sample size mismatch')
                if self.predictor == 3:  # PREDICTOR.FLOATINGPOINT
                    # the floating-point horizontal differencing decoder
                    # needs the raw byte order
                    typecode = dtype.char

                def unpack(data, typecode=typecode, out=None):
                    try:
                        # read only numpy array
                        return numpy.frombuffer(data, typecode)
                    except ValueError:
                        # strips may be missing EOI
                        # log.warning('TiffPage.asarray: ...')
                        bps = bitspersample // 8
                        xlen = (len(data) // bps) * bps
                        return numpy.frombuffer(data[:xlen], typecode)

            elif isinstance(bitspersample, tuple):

                def unpack(data, out=None):
                    return unpack_rgb(data, typecode, bitspersample)

            else:

                def unpack(data, out=None):
                    return packints_decode(data, typecode, bitspersample,
                                           runlen)

            # TODO: store decode function for future use
            # TODO: unify tile and strip decoding
            if istiled:
                unpredict = TIFF.UNPREDICTORS[self.predictor]

                def decode(tile, tileindex):
                    return tile_decode(tile, tileindex, tileshape, tiledshape,
                                       lsb2msb, decompress, unpack, unpredict,
                                       result[0])

                tileiter = buffered_read(fh, lock, offsets, bytecounts)
                if maxworkers is None:
                    maxworkers = 0 if self.compression > 1 else 1
                if maxworkers == 0:
                    import multiprocessing  # noqa: delay import
                    maxworkers = multiprocessing.cpu_count() // 2
                if maxworkers < 2:
                    for i, tile in enumerate(tileiter):
                        decode(tile, i)
                else:
                    # decode first tile un-threaded to catch exceptions
                    decode(next(tileiter), 0)
                    with ThreadPoolExecutor(maxworkers) as executor:
                        executor.map(decode, tileiter, range(1, len(offsets)))

            else:
                stripsize = self.rowsperstrip * self.imagewidth
                if self.planarconfig == 1:
                    stripsize *= self.samplesperpixel
                outsize = stripsize * self.dtype.itemsize
                result = result.reshape(-1)
                index = 0
                for strip in buffered_read(fh, lock, offsets, bytecounts):
                    if lsb2msb:
                        strip = bitorder_decode(strip, out=strip)
                    strip = decompress(strip, out=outsize)
                    strip = unpack(strip)
                    size = min(result.size, strip.size, stripsize,
                               result.size - index)
                    result[index:index+size] = strip[:size]
                    del strip
                    index += size

        result.shape = self._shape

        if self.predictor != 1 and not (istiled and not self.is_contiguous):
            unpredict = TIFF.UNPREDICTORS[self.predictor]
            result = unpredict(result, axis=-2, out=result)

        if squeeze:
            try:
                result.shape = self.shape
            except ValueError:
                log.warning('TiffPage.asarray: failed to reshape %s to %s',
                            result.shape, self.shape)

        if closed:
            # TODO: file should remain open if an exception occurred above
            fh.close()
        return result

    def asrgb(self, uint8=False, alpha=None, colormap=None,
              dmin=None, dmax=None, **kwargs):
        """Return image data as RGB(A).

        Work in progress.

        """
        data = self.asarray(**kwargs)
        self = self.keyframe  # self or keyframe
        photometric = self.photometric
        PHOTOMETRIC = TIFF.PHOTOMETRIC

        if photometric == PHOTOMETRIC.PALETTE:
            colormap = self.colormap
            if (colormap.shape[1] < 2**self.bitspersample or
                    self.dtype.char not in 'BH'):
                raise ValueError('cannot apply colormap')
            if uint8:
                if colormap.max() > 255:
                    colormap >>= 8
                colormap = colormap.astype('uint8')
            if 'S' in self.axes:
                data = data[..., 0] if self.planarconfig == 1 else data[0]
            data = apply_colormap(data, colormap)

        elif photometric == PHOTOMETRIC.RGB:
            if 'ExtraSamples' in self.tags:
                if alpha is None:
                    alpha = TIFF.EXTRASAMPLE
                extrasamples = self.extrasamples
                if self.tags['ExtraSamples'].count == 1:
                    extrasamples = (extrasamples,)
                for i, exs in enumerate(extrasamples):
                    if exs in alpha:
                        if self.planarconfig == 1:
                            data = data[..., [0, 1, 2, 3+i]]
                        else:
                            data = data[:, [0, 1, 2, 3+i]]
                        break
            else:
                if self.planarconfig == 1:
                    data = data[..., :3]
                else:
                    data = data[:, :3]
            # TODO: convert to uint8?

        elif photometric == PHOTOMETRIC.MINISBLACK:
            raise NotImplementedError()
        elif photometric == PHOTOMETRIC.MINISWHITE:
            raise NotImplementedError()
        elif photometric == PHOTOMETRIC.SEPARATED:
            raise NotImplementedError()
        else:
            raise NotImplementedError()
        return data

    def _gettags(self, codes=None, lock=None):
        """Return list of (code, TiffTag)."""
        tags = []
        for tag in self.tags.values():
            code = tag.code
            if not codes or code in codes:
                tags.append((code, tag))
        return tags

    def aspage(self):
        """Return self."""
        return self

    @property
    def keyframe(self):
        """Return keyframe, self."""
        return self

    @keyframe.setter
    def keyframe(self, index):
        """Set keyframe, NOP."""
        return

    @lazyattr
    def pages(self):
        """Return sequence of sub-pages (SubIFDs)."""
        if 'SubIFDs' not in self.tags:
            return tuple()
        return TiffPages(self)

    @property
    def hash(self):
        """Return checksum to identify pages in same series."""
        return hash(
            self._shape + (
                self.tilewidth, self.tilelength, self.tiledepth,
                self.bitspersample, self.fillorder, self.predictor,
                self.extrasamples, self.photometric, self.compression,
                self.planarconfig))

    @lazyattr
    def _offsetscounts(self):
        """Return simplified offsets and bytecounts."""
        if self.is_contiguous:
            offset, bytecount = self.is_contiguous
            return [offset], [bytecount]
        if self.is_tiled:
            return self.dataoffsets, self.databytecounts
        return clean_offsetscounts(self.dataoffsets, self.databytecounts)

    @lazyattr
    def is_contiguous(self):
        """Return offset and size of contiguous data, else None.

        Excludes prediction and fill_order.

        """
        if (self.compression != 1
                or self.bitspersample not in (8, 16, 32, 64)):
            return None
        if 'TileWidth' in self.tags:
            if (self.imagewidth != self.tilewidth or
                    self.imagelength % self.tilelength or
                    self.tilewidth % 16 or self.tilelength % 16):
                return None
            if ('ImageDepth' in self.tags and 'TileDepth' in self.tags and
                    (self.imagelength != self.tilelength or
                     self.imagedepth % self.tiledepth)):
                return None

        offsets = self.dataoffsets
        bytecounts = self.databytecounts
        if len(offsets) == 1:
            return offsets[0], bytecounts[0]
        if self.is_stk or all((offsets[i] + bytecounts[i] == offsets[i+1] or
                               bytecounts[i+1] == 0)  # no data/ignore offset
                              for i in range(len(offsets)-1)):
            return offsets[0], sum(bytecounts)
        return None

    @lazyattr
    def is_final(self):
        """Return if page's image data are stored in final form.

        Excludes byte-swapping.

        """
        return (self.is_contiguous and self.fillorder == 1 and
                self.predictor == 1 and not self.is_subsampled)

    @lazyattr
    def is_memmappable(self):
        """Return if page's image data in file can be memory-mapped."""
        return (self.parent.filehandle.is_file and self.is_final and
                # (self.bitspersample == 8 or self.parent.isnative) and
                self.is_contiguous[0] % self.dtype.itemsize == 0)  # aligned?

    def __str__(self, detail=0, width=79):
        """Return string containing information about page."""
        if self.keyframe != self:
            return TiffFrame.__str__(self, detail, width)
        attr = ''
        for name in ('memmappable', 'final', 'contiguous'):
            attr = getattr(self, 'is_'+name)
            if attr:
                attr = name.upper()
                break
        info = '  '.join(s.lower() for s in (
            'x'.join(str(i) for i in self.shape),
            '%s%s' % (TIFF.SAMPLEFORMAT(self.sampleformat).name,
                      self.bitspersample),
            ' '.join(i for i in (
                TIFF.PHOTOMETRIC(self.photometric).name,
                'REDUCED' if self.is_reduced else '',
                'MASK' if self.is_mask else '',
                'TILED' if self.is_tiled else '',
                self.compression.name if self.compression != 1 else '',
                self.planarconfig.name if self.planarconfig != 1 else '',
                self.predictor.name if self.predictor != 1 else '',
                self.fillorder.name if self.fillorder != 1 else '',
                ) + tuple(f.upper() for f in self.flags) + (attr,)
                     if i)
            ) if s)
        info = 'TiffPage %i @%i  %s' % (self.index, self.offset, info)
        if detail <= 0:
            return info
        info = [info]
        tags = self.tags
        tlines = []
        vlines = []
        for tag in sorted(tags.values(), key=lambda x: x.code):
            value = tag.__str__(width=width+1)
            tlines.append(value[:width].strip())
            if detail > 1 and len(value) > width:
                name = tag.name.upper()
                if detail <= 2 and ('COUNTS' in name or 'OFFSETS' in name):
                    value = pformat(tag.value, width=width, height=detail*4)
                else:
                    value = pformat(tag.value, width=width, height=detail*12)
                vlines.append('%s\n%s' % (tag.name, value))
        info.append('\n'.join(tlines))
        if detail > 1:
            info.append('\n\n'.join(vlines))
            for name in ('ndpi',):
                name = name + '_tags'
                attr = getattr(self, name, False)
                if attr:
                    info.append('%s\n%s' % (name.upper(), pformat(attr)))
        if detail > 3:
            try:
                info.append('DATA\n%s' % pformat(
                    self.asarray(), width=width, height=detail*8))
            except Exception:
                pass
        return '\n\n'.join(info)

    @lazyattr
    def flags(self):
        """Return set of flags."""
        return set((name.lower() for name in sorted(TIFF.FILE_FLAGS)
                    if getattr(self, 'is_' + name)))

    @property
    def ndim(self):
        """Return number of array dimensions."""
        return len(self.shape)

    @property
    def size(self):
        """Return number of elements in array."""
        return product(self.shape)

    @lazyattr
    def andor_tags(self):
        """Return consolidated metadata from Andor tags as dict.

        Remove Andor tags from self.tags.

        """
        if not self.is_andor:
            return None
        tags = self.tags
        result = {'Id': tags['AndorId'].value}
        for tag in list(self.tags.values()):
            code = tag.code
            if not 4864 < code < 5031:
                continue
            value = tag.value
            name = tag.name[5:] if len(tag.name) > 5 else tag.name
            result[name] = value
            del tags[tag.name]
        return result

    @lazyattr
    def epics_tags(self):
        """Return consolidated metadata from EPICS areaDetector tags as dict.

        Remove areaDetector tags from self.tags.

        """
        if not self.is_epics:
            return None
        result = {}
        tags = self.tags
        for tag in list(self.tags.values()):
            code = tag.code
            if not 65000 <= code < 65500:
                continue
            value = tag.value
            if code == 65000:
                result['timeStamp'] = datetime.datetime.fromtimestamp(
                    float(value))
            elif code == 65001:
                result['uniqueID'] = int(value)
            elif code == 65002:
                result['epicsTSSec'] = int(value)
            elif code == 65003:
                result['epicsTSNsec'] = int(value)
            else:
                key, value = value.split(':', 1)
                result[key] = astype(value)
            del tags[tag.name]
        return result

    @lazyattr
    def ndpi_tags(self):
        """Return consolidated metadata from Hamamatsu NDPI as dict."""
        if not self.is_ndpi:
            return None
        tags = self.tags
        result = {}
        for name in ('Make', 'Model', 'Software'):
            result[name] = tags[name].value
        for code, name in TIFF.NDPI_TAGS.items():
            code = str(code)
            if code in tags:
                result[name] = tags[code].value
                # del tags[code]
        return result

    @lazyattr
    def geotiff_tags(self):
        """Return consolidated metadata from GeoTIFF tags as dict."""
        if not self.is_geotiff:
            return None
        tags = self.tags

        gkd = tags['GeoKeyDirectoryTag'].value
        if gkd[0] != 1:
            log.warning('GeoTIFF tags: invalid GeoKeyDirectoryTag')
            return {}

        result = {
            'KeyDirectoryVersion': gkd[0],
            'KeyRevision': gkd[1],
            'KeyRevisionMinor': gkd[2],
            # 'NumberOfKeys': gkd[3],
        }
        # deltags = ['GeoKeyDirectoryTag']
        geokeys = TIFF.GEO_KEYS
        geocodes = TIFF.GEO_CODES
        for index in range(gkd[3]):
            keyid, tagid, count, offset = gkd[4 + index * 4: index * 4 + 8]
            keyid = geokeys.get(keyid, keyid)
            if tagid == 0:
                value = offset
            else:
                tagname = TIFF.TAGS[tagid]
                # deltags.append(tagname)
                value = tags[tagname].value[offset: offset + count]
                if tagid == 34737 and count > 1 and value[-1] == '|':
                    value = value[:-1]
                value = value if count > 1 else value[0]
            if keyid in geocodes:
                try:
                    value = geocodes[keyid](value)
                except Exception:
                    pass
            result[keyid] = value

        if 'IntergraphMatrixTag' in tags:
            value = tags['IntergraphMatrixTag'].value
            value = numpy.array(value)
            if len(value) == 16:
                value = value.reshape((4, 4)).tolist()
            result['IntergraphMatrix'] = value
        if 'ModelPixelScaleTag' in tags:
            value = numpy.array(tags['ModelPixelScaleTag'].value).tolist()
            result['ModelPixelScale'] = value
        if 'ModelTiepointTag' in tags:
            value = tags['ModelTiepointTag'].value
            value = numpy.array(value).reshape((-1, 6)).squeeze().tolist()
            result['ModelTiepoint'] = value
        if 'ModelTransformationTag' in tags:
            value = tags['ModelTransformationTag'].value
            value = numpy.array(value).reshape((4, 4)).tolist()
            result['ModelTransformation'] = value
        # if 'ModelPixelScaleTag' in tags and 'ModelTiepointTag' in tags:
        #     sx, sy, sz = tags['ModelPixelScaleTag'].value
        #     tiepoints = tags['ModelTiepointTag'].value
        #     transforms = []
        #     for tp in range(0, len(tiepoints), 6):
        #         i, j, k, x, y, z = tiepoints[tp:tp+6]
        #         transforms.append([
        #             [sx, 0.0, 0.0, x - i * sx],
        #             [0.0, -sy, 0.0, y + j * sy],
        #             [0.0, 0.0, sz, z - k * sz],
        #             [0.0, 0.0, 0.0, 1.0]])
        #     if len(tiepoints) == 6:
        #         transforms = transforms[0]
        #     result['ModelTransformation'] = transforms

        if 'RPCCoefficientTag' in tags:
            rpcc = tags['RPCCoefficientTag'].value
            result['RPCCoefficient'] = {
                'ERR_BIAS': rpcc[0],
                'ERR_RAND': rpcc[1],
                'LINE_OFF': rpcc[2],
                'SAMP_OFF': rpcc[3],
                'LAT_OFF': rpcc[4],
                'LONG_OFF': rpcc[5],
                'HEIGHT_OFF': rpcc[6],
                'LINE_SCALE': rpcc[7],
                'SAMP_SCALE': rpcc[8],
                'LAT_SCALE': rpcc[9],
                'LONG_SCALE': rpcc[10],
                'HEIGHT_SCALE': rpcc[11],
                'LINE_NUM_COEFF': rpcc[12:33],
                'LINE_DEN_COEFF ': rpcc[33:53],
                'SAMP_NUM_COEFF': rpcc[53:73],
                'SAMP_DEN_COEFF': rpcc[73:]}

        return result

    @property
    def is_reduced(self):
        """Page is reduced image of another image."""
        return self.subfiletype & 0b1

    @property
    def is_multipage(self):
        """Page is part of multi-page image."""
        return self.subfiletype & 0b10

    @property
    def is_mask(self):
        """Page is transparency mask for another image."""
        return self.subfiletype & 0b100

    @property
    def is_mrc(self):
        """Page is part of Mixed Raster Content."""
        return self.subfiletype & 0b1000

    @property
    def is_tiled(self):
        """Page contains tiled image."""
        return 'TileWidth' in self.tags

    @property
    def is_subsampled(self):
        """Page contains chroma subsampled image."""
        return ('YCbCrSubSampling' in self.tags and
                self.tags['YCbCrSubSampling'].value != (1, 1))

    @lazyattr
    def is_imagej(self):
        """Return ImageJ description if exists, else None."""
        for description in (self.description, self.description1):
            if not description:
                return None
            if description[:7] == 'ImageJ=':
                return description
        return None

    @lazyattr
    def is_shaped(self):
        """Return description containing array shape if exists, else None."""
        for description in (self.description, self.description1):
            if not description:
                return None
            if description[:1] == '{' and '"shape":' in description:
                return description
            if description[:6] == 'shape=':
                return description
        return None

    @property
    def is_mdgel(self):
        """Page contains MDFileTag tag."""
        return 'MDFileTag' in self.tags

    @property
    def is_mediacy(self):
        """Page contains Media Cybernetics Id tag."""
        return ('MC_Id' in self.tags and
                self.tags['MC_Id'].value[:7] == b'MC TIFF')

    @property
    def is_stk(self):
        """Page contains UIC2Tag tag."""
        return 'UIC2tag' in self.tags

    @property
    def is_lsm(self):
        """Page contains CZ_LSMINFO tag."""
        return 'CZ_LSMINFO' in self.tags

    @property
    def is_fluoview(self):
        """Page contains FluoView MM_STAMP tag."""
        return 'MM_Stamp' in self.tags

    @property
    def is_nih(self):
        """Page contains NIH image header."""
        return 'NIHImageHeader' in self.tags

    @property
    def is_sgi(self):
        """Page contains SGI image and tile depth tags."""
        return 'ImageDepth' in self.tags and 'TileDepth' in self.tags

    @property
    def is_vista(self):
        """Software tag is 'ISS Vista'."""
        return self.software == 'ISS Vista'

    @property
    def is_metaseries(self):
        """Page contains MDS MetaSeries metadata in ImageDescription tag."""
        if self.index > 1 or self.software != 'MetaSeries':
            return False
        d = self.description
        return d.startswith('<MetaData>') and d.endswith('</MetaData>')

    @property
    def is_ome(self):
        """Page contains OME-XML in ImageDescription tag."""
        if self.index > 1 or not self.description:
            return False
        d = self.description
        return d[:14] == '<?xml version=' and d[-6:] == '</OME>'

    @property
    def is_scn(self):
        """Page contains Leica SCN XML in ImageDescription tag."""
        if self.index > 1 or not self.description:
            return False
        d = self.description
        return d[:14] == '<?xml version=' and d[-6:] == '</scn>'

    @property
    def is_micromanager(self):
        """Page contains Micro-Manager metadata."""
        return 'MicroManagerMetadata' in self.tags

    @property
    def is_andor(self):
        """Page contains Andor Technology tags."""
        return 'AndorId' in self.tags

    @property
    def is_pilatus(self):
        """Page contains Pilatus tags."""
        return (self.software[:8] == 'TVX TIFF' and
                self.description[:2] == '# ')

    @property
    def is_epics(self):
        """Page contains EPICS areaDetector tags."""
        return (self.description == 'EPICS areaDetector' or
                self.software == 'EPICS areaDetector')

    @property
    def is_tvips(self):
        """Page contains TVIPS metadata."""
        return 'TVIPS' in self.tags

    @property
    def is_fei(self):
        """Page contains SFEG or HELIOS metadata."""
        return 'FEI_SFEG' in self.tags or 'FEI_HELIOS' in self.tags

    @property
    def is_sem(self):
        """Page contains Zeiss SEM metadata."""
        return 'CZ_SEM' in self.tags

    @property
    def is_svs(self):
        """Page contains Aperio metadata."""
        return self.description[:20] == 'Aperio Image Library'

    @property
    def is_scanimage(self):
        """Page contains ScanImage metadata."""
        return (self.description[:12] == 'state.config' or
                self.software[:22] == 'SI.LINE_FORMAT_VERSION' or
                'scanimage.SI' in self.description[-256:])

    @property
    def is_qpi(self):
        """Page contains PerkinElmer tissue images metadata."""
        # The ImageDescription tag contains XML with a top-level
        # <PerkinElmer-QPI-ImageDescription> element
        return self.software[:15] == 'PerkinElmer-QPI'

    @property
    def is_geotiff(self):
        """Page contains GeoTIFF metadata."""
        return 'GeoKeyDirectoryTag' in self.tags

    @property
    def is_sis(self):
        """Page contains Olympus SIS metadata."""
        return 'OlympusSIS' in self.tags or 'OlympusINI' in self.tags

    @lazyattr  # must not be property; tag 65420 is later removed
    def is_ndpi(self):
        """Page contains NDPI metadata."""
        return '65420' in self.tags and 'Make' in self.tags


class TiffFrame(object):
    """Lightweight TIFF image file directory (IFD).

    Only a limited number of tag values are read from file, e.g. StripOffsets,
    and StripByteCounts. Other tag values are assumed to be identical with a
    specified TiffPage instance, the keyframe.

    TiffFrame is intended to reduce resource usage and speed up reading image
    data from file, not for introspection of metadata.

    Not compatible with Python 2.

    """
    __slots__ = 'index', 'parent', 'offset', '_offsetscounts', '_keyframe'

    is_mdgel = False
    pages = None
    tags = {}

    def __init__(self, parent, index, offset=None, keyframe=None,
                 offsets=None, bytecounts=None):
        """Initialize TiffFrame from file or values.

        The file handle position must be at the offset to a valid IFD.

        """
        self._keyframe = None
        self.parent = parent
        self.index = index
        self.offset = offset

        if offsets is not None:
            # initialize "virtual frame" from offsets and bytecounts
            self._offsetscounts = offsets, bytecounts
            self._keyframe = keyframe
            return

        if offset is None:
            self.offset = parent.filehandle.tell()
        else:
            parent.filehandle.seek(offset)

        if keyframe is None:
            tags = {273, 279, 324, 325}
        elif keyframe.is_contiguous:
            tags = {256, 273, 324}
        else:
            tags = {256, 273, 279, 324, 325}

        dataoffsets = databytecounts = []

        for code, tag in self._gettags(tags):
            if code == 273 or code == 324:
                dataoffsets = list(tag.value)
            elif code == 279 or code == 325:
                databytecounts = list(tag.value)
            elif code == 256 and keyframe.imagewidth != tag.value:
                raise RuntimeError('incompatible keyframe')
            # elif code == 270:
            #     tagname = tag.name
            #     if tagname not in tags:
            #         tags[tagname] = bytes2str(tag.value)
            #     elif 'ImageDescription1' not in tags:
            #         tags['ImageDescription1'] = bytes2str(tag.value)
            # else:
            #     tags[tag.name] = tag.value

        if not dataoffsets:
            log.warning('TiffFrame %i is missing required tags', index)

        self._offsetscounts = dataoffsets, databytecounts

        if keyframe is not None:
            self.keyframe = keyframe

    def _gettags(self, codes=None, lock=None):
        """Return list of (code, TiffTag) from file."""
        fh = self.parent.filehandle
        tiff = self.parent.tiff
        unpack = struct.unpack
        lock = NullContext() if lock is None else lock
        tags = []

        with lock:
            fh.seek(self.offset)
            try:
                tagno = unpack(tiff.tagnoformat, fh.read(tiff.tagnosize))[0]
                if tagno > 4096:
                    raise TiffFileError('suspicious number of tags')
            except Exception:
                raise TiffFileError(
                    'corrupted page list at offset %i' % self.offset)

            tagoffset = self.offset + tiff.tagnosize  # fh.tell()
            tagsize = tiff.tagsize
            tagindex = -tagsize
            codeformat = tiff.tagformat1[:2]
            tagbytes = fh.read(tagsize * tagno)

            for _ in range(tagno):
                tagindex += tagsize
                code = unpack(codeformat, tagbytes[tagindex:tagindex+2])[0]
                if codes and code not in codes:
                    continue
                try:
                    tag = TiffTag(self.parent,
                                  tagbytes[tagindex:tagindex+tagsize],
                                  tagoffset+tagindex)
                except TiffFileError as exc:
                    log.warning('%s: %s', exc.__class__.__name__, exc)
                    continue
                tags.append((code, tag))

        return tags

    def aspage(self):
        """Return TiffPage from file."""
        if self.offset is None:
            raise ValueError('cannot return virtual frame as page.')
        self.parent.filehandle.seek(self.offset)
        return TiffPage(self.parent, index=self.index)

    def asarray(self, *args, **kwargs):
        """Read image data from file and return as numpy array."""
        # TODO: fix TypeError on Python 2
        #   "TypeError: unbound method asarray() must be called with TiffPage
        #   instance as first argument (got TiffFrame instance instead)"
        if self._keyframe is None:
            raise RuntimeError('keyframe not set')
        kwargs['validate'] = False
        return TiffPage.asarray(self, *args, **kwargs)

    def asrgb(self, *args, **kwargs):
        """Read image data from file and return RGB image as numpy array."""
        if self._keyframe is None:
            raise RuntimeError('keyframe not set')
        kwargs['validate'] = False
        return TiffPage.asrgb(self, *args, **kwargs)

    @property
    def keyframe(self):
        """Return keyframe."""
        return self._keyframe

    @keyframe.setter
    def keyframe(self, keyframe):
        """Set keyframe."""
        if self._keyframe == keyframe:
            return
        if self._keyframe is not None:
            raise RuntimeError('cannot reset keyframe')
        if len(self._offsetscounts[0]) != len(keyframe.dataoffsets):
            raise RuntimeError('incompatible keyframe')
        if keyframe.is_tiled:
            pass
        if keyframe.is_contiguous:
            self._offsetscounts = ([self._offsetscounts[0][0]],
                                   [keyframe.is_contiguous[1]])
        else:
            self._offsetscounts = clean_offsetscounts(*self._offsetscounts)
        self._keyframe = keyframe

    @property
    def is_contiguous(self):
        """Return offset and size of contiguous data, else None."""
        if self._keyframe is None:
            raise RuntimeError('keyframe not set')
        if self._keyframe.is_contiguous:
            return self._offsetscounts[0][0], self._keyframe.is_contiguous[1]
        return None

    @property
    def is_memmappable(self):
        """Return if page's image data in file can be memory-mapped."""
        if self._keyframe is None:
            raise RuntimeError('keyframe not set')
        return self._keyframe.is_memmappable

    @property
    def hash(self):
        """Return checksum to identify pages in same series."""
        if self._keyframe is None:
            raise RuntimeError('keyframe not set')
        return self._keyframe.hash

    def __getattr__(self, name):
        """Return attribute from keyframe."""
        if name in TIFF.FRAME_ATTRS:
            return getattr(self._keyframe, name)
        # this error could be raised because an AttributeError was
        # raised inside a @property function
        raise AttributeError("'%s' object has no attribute '%s'" %
                             (self.__class__.__name__, name))

    def __str__(self, detail=0, width=79):
        """Return string containing information about frame."""
        if self._keyframe is None:
            info = ''
            kf = None
        else:
            info = '  '.join(s for s in ('x'.join(str(i) for i in self.shape),
                                         str(self.dtype)))
            kf = TiffPage.__str__(self._keyframe, width=width-11)
        if detail > 3:
            of, bc = self._offsetscounts
            of = pformat(of, width=width-9, height=detail-3)
            bc = pformat(bc, width=width-13, height=detail-3)
            info = '\n Keyframe %s\n Offsets %s\n Bytecounts %s' % (
                kf, of, bc)
        return 'TiffFrame %i @%s  %s' % (self.index, self.offset, info)


class TiffTag(object):
    """TIFF tag structure.

    Attributes
    ----------
    name : string
        Name of tag.
    code : int
        Decimal code of tag.
    dtype : str
        Datatype of tag data. One of TIFF DATA_FORMATS.
    count : int
        Number of values.
    value : various types
        Tag data as Python object.
    ImageSourceData : int
        Location of value in file.

    All attributes are read-only.

    """
    __slots__ = ('code', 'count', 'dtype', 'value', 'valueoffset')

    def __init__(self, parent, tagheader, tagoffset):
        """Initialize instance from tag header."""
        fh = parent.filehandle
        tiff = parent.tiff
        byteorder = tiff.byteorder
        offsetsize = tiff.offsetsize
        unpack = struct.unpack

        self.valueoffset = tagoffset + offsetsize + 4
        code, type_ = unpack(tiff.tagformat1, tagheader[:4])
        count, value = unpack(tiff.tagformat2, tagheader[4:])

        try:
            dtype = TIFF.DATA_FORMATS[type_]
        except KeyError:
            raise TiffFileError('unknown tag data type %i' % type_)

        fmt = '%s%i%s' % (byteorder, count * int(dtype[0]), dtype[1])
        size = struct.calcsize(fmt)
        if size > offsetsize or code in TIFF.TAG_READERS:
            self.valueoffset = offset = unpack(tiff.offsetformat, value)[0]
            if offset < 8 or offset > fh.size - size:
                raise TiffFileError('invalid tag value offset')
            # if offset % 2:
            #     log.warning('TiffTag: value does not begin on word boundary')
            fh.seek(offset)
            if code in TIFF.TAG_READERS:
                readfunc = TIFF.TAG_READERS[code]
                value = readfunc(fh, byteorder, dtype, count, offsetsize)
            elif type_ == 7 or (count > 1 and dtype[-1] == 'B'):
                value = read_bytes(fh, byteorder, dtype, count, offsetsize)
            elif code in TIFF.TAGS or dtype[-1] == 's':
                value = unpack(fmt, fh.read(size))
            else:
                value = read_numpy(fh, byteorder, dtype, count, offsetsize)
        elif dtype[-1] == 'B' or type_ == 7:
            value = value[:size]
        else:
            value = unpack(fmt, value[:size])

        process = (code not in TIFF.TAG_READERS and code not in TIFF.TAG_TUPLE
                   and type_ != 7)
        if process and dtype[-1] == 's' and isinstance(value[0], bytes):
            # TIFF ASCII fields can contain multiple strings,
            #   each terminated with a NUL
            value = value[0]
            try:
                value = bytes2str(stripascii(value).strip())
            except UnicodeDecodeError:
                # TODO: this doesn't work on Python 2
                log.warning(
                    'TiffTag %i: coercing invalid ASCII to bytes', code)
                dtype = '1B'
        else:
            if code in TIFF.TAG_ENUM:
                t = TIFF.TAG_ENUM[code]
                try:
                    value = tuple(t(v) for v in value)
                except ValueError as exc:
                    log.warning('TiffTag  %i: %s', code, str(exc))
            if process:
                if len(value) == 1:
                    value = value[0]

        self.code = code
        self.dtype = dtype
        self.count = count
        self.value = value

    @property
    def name(self):
        """Return name of tag from TIFF.TAGS registry."""
        try:
            return TIFF.TAGS[self.code]
        except KeyError:
            return str(self.code)

    def _fix_lsm_bitspersample(self, parent):
        """Correct LSM bitspersample tag.

        Old LSM writers may use a separate region for two 16-bit values,
        although they fit into the tag value element of the tag.

        """
        if self.code != 258 or self.count != 2:
            return
        # TODO: test this case; need example file
        log.warning('TiffTag %i: correcting LSM bitspersample tag', self.code)
        value = struct.pack('<HH', *self.value)
        self.valueoffset = struct.unpack('<I', value)[0]
        parent.filehandle.seek(self.valueoffset)
        self.value = struct.unpack('<HH', parent.filehandle.read(4))

    def __str__(self, detail=0, width=79):
        """Return string containing information about tag."""
        height = 1 if detail <= 0 else 8 * detail
        tcode = '%i%s' % (self.count * int(self.dtype[0]), self.dtype[1])
        if self.name == str(self.code):
            codename = self.name
        else:
            codename = '%i %s' % (self.code, self.name)
        line = 'TiffTag %s %s @%i  ' % (codename, tcode,
                                        self.valueoffset)[:width]

        if self.code in TIFF.TAG_ENUM:
            if self.count == 1:
                value = TIFF.TAG_ENUM[self.code](self.value).name
            else:
                value = pformat(tuple(v.name for v in self.value))
        else:
            value = pformat(self.value, width=width, height=height)

        if detail <= 0:
            line += value
            line = line[:width]
        else:
            line += '\n' + value
        return line


class TiffPageSeries(object):
    """Series of TIFF pages with compatible shape and data type.

    Attributes
    ----------
    pages : list of TiffPage
        Sequence of TiffPages in series.
    dtype : numpy.dtype
        Data type (native byte order) of the image array in series.
    shape : tuple
        Dimensions of the image array in series.
    axes : str
        Labels of axes in shape. See TiffPage.axes.
    offset : int or None
        Position of image data in file if memory-mappable, else None.

    """
    def __init__(self, pages, shape, dtype, axes, parent=None, name=None,
                 transform=None, kind=None, truncated=False):
        """Initialize instance."""
        self.index = 0
        self._pages = pages  # might contain only first of contiguous pages
        self.shape = tuple(shape)
        self.axes = ''.join(axes)
        self.dtype = numpy.dtype(dtype)
        self.kind = kind if kind else ''
        self.name = name if name else ''
        self.transform = transform
        if parent:
            self.parent = parent
        elif pages:
            self.parent = pages[0].parent
        else:
            self.parent = None
        if len(pages) == 1 and not truncated:
            self._len = int(product(self.shape) // product(pages[0].shape))
        else:
            self._len = len(pages)

    def asarray(self, out=None):
        """Return image data from series of TIFF pages as numpy array."""
        if self.parent:
            result = self.parent.asarray(series=self, out=out)
            if self.transform is not None:
                result = self.transform(result)
            return result
        return None

    @lazyattr
    def offset(self):
        """Return offset to series data in file, if any."""
        if not self._pages:
            return None

        pos = 0
        for page in self._pages:
            if page is None:
                return None
            if not page.is_final:
                return None
            if not pos:
                pos = page.is_contiguous[0] + page.is_contiguous[1]
                continue
            if pos != page.is_contiguous[0]:
                return None
            pos += page.is_contiguous[1]

        page = self._pages[0]
        offset = page.is_contiguous[0]
        if (page.is_imagej or page.is_shaped) and len(self._pages) == 1:
            # truncated files
            return offset
        if pos == offset + product(self.shape) * self.dtype.itemsize:
            return offset
        return None

    @property
    def ndim(self):
        """Return number of array dimensions."""
        return len(self.shape)

    @property
    def size(self):
        """Return number of elements in array."""
        return int(product(self.shape))

    @property
    def pages(self):
        """Return sequence of all pages in series."""
        # a workaround to keep the old interface working
        return self

    def _getitem(self, key):
        """Return specified page of series from cache or file."""
        key = int(key)
        if key < 0:
            key %= self._len
        if len(self._pages) == 1 and 0 < key < self._len:
            index = self._pages[0].index
            return self.parent.pages._getitem(index + key)
        return self._pages[key]

    def __getitem__(self, key):
        """Return specified page(s)."""
        getitem = self._getitem
        if isinstance(key, inttypes):
            return getitem(key)
        if isinstance(key, slice):
            return [getitem(i) for i in range(*key.indices(self._len))]
        if isinstance(key, Iterable):
            return [getitem(k) for k in key]
        raise TypeError('key must be an integer, slice, or iterable')

    def __iter__(self):
        """Return iterator over pages in series."""
        if len(self._pages) == self._len:
            for page in self._pages:
                yield page
        else:
            pages = self.parent.pages
            index = self._pages[0].index
            for i in range(self._len):
                yield pages[index + i]

    def __len__(self):
        """Return number of pages in series."""
        return self._len

    def __str__(self):
        """Return string with information about series."""
        s = '  '.join(s for s in (
            snipstr("'%s'" % self.name, 20) if self.name else '',
            'x'.join(str(i) for i in self.shape),
            str(self.dtype),
            self.axes,
            self.kind,
            '%i Pages' % len(self.pages),
            ('Offset=%i' % self.offset) if self.offset else '') if s)
        return 'TiffPageSeries %i  %s' % (self.index, s)


class TiffSequence(object):
    """Sequence of TIFF files.

    The image data in all files must match shape, dtype, etc.

    Attributes
    ----------
    files : list
        List of file names.
    shape : tuple
        Shape of image sequence. Excludes shape of image array.
    axes : str
        Labels of axes in shape.

    """
    _patterns = {
        'axes': r"""
            # matches Olympus OIF and Leica TIFF series
            _?(?:(q|l|p|a|c|t|x|y|z|ch|tp)(\d{1,4}))
            _?(?:(q|l|p|a|c|t|x|y|z|ch|tp)(\d{1,4}))?
            _?(?:(q|l|p|a|c|t|x|y|z|ch|tp)(\d{1,4}))?
            _?(?:(q|l|p|a|c|t|x|y|z|ch|tp)(\d{1,4}))?
            _?(?:(q|l|p|a|c|t|x|y|z|ch|tp)(\d{1,4}))?
            _?(?:(q|l|p|a|c|t|x|y|z|ch|tp)(\d{1,4}))?
            _?(?:(q|l|p|a|c|t|x|y|z|ch|tp)(\d{1,4}))?
            """}

    class ParseError(Exception):
        """Custom TiffSequence parser error."""

    def __init__(self, files=None, container=None, sort=None, pattern=None,
                 imread=None):
        """Initialize instance from multiple files.

        Parameters
        ----------
        files : str, pathlib.Path, or sequence thereof
            Glob filename pattern or sequence of file names.
            Default is '\\*.tif'.
            Binary streams are not supported.
        container : str or container instance
            Name or open instance of ZIP file in which files are stored.
        sort : function
            Sort function used to sort file names when 'files' is a  pattern.
            The default (None) is natural_sorted. Use sort=False to disable
            sorting.
        pattern : str
            Regular expression pattern that matches axes names and sequence
            indices in file names. By default (None), no pattern matching is
            performed. The predefined 'axes' pattern matches Olympus OIF and
            Leica TIFF series.
        imread : function or class
            Image read function or class with asarray function returning numpy
            array from single file.

        """
        if sort is None:
            sort = natural_sorted
        if files is None:
            files = '*.tif'
        self._container = container
        if container:
            import fnmatch  # noqa
            if isinstance(container, basestring):
                import zipfile  # noqa
                self._container = zipfile.ZipFile(container)
            elif not hasattr(self._container, 'open'):
                raise ValueError('invalid container')
            if isinstance(files, basestring):
                files = fnmatch.filter(self._container.namelist(), files)
                if sort:
                    files = sort(files)
        else:
            if isinstance(files, pathlib.Path):
                files = str(files)
            if isinstance(files, basestring):
                files = glob.glob(files)
                if sort:
                    files = sort(files)
            if not files:
                raise ValueError('no files found')

        files = list(files)
        if not files:
            raise ValueError('no files found')
        if isinstance(files[0], pathlib.Path):
            files = [str(pathlib.Path(f)) for f in files]
        elif not isinstance(files[0], basestring):
            raise ValueError('not a file name')
        self.files = files

        if imread is None:
            imread = TiffFile

        if hasattr(imread, 'asarray'):
            # redefine imread to use asarray from class
            _imread0 = imread

            def imread(fname, **kwargs):
                with _imread0(fname) as im:
                    return im.asarray(**kwargs)

        if container:
            # redefine imread to read from container
            _imread1 = imread

            def imread(fname, **kwargs):
                with self._container.open(fname) as fh:
                    with io.BytesIO(fh.read()) as fh2:
                        return _imread1(fh2, **kwargs)

        self.imread = imread
        self.axes = 'I'
        self.shape = (len(files),)
        self._startindex = (0,)
        self._indices = tuple((i,) for i in range(len(files)))

        self.pattern = self._patterns.get(pattern, pattern)
        if self.pattern:
            try:
                self._parse()
                if not self.axes:
                    self.axes = 'I'
            except TiffSequence.ParseError:
                pass

    def __str__(self):
        """Return string with information about image sequence."""
        return '\n'.join([
            str(self._container) if self._container else self.files[0],
            ' size: %i' % len(self.files),
            ' axes: %s' % self.axes,
            ' shape: %s' % str(self.shape)])

    def __len__(self):
        return len(self.files)

    def __enter__(self):
        return self

    def __exit__(self, exc_type, exc_value, traceback):
        self.close()

    def close(self):
        if self._container:
            self._container.close()
        self._container = None

    def asarray(self, file=None, out=None, **kwargs):
        """Read image data from files and return as numpy array.

        The kwargs parameters are passed to the imread function.

        Raise IndexError or ValueError if image shapes do not match.

        """
        if file is not None:
            if isinstance(file, int):
                return self.imread(self.files[file], **kwargs)
            return self.imread(file, **kwargs)

        im = self.imread(self.files[0], **kwargs)
        shape = self.shape + im.shape
        result = create_output(out, shape, dtype=im.dtype)
        result = result.reshape(-1, *im.shape)
        for index, fname in zip(self._indices, self.files):
            index = [i-j for i, j in zip(index, self._startindex)]
            index = numpy.ravel_multi_index(index, self.shape)
            im = self.imread(fname, **kwargs)
            result[index] = im
        result.shape = shape
        return result

    def _parse(self):
        """Get axes and shape from file names."""
        if not self.pattern:
            raise TiffSequence.ParseError('invalid pattern')
        pattern = re.compile(self.pattern, re.IGNORECASE | re.VERBOSE)
        matches = pattern.findall(os.path.split(self.files[0])[-1])
        if not matches:
            raise TiffSequence.ParseError('pattern does not match file names')
        matches = matches[-1]
        if len(matches) % 2:
            raise TiffSequence.ParseError(
                'pattern does not match axis name and index')
        axes = ''.join(m for m in matches[::2] if m)
        if not axes:
            raise TiffSequence.ParseError('pattern does not match file names')

        indices = []
        for fname in self.files:
            fname = os.path.split(fname)[-1]
            matches = pattern.findall(fname)[-1]
            if axes != ''.join(m for m in matches[::2] if m):
                raise ValueError('axes do not match within image sequence')
            indices.append([int(m) for m in matches[1::2] if m])
        shape = tuple(numpy.max(indices, axis=0))
        startindex = tuple(numpy.min(indices, axis=0))
        shape = tuple(i-j+1 for i, j in zip(shape, startindex))
        if product(shape) != len(self.files):
            log.warning(
                'TiffSequence: files are missing. Missing data are zeroed')

        self.axes = axes.upper()
        self.shape = shape
        self._indices = indices
        self._startindex = startindex


class FileHandle(object):
    """Binary file handle.

    A limited, special purpose file handler that can:

    * handle embedded files (for CZI within CZI files)
    * re-open closed files (for multi-file formats, such as OME-TIFF)
    * read and write numpy arrays and records from file like objects

    Only 'rb' and 'wb' modes are supported. Concurrently reading and writing
    of the same stream is untested.

    When initialized from another file handle, do not use it unless this
    FileHandle is closed.

    Attributes
    ----------
    name : str
        Name of the file.
    path : str
        Absolute path to file.
    size : int
        Size of file in bytes.
    is_file : bool
        If True, file has a filno and can be memory-mapped.

    All attributes are read-only.

    """
    __slots__ = ('_fh', '_file', '_mode', '_name', '_dir', '_lock',
                 '_offset', '_size', '_close', 'is_file')

    def __init__(self, file, mode='rb', name=None, offset=None, size=None):
        """Initialize file handle from file name or another file handle.

        Parameters
        ----------
        file : str, pathlib.Path, binary stream, or FileHandle
            File name or seekable binary stream, such as an open file
            or BytesIO.
        mode : str
            File open mode in case 'file' is a file name. Must be 'rb' or 'wb'.
        name : str
            Optional name of file in case 'file' is a binary stream.
        offset : int
            Optional start position of embedded file. By default, this is
            the current file position.
        size : int
            Optional size of embedded file. By default, this is the number
            of bytes from the 'offset' to the end of the file.

        """
        self._file = file
        self._fh = None
        self._mode = mode
        self._name = name
        self._dir = ''
        self._offset = offset
        self._size = size
        self._close = True
        self.is_file = False
        self._lock = NullContext()
        self.open()

    def open(self):
        """Open or re-open file."""
        if self._fh:
            return  # file is open

        if isinstance(self._file, pathlib.Path):
            self._file = str(self._file)
        if isinstance(self._file, basestring):
            # file name
            self._file = os.path.realpath(self._file)
            self._dir, self._name = os.path.split(self._file)
            self._fh = open(self._file, self._mode)
            self._close = True
            if self._offset is None:
                self._offset = 0
        elif isinstance(self._file, FileHandle):
            # FileHandle
            self._fh = self._file._fh
            if self._offset is None:
                self._offset = 0
            self._offset += self._file._offset
            self._close = False
            if not self._name:
                if self._offset:
                    name, ext = os.path.splitext(self._file._name)
                    self._name = '%s@%i%s' % (name, self._offset, ext)
                else:
                    self._name = self._file._name
            if self._mode and self._mode != self._file._mode:
                raise ValueError('FileHandle has wrong mode')
            self._mode = self._file._mode
            self._dir = self._file._dir
        elif hasattr(self._file, 'seek'):
            # binary stream: open file, BytesIO
            try:
                self._file.tell()
            except Exception:
                raise ValueError('binary stream is not seekable')
            self._fh = self._file
            if self._offset is None:
                self._offset = self._file.tell()
            self._close = False
            if not self._name:
                try:
                    self._dir, self._name = os.path.split(self._fh.name)
                except AttributeError:
                    self._name = 'Unnamed binary stream'
            try:
                self._mode = self._fh.mode
            except AttributeError:
                pass
        else:
            raise ValueError('The first parameter must be a file name, '
                             'seekable binary stream, or FileHandle')

        if self._offset:
            self._fh.seek(self._offset)

        if self._size is None:
            pos = self._fh.tell()
            self._fh.seek(self._offset, 2)
            self._size = self._fh.tell()
            self._fh.seek(pos)

        try:
            self._fh.fileno()
            self.is_file = True
        except Exception:
            self.is_file = False

    def read(self, size=-1):
        """Read 'size' bytes from file, or until EOF is reached."""
        if size < 0 and self._offset:
            size = self._size
        return self._fh.read(size)

    def readinto(self, b):
        """Read up to len(b) bytes into b, and return number of bytes read."""
        return self._fh.readinto(b)

    def write(self, bytestring):
        """Write bytestring to file."""
        return self._fh.write(bytestring)

    def flush(self):
        """Flush write buffers if applicable."""
        return self._fh.flush()

    def memmap_array(self, dtype, shape, offset=0, mode='r', order='C'):
        """Return numpy.memmap of data stored in file."""
        if not self.is_file:
            raise ValueError('Cannot memory-map file without fileno')
        return numpy.memmap(self._fh, dtype=dtype, mode=mode,
                            offset=self._offset + offset,
                            shape=shape, order=order)

    def read_array(self, dtype, count=-1, out=None):
        """Return numpy array from file in native byte order."""
        fh = self._fh
        dtype = numpy.dtype(dtype)

        if count < 0:
            size = self._size if out is None else out.nbytes
            count = size // dtype.itemsize
        else:
            size = count * dtype.itemsize

        result = numpy.empty(count, dtype) if out is None else out

        if result.nbytes != size:
            raise ValueError('size mismatch')

        n = fh.readinto(result)
        if n != size:
            raise ValueError('failed to read %i bytes' % size)

        if not result.dtype.isnative:
            if not dtype.isnative:
                result.byteswap(True)
            result = result.newbyteorder()
        elif result.dtype.isnative != dtype.isnative:
            result.byteswap(True)

        if out is not None:
            if hasattr(out, 'flush'):
                out.flush()

        return result

    def read_record(self, dtype, shape=1, byteorder=None):
        """Return numpy record from file."""
        rec = numpy.rec
        try:
            record = rec.fromfile(self._fh, dtype, shape, byteorder=byteorder)
        except Exception:
            dtype = numpy.dtype(dtype)
            if shape is None:
                shape = self._size // dtype.itemsize
            size = product(sequence(shape)) * dtype.itemsize
            data = self._fh.read(size)
            record = rec.fromstring(data, dtype, shape, byteorder=byteorder)
        return record[0] if shape == 1 else record

    def write_empty(self, size):
        """Append size bytes to file. Position must be at end of file."""
        if size < 1:
            return
        self._fh.seek(size-1, 1)
        self._fh.write(b'\x00')

    def write_array(self, data):
        """Write numpy array to binary file."""
        try:
            data.tofile(self._fh)
        except Exception:
            # BytesIO
            self._fh.write(data.tostring())

    def tell(self):
        """Return file's current position."""
        return self._fh.tell() - self._offset

    def seek(self, offset, whence=0):
        """Set file's current position."""
        if self._offset:
            if whence == 0:
                self._fh.seek(self._offset + offset, whence)
                return
            if whence == 2 and self._size > 0:
                self._fh.seek(self._offset + self._size + offset, 0)
                return
        self._fh.seek(offset, whence)

    def close(self):
        """Close file."""
        if self._close and self._fh:
            self._fh.close()
            self._fh = None

    def __enter__(self):
        return self

    def __exit__(self, exc_type, exc_value, traceback):
        self.close()

    def __getattr__(self, name):
        """Return attribute from underlying file object."""
        if self._offset:
            warnings.warn(
                "FileHandle: '%s' not implemented for embedded files" % name)
        return getattr(self._fh, name)

    @property
    def name(self):
        return self._name

    @property
    def dirname(self):
        return self._dir

    @property
    def path(self):
        return os.path.join(self._dir, self._name)

    @property
    def size(self):
        return self._size

    @property
    def closed(self):
        return self._fh is None

    @property
    def lock(self):
        return self._lock

    @lock.setter
    def lock(self, value):
        self._lock = threading.RLock() if value else NullContext()


class NullContext(object):
    """Null context manager.

    >>> with NullContext():
    ...     pass

    """
    def __enter__(self):
        return self

    def __exit__(self, exc_type, exc_value, traceback):
        pass


class OpenFileCache(object):
    """Keep files open."""

    __slots__ = ('files', 'past', 'lock', 'size')

    def __init__(self, size, lock=None):
        """Initialize open file cache."""
        self.past = []  # FIFO of opened files
        self.files = {}  # refcounts of opened files
        self.lock = NullContext() if lock is None else lock
        self.size = int(size)

    def open(self, filehandle):
        """Re-open file if necessary."""
        with self.lock:
            if filehandle in self.files:
                self.files[filehandle] += 1
            elif filehandle.closed:
                filehandle.open()
                self.files[filehandle] = 1
                self.past.append(filehandle)

    def close(self, filehandle):
        """Close openend file if no longer used."""
        with self.lock:
            if filehandle in self.files:
                self.files[filehandle] -= 1
                # trim the file cache
                index = 0
                size = len(self.past)
                while size > self.size and index < size:
                    filehandle = self.past[index]
                    if self.files[filehandle] == 0:
                        filehandle.close()
                        del self.files[filehandle]
                        del self.past[index]
                        size -= 1
                    else:
                        index += 1

    def clear(self):
        """Close all opened files if not in use."""
        with self.lock:
            for filehandle, refcount in list(self.files.items()):
                if refcount == 0:
                    filehandle.close()
                    del self.files[filehandle]
                    del self.past[self.past.index(filehandle)]


class LazyConst(object):
    """Class whose attributes are computed on first access from its methods."""
    def __init__(self, cls):
        self._cls = cls
        self.__doc__ = getattr(cls, '__doc__')

    def __getattr__(self, name):
        func = getattr(self._cls, name)
        if not callable(func):
            return func
        try:
            value = func()
        except TypeError:
            # Python 2 unbound method
            value = func.__func__()
        setattr(self, name, value)
        return value


@LazyConst
class TIFF(object):
    """Namespace for module constants."""

    def CLASSIC_LE():
        class ClassicTiffLe(object):
            __slots__ = []
            version = 42
            byteorder = '<'
            offsetsize = 4
            offsetformat = '<I'
            ifdoffsetsize = 4
            ifdoffsetformat = '<I'
            tagnosize = 2
            tagnoformat = '<H'
            tagsize = 12
            tagformat1 = '<HH'
            tagformat2 = '<I4s'

        return ClassicTiffLe

    def CLASSIC_BE():
        class ClassicTiffBe(object):
            __slots__ = []
            version = 42
            byteorder = '>'
            offsetsize = 4
            offsetformat = '>I'
            ifdoffsetsize = 4
            ifdoffsetformat = '>I'
            tagnosize = 2
            tagnoformat = '>H'
            tagsize = 12
            tagformat1 = '>HH'
            tagformat2 = '>I4s'

        return ClassicTiffBe

    def BIG_LE():
        class BigTiffLe(object):
            __slots__ = []
            version = 43
            byteorder = '<'
            offsetsize = 8
            offsetformat = '<Q'
            ifdoffsetsize = 8
            ifdoffsetformat = '<Q'
            tagnosize = 8
            tagnoformat = '<Q'
            tagsize = 20
            tagformat1 = '<HH'
            tagformat2 = '<Q8s'

        return BigTiffLe

    def BIG_BE():
        class BigTiffBe(object):
            __slots__ = []
            version = 43
            byteorder = '>'
            offsetsize = 8
            offsetformat = '>Q'
            ifdoffsetsize = 8
            ifdoffsetformat = '>Q'
            tagnosize = 8
            tagnoformat = '>Q'
            tagsize = 20
            tagformat1 = '>HH'
            tagformat2 = '>Q8s'

        return BigTiffBe

    def NDPI_LE():
        class NdpiTiffLe(object):
            __slots__ = []
            version = 42
            byteorder = '<'
            offsetsize = 4
            offsetformat = '<I'
            ifdoffsetsize = 8  # NDPI uses 8 bytes IFD offsets
            ifdoffsetformat = '<Q'
            tagnosize = 2
            tagnoformat = '<H'
            tagsize = 12
            tagformat1 = '<HH'
            tagformat2 = '<I4s'

        return NdpiTiffLe

    def TAGS():
        # TIFF tag codes and names from TIFF6, TIFF/EP, EXIF, and other specs
        return {
            11: 'ProcessingSoftware',
            254: 'NewSubfileType',
            255: 'SubfileType',
            256: 'ImageWidth',
            257: 'ImageLength',
            258: 'BitsPerSample',
            259: 'Compression',
            262: 'PhotometricInterpretation',
            263: 'Thresholding',
            264: 'CellWidth',
            265: 'CellLength',
            266: 'FillOrder',
            269: 'DocumentName',
            270: 'ImageDescription',
            271: 'Make',
            272: 'Model',
            273: 'StripOffsets',
            274: 'Orientation',
            277: 'SamplesPerPixel',
            278: 'RowsPerStrip',
            279: 'StripByteCounts',
            280: 'MinSampleValue',
            281: 'MaxSampleValue',
            282: 'XResolution',
            283: 'YResolution',
            284: 'PlanarConfiguration',
            285: 'PageName',
            286: 'XPosition',
            287: 'YPosition',
            288: 'FreeOffsets',
            289: 'FreeByteCounts',
            290: 'GrayResponseUnit',
            291: 'GrayResponseCurve',
            292: 'T4Options',
            293: 'T6Options',
            296: 'ResolutionUnit',
            297: 'PageNumber',
            300: 'ColorResponseUnit',
            301: 'TransferFunction',
            305: 'Software',
            306: 'DateTime',
            315: 'Artist',
            316: 'HostComputer',
            317: 'Predictor',
            318: 'WhitePoint',
            319: 'PrimaryChromaticities',
            320: 'ColorMap',
            321: 'HalftoneHints',
            322: 'TileWidth',
            323: 'TileLength',
            324: 'TileOffsets',
            325: 'TileByteCounts',
            326: 'BadFaxLines',
            327: 'CleanFaxData',
            328: 'ConsecutiveBadFaxLines',
            330: 'SubIFDs',
            332: 'InkSet',
            333: 'InkNames',
            334: 'NumberOfInks',
            336: 'DotRange',
            337: 'TargetPrinter',
            338: 'ExtraSamples',
            339: 'SampleFormat',
            340: 'SMinSampleValue',
            341: 'SMaxSampleValue',
            342: 'TransferRange',
            343: 'ClipPath',
            344: 'XClipPathUnits',
            345: 'YClipPathUnits',
            346: 'Indexed',
            347: 'JPEGTables',
            351: 'OPIProxy',
            400: 'GlobalParametersIFD',
            401: 'ProfileType',
            402: 'FaxProfile',
            403: 'CodingMethods',
            404: 'VersionYear',
            405: 'ModeNumber',
            433: 'Decode',
            434: 'DefaultImageColor',
            435: 'T82Options',
            437: 'JPEGTables_',  # 347
            512: 'JPEGProc',
            513: 'JPEGInterchangeFormat',
            514: 'JPEGInterchangeFormatLength',
            515: 'JPEGRestartInterval',
            517: 'JPEGLosslessPredictors',
            518: 'JPEGPointTransforms',
            519: 'JPEGQTables',
            520: 'JPEGDCTables',
            521: 'JPEGACTables',
            529: 'YCbCrCoefficients',
            530: 'YCbCrSubSampling',
            531: 'YCbCrPositioning',
            532: 'ReferenceBlackWhite',
            559: 'StripRowCounts',
            700: 'XMP',  # XMLPacket
            769: 'GDIGamma',  # GDI+
            770: 'ICCProfileDescriptor',  # GDI+
            771: 'SRGBRenderingIntent',  # GDI+
            800: 'ImageTitle',  # GDI+
            999: 'USPTO_Miscellaneous',
            4864: 'AndorId',  # TODO: Andor Technology 4864 - 5030
            4869: 'AndorTemperature',
            4876: 'AndorExposureTime',
            4878: 'AndorKineticCycleTime',
            4879: 'AndorAccumulations',
            4881: 'AndorAcquisitionCycleTime',
            4882: 'AndorReadoutTime',
            4884: 'AndorPhotonCounting',
            4885: 'AndorEmDacLevel',
            4890: 'AndorFrames',
            4896: 'AndorHorizontalFlip',
            4897: 'AndorVerticalFlip',
            4898: 'AndorClockwise',
            4899: 'AndorCounterClockwise',
            4904: 'AndorVerticalClockVoltage',
            4905: 'AndorVerticalShiftSpeed',
            4907: 'AndorPreAmpSetting',
            4908: 'AndorCameraSerial',
            4911: 'AndorActualTemperature',
            4912: 'AndorBaselineClamp',
            4913: 'AndorPrescans',
            4914: 'AndorModel',
            4915: 'AndorChipSizeX',
            4916: 'AndorChipSizeY',
            4944: 'AndorBaselineOffset',
            4966: 'AndorSoftwareVersion',
            18246: 'Rating',
            18247: 'XP_DIP_XML',
            18248: 'StitchInfo',
            18249: 'RatingPercent',
            20481: 'ResolutionXUnit',  # GDI+
            20482: 'ResolutionYUnit',  # GDI+
            20483: 'ResolutionXLengthUnit',  # GDI+
            20484: 'ResolutionYLengthUnit',  # GDI+
            20485: 'PrintFlags',  # GDI+
            20486: 'PrintFlagsVersion',  # GDI+
            20487: 'PrintFlagsCrop',  # GDI+
            20488: 'PrintFlagsBleedWidth',  # GDI+
            20489: 'PrintFlagsBleedWidthScale',  # GDI+
            20490: 'HalftoneLPI',  # GDI+
            20491: 'HalftoneLPIUnit',  # GDI+
            20492: 'HalftoneDegree',  # GDI+
            20493: 'HalftoneShape',  # GDI+
            20494: 'HalftoneMisc',  # GDI+
            20495: 'HalftoneScreen',  # GDI+
            20496: 'JPEGQuality',  # GDI+
            20497: 'GridSize',  # GDI+
            20498: 'ThumbnailFormat',  # GDI+
            20499: 'ThumbnailWidth',  # GDI+
            20500: 'ThumbnailHeight',  # GDI+
            20501: 'ThumbnailColorDepth',  # GDI+
            20502: 'ThumbnailPlanes',  # GDI+
            20503: 'ThumbnailRawBytes',  # GDI+
            20504: 'ThumbnailSize',  # GDI+
            20505: 'ThumbnailCompressedSize',  # GDI+
            20506: 'ColorTransferFunction',  # GDI+
            20507: 'ThumbnailData',
            20512: 'ThumbnailImageWidth',  # GDI+
            20513: 'ThumbnailImageHeight',  # GDI+
            20514: 'ThumbnailBitsPerSample',  # GDI+
            20515: 'ThumbnailCompression',
            20516: 'ThumbnailPhotometricInterp',  # GDI+
            20517: 'ThumbnailImageDescription',  # GDI+
            20518: 'ThumbnailEquipMake',  # GDI+
            20519: 'ThumbnailEquipModel',  # GDI+
            20520: 'ThumbnailStripOffsets',  # GDI+
            20521: 'ThumbnailOrientation',  # GDI+
            20522: 'ThumbnailSamplesPerPixel',  # GDI+
            20523: 'ThumbnailRowsPerStrip',  # GDI+
            20524: 'ThumbnailStripBytesCount',  # GDI+
            20525: 'ThumbnailResolutionX',
            20526: 'ThumbnailResolutionY',
            20527: 'ThumbnailPlanarConfig',  # GDI+
            20528: 'ThumbnailResolutionUnit',
            20529: 'ThumbnailTransferFunction',
            20530: 'ThumbnailSoftwareUsed',  # GDI+
            20531: 'ThumbnailDateTime',  # GDI+
            20532: 'ThumbnailArtist',  # GDI+
            20533: 'ThumbnailWhitePoint',  # GDI+
            20534: 'ThumbnailPrimaryChromaticities',  # GDI+
            20535: 'ThumbnailYCbCrCoefficients',  # GDI+
            20536: 'ThumbnailYCbCrSubsampling',  # GDI+
            20537: 'ThumbnailYCbCrPositioning',
            20538: 'ThumbnailRefBlackWhite',  # GDI+
            20539: 'ThumbnailCopyRight',  # GDI+
            20545: 'InteroperabilityIndex',
            20546: 'InteroperabilityVersion',
            20624: 'LuminanceTable',
            20625: 'ChrominanceTable',
            20736: 'FrameDelay',  # GDI+
            20737: 'LoopCount',  # GDI+
            20738: 'GlobalPalette',  # GDI+
            20739: 'IndexBackground',  # GDI+
            20740: 'IndexTransparent',  # GDI+
            20752: 'PixelUnit',  # GDI+
            20753: 'PixelPerUnitX',  # GDI+
            20754: 'PixelPerUnitY',  # GDI+
            20755: 'PaletteHistogram',  # GDI+
            28672: 'SonyRawFileType',  # Sony ARW
            28722: 'VignettingCorrParams',  # Sony ARW
            28725: 'ChromaticAberrationCorrParams',  # Sony ARW
            28727: 'DistortionCorrParams',  # Sony ARW
            # Private tags >= 32768
            32781: 'ImageID',
            32931: 'WangTag1',
            32932: 'WangAnnotation',
            32933: 'WangTag3',
            32934: 'WangTag4',
            32953: 'ImageReferencePoints',
            32954: 'RegionXformTackPoint',
            32955: 'WarpQuadrilateral',
            32956: 'AffineTransformMat',
            32995: 'Matteing',
            32996: 'DataType',  # use SampleFormat
            32997: 'ImageDepth',
            32998: 'TileDepth',
            33300: 'ImageFullWidth',
            33301: 'ImageFullLength',
            33302: 'TextureFormat',
            33303: 'TextureWrapModes',
            33304: 'FieldOfViewCotangent',
            33305: 'MatrixWorldToScreen',
            33306: 'MatrixWorldToCamera',
            33405: 'Model2',
            33421: 'CFARepeatPatternDim',
            33422: 'CFAPattern',
            33423: 'BatteryLevel',
            33424: 'KodakIFD',
            33434: 'ExposureTime',
            33437: 'FNumber',
            33432: 'Copyright',
            33445: 'MDFileTag',
            33446: 'MDScalePixel',
            33447: 'MDColorTable',
            33448: 'MDLabName',
            33449: 'MDSampleInfo',
            33450: 'MDPrepDate',
            33451: 'MDPrepTime',
            33452: 'MDFileUnits',
            33471: 'OlympusINI',
            33550: 'ModelPixelScaleTag',
            33560: 'OlympusSIS',  # see also 33471 and 34853
            33589: 'AdventScale',
            33590: 'AdventRevision',
            33628: 'UIC1tag',  # Metamorph  Universal Imaging Corp STK
            33629: 'UIC2tag',
            33630: 'UIC3tag',
            33631: 'UIC4tag',
            33723: 'IPTCNAA',
            33858: 'ExtendedTagsOffset',  # DEFF points IFD with private tags
            33918: 'IntergraphPacketData',  # INGRPacketDataTag
            33919: 'IntergraphFlagRegisters',  # INGRFlagRegisters
            33920: 'IntergraphMatrixTag',  # IrasBTransformationMatrix
            33921: 'INGRReserved',
            33922: 'ModelTiepointTag',
            33923: 'LeicaMagic',
            34016: 'Site',  # 34016..34032 ANSI IT8 TIFF/IT
            34017: 'ColorSequence',
            34018: 'IT8Header',
            34019: 'RasterPadding',
            34020: 'BitsPerRunLength',
            34021: 'BitsPerExtendedRunLength',
            34022: 'ColorTable',
            34023: 'ImageColorIndicator',
            34024: 'BackgroundColorIndicator',
            34025: 'ImageColorValue',
            34026: 'BackgroundColorValue',
            34027: 'PixelIntensityRange',
            34028: 'TransparencyIndicator',
            34029: 'ColorCharacterization',
            34030: 'HCUsage',
            34031: 'TrapIndicator',
            34032: 'CMYKEquivalent',
            34118: 'CZ_SEM',  # Zeiss SEM
            34152: 'AFCP_IPTC',
            34232: 'PixelMagicJBIGOptions',  # EXIF, also TI FrameCount
            34263: 'JPLCartoIFD',
            34122: 'IPLAB',  # number of images
            34264: 'ModelTransformationTag',
            34306: 'WB_GRGBLevels',  # Leaf MOS
            34310: 'LeafData',
            34361: 'MM_Header',
            34362: 'MM_Stamp',
            34363: 'MM_Unknown',
            34377: 'ImageResources',  # Photoshop
            34386: 'MM_UserBlock',
            34412: 'CZ_LSMINFO',
            34665: 'ExifTag',
            34675: 'InterColorProfile',  # ICCProfile
            34680: 'FEI_SFEG',  #
            34682: 'FEI_HELIOS',  #
            34683: 'FEI_TITAN',  #
            34687: 'FXExtensions',
            34688: 'MultiProfiles',
            34689: 'SharedData',
            34690: 'T88Options',
            34710: 'MarCCD',  # offset to MarCCD header
            34732: 'ImageLayer',
            34735: 'GeoKeyDirectoryTag',
            34736: 'GeoDoubleParamsTag',
            34737: 'GeoAsciiParamsTag',
            34750: 'JBIGOptions',
            34821: 'PIXTIFF',  # ? Pixel Translations Inc
            34850: 'ExposureProgram',
            34852: 'SpectralSensitivity',
            34853: 'GPSTag',  # GPSIFD  also OlympusSIS2
            34855: 'ISOSpeedRatings',
            34856: 'OECF',
            34857: 'Interlace',
            34858: 'TimeZoneOffset',
            34859: 'SelfTimerMode',
            34864: 'SensitivityType',
            34865: 'StandardOutputSensitivity',
            34866: 'RecommendedExposureIndex',
            34867: 'ISOSpeed',
            34868: 'ISOSpeedLatitudeyyy',
            34869: 'ISOSpeedLatitudezzz',
            34908: 'HylaFAXFaxRecvParams',
            34909: 'HylaFAXFaxSubAddress',
            34910: 'HylaFAXFaxRecvTime',
            34911: 'FaxDcs',
            34929: 'FedexEDR',
            34954: 'LeafSubIFD',
            34959: 'Aphelion1',
            34960: 'Aphelion2',
            34961: 'AphelionInternal',  # ADCIS
            36864: 'ExifVersion',
            36867: 'DateTimeOriginal',
            36868: 'DateTimeDigitized',
            36873: 'GooglePlusUploadCode',
            36880: 'OffsetTime',
            36881: 'OffsetTimeOriginal',
            36882: 'OffsetTimeDigitized',
            # TODO: Pilatus/CHESS/TV6 36864..37120 conflicting with Exif tags
            # 36864: 'TVX ?',
            # 36865: 'TVX_NumExposure',
            # 36866: 'TVX_NumBackground',
            # 36867: 'TVX_ExposureTime',
            # 36868: 'TVX_BackgroundTime',
            # 36870: 'TVX ?',
            # 36873: 'TVX_SubBpp',
            # 36874: 'TVX_SubWide',
            # 36875: 'TVX_SubHigh',
            # 36876: 'TVX_BlackLevel',
            # 36877: 'TVX_DarkCurrent',
            # 36878: 'TVX_ReadNoise',
            # 36879: 'TVX_DarkCurrentNoise',
            # 36880: 'TVX_BeamMonitor',
            # 37120: 'TVX_UserVariables',  # A/D values
            37121: 'ComponentsConfiguration',
            37122: 'CompressedBitsPerPixel',
            37377: 'ShutterSpeedValue',
            37378: 'ApertureValue',
            37379: 'BrightnessValue',
            37380: 'ExposureBiasValue',
            37381: 'MaxApertureValue',
            37382: 'SubjectDistance',
            37383: 'MeteringMode',
            37384: 'LightSource',
            37385: 'Flash',
            37386: 'FocalLength',
            37387: 'FlashEnergy_',  # 37387
            37388: 'SpatialFrequencyResponse_',  # 37388
            37389: 'Noise',
            37390: 'FocalPlaneXResolution',
            37391: 'FocalPlaneYResolution',
            37392: 'FocalPlaneResolutionUnit',
            37393: 'ImageNumber',
            37394: 'SecurityClassification',
            37395: 'ImageHistory',
            37396: 'SubjectLocation',
            37397: 'ExposureIndex',
            37398: 'TIFFEPStandardID',
            37399: 'SensingMethod',
            37434: 'CIP3DataFile',
            37435: 'CIP3Sheet',
            37436: 'CIP3Side',
            37439: 'StoNits',
            37500: 'MakerNote',
            37510: 'UserComment',
            37520: 'SubsecTime',
            37521: 'SubsecTimeOriginal',
            37522: 'SubsecTimeDigitized',
            37679: 'MODIText',  # Microsoft Office Document Imaging
            37680: 'MODIOLEPropertySetStorage',
            37681: 'MODIPositioning',
            37706: 'TVIPS',  # offset to TemData structure
            37707: 'TVIPS1',
            37708: 'TVIPS2',  # same TemData structure as undefined
            37724: 'ImageSourceData',  # Photoshop
            37888: 'Temperature',
            37889: 'Humidity',
            37890: 'Pressure',
            37891: 'WaterDepth',
            37892: 'Acceleration',
            37893: 'CameraElevationAngle',
            40001: 'MC_IpWinScal',  # Media Cybernetics
            # 40001: 'RecipName',  # MS FAX
            40002: 'RecipNumber',
            40003: 'SenderName',
            40004: 'Routing',
            40005: 'CallerId',
            40006: 'TSID',
            40007: 'CSID',
            40008: 'FaxTime',
            40100: 'MC_IdOld',
            40106: 'MC_Unknown',
            40965: 'InteroperabilityTag',  # InteropOffset
            40091: 'XPTitle',
            40092: 'XPComment',
            40093: 'XPAuthor',
            40094: 'XPKeywords',
            40095: 'XPSubject',
            40960: 'FlashpixVersion',
            40961: 'ColorSpace',
            40962: 'PixelXDimension',
            40963: 'PixelYDimension',
            40964: 'RelatedSoundFile',
            40976: 'SamsungRawPointersOffset',
            40977: 'SamsungRawPointersLength',
            41217: 'SamsungRawByteOrder',
            41218: 'SamsungRawUnknown',
            41483: 'FlashEnergy',
            41484: 'SpatialFrequencyResponse',
            41485: 'Noise_',  # 37389
            41486: 'FocalPlaneXResolution_',  # 37390
            41487: 'FocalPlaneYResolution_',  # 37391
            41488: 'FocalPlaneResolutionUnit_',  # 37392
            41489: 'ImageNumber_',  # 37393
            41490: 'SecurityClassification_',  # 37394
            41491: 'ImageHistory_',  # 37395
            41492: 'SubjectLocation_',  # 37395
            41493: 'ExposureIndex_ ',  # 37397
            41494: 'TIFF-EPStandardID',
            41495: 'SensingMethod_',  # 37399
            41728: 'FileSource',
            41729: 'SceneType',
            41730: 'CFAPattern_',  # 33422
            41985: 'CustomRendered',
            41986: 'ExposureMode',
            41987: 'WhiteBalance',
            41988: 'DigitalZoomRatio',
            41989: 'FocalLengthIn35mmFilm',
            41990: 'SceneCaptureType',
            41991: 'GainControl',
            41992: 'Contrast',
            41993: 'Saturation',
            41994: 'Sharpness',
            41995: 'DeviceSettingDescription',
            41996: 'SubjectDistanceRange',
            42016: 'ImageUniqueID',
            42032: 'CameraOwnerName',
            42033: 'BodySerialNumber',
            42034: 'LensSpecification',
            42035: 'LensMake',
            42036: 'LensModel',
            42037: 'LensSerialNumber',
            42112: 'GDAL_METADATA',
            42113: 'GDAL_NODATA',
            42240: 'Gamma',
            43314: 'NIHImageHeader',
            44992: 'ExpandSoftware',
            44993: 'ExpandLens',
            44994: 'ExpandFilm',
            44995: 'ExpandFilterLens',
            44996: 'ExpandScanner',
            44997: 'ExpandFlashLamp',
            48129: 'PixelFormat',  # HDP and WDP
            48130: 'Transformation',
            48131: 'Uncompressed',
            48132: 'ImageType',
            48256: 'ImageWidth_',  # 256
            48257: 'ImageHeight_',
            48258: 'WidthResolution',
            48259: 'HeightResolution',
            48320: 'ImageOffset',
            48321: 'ImageByteCount',
            48322: 'AlphaOffset',
            48323: 'AlphaByteCount',
            48324: 'ImageDataDiscard',
            48325: 'AlphaDataDiscard',
            50003: 'KodakAPP3',
            50215: 'OceScanjobDescription',
            50216: 'OceApplicationSelector',
            50217: 'OceIdentificationNumber',
            50218: 'OceImageLogicCharacteristics',
            50255: 'Annotations',
            50288: 'MC_Id',  # Media Cybernetics
            50289: 'MC_XYPosition',
            50290: 'MC_ZPosition',
            50291: 'MC_XYCalibration',
            50292: 'MC_LensCharacteristics',
            50293: 'MC_ChannelName',
            50294: 'MC_ExcitationWavelength',
            50295: 'MC_TimeStamp',
            50296: 'MC_FrameProperties',
            50341: 'PrintImageMatching',
            50495: 'PCO_RAW',  # TODO: PCO CamWare
            50547: 'OriginalFileName',
            50560: 'USPTO_OriginalContentType',  # US Patent Office
            50561: 'USPTO_RotationCode',
            50648: 'CR2Unknown1',
            50649: 'CR2Unknown2',
            50656: 'CR2CFAPattern',
            50674: 'LercParameters',  # ESGI 50674 .. 50677
            50706: 'DNGVersion',  # DNG 50706 .. 51112
            50707: 'DNGBackwardVersion',
            50708: 'UniqueCameraModel',
            50709: 'LocalizedCameraModel',
            50710: 'CFAPlaneColor',
            50711: 'CFALayout',
            50712: 'LinearizationTable',
            50713: 'BlackLevelRepeatDim',
            50714: 'BlackLevel',
            50715: 'BlackLevelDeltaH',
            50716: 'BlackLevelDeltaV',
            50717: 'WhiteLevel',
            50718: 'DefaultScale',
            50719: 'DefaultCropOrigin',
            50720: 'DefaultCropSize',
            50721: 'ColorMatrix1',
            50722: 'ColorMatrix2',
            50723: 'CameraCalibration1',
            50724: 'CameraCalibration2',
            50725: 'ReductionMatrix1',
            50726: 'ReductionMatrix2',
            50727: 'AnalogBalance',
            50728: 'AsShotNeutral',
            50729: 'AsShotWhiteXY',
            50730: 'BaselineExposure',
            50731: 'BaselineNoise',
            50732: 'BaselineSharpness',
            50733: 'BayerGreenSplit',
            50734: 'LinearResponseLimit',
            50735: 'CameraSerialNumber',
            50736: 'LensInfo',
            50737: 'ChromaBlurRadius',
            50738: 'AntiAliasStrength',
            50739: 'ShadowScale',
            50740: 'DNGPrivateData',
            50741: 'MakerNoteSafety',
            50752: 'RawImageSegmentation',
            50778: 'CalibrationIlluminant1',
            50779: 'CalibrationIlluminant2',
            50780: 'BestQualityScale',
            50781: 'RawDataUniqueID',
            50784: 'AliasLayerMetadata',
            50827: 'OriginalRawFileName',
            50828: 'OriginalRawFileData',
            50829: 'ActiveArea',
            50830: 'MaskedAreas',
            50831: 'AsShotICCProfile',
            50832: 'AsShotPreProfileMatrix',
            50833: 'CurrentICCProfile',
            50834: 'CurrentPreProfileMatrix',
            50838: 'IJMetadataByteCounts',
            50839: 'IJMetadata',
            50844: 'RPCCoefficientTag',
            50879: 'ColorimetricReference',
            50885: 'SRawType',
            50898: 'PanasonicTitle',
            50899: 'PanasonicTitle2',
            50908: 'RSID',  # DGIWG
            50909: 'GEO_METADATA',  # DGIWG XML
            50931: 'CameraCalibrationSignature',
            50932: 'ProfileCalibrationSignature',
            50933: 'ProfileIFD',
            50934: 'AsShotProfileName',
            50935: 'NoiseReductionApplied',
            50936: 'ProfileName',
            50937: 'ProfileHueSatMapDims',
            50938: 'ProfileHueSatMapData1',
            50939: 'ProfileHueSatMapData2',
            50940: 'ProfileToneCurve',
            50941: 'ProfileEmbedPolicy',
            50942: 'ProfileCopyright',
            50964: 'ForwardMatrix1',
            50965: 'ForwardMatrix2',
            50966: 'PreviewApplicationName',
            50967: 'PreviewApplicationVersion',
            50968: 'PreviewSettingsName',
            50969: 'PreviewSettingsDigest',
            50970: 'PreviewColorSpace',
            50971: 'PreviewDateTime',
            50972: 'RawImageDigest',
            50973: 'OriginalRawFileDigest',
            50974: 'SubTileBlockSize',
            50975: 'RowInterleaveFactor',
            50981: 'ProfileLookTableDims',
            50982: 'ProfileLookTableData',
            51008: 'OpcodeList1',
            51009: 'OpcodeList2',
            51022: 'OpcodeList3',
            51023: 'FibicsXML',  #
            51041: 'NoiseProfile',
            51043: 'TimeCodes',
            51044: 'FrameRate',
            51058: 'TStop',
            51081: 'ReelName',
            51089: 'OriginalDefaultFinalSize',
            51090: 'OriginalBestQualitySize',
            51091: 'OriginalDefaultCropSize',
            51105: 'CameraLabel',
            51107: 'ProfileHueSatMapEncoding',
            51108: 'ProfileLookTableEncoding',
            51109: 'BaselineExposureOffset',
            51110: 'DefaultBlackRender',
            51111: 'NewRawImageDigest',
            51112: 'RawToPreviewGain',
            51125: 'DefaultUserCrop',
            51123: 'MicroManagerMetadata',
            51159: 'ZIFmetadata',  # Objective Pathology Services
            51160: 'ZIFannotations',  # Objective Pathology Services
            59932: 'Padding',
            59933: 'OffsetSchema',
            # Reusable Tags 65000-65535
            # 65000:  Dimap_Document XML
            # 65000-65112:  Photoshop Camera RAW EXIF tags
            # 65000: 'OwnerName',
            # 65001: 'SerialNumber',
            # 65002: 'Lens',
            # 65024: 'KDC_IFD',
            # 65100: 'RawFile',
            # 65101: 'Converter',
            # 65102: 'WhiteBalance',
            # 65105: 'Exposure',
            # 65106: 'Shadows',
            # 65107: 'Brightness',
            # 65108: 'Contrast',
            # 65109: 'Saturation',
            # 65110: 'Sharpness',
            # 65111: 'Smoothness',
            # 65112: 'MoireFilter',
            65200: 'FlexXML',
        }

    def TAG_NAMES():
        return {v: c for c, v in TIFF.TAGS.items()}

    def TAG_READERS():
        # Map TIFF tag codes to import functions
        return {
            320: read_colormap,
            # 700: read_bytes,  # read_utf8,
            # 34377: read_bytes,
            33723: read_bytes,
            # 34675: read_bytes,
            33628: read_uic1tag,  # Universal Imaging Corp STK
            33629: read_uic2tag,
            33630: read_uic3tag,
            33631: read_uic4tag,
            34118: read_cz_sem,  # Carl Zeiss SEM
            34361: read_mm_header,  # Olympus FluoView
            34362: read_mm_stamp,
            34363: read_numpy,  # MM_Unknown
            34386: read_numpy,  # MM_UserBlock
            34412: read_cz_lsminfo,  # Carl Zeiss LSM
            34680: read_fei_metadata,  # S-FEG
            34682: read_fei_metadata,  # Helios NanoLab
            37706: read_tvips_header,  # TVIPS EMMENU
            37724: read_bytes,  # ImageSourceData
            33923: read_bytes,  # read_leica_magic
            43314: read_nih_image_header,
            # 40001: read_bytes,
            40100: read_bytes,
            50288: read_bytes,
            50296: read_bytes,
            50839: read_bytes,
            51123: read_json,
            33471: read_sis_ini,
            33560: read_sis,
            34665: read_exif_ifd,
            34853: read_gps_ifd,  # conflicts with OlympusSIS
            40965: read_interoperability_ifd,
        }

    def TAG_TUPLE():
        # Tags whose values must be stored as tuples
        return frozenset((273, 279, 324, 325, 330, 530, 531, 34736))

    def TAG_ATTRIBUTES():
        #  Map tag codes to TiffPage attribute names
        return {
            'ImageWidth': 'imagewidth',
            'ImageLength': 'imagelength',
            'BitsPerSample': 'bitspersample',
            'Compression': 'compression',
            'PlanarConfiguration': 'planarconfig',
            'FillOrder': 'fillorder',
            'PhotometricInterpretation': 'photometric',
            'ColorMap': 'colormap',
            'ImageDescription': 'description',
            'ImageDescription1': 'description1',
            'SamplesPerPixel': 'samplesperpixel',
            'RowsPerStrip': 'rowsperstrip',
            'Software': 'software',
            'Predictor': 'predictor',
            'TileWidth': 'tilewidth',
            'TileLength': 'tilelength',
            'ExtraSamples': 'extrasamples',
            'SampleFormat': 'sampleformat',
            'ImageDepth': 'imagedepth',
            'TileDepth': 'tiledepth',
            'NewSubfileType': 'subfiletype',
        }

    def TAG_ENUM():
        return {
            # 254: TIFF.FILETYPE,
            255: TIFF.OFILETYPE,
            259: TIFF.COMPRESSION,
            262: TIFF.PHOTOMETRIC,
            263: TIFF.THRESHHOLD,
            266: TIFF.FILLORDER,
            274: TIFF.ORIENTATION,
            284: TIFF.PLANARCONFIG,
            290: TIFF.GRAYRESPONSEUNIT,
            # 292: TIFF.GROUP3OPT,
            # 293: TIFF.GROUP4OPT,
            296: TIFF.RESUNIT,
            300: TIFF.COLORRESPONSEUNIT,
            317: TIFF.PREDICTOR,
            338: TIFF.EXTRASAMPLE,
            339: TIFF.SAMPLEFORMAT,
            # 512: TIFF.JPEGPROC,
            # 531: TIFF.YCBCRPOSITION,
        }

    def FILETYPE():
        class FILETYPE(enum.IntFlag):
            # Python 3.6 only
            UNDEFINED = 0
            REDUCEDIMAGE = 1
            PAGE = 2
            MASK = 4
        return FILETYPE

    def OFILETYPE():
        class OFILETYPE(enum.IntEnum):
            UNDEFINED = 0
            IMAGE = 1
            REDUCEDIMAGE = 2
            PAGE = 3
        return OFILETYPE

    def COMPRESSION():
        class COMPRESSION(enum.IntEnum):
            NONE = 1  # Uncompressed
            CCITTRLE = 2  # CCITT 1D
            CCITT_T4 = 3  # 'T4/Group 3 Fax',
            CCITT_T6 = 4  # 'T6/Group 4 Fax',
            LZW = 5
            OJPEG = 6  # old-style JPEG
            JPEG = 7
            ADOBE_DEFLATE = 8
            JBIG_BW = 9
            JBIG_COLOR = 10
            JPEG_99 = 99
            KODAK_262 = 262
            NEXT = 32766
            SONY_ARW = 32767
            PACKED_RAW = 32769
            SAMSUNG_SRW = 32770
            CCIRLEW = 32771
            SAMSUNG_SRW2 = 32772
            PACKBITS = 32773
            THUNDERSCAN = 32809
            IT8CTPAD = 32895
            IT8LW = 32896
            IT8MP = 32897
            IT8BL = 32898
            PIXARFILM = 32908
            PIXARLOG = 32909
            DEFLATE = 32946
            DCS = 32947
            APERIO_JP2000_YCBC = 33003  # Leica Aperio
            APERIO_JP2000_RGB = 33005  # Leica Aperio
            JBIG = 34661
            SGILOG = 34676
            SGILOG24 = 34677
            JPEG2000 = 34712
            NIKON_NEF = 34713
            JBIG2 = 34715
            MDI_BINARY = 34718  # Microsoft Document Imaging
            MDI_PROGRESSIVE = 34719  # Microsoft Document Imaging
            MDI_VECTOR = 34720  # Microsoft Document Imaging
            LERC = 34887  # ESRI Lerc
            JPEG_LOSSY = 34892
            LZMA = 34925
            ZSTD_DEPRECATED = 34926
            WEBP_DEPRECATED = 34927
            OPS_PNG = 34933  # Objective Pathology Services
            OPS_JPEGXR = 34934  # Objective Pathology Services
            ZSTD = 50000
            WEBP = 50001
            PIXTIFF = 50013
            KODAK_DCR = 65000
            PENTAX_PEF = 65535
            # def __bool__(self): return self != 1  # Python 3.6+ only
        return COMPRESSION

    def PHOTOMETRIC():
        class PHOTOMETRIC(enum.IntEnum):
            MINISWHITE = 0
            MINISBLACK = 1
            RGB = 2
            PALETTE = 3
            MASK = 4
            SEPARATED = 5  # CMYK
            YCBCR = 6
            CIELAB = 8
            ICCLAB = 9
            ITULAB = 10
            CFA = 32803  # Color Filter Array
            LOGL = 32844
            LOGLUV = 32845
            LINEAR_RAW = 34892
        return PHOTOMETRIC

    def THRESHHOLD():
        class THRESHHOLD(enum.IntEnum):
            BILEVEL = 1
            HALFTONE = 2
            ERRORDIFFUSE = 3
        return THRESHHOLD

    def FILLORDER():
        class FILLORDER(enum.IntEnum):
            MSB2LSB = 1
            LSB2MSB = 2
        return FILLORDER

    def ORIENTATION():
        class ORIENTATION(enum.IntEnum):
            TOPLEFT = 1
            TOPRIGHT = 2
            BOTRIGHT = 3
            BOTLEFT = 4
            LEFTTOP = 5
            RIGHTTOP = 6
            RIGHTBOT = 7
            LEFTBOT = 8
        return ORIENTATION

    def PLANARCONFIG():
        class PLANARCONFIG(enum.IntEnum):
            CONTIG = 1
            SEPARATE = 2
        return PLANARCONFIG

    def GRAYRESPONSEUNIT():
        class GRAYRESPONSEUNIT(enum.IntEnum):
            _10S = 1
            _100S = 2
            _1000S = 3
            _10000S = 4
            _100000S = 5
        return GRAYRESPONSEUNIT

    def GROUP4OPT():
        class GROUP4OPT(enum.IntEnum):
            UNCOMPRESSED = 2
        return GROUP4OPT

    def RESUNIT():
        class RESUNIT(enum.IntEnum):
            NONE = 1
            INCH = 2
            CENTIMETER = 3
            # def __bool__(self): return self != 1  # Python 3.6 only
        return RESUNIT

    def COLORRESPONSEUNIT():
        class COLORRESPONSEUNIT(enum.IntEnum):
            _10S = 1
            _100S = 2
            _1000S = 3
            _10000S = 4
            _100000S = 5
        return COLORRESPONSEUNIT

    def PREDICTOR():
        class PREDICTOR(enum.IntEnum):
            NONE = 1
            HORIZONTAL = 2
            FLOATINGPOINT = 3
            # def __bool__(self): return self != 1  # Python 3.6 only
        return PREDICTOR

    def EXTRASAMPLE():
        class EXTRASAMPLE(enum.IntEnum):
            UNSPECIFIED = 0
            ASSOCALPHA = 1
            UNASSALPHA = 2
        return EXTRASAMPLE

    def SAMPLEFORMAT():
        class SAMPLEFORMAT(enum.IntEnum):
            UINT = 1
            INT = 2
            IEEEFP = 3
            VOID = 4
            COMPLEXINT = 5
            COMPLEXIEEEFP = 6
        return SAMPLEFORMAT

    def DATATYPES():
        class DATATYPES(enum.IntEnum):
            NOTYPE = 0
            BYTE = 1
            ASCII = 2
            SHORT = 3
            LONG = 4
            RATIONAL = 5
            SBYTE = 6
            UNDEFINED = 7
            SSHORT = 8
            SLONG = 9
            SRATIONAL = 10
            FLOAT = 11
            DOUBLE = 12
            IFD = 13
            UNICODE = 14
            COMPLEX = 15
            LONG8 = 16
            SLONG8 = 17
            IFD8 = 18
        return DATATYPES

    def DATA_FORMATS():
        # Map TIFF DATATYPES to Python struct formats
        return {
            1: '1B',   # BYTE 8-bit unsigned integer.
            2: '1s',   # ASCII 8-bit byte that contains a 7-bit ASCII code;
                       #   the last byte must be NULL (binary zero).
            3: '1H',   # SHORT 16-bit (2-byte) unsigned integer
            4: '1I',   # LONG 32-bit (4-byte) unsigned integer.
            5: '2I',   # RATIONAL Two LONGs: the first represents the numerator
                       #   of a fraction; the second, the denominator.
            6: '1b',   # SBYTE An 8-bit signed (twos-complement) integer.
            7: '1B',   # UNDEFINED An 8-bit byte that may contain anything,
                       #   depending on the definition of the field.
            8: '1h',   # SSHORT A 16-bit (2-byte) signed (twos-complement)
                       #   integer.
            9: '1i',   # SLONG A 32-bit (4-byte) signed (twos-complement)
                       #   integer.
            10: '2i',  # SRATIONAL Two SLONGs: the first represents the
                       #   numerator of a fraction, the second the denominator.
            11: '1f',  # FLOAT Single precision (4-byte) IEEE format.
            12: '1d',  # DOUBLE Double precision (8-byte) IEEE format.
            13: '1I',  # IFD unsigned 4 byte IFD offset.
            # 14: '',  # UNICODE
            # 15: '',  # COMPLEX
            16: '1Q',  # LONG8 unsigned 8 byte integer (BigTiff)
            17: '1q',  # SLONG8 signed 8 byte integer (BigTiff)
            18: '1Q',  # IFD8 unsigned 8 byte IFD offset (BigTiff)
        }

    def DATA_DTYPES():
        # Map numpy dtypes to TIFF DATATYPES
        return {'B': 1, 's': 2, 'H': 3, 'I': 4, '2I': 5, 'b': 6,
                'h': 8, 'i': 9, '2i': 10, 'f': 11, 'd': 12, 'Q': 16, 'q': 17}

    def SAMPLE_DTYPES():
        # Map TIFF SampleFormats and BitsPerSample to numpy dtype
        return {
            # UINT
            (1, 1): '?',  # bitmap
            (1, 2): 'B',
            (1, 3): 'B',
            (1, 4): 'B',
            (1, 5): 'B',
            (1, 6): 'B',
            (1, 7): 'B',
            (1, 8): 'B',
            (1, 9): 'H',
            (1, 10): 'H',
            (1, 11): 'H',
            (1, 12): 'H',
            (1, 13): 'H',
            (1, 14): 'H',
            (1, 15): 'H',
            (1, 16): 'H',
            (1, 17): 'I',
            (1, 18): 'I',
            (1, 19): 'I',
            (1, 20): 'I',
            (1, 21): 'I',
            (1, 22): 'I',
            (1, 23): 'I',
            (1, 24): 'I',
            (1, 25): 'I',
            (1, 26): 'I',
            (1, 27): 'I',
            (1, 28): 'I',
            (1, 29): 'I',
            (1, 30): 'I',
            (1, 31): 'I',
            (1, 32): 'I',
            (1, 64): 'Q',
            # VOID : treat as UINT
            (4, 1): '?',  # bitmap
            (4, 2): 'B',
            (4, 3): 'B',
            (4, 4): 'B',
            (4, 5): 'B',
            (4, 6): 'B',
            (4, 7): 'B',
            (4, 8): 'B',
            (4, 9): 'H',
            (4, 10): 'H',
            (4, 11): 'H',
            (4, 12): 'H',
            (4, 13): 'H',
            (4, 14): 'H',
            (4, 15): 'H',
            (4, 16): 'H',
            (4, 17): 'I',
            (4, 18): 'I',
            (4, 19): 'I',
            (4, 20): 'I',
            (4, 21): 'I',
            (4, 22): 'I',
            (4, 23): 'I',
            (4, 24): 'I',
            (4, 25): 'I',
            (4, 26): 'I',
            (4, 27): 'I',
            (4, 28): 'I',
            (4, 29): 'I',
            (4, 30): 'I',
            (4, 31): 'I',
            (4, 32): 'I',
            (4, 64): 'Q',
            # INT
            (2, 8): 'b',
            (2, 16): 'h',
            (2, 32): 'i',
            (2, 64): 'q',
            # IEEEFP : 24 bit not supported by numpy
            (3, 16): 'e',
            # (3, 24): '',  #
            (3, 32): 'f',
            (3, 64): 'd',
            # COMPLEXIEEEFP
            (6, 64): 'F',
            (6, 128): 'D',
            # RGB565
            (1, (5, 6, 5)): 'B',
            # COMPLEXINT : not supported by numpy
        }

    def PREDICTORS():
        # Map PREDICTOR to predictor encode functions
        if imagecodecs is None:
            return {
                None: identityfunc,
                1: identityfunc,
                2: delta_encode,
            }
        return {
            None: imagecodecs.none_encode,
            1: imagecodecs.none_encode,
            2: imagecodecs.delta_encode,
            3: imagecodecs.floatpred_encode,
        }

    def UNPREDICTORS():
        # Map PREDICTOR to predictor decode functions
        if imagecodecs is None:
            return {
                None: identityfunc,
                1: identityfunc,
                2: delta_decode,
            }
        return {
            None: imagecodecs.none_decode,
            1: imagecodecs.none_decode,
            2: imagecodecs.delta_decode,
            3: imagecodecs.floatpred_decode,
        }

    def COMPESSORS():
        # Map COMPRESSION to compress functions
        if hasattr(imagecodecs, 'zlib_encode'):
            return {
                None: imagecodecs.none_encode,
                1: imagecodecs.none_encode,
                8: imagecodecs.zlib_encode,
                32946: imagecodecs.zlib_encode,
                32773: imagecodecs.packbits_encode,
                34925: imagecodecs.lzma_encode,
                50000: imagecodecs.zstd_encode,
                50001: imagecodecs.webp_encode
            }

        def zlib_encode(data, level=6, out=None):
            """Compress Zlib DEFLATE."""
            return zlib.compress(data, level)

        if imagecodecs is None:
            return {
                None: identityfunc,
                1: identityfunc,
                8: zlib_encode,
                32946: zlib_encode,
                # 34925: lzma.compress
            }

        return {
            None: imagecodecs.none_encode,
            1: imagecodecs.none_encode,
            8: zlib_encode,
            32946: zlib_encode,
            32773: imagecodecs.packbits_encode,
            }

    def DECOMPESSORS():
        # Map COMPRESSION to decompress functions
        if hasattr(imagecodecs, 'zlib_decode'):
            return {
                None: imagecodecs.none_decode,
                1: imagecodecs.none_decode,
                5: imagecodecs.lzw_decode,
                6: imagecodecs.jpeg_decode,
                7: imagecodecs.jpeg_decode,
                8: imagecodecs.zlib_decode,
                32946: imagecodecs.zlib_decode,
                32773: imagecodecs.packbits_decode,
                # 34892: imagecodecs.jpeg_decode,  # DNG lossy
                34925: imagecodecs.lzma_decode,
                34926: imagecodecs.zstd_decode,  # deprecated
                34927: imagecodecs.webp_decode,  # deprecated
                33003: imagecodecs.j2k_decode,
                33005: imagecodecs.j2k_decode,
                34712: imagecodecs.j2k_decode,
                34933: imagecodecs.png_decode,
                34934: imagecodecs.jxr_decode,
                50000: imagecodecs.zstd_decode,
                50001: imagecodecs.webp_decode,
            }

        def zlib_decode(data, out=None):
            """Decompress Zlib DEFLATE."""
            return zlib.decompress(data)

        if imagecodecs is None:
            return {
                None: identityfunc,
                1: identityfunc,
                8: zlib_decode,
                32946: zlib_decode,
                # 34925: lzma.decompress
            }

        return {
            None: imagecodecs.none_decode,
            1: imagecodecs.none_decode,
            5: imagecodecs.lzw_decode,
            8: zlib_decode,
            32946: zlib_decode,
            32773: imagecodecs.packbits_decode,
        }

    def FRAME_ATTRS():
        # Attributes that a TiffFrame shares with its keyframe
        return set('shape ndim size dtype axes is_final'.split())

    def FILE_FLAGS():
        # TiffFile and TiffPage 'is_\*' attributes
        exclude = set('reduced mask final memmappable '
                      'contiguous tiled subsampled'.split())
        return set(a[3:] for a in dir(TiffPage)
                   if a[:3] == 'is_' and a[3:] not in exclude)

    def FILE_EXTENSIONS():
        # TIFF file extensions
        return tuple('tif tiff ome.tif lsm stk qpi pcoraw '
                     'gel seq svs zif ndpi bif tf8 tf2 btf'.split())

    def FILEOPEN_FILTER():
        # String for use in Windows File Open box
        return [('%s files' % ext.upper(), '*.%s' % ext)
                for ext in TIFF.FILE_EXTENSIONS] + [('allfiles', '*')]

    def AXES_LABELS():
        # TODO: is there a standard for character axes labels?
        axes = {
            'X': 'width',
            'Y': 'height',
            'Z': 'depth',
            'S': 'sample',  # rgb(a)
            'I': 'series',  # general sequence, plane, page, IFD
            'T': 'time',
            'C': 'channel',  # color, emission wavelength
            'A': 'angle',
            'P': 'phase',  # formerly F    # P is Position in LSM!
            'R': 'tile',  # region, point, mosaic
            'H': 'lifetime',  # histogram
            'E': 'lambda',  # excitation wavelength
            'L': 'exposure',  # lux
            'V': 'event',
            'Q': 'other',
            'M': 'mosaic',  # LSM 6
        }
        axes.update(dict((v, k) for k, v in axes.items()))
        return axes

    def NDPI_TAGS():
        # 65420 - 65458  Private Hamamatsu NDPI tags
        tags = dict((code, str(code)) for code in range(65420, 65459))
        tags.update({
            65420: 'FileFormat',
            65421: 'Magnification',  # SourceLens
            65422: 'XOffsetFromSlideCentre',
            65423: 'YOffsetFromSlideCentre',
            65424: 'ZOffsetFromSlideCentre',
            65427: 'UserLabel',
            65428: 'AuthCode',  # ?
            65442: 'ScannerSerialNumber',
            65449: 'Comments',
            65447: 'BlankLanes',
            65434: 'Fluorescence',
        })
        return tags

    def EXIF_TAGS():
        tags = {
            # 65000 - 65112  Photoshop Camera RAW EXIF tags
            65000: 'OwnerName',
            65001: 'SerialNumber',
            65002: 'Lens',
            65100: 'RawFile',
            65101: 'Converter',
            65102: 'WhiteBalance',
            65105: 'Exposure',
            65106: 'Shadows',
            65107: 'Brightness',
            65108: 'Contrast',
            65109: 'Saturation',
            65110: 'Sharpness',
            65111: 'Smoothness',
            65112: 'MoireFilter',
        }
        tags.update(TIFF.TAGS)
        return tags

    def GPS_TAGS():
        return {
            0: 'GPSVersionID',
            1: 'GPSLatitudeRef',
            2: 'GPSLatitude',
            3: 'GPSLongitudeRef',
            4: 'GPSLongitude',
            5: 'GPSAltitudeRef',
            6: 'GPSAltitude',
            7: 'GPSTimeStamp',
            8: 'GPSSatellites',
            9: 'GPSStatus',
            10: 'GPSMeasureMode',
            11: 'GPSDOP',
            12: 'GPSSpeedRef',
            13: 'GPSSpeed',
            14: 'GPSTrackRef',
            15: 'GPSTrack',
            16: 'GPSImgDirectionRef',
            17: 'GPSImgDirection',
            18: 'GPSMapDatum',
            19: 'GPSDestLatitudeRef',
            20: 'GPSDestLatitude',
            21: 'GPSDestLongitudeRef',
            22: 'GPSDestLongitude',
            23: 'GPSDestBearingRef',
            24: 'GPSDestBearing',
            25: 'GPSDestDistanceRef',
            26: 'GPSDestDistance',
            27: 'GPSProcessingMethod',
            28: 'GPSAreaInformation',
            29: 'GPSDateStamp',
            30: 'GPSDifferential',
            31: 'GPSHPositioningError',
        }

    def IOP_TAGS():
        return {
            1: 'InteroperabilityIndex',
            2: 'InteroperabilityVersion',
            4096: 'RelatedImageFileFormat',
            4097: 'RelatedImageWidth',
            4098: 'RelatedImageLength',
        }

    def GEO_KEYS():
        return {
            1024: 'GTModelTypeGeoKey',
            1025: 'GTRasterTypeGeoKey',
            1026: 'GTCitationGeoKey',
            2048: 'GeographicTypeGeoKey',
            2049: 'GeogCitationGeoKey',
            2050: 'GeogGeodeticDatumGeoKey',
            2051: 'GeogPrimeMeridianGeoKey',
            2052: 'GeogLinearUnitsGeoKey',
            2053: 'GeogLinearUnitSizeGeoKey',
            2054: 'GeogAngularUnitsGeoKey',
            2055: 'GeogAngularUnitsSizeGeoKey',
            2056: 'GeogEllipsoidGeoKey',
            2057: 'GeogSemiMajorAxisGeoKey',
            2058: 'GeogSemiMinorAxisGeoKey',
            2059: 'GeogInvFlatteningGeoKey',
            2060: 'GeogAzimuthUnitsGeoKey',
            2061: 'GeogPrimeMeridianLongGeoKey',
            2062: 'GeogTOWGS84GeoKey',
            3059: 'ProjLinearUnitsInterpCorrectGeoKey',  # GDAL
            3072: 'ProjectedCSTypeGeoKey',
            3073: 'PCSCitationGeoKey',
            3074: 'ProjectionGeoKey',
            3075: 'ProjCoordTransGeoKey',
            3076: 'ProjLinearUnitsGeoKey',
            3077: 'ProjLinearUnitSizeGeoKey',
            3078: 'ProjStdParallel1GeoKey',
            3079: 'ProjStdParallel2GeoKey',
            3080: 'ProjNatOriginLongGeoKey',
            3081: 'ProjNatOriginLatGeoKey',
            3082: 'ProjFalseEastingGeoKey',
            3083: 'ProjFalseNorthingGeoKey',
            3084: 'ProjFalseOriginLongGeoKey',
            3085: 'ProjFalseOriginLatGeoKey',
            3086: 'ProjFalseOriginEastingGeoKey',
            3087: 'ProjFalseOriginNorthingGeoKey',
            3088: 'ProjCenterLongGeoKey',
            3089: 'ProjCenterLatGeoKey',
            3090: 'ProjCenterEastingGeoKey',
            3091: 'ProjFalseOriginNorthingGeoKey',
            3092: 'ProjScaleAtNatOriginGeoKey',
            3093: 'ProjScaleAtCenterGeoKey',
            3094: 'ProjAzimuthAngleGeoKey',
            3095: 'ProjStraightVertPoleLongGeoKey',
            3096: 'ProjRectifiedGridAngleGeoKey',
            4096: 'VerticalCSTypeGeoKey',
            4097: 'VerticalCitationGeoKey',
            4098: 'VerticalDatumGeoKey',
            4099: 'VerticalUnitsGeoKey',
        }

    def GEO_CODES():
        try:
            from .tifffile_geodb import GEO_CODES  # delayed import
        except (ImportError, ValueError):
            try:
                from tifffile_geodb import GEO_CODES  # delayed import
            except (ImportError, ValueError):
                GEO_CODES = {}
        return GEO_CODES

    def CZ_LSMINFO():
        return [
            ('MagicNumber', 'u4'),
            ('StructureSize', 'i4'),
            ('DimensionX', 'i4'),
            ('DimensionY', 'i4'),
            ('DimensionZ', 'i4'),
            ('DimensionChannels', 'i4'),
            ('DimensionTime', 'i4'),
            ('DataType', 'i4'),  # DATATYPES
            ('ThumbnailX', 'i4'),
            ('ThumbnailY', 'i4'),
            ('VoxelSizeX', 'f8'),
            ('VoxelSizeY', 'f8'),
            ('VoxelSizeZ', 'f8'),
            ('OriginX', 'f8'),
            ('OriginY', 'f8'),
            ('OriginZ', 'f8'),
            ('ScanType', 'u2'),
            ('SpectralScan', 'u2'),
            ('TypeOfData', 'u4'),  # TYPEOFDATA
            ('OffsetVectorOverlay', 'u4'),
            ('OffsetInputLut', 'u4'),
            ('OffsetOutputLut', 'u4'),
            ('OffsetChannelColors', 'u4'),
            ('TimeIntervall', 'f8'),
            ('OffsetChannelDataTypes', 'u4'),
            ('OffsetScanInformation', 'u4'),  # SCANINFO
            ('OffsetKsData', 'u4'),
            ('OffsetTimeStamps', 'u4'),
            ('OffsetEventList', 'u4'),
            ('OffsetRoi', 'u4'),
            ('OffsetBleachRoi', 'u4'),
            ('OffsetNextRecording', 'u4'),
            # LSM 2.0 ends here
            ('DisplayAspectX', 'f8'),
            ('DisplayAspectY', 'f8'),
            ('DisplayAspectZ', 'f8'),
            ('DisplayAspectTime', 'f8'),
            ('OffsetMeanOfRoisOverlay', 'u4'),
            ('OffsetTopoIsolineOverlay', 'u4'),
            ('OffsetTopoProfileOverlay', 'u4'),
            ('OffsetLinescanOverlay', 'u4'),
            ('ToolbarFlags', 'u4'),
            ('OffsetChannelWavelength', 'u4'),
            ('OffsetChannelFactors', 'u4'),
            ('ObjectiveSphereCorrection', 'f8'),
            ('OffsetUnmixParameters', 'u4'),
            # LSM 3.2, 4.0 end here
            ('OffsetAcquisitionParameters', 'u4'),
            ('OffsetCharacteristics', 'u4'),
            ('OffsetPalette', 'u4'),
            ('TimeDifferenceX', 'f8'),
            ('TimeDifferenceY', 'f8'),
            ('TimeDifferenceZ', 'f8'),
            ('InternalUse1', 'u4'),
            ('DimensionP', 'i4'),
            ('DimensionM', 'i4'),
            ('DimensionsReserved', '16i4'),
            ('OffsetTilePositions', 'u4'),
            ('', '9u4'),  # Reserved
            ('OffsetPositions', 'u4'),
            # ('', '21u4'),  # must be 0
        ]

    def CZ_LSMINFO_READERS():
        # Import functions for CZ_LSMINFO sub-records
        # TODO: read more CZ_LSMINFO sub-records
        return {
            'ScanInformation': read_lsm_scaninfo,
            'TimeStamps': read_lsm_timestamps,
            'EventList': read_lsm_eventlist,
            'ChannelColors': read_lsm_channelcolors,
            'Positions': read_lsm_floatpairs,
            'TilePositions': read_lsm_floatpairs,
            'VectorOverlay': None,
            'InputLut': None,
            'OutputLut': None,
            'TimeIntervall': None,
            'ChannelDataTypes': None,
            'KsData': None,
            'Roi': None,
            'BleachRoi': None,
            'NextRecording': None,
            'MeanOfRoisOverlay': None,
            'TopoIsolineOverlay': None,
            'TopoProfileOverlay': None,
            'ChannelWavelength': None,
            'SphereCorrection': None,
            'ChannelFactors': None,
            'UnmixParameters': None,
            'AcquisitionParameters': None,
            'Characteristics': None,
        }

    def CZ_LSMINFO_SCANTYPE():
        # Map CZ_LSMINFO.ScanType to dimension order
        return {
            0: 'XYZCT',  # 'Stack' normal x-y-z-scan
            1: 'XYZCT',  # 'Z-Scan' x-z-plane Y=1
            2: 'XYZCT',  # 'Line'
            3: 'XYTCZ',  # 'Time Series Plane' time series x-y  XYCTZ ? Z=1
            4: 'XYZTC',  # 'Time Series z-Scan' time series x-z
            5: 'XYTCZ',  # 'Time Series Mean-of-ROIs'
            6: 'XYZTC',  # 'Time Series Stack' time series x-y-z
            7: 'XYCTZ',  # Spline Scan
            8: 'XYCZT',  # Spline Plane x-z
            9: 'XYTCZ',  # Time Series Spline Plane x-z
            10: 'XYZCT',  # 'Time Series Point' point mode
        }

    def CZ_LSMINFO_DIMENSIONS():
        # Map dimension codes to CZ_LSMINFO attribute
        return {
            'X': 'DimensionX',
            'Y': 'DimensionY',
            'Z': 'DimensionZ',
            'C': 'DimensionChannels',
            'T': 'DimensionTime',
            'P': 'DimensionP',
            'M': 'DimensionM',
        }

    def CZ_LSMINFO_DATATYPES():
        # Description of CZ_LSMINFO.DataType
        return {
            0: 'varying data types',
            1: '8 bit unsigned integer',
            2: '12 bit unsigned integer',
            5: '32 bit float',
        }

    def CZ_LSMINFO_TYPEOFDATA():
        # Description of CZ_LSMINFO.TypeOfData
        return {
            0: 'Original scan data',
            1: 'Calculated data',
            2: '3D reconstruction',
            3: 'Topography height map',
        }

    def CZ_LSMINFO_SCANINFO_ARRAYS():
        return {
            0x20000000: 'Tracks',
            0x30000000: 'Lasers',
            0x60000000: 'DetectionChannels',
            0x80000000: 'IlluminationChannels',
            0xa0000000: 'BeamSplitters',
            0xc0000000: 'DataChannels',
            0x11000000: 'Timers',
            0x13000000: 'Markers',
        }

    def CZ_LSMINFO_SCANINFO_STRUCTS():
        return {
            # 0x10000000: 'Recording',
            0x40000000: 'Track',
            0x50000000: 'Laser',
            0x70000000: 'DetectionChannel',
            0x90000000: 'IlluminationChannel',
            0xb0000000: 'BeamSplitter',
            0xd0000000: 'DataChannel',
            0x12000000: 'Timer',
            0x14000000: 'Marker',
        }

    def CZ_LSMINFO_SCANINFO_ATTRIBUTES():
        return {
            # Recording
            0x10000001: 'Name',
            0x10000002: 'Description',
            0x10000003: 'Notes',
            0x10000004: 'Objective',
            0x10000005: 'ProcessingSummary',
            0x10000006: 'SpecialScanMode',
            0x10000007: 'ScanType',
            0x10000008: 'ScanMode',
            0x10000009: 'NumberOfStacks',
            0x1000000a: 'LinesPerPlane',
            0x1000000b: 'SamplesPerLine',
            0x1000000c: 'PlanesPerVolume',
            0x1000000d: 'ImagesWidth',
            0x1000000e: 'ImagesHeight',
            0x1000000f: 'ImagesNumberPlanes',
            0x10000010: 'ImagesNumberStacks',
            0x10000011: 'ImagesNumberChannels',
            0x10000012: 'LinscanXySize',
            0x10000013: 'ScanDirection',
            0x10000014: 'TimeSeries',
            0x10000015: 'OriginalScanData',
            0x10000016: 'ZoomX',
            0x10000017: 'ZoomY',
            0x10000018: 'ZoomZ',
            0x10000019: 'Sample0X',
            0x1000001a: 'Sample0Y',
            0x1000001b: 'Sample0Z',
            0x1000001c: 'SampleSpacing',
            0x1000001d: 'LineSpacing',
            0x1000001e: 'PlaneSpacing',
            0x1000001f: 'PlaneWidth',
            0x10000020: 'PlaneHeight',
            0x10000021: 'VolumeDepth',
            0x10000023: 'Nutation',
            0x10000034: 'Rotation',
            0x10000035: 'Precession',
            0x10000036: 'Sample0time',
            0x10000037: 'StartScanTriggerIn',
            0x10000038: 'StartScanTriggerOut',
            0x10000039: 'StartScanEvent',
            0x10000040: 'StartScanTime',
            0x10000041: 'StopScanTriggerIn',
            0x10000042: 'StopScanTriggerOut',
            0x10000043: 'StopScanEvent',
            0x10000044: 'StopScanTime',
            0x10000045: 'UseRois',
            0x10000046: 'UseReducedMemoryRois',
            0x10000047: 'User',
            0x10000048: 'UseBcCorrection',
            0x10000049: 'PositionBcCorrection1',
            0x10000050: 'PositionBcCorrection2',
            0x10000051: 'InterpolationY',
            0x10000052: 'CameraBinning',
            0x10000053: 'CameraSupersampling',
            0x10000054: 'CameraFrameWidth',
            0x10000055: 'CameraFrameHeight',
            0x10000056: 'CameraOffsetX',
            0x10000057: 'CameraOffsetY',
            0x10000059: 'RtBinning',
            0x1000005a: 'RtFrameWidth',
            0x1000005b: 'RtFrameHeight',
            0x1000005c: 'RtRegionWidth',
            0x1000005d: 'RtRegionHeight',
            0x1000005e: 'RtOffsetX',
            0x1000005f: 'RtOffsetY',
            0x10000060: 'RtZoom',
            0x10000061: 'RtLinePeriod',
            0x10000062: 'Prescan',
            0x10000063: 'ScanDirectionZ',
            # Track
            0x40000001: 'MultiplexType',  # 0 After Line; 1 After Frame
            0x40000002: 'MultiplexOrder',
            0x40000003: 'SamplingMode',  # 0 Sample; 1 Line Avg; 2 Frame Avg
            0x40000004: 'SamplingMethod',  # 1 Mean; 2 Sum
            0x40000005: 'SamplingNumber',
            0x40000006: 'Acquire',
            0x40000007: 'SampleObservationTime',
            0x4000000b: 'TimeBetweenStacks',
            0x4000000c: 'Name',
            0x4000000d: 'Collimator1Name',
            0x4000000e: 'Collimator1Position',
            0x4000000f: 'Collimator2Name',
            0x40000010: 'Collimator2Position',
            0x40000011: 'IsBleachTrack',
            0x40000012: 'IsBleachAfterScanNumber',
            0x40000013: 'BleachScanNumber',
            0x40000014: 'TriggerIn',
            0x40000015: 'TriggerOut',
            0x40000016: 'IsRatioTrack',
            0x40000017: 'BleachCount',
            0x40000018: 'SpiCenterWavelength',
            0x40000019: 'PixelTime',
            0x40000021: 'CondensorFrontlens',
            0x40000023: 'FieldStopValue',
            0x40000024: 'IdCondensorAperture',
            0x40000025: 'CondensorAperture',
            0x40000026: 'IdCondensorRevolver',
            0x40000027: 'CondensorFilter',
            0x40000028: 'IdTransmissionFilter1',
            0x40000029: 'IdTransmission1',
            0x40000030: 'IdTransmissionFilter2',
            0x40000031: 'IdTransmission2',
            0x40000032: 'RepeatBleach',
            0x40000033: 'EnableSpotBleachPos',
            0x40000034: 'SpotBleachPosx',
            0x40000035: 'SpotBleachPosy',
            0x40000036: 'SpotBleachPosz',
            0x40000037: 'IdTubelens',
            0x40000038: 'IdTubelensPosition',
            0x40000039: 'TransmittedLight',
            0x4000003a: 'ReflectedLight',
            0x4000003b: 'SimultanGrabAndBleach',
            0x4000003c: 'BleachPixelTime',
            # Laser
            0x50000001: 'Name',
            0x50000002: 'Acquire',
            0x50000003: 'Power',
            # DetectionChannel
            0x70000001: 'IntegrationMode',
            0x70000002: 'SpecialMode',
            0x70000003: 'DetectorGainFirst',
            0x70000004: 'DetectorGainLast',
            0x70000005: 'AmplifierGainFirst',
            0x70000006: 'AmplifierGainLast',
            0x70000007: 'AmplifierOffsFirst',
            0x70000008: 'AmplifierOffsLast',
            0x70000009: 'PinholeDiameter',
            0x7000000a: 'CountingTrigger',
            0x7000000b: 'Acquire',
            0x7000000c: 'PointDetectorName',
            0x7000000d: 'AmplifierName',
            0x7000000e: 'PinholeName',
            0x7000000f: 'FilterSetName',
            0x70000010: 'FilterName',
            0x70000013: 'IntegratorName',
            0x70000014: 'ChannelName',
            0x70000015: 'DetectorGainBc1',
            0x70000016: 'DetectorGainBc2',
            0x70000017: 'AmplifierGainBc1',
            0x70000018: 'AmplifierGainBc2',
            0x70000019: 'AmplifierOffsetBc1',
            0x70000020: 'AmplifierOffsetBc2',
            0x70000021: 'SpectralScanChannels',
            0x70000022: 'SpiWavelengthStart',
            0x70000023: 'SpiWavelengthStop',
            0x70000026: 'DyeName',
            0x70000027: 'DyeFolder',
            # IlluminationChannel
            0x90000001: 'Name',
            0x90000002: 'Power',
            0x90000003: 'Wavelength',
            0x90000004: 'Aquire',
            0x90000005: 'DetchannelName',
            0x90000006: 'PowerBc1',
            0x90000007: 'PowerBc2',
            # BeamSplitter
            0xb0000001: 'FilterSet',
            0xb0000002: 'Filter',
            0xb0000003: 'Name',
            # DataChannel
            0xd0000001: 'Name',
            0xd0000003: 'Acquire',
            0xd0000004: 'Color',
            0xd0000005: 'SampleType',
            0xd0000006: 'BitsPerSample',
            0xd0000007: 'RatioType',
            0xd0000008: 'RatioTrack1',
            0xd0000009: 'RatioTrack2',
            0xd000000a: 'RatioChannel1',
            0xd000000b: 'RatioChannel2',
            0xd000000c: 'RatioConst1',
            0xd000000d: 'RatioConst2',
            0xd000000e: 'RatioConst3',
            0xd000000f: 'RatioConst4',
            0xd0000010: 'RatioConst5',
            0xd0000011: 'RatioConst6',
            0xd0000012: 'RatioFirstImages1',
            0xd0000013: 'RatioFirstImages2',
            0xd0000014: 'DyeName',
            0xd0000015: 'DyeFolder',
            0xd0000016: 'Spectrum',
            0xd0000017: 'Acquire',
            # Timer
            0x12000001: 'Name',
            0x12000002: 'Description',
            0x12000003: 'Interval',
            0x12000004: 'TriggerIn',
            0x12000005: 'TriggerOut',
            0x12000006: 'ActivationTime',
            0x12000007: 'ActivationNumber',
            # Marker
            0x14000001: 'Name',
            0x14000002: 'Description',
            0x14000003: 'TriggerIn',
            0x14000004: 'TriggerOut',
        }

    def NIH_IMAGE_HEADER():
        return [
            ('FileID', 'a8'),
            ('nLines', 'i2'),
            ('PixelsPerLine', 'i2'),
            ('Version', 'i2'),
            ('OldLutMode', 'i2'),
            ('OldnColors', 'i2'),
            ('Colors', 'u1', (3, 32)),
            ('OldColorStart', 'i2'),
            ('ColorWidth', 'i2'),
            ('ExtraColors', 'u2', (6, 3)),
            ('nExtraColors', 'i2'),
            ('ForegroundIndex', 'i2'),
            ('BackgroundIndex', 'i2'),
            ('XScale', 'f8'),
            ('Unused2', 'i2'),
            ('Unused3', 'i2'),
            ('UnitsID', 'i2'),  # NIH_UNITS_TYPE
            ('p1', [('x', 'i2'), ('y', 'i2')]),
            ('p2', [('x', 'i2'), ('y', 'i2')]),
            ('CurveFitType', 'i2'),  # NIH_CURVEFIT_TYPE
            ('nCoefficients', 'i2'),
            ('Coeff', 'f8', 6),
            ('UMsize', 'u1'),
            ('UM', 'a15'),
            ('UnusedBoolean', 'u1'),
            ('BinaryPic', 'b1'),
            ('SliceStart', 'i2'),
            ('SliceEnd', 'i2'),
            ('ScaleMagnification', 'f4'),
            ('nSlices', 'i2'),
            ('SliceSpacing', 'f4'),
            ('CurrentSlice', 'i2'),
            ('FrameInterval', 'f4'),
            ('PixelAspectRatio', 'f4'),
            ('ColorStart', 'i2'),
            ('ColorEnd', 'i2'),
            ('nColors', 'i2'),
            ('Fill1', '3u2'),
            ('Fill2', '3u2'),
            ('Table', 'u1'),  # NIH_COLORTABLE_TYPE
            ('LutMode', 'u1'),  # NIH_LUTMODE_TYPE
            ('InvertedTable', 'b1'),
            ('ZeroClip', 'b1'),
            ('XUnitSize', 'u1'),
            ('XUnit', 'a11'),
            ('StackType', 'i2'),  # NIH_STACKTYPE_TYPE
            # ('UnusedBytes', 'u1', 200)
        ]

    def NIH_COLORTABLE_TYPE():
        return ('CustomTable', 'AppleDefault', 'Pseudo20', 'Pseudo32',
                'Rainbow', 'Fire1', 'Fire2', 'Ice', 'Grays', 'Spectrum')

    def NIH_LUTMODE_TYPE():
        return ('PseudoColor', 'OldAppleDefault', 'OldSpectrum', 'GrayScale',
                'ColorLut', 'CustomGrayscale')

    def NIH_CURVEFIT_TYPE():
        return ('StraightLine', 'Poly2', 'Poly3', 'Poly4', 'Poly5', 'ExpoFit',
                'PowerFit', 'LogFit', 'RodbardFit', 'SpareFit1',
                'Uncalibrated', 'UncalibratedOD')

    def NIH_UNITS_TYPE():
        return ('Nanometers', 'Micrometers', 'Millimeters', 'Centimeters',
                'Meters', 'Kilometers', 'Inches', 'Feet', 'Miles', 'Pixels',
                'OtherUnits')

    def NIH_STACKTYPE_TYPE():
        return ('VolumeStack', 'RGBStack', 'MovieStack', 'HSVStack')

    def TVIPS_HEADER_V1():
        # TVIPS TemData structure from EMMENU Help file
        return [
            ('Version', 'i4'),
            ('CommentV1', 'a80'),
            ('HighTension', 'i4'),
            ('SphericalAberration', 'i4'),
            ('IlluminationAperture', 'i4'),
            ('Magnification', 'i4'),
            ('PostMagnification', 'i4'),
            ('FocalLength', 'i4'),
            ('Defocus', 'i4'),
            ('Astigmatism', 'i4'),
            ('AstigmatismDirection', 'i4'),
            ('BiprismVoltage', 'i4'),
            ('SpecimenTiltAngle', 'i4'),
            ('SpecimenTiltDirection', 'i4'),
            ('IlluminationTiltDirection', 'i4'),
            ('IlluminationTiltAngle', 'i4'),
            ('ImageMode', 'i4'),
            ('EnergySpread', 'i4'),
            ('ChromaticAberration', 'i4'),
            ('ShutterType', 'i4'),
            ('DefocusSpread', 'i4'),
            ('CcdNumber', 'i4'),
            ('CcdSize', 'i4'),
            ('OffsetXV1', 'i4'),
            ('OffsetYV1', 'i4'),
            ('PhysicalPixelSize', 'i4'),
            ('Binning', 'i4'),
            ('ReadoutSpeed', 'i4'),
            ('GainV1', 'i4'),
            ('SensitivityV1', 'i4'),
            ('ExposureTimeV1', 'i4'),
            ('FlatCorrected', 'i4'),
            ('DeadPxCorrected', 'i4'),
            ('ImageMean', 'i4'),
            ('ImageStd', 'i4'),
            ('DisplacementX', 'i4'),
            ('DisplacementY', 'i4'),
            ('DateV1', 'i4'),
            ('TimeV1', 'i4'),
            ('ImageMin', 'i4'),
            ('ImageMax', 'i4'),
            ('ImageStatisticsQuality', 'i4'),
        ]

    def TVIPS_HEADER_V2():
        return [
            ('ImageName', 'V160'),  # utf16
            ('ImageFolder', 'V160'),
            ('ImageSizeX', 'i4'),
            ('ImageSizeY', 'i4'),
            ('ImageSizeZ', 'i4'),
            ('ImageSizeE', 'i4'),
            ('ImageDataType', 'i4'),
            ('Date', 'i4'),
            ('Time', 'i4'),
            ('Comment', 'V1024'),
            ('ImageHistory', 'V1024'),
            ('Scaling', '16f4'),
            ('ImageStatistics', '16c16'),
            ('ImageType', 'i4'),
            ('ImageDisplaType', 'i4'),
            ('PixelSizeX', 'f4'),  # distance between two px in x, [nm]
            ('PixelSizeY', 'f4'),  # distance between two px in y, [nm]
            ('ImageDistanceZ', 'f4'),
            ('ImageDistanceE', 'f4'),
            ('ImageMisc', '32f4'),
            ('TemType', 'V160'),
            ('TemHighTension', 'f4'),
            ('TemAberrations', '32f4'),
            ('TemEnergy', '32f4'),
            ('TemMode', 'i4'),
            ('TemMagnification', 'f4'),
            ('TemMagnificationCorrection', 'f4'),
            ('PostMagnification', 'f4'),
            ('TemStageType', 'i4'),
            ('TemStagePosition', '5f4'),  # x, y, z, a, b
            ('TemImageShift', '2f4'),
            ('TemBeamShift', '2f4'),
            ('TemBeamTilt', '2f4'),
            ('TilingParameters', '7f4'),  # 0: tiling? 1:x 2:y 3: max x
                                          # 4: max y 5: overlap x 6: overlap y
            ('TemIllumination', '3f4'),  # 0: spotsize 1: intensity
            ('TemShutter', 'i4'),
            ('TemMisc', '32f4'),
            ('CameraType', 'V160'),
            ('PhysicalPixelSizeX', 'f4'),
            ('PhysicalPixelSizeY', 'f4'),
            ('OffsetX', 'i4'),
            ('OffsetY', 'i4'),
            ('BinningX', 'i4'),
            ('BinningY', 'i4'),
            ('ExposureTime', 'f4'),
            ('Gain', 'f4'),
            ('ReadoutRate', 'f4'),
            ('FlatfieldDescription', 'V160'),
            ('Sensitivity', 'f4'),
            ('Dose', 'f4'),
            ('CamMisc', '32f4'),
            ('FeiMicroscopeInformation', 'V1024'),
            ('FeiSpecimenInformation', 'V1024'),
            ('Magic', 'u4'),
        ]

    def MM_HEADER():
        # Olympus FluoView MM_Header
        MM_DIMENSION = [
            ('Name', 'a16'),
            ('Size', 'i4'),
            ('Origin', 'f8'),
            ('Resolution', 'f8'),
            ('Unit', 'a64')]
        return [
            ('HeaderFlag', 'i2'),
            ('ImageType', 'u1'),
            ('ImageName', 'a257'),
            ('OffsetData', 'u4'),
            ('PaletteSize', 'i4'),
            ('OffsetPalette0', 'u4'),
            ('OffsetPalette1', 'u4'),
            ('CommentSize', 'i4'),
            ('OffsetComment', 'u4'),
            ('Dimensions', MM_DIMENSION, 10),
            ('OffsetPosition', 'u4'),
            ('MapType', 'i2'),
            ('MapMin', 'f8'),
            ('MapMax', 'f8'),
            ('MinValue', 'f8'),
            ('MaxValue', 'f8'),
            ('OffsetMap', 'u4'),
            ('Gamma', 'f8'),
            ('Offset', 'f8'),
            ('GrayChannel', MM_DIMENSION),
            ('OffsetThumbnail', 'u4'),
            ('VoiceField', 'i4'),
            ('OffsetVoiceField', 'u4'),
        ]

    def MM_DIMENSIONS():
        # Map FluoView MM_Header.Dimensions to axes characters
        return {
            'X': 'X',
            'Y': 'Y',
            'Z': 'Z',
            'T': 'T',
            'CH': 'C',
            'WAVELENGTH': 'C',
            'TIME': 'T',
            'XY': 'R',
            'EVENT': 'V',
            'EXPOSURE': 'L',
        }

    def UIC_TAGS():
        # Map Universal Imaging Corporation MetaMorph internal tag ids to
        # name and type
        from fractions import Fraction  # delayed import

        return [
            ('AutoScale', int),
            ('MinScale', int),
            ('MaxScale', int),
            ('SpatialCalibration', int),
            ('XCalibration', Fraction),
            ('YCalibration', Fraction),
            ('CalibrationUnits', str),
            ('Name', str),
            ('ThreshState', int),
            ('ThreshStateRed', int),
            ('tagid_10', None),  # undefined
            ('ThreshStateGreen', int),
            ('ThreshStateBlue', int),
            ('ThreshStateLo', int),
            ('ThreshStateHi', int),
            ('Zoom', int),
            ('CreateTime', julian_datetime),
            ('LastSavedTime', julian_datetime),
            ('currentBuffer', int),
            ('grayFit', None),
            ('grayPointCount', None),
            ('grayX', Fraction),
            ('grayY', Fraction),
            ('grayMin', Fraction),
            ('grayMax', Fraction),
            ('grayUnitName', str),
            ('StandardLUT', int),
            ('wavelength', int),
            ('StagePosition', '(%i,2,2)u4'),  # N xy positions as fract
            ('CameraChipOffset', '(%i,2,2)u4'),  # N xy offsets as fract
            ('OverlayMask', None),
            ('OverlayCompress', None),
            ('Overlay', None),
            ('SpecialOverlayMask', None),
            ('SpecialOverlayCompress', None),
            ('SpecialOverlay', None),
            ('ImageProperty', read_uic_image_property),
            ('StageLabel', '%ip'),  # N str
            ('AutoScaleLoInfo', Fraction),
            ('AutoScaleHiInfo', Fraction),
            ('AbsoluteZ', '(%i,2)u4'),  # N fractions
            ('AbsoluteZValid', '(%i,)u4'),  # N long
            ('Gamma', 'I'),  # 'I' uses offset
            ('GammaRed', 'I'),
            ('GammaGreen', 'I'),
            ('GammaBlue', 'I'),
            ('CameraBin', '2I'),
            ('NewLUT', int),
            ('ImagePropertyEx', None),
            ('PlaneProperty', int),
            ('UserLutTable', '(256,3)u1'),
            ('RedAutoScaleInfo', int),
            ('RedAutoScaleLoInfo', Fraction),
            ('RedAutoScaleHiInfo', Fraction),
            ('RedMinScaleInfo', int),
            ('RedMaxScaleInfo', int),
            ('GreenAutoScaleInfo', int),
            ('GreenAutoScaleLoInfo', Fraction),
            ('GreenAutoScaleHiInfo', Fraction),
            ('GreenMinScaleInfo', int),
            ('GreenMaxScaleInfo', int),
            ('BlueAutoScaleInfo', int),
            ('BlueAutoScaleLoInfo', Fraction),
            ('BlueAutoScaleHiInfo', Fraction),
            ('BlueMinScaleInfo', int),
            ('BlueMaxScaleInfo', int),
            # ('OverlayPlaneColor', read_uic_overlay_plane_color),
        ]

    def PILATUS_HEADER():
        # PILATUS CBF Header Specification, Version 1.4
        # Map key to [value_indices], type
        return {
            'Detector': ([slice(1, None)], str),
            'Pixel_size': ([1, 4], float),
            'Silicon': ([3], float),
            'Exposure_time': ([1], float),
            'Exposure_period': ([1], float),
            'Tau': ([1], float),
            'Count_cutoff': ([1], int),
            'Threshold_setting': ([1], float),
            'Gain_setting': ([1, 2], str),
            'N_excluded_pixels': ([1], int),
            'Excluded_pixels': ([1], str),
            'Flat_field': ([1], str),
            'Trim_file': ([1], str),
            'Image_path': ([1], str),
            # optional
            'Wavelength': ([1], float),
            'Energy_range': ([1, 2], float),
            'Detector_distance': ([1], float),
            'Detector_Voffset': ([1], float),
            'Beam_xy': ([1, 2], float),
            'Flux': ([1], str),
            'Filter_transmission': ([1], float),
            'Start_angle': ([1], float),
            'Angle_increment': ([1], float),
            'Detector_2theta': ([1], float),
            'Polarization': ([1], float),
            'Alpha': ([1], float),
            'Kappa': ([1], float),
            'Phi': ([1], float),
            'Phi_increment': ([1], float),
            'Chi': ([1], float),
            'Chi_increment': ([1], float),
            'Oscillation_axis': ([slice(1, None)], str),
            'N_oscillations': ([1], int),
            'Start_position': ([1], float),
            'Position_increment': ([1], float),
            'Shutter_time': ([1], float),
            'Omega': ([1], float),
            'Omega_increment': ([1], float)
        }

    def ALLOCATIONGRANULARITY():
        # alignment for writing contiguous data to TIFF
        import mmap  # delayed import
        return mmap.ALLOCATIONGRANULARITY


def read_tags(fh, byteorder, offsetsize, tagnames, customtags=None,
              maxifds=None):
    """Read tags from chain of IFDs and return as list of dicts.

    The file handle position must be at a valid IFD header.

    """
    if offsetsize == 4:
        offsetformat = byteorder+'I'
        tagnosize = 2
        tagnoformat = byteorder+'H'
        tagsize = 12
        tagformat1 = byteorder+'HH'
        tagformat2 = byteorder+'I4s'
    elif offsetsize == 8:
        offsetformat = byteorder+'Q'
        tagnosize = 8
        tagnoformat = byteorder+'Q'
        tagsize = 20
        tagformat1 = byteorder+'HH'
        tagformat2 = byteorder+'Q8s'
    else:
        raise ValueError('invalid offset size')

    if customtags is None:
        customtags = {}
    if maxifds is None:
        maxifds = 2**32

    result = []
    unpack = struct.unpack
    offset = fh.tell()
    while len(result) < maxifds:
        # loop over IFDs
        try:
            tagno = unpack(tagnoformat, fh.read(tagnosize))[0]
            if tagno > 4096:
                raise TiffFileError('suspicious number of tags')
        except Exception:
            log.warning('read_tags: corrupted tag list at offset %i', offset)
            break

        tags = {}
        data = fh.read(tagsize * tagno)
        pos = fh.tell()
        index = 0
        for _ in range(tagno):
            code, type_ = unpack(tagformat1, data[index:index+4])
            count, value = unpack(tagformat2, data[index+4:index+tagsize])
            index += tagsize
            name = tagnames.get(code, str(code))
            try:
                dtype = TIFF.DATA_FORMATS[type_]
            except KeyError:
                raise TiffFileError('unknown tag data type %i' % type_)

            fmt = '%s%i%s' % (byteorder, count * int(dtype[0]), dtype[1])
            size = struct.calcsize(fmt)
            if size > offsetsize or code in customtags:
                offset = unpack(offsetformat, value)[0]
                if offset < 8 or offset > fh.size - size:
                    raise TiffFileError('invalid tag value offset %i' % offset)
                fh.seek(offset)
                if code in customtags:
                    readfunc = customtags[code][1]
                    value = readfunc(fh, byteorder, dtype, count, offsetsize)
                elif type_ == 7 or (count > 1 and dtype[-1] == 'B'):
                    value = read_bytes(fh, byteorder, dtype, count, offsetsize)
                elif code in tagnames or dtype[-1] == 's':
                    value = unpack(fmt, fh.read(size))
                else:
                    value = read_numpy(fh, byteorder, dtype, count, offsetsize)
            elif dtype[-1] == 'B' or type_ == 7:
                value = value[:size]
            else:
                value = unpack(fmt, value[:size])

            if code not in customtags and code not in TIFF.TAG_TUPLE:
                if len(value) == 1:
                    value = value[0]
            if type_ != 7 and dtype[-1] == 's' and isinstance(value, bytes):
                # TIFF ASCII fields can contain multiple strings,
                #   each terminated with a NUL
                try:
                    value = bytes2str(stripascii(value).strip())
                except UnicodeDecodeError:
                    log.warning(
                        'read_tags: coercing invalid ASCII to bytes (tag %i)',
                        code)

            tags[name] = value

        result.append(tags)
        # read offset to next page
        fh.seek(pos)
        offset = unpack(offsetformat, fh.read(offsetsize))[0]
        if offset == 0:
            break
        if offset >= fh.size:
            log.warning('read_tags: invalid page offset (%i)', offset)
            break
        fh.seek(offset)

    if result and maxifds == 1:
        result = result[0]
    return result


def read_exif_ifd(fh, byteorder, dtype, count, offsetsize):
    """Read EXIF tags from file and return as dict."""
    exif = read_tags(fh, byteorder, offsetsize, TIFF.EXIF_TAGS, maxifds=1)
    for name in ('ExifVersion', 'FlashpixVersion'):
        try:
            exif[name] = bytes2str(exif[name])
        except Exception:
            pass
    if 'UserComment' in exif:
        idcode = exif['UserComment'][:8]
        try:
            if idcode == b'ASCII\x00\x00\x00':
                exif['UserComment'] = bytes2str(exif['UserComment'][8:])
            elif idcode == b'UNICODE\x00':
                exif['UserComment'] = exif['UserComment'][8:].decode('utf-16')
        except Exception:
            pass
    return exif


def read_gps_ifd(fh, byteorder, dtype, count, offsetsize):
    """Read GPS tags from file and return as dict."""
    return read_tags(fh, byteorder, offsetsize, TIFF.GPS_TAGS, maxifds=1)


def read_interoperability_ifd(fh, byteorder, dtype, count, offsetsize):
    """Read Interoperability tags from file and return as dict."""
    tag_names = {1: 'InteroperabilityIndex'}
    return read_tags(fh, byteorder, offsetsize, tag_names, maxifds=1)


def read_bytes(fh, byteorder, dtype, count, offsetsize):
    """Read tag data from file and return as byte string."""
    dtype = 'B' if dtype[-1] == 's' else byteorder+dtype[-1]
    count *= numpy.dtype(dtype).itemsize
    data = fh.read(count)
    if len(data) != count:
        log.warning('read_bytes: failed to read all bytes (%i < %i)',
                    len(data), count)
    return data


def read_utf8(fh, byteorder, dtype, count, offsetsize):
    """Read tag data from file and return as unicode string."""
    return fh.read(count).decode('utf-8')


def read_numpy(fh, byteorder, dtype, count, offsetsize):
    """Read tag data from file and return as numpy array."""
    dtype = 'b' if dtype[-1] == 's' else byteorder+dtype[-1]
    return fh.read_array(dtype, count)


def read_colormap(fh, byteorder, dtype, count, offsetsize):
    """Read ColorMap data from file and return as numpy array."""
    cmap = fh.read_array(byteorder+dtype[-1], count)
    cmap.shape = (3, -1)
    return cmap


def read_json(fh, byteorder, dtype, count, offsetsize):
    """Read JSON tag data from file and return as object."""
    data = fh.read(count)
    try:
        return json.loads(unicode(stripnull(data), 'utf-8'))
    except ValueError:
        log.warning('read_json: invalid JSON')


def read_mm_header(fh, byteorder, dtype, count, offsetsize):
    """Read FluoView mm_header tag from file and return as dict."""
    mmh = fh.read_record(TIFF.MM_HEADER, byteorder=byteorder)
    mmh = recarray2dict(mmh)
    mmh['Dimensions'] = [
        (bytes2str(d[0]).strip(), d[1], d[2], d[3], bytes2str(d[4]).strip())
        for d in mmh['Dimensions']]
    d = mmh['GrayChannel']
    mmh['GrayChannel'] = (
        bytes2str(d[0]).strip(), d[1], d[2], d[3], bytes2str(d[4]).strip())
    return mmh


def read_mm_stamp(fh, byteorder, dtype, count, offsetsize):
    """Read FluoView mm_stamp tag from file and return as numpy.ndarray."""
    return fh.read_array(byteorder+'f8', 8)


def read_uic1tag(fh, byteorder, dtype, count, offsetsize, planecount=None):
    """Read MetaMorph STK UIC1Tag from file and return as dict.

    Return empty dictionary if planecount is unknown.

    """
    assert dtype in ('2I', '1I') and byteorder == '<'
    result = {}
    if dtype == '2I':
        # pre MetaMorph 2.5 (not tested)
        values = fh.read_array('<u4', 2*count).reshape(count, 2)
        result = {'ZDistance': values[:, 0] / values[:, 1]}
    elif planecount:
        for _ in range(count):
            tagid = struct.unpack('<I', fh.read(4))[0]
            if tagid in (28, 29, 37, 40, 41):
                # silently skip unexpected tags
                fh.read(4)
                continue
            name, value = read_uic_tag(fh, tagid, planecount, offset=True)
            result[name] = value
    return result


def read_uic2tag(fh, byteorder, dtype, planecount, offsetsize):
    """Read MetaMorph STK UIC2Tag from file and return as dict."""
    assert dtype == '2I' and byteorder == '<'
    values = fh.read_array('<u4', 6*planecount).reshape(planecount, 6)
    return {
        'ZDistance': values[:, 0] / values[:, 1],
        'DateCreated': values[:, 2],  # julian days
        'TimeCreated': values[:, 3],  # milliseconds
        'DateModified': values[:, 4],  # julian days
        'TimeModified': values[:, 5]}  # milliseconds


def read_uic3tag(fh, byteorder, dtype, planecount, offsetsize):
    """Read MetaMorph STK UIC3Tag from file and return as dict."""
    assert dtype == '2I' and byteorder == '<'
    values = fh.read_array('<u4', 2*planecount).reshape(planecount, 2)
    return {'Wavelengths': values[:, 0] / values[:, 1]}


def read_uic4tag(fh, byteorder, dtype, planecount, offsetsize):
    """Read MetaMorph STK UIC4Tag from file and return as dict."""
    assert dtype == '1I' and byteorder == '<'
    result = {}
    while True:
        tagid = struct.unpack('<H', fh.read(2))[0]
        if tagid == 0:
            break
        name, value = read_uic_tag(fh, tagid, planecount, offset=False)
        result[name] = value
    return result


def read_uic_tag(fh, tagid, planecount, offset):
    """Read a single UIC tag value from file and return tag name and value.

    UIC1Tags use an offset.

    """
    def read_int(count=1):
        value = struct.unpack('<%iI' % count, fh.read(4*count))
        return value[0] if count == 1 else value

    try:
        name, dtype = TIFF.UIC_TAGS[tagid]
    except IndexError:
        # unknown tag
        return '_TagId%i' % tagid, read_int()

    Fraction = TIFF.UIC_TAGS[4][1]

    if offset:
        pos = fh.tell()
        if dtype not in (int, None):
            off = read_int()
            if off < 8:
                if dtype is str:
                    return name, ''
                log.warning("read_uic_tag: invalid offset for tag '%s' (%i)",
                            name, off)
                return name, off
            fh.seek(off)

    if dtype is None:
        # skip
        name = '_' + name
        value = read_int()
    elif dtype is int:
        # int
        value = read_int()
    elif dtype is Fraction:
        # fraction
        value = read_int(2)
        value = value[0] / value[1]
    elif dtype is julian_datetime:
        # datetime
        value = julian_datetime(*read_int(2))
    elif dtype is read_uic_image_property:
        # ImagePropertyEx
        value = read_uic_image_property(fh)
    elif dtype is str:
        # pascal string
        size = read_int()
        if 0 <= size < 2**10:
            value = struct.unpack('%is' % size, fh.read(size))[0][:-1]
            value = bytes2str(stripnull(value))
        elif offset:
            value = ''
            log.warning("read_uic_tag: corrupt string in tag '%s'", name)
        else:
            raise ValueError('read_uic_tag: invalid string size %i' % size)
    elif dtype == '%ip':
        # sequence of pascal strings
        value = []
        for _ in range(planecount):
            size = read_int()
            if 0 <= size < 2**10:
                string = struct.unpack('%is' % size, fh.read(size))[0][:-1]
                string = bytes2str(stripnull(string))
                value.append(string)
            elif offset:
                log.warning("read_uic_tag: corrupt string in tag '%s'", name)
            else:
                raise ValueError('read_uic_tag: invalid string size: %i' %
                                 size)
    else:
        # struct or numpy type
        dtype = '<' + dtype
        if '%i' in dtype:
            dtype = dtype % planecount
        if '(' in dtype:
            # numpy type
            value = fh.read_array(dtype, 1)[0]
            if value.shape[-1] == 2:
                # assume fractions
                value = value[..., 0] / value[..., 1]
        else:
            # struct format
            value = struct.unpack(dtype, fh.read(struct.calcsize(dtype)))
            if len(value) == 1:
                value = value[0]

    if offset:
        fh.seek(pos + 4)

    return name, value


def read_uic_image_property(fh):
    """Read UIC ImagePropertyEx tag from file and return as dict."""
    # TODO: test this
    size = struct.unpack('B', fh.read(1))[0]
    name = struct.unpack('%is' % size, fh.read(size))[0][:-1]
    flags, prop = struct.unpack('<IB', fh.read(5))
    if prop == 1:
        value = struct.unpack('II', fh.read(8))
        value = value[0] / value[1]
    else:
        size = struct.unpack('B', fh.read(1))[0]
        value = struct.unpack('%is' % size, fh.read(size))[0]
    return dict(name=name, flags=flags, value=value)


def read_cz_lsminfo(fh, byteorder, dtype, count, offsetsize):
    """Read CZ_LSMINFO tag from file and return as dict."""
    assert byteorder == '<'
    magic_number, structure_size = struct.unpack('<II', fh.read(8))
    if magic_number not in (50350412, 67127628):
        raise ValueError('invalid CZ_LSMINFO structure')
    fh.seek(-8, 1)

    if structure_size < numpy.dtype(TIFF.CZ_LSMINFO).itemsize:
        # adjust structure according to structure_size
        lsminfo = []
        size = 0
        for name, dtype in TIFF.CZ_LSMINFO:
            size += numpy.dtype(dtype).itemsize
            if size > structure_size:
                break
            lsminfo.append((name, dtype))
    else:
        lsminfo = TIFF.CZ_LSMINFO

    lsminfo = fh.read_record(lsminfo, byteorder=byteorder)
    lsminfo = recarray2dict(lsminfo)

    # read LSM info subrecords at offsets
    for name, reader in TIFF.CZ_LSMINFO_READERS.items():
        if reader is None:
            continue
        offset = lsminfo.get('Offset' + name, 0)
        if offset < 8:
            continue
        fh.seek(offset)
        try:
            lsminfo[name] = reader(fh)
        except ValueError:
            pass
    return lsminfo


def read_lsm_floatpairs(fh):
    """Read LSM sequence of float pairs from file and return as list."""
    size = struct.unpack('<i', fh.read(4))[0]
    return fh.read_array('<2f8', count=size)


def read_lsm_positions(fh):
    """Read LSM positions from file and return as list."""
    size = struct.unpack('<I', fh.read(4))[0]
    return fh.read_array('<2f8', count=size)


def read_lsm_timestamps(fh):
    """Read LSM time stamps from file and return as list."""
    size, count = struct.unpack('<ii', fh.read(8))
    if size != (8 + 8 * count):
        log.warning('read_lsm_timestamps: invalid LSM TimeStamps block')
        return []
    # return struct.unpack('<%dd' % count, fh.read(8*count))
    return fh.read_array('<f8', count=count)


def read_lsm_eventlist(fh):
    """Read LSM events from file and return as list of (time, type, text)."""
    count = struct.unpack('<II', fh.read(8))[1]
    events = []
    while count > 0:
        esize, etime, etype = struct.unpack('<IdI', fh.read(16))
        etext = bytes2str(stripnull(fh.read(esize - 16)))
        events.append((etime, etype, etext))
        count -= 1
    return events


def read_lsm_channelcolors(fh):
    """Read LSM ChannelColors structure from file and return as dict."""
    result = {'Mono': False, 'Colors': [], 'ColorNames': []}
    pos = fh.tell()
    (size, ncolors, nnames,
     coffset, noffset, mono) = struct.unpack('<IIIIII', fh.read(24))
    if ncolors != nnames:
        log.warning(
            'read_lsm_channelcolors: invalid LSM ChannelColors structure')
        return result
    result['Mono'] = bool(mono)
    # Colors
    fh.seek(pos + coffset)
    colors = fh.read_array('uint8', count=ncolors*4).reshape((ncolors, 4))
    result['Colors'] = colors.tolist()
    # ColorNames
    fh.seek(pos + noffset)
    buffer = fh.read(size - noffset)
    names = []
    while len(buffer) > 4:
        size = struct.unpack('<I', buffer[:4])[0]
        names.append(bytes2str(buffer[4:3+size]))
        buffer = buffer[4+size:]
    result['ColorNames'] = names
    return result


def read_lsm_scaninfo(fh):
    """Read LSM ScanInfo structure from file and return as dict."""
    block = {}
    blocks = [block]
    unpack = struct.unpack
    if struct.unpack('<I', fh.read(4))[0] != 0x10000000:
        # not a Recording sub block
        log.warning('read_lsm_scaninfo: invalid LSM ScanInfo structure')
        return block
    fh.read(8)
    while True:
        entry, dtype, size = unpack('<III', fh.read(12))
        if dtype == 2:
            # ascii
            value = bytes2str(stripnull(fh.read(size)))
        elif dtype == 4:
            # long
            value = unpack('<i', fh.read(4))[0]
        elif dtype == 5:
            # rational
            value = unpack('<d', fh.read(8))[0]
        else:
            value = 0
        if entry in TIFF.CZ_LSMINFO_SCANINFO_ARRAYS:
            blocks.append(block)
            name = TIFF.CZ_LSMINFO_SCANINFO_ARRAYS[entry]
            newobj = []
            block[name] = newobj
            block = newobj
        elif entry in TIFF.CZ_LSMINFO_SCANINFO_STRUCTS:
            blocks.append(block)
            newobj = {}
            block.append(newobj)
            block = newobj
        elif entry in TIFF.CZ_LSMINFO_SCANINFO_ATTRIBUTES:
            name = TIFF.CZ_LSMINFO_SCANINFO_ATTRIBUTES[entry]
            block[name] = value
        elif entry == 0xffffffff:
            # end sub block
            block = blocks.pop()
        else:
            # unknown entry
            block['Entry0x%x' % entry] = value
        if not blocks:
            break
    return block


def read_sis(fh, byteorder, dtype, count, offsetsize):
    """Read OlympusSIS structure and return as dict.

    No specification is avaliable. Only few fields are known.

    """
    result = {}

    (magic, _, minute, hour, day, month, year, _, name, tagcount
     ) = struct.unpack('<4s6shhhhh6s32sh', fh.read(60))

    if magic != b'SIS0':
        raise ValueError('invalid OlympusSIS structure')

    result['name'] = bytes2str(stripnull(name))
    try:
        result['datetime'] = datetime.datetime(1900+year, month+1, day,
                                               hour, minute)
    except ValueError:
        pass

    data = fh.read(8 * tagcount)
    for i in range(0, tagcount*8, 8):
        tagtype, count, offset = struct.unpack('<hhI', data[i:i+8])
        fh.seek(offset)
        if tagtype == 1:
            # general data
            (_, lenexp, xcal, ycal, _, mag, _, camname, pictype,
             ) = struct.unpack('<10shdd8sd2s34s32s', fh.read(112))  # 220
            m = math.pow(10, lenexp)
            result['pixelsizex'] = xcal * m
            result['pixelsizey'] = ycal * m
            result['magnification'] = mag
            result['cameraname'] = bytes2str(stripnull(camname))
            result['picturetype'] = bytes2str(stripnull(pictype))
        elif tagtype == 10:
            # channel data
            continue
            # TODO: does not seem to work?
            # (length, _, exptime, emv, _, camname, _, mictype,
            #  ) = struct.unpack('<h22sId4s32s48s32s', fh.read(152))  # 720
            # result['exposuretime'] = exptime
            # result['emvoltage'] = emv
            # result['cameraname2'] = bytes2str(stripnull(camname))
            # result['microscopename'] = bytes2str(stripnull(mictype))

    return result


def read_sis_ini(fh, byteorder, dtype, count, offsetsize):
    """Read OlympusSIS INI string and return as dict."""
    inistr = fh.read(count)
    inistr = bytes2str(stripnull(inistr))
    try:
        return olympusini_metadata(inistr)
    except Exception as exc:
        log.warning('olympusini_metadata: %s: %s', exc.__class__.__name__, exc)
        return {}


def read_tvips_header(fh, byteorder, dtype, count, offsetsize):
    """Read TVIPS EM-MENU headers and return as dict."""
    result = {}
    header = fh.read_record(TIFF.TVIPS_HEADER_V1, byteorder=byteorder)
    for name, typestr in TIFF.TVIPS_HEADER_V1:
        result[name] = header[name].tolist()
    if header['Version'] == 2:
        header = fh.read_record(TIFF.TVIPS_HEADER_V2, byteorder=byteorder)
        if header['Magic'] != int(0xaaaaaaaa):
            log.warning('read_tvips_header: invalid TVIPS v2 magic number')
            return {}
        # decode utf16 strings
        for name, typestr in TIFF.TVIPS_HEADER_V2:
            if typestr.startswith('V'):
                s = header[name].tostring().decode('utf16', errors='ignore')
                result[name] = stripnull(s, null='\0')
            else:
                result[name] = header[name].tolist()
        # convert nm to m
        for axis in 'XY':
            header['PhysicalPixelSize' + axis] /= 1e9
            header['PixelSize' + axis] /= 1e9
    elif header.version != 1:
        log.warning('read_tvips_header: unknown TVIPS header version')
        return {}
    return result


def read_fei_metadata(fh, byteorder, dtype, count, offsetsize):
    """Read FEI SFEG/HELIOS headers and return as dict."""
    result = {}
    section = {}
    data = bytes2str(stripnull(fh.read(count)))
    for line in data.splitlines():
        line = line.strip()
        if line.startswith('['):
            section = {}
            result[line[1:-1]] = section
            continue
        try:
            key, value = line.split('=')
        except ValueError:
            continue
        section[key] = astype(value)
    return result


def read_cz_sem(fh, byteorder, dtype, count, offsetsize):
    """Read Zeiss SEM tag and return as dict.

    See https://sourceforge.net/p/gwyddion/mailman/message/29275000/ for
    unnamed values.

    """
    result = {'': ()}
    key = None
    data = bytes2str(stripnull(fh.read(count)))
    for line in data.splitlines():
        if line.isupper():
            key = line.lower()
        elif key:
            try:
                name, value = line.split('=')
            except ValueError:
                try:
                    name, value = line.split(':', 1)
                except Exception:
                    continue
            value = value.strip()
            unit = ''
            try:
                v, u = value.split()
                number = astype(v, (int, float))
                if number != v:
                    value = number
                    unit = u
            except Exception:
                number = astype(value, (int, float))
                if number != value:
                    value = number
                if value in ('No', 'Off'):
                    value = False
                elif value in ('Yes', 'On'):
                    value = True
            result[key] = (name.strip(), value)
            if unit:
                result[key] += (unit,)
            key = None
        else:
            result[''] += (astype(line, (int, float)),)
    return result


def read_nih_image_header(fh, byteorder, dtype, count, offsetsize):
    """Read NIH_IMAGE_HEADER tag from file and return as dict."""
    a = fh.read_record(TIFF.NIH_IMAGE_HEADER, byteorder=byteorder)
    a = a.newbyteorder(byteorder)
    a = recarray2dict(a)
    a['XUnit'] = a['XUnit'][:a['XUnitSize']]
    a['UM'] = a['UM'][:a['UMsize']]
    return a


def read_scanimage_metadata(fh):
    """Read ScanImage BigTIFF v3 static and ROI metadata from open file.

    Return non-varying frame data as dict and ROI group data as JSON.

    The settings can be used to read image data and metadata without parsing
    the TIFF file.

    Raise ValueError if file does not contain valid ScanImage v3 metadata.

    """
    fh.seek(0)
    try:
        byteorder, version = struct.unpack('<2sH', fh.read(4))
        if byteorder != b'II' or version != 43:
            raise Exception
        fh.seek(16)
        magic, version, size0, size1 = struct.unpack('<IIII', fh.read(16))
        if magic != 117637889 or version != 3:
            raise Exception
    except Exception:
        raise ValueError('not a ScanImage BigTIFF v3 file')

    frame_data = matlabstr2py(bytes2str(fh.read(size0)[:-1]))
    roi_data = read_json(fh, '<', None, size1, None) if size1 > 1 else {}
    return frame_data, roi_data


def read_micromanager_metadata(fh):
    """Read MicroManager non-TIFF settings from open file and return as dict.

    The settings can be used to read image data without parsing the TIFF file.

    Raise ValueError if the file does not contain valid MicroManager metadata.

    """
    fh.seek(0)
    try:
        byteorder = {b'II': '<', b'MM': '>'}[fh.read(2)]
    except IndexError:
        raise ValueError('not a MicroManager TIFF file')

    result = {}
    fh.seek(8)
    (index_header, index_offset, display_header, display_offset,
     comments_header, comments_offset, summary_header, summary_length
     ) = struct.unpack(byteorder + 'IIIIIIII', fh.read(32))

    if summary_header != 2355492:
        raise ValueError('invalid MicroManager summary header')
    result['Summary'] = read_json(fh, byteorder, None, summary_length, None)

    if index_header != 54773648:
        raise ValueError('invalid MicroManager index header')
    fh.seek(index_offset)
    header, count = struct.unpack(byteorder + 'II', fh.read(8))
    if header != 3453623:
        raise ValueError('invalid MicroManager index header')
    data = struct.unpack(byteorder + 'IIIII'*count, fh.read(20*count))
    result['IndexMap'] = {'Channel': data[::5],
                          'Slice': data[1::5],
                          'Frame': data[2::5],
                          'Position': data[3::5],
                          'Offset': data[4::5]}

    if display_header != 483765892:
        raise ValueError('invalid MicroManager display header')
    fh.seek(display_offset)
    header, count = struct.unpack(byteorder + 'II', fh.read(8))
    if header != 347834724:
        raise ValueError('invalid MicroManager display header')
    result['DisplaySettings'] = read_json(fh, byteorder, None, count, None)

    if comments_header != 99384722:
        raise ValueError('invalid MicroManager comments header')
    fh.seek(comments_offset)
    header, count = struct.unpack(byteorder + 'II', fh.read(8))
    if header != 84720485:
        raise ValueError('invalid MicroManager comments header')
    result['Comments'] = read_json(fh, byteorder, None, count, None)

    return result


def read_metaseries_catalog(fh):
    """Read MetaSeries non-TIFF hint catalog from file.

    Raise ValueError if the file does not contain a valid hint catalog.

    """
    # TODO: implement read_metaseries_catalog
    raise NotImplementedError()


def imagej_metadata_tag(metadata, byteorder):
    """Return IJMetadata and IJMetadataByteCounts tags from metadata dict.

    The tags can be passed to the TiffWriter.save function as extratags.

    The metadata dict may contain the following keys and values:

        Info : str
            Human-readable information as string.
        Labels : sequence of str
            Human-readable labels for each channel.
        Ranges : sequence of doubles
            Lower and upper values for each channel.
        LUTs : sequence of (3, 256) uint8 ndarrays
            Color palettes for each channel.
        Plot : bytes
            Undocumented ImageJ internal format.
        ROI: bytes
            Undocumented ImageJ internal region of interest format.
        Overlays : bytes
            Undocumented ImageJ internal format.

    """
    header = [{'>': b'IJIJ', '<': b'JIJI'}[byteorder]]
    bytecounts = [0]
    body = []

    def _string(data, byteorder):
        return data.encode('utf-16' + {'>': 'be', '<': 'le'}[byteorder])

    def _doubles(data, byteorder):
        return struct.pack(byteorder+('d' * len(data)), *data)

    def _ndarray(data, byteorder):
        return data.tobytes()

    def _bytes(data, byteorder):
        return data

    metadata_types = (
        ('Info', b'info', 1, _string),
        ('Labels', b'labl', None, _string),
        ('Ranges', b'rang', 1, _doubles),
        ('LUTs', b'luts', None, _ndarray),
        ('Plot', b'plot', 1, _bytes),
        ('ROI', b'roi ', 1, _bytes),
        ('Overlays', b'over', None, _bytes))

    for key, mtype, count, func in metadata_types:
        if key.lower() in metadata:
            key = key.lower()
        elif key not in metadata:
            continue
        if byteorder == '<':
            mtype = mtype[::-1]
        values = metadata[key]
        if count is None:
            count = len(values)
        else:
            values = [values]
        header.append(mtype + struct.pack(byteorder+'I', count))
        for value in values:
            data = func(value, byteorder)
            body.append(data)
            bytecounts.append(len(data))

    if not body:
        return ()
    body = b''.join(body)
    header = b''.join(header)
    data = header + body
    bytecounts[0] = len(header)
    bytecounts = struct.pack(byteorder+('I' * len(bytecounts)), *bytecounts)
    return ((50839, 'B', len(data), data, True),
            (50838, 'I', len(bytecounts)//4, bytecounts, True))


def imagej_metadata(data, bytecounts, byteorder):
    """Return IJMetadata tag value as dict.

    The 'Info' string can have multiple formats, e.g. OIF or ScanImage,
    that might be parsed into dicts using the matlabstr2py or
    oiffile.SettingsFile functions.

    """
    def _string(data, byteorder):
        return data.decode('utf-16' + {'>': 'be', '<': 'le'}[byteorder])

    def _doubles(data, byteorder):
        return struct.unpack(byteorder+('d' * (len(data) // 8)), data)

    def _lut(data, byteorder):
        return numpy.frombuffer(data, 'uint8').reshape(-1, 256)

    def _bytes(data, byteorder):
        return data

    metadata_types = {  # big-endian
        b'info': ('Info', _string),
        b'labl': ('Labels', _string),
        b'rang': ('Ranges', _doubles),
        b'luts': ('LUTs', _lut),
        b'plot': ('Plots', _bytes),
        b'roi ': ('ROI', _bytes),
        b'over': ('Overlays', _bytes)}
    metadata_types.update(  # little-endian
        dict((k[::-1], v) for k, v in metadata_types.items()))

    if not bytecounts:
        raise ValueError('no ImageJ metadata')

    if not data[:4] in (b'IJIJ', b'JIJI'):
        raise ValueError('invalid ImageJ metadata')

    header_size = bytecounts[0]
    if header_size < 12 or header_size > 804:
        raise ValueError('invalid ImageJ metadata header size')

    ntypes = (header_size - 4) // 8
    header = struct.unpack(byteorder+'4sI'*ntypes, data[4:4+ntypes*8])
    pos = 4 + ntypes * 8
    counter = 0
    result = {}
    for mtype, count in zip(header[::2], header[1::2]):
        values = []
        name, func = metadata_types.get(mtype, (bytes2str(mtype), read_bytes))
        for _ in range(count):
            counter += 1
            pos1 = pos + bytecounts[counter]
            values.append(func(data[pos:pos1], byteorder))
            pos = pos1
        result[name.strip()] = values[0] if count == 1 else values
    return result


def imagej_description_metadata(description):
    """Return metatata from ImageJ image description as dict.

    Raise ValueError if not a valid ImageJ description.

    >>> description = 'ImageJ=1.11a\\nimages=510\\nhyperstack=true\\n'
    >>> imagej_description_metadata(description)  # doctest: +SKIP
    {'ImageJ': '1.11a', 'images': 510, 'hyperstack': True}

    """
    def _bool(val):
        return {'true': True, 'false': False}[val.lower()]

    result = {}
    for line in description.splitlines():
        try:
            key, val = line.split('=')
        except Exception:
            continue
        key = key.strip()
        val = val.strip()
        for dtype in (int, float, _bool):
            try:
                val = dtype(val)
                break
            except Exception:
                pass
        result[key] = val

    if 'ImageJ' not in result:
        raise ValueError('not a ImageJ image description')
    return result


def imagej_description(shape, rgb=None, colormaped=False, version=None,
                       hyperstack=None, mode=None, loop=None, **kwargs):
    """Return ImageJ image description from data shape.

    ImageJ can handle up to 6 dimensions in order TZCYXS.

    >>> imagej_description((51, 5, 2, 196, 171))  # doctest: +SKIP
    ImageJ=1.11a
    images=510
    channels=2
    slices=5
    frames=51
    hyperstack=true
    mode=grayscale
    loop=false

    """
    if colormaped:
        raise NotImplementedError('ImageJ colormapping not supported')
    if version is None:
        version = '1.11a'
    shape = imagej_shape(shape, rgb=rgb)
    rgb = shape[-1] in (3, 4)

    result = ['ImageJ=%s' % version]
    append = []
    result.append('images=%i' % product(shape[:-3]))
    if hyperstack is None:
        hyperstack = True
        append.append('hyperstack=true')
    else:
        append.append('hyperstack=%s' % bool(hyperstack))
    if shape[2] > 1:
        result.append('channels=%i' % shape[2])
    if mode is None and not rgb:
        mode = 'grayscale'
    if hyperstack and mode:
        append.append('mode=%s' % mode)
    if shape[1] > 1:
        result.append('slices=%i' % shape[1])
    if shape[0] > 1:
        result.append('frames=%i' % shape[0])
        if loop is None:
            append.append('loop=false')
    if loop is not None:
        append.append('loop=%s' % bool(loop))
    for key, value in kwargs.items():
        append.append('%s=%s' % (key.lower(), value))

    return '\n'.join(result + append + [''])


def imagej_shape(shape, rgb=None):
    """Return shape normalized to 6D ImageJ hyperstack TZCYXS.

    Raise ValueError if not a valid ImageJ hyperstack shape.

    >>> imagej_shape((2, 3, 4, 5, 3), False)
    (2, 3, 4, 5, 3, 1)

    """
    shape = tuple(int(i) for i in shape)
    ndim = len(shape)
    if 1 > ndim > 6:
        raise ValueError('invalid ImageJ hyperstack: not 2 to 6 dimensional')
    if rgb is None:
        rgb = shape[-1] in (3, 4) and ndim > 2
    if rgb and shape[-1] not in (3, 4):
        raise ValueError('invalid ImageJ hyperstack: not a RGB image')
    if not rgb and ndim == 6 and shape[-1] != 1:
        raise ValueError('invalid ImageJ hyperstack: not a non-RGB image')
    if rgb or shape[-1] == 1:
        return (1, ) * (6 - ndim) + shape
    return (1, ) * (5 - ndim) + shape + (1,)


def json_description(shape, **metadata):
    """Return JSON image description from data shape and other metadata.

    Return UTF-8 encoded JSON.

    >>> json_description((256, 256, 3), axes='YXS')  # doctest: +SKIP
    b'{"shape": [256, 256, 3], "axes": "YXS"}'

    """
    metadata.update(shape=shape)
    return json.dumps(metadata)  # .encode('utf-8')


def json_description_metadata(description):
    """Return metatata from JSON formated image description as dict.

    Raise ValuError if description is of unknown format.

    >>> description = '{"shape": [256, 256, 3], "axes": "YXS"}'
    >>> json_description_metadata(description)  # doctest: +SKIP
    {'shape': [256, 256, 3], 'axes': 'YXS'}
    >>> json_description_metadata('shape=(256, 256, 3)')
    {'shape': (256, 256, 3)}

    """
    if description[:6] == 'shape=':
        # old-style 'shaped' description; not JSON
        shape = tuple(int(i) for i in description[7:-1].split(','))
        return dict(shape=shape)
    if description[:1] == '{' and description[-1:] == '}':
        # JSON description
        return json.loads(description)
    raise ValueError('invalid JSON image description', description)


def fluoview_description_metadata(description, ignoresections=None):
    """Return metatata from FluoView image description as dict.

    The FluoView image description format is unspecified. Expect failures.

    >>> descr = ('[Intensity Mapping]\\nMap Ch0: Range=00000 to 02047\\n'
    ...          '[Intensity Mapping End]')
    >>> fluoview_description_metadata(descr)
    {'Intensity Mapping': {'Map Ch0: Range': '00000 to 02047'}}

    """
    if not description.startswith('['):
        raise ValueError('invalid FluoView image description')
    if ignoresections is None:
        ignoresections = {'Region Info (Fields)', 'Protocol Description'}

    result = {}
    sections = [result]
    comment = False
    for line in description.splitlines():
        if not comment:
            line = line.strip()
        if not line:
            continue
        if line[0] == '[':
            if line[-5:] == ' End]':
                # close section
                del sections[-1]
                section = sections[-1]
                name = line[1:-5]
                if comment:
                    section[name] = '\n'.join(section[name])
                if name[:4] == 'LUT ':
                    a = numpy.array(section[name], dtype='uint8')
                    a.shape = -1, 3
                    section[name] = a
                continue
            # new section
            comment = False
            name = line[1:-1]
            if name[:4] == 'LUT ':
                section = []
            elif name in ignoresections:
                section = []
                comment = True
            else:
                section = {}
            sections.append(section)
            result[name] = section
            continue
        # add entry
        if comment:
            section.append(line)
            continue
        line = line.split('=', 1)
        if len(line) == 1:
            section[line[0].strip()] = None
            continue
        key, value = line
        if key[:4] == 'RGB ':
            section.extend(int(rgb) for rgb in value.split())
        else:
            section[key.strip()] = astype(value.strip())
    return result


def pilatus_description_metadata(description):
    """Return metatata from Pilatus image description as dict.

    Return metadata from Pilatus pixel array detectors by Dectris, created
    by camserver or TVX software.

    >>> pilatus_description_metadata('# Pixel_size 172e-6 m x 172e-6 m')
    {'Pixel_size': (0.000172, 0.000172)}

    """
    result = {}
    if not description.startswith('# '):
        return result
    for c in '#:=,()':
        description = description.replace(c, ' ')
    for line in description.split('\n'):
        if line[:2] != '  ':
            continue
        line = line.split()
        name = line[0]
        if line[0] not in TIFF.PILATUS_HEADER:
            try:
                result['DateTime'] = datetime.datetime.strptime(
                    ' '.join(line), '%Y-%m-%dT%H %M %S.%f')
            except Exception:
                result[name] = ' '.join(line[1:])
            continue
        indices, dtype = TIFF.PILATUS_HEADER[line[0]]
        if isinstance(indices[0], slice):
            # assumes one slice
            values = line[indices[0]]
        else:
            values = [line[i] for i in indices]
        if dtype is float and values[0] == 'not':
            values = ['NaN']
        values = tuple(dtype(v) for v in values)
        if dtype == str:
            values = ' '.join(values)
        elif len(values) == 1:
            values = values[0]
        result[name] = values
    return result


def svs_description_metadata(description):
    """Return metatata from Aperio image description as dict.

    The Aperio image description format is unspecified. Expect failures.

    >>> svs_description_metadata('Aperio Image Library v1.0')
    {'Aperio Image Library': 'v1.0'}

    """
    if not description.startswith('Aperio Image Library '):
        raise ValueError('invalid Aperio image description')
    result = {}
    lines = description.split('\n')
    key, value = lines[0].strip().rsplit(None, 1)  # 'Aperio Image Library'
    result[key.strip()] = value.strip()
    if len(lines) == 1:
        return result
    items = lines[1].split('|')
    result[''] = items[0].strip()  # TODO: parse this?
    for item in items[1:]:
        key, value = item.split(' = ')
        result[key.strip()] = astype(value.strip())
    return result


def stk_description_metadata(description):
    """Return metadata from MetaMorph image description as list of dict.

    The MetaMorph image description format is unspecified. Expect failures.

    """
    description = description.strip()
    if not description:
        return []
    try:
        description = bytes2str(description)
    except UnicodeDecodeError as exc:
        log.warning('stk_description_metadata: %s: %s',
                    exc.__class__.__name__, exc)
        return []
    result = []
    for plane in description.split('\x00'):
        d = {}
        for line in plane.split('\r\n'):
            line = line.split(':', 1)
            if len(line) > 1:
                name, value = line
                d[name.strip()] = astype(value.strip())
            else:
                value = line[0].strip()
                if value:
                    if '' in d:
                        d[''].append(value)
                    else:
                        d[''] = [value]
        result.append(d)
    return result


def metaseries_description_metadata(description):
    """Return metatata from MetaSeries image description as dict."""
    if not description.startswith('<MetaData>'):
        raise ValueError('invalid MetaSeries image description')

    from xml.etree import cElementTree as etree  # delayed import
    root = etree.fromstring(description)
    types = {'float': float, 'int': int,
             'bool': lambda x: asbool(x, 'on', 'off')}

    def parse(root, result):
        # recursive
        for child in root:
            attrib = child.attrib
            if not attrib:
                result[child.tag] = parse(child, {})
                continue
            if 'id' in attrib:
                i = attrib['id']
                t = attrib['type']
                v = attrib['value']
                if t in types:
                    result[i] = types[t](v)
                else:
                    result[i] = v
        return result

    adict = parse(root, {})
    if 'Description' in adict:
        adict['Description'] = adict['Description'].replace('&#13;&#10;', '\n')
    return adict


def scanimage_description_metadata(description):
    """Return metatata from ScanImage image description as dict."""
    return matlabstr2py(description)


def scanimage_artist_metadata(artist):
    """Return metatata from ScanImage artist tag as dict."""
    try:
        return json.loads(artist)
    except ValueError as exc:
        log.warning('scanimage_artist_metadata: %s: %s',
                    exc.__class__.__name__, exc)


def olympusini_metadata(inistr):
    """Return OlympusSIS metadata from INI string.

    No documentation is available.

    """
    def keyindex(key):
        # split key into name and index
        index = 0
        i = len(key.rstrip('0123456789'))
        if i < len(key):
            index = int(key[i:]) - 1
            key = key[:i]
        return key, index

    result = {}
    bands = []
    zpos = None
    tpos = None
    for line in inistr.splitlines():
        line = line.strip()
        if line == '' or line[0] == ';':
            continue
        if line[0] == '[' and line[-1] == ']':
            section_name = line[1:-1]
            result[section_name] = section = {}
            if section_name == 'Dimension':
                result['axes'] = axes = []
                result['shape'] = shape = []
            elif section_name == 'ASD':
                result[section_name] = []
            elif section_name == 'Z':
                if 'Dimension' in result:
                    result[section_name]['ZPos'] = zpos = []
            elif section_name == 'Time':
                if 'Dimension' in result:
                    result[section_name]['TimePos'] = tpos = []
            elif section_name == 'Band':
                nbands = result['Dimension']['Band']
                bands = [{'LUT': []} for i in range(nbands)]
                result[section_name] = bands
                iband = 0
        else:
            key, value = line.split('=')
            if value.strip() == '':
                value = None
            elif ',' in value:
                value = tuple(astype(v) for v in value.split(','))
            else:
                value = astype(value)

            if section_name == 'Dimension':
                section[key] = value
                axes.append(key)
                shape.append(value)
            elif section_name == 'ASD':
                if key == 'Count':
                    result['ASD'] = [{}] * value
                else:
                    key, index = keyindex(key)
                    result['ASD'][index][key] = value
            elif section_name == 'Band':
                if key[:3] == 'LUT':
                    lut = bands[iband]['LUT']
                    value = struct.pack('<I', value)
                    lut.append(
                        [ord(value[0:1]), ord(value[1:2]), ord(value[2:3])])
                else:
                    key, iband = keyindex(key)
                    bands[iband][key] = value
            elif key[:4] == 'ZPos' and zpos is not None:
                zpos.append(value)
            elif key[:7] == 'TimePos' and tpos is not None:
                tpos.append(value)
            else:
                section[key] = value

    if 'axes' in result:
        sisaxes = {'Band': 'C'}
        axes = []
        shape = []
        for i, x in zip(result['shape'], result['axes']):
            if i > 1:
                axes.append(sisaxes.get(x, x[0].upper()))
                shape.append(i)
        result['axes'] = ''.join(axes)
        result['shape'] = tuple(shape)
    try:
        result['Z']['ZPos'] = numpy.array(
            result['Z']['ZPos'][:result['Dimension']['Z']], 'float64')
    except Exception:
        pass
    try:
        result['Time']['TimePos'] = numpy.array(
            result['Time']['TimePos'][:result['Dimension']['Time']], 'int32')
    except Exception:
        pass
    for band in bands:
        band['LUT'] = numpy.array(band['LUT'], 'uint8')

    return result


def tile_decode(tile, tileindex, tileshape, tiledshape,
                lsb2msb, decompress, unpack, unpredict, out):
    """Decode tile segment bytes into 5D output array."""
    _, imagedepth, imagelength, imagewidth, _ = out.shape
    tileddepth, tiledlength, tiledwidth = tiledshape
    tiledepth, tilelength, tilewidth, samples = tileshape
    tilesize = tiledepth * tilelength * tilewidth * samples

    pl = tileindex // (tiledwidth * tiledlength * tileddepth)
    td = (tileindex // (tiledwidth * tiledlength)) % tileddepth * tiledepth
    tl = (tileindex // tiledwidth) % tiledlength * tilelength
    tw = tileindex % tiledwidth * tilewidth

    if tile:
        if lsb2msb:
            tile = bitorder_decode(tile, out=tile)
        tile = decompress(tile)
        tile = unpack(tile)
        # decompression / unpacking might return too many bytes
        tile = tile[:tilesize]
        try:
            # complete tile according to TIFF specification
            tile.shape = tileshape
        except ValueError:
            # tile fills remaining space; found in some JPEG compressed slides
            s = (min(imagedepth - td, tiledepth),
                 min(imagelength - tl, tilelength),
                 min(imagewidth - tw, tilewidth),
                 samples)
            try:
                tile.shape = s
            except ValueError:
                # incomplete tile; see gdal issue #1179
                log.warning('tile_decode: incomplete tile %s %s',
                            tile.shape, tileshape)
                t = numpy.zeros(tilesize, tile.dtype)
                s = min(tile.size, tilesize)
                t[:s] = tile[:s]
                tile = t.reshape(tileshape)

        tile = unpredict(tile, axis=-2, out=tile)
        out[pl, td:td+tiledepth, tl:tl+tilelength, tw:tw+tilewidth] = (
            tile[:imagedepth-td, :imagelength-tl, :imagewidth-tw])
    else:
        out[pl, td:td+tiledepth, tl:tl+tilelength, tw:tw+tilewidth] = 0


def unpack_rgb(data, dtype=None, bitspersample=None, rescale=True):
    """Return array from byte string containing packed samples.

    Use to unpack RGB565 or RGB555 to RGB888 format.

    Parameters
    ----------
    data : byte str
        The data to be decoded. Samples in each pixel are stored consecutively.
        Pixels are aligned to 8, 16, or 32 bit boundaries.
    dtype : numpy.dtype
        The sample data type. The byteorder applies also to the data stream.
    bitspersample : tuple
        Number of bits for each sample in a pixel.
    rescale : bool
        Upscale samples to the number of bits in dtype.

    Returns
    -------
    numpy.ndarray
        Flattened array of unpacked samples of native dtype.

    Examples
    --------
    >>> data = struct.pack('BBBB', 0x21, 0x08, 0xff, 0xff)
    >>> print(unpack_rgb(data, '<B', (5, 6, 5), False))
    [ 1  1  1 31 63 31]
    >>> print(unpack_rgb(data, '<B', (5, 6, 5)))
    [  8   4   8 255 255 255]
    >>> print(unpack_rgb(data, '<B', (5, 5, 5)))
    [ 16   8   8 255 255 255]

    """
    if bitspersample is None:
        bitspersample = (5, 6, 5)
    if dtype is None:
        dtype = '<B'
    dtype = numpy.dtype(dtype)
    bits = int(numpy.sum(bitspersample))
    if not (bits <= 32 and all(i <= dtype.itemsize*8 for i in bitspersample)):
        raise ValueError('sample size not supported: %s' % str(bitspersample))
    dt = next(i for i in 'BHI' if numpy.dtype(i).itemsize*8 >= bits)
    data = numpy.frombuffer(data, dtype.byteorder+dt)
    result = numpy.empty((data.size, len(bitspersample)), dtype.char)
    for i, bps in enumerate(bitspersample):
        t = data >> int(numpy.sum(bitspersample[i+1:]))
        t &= int('0b'+'1'*bps, 2)
        if rescale:
            o = ((dtype.itemsize * 8) // bps + 1) * bps
            if o > data.dtype.itemsize * 8:
                t = t.astype('I')
            t *= (2**o - 1) // (2**bps - 1)
            t //= 2**(o - (dtype.itemsize * 8))
        result[:, i] = t
    return result.reshape(-1)


def delta_encode(data, axis=-1, out=None):
    """Encode Delta."""
    if isinstance(data, (bytes, bytearray)):
        data = numpy.frombuffer(data, dtype='u1')
        diff = numpy.diff(data, axis=0)
        return numpy.insert(diff, 0, data[0]).tobytes()

    dtype = data.dtype
    if dtype.kind == 'f':
        data = data.view('u%i' % dtype.itemsize)

    diff = numpy.diff(data, axis=axis)
    key = [slice(None)] * data.ndim
    key[axis] = 0
    diff = numpy.insert(diff, 0, data[tuple(key)], axis=axis)

    if dtype.kind == 'f':
        return diff.view(dtype)
    return diff


def delta_decode(data, axis=-1, out=None):
    """Decode Delta."""
    if out is not None and not out.flags.writeable:
        out = None
    if isinstance(data, (bytes, bytearray)):
        data = numpy.frombuffer(data, dtype='u1')
        return numpy.cumsum(data, axis=0, dtype='u1', out=out).tobytes()
    if data.dtype.kind == 'f':
        view = data.view('u%i' % data.dtype.itemsize)
        view = numpy.cumsum(view, axis=axis, dtype=view.dtype)
        return view.view(data.dtype)
    return numpy.cumsum(data, axis=axis, dtype=data.dtype, out=out)


def bitorder_decode(data, out=None, _bitorder=[]):
    """Reverse bits in each byte of byte string or numpy array.

    Decode data where pixels with lower column values are stored in the
    lower-order bits of the bytes (TIFF FillOrder is LSB2MSB).

    Parameters
    ----------
    data : byte string or ndarray
        The data to be bit reversed. If byte string, a new bit-reversed byte
        string is returned. Numpy arrays are bit-reversed in-place.

    Examples
    --------
    >>> bitorder_decode(b'\\x01\\x64')
    b'\\x80&'
    >>> data = numpy.array([1, 666], dtype='uint16')
    >>> bitorder_decode(data)
    >>> data
    array([  128, 16473], dtype=uint16)

    """
    if not _bitorder:
        _bitorder.append(
            b'\x00\x80@\xc0 \xa0`\xe0\x10\x90P\xd00\xb0p\xf0\x08\x88H\xc8('
            b'\xa8h\xe8\x18\x98X\xd88\xb8x\xf8\x04\x84D\xc4$\xa4d\xe4\x14'
            b'\x94T\xd44\xb4t\xf4\x0c\x8cL\xcc,\xacl\xec\x1c\x9c\\\xdc<\xbc|'
            b'\xfc\x02\x82B\xc2"\xa2b\xe2\x12\x92R\xd22\xb2r\xf2\n\x8aJ\xca*'
            b'\xaaj\xea\x1a\x9aZ\xda:\xbaz\xfa\x06\x86F\xc6&\xa6f\xe6\x16'
            b'\x96V\xd66\xb6v\xf6\x0e\x8eN\xce.\xaen\xee\x1e\x9e^\xde>\xbe~'
            b'\xfe\x01\x81A\xc1!\xa1a\xe1\x11\x91Q\xd11\xb1q\xf1\t\x89I\xc9)'
            b'\xa9i\xe9\x19\x99Y\xd99\xb9y\xf9\x05\x85E\xc5%\xa5e\xe5\x15'
            b'\x95U\xd55\xb5u\xf5\r\x8dM\xcd-\xadm\xed\x1d\x9d]\xdd=\xbd}'
            b'\xfd\x03\x83C\xc3#\xa3c\xe3\x13\x93S\xd33\xb3s\xf3\x0b\x8bK'
            b'\xcb+\xabk\xeb\x1b\x9b[\xdb;\xbb{\xfb\x07\x87G\xc7\'\xa7g\xe7'
            b'\x17\x97W\xd77\xb7w\xf7\x0f\x8fO\xcf/\xafo\xef\x1f\x9f_'
            b'\xdf?\xbf\x7f\xff')
        _bitorder.append(numpy.frombuffer(_bitorder[0], dtype='uint8'))
    try:
        view = data.view('uint8')
        numpy.take(_bitorder[1], view, out=view)
        return data
    except AttributeError:
        return data.translate(_bitorder[0])
    except ValueError:
        raise NotImplementedError('slices of arrays not supported')
    return None


def packints_decode(data, dtype, numbits, runlen=0, out=None):
    """Decompress byte string to array of integers.

    This implementation only handles itemsizes 1, 8, 16, 32, and 64 bits.
    Install the imagecodecs package for decoding other integer sizes.

    Parameters
    ----------
    data : byte str
        Data to decompress.
    dtype : numpy.dtype or str
        A numpy boolean or integer type.
    numbits : int
        Number of bits per integer.
    runlen : int
        Number of consecutive integers, after which to start at next byte.

    Examples
    --------
    >>> packints_decode(b'a', 'B', 1)
    array([0, 1, 1, 0, 0, 0, 0, 1], dtype=uint8)

    """
    if numbits == 1:  # bitarray
        data = numpy.frombuffer(data, '|B')
        data = numpy.unpackbits(data)
        if runlen % 8:
            data = data.reshape(-1, runlen + (8 - runlen % 8))
            data = data[:, :runlen].reshape(-1)
        return data.astype(dtype)
    if numbits in (8, 16, 32, 64):
        return numpy.frombuffer(data, dtype)
    raise NotImplementedError('unpacking %s-bit integers to %s not supported'
                              % (numbits, numpy.dtype(dtype)))


if imagecodecs is not None:
    bitorder_decode = imagecodecs.bitorder_decode  # noqa
    packints_decode = imagecodecs.packints_decode  # noqa


def decode_lzw(encoded):
    """Decompress LZW encoded byte string."""
    warnings.warn(
        'The decode_lzw function was removed from the tifffile package.\n'
        'Use the lzw_decode function from the imagecodecs package instead.')
    return imagecodecs.lzw_decode(encoded)


def apply_colormap(image, colormap, contig=True):
    """Return palette-colored image.

    The image values are used to index the colormap on axis 1. The returned
    image is of shape image.shape+colormap.shape[0] and dtype colormap.dtype.

    Parameters
    ----------
    image : numpy.ndarray
        Indexes into the colormap.
    colormap : numpy.ndarray
        RGB lookup table aka palette of shape (3, 2**bits_per_sample).
    contig : bool
        If True, return a contiguous array.

    Examples
    --------
    >>> image = numpy.arange(256, dtype='uint8')
    >>> colormap = numpy.vstack([image, image, image]).astype('uint16') * 256
    >>> apply_colormap(image, colormap)[-1]
    array([65280, 65280, 65280], dtype=uint16)

    """
    image = numpy.take(colormap, image, axis=1)
    image = numpy.rollaxis(image, 0, image.ndim)
    if contig:
        image = numpy.ascontiguousarray(image)
    return image


def reorient(image, orientation):
    """Return reoriented view of image array.

    Parameters
    ----------
    image : numpy.ndarray
        Non-squeezed output of asarray() functions.
        Axes -3 and -2 must be image length and width respectively.
    orientation : int or str
        One of TIFF.ORIENTATION names or values.

    """
    orient = TIFF.ORIENTATION
    orientation = enumarg(orient, orientation)

    if orientation == orient.TOPLEFT:
        return image
    if orientation == orient.TOPRIGHT:
        return image[..., ::-1, :]
    if orientation == orient.BOTLEFT:
        return image[..., ::-1, :, :]
    if orientation == orient.BOTRIGHT:
        return image[..., ::-1, ::-1, :]
    if orientation == orient.LEFTTOP:
        return numpy.swapaxes(image, -3, -2)
    if orientation == orient.RIGHTTOP:
        return numpy.swapaxes(image, -3, -2)[..., ::-1, :]
    if orientation == orient.RIGHTBOT:
        return numpy.swapaxes(image, -3, -2)[..., ::-1, :, :]
    if orientation == orient.LEFTBOT:
        return numpy.swapaxes(image, -3, -2)[..., ::-1, ::-1, :]
    return image


def repeat_nd(a, repeats):
    """Return read-only view into input array with elements repeated.

    Zoom nD image by integer factors using nearest neighbor interpolation
    (box filter).

    Parameters
    ----------
    a : array_like
        Input array.
    repeats : sequence of int
        The number of repetitions to apply along each dimension of input array.

    Examples
    --------
    >>> repeat_nd([[1, 2], [3, 4]], (2, 2))
    array([[1, 1, 2, 2],
           [1, 1, 2, 2],
           [3, 3, 4, 4],
           [3, 3, 4, 4]])

    """
    a = numpy.asarray(a)
    reshape = []
    shape = []
    strides = []
    for i, j, k in zip(a.strides, a.shape, repeats):
        shape.extend((j, k))
        strides.extend((i, 0))
        reshape.append(j * k)
    return numpy.lib.stride_tricks.as_strided(
        a, shape, strides, writeable=False).reshape(reshape)


def reshape_nd(data_or_shape, ndim):
    """Return image array or shape with at least ndim dimensions.

    Prepend 1s to image shape as necessary.

    >>> reshape_nd(numpy.empty(0), 1).shape
    (0,)
    >>> reshape_nd(numpy.empty(1), 2).shape
    (1, 1)
    >>> reshape_nd(numpy.empty((2, 3)), 3).shape
    (1, 2, 3)
    >>> reshape_nd(numpy.empty((3, 4, 5)), 3).shape
    (3, 4, 5)
    >>> reshape_nd((2, 3), 3)
    (1, 2, 3)

    """
    is_shape = isinstance(data_or_shape, tuple)
    shape = data_or_shape if is_shape else data_or_shape.shape
    if len(shape) >= ndim:
        return data_or_shape
    shape = (1,) * (ndim - len(shape)) + shape
    return shape if is_shape else data_or_shape.reshape(shape)


def squeeze_axes(shape, axes, skip=None):
    """Return shape and axes with single-dimensional entries removed.

    Remove unused dimensions unless their axes are listed in 'skip'.

    >>> squeeze_axes((5, 1, 2, 1, 1), 'TZYXC')
    ((5, 2, 1), 'TYX')

    """
    if len(shape) != len(axes):
        raise ValueError('dimensions of axes and shape do not match')
    if skip is None:
        skip = 'XY'
    shape, axes = zip(*(i for i in zip(shape, axes)
                        if i[0] > 1 or i[1] in skip))
    return tuple(shape), ''.join(axes)


def transpose_axes(image, axes, asaxes=None):
    """Return image with its axes permuted to match specified axes.

    A view is returned if possible.

    >>> transpose_axes(numpy.zeros((2, 3, 4, 5)), 'TYXC', asaxes='CTZYX').shape
    (5, 2, 1, 3, 4)

    """
    for ax in axes:
        if ax not in asaxes:
            raise ValueError('unknown axis %s' % ax)
    # add missing axes to image
    if asaxes is None:
        asaxes = 'CTZYX'
    shape = image.shape
    for ax in reversed(asaxes):
        if ax not in axes:
            axes = ax + axes
            shape = (1,) + shape
    image = image.reshape(shape)
    # transpose axes
    image = image.transpose([axes.index(ax) for ax in asaxes])
    return image


def reshape_axes(axes, shape, newshape, unknown=None):
    """Return axes matching new shape.

    By default, unknown dimensions are labelled 'Q'.

    >>> reshape_axes('YXS', (219, 301, 1), (219, 301))
    'YX'
    >>> reshape_axes('IYX', (12, 219, 301), (3, 4, 219, 1, 301, 1))
    'QQYQXQ'

    """
    shape = tuple(shape)
    newshape = tuple(newshape)
    if len(axes) != len(shape):
        raise ValueError('axes do not match shape')

    size = product(shape)
    newsize = product(newshape)
    if size != newsize:
        raise ValueError('cannot reshape %s to %s' % (shape, newshape))
    if not axes or not newshape:
        return ''

    lendiff = max(0, len(shape) - len(newshape))
    if lendiff:
        newshape = newshape + (1,) * lendiff

    i = len(shape)-1
    prodns = 1
    prods = 1
    result = []
    for ns in newshape[::-1]:
        prodns *= ns
        while i > 0 and shape[i] == 1 and ns != 1:
            i -= 1
        if ns == shape[i] and prodns == prods*shape[i]:
            prods *= shape[i]
            result.append(axes[i])
            i -= 1
        elif unknown:
            result.append(unknown)
        else:
            unknown = 'Q'
            result.append(unknown)

    return ''.join(reversed(result[lendiff:]))


def stack_pages(pages, out=None, maxworkers=None, **kwargs):
    """Read data from sequence of TiffPage and stack them vertically.

    Additional parameters are passsed to the TiffPage.asarray function.

    """
    npages = len(pages)
    if npages == 0:
        raise ValueError('no pages')

    if npages == 1:
        kwargs['maxworkers'] = maxworkers
        return pages[0].asarray(out=out, **kwargs)

    page0 = next(p for p in pages if p is not None).keyframe
    page0.asarray(validate=None)  # ThreadPoolExecutor swallows exceptions
    shape = (npages,) + page0.shape
    dtype = page0.dtype
    out = create_output(out, shape, dtype)

    if maxworkers is None:
        if page0.compression > 1:
            if page0.is_tiled:
                maxworkers = 1
                kwargs['maxworkers'] = 0
            else:
                maxworkers = 0
        else:
            maxworkers = 1
    if maxworkers == 0:
        import multiprocessing  # noqa: delay import
        maxworkers = multiprocessing.cpu_count() // 2
    if maxworkers > 1:
        kwargs['maxworkers'] = 1

    page0.parent.filehandle.lock = maxworkers > 1

    filecache = OpenFileCache(size=max(4, maxworkers),
                              lock=page0.parent.filehandle.lock)

    def func(page, index, out=out, filecache=filecache, kwargs=kwargs):
        """Read, decode, and copy page data."""
        if page is not None:
            filecache.open(page.parent.filehandle)
            out[index] = page.asarray(lock=filecache.lock, reopen=False,
                                      validate=False, **kwargs)
            filecache.close(page.parent.filehandle)

    if maxworkers < 2:
        for i, page in enumerate(pages):
            func(page, i)
    else:
        # TODO: add exception handling
        with ThreadPoolExecutor(maxworkers) as executor:
            executor.map(func, pages, range(npages))

    filecache.clear()
    page0.parent.filehandle.lock = None

    return out


def clean_offsetscounts(offsets, counts):
    """Return cleaned offsets and byte counts.

    Remove zero offsets and counts.
    Use to sanitize StripOffsets and StripByteCounts tag values.

    """
    # TODO: cythonize this
    offsets = list(offsets)
    counts = list(counts)
    size = len(offsets)
    if size != len(counts):
        raise ValueError('StripOffsets and StripByteCounts mismatch')
    j = 0
    for i, (o, b) in enumerate(zip(offsets, counts)):
        if b > 0:
            if o > 0:
                if i > j:
                    offsets[j] = o
                    counts[j] = b
                j += 1
                continue
            raise ValueError('invalid offset')
        log.warning('clean_offsetscounts: empty bytecount')
    if size == len(offsets):
        return offsets, counts
    if j == 0:
        return [offsets[0]], [counts[0]]
    return offsets[:j], counts[:j]


def buffered_read(fh, lock, offsets, bytecounts, buffersize=None):
    """Return iterator over segments read from file."""
    if buffersize is None:
        buffersize = 2**26
    length = len(offsets)
    i = 0
    while i < length:
        data = []
        with lock:
            size = 0
            while size < buffersize and i < length:
                fh.seek(offsets[i])
                bytecount = bytecounts[i]
                data.append(fh.read(bytecount))
                # buffer = bytearray(bytecount)
                # n = fh.readinto(buffer)
                # data.append(buffer[:n])
                size += bytecount
                i += 1
        for segment in data:
            yield segment


def create_output(out, shape, dtype, mode='w+', suffix=None):
    """Return numpy array where image data of shape and dtype can be copied.

    The 'out' parameter may have the following values or types:

    None
        An empty array of shape and dtype is created and returned.
    numpy.ndarray
        An existing writable array of compatible dtype and shape. A view of
        the same array is returned after verification.
    'memmap' or 'memmap:tempdir'
        A memory-map to an array stored in a temporary binary file on disk
        is created and returned.
    str or open file
        The file name or file object used to create a memory-map to an array
        stored in a binary file on disk. The created memory-mapped array is
        returned.

    """
    if out is None:
        return numpy.zeros(shape, dtype)
    if isinstance(out, str) and out[:6] == 'memmap':
        import tempfile  # noqa: delay import
        tempdir = out[7:] if len(out) > 7 else None
        if suffix is None:
            suffix = '.memmap'
        with tempfile.NamedTemporaryFile(dir=tempdir, suffix=suffix) as fh:
            return numpy.memmap(fh, shape=shape, dtype=dtype, mode=mode)
    if isinstance(out, numpy.ndarray):
        if product(shape) != product(out.shape):
            raise ValueError('incompatible output shape')
        if not numpy.can_cast(dtype, out.dtype):
            raise ValueError('incompatible output dtype')
        return out.reshape(shape)
    if isinstance(out, pathlib.Path):
        out = str(out)
    return numpy.memmap(out, shape=shape, dtype=dtype, mode=mode)


def matlabstr2py(string):
    """Return Python object from Matlab string representation.

    Return str, bool, int, float, list (Matlab arrays or cells), or
    dict (Matlab structures) types.

    Use to access ScanImage metadata.

    >>> matlabstr2py('1')
    1
    >>> matlabstr2py("['x y z' true false; 1 2.0 -3e4; NaN Inf @class]")
    [['x y z', True, False], [1, 2.0, -30000.0], [nan, inf, '@class']]
    >>> d = matlabstr2py("SI.hChannels.channelType = {'stripe' 'stripe'}\\n"
    ...                  "SI.hChannels.channelsActive = 2")
    >>> d['SI.hChannels.channelType']
    ['stripe', 'stripe']

    """
    # TODO: handle invalid input
    # TODO: review unboxing of multidimensional arrays

    def lex(s):
        # return sequence of tokens from matlab string representation
        tokens = ['[']
        while True:
            t, i = next_token(s)
            if t is None:
                break
            if t == ';':
                tokens.extend((']', '['))
            elif t == '[':
                tokens.extend(('[', '['))
            elif t == ']':
                tokens.extend((']', ']'))
            else:
                tokens.append(t)
            s = s[i:]
        tokens.append(']')
        return tokens

    def next_token(s):
        # return next token in matlab string
        length = len(s)
        if length == 0:
            return None, 0
        i = 0
        while i < length and s[i] == ' ':
            i += 1
        if i == length:
            return None, i
        if s[i] in '{[;]}':
            return s[i], i + 1
        if s[i] == "'":
            j = i + 1
            while j < length and s[j] != "'":
                j += 1
            return s[i: j+1], j + 1
        if s[i] == '<':
            j = i + 1
            while j < length and s[j] != '>':
                j += 1
            return s[i: j+1], j + 1
        j = i
        while j < length and not s[j] in ' {[;]}':
            j += 1
        return s[i:j], j

    def value(s, fail=False):
        # return Python value of token
        s = s.strip()
        if not s:
            return s
        if len(s) == 1:
            try:
                return int(s)
            except Exception:
                if fail:
                    raise ValueError()
                return s
        if s[0] == "'":
            if fail and s[-1] != "'" or "'" in s[1:-1]:
                raise ValueError()
            return s[1:-1]
        if s[0] == '<':
            if fail and s[-1] != '>' or '<' in s[1:-1]:
                raise ValueError()
            return s
        if fail and any(i in s for i in " ';[]{}"):
            raise ValueError()
        if s[0] == '@':
            return s
        if s in ('true', 'True'):
            return True
        if s in ('false', 'False'):
            return False
        if s[:6] == 'zeros(':
            return numpy.zeros([int(i) for i in s[6:-1].split(',')]).tolist()
        if s[:5] == 'ones(':
            return numpy.ones([int(i) for i in s[5:-1].split(',')]).tolist()
        if '.' in s or 'e' in s:
            try:
                return float(s)
            except Exception:
                pass
        try:
            return int(s)
        except Exception:
            pass
        try:
            return float(s)  # nan, inf
        except Exception:
            if fail:
                raise ValueError()
        return s

    def parse(s):
        # return Python value from string representation of Matlab value
        s = s.strip()
        try:
            return value(s, fail=True)
        except ValueError:
            pass
        result = add2 = []
        levels = [add2]
        for t in lex(s):
            if t in '[{':
                add2 = []
                levels.append(add2)
            elif t in ']}':
                x = levels.pop()
                if len(x) == 1 and isinstance(x[0], (list, str)):
                    x = x[0]
                add2 = levels[-1]
                add2.append(x)
            else:
                add2.append(value(t))
        if len(result) == 1 and isinstance(result[0], (list, str)):
            result = result[0]
        return result

    if '\r' in string or '\n' in string:
        # structure
        d = {}
        for line in string.splitlines():
            line = line.strip()
            if not line or line[0] == '%':
                continue
            k, v = line.split('=', 1)
            k = k.strip()
            if any(c in k for c in " ';[]{}<>"):
                continue
            d[k] = parse(v)
        return d
    return parse(string)


def stripnull(string, null=b'\x00'):
    """Return string truncated at first null character.

    Clean NULL terminated C strings. For unicode strings use null='\\0'.

    >>> stripnull(b'string\\x00')
    b'string'
    >>> stripnull('string\\x00', null='\\0')
    'string'

    """
    i = string.find(null)
    return string if (i < 0) else string[:i]


def stripascii(string):
    """Return string truncated at last byte that is 7-bit ASCII.

    Clean NULL separated and terminated TIFF strings.

    >>> stripascii(b'string\\x00string\\n\\x01\\x00')
    b'string\\x00string\\n'
    >>> stripascii(b'\\x00')
    b''

    """
    # TODO: pythonize this
    i = len(string)
    while i:
        i -= 1
        if 8 < byte2int(string[i]) < 127:
            break
    else:
        i = -1
    return string[:i+1]


def asbool(value, true=(b'true', u'true'), false=(b'false', u'false')):
    """Return string as bool if possible, else raise TypeError.

    >>> asbool(b' False ')
    False

    """
    value = value.strip().lower()
    if value in true:  # might raise UnicodeWarning/BytesWarning
        return True
    if value in false:
        return False
    raise TypeError()


def astype(value, types=None):
    """Return argument as one of types if possible.

    >>> astype('42')
    42
    >>> astype('3.14')
    3.14
    >>> astype('True')
    True
    >>> astype(b'Neee-Wom')
    'Neee-Wom'

    """
    if types is None:
        types = int, float, asbool, bytes2str
    for typ in types:
        try:
            return typ(value)
        except (ValueError, AttributeError, TypeError, UnicodeEncodeError):
            pass
    return value


def format_size(size, threshold=1536):
    """Return file size as string from byte size.

    >>> format_size(1234)
    '1234 B'
    >>> format_size(12345678901)
    '11.50 GiB'

    """
    if size < threshold:
        return "%i B" % size
    for unit in ('KiB', 'MiB', 'GiB', 'TiB', 'PiB'):
        size /= 1024.0
        if size < threshold:
            return "%.2f %s" % (size, unit)
    return 'ginormous'


def identityfunc(arg, *args, **kwargs):
    """Single argument identity function.

    >>> identityfunc('arg')
    'arg'

    """
    return arg


def nullfunc(*args, **kwargs):
    """Null function.

    >>> nullfunc('arg', kwarg='kwarg')

    """
    return


def sequence(value):
    """Return tuple containing value if value is not a tuple or list.

    >>> sequence(1)
    (1,)
    >>> sequence([1])
    [1]
    >>> sequence('ab')
    ('ab',)

    """
    return value if isinstance(value, (tuple, list)) else (value,)


def product(iterable):
    """Return product of sequence of numbers.

    Equivalent of functools.reduce(operator.mul, iterable, 1).
    Multiplying numpy integers might overflow.

    >>> product([2**8, 2**30])
    274877906944
    >>> product([])
    1

    """
    prod = 1
    for i in iterable:
        prod *= i
    return prod


def natural_sorted(iterable):
    """Return human sorted list of strings.

    E.g. for sorting file names.

    >>> natural_sorted(['f1', 'f2', 'f10'])
    ['f1', 'f2', 'f10']

    """
    def sortkey(x):
        return [(int(c) if c.isdigit() else c) for c in re.split(numbers, x)]

    numbers = re.compile(r'(\d+)')
    return sorted(iterable, key=sortkey)


def excel_datetime(timestamp, epoch=None):
    """Return datetime object from timestamp in Excel serial format.

    Convert LSM time stamps.

    >>> excel_datetime(40237.029999999795)
    datetime.datetime(2010, 2, 28, 0, 43, 11, 999982)

    """
    if epoch is None:
        epoch = datetime.datetime.fromordinal(693594)
    return epoch + datetime.timedelta(timestamp)


def julian_datetime(julianday, milisecond=0):
    """Return datetime from days since 1/1/4713 BC and ms since midnight.

    Convert Julian dates according to MetaMorph.

    >>> julian_datetime(2451576, 54362783)
    datetime.datetime(2000, 2, 2, 15, 6, 2, 783)

    """
    if julianday <= 1721423:
        # no datetime before year 1
        return None

    a = julianday + 1
    if a > 2299160:
        alpha = math.trunc((a - 1867216.25) / 36524.25)
        a += 1 + alpha - alpha // 4
    b = a + (1524 if a > 1721423 else 1158)
    c = math.trunc((b - 122.1) / 365.25)
    d = math.trunc(365.25 * c)
    e = math.trunc((b - d) / 30.6001)

    day = b - d - math.trunc(30.6001 * e)
    month = e - (1 if e < 13.5 else 13)
    year = c - (4716 if month > 2.5 else 4715)

    hour, milisecond = divmod(milisecond, 1000 * 60 * 60)
    minute, milisecond = divmod(milisecond, 1000 * 60)
    second, milisecond = divmod(milisecond, 1000)

    return datetime.datetime(year, month, day,
                             hour, minute, second, milisecond)


def byteorder_isnative(byteorder):
    """Return if byteorder matches the system's byteorder.

    >>> byteorder_isnative('=')
    True

    """
    if byteorder in ('=', sys.byteorder):
        return True
    keys = {'big': '>', 'little': '<'}
    return keys.get(byteorder, byteorder) == keys[sys.byteorder]


def recarray2dict(recarray):
    """Return numpy.recarray as dict."""
    # TODO: subarrays
    result = {}
    for descr, value in zip(recarray.dtype.descr, recarray):
        name, dtype = descr[:2]
        if dtype[1] == 'S':
            value = bytes2str(stripnull(value))
        elif value.ndim < 2:
            value = value.tolist()
        result[name] = value
    return result


def xml2dict(xml, sanitize=True, prefix=None):
    """Return XML as dict.

    >>> xml2dict('<?xml version="1.0" ?><root attr="name"><key>1</key></root>')
    {'root': {'key': 1, 'attr': 'name'}}

    """
    from xml.etree import cElementTree as etree  # delayed import

    at = tx = ''
    if prefix:
        at, tx = prefix

    def astype(value):
        # return value as int, float, bool, or str
        for t in (int, float, asbool):
            try:
                return t(value)
            except Exception:
                pass
        return value

    def etree2dict(t):
        # adapted from https://stackoverflow.com/a/10077069/453463
        key = t.tag
        if sanitize:
            key = key.rsplit('}', 1)[-1]
        d = {key: {} if t.attrib else None}
        children = list(t)
        if children:
            dd = collections.defaultdict(list)
            for dc in map(etree2dict, children):
                for k, v in dc.items():
                    dd[k].append(astype(v))
            d = {key: {k: astype(v[0]) if len(v) == 1 else astype(v)
                       for k, v in dd.items()}}
        if t.attrib:
            d[key].update((at + k, astype(v)) for k, v in t.attrib.items())
        if t.text:
            text = t.text.strip()
            if children or t.attrib:
                if text:
                    d[key][tx + 'value'] = astype(text)
            else:
                d[key] = astype(text)
        return d

    return etree2dict(etree.fromstring(xml))


def hexdump(bytestr, width=75, height=24, snipat=-2, modulo=2, ellipsis=None):
    """Return hexdump representation of byte string.

    >>> hexdump(binascii.unhexlify('49492a00080000000e00fe0004000100'))
    '49 49 2a 00 08 00 00 00 0e 00 fe 00 04 00 01 00 II*.............'

    """
    size = len(bytestr)
    if size < 1 or width < 2 or height < 1:
        return ''
    if height == 1:
        addr = b''
        bytesperline = min(modulo * (((width - len(addr)) // 4) // modulo),
                           size)
        if bytesperline < 1:
            return ''
        nlines = 1
    else:
        addr = b'%%0%ix: ' % len(b'%x' % size)
        bytesperline = min(modulo * (((width - len(addr % 1)) // 4) // modulo),
                           size)
        if bytesperline < 1:
            return ''
        width = 3*bytesperline + len(addr % 1)
        nlines = (size - 1) // bytesperline + 1

    if snipat is None or snipat == 1:
        snipat = height
    elif 0 < abs(snipat) < 1:
        snipat = int(math.floor(height * snipat))
    if snipat < 0:
        snipat += height

    if height == 1 or nlines == 1:
        blocks = [(0, bytestr[:bytesperline])]
        addr = b''
        height = 1
        width = 3 * bytesperline
    elif height is None or nlines <= height:
        blocks = [(0, bytestr)]
    elif snipat <= 0:
        start = bytesperline * (nlines - height)
        blocks = [(start, bytestr[start:])]  # (start, None)
    elif snipat >= height or height < 3:
        end = bytesperline * height
        blocks = [(0, bytestr[:end])]  # (end, None)
    else:
        end1 = bytesperline * snipat
        end2 = bytesperline * (height - snipat - 1)
        blocks = [(0, bytestr[:end1]),
                  (size-end1-end2, None),
                  (size-end2, bytestr[size-end2:])]

    ellipsis = b'...' if ellipsis is None else str2bytes(ellipsis)
    result = []
    for start, bytestr in blocks:
        if bytestr is None:
            result.append(ellipsis)  # 'skip %i bytes' % start)
            continue
        hexstr = binascii.hexlify(bytestr)
        strstr = re.sub(br'[^\x20-\x7f]', b'../fd_deeploc_core', bytestr)
        for i in range(0, len(bytestr), bytesperline):
            h = hexstr[2*i:2*i+bytesperline*2]
            r = (addr % (i + start)) if height > 1 else addr
            r += b' '.join(h[i:i+2] for i in range(0, 2*bytesperline, 2))
            r += b' ' * (width - len(r))
            r += strstr[i:i+bytesperline]
            result.append(r)
    result = b'\n'.join(result)
    if sys.version_info[0] == 3:
        result = result.decode('ascii')
    return result


def isprintable(string):
    """Return if all characters in string are printable.

    >>> isprintable('abc')
    True
    >>> isprintable(b'\01')
    False

    """
    string = string.strip()
    if not string:
        return True
    if sys.version_info[0] == 3:
        try:
            return string.isprintable()
        except Exception:
            pass
        try:
            return string.decode('utf-8').isprintable()
        except Exception:
            pass
    else:
        if string.isalnum():
            return True
        printable = ('0123456789abcdefghijklmnopqrstuvwxyzABCDEFGHIJKLMNOPQRST'
                     'UVWXYZ!"#$%&\'()*+,-./:;<=>?@[\\]^_`{|}~ \t\n\r\x0b\x0c')
        return all(c in printable for c in string)


def clean_whitespace(string, compact=False):
    """Return string with compressed whitespace."""
    for a, b in (('\r\n', '\n'), ('\r', '\n'), ('\n\n', '\n'),
                 ('\t', ' '), ('  ', ' ')):
        string = string.replace(a, b)
    if compact:
        for a, b in (('\n', ' '), ('[ ', '['),
                     ('  ', ' '), ('  ', ' '), ('  ', ' ')):
            string = string.replace(a, b)
    return string.strip()


def pformat_xml(xml):
    """Return pretty formatted XML."""
    try:
        from lxml import etree  # delayed import
        if not isinstance(xml, bytes):
            xml = xml.encode('utf-8')
        xml = etree.parse(io.BytesIO(xml))
        xml = etree.tostring(xml, pretty_print=True, xml_declaration=True,
                             encoding=xml.docinfo.encoding)
        xml = bytes2str(xml)
    except Exception:
        if isinstance(xml, bytes):
            xml = bytes2str(xml)
        xml = xml.replace('><', '>\n<')
    return xml.replace('  ', ' ').replace('\t', ' ')


def pformat(arg, width=79, height=24, compact=True):
    """Return pretty formatted representation of object as string.

    Whitespace might be altered.

    """
    if height is None or height < 1:
        height = 1024
    if width is None or width < 1:
        width = 256

    npopt = numpy.get_printoptions()
    numpy.set_printoptions(threshold=100, linewidth=width)

    if isinstance(arg, basestring):
        if arg[:5].lower() in ('<?xml', b'<?xml'):
            if isinstance(arg, bytes):
                arg = bytes2str(arg)
            if height == 1:
                arg = arg[:4*width]
            else:
                arg = pformat_xml(arg)
        elif isinstance(arg, bytes):
            if isprintable(arg):
                arg = bytes2str(arg)
                arg = clean_whitespace(arg)
            else:
                numpy.set_printoptions(**npopt)
                return hexdump(arg, width=width, height=height, modulo=1)
        arg = arg.rstrip()
    elif isinstance(arg, numpy.record):
        arg = arg.pprint()
    else:
        import pprint  # delayed import
        compact = {} if sys.version_info[0] == 2 else dict(compact=compact)
        arg = pprint.pformat(arg, width=width, **compact)

    numpy.set_printoptions(**npopt)

    if height == 1:
        arg = clean_whitespace(arg, compact=True)
        return arg[:width]

    argl = list(arg.splitlines())
    if len(argl) > height:
        arg = '\n'.join(argl[:height//2] + ['...'] + argl[-height//2:])
    return arg


def snipstr(string, width=79, snipat=None, ellipsis='...'):
    """Return string cut to specified length.

    >>> snipstr('abcdefghijklmnop', 8)
    'abc...op'

    """
    if snipat is None:
        snipat = 0.5
    if ellipsis is None:
        if isinstance(string, bytes):
            ellipsis = b'...'
        else:
            ellipsis = u'\u2026'  # does not print on win-py3.5
    esize = len(ellipsis)

    splitlines = string.splitlines()
    # TODO: finish and test multiline snip

    result = []
    for line in splitlines:
        if line is None:
            result.append(ellipsis)
            continue
        linelen = len(line)
        if linelen <= width:
            result.append(string)
            continue

        split = snipat
        if split is None or split == 1:
            split = linelen
        elif 0 < abs(split) < 1:
            split = int(math.floor(linelen * split))
        if split < 0:
            split += linelen
            if split < 0:
                split = 0

        if esize == 0 or width < esize + 1:
            if split <= 0:
                result.append(string[-width:])
            else:
                result.append(string[:width])
        elif split <= 0:
            result.append(ellipsis + string[esize-width:])
        elif split >= linelen or width < esize + 4:
            result.append(string[:width-esize] + ellipsis)
        else:
            splitlen = linelen - width + esize
            end1 = split - splitlen // 2
            end2 = end1 + splitlen
            result.append(string[:end1] + ellipsis + string[end2:])

    if isinstance(string, bytes):
        return b'\n'.join(result)
    return '\n'.join(result)


def enumarg(enum, arg):
    """Return enum member from its name or value.

    >>> enumarg(TIFF.PHOTOMETRIC, 2)
    <PHOTOMETRIC.RGB: 2>
    >>> enumarg(TIFF.PHOTOMETRIC, 'RGB')
    <PHOTOMETRIC.RGB: 2>

    """
    try:
        return enum(arg)
    except Exception:
        try:
            return enum[arg.upper()]
        except Exception:
            raise ValueError('invalid argument %s' % arg)


def parse_kwargs(kwargs, *keys, **keyvalues):
    """Return dict with keys from keys|keyvals and values from kwargs|keyvals.

    Existing keys are deleted from kwargs.

    >>> kwargs = {'one': 1, 'two': 2, 'four': 4}
    >>> kwargs2 = parse_kwargs(kwargs, 'two', 'three', four=None, five=5)
    >>> kwargs == {'one': 1}
    True
    >>> kwargs2 == {'two': 2, 'four': 4, 'five': 5}
    True

    """
    result = {}
    for key in keys:
        if key in kwargs:
            result[key] = kwargs[key]
            del kwargs[key]
    for key, value in keyvalues.items():
        if key in kwargs:
            result[key] = kwargs[key]
            del kwargs[key]
        else:
            result[key] = value
    return result


def update_kwargs(kwargs, **keyvalues):
    """Update dict with keys and values if keys do not already exist.

    >>> kwargs = {'one': 1, }
    >>> update_kwargs(kwargs, one=None, two=2)
    >>> kwargs == {'one': 1, 'two': 2}
    True

    """
    for key, value in keyvalues.items():
        if key not in kwargs:
            kwargs[key] = value


def validate_jhove(filename, jhove=None, ignore=None):
    """Validate TIFF file using jhove -m TIFF-hul.

    Raise ValueError if jhove outputs an error message unless the message
    contains one of the strings in 'ignore'.

    JHOVE does not support bigtiff or more than 50 IFDs.

    See `JHOVE TIFF-hul Module <http://jhove.sourceforge.net/tiff-hul.html>`_

    """
    import subprocess  # noqa: delayed import
    if ignore is None:
        ignore = ['More than 50 IFDs']
    if jhove is None:
        jhove = 'jhove'
    out = subprocess.check_output([jhove, filename, '-m', 'TIFF-hul'])
    if b'ErrorMessage: ' in out:
        for line in out.splitlines():
            line = line.strip()
            if line.startswith(b'ErrorMessage: '):
                error = line[14:].decode('utf8')
                for i in ignore:
                    if i in error:
                        break
                else:
                    raise ValueError(error)
                break


def lsm2bin(lsmfile, binfile=None, tile=None, verbose=True):
    """Convert [MP]TZCYX LSM file to series of BIN files.

    One BIN file containing 'ZCYX' data are created for each position, time,
    and tile. The position, time, and tile indices are encoded at the end
    of the filenames.

    """
    verbose = print_ if verbose else nullfunc

    if tile is None:
        tile = (256, 256)

    if binfile is None:
        binfile = lsmfile
    elif binfile.lower() == 'none':
        binfile = None
    if binfile:
        binfile += '_(z%ic%iy%ix%i)_m%%ip%%it%%03iy%%ix%%i.bin'

    verbose('\nOpening LSM file... ', end='', flush=True)
    start_time = time.time()

    with TiffFile(lsmfile) as lsm:
        if not lsm.is_lsm:
            verbose('\n', lsm, flush=True)
            raise ValueError('not a LSM file')
        series = lsm.series[0]  # first series contains the image data
        shape = series.shape
        axes = series.axes
        dtype = series.dtype
        size = product(shape) * dtype.itemsize

        verbose('%.3f s' % (time.time() - start_time))
        # verbose(lsm, flush=True)
        verbose('Image\n  axes:  %s\n  shape: %s\n  dtype: %s\n  size:  %s'
                % (axes, shape, dtype, format_size(size)), flush=True)
        if not series.axes.endswith('TZCYX'):
            raise ValueError('not a *TZCYX LSM file')

        verbose('Copying image from LSM to BIN files', end='', flush=True)
        start_time = time.time()
        tiles = shape[-2] // tile[-2], shape[-1] // tile[-1]
        if binfile:
            binfile = binfile % (shape[-4], shape[-3], tile[0], tile[1])
        shape = (1,) * (7-len(shape)) + shape
        # cache for ZCYX stacks and output files
        data = numpy.empty(shape[3:], dtype=dtype)
        out = numpy.empty((shape[-4], shape[-3], tile[0], tile[1]),
                          dtype=dtype)
        # iterate over Tiff pages containing data
        pages = iter(series.pages)
        for m in range(shape[0]):  # mosaic axis
            for p in range(shape[1]):  # position axis
                for t in range(shape[2]):  # time axis
                    for z in range(shape[3]):  # z slices
                        data[z] = next(pages).asarray()
                    for y in range(tiles[0]):  # tile y
                        for x in range(tiles[1]):  # tile x
                            out[:] = data[...,
                                          y*tile[0]:(y+1)*tile[0],
                                          x*tile[1]:(x+1)*tile[1]]
                            if binfile:
                                out.tofile(binfile % (m, p, t, y, x))
                            verbose('.', end='', flush=True)
        verbose(' %.3f s' % (time.time() - start_time))


def imshow(data, photometric=None, planarconfig=None, bitspersample=None,
           interpolation=None, cmap=None, vmin=None, vmax=None,
           figure=None, title=None, dpi=96, subplot=None, maxdim=None,
           **kwargs):
    """Plot n-dimensional images using matplotlib.pyplot.

    Return figure, subplot and plot axis.
    Requires pyplot already imported C{from matplotlib import pyplot}.

    Parameters
    ----------
    data : nd array
        The image data.
    photometric : {'MINISWHITE', 'MINISBLACK', 'RGB', or 'PALETTE'}
        The color space of the image data.
    planarconfig : {'CONTIG' or 'SEPARATE'}
        Defines how components of each pixel are stored.
    bitspersample : int
        Number of bits per channel in integer RGB images.
    interpolation : str
        The image interpolation method used in matplotlib.imshow. By default,
        'nearest' will be used for image dimensions <= 512, else 'bilinear'.
    cmap : str or matplotlib.colors.Colormap
        The colormap maps non-RGBA scalar data to colors.
    vmin, vmax : scalar
        Data range covered by the colormap. By default, the complete
        range of the data is covered.
    figure : matplotlib.figure.Figure
        Matplotlib figure to use for plotting.
    title : str
        Window and subplot title.
    subplot : int
        A matplotlib.pyplot.subplot axis.
    maxdim : int
        Maximum image width and length.
    kwargs : dict
        Additional arguments for matplotlib.pyplot.imshow.

    """
    # TODO: rewrite detection of isrgb, iscontig
    # TODO: use planarconfig
    if photometric is None:
        photometric = 'RGB'
    if maxdim is None:
        maxdim = 2**16
    isrgb = photometric in ('RGB', 'YCBCR')  # 'PALETTE', 'YCBCR'

    if data.dtype == 'float16':
        data = data.astype('float32')

    if data.dtype.kind == 'b':
        isrgb = False

    if isrgb and not (data.shape[-1] in (3, 4) or (
            data.ndim > 2 and data.shape[-3] in (3, 4))):
        isrgb = False
        photometric = 'MINISBLACK'

    data = data.squeeze()
    if photometric in ('MINISWHITE', 'MINISBLACK', None):
        data = reshape_nd(data, 2)
    else:
        data = reshape_nd(data, 3)

    dims = data.ndim
    if dims < 2:
        raise ValueError('not an image')
    if dims == 2:
        dims = 0
        isrgb = False
    else:
        if isrgb and data.shape[-3] in (3, 4):
            data = numpy.swapaxes(data, -3, -2)
            data = numpy.swapaxes(data, -2, -1)
        elif not isrgb and (data.shape[-1] < data.shape[-2] // 8 and
                            data.shape[-1] < data.shape[-3] // 8 and
                            data.shape[-1] < 5):
            data = numpy.swapaxes(data, -3, -1)
            data = numpy.swapaxes(data, -2, -1)
        isrgb = isrgb and data.shape[-1] in (3, 4)
        dims -= 3 if isrgb else 2

    if interpolation is None:
        threshold = 512
    elif isinstance(interpolation, int):
        threshold = interpolation
    else:
        threshold = 0

    if isrgb:
        data = data[..., :maxdim, :maxdim, :maxdim]
        if threshold:
            if (data.shape[-2] > threshold or data.shape[-3] > threshold):
                interpolation = 'bilinear'
            else:
                interpolation = 'nearest'
    else:
        data = data[..., :maxdim, :maxdim]
        if threshold:
            if (data.shape[-1] > threshold or data.shape[-2] > threshold):
                interpolation = 'bilinear'
            else:
                interpolation = 'nearest'

    if photometric == 'PALETTE' and isrgb:
        datamax = data.max()
        if datamax > 255:
            data = data >> 8  # possible precision loss
        data = data.astype('B')
    elif data.dtype.kind in 'ui':
        if not (isrgb and data.dtype.itemsize <= 1) or bitspersample is None:
            try:
                bitspersample = int(math.ceil(math.log(data.max(), 2)))
            except Exception:
                bitspersample = data.dtype.itemsize * 8
        elif not isinstance(bitspersample, inttypes):
            # bitspersample can be tuple, e.g. (5, 6, 5)
            bitspersample = data.dtype.itemsize * 8
        datamax = 2**bitspersample
        if isrgb:
            if bitspersample < 8:
                data = data << (8 - bitspersample)
            elif bitspersample > 8:
                data = data >> (bitspersample - 8)  # precision loss
            data = data.astype('B')
    elif data.dtype.kind == 'f':
        datamax = data.max()
        if isrgb and datamax > 1.0:
            if data.dtype.char == 'd':
                data = data.astype('f')
                data /= datamax
            else:
                data = data / datamax
    elif data.dtype.kind == 'b':
        datamax = 1
    elif data.dtype.kind == 'c':
        data = numpy.absolute(data)
        datamax = data.max()

    if isrgb:
        vmin = 0
    else:
        if vmax is None:
            vmax = datamax
        if vmin is None:
            if data.dtype.kind == 'i':
                dtmin = numpy.iinfo(data.dtype).min
                vmin = numpy.min(data)
                if vmin == dtmin:
                    vmin = numpy.min(data[data > dtmin])
            elif data.dtype.kind == 'f':
                dtmin = numpy.finfo(data.dtype).min
                vmin = numpy.min(data)
                if vmin == dtmin:
                    vmin = numpy.min(data[data > dtmin])
            else:
                vmin = 0

    pyplot = sys.modules['matplotlib.pyplot']

    if figure is None:
        pyplot.rc('font', family='sans-serif', weight='normal', size=8)
        figure = pyplot.figure(dpi=dpi, figsize=(10.3, 6.3), frameon=True,
                               facecolor='1.0', edgecolor='w')
        try:
            figure.canvas.manager.window.title(title)
        except Exception:
            pass
        size = len(title.splitlines()) if title else 1
        pyplot.subplots_adjust(bottom=0.03*(dims+2), top=0.98-size*0.03,
                               left=0.1, right=0.95, hspace=0.05, wspace=0.0)
    if subplot is None:
        subplot = 111
    subplot = pyplot.subplot(subplot)
    subplot.set_facecolor((0, 0, 0))

    if title:
        try:
            title = unicode(title, 'Windows-1252')
        except TypeError:
            pass
        pyplot.title(title, size=11)

    if cmap is None:
        if data.dtype.char == '?':
            cmap = 'gray'
        elif data.dtype.kind in 'buf' or vmin == 0:
            cmap = 'viridis'
        else:
            cmap = 'coolwarm'
        if photometric == 'MINISWHITE':
            cmap += '_r'

    image = pyplot.imshow(numpy.atleast_2d(data[(0,) * dims].squeeze()),
                          vmin=vmin, vmax=vmax, cmap=cmap,
                          interpolation=interpolation, **kwargs)

    if not isrgb:
        pyplot.colorbar()  # panchor=(0.55, 0.5), fraction=0.05

    def format_coord(x, y):
        # callback function to format coordinate display in toolbar
        x = int(x + 0.5)
        y = int(y + 0.5)
        try:
            if dims:
                return '%s @ %s [%4i, %4i]' % (
                    curaxdat[1][y, x], current, y, x)
            return '%s @ [%4i, %4i]' % (data[y, x], y, x)
        except IndexError:
            return ''

    def none(event):
        return ''

    subplot.format_coord = format_coord
    image.get_cursor_data = none
    image.format_cursor_data = none

    if dims:
        current = list((0,) * dims)
        curaxdat = [0, data[tuple(current)].squeeze()]
        sliders = [pyplot.Slider(
            pyplot.axes([0.125, 0.03*(axis+1), 0.725, 0.025]),
            'Dimension %i' % axis, 0, data.shape[axis]-1, 0, facecolor='0.5',
            valfmt='%%.0f [%i]' % data.shape[axis]) for axis in range(dims)]
        for slider in sliders:
            slider.drawon = False

        def set_image(current, sliders=sliders, data=data):
            # change image and redraw canvas
            curaxdat[1] = data[tuple(current)].squeeze()
            image.set_data(curaxdat[1])
            for ctrl, index in zip(sliders, current):
                ctrl.eventson = False
                ctrl.set_val(index)
                ctrl.eventson = True
            figure.canvas.draw()

        def on_changed(index, axis, data=data, current=current):
            # callback function for slider change event
            index = int(round(index))
            curaxdat[0] = axis
            if index == current[axis]:
                return
            if index >= data.shape[axis]:
                index = 0
            elif index < 0:
                index = data.shape[axis] - 1
            current[axis] = index
            set_image(current)

        def on_keypressed(event, data=data, current=current):
            # callback function for key press event
            key = event.key
            axis = curaxdat[0]
            if str(key) in '0123456789':
                on_changed(key, axis)
            elif key == 'right':
                on_changed(current[axis] + 1, axis)
            elif key == 'left':
                on_changed(current[axis] - 1, axis)
```
