## Supplementary Software for "Field dependent deep learning enables high-throughput whole-cell 3D super-resolution imaging": perlin_noise.py

```
"""Perlin noise implementation."""
# Licensed under ISC
from itertools import product
import math
import random


def smoothstep(t):
    """Smooth curve with a zero derivative at 0 and 1, making it useful for
    interpolating.
    """
    return 6*(t**5) - 15*(t**4) + 10*(t**3)  #t * t * (3. - 2. * t)


def lerp(t, a, b):
    """Linear interpolation between a and b, given a fraction t."""
    return a + t * (b - a)


class PerlinNoiseFactory(object):
    """Callable that produces Perlin noise for an arbitrary point in an
    arbitrary number of dimensions.  The underlying grid is aligned with the
    integers.
    There is no limit to the coordinates used; new gradients are generated on
    the fly as necessary.
    """

    def __init__(self, dimension, octaves=1, tile=(), unbias=False):
        """Create a new Perlin noise factory in the given number of dimensions,
        which should be an integer and at least 1.
        More octaves create a foggier and more-detailed noise pattern.  More
        than 4 octaves is rather excessive.
        ``tile`` can be used to make a seamlessly tiling pattern.  For example:
            pnf = PerlinNoiseFactory(2, tile=(0, 3))
        This will produce noise that tiles every 3 units vertically, but never
        tiles horizontally.
        If ``unbias`` is true, the smoothstep function will be applied to the
        output before returning it, to counteract some of Perlin noise's
        significant bias towards the center of its output range.
        """
        self.dimension = dimension
        self.octaves = octaves
        self.tile = tile + (0,) * dimension
        self.unbias = unbias

        # For n dimensions, the range of Perlin noise is ±sqrt(n)/2; multiply
        # by this to scale to ±1
        self.scale_factor = 2 * dimension ** -0.5

        self.gradient = {}

    def _generate_gradient(self):
        # Generate a random unit vector at each grid point -- this is the
        # "gradient" vector, in that the grid tile slopes towards it

        # 1 dimension is special, since the only unit vector is trivial;
        # instead, use a slope between -1 and 1
        if self.dimension == 1:
            return (random.uniform(-1, 1),)

        # Generate a random point on the surface of the unit n-hypersphere;
        # this is the same as a random unit vector in n dimensions.  Thanks
        # to: http://mathworld.wolfram.com/SpherePointPicking.html
        # Pick n normal random variables with stddev 1
        random_point = [random.gauss(0, 1) for _ in range(self.dimension)]
        # Then scale the result to a unit vector
        scale = sum(n * n for n in random_point) ** -0.5
        return tuple(coord * scale for coord in random_point)

    def get_plain_noise(self, *point):
        """Get plain noise for a single point, without taking into account
        either octaves or tiling.
        """
        if len(point) != self.dimension:
            raise ValueError("Expected {} values, got {}".format(
                self.dimension, len(point)))

        # Build a list of the (min, max) bounds in each dimension
        grid_coords = []
        for coord in point:
            min_coord = math.floor(coord)
            max_coord = min_coord + 1
            grid_coords.append((min_coord, max_coord))

        # Compute the dot product of each gradient vector and the point's
        # distance from the corresponding grid point.  This gives you each
        # gradient's "influence" on the chosen point.
        dots = []
        for grid_point in product(*grid_coords):
            if grid_point not in self.gradient:
                self.gradient[grid_point] = self._generate_gradient()
            gradient = self.gradient[grid_point]

            dot = 0
            for i in range(self.dimension):
                dot += gradient[i] * (point[i] - grid_point[i])
            dots.append(dot)

        # Interpolate all those dot products together.  The interpolation is
        # done with smoothstep to smooth out the slope as you pass from one
        # grid cell into the next.
        # Due to the way product() works, dot products are ordered such that
        # the last dimension alternates: (..., min), (..., max), etc.  So we
        # can interpolate adjacent pairs to "collapse" that last dimension.  Then
        # the results will alternate in their second-to-last dimension, and so
        # forth, until we only have a single value left.
        dim = self.dimension
        while len(dots) > 1:
            dim -= 1
            s = smoothstep(point[dim] - grid_coords[dim][0])

            next_dots = []
            while dots:
                next_dots.append(lerp(s, dots.pop(0), dots.pop(0)))

            dots = next_dots

        return dots[0] * self.scale_factor

    def __call__(self, *point):
        """Get the value of this Perlin noise function at the given point.  The
        number of values given should match the number of dimensions.
        """
        ret = 0
        for o in range(self.octaves):
            o2 = 1 << o
            new_point = []
            for i, coord in enumerate(point):
                coord *= o2
                if self.tile[i]:
                    coord %= self.tile[i] * o2
                new_point.append(coord)
            ret += self.get_plain_noise(*new_point) / o2

        # Need to scale n back down since adding all those extra octaves has
        # probably expanded it beyond ±1
        # 1 octave: ±1
        # 2 octaves: ±1½
        # 3 octaves: ±1¾
        ret /= 2 - 2 ** (1 - self.octaves)

        if self.unbias:
            # The output of the plain Perlin noise algorithm has a fairly
            # strong bias towards the center due to the central limit theorem
            # -- in fact the top and bottom 1/8 virtually never happen.  That's
            # a quarter of our entire output range!  If only we had a function
            # in [0..1] that could introduce a bias towards the endpoints...
            r = (ret + 1) / 2
            # Doing it this many times is a completely made-up heuristic.
            for _ in range(int(self.octaves / 2 + 0.5)):
                r = smoothstep(r)
            ret = r * 2 - 1

        return ret
```
