## Supplementary Software for "Field dependent deep learning enables high-throughput whole-cell 3D super-resolution imaging": plot_funcs.py

```
from matplotlib import pyplot as plt
from matplotlib import gridspec
import pickle
import numpy as np
import seaborn as sns
import os
import scipy.signal

from matplotlib.colors import hsv_to_rgb
from matplotlib.colors import rgb_to_hsv
import cv2
import PIL
from PIL import ImageEnhance
from PIL import ImageTk

from local_utils.utils import *
from local_utils.csv_utils import *


def plot_od(od, label=None, col=None):
    """Produces a line plot from a ordered dictionary as used to store training process in the Model class
    
    Parameters
    ----------
    od: OrderedDict of floats
        DECODE model
    label: str
        Label
    col: 'str'
        Color
    """
    plt.plot(*zip(*sorted(od.items())), label=label, color=col)


def create_3d_hist(preds, z_clip=None, pix_size=5, sigma=3, contrast_fac=10, clip_density=100):
    """Produces a coloured histogram to display 3D reconstructions.
    
    Parameters
    ----------
    preds: list
        List of localizations with columns: 'localization', 'frame', 'x', 'y', 'z'
    z_clip: list of ints
        Clips the the z values at the given lower and upper limit to control the colorrange. 
    pix_size: float
        Size of the pixel (nano meter) in the reconstruction plot
    sigma:
        Size of Gaussian used for blurring
    constrast fact: float
        Contrast can be scaled with this variable if the output image is to bright/dark
    clip_density: float 
        Percentile between 0 and 100. Artifacts that produce extremely dense regions in the histrogram can 
        mess up the contrast scaling. This parameter can be used to exclude the brightest regions. 
        
    Returns
    -------
    Image: PIL image
        Coloured histogram of 3D reconstruction
    """
    # adjust colormap
    lin_hue = np.linspace(0, 1, 256)
    cmap = plt.get_cmap('jet', lut=256);
    cmap = cmap(lin_hue)
    cmap_hsv = rgb_to_hsv(cmap[:, :3])
    storm_hue = cmap_hsv[:, 0]
    _, b = np.unique(storm_hue, return_index=True)
    storm_hue = [storm_hue[index] for index in sorted(b)]
    n_val = len(storm_hue)
    storm_hue = np.interp(np.linspace(0, n_val, 256), np.arange(n_val), storm_hue)

    x_pos = np.clip(np.array(preds)[:, 2], 0, np.inf)
    y_pos = np.clip(np.array(preds)[:, 3], 0, np.inf)
    z_pos = np.array(preds)[:, 4]

    min_z = min(z_pos)
    max_z = max(z_pos)

    if z_clip is not None:
        z_pos[z_pos < z_clip[0]] = z_clip[0]
        z_pos[z_pos > z_clip[1]] = z_clip[1]
        zc_val = (z_pos - z_clip[0]) / (z_clip[1] - z_clip[0])

    else:
        zc_val = (z_pos - min_z) / (max_z - min_z)

    z_hue = np.interp(zc_val, lin_hue, storm_hue)

    nx = int((np.max(x_pos)) // pix_size + 1)
    ny = int((np.max(y_pos)) // pix_size + 1)
    dims = (nx, ny)

    x_vals = np.array(x_pos // pix_size, dtype='int')
    y_vals = np.array(y_pos // pix_size, dtype='int')

    lin_idx = np.ravel_multi_index((x_vals, y_vals), dims)
    density = np.bincount(lin_idx, weights=np.ones(len(lin_idx)), minlength=np.prod(dims)).reshape(dims)
    density = np.clip(density, 0, np.percentile(density, clip_density))
    zsum = np.bincount(lin_idx, weights=z_hue, minlength=np.prod(dims)).reshape(dims)
    zavg = zsum / density
    zavg[np.isnan(zavg)] = 0

    hue = zavg[:, :, None]
    sat = np.ones(density.shape)[:, :, None]
    val = (density / np.max(density))[:, :, None]
    sr_HSV = np.concatenate((hue, sat, val), 2)
    sr_RGB = hsv_to_rgb(sr_HSV)
    # %have to gaussian blur in rgb domain
    sr_RGBblur = cv2.GaussianBlur(sr_RGB, (11, 11), sigma / pix_size)
    sr_HSVblur = rgb_to_hsv(sr_RGBblur)

    val = sr_HSVblur[:, :, 2]

    sr_HSVfinal = np.concatenate((sr_HSVblur[:, :, :2], val[:, :, None]), 2)
    sr_RGBfinal = hsv_to_rgb(sr_HSVfinal)

    sr_Im = PIL.Image.fromarray(np.array(np.round(sr_RGBfinal * 256), dtype='uint8'))
    enhancer = ImageEnhance.Contrast(sr_Im)
    sr_Im = enhancer.enhance(contrast_fac)

    return sr_Im.transpose(PIL.Image.TRANSPOSE)


def create_2d_hist(preds, pix_size=5, sigma=3, contrast_fac=2, clip_density=100):
    """Produces a coloured histogram to display 3D reconstructions.
    
    Parameters
    ----------
    preds: list
        List of localizations with columns: 'localization', 'frame', 'x', 'y', 'z'
    pix_size: float
        Size of the pixel (nano meter) in the reconstruction plot
    sigma:
        Size of Gaussian used for blurring
    constrast fact: float
        Contrast can be scaled with this variable if the output image is to bright/dark
    clip_density: float 
        Percentile between 0 and 100. Artifacts that produce extremely dense regions in the histrogram can 
        mess up the contrast scaling. This parameter can be used to exclude the brightest regions. 
        
    Returns
    -------
    sr_blur: array
        Histogram of 2D reconstruction
    """

    x_pos = np.clip(np.array(preds)[:, 2], 0, np.inf)
    y_pos = np.clip(np.array(preds)[:, 3], 0, np.inf)

    nx = int((np.max(x_pos)) // pix_size + 1)
    ny = int((np.max(y_pos)) // pix_size + 1)

    dims = (nx, ny)

    x_vals = np.array(x_pos // pix_size, dtype='int')
    y_vals = np.array(y_pos // pix_size, dtype='int')

    lin_idx = np.ravel_multi_index((x_vals, y_vals), dims)
    density = np.bincount(lin_idx, weights=np.ones(len(lin_idx)), minlength=np.prod(dims)).reshape(dims)
    density = np.clip(density, 0, np.percentile(density, clip_density))

    val = (density / np.max(density)).T[:, :, None]

    sr_blur = cv2.GaussianBlur(val, (3, 3), sigma / pix_size)
    sr_blur = np.clip(sr_blur, 0, sr_blur.max() / contrast_fac)

    return sr_blur


def plot_sample_predictions(model, plot_infs, eval_csv, plot_num, fov_size, pixel_size):

    h, w = plot_infs[0]['raw_img'].shape[0], plot_infs[0]['raw_img'].shape[1]
    rows, columns = plot_infs[0]['rows'], plot_infs[0]['columns']
    win_size = plot_infs[0]['win_size']

    img_infs = {}
    for k in plot_infs[1]:
        img_infs[k] = np.zeros([h, w])
        for i in range(len(plot_infs)-1):
            row_start = i // columns * win_size
            column_start = i % columns * win_size

            img_infs[k][row_start:row_start+win_size if row_start+win_size < h else h,
                        column_start:column_start+win_size if column_start+win_size < w else w] = plot_infs[i+1][k]

    fig1, ax_arr = plt.subplots(4, 3, figsize=(9,12), constrained_layout=True)
    # plt.subplots_adjust(wspace=0.5, hspace=0.5)
    ax = []
    datas = []
    plts = []
    for i in ax_arr:
        for j in i:
            ax.append(j)

    datas.append(plot_infs[0]['raw_img'])
    # datas.append(img_infs['Samples_ps'])  # 决定分子最终所在的像素位置的布尔型图
    datas.append(plot_infs[0]['raw_img'])
    datas.append(img_infs['Probs'])
    datas.append(img_infs['XO'])
    datas.append(img_infs['YO'])
    datas.append(img_infs['ZO'])
    datas.append(img_infs['Int'])
    datas.append(img_infs['XO_sig'])
    datas.append(img_infs['YO_sig'])
    datas.append(img_infs['ZO_sig'])
    if model.psf_pred:
        datas.append(img_infs['BG'])
        datas.append(scipy.signal.medfilt2d(plot_infs[0]['raw_img'] - img_infs['BG'], kernel_size=9))
    titles = ['Image', 'deterministic locs', 'Inferred probabilities',  'Inferred x-offset',
              'Inferred y-offset', 'Inferred z-offset', 'Intensity', 'X-offset_uncertainty',
              'Y-offset_uncertainty', 'Z-offset_uncertainty', 'Predicted raw image', 'Background']

    for i in range(len(datas)):  # fushuang
        plts.append(ax[i].imshow(datas[i]))
        ax[i].set_title(titles[i], fontsize=10)
        plt.colorbar(plts[i], ax=ax[i], fraction=0.046, pad=0.04)

    for x,y in zip(img_infs['Samples_ps'].nonzero()[1],img_infs['Samples_ps'].nonzero()[0]):
        ax[1].add_patch(plt.Circle((x, y), radius=1.5, color='cyan', fill=True, lw=0.5, alpha=0.8))

    if eval_csv is not None:
        # 以ground truth的pixel近似为中心画圈圈
        truth_map = get_pixel_truth(eval_csv, plot_num , fov_size, pixel_size)
        for x, y in zip(truth_map.nonzero()[0], truth_map.nonzero()[1]):
            # circ = plt.Circle((x, y), radius=3, color='g', fill=False, lw=0.5)
            ax[1].add_patch(plt.Circle((x, y), radius=3, color='g', fill=False, lw=0.5))
            ax[2].add_patch(plt.Circle((x, y), radius=3, color='g', fill=False, lw=0.5))

    fig_fs,ax_fs=plt.subplots(1,1,figsize=(6,6),constrained_layout=True)
    plt.colorbar(ax_fs.imshow(datas[0], cmap='gray'), ax=ax_fs, fraction=0.046, pad=0.04)
    for x,y in zip(img_infs['Samples_ps'].nonzero()[1],img_infs['Samples_ps'].nonzero()[0]):
        ax_fs.add_patch(plt.Circle((x, y), radius=1.5, color='cyan', fill=True, lw=0.5, alpha=0.8))


    # plt.figure()
    # plt.imshow(datas[2])
    # plt.figure()
    # plt.imshow(datas[1])
    # plt.tight_layout()
    plt.show()


def plot_train_record(model):
    plt.figure(figsize=(9,6),constrained_layout=True)
    plt.subplot(3,3,1);plot_od(model.recorder['rmse_lat']);plt.xlabel('iterations');plt.ylabel('RMSE_Lateral')
    plt.subplot(3,3,2);plot_od(model.recorder['rmse_ax']);plt.xlabel('iterations');plt.ylabel('RMSE_Axial')
    plt.subplot(3,3,3);plot_od(model.recorder['rmse_vol']);plt.xlabel('iterations');plt.ylabel('RMSE_Voxel')
    plt.subplot(3,3,4);plot_od(model.recorder['eff_lat']);plt.xlabel('iterations');plt.ylabel('lateral efficiency')
    plt.subplot(3,3,5);plot_od(model.recorder['eff_3d']);plt.xlabel('iterations');plt.ylabel('3D efficiency')
    plt.subplot(3,3,6);plot_od(model.recorder['recall']);plt.xlabel('iterations');plt.ylabel('recall')
    plt.subplot(3,3,7);plot_od(model.recorder['precision']);plt.xlabel('iterations');plt.ylabel('precision')
    plt.subplot(3,3,8);plot_od(model.recorder['jaccard']);plt.xlabel('iterations');plt.ylabel('jaccard')
    plt.subplot(3,3,9);plot_od(model.recorder['cost_hist']);plt.xlabel('iterations');plt.ylabel('cost')
    # plt.subplots_adjust(wspace=0.5,hspace=0.5)
    # plt.tight_layout()
    plt.show()

def plot_preds_distribution(preds,preds_final):
    fig,axes = plt.subplots(2,2)
    axes[0,0].hist(np.array(preds)[:, 6], bins=50)
    axes[0,0].axvspan(np.array(preds_final)[:, 6].min(),np.array(preds_final)[:, 6].max(),color='green', alpha=0.1)
    axes[0,0].set_xlabel(r'$nms-p$')
    axes[0,0].set_ylabel('counts')

    axes[0,1].hist(np.array(preds)[:, 7], bins=50)
    axes[0,1].axvspan(0, np.array(preds_final)[:, 7].max(),color='green', alpha=0.1)
    axes[0,1].set_xlabel(r'$\sigma_x$ [nm]')
    axes[0,1].set_ylabel('counts')

    axes[1,0].hist(np.array(preds)[:, 8], bins=50)
    axes[1,0].axvspan(0, np.array(preds_final)[:, 8].max(),color='green', alpha=0.1)
    axes[1,0].set_xlabel(r'$\sigma_y$ [nm]')
    axes[1,0].set_ylabel('counts')

    axes[1,1].hist(np.array(preds)[:, 9], bins=50)
    axes[1,1].axvspan(0, np.array(preds_final)[:, 9].max(),color='green', alpha=0.1)
    axes[1,1].set_xlabel(r'$\sigma_z$ [nm]')
    axes[1,1].set_ylabel('counts')
    plt.tight_layout()
    plt.show()
    return fig,axes
```
