## Supplementary Software for "Field dependent deep learning enables high-throughput whole-cell 3D super-resolution imaging": utils.py

```
import numpy as np
import torch 
import csv
import copy
#import scipy.stats as stats
import scipy.stats
from matplotlib import pyplot as plt
from operator import itemgetter
import random
from local_utils import *

def gpu(x):
    '''Transforms numpy array or torch tensor it torch.cuda.FloatTensor'''
    if isinstance(x, np.ndarray):
        return torch.cuda.FloatTensor(x.astype('float32'))
    else:
        return torch.cuda.FloatTensor(x)

def cpu(x):
    '''Transforms torch tensor into numpy array'''
    return x.cpu().detach().numpy()

def softp(x):
    '''Returns softplus(x)'''
    return(np.log(1+np.exp(x)))

def sigmoid(x):
    '''Returns sigmoid(x)'''
    return 1 / (1 + np.exp(-x))

def inv_softp(x):
    '''Returns inverse softplus(x)'''
    return np.log(np.exp(x)-1)

def inv_sigmoid(x):
    '''Returns inverse sigmoid(x)'''
    return -np.log(1/x-1)

def torch_arctanh(x):
    '''Returns arctanh(x) for tensor input'''
    return 0.5*torch.log(1+x) - 0.5*torch.log(1-x)

def torch_softp(x):
    '''Returns softplus(x) for tensor input'''
    return (torch.log(1+torch.exp(x)))

def flip_filt(filt):
    '''Returns filter flipped over x and y dimension'''
    return np.ascontiguousarray(filt[...,::-1,::-1])


def get_bg_stats(images, percentile=10,plot=False,xlim=None, floc=0):
    """Infers the parameters of a gamma distribution that fit the background of SMLM recordings. 
    Identifies the darkest pixels from the averaged images as background and fits a gamma distribution to the histogram of intensity values.
    
    Parameters
    ----------
    images: array
        3D array of recordings
    percentile: float
        Percentile between 0 and 100. Sets the percentage of pixels that are assumed to only containg background activity (i.e. no fluorescent signal)
    plot: bool
        If true produces a plot of the histogram and fit
    xlim: list of floats
        Sets xlim of the plot
    floc: float
        Baseline for the the gamma fit. Equal to fitting gamma to (x - floc)
        
    Returns
    -------
    mean, scale: float
        Mean and scale parameter of the gamma fit
    """
    # 确保图片为正值
    ind = np.where(images <= 0)
    images[ind] = 1

    # 先将图像中每个位置求全部帧的平均，然后选出10%位置的值，然后得到小于这个值的位置的坐标
    map_empty = np.where(images.mean(0) < np.percentile(images.mean(0),percentile))
    # 取出imagestack中这些位置的所有值
    pixel_vals = images[:,map_empty[0],map_empty[1]].reshape(-1)
    # 调用scipy库的gamma拟合,返回的是alpha和scale=1/beta
    fit_alpha, fit_loc, fit_beta=scipy.stats.gamma.fit(pixel_vals,floc=floc)
    
    if plot:
        plt.figure(constrained_layout=True)
        if xlim is None: 
            low,high = pixel_vals.min(),pixel_vals.max()
        else:
            low,high = xlim[0],xlim[1]

        _ = plt.hist(pixel_vals, bins=np.linspace(low,high), histtype='step', label='data')
        _ = plt.hist(np.random.gamma(shape=fit_alpha,scale=fit_beta,size=len(pixel_vals))+floc, bins=np.linspace(low,high), histtype='step', label='fit')
        plt.xlim(low,high)
        plt.legend()
        # plt.tight_layout()
        plt.show()
    return fit_alpha*fit_beta,fit_beta  # 返回gamma分布的期望


def get_window_map(img, winsize=40, percentile=20):
    """Helper function 
    
    Parameters
    ----------
    images: array
        3D array of recordings
    percentile: float
        Percentile between 0 and 100. Sets the percentage of pixels that are assumed to only containg background activity (i.e. no fluorescent signal)
    plot: bool
        If true produces a plot of the histogram and fit
    xlim: list of floats
        Sets xlim of the plot
    floc: float
        Baseline for the the gamma fit. Equal to fitting gamma to (x - floc)
        
    Returns
    -------
    binmap: array
        Mean and scale parameter of the gamma fit
    """      
    img = img.mean(0)  # 按第一维求平均,得到[64 64]
    res = np.zeros([int(img.shape[0]-winsize), int(img.shape[1]-winsize)])  # [64-40，64-40]的零矩阵
    for i in range(res.shape[0]):  # 0-24
        for j in range(res.shape[1]):
            res[i,j] = img[i:i+int(winsize), j:j+int(winsize)].mean()  # 以i j出发求[40 40]区域内的平均值
    thresh = np.percentile(res,percentile)  # 从小到大，第percentile%的值，也就是还有percentile%比这个值小
    binmap = np.zeros_like(res)
    binmap[res>thresh] = 1  # 图像中intensity大于20%的都设为1，应该是表示该处有分子荧光
    return binmap


def find_frame(molecule_list, frame_nbr):
    """find the molecule list index of the specified frame_number then plus 1 (the molecule list is sorted)"""
    list_index = None
    for i,molecule in enumerate(molecule_list):
        if molecule[1] > frame_nbr:
            list_index = i
            break
    return list_index


def get_pixel_truth(eval_csv, ind, field_size, pixel_size):
    """draw a pixel-wise binary map of the groudtruth"""
    eval_list = []
    if isinstance(eval_csv, str):
        with open(eval_csv, 'r') as csvfile:
            reader = csv.reader(csvfile, delimiter=',')
            for row in reader:
                if 'truth' not in row[0]:
                    eval_list.append([float(r) for r in row])
    else:
        for r in eval_csv:
            eval_list.append([i for i in r])
    eval_list = sorted(eval_list, key=itemgetter(1))  # csv文件按frame升序来排列

    molecule_list=[]
    for i in range(len(eval_list)):
        if eval_list[i][1]==ind:
            molecule_list.append([round(eval_list[i][2]/pixel_size[0]),round(eval_list[i][3]/pixel_size[1])])
        if eval_list[i][1]>ind:
            break

    truth_map = np.zeros([round(field_size[0] / pixel_size[0]), round(field_size[1] // pixel_size[1])])
    for molecule in molecule_list:
        truth_map[molecule[0],molecule[1]]=1

    return truth_map


def setup_seed(seed):
    torch.manual_seed(seed)
    torch.cuda.manual_seed_all(seed)
    np.random.seed(seed)
    random.seed(seed)
    torch.backends.cudnn.deterministic = True
    torch.backends.cudnn.benchmark = False

def otf_gauss2D(shape=(3,3),Isigmax=0.5,Isigmay=0.5):
    """
    2D gaussian mask - should give the same result as MATLAB's otf rescale
    """
    m,n = [(ss-1.)/2. for ss in shape]
    y,x = np.ogrid[-m:m+1,-n:n+1]
    h = np.exp( -(x*x)/(2.*Isigmax*Isigmax+1e-6)- (y*y)/(2.*Isigmay*Isigmay+1e-6) )
    h[ h < np.finfo(h.dtype).eps*h.max() ] = 0
    sumh = h.sum()
    if sumh != 0:
        h /= sumh
    return h

def read_first_size_gb_tiff(image_path, size_gb=4):
    with TiffFile(image_path, is_ome=False) as tif:
        total_shape = tif.series[0].shape
        occu_mem = total_shape[0] * total_shape[1] * total_shape[2] * 16 / (1024 ** 3) / 8
        if occu_mem<size_gb:
            index_img=total_shape[0]
        else:
            index_img = int(size_gb/occu_mem*total_shape[0])
        images = tif.asarray(key=range(0,index_img), series=0)
    print("read first %d images" % (images.shape[0]))
    return images
```
