## Supplementary Software for "Field dependent deep learning enables high-throughput whole-cell 3D super-resolution imaging": _PSF_vector_gpu.py

```
from ctypes import *

import numpy as np
import torch.nn as nn

from local_utils.utils import *


class PSF_VECTOR_GPU:

    def sim_psf_vector(self, S, X_os, Y_os, Z, aber_map, field_xy, robust_training=False):
        """Generate the PSFs based on C code

        Parameters
        ----------
        S:
            Coordinates of the pixel which has a molecule [row, column]
        X_os:
            Sub_pixel x offset
        Y_os:
            Sub_pixel y offset
        Z:
            Z position
        aber_map:
            Aberration map
        field_xy:
            Selected sub-area [x_start, x_end, y_start, y_end] of the whole aberration map, it should be noted
            x corresponds to column, y corresponds to the row
        robust_training: bool
            If True, add small zernike disturbance to the simulated PSFs
        """
        try:
            psf = CDLL("PSF_vector_gpu/psf_simu_gpu.dll", winmode=0)
        except FileNotFoundError:
            psf = CDLL("../../PSF_vector_gpu/psf_simu_gpu.dll", winmode=0)

        # psf = CDLL("F:/projects/FS_work/psf_simu_gpu_LTD/x64/Debug/psf_simu_gpu.dll", winmode=0)

        class sysParas_(Structure):
            _fields_ = [
                ('aberrations_', POINTER(c_float)),
                ('NA_', c_float),
                ('refmed_', c_float),
                ('refcov_', c_float),
                ('refimm_', c_float),
                ('lambdaX_', c_float),
                ('objStage0_', c_float),
                ('zemit0_', c_float),
                ('pixelSizeX_', c_float),
                ('pixelSizeY_', c_float),
                ('sizeX_', c_float),
                ('sizeY_', c_float),
                ('PupilSize_', c_float),
                ('Npupil_', c_float),
                ('zernikeModesN_', c_int),
                ('xemit_', POINTER(c_float)),
                ('yemit_', POINTER(c_float)),
                ('zemit_', POINTER(c_float)),
                ('objstage_', POINTER(c_float)),
                ('aberrationsParas_', POINTER(c_float)),
                ('psfOut_', POINTER(c_float)),
                ('aberrationOut_', POINTER(c_float)),
                ('Nmol_', c_int),
                ('showAberrationNumber_', c_int)
            ]

        sP = sysParas_()

        # In pyInterfacePSFfSimu.cPSFf(), the x y positions should be inverse to ensure the input X_os corresponds to
        # the column
        npXemit = np.array(np.squeeze(Y_os) * self.psf_pars['pixel_size_xy'][1], dtype=np.float32)  # nm
        npYemit = np.array(np.squeeze(X_os) * self.psf_pars['pixel_size_xy'][0], dtype=np.float32)
        npZemit = np.array(1 * np.squeeze(Z), dtype=np.float32)
        npObjStage = np.array(0 * (np.squeeze(Z)), dtype=np.float32)

        npSizeX = self.psf_pars['psf_size']
        npSizeY = self.psf_pars['psf_size']
        sP.Npupil_ = 64
        sP.Nmol_ = npXemit.size
        sP.NA_ = self.psf_pars['NA']
        sP.refmed_ = self.psf_pars['refmed']
        sP.refcov_ = self.psf_pars['refcov']
        sP.refimm_ = self.psf_pars['refimm']
        sP.lambdaX_ = self.psf_pars['lambda']
        sP.objStage0_ = self.psf_pars['initial_obj_stage']
        sP.zemit0_ = -1 * sP.refmed_ / sP.refimm_ * (sP.objStage0_)
        sP.pixelSizeX_ = self.psf_pars['pixel_size_xy'][1]
        sP.pixelSizeY_ = self.psf_pars['pixel_size_xy'][0]
        sP.zernikeModesN_ = 21
        sP.sizeX_ = npSizeX
        sP.sizeY_ = npSizeY
        sP.PupilSize_ = 1.0
        sP.showAberrationNumber_ = 1
        zernikeModes = np.array([2, -2, 0, 2, 2, 0, 3, -1, 0, 3, 1, 0, 4, 0, 0, 3, -3, 0, 3, 3, 0,
                                 4, -2, 0, 4, 2, 0, 5, -1, 0, 5, 1, 0, 6, 0, 0, 4, -4, 0, 4, 4, 0,
                                 5, -3, 0, 5, 3, 0, 6, -2, 0, 6, 2, 0, 7, 1, 0, 7, -1, 0, 8, 0, 0],
                                dtype=np.float32).reshape(21, 3).T
        zernikeModes[2, :] = zernikeModes[2, :] * sP.lambdaX_
        zernikeModes = zernikeModes.reshape(3, 21).flatten()

        sP.aberrations_ = zernikeModes.ctypes.data_as(POINTER(c_float))

        Nmol = sP.Nmol_
        # npXYZOBuffer = np.empty((4, Nmol), dtype=np.float32, order='C')
        npPSFBuffer = np.empty((Nmol, npSizeX, npSizeY), dtype=np.float32, order='C')
        npWaberrationBuffer = np.empty((int(sP.Npupil_), int(sP.Npupil_)), dtype=np.float32, order='C')
        aberrationsParas = np.empty((Nmol, sP.zernikeModesN_), dtype=np.float32, order='C')

        sP.xemit_ = npXemit.ctypes.data_as(POINTER(c_float))
        sP.yemit_ = npYemit.ctypes.data_as(POINTER(c_float))
        sP.zemit_ = npZemit.ctypes.data_as(POINTER(c_float))
        sP.objstage_ = npObjStage.ctypes.data_as(POINTER(c_float))

        aber_map_crop = aber_map[field_xy[2]:field_xy[3] + 1, field_xy[0]:field_xy[1] + 1, :]
        # robust_training means training data will randomly generate aberrations to the PSF, as we can not
        # measure the aberration accurately, and model mismatch will cause many artifacts, we think generate
        # data with some random aberrations can help network generalize
        if robust_training:
            for n in range(0, Nmol):
                aberrationsParas[n] = aber_map_crop[S[n, 1], S[n, 2], 0:21] + \
                                      np.random.normal(loc=0, scale=0.01, size=21)
        else:
            for n in range(0, Nmol):
                aberrationsParas[n] = aber_map_crop[S[n, 1], S[n, 2], 0:21]

        aberrationsParas = aberrationsParas * sP.lambdaX_  # multiply with lambda
        sP.aberrationsParas_ = aberrationsParas.ctypes.data_as(POINTER(c_float))
        sP.psfOut_ = npPSFBuffer.ctypes.data_as(POINTER(c_float))

        sP.aberrationOut_ = npWaberrationBuffer.ctypes.data_as(POINTER(c_float))
        psf.vectorPSFF1(sP)  # run

        if np.size(np.where(np.isnan(npPSFBuffer))) != 0:
            print('nan in the gpu psf!!!', np.where(np.isnan(npPSFBuffer)))

        # otf rescale
        npPSFBuffer = gpu(npPSFBuffer)
        for i in range(len(npPSFBuffer)):
            h = gpu(otf_gauss2D(shape=[5, 5], Isigmax=aber_map_crop[S[i, 1], S[i, 2], 21],
                                Isigmay=aber_map_crop[S[i, 1], S[i, 2], 22])).reshape([1, 1, 5, 5])
            tmp = nn.functional.conv2d(
                npPSFBuffer[i].reshape(1, 1, self.psf_pars['psf_size'], self.psf_pars['psf_size'])
                , h, padding=2, stride=1)
            npPSFBuffer[i] = tmp.reshape(self.psf_pars['psf_size'], self.psf_pars['psf_size'])

        return npPSFBuffer
```
