## Supplementary Software for "Field dependent deep learning enables high-throughput whole-cell 3D super-resolution imaging": Example_SMLM.m

```
% Copyright (c) 2022 Li Lab, Southern University of Science and Technology, Shenzhen
% author: Li-Lab
%
% date: 2022.05.05
% Tested with CUDA 10.1 (Express installation) and Matlab R2019a
%%
clear all
clc
addpath source
frame_size = 64;  % pixel, the image size
N_pixels = 27;     % pixel, the psf size of beads
pixel_size = 100; %nm, pixel size of the image
Nmol = 1;
Nphotons = 30000 +5000*rand(1,Nmol); %simulated the photons per beads
bg = 100 + 100*rand(1,Nmol);         %simulated the background of image

z_depth = 2000;               %nm, the depth of bead stack
z_step =50;                   %nm, the step of bead stack in each frame

%% parameters for PSF model used for simulated
paraSim.NA = 1.35;                                                      % numerical aperture of objective lens             
paraSim.refmed = 1.34;                                                  % refractive index of sample medium
paraSim.refcov = 1.525;                                                 % refractive index of converslip
paraSim.refimm = 1.406;                                                 % refractive index of immersion oil
paraSim.lambda = 680;                                                   % wavelength of emission
paraSim.objStage0 = -0;                                              % nm, initial objStage0 position,relative to focus at coverslip
paraSim.zemit0 = -1*paraSim.refmed/paraSim.refimm*(paraSim.objStage0);  % reference emitter z position, nm, distance of molecule to coverslip
paraSim.pixelSizeX = pixel_size;                                        % nm, pixel size of the image
paraSim.pixelSizeY = pixel_size;                                        % nm, pixel size of the image
paraSim.Npupil = 64;                                                    % sampling at the pupil plane
%simulated zernike polynomial coefficients for an astigmatic PSF
paraSim.aberrations = [ 2.0000   -2.0000    0.3706                      
                        2.0000    2.0000   72.7186
                        3.0000   -1.0000   19.0738
                        3.0000    1.0000    1.5904
                        4.0000         0   -1.2268
                        3.0000   -3.0000   -2.1291
                        3.0000    3.0000    2.9314
                        4.0000   -2.0000   -2.0293
                        4.0000    2.0000   -7.1634
                        5.0000   -1.0000    0.1961
                        5.0000    1.0000   -8.0279
                        6.0000         0    8.9532
                        4.0000   -4.0000    0.2143
                        4.0000    4.0000   -0.1633
                        5.0000   -3.0000    0.1333
                        5.0000    3.0000    0.2868
                        6.0000   -2.0000    0.8603
                        6.0000    2.0000   -0.9256
                        7.0000    1.0000   -2.2734
                        7.0000   -1.0000   -0.3728
                        8.0000         0   11.1233];
                    
paraSim.Nmol = 1; 
paraSim.sizeX = N_pixels;
paraSim.sizeY = N_pixels;
paraSim.xemit = 0;
paraSim.yemit = 0;
paraSim.zemit = 0;
paraSim.objStage = 0;

z=z_depth/2:-z_step:-z_depth/2;
stacks_num = length(z);
PSF_stack =zeros(N_pixels,N_pixels,stacks_num);
PSF_stack =[];
for idx_stack = 1:stacks_num
    paraSim.aberrationsParas=[];
    for j = 1:1
        paraSim.objStage = -((idx_stack-1)*z_step-z_depth/2);
        paraSim.showAberrationNumber=j;
        paraSim.aberrationsParas(j,:) = paraSim.aberrations(:,3); 
       % GPU simulated
%         [PSFs, Waberration]=psf_gpu_simu2(paraSim); 
        % CPU simulated
        [PSFs, Waberration]=psf_simu2_floatC(paraSim);
    end
    PSFs = PSFs.*Nphotons+bg;  
    PSF_stack(:,:,idx_stack)=PSFs;
end
PSF_stack = poissrnd(PSF_stack);
%%  plot PSF stack 

f = figure
initPosition = f.Position;
f.Position = [initPosition(1), initPosition(2)-900+initPosition(4),900, 900];

for k=1:length(PSF_stack)
    subplot(7,7,k);
    imagesc(PSF_stack(:,:,k)) ;
    sub_title=title([ num2str((k-1)*z_step-z_depth/2),' nm']);
    set(gca,'FontName','time','FontSize',10,'FontWeight','bold');
    set(sub_title,'FontName','time','FontSize',12,'LineWidth',3,'FontWeight','bold'); 
end

%%  parameters for PSF model used for fit
paraFit.NA = paraSim.NA;                                                    % numerical aperture of objective lens             
paraFit.refmed = paraSim.refmed;                                            % refractive index of sample medium
paraFit.refcov = paraSim.refcov;                                            % refractive index of converslip
paraFit.refimm =paraSim. refimm;                                            % refractive index of immersion oil
paraFit.lambda = paraSim.lambda;                                             % wavelength of emission
paraFit.zemit0 = paraSim.zemit0;                                            % reference emitter z position, nm, distance of molecule to coverslip
paraFit.objStage0 = paraSim.objStage0;                                            %  nm, initial objStage0 position,relative to focus at coverslip
paraFit. pixelSizeX = paraSim.pixelSizeX;                                        % nm, pixel size of the image
paraFit. pixelSizeY = paraSim.pixelSizeX;                                        % nm, pixel size of the image
paraFit.Npupil =paraSim.Npupil;                                             % sampling at the pupil plane
paraFit.sizeX = size(PSF_stack,1);
paraFit.sizeY = size(PSF_stack,2);
paraFit.sizeZ = size(PSF_stack,3);
%initializing zernike polynomial coefficients
paraFit.aberrations = [2,-2,0.0; 2,2,0.0; 3,-1,0.0; 3,1,0.0; 4,0,0.0; 3,-3,0.0; 3,3,0.0; 4,-2,0.0; 4,2,0.0; 5,-1,0.0; 5,1,0.0; 6,0,0.0; 4,-4,0.0; 4,4,0.0;  5,-3,0.0; 5,3,0.0;  6,-2,0.0; 6,2,0.0; 7,1,0.0; 7,-1,0.00; 8,0,0];

%% initial parameters for fit
Nz = size(PSF_stack,3);
Npixel = size(PSF_stack,1);
numAberrations = size(paraFit.aberrations,1);
shared = [ones(1,numAberrations) 1 1 1 0 0];  % 1 is shared parameters between z slices, 0 is free parameters between z slices, only consider  [x, y, z, I, bg]

sumShared = sum(shared);
numparams = 26*Nz-sumShared*(Nz-1);
thetainit = zeros(numparams,1);
bg = zeros(1,Nz);
Nph = zeros(1,Nz);
x0 = zeros(1,Nz);
y0 = zeros(1,Nz);
z0 = zeros(1,Nz);

% center of mass with nm unit
ImageSizex = paraFit.pixelSizeX*Npixel/2;
ImageSizey = paraFit.pixelSizeY*Npixel/2;

DxImage = 2*ImageSizex/paraFit.sizeX;
DyImage = 2*ImageSizey/paraFit.sizeY;
ximagelin = -ImageSizex+DxImage/2:DxImage:ImageSizex;
yimagelin = -ImageSizey+DyImage/2:DyImage:ImageSizey;
[YImage,XImage] = meshgrid(yimagelin,ximagelin);
for i = 1:Nz
    dTemp = PSF_stack(:,:,i);
    bg(i) = min(dTemp(:));
    bg(i) = max(bg(i),1);
    Nph(i) = sum(sum(dTemp-bg(i)));
    x0(i) = sum(sum(XImage.*dTemp))/Nph(i);
    y0(i) = sum(sum(YImage.*dTemp))/Nph(i);
    z0(i) = paraSim.zemit/2;%
end

allTheta = zeros(numAberrations+5,Nz);
allTheta(numAberrations+1,:)=x0';
allTheta(numAberrations+2,:)=y0';
allTheta(numAberrations+3,:)=z0';
allTheta(numAberrations+4,:)=Nph';
allTheta(numAberrations+5,:)=bg';
allTheta(1:numAberrations,:) = repmat(paraFit.aberrations(:,3),[1 Nz]);

map = zeros(numparams,3);
n=1;
for i = 1:numAberrations+5
    if shared(i)==1
        map(n,1)= 1;
        map(n,2)=i;
        map(n,3)=0;
        n = n+1;
    elseif shared(i)==0
        for j = 1:Nz
            map(n,1)=0;
            map(n,2)=i;
            map(n,3)=j;
            n = n+1;
        end
    end
end


for i = 1:numparams
    if map(i,1)==1
        thetainit(i)= mean(allTheta(map(i,2),:));
    elseif map(i,1)==0
         thetainit(i) = allTheta(map(i,2),map(i,3));
    end
end

% we assume that parameters for zernike coefficiens are always linked
zernikecoefsmax = 0.25*paraFit.lambda*ones(numAberrations,1);
paraFit.maxJump = [zernikecoefsmax',paraFit.pixelSizeX*ones(1,max(Nz*double(shared(numAberrations+1)==0),1)),paraFit.pixelSizeY*ones(1,max(Nz*double(shared(numAberrations+2)==0),1)),500*ones(1,max(Nz*double(shared(numAberrations+3)==0),1)),2*max(Nph(:)).*ones(1,max(Nz*double(shared(numAberrations+4)==0),1)),100*ones(1,max(Nz*double(shared(numAberrations+5)==0),1))];

%% fitting simulated bead stack
paraFit.numparams = numparams;
paraFit.numAberrations = numAberrations;
paraFit.zemitStack = zeros(size(PSF_stack,3),1); % move emitter

zmax=(paraFit.sizeZ-1)*z_step/2;
paraFit.objStageStack=zmax:-z_step:-zmax;
paraFit.ztype = 'stage';
paraFit.map = map;
paraFit.Nitermax = 75;
paraFitCell= struct2cell(paraFit);

tempGauss1 = 0;


% GPU fitting
if gpuDeviceCount>0
    [P,model,err] = MLE_FitAbberation_Final_GPU_float(PSF_stack,thetainit,paraFitCell,shared,0.1,tempGauss1);
else
    % CPU fitting
    [P,model,err] = MLE_FitAbberation_Final_CPU(PSF_stack,thetainit,paraFitCell,shared,0.1,tempGauss1);
end
    
%% plot fitted pupil function
PupilSize = 1.0;
Npupil = paraFit.Npupil;
DxyPupil = 2*PupilSize/Npupil;
XYPupil = -PupilSize+DxyPupil/2:DxyPupil:PupilSize;
[YPupil,XPupil] = meshgrid(XYPupil,XYPupil);
ApertureMask = double((XPupil.^2+YPupil.^2)<1.0);

Waberration0 = zeros(size(XPupil));
orders = paraFit.aberrations(:,1:2);
zernikecoefs = P(1:21);
normfac = sqrt(2*(orders(:,1)+1)./(1+double(orders(:,2)==0)));
zernikecoefs = normfac.*zernikecoefs;
allzernikes = get_zernikefunctions(orders,XPupil,YPupil);
for j = 1:numel(zernikecoefs)
  Waberration0 = Waberration0+zernikecoefs(j)*squeeze(allzernikes(j,:,:));  
end
Waberration0 = Waberration0.*ApertureMask;

axpupil = axes(figure);
imagesc(axpupil,Waberration0)
axis(axpupil,'equal')
axis(axpupil,'tight')
ylabel('Y(pixel)') ,xlabel('X(pixel)')
c_title=colorbar;
title(c_title,'nm')

%% plot fitted zernike polynomial coefficients
axMode =axes(figure);
bar(axMode,[paraSim.aberrations(:,3),P(1:21)]);
legend(axMode,{'Simulated','Fitting Result'})
for k=size(orders):-1:1
    axn{k}=[num2str(orders(k,1)) ',' num2str(orders(k,2))];
end
axMode.XTick=1:length(axn);
axMode.XTickLabel=axn;
xtickangle(45)
ylabel('zernike rms value(nm)') ,xlabel('zernike mode')
disp('...Successfully finished!');
```
