## Supplementary Software for "Field dependent deep learning enables high-throughput whole-cell 3D super-resolution imaging": cztfunc.m

```
function dataout = cztfunc(datain,A,B,D)
% This function evaluates the FT via the czt-algorithm
% arguments: datain = input data, dimensions K x N
%            A,B,D = auxiliary vectors computed in prechirpz, must have
%            lengths N, M, and L=N+M-1
% function value: dataout = output data, dimensions K x M
%
% Sjoerd Stallinga, TU Delft

% (C) Copyright 2018
% All rights reserved
% Department of Imaging Physics
% Faculty of Applied Sciences
% Delft University of Technology
% Delft, The Netherlands   

N = length(A);
M = length(B);
L = length(D);
K = size(datain,1);
Amt = repmat(A,K,1);
Bmt = repmat(B,K,1);
Dmt = repmat(D,K,1);

cztin =  zeros(K,L);
cztin(:,1:N)= Amt.*datain;
temp = Dmt.*fft(cztin,[],2);
cztout = ifft(temp,[],2);
dataout = Bmt.*cztout(:,1:M);
  
end
```
