## Supplementary Software for "Field dependent deep learning enables high-throughput whole-cell 3D super-resolution imaging": get_zernikefunctions.m

```
function allzernikes = get_zernikefunctions(orders,x,y)
% This function computes the Zernike basis aberration functions for the
% Zernike orders given in the input argument orders.
%
% Sjoerd Stallinga, TU Delft

% (C) Copyright 2018
% All rights reserved
% Department of Imaging Physics
% Faculty of Applied Sciences
% Delft University of Technology
% Delft, The Netherlands   

zersize = size(orders);
Nzer = zersize(1);
radormax = max(orders(:,1));
azormax = max(abs(orders(:,2)));
[Nx,Ny] = size(x);

% Evaluation of the radial Zernike polynomials using the recursion relation for
% the Jacobi polynomials.
zerpol = zeros(radormax+1,azormax+1,Nx,Ny);
rhosq = x.^2+y.^2;
rho = sqrt(rhosq);
zerpol(1,1,:,:) = ones(size(x));
for jm = 1:azormax+2
  m = jm-1;
  if (m>0)
    zerpol(jm,jm,:,:) = rho.*squeeze(zerpol(jm-1,jm-1,:,:));
  end
  zerpol(jm+2,jm,:,:) = ((m+2)*rhosq-m-1).*squeeze(zerpol(jm,jm,:,:));
  for p = 2:radormax-m+2
     n = m+2*p;
     jn = n+1;
     zerpol(jn,jm,:,:) = (2*(n-1)*(n*(n-2)*(2*rhosq-1)-m^2).*squeeze(zerpol(jn-2,jm,:,:))-...
         n*(n+m-2)*(n-m-2)*squeeze(zerpol(jn-4,jm,:,:)))/((n-2)*(n+m)*(n-m));
  end
end

% Computation of the Zernike aberration functions from the radial Zernike
% polynomials and the azimuthal factors
phi = atan2(y,x);
allzernikes = zeros(Nzer,Nx,Ny);
for j =1:Nzer
  n = orders(j,1);
  m = orders(j,2);
  if m>=0
    allzernikes(j,:,:) = squeeze(zerpol(n+1,m+1,:,:)).*cos(m*phi);
  else
    allzernikes(j,:,:) = squeeze(zerpol(n+1,-m+1,:,:)).*sin(-m*phi);
  end
end

end
```
