## Supplementary Software for "Field dependent deep learning enables high-throughput whole-cell 3D super-resolution imaging": kernel_cholesky.m

```
function [L,U,errFlag] = kernel_cholesky(A,n)
L = zeros(n,n);
U = zeros(n,n);
%A = double(A);
errFlag = 0;
for i = 0:n-1
    for j = 0:i
        s = 0;
        for k = 0:j-1
            s = s + U(i*n+k+1)*U(j*n+k+1);
        end
        
        if i==j
            if A(i*n+i+1)-s>=0
                U(i*n+j+1)=sqrt(A(i*n+i+1)-s);
                L(j*n+i+1)=U(i*n+j+1);
            else
                errFlag = 1;
                break
            end
            
        else
            U(i*n+j+1) = 1/U(j*n+j+1)*(A(i*n+j+1)-s);
            L(j*n+i+1)=U(i*n+j+1);
            
        end
        
    end
end
end
```
