## Supplementary Software for "Field dependent deep learning enables high-throughput whole-cell 3D super-resolution imaging": kernel_DerivativeAberration_vectorPSF.m

```
function [dudt,model] =  kernel_DerivativeAberration_vectorPSF(theta,parameters,map)

% map[shared parameterID channelID]

lambda = parameters.lambda;
zemit0 = parameters.zemit0;
objStage0= parameters.objStage0; 
zemitStack = parameters.zemitStack;
objStageStack = parameters.objStageStack;
sizeX = parameters.sizeX;
sizeY = parameters.sizeY; 
sizeZ = parameters.sizeZ;
XPupil=parameters.XPupil;
PolarizationVector=parameters.PolarizationVector;
Amplitude=parameters.Amplitude;
wavevector=parameters.wavevector;
wavevectorzmed=parameters.wavevectorzmed;
normfac=parameters.normfac;
allzernikes=parameters.allzernikes;
Ax=parameters.Ax;
Bx=parameters.Bx;
Dx=parameters.Dx;
Ay=parameters.Ay;
By=parameters.By;
Dy=parameters.Dy;
normIntensity = parameters.normIntensity;

numparams = parameters.numparams;
numAberrations = parameters.numAberrations;
ztype = parameters.ztype;

parameterID = numAberrations+5;
channelID = parameters.sizeZ;

repTheta = zeros(parameterID,channelID);

for i = 1:numparams
    if map(i,1)==0
        repTheta(map(i,2),map(i,3)) = theta(i);
    elseif map(i,1)==1
        repTheta(map(i,2),1:channelID) = theta(i)*ones(1,channelID);
    end
end


% calculation aberration function
Waberration = zeros(size(XPupil));
zernikecoefs = squeeze(repTheta(1:numAberrations,1));
zernikecoefs = normfac.*zernikecoefs;
for j = 1:numel(zernikecoefs)
  Waberration = Waberration+zernikecoefs(j)*squeeze(allzernikes(j,:,:));  
end

PhaseFactor = exp(2*pi*1i*Waberration/lambda);


numders = 3+numAberrations;

FieldMatrix = cell(2,3,sizeZ);
FieldMatrixDerivatives = cell(2,3,sizeZ,numders);

xemit = repTheta(numAberrations+1,:);
yemit = repTheta(numAberrations+2,:);
zemit = repTheta(numAberrations+3,:);
if strcmp(ztype,'stage')
    objStage = repTheta(numAberrations+3,:)'+objStageStack(:)+objStage0(:);
    zemit = zemitStack(:)+zemit0(:);
elseif strcmp(ztype,'emitter')
    objStage = objStageStack(:)+objStage0(:);
    zemit = repTheta(numAberrations+3,:)'+zemitStack(:)+zemit0(:);
end


Nph = repTheta(numAberrations+4,:);
bg = repTheta(numAberrations+5,:);

% numparams = 26;
PSF = zeros(sizeX,sizeY,sizeZ);
PSFderivatives = zeros(sizeX,sizeY,sizeZ,numders);


for jz = 1:sizeZ
    % xyz induced phase contribution
    Wxyz= (-1*xemit(jz))*wavevector{1}+(-1*yemit(jz))*wavevector{2}+(zemit(jz))*wavevectorzmed;

    PositionPhaseMask = exp(1i*(Wxyz+(objStage(jz))*wavevector{3}));
  
  for itel = 1:2
    for jtel = 1:3    
      % pupil functions and FT to matrix elements
      PupilMatrix = Amplitude.*PhaseFactor.*PolarizationVector{itel,jtel};
      PupilFunction = PositionPhaseMask.*PupilMatrix;
      IntermediateImage = transpose(cztfunc(PupilFunction,Ay,By,Dy));
      FieldMatrix{itel,jtel,jz} = transpose(cztfunc(IntermediateImage,Ax,Bx,Dx));
      
       % pupil functions for xy-derivatives and FT to matrix elements
      for jder = 1:2
        PupilFunction = 1i*wavevector{jder}.*PositionPhaseMask.*PupilMatrix;
        IntermediateImage = transpose(cztfunc(PupilFunction,Ay,By,Dy));
        FieldMatrixDerivatives{itel,jtel,jz,numAberrations+jder} = transpose(cztfunc(IntermediateImage,Ax,Bx,Dx));
      end
      
      % pupil functions for z-derivative and FT to matrix elements
      if strcmp(ztype,'stage') % find stage position
          PupilFunction = 1i*wavevector{3}.*PositionPhaseMask.*PupilMatrix;
      elseif strcmp(ztype,'emitter') % find emitter position
          PupilFunction = 1i*wavevectorzmed.*PositionPhaseMask.*PupilMatrix;
      end
      IntermediateImage = transpose(cztfunc(PupilFunction,Ay,By,Dy));
      FieldMatrixDerivatives{itel,jtel,jz,numAberrations+3} = transpose(cztfunc(IntermediateImage,Ax,Bx,Dx));
      
       % pupil functions for Zernike mode-derivative and FT to matrix elements
      for jzer = 1:numAberrations
          PupilFunction = (2*pi*1i*normfac(jzer)*squeeze(allzernikes(jzer,:,:))/lambda).*PositionPhaseMask.*PupilMatrix;
          IntermediateImage = transpose(cztfunc(PupilFunction,Ay,By,Dy));
          FieldMatrixDerivatives{itel,jtel,jz,jzer} = transpose(cztfunc(IntermediateImage,Ax,Bx,Dx));
      end
      
      
    end
  end
end


for jz = 1:sizeZ
    for jtel = 1:3
        for itel = 1:2
            PSF(:,:,jz) = PSF(:,:,jz) + (1/3)*abs(FieldMatrix{itel,jtel,jz}).^2;
            for jder = 1:numders
                PSFderivatives(:,:,jz,jder) = PSFderivatives(:,:,jz,jder) +...
                    (2/3)*real(conj(FieldMatrix{itel,jtel,jz}).*FieldMatrixDerivatives{itel,jtel,jz,jder});
            end
        end
    end
end


PSF = PSF/normIntensity;
PSFderivatives = PSFderivatives/normIntensity;

model = PSF;
for i = 1:sizeZ
    model(:,:,i) = Nph(i)*PSF(:,:,i)+bg(i);
end

dudt = zeros(sizeX,sizeY,sizeZ,numAberrations+5);
for i = 1:sizeZ
 for jp = 1:2
    dudt(:,:,i,numAberrations+jp) = -1*Nph(i)*PSFderivatives(:,:,i,numAberrations+jp);
 end
 for jp = 1:numAberrations
    dudt(:,:,i,jp) = Nph(i)*PSFderivatives(:,:,i,jp);
 end
  dudt(:,:,i,numAberrations+3) = Nph(i)*PSFderivatives(:,:,i,numAberrations+3);
end


dudt(:,:,:,numAberrations+4) = PSF;
dudt(:,:,:,numAberrations+5) = ones(size(PSF));
  
% dudt = PSFderivatives;
```
