## Supplementary Software for "Field dependent deep learning enables high-throughput whole-cell 3D super-resolution imaging": kernel_luEvaluate.m

```
function x = kernel_luEvaluate(L,U,b,n)
%Ax = b -> LUx = b. Then y is defined to be Ux
x = zeros(n,1);
y = zeros(n,1);
%Forward solve Ly = b
for i = 0:n-1
    y(i+1)=b(i+1);
    for j = 0:i-1
        y(i+1) = y(i+1)-L(j*n+i+1)*y(j+1);
    end
    y(i+1) = y(i+1)/L(i*n+i+1);
end
%Backward solve Ux = y
for i = n-1:-1:0
    x(i+1)=y(i+1);
    for j = i+1:n-1
        x(i+1) = x(i+1)-U(j*n+i+1)*x(j+1);
    end
    x(i+1) = x(i+1)/U(i*n+i+1);
end
```
