## Supplementary Software for "Field dependent deep learning enables high-throughput whole-cell 3D super-resolution imaging": MLE_FitAbberation_Final.m

```
function [P,modelOut,err] = MLE_FitAbberation_Final(data,theta0,parameters,shared)
% common term for calculate PSF and PSF derivatives
NA = parameters.NA;
refmed = parameters.refmed;
refcov = parameters.refcov;
refimm = parameters.refimm;
lambda = parameters.lambda;
Npupil = parameters.Npupil;
sizeX = parameters.sizeX;
sizeY = parameters.sizeY;
pixelSizeX = parameters.pixelSizeX;
pixelSizeY = parameters.pixelSizeY;
xrange = pixelSizeX*sizeX/2;
yrange = pixelSizeY*sizeY/2;

% pupil radius (in diffraction units) and pupil coordinate sampling
PupilSize = 1.0;
DxyPupil = 2*PupilSize/Npupil;
XYPupil = -PupilSize+DxyPupil/2:DxyPupil:PupilSize;
[YPupil,XPupil] = meshgrid(XYPupil,XYPupil);

% calculation of relevant Fresnel-coefficients for the interfaces
% between the medium and the cover slip and between the cover slip
% and the immersion fluid
CosThetaMed = sqrt(1-(XPupil.^2+YPupil.^2)*NA^2/refmed^2);
CosThetaCov = sqrt(1-(XPupil.^2+YPupil.^2)*NA^2/refcov^2);
CosThetaImm = sqrt(1-(XPupil.^2+YPupil.^2)*NA^2/refimm^2);
%need check again, compared to original equation, FresnelPmedcov is
%multipiled by refmed
FresnelPmedcov = 2*refmed*CosThetaMed./(refmed*CosThetaCov+refcov*CosThetaMed);
FresnelSmedcov = 2*refmed*CosThetaMed./(refmed*CosThetaMed+refcov*CosThetaCov);
FresnelPcovimm = 2*refcov*CosThetaCov./(refcov*CosThetaImm+refimm*CosThetaCov);
FresnelScovimm = 2*refcov*CosThetaCov./(refcov*CosThetaCov+refimm*CosThetaImm);
FresnelP = FresnelPmedcov.*FresnelPcovimm;
FresnelS = FresnelSmedcov.*FresnelScovimm;

% Apoidization for sine condition
apoid = sqrt(CosThetaImm)./CosThetaMed;
% definition aperture
ApertureMask = double((XPupil.^2+YPupil.^2)<1.0);
Amplitude = ApertureMask.*apoid;


% setting of vectorial functions
Phi = atan2(YPupil,XPupil);
CosPhi = cos(Phi);
SinPhi = sin(Phi);
CosTheta = sqrt(1-(XPupil.^2+YPupil.^2)*NA^2/refmed^2);
SinTheta = sqrt(1-CosTheta.^2);

pvec{1} = FresnelP.*CosTheta.*CosPhi;
pvec{2} = FresnelP.*CosTheta.*SinPhi;
pvec{3} = -FresnelP.*SinTheta;
svec{1} = -FresnelS.*SinPhi;
svec{2} = FresnelS.*CosPhi;
svec{3} = 0;

PolarizationVector = cell(2,3);
for jtel = 1:3
  PolarizationVector{1,jtel} = CosPhi.*pvec{jtel}-SinPhi.*svec{jtel};
  PolarizationVector{2,jtel} = SinPhi.*pvec{jtel}+CosPhi.*svec{jtel};
end

wavevector = cell(1,3);
wavevector{1} = (2*pi*NA/lambda)*XPupil;
wavevector{2} = (2*pi*NA/lambda)*YPupil;
wavevector{3} = (2*pi*refimm/lambda)*CosThetaImm;
wavevectorzmed = (2*pi*refmed/lambda)*CosThetaMed; 

orders = parameters.aberrations(:,1:2);
normfac = sqrt(2*(orders(:,1)+1)./(1+double(orders(:,2)==0)));
allzernikes = get_zernikefunctions(orders,XPupil,YPupil);

ImageSizex = xrange*NA/lambda;
ImageSizey = yrange*NA/lambda;

[Ax,Bx,Dx] = prechirpz(PupilSize,ImageSizex,Npupil,sizeX);
[Ay,By,Dy] = prechirpz(PupilSize,ImageSizey,Npupil,sizeY);

parameters.XPupil = XPupil;
parameters.PolarizationVector = PolarizationVector;
parameters.Amplitude = Amplitude;
parameters.wavevector = wavevector;
parameters.wavevectorzmed = wavevectorzmed;
parameters.orders = orders;
parameters.normfac = normfac;
parameters.allzernikes = allzernikes;

parameters.Ax = Ax;
parameters.Bx = Bx;
parameters.Dx = Dx;
parameters.Ay = Ay;
parameters.By = By;
parameters.Dy = Dy;

% calculate intensity normalization function using the PSFs at focus
% position without any aberration
FieldMatrix = cell(2,3);
for itel = 1:2
  for jtel = 1:3
    PupilFunction = Amplitude.*PolarizationVector{itel,jtel};
    IntermediateImage = transpose(cztfunc(PupilFunction,Ay,By,Dy));
    FieldMatrix{itel,jtel} = transpose(cztfunc(IntermediateImage,Ax,Bx,Dx));
  end
end

intFocus = zeros(sizeX,sizeY);
for jtel = 1:3
    for itel = 1:2
        intFocus = intFocus + (1/3)*abs(FieldMatrix{itel,jtel}).^2;
    end
end

normIntensity = sum(intFocus(:));
parameters.normIntensity = normIntensity;


% prepare parameters for fit
NV = parameters.numparams;
numAberrations = parameters.numAberrations;
maxJump = parameters.maxJump;
iterations = parameters.Nitermax;
oldErr = 1e13;
tolerance=1e-6;
newLambda = 0.1;
newUpdate = 1e13*ones(NV,1);

% zernikecoefsmax = 0.25*parameters.lambda*ones(numAberrations,1);
% maxJump = [zernikecoefsmax',pixelSizeX*ones(1,max(sizeZ*double(shared(numAberrations+1)==0),1)),pixelSizeY*ones(1,max(sizeZ*double(shared(numAberrations+2)==0),1)),500*ones(1,max(sizeZ*double(shared(numAberrations+3)==0),1)),2*max(Nph(:)).*ones(1,max(sizeZ*double(shared(numAberrations+4)==0),1)),100*ones(1,max(sizeZ*double(shared(numAberrations+5)==0),1))];
newTheta = theta0;
errFlag = 0;
noChannels = size(data,3);
jacobian = single(zeros(NV,1));
hessian = single(zeros(NV,NV));

map = parameters.map;
data(data<=0) = 1e3*eps;
[newDudt,model] =  kernel_DerivativeAberration_vectorPSF(newTheta,parameters,map);
%need check negative
newErr = 2*((model-data)-data.*log(model./data));
newErr = sum(newErr(:));


t1 = single(1-data./model);

n = 1;
for i = 1:26
    temp = t1.*squeeze(newDudt(:,:,:,i));
    if shared(i) ==0
        for j=1:noChannels
            temp1 = temp(:,:,j);
            jacobian(n)=sum(temp1(:));
            n=n+1;
        end
        
    elseif shared(i)==1
        jacobian(n) = sum(temp(:));
        n = n+1;
    end
end

t2 = data./model.^2;


for m = 1:NV
    mm = map(m,2);
    mmShared = map(m,1);
    mmChannel = map (m,3);
    temp1 = squeeze(newDudt(:,:,:,mm));
    for n = m:NV
        nn = map(n,2);
        nnShared = map(n,1);
        nnChannel = map(n,3);
        temp2 = squeeze(newDudt(:,:,:,nn));
        if mmShared ==1 &&nnShared ==1
            temp = t2.*temp1.*temp2;
            hessian((m-1)*NV+n) = sum(temp(:));
            hessian((n-1)*NV+m) = hessian((m-1)*NV+n);
        elseif mmShared ==1&&nnShared==0
            temp = t2(:,:,nnChannel).*temp1(:,:,nnChannel).*temp2(:,:,nnChannel);
            hessian((m-1)*NV+n) = sum(temp(:));
            hessian((n-1)*NV+m) = hessian((m-1)*NV+n);
            
        elseif nnShared ==1&&mmShared==0
            temp = t2(:,:,mmChannel).*temp1(:,:,mmChannel).*temp2(:,:,mmChannel);
            hessian((m-1)*NV+n) = sum(temp(:));
            hessian((n-1)*NV+m) = hessian((m-1)*NV+n);
        elseif mmShared ==0&&nnShared==0
            if mmChannel == nnChannel
                temp = t2(:,:,mmChannel).*temp1(:,:,mmChannel).*temp2(:,:,mmChannel);
                hessian((m-1)*NV+n) = sum(temp(:));
                hessian((n-1)*NV+m) = hessian((m-1)*NV+n);
            end
            
        end
        
    end
end


oldLambda = newLambda;
oldTheta = newTheta;

for kk =1:iterations
    kk
    if abs((newErr-oldErr)/newErr)<tolerance
        break;
    else
        if newErr>1*oldErr
            newLambda = oldLambda;
            newTheta = oldTheta;
            newErr = oldErr;
            
            mult = (1 + newLambda*10)/(1 + newLambda);
            newLambda = 10*newLambda;
            
        elseif newErr<oldErr
            if errFlag == 0
                newLambda = 0.1*newLambda;
                mult = 1 + newLambda;
            end
        end
        
        
        for i = 0:NV-1
            hessian(i*NV+i+1) = hessian(i*NV+i+1)*mult;
        end
        
        
        [L,U,errFlag] = kernel_cholesky(hessian,NV);
        if errFlag ==0
            oldLambda = newLambda;
            oldUpdate = newUpdate;
            
            oldTheta = newTheta;
            oldErr = newErr;
            
            newUpdate = kernel_luEvaluate(L,U,jacobian,NV);

            for ll =1:NV
%                 if newUpdate(ll)/oldUpdate(ll)<-0.5
%                     maxJump(ll) = maxJump(ll)*0.5;
%                 end
                newUpdate(ll) = newUpdate(ll)/(1+abs(newUpdate(ll)/maxJump(ll)));
                newTheta(ll) = newTheta(ll)-newUpdate(ll);
            end

            
            temp = newTheta(1:numAberrations);
            temp(temp<-150) = -150;
            temp(temp>150) = 150;
            newTheta(1:numAberrations)=temp;
            
%             n=numAberrations+1;
%             for i = 1:5
%                 if shared(i)==1
%                     %                         maxJump(n)=mean(maxJump_Init(i,:));
%                     switch i
%                         case 1
%                         case 2
%                             newThetaAll(n)= max(newThetaAll(n),(sz-1)/2-sz/4.0);
%                             newThetaAll(n) = min(newThetaAll(n),(sz-1)/2+sz/4.0);
%                         case 3
%                             newThetaAll(n) = max(newThetaAll(n), 0);
%                             newThetaAll(n) = min(newThetaAll(n), spline_zsize);
%                         case 4
%                             newThetaAll(n) = max(newThetaAll(n), 1);
%                         case 5
%                             newThetaAll(n) = max(newThetaAll(n), 0.01);
%                     end
%                     
%                     for j = 1:noChannels
%                         newTheta(i,j)=newThetaAll(n)+dT(i,j);
%                     end
%                 else
%                     for j= 1:noChannels
%                         %                             newThetaAll(n+j-1)=newTheta(i,1);
%                         switch i
%                             case 1
%                             case 2
%                                 newThetaAll(n+j-1)= max(newThetaAll(n+j-1),(sz-1)/2-sz/4.0);
%                                 newThetaAll(n+j-1) = min(newThetaAll(n+j-1),(sz-1)/2+sz/4.0);
%                             case 3
%                                 newThetaAll(n+j-1) = max(newThetaAll(n+j-1), 0);
%                                 newThetaAll(n+j-1) = min(newThetaAll(n+j-1), spline_zsize);
%                             case 4
%                                 newThetaAll(n+j-1) = max(newThetaAll(n+j-1), 1);
%                             case 5
%                                 newThetaAll(n+j-1) = max(newThetaAll(n+j-1), 0.01);
%                         end
%                         newTheta(i,j)=newThetaAll(n+j-1);
%                         
%                     end
%                     n = n+j-1;
%                 end
%                 n=n+1;
%             end
            
            jacobian = single(zeros(NV,1));
            hessian = single(zeros(NV,NV));
            modelOut = model;
            tic
            [newDudt,model] =  kernel_DerivativeAberration_vectorPSF(newTheta,parameters,map);
            toc
            %need check negative
            newErr = 2*((model-data)-data.*log(model./data));
            newErr = sum(newErr(:));
            err = newErr;
            t1 = single(1-data./model);
            
            n = 1;
            for i = 1:26
                temp = t1.*squeeze(newDudt(:,:,:,i));
                if shared(i) ==0
                    for j=1:noChannels
                        temp1 = temp(:,:,j);
                        jacobian(n)=sum(temp1(:));
                        n=n+1;
                    end
                    
                elseif shared(i)==1
                    jacobian(n) = sum(temp(:));
                    n = n+1;
                end
            end
            
            t2 = data./model.^2;
            
%             map = zeros(NV,2);
%             n=1;
%             for i = 1:26
%                 if shared(i)==1
%                     map(n,1)= 1;
%                     map(n,2)=i;
%                     map(n,3)=0;
%                     n = n+1;
%                 elseif shared(i)==0
%                     for j = 1:noChannels
%                         map(n,1)=0;
%                         map(n,2)=i;
%                         map(n,3)=j;
%                         n = n+1;
%                     end
%                 end
%             end
            
            for m = 1:NV
                mm = map(m,2);
                mmShared = map(m,1);
                mmChannel = map (m,3);
                temp1 = squeeze(newDudt(:,:,:,mm));
                for n = m:NV
                    nn = map(n,2);
                    nnShared = map(n,1);
                    nnChannel = map(n,3);
                    temp2 = squeeze(newDudt(:,:,:,nn));
                    if mmShared ==1 &&nnShared ==1
                        temp = t2.*temp1.*temp2;
                        hessian((m-1)*NV+n) = sum(temp(:));
                        hessian((n-1)*NV+m) = hessian((m-1)*NV+n);
                    elseif mmShared ==1&&nnShared==0
                        temp = t2(:,:,nnChannel).*temp1(:,:,nnChannel).*temp2(:,:,nnChannel);
                        hessian((m-1)*NV+n) = sum(temp(:));
                        hessian((n-1)*NV+m) = hessian((m-1)*NV+n);
                        
                    elseif nnShared ==1&&mmShared==0
                        temp = t2(:,:,mmChannel).*temp1(:,:,mmChannel).*temp2(:,:,mmChannel);
                        hessian((m-1)*NV+n) = sum(temp(:));
                        hessian((n-1)*NV+m) = hessian((m-1)*NV+n);
                    elseif mmShared ==0&&nnShared==0
                        if mmChannel == nnChannel
                            temp = t2(:,:,mmChannel).*temp1(:,:,mmChannel).*temp2(:,:,mmChannel);
                            hessian((m-1)*NV+n) = sum(temp(:));
                            hessian((n-1)*NV+m) = hessian((m-1)*NV+n);
                        end
                        
                    end
                    
                end
            end
        else
            mult = (1 + newLambda*10)/(1 + newLambda);
            newLambda = 10*newLambda;
            disp('CHOLERRER')
        end
    end
     
end

P = newTheta;
P(end+1) = kk;
end
```
