## Supplementary Software for "Field dependent deep learning enables high-throughput whole-cell 3D super-resolution imaging": prechirpz.m

```
function [A,B,D] = prechirpz(xsize,qsize,N,M)
% This function evaluates the auxiliary vectors for the evaluation of the
% FT via the czt-algorithm
% arguments: xsize = window in real space abs(x)<xsize
%            qsize = window in Fourier space abs(q)<qsize
%            N = # sample points in real space (even)
%            M = # sample points in Fourier space (odd)
% function value: A,B,D = auxiliary vectors of lengths N, M, and L=N+M-1
%
% Sjoerd Stallinga, TU Delft

% (C) Copyright 2018
% All rights reserved
% Department of Imaging Physics
% Faculty of Applied Sciences
% Delft University of Technology
% Delft, The Netherlands   

L = N+M-1;
sigma = 2*pi*xsize*qsize/N/M;
Afac = exp(2*1i*sigma*(1-M));
Bfac = exp(2*1i*sigma*(1-N));
sqW = exp(2*1i*sigma);
W = sqW^2;
Gfac = (2*xsize/N)*exp(1i*sigma*(1-N)*(1-M));

Utmp = zeros(1,N);
A = zeros(1,N);
Utmp(1) = sqW*Afac;
A(1) = 1.0;
for i=2:N
  A(i) = Utmp(i-1)*A(i-1);
  Utmp(i) = Utmp(i-1)*W;
end
  
Utmp = zeros(1,M);
B = ones(1,M);
Utmp(1) = sqW*Bfac;
B(1) = Gfac;
for i=2:M
  B(i) = Utmp(i-1)*B(i-1);
  Utmp(i) = Utmp(i-1)*W;
end

Utmp = zeros(1,max(N,M)+1);
Vtmp = zeros(1,max(N,M)+1);
Utmp(1) = sqW;
Vtmp(1) = 1.0;
for i=2:max(N,M)+1
  Vtmp(i) = Utmp(i-1)*Vtmp(i-1);
  Utmp(i) = Utmp(i-1)*W;
end
D = ones(1,L);
for i=1:M
  D(i) = conj(Vtmp(i));
end
for i=1:N
  D(L+1-i) = conj(Vtmp(i+1));
end
  
D = fft(D);

end
```
