## Supplementary Software for "Field dependent deep learning enables high-throughput whole-cell 3D super-resolution imaging": calibrate_abr_map.m

```
%%
%
% Copyright (c) 2022 Li Lab, Southern University of Science and Technology, Shenzhen.
% 
%%
function [SXY,beadpos,parameters]=calibrate_abr_map(p)

if ~isfield(p,'xrange')
    p.xrange=[-inf inf]; p.yrange=[-inf inf]; 
end


if ~isfield(p,'smoothxy')
    p.smoothxy=0;
end

if ~isfield(p,'isglobalfit')
    p.isglobalfit=0;
end

if ~isfield(p,'filechannel')
    p.filechannel=1;
end

%get bead positions
p.status.String='Load files and segment beads';drawnow

if ~isfield(p,'tabgroup')
    f=figure('Name','Bead stacks calibration');
    p.tabgroup=uitabgroup(f);
    calibrationfigure=f;
else
    calibrationfigure=p.tabgroup.Parent;
end
%get beads from images
[beads,p]=images2beads_globalfit(p);
imageRoi=p.roi{1};

%get positions of beads
for k=length(beads):-1:1
    beadposx(k)=beads(k).pos(1);
    beadposy(k)=beads(k).pos(2);
end

%if only to take beads in a certain range, remove others
if isfield(p,'fov')&&~isempty(p.fov)
    indbad=beadposx<p.fov(1)| beadposx>p.fov(3)|beadposy<p.fov(2)|beadposy>p.fov(4);
    beads=beads(~indbad);
end

if isempty(beads)
    warndlg('Could not find and segment any bead. ROI size too large?')
    p.status.String='error: could not find and segment any bead...';
    return
end

p.midpoint=round(size(beads(1).stack.image,3)/2); %reference for beads
p.ploton=false;

f0g=p.midpoint;
for k=1:length(beads)
    beads(k).f0=f0g;
end

%get positions of beads
for k=length(beads):-1:1
    beadposxs(k)=beads(k).pos(1);
    beadposys(k)=beads(k).pos(2);
    beadfilenumber(k)=beads(k).filenumber;
end

%spatially dependent calibration
tgmain=p.tabgroup;
for X=1:length(p.xrange)-1
    for Y=1:length(p.yrange)-1
        if length(p.xrange)>2||length(p.yrange)>2
            ht=uitab(tgmain,'Title',['X' num2str(X) 'Y' num2str(Y)]);
            p.tabgroup=uitabgroup(ht);
        end
        
        indgood=beadposxs< p.xrange(X+1) & beadposxs>p.xrange(X) & beadposys<p.yrange(Y+1) & beadposys>p.yrange(Y);
        beadsh=beads(indgood);
        
        for k=1:max(beadfilenumber)
            indfile=(beadfilenumber==k)&indgood;
            p.fileax(k).NextPlot='add';
            scatter(p.fileax(k),beadposxs(indfile),beadposys(indfile),60,[1 1 1]);
            scatter(p.fileax(k),beadposxs(indfile),beadposys(indfile),50);
        end
        if isempty(beadsh)
            disp(['no beads found in part' num2str(p.xrange(X:X+1)) ', ' num2str(p.yrange(Y:Y+1))])
            continue
        end
        
        indgoodc=true(size(beadsh));
        gausscal=[];
        p.ax_z=[];

        % get beads calibration
        p.status.String='get beads calibration';drawnow
        [csplinecal,indgoods,beadpos{X,Y},~,testallrois,beadspos]=getstackcal_g(beadsh(indgoodc),p);
        
        for f=1:max(beadpos{X,Y}.filenumber(:))
            indfile=(beadpos{X,Y}.filenumber==f);
            p.fileax(f).NextPlot='add';
            plot(p.fileax(f),beadpos{X,Y}.xim(indfile),beadpos{X,Y}.yim(indfile),'m+');
        end
        
        icf=find(indgoodc);
        icfs=icf(indgoods);
        for k=1:length(csplinecal.cspline.coeff)
            cspline.coeff{k}=single(csplinecal.cspline.coeff{k});
        end
        cspline.dz=csplinecal.cspline.dz;
        cspline.z0=csplinecal.cspline.z0;
        cspline.x0=csplinecal.cspline.x0;
        cspline.global.isglobal=p.isglobalfit;
        cspline.mirror=csplinecal.cspline.mirror;

        gausscal=[];
        gauss_sx2_sy2=[];
        gauss_zfit=[];
        p.ax_sxsy=[];
            
        cspline_all=csplinecal;
        cspline_all=[];
        PSF=csplinecal.PSF;
        SXY(X,Y)=struct('cspline',cspline,'posind',[X,Y],'EMon',p.emgain,'PSF',{PSF});
        
%         SXY(X,Y)=struct('gausscal',gausscal,'cspline_all',cspline_all,'gauss_sx2_sy2',gauss_sx2_sy2,'gauss_zfit',gauss_zfit,...
%             'cspline',cspline,'Xrangeall',p.xrange+imageRoi(1),'Yrangeall',p.yrange+imageRoi(2),'Xrange',p.xrange([X X+1])+imageRoi(1),...
%             'Yrange',p.yrange([Y Y+1])+imageRoi(2),'posind',[X,Y],'EMon',p.emgain,'PSF',{PSF});        
        
        % ZERNIKE fitting    
        axzernike=axes(uitab(p.tabgroup,'Title','Zernikefit'));
        axPupil=axes(uitab(p.tabgroup,'Title','Pupil'));
        axMode=axes(uitab(p.tabgroup,'Title','ZernikeModel'));   
       
        stack=csplinecal.PSF{1}; %this would be the average... not sure if good.
        mp=ceil(size(stack,1)/2);
        rxy=floor(p.ROIxy/2);
        zborder=round(100/p.dz); %from alignment: outside is bad.
        stack=stack(mp-rxy:mp+rxy,mp-rxy:mp+rxy,zborder+1:end-zborder);

        %arbitrary
        stack=stack*1000; %random photons, before normalized to maximum pixel
        psfrescale = p.psfrescale;

        p.zernikefit.dz=p.dz;
        
        [SXY(X,Y).zernikefit,PSFZernike,aberrations_avg1,RRSE,MAPE]=zernikefitBeadstack(stack,p.zernikefit,psfrescale,axzernike,axPupil,axMode);
        coeffZ=Spline3D_interp(PSFZernike);
        axzernikef=axes(uitab(p.tabgroup,'Title','Zval'));
        p.z0=size(coeffZ,3)/2;
        posbeads=testfit_spline(testallrois,{coeffZ},0,p,{},axzernikef);
        beads_zrnikecoeff.aberrations_std=p.aberrationsstd;
        beads_zrnikecoeff.mapsmooth=p.mapsmooth;
        beads_zrnikecoeff.modelrrse=p.modelrrse;
        beads_zrnikecoeff.localroi=p.localroi;
        stack_avg = stack;
        beads_roi_size = size(csplinecal.PSF_all{1});
        beads_num = beads_roi_size(4);
        stack_all = csplinecal.PSF_all{1};
        if p.zernikefit.calculatefdmaps
            for k = 1:beads_num
                p.status.String=['calculate zernike parameters of individual PSFs: ' num2str(k) ' of ' num2str(beads_num)]; drawnow
                
%                 k
%                 beadspos{k}               
                if mod(k,100) == 0
                    gpuCount = gpuDeviceCount;
                    for i=1:gpuCount
                        gpu = gpuDevice(i)
                        gpu.AvailableMemory;
                    end
                end
                stack = stack_all(:,:,:,k);
                beadspos_single = beadspos{k};
                mp=ceil(size(stack,1)/2);
                rxy=floor(p.ROIxy/2);
                zborder=round(100/p.dz); %from alignment: outside is bad.
                stack=stack(mp-rxy:mp+rxy,mp-rxy:mp+rxy,zborder+1:end-zborder);

                zcrop = zeros(1,length(stack(1,1,:)));
                zcrop(1:floor(length(stack(1,1,:))/2))=-1;
                zcrop(ceil(length(stack(1,1,:))/2+1):end) = 1;

                stack_flter=logical(sum(sum(stack)));
                z0_shift = sum(zcrop.*stack_flter(1,:))*p.dz;
                stack = stack(:,:,stack_flter);%filter out frames with all zeros in stack
                stack_avg_single = stack_avg(:,:,logical(sum(sum(stack))));%filter out frames with all zeros in stack
                %arbitrary
                stack=stack*1000; %random photons, before normalized to maximum pixel
                p.zernikefit.dz=p.dz;
                
                [zernikecoeff_P,Waberration,PSFzernike_single,sigmaxy,RRSE,MAPE,z_pos] = zernikefitBeadstack_all(stack,stack_avg_single,p.zernikefit,psfrescale,axzernike,axPupil,beadspos_single,aberrations_avg1,z0_shift); 

                beads_zrnikecoeff.aberrations1 = aberrations_avg1;
                beads_zrnikecoeff.RRSE.RRSE1(k)= RRSE.RRSE1;
                beads_zrnikecoeff.RRSE.STD_aberrations1(k)= RRSE.STD_aberrations1;
                beads_zrnikecoeff.MAPE.MAPE1(k)= MAPE.MAPE1;
                beads_zrnikecoeff.MAPE.MAPE_aberrations1(k)= MAPE.MAPE_aberrations1;

                beads_zrnikecoeff.pos{k}= beadspos{k};
                beads_zrnikecoeff.sigmaxy{k}= sigmaxy;
                beads_zrnikecoeff.z_pos{k}=z_pos;

                beads_zrnikecoeff.coeff_P{k}= zernikecoeff_P;
                beads_zrnikecoeff.value_P{k} = zernikecoeff_P(:,3);

                beads_zrnikecoeff.Waberration{k} = Waberration;  %just for analyse
            end
            frame_roi = beads.roi;
            beads_zrnikecoeff.roi = frame_roi(3:4);
            beads_zrnikecoeff.model = zernikecoeff_P(:,1:2);
        end
    end
end

axcrlb=axes(uitab(p.tabgroup,'Title','CRLB'));
plotCRLBcsplinePSF(csplinecal.cspline,p.zernikefit,axcrlb);
parameters=myrmfield(p,{'tabgroup','status','ax_z','ax_sxsy','fileax'});
lambda = parameters.zernikefit.lambda;
aber_map=zeros(imageRoi(3),imageRoi(4),21);
if p.zernikefit.calculatefdmaps
    axDist=axes(uitab(p.tabgroup,'Title','BeadsDistribution'));
    axMap=axes(uitab(p.tabgroup,'Title','ZernikeMap'));
    maps_gauss = zernikecoeffMap(p,beads_zrnikecoeff,axMap,axDist); 
    p.status.String='save calibration';drawnow 
    for i = 1:21
        aber_map(:,:,i) = maps_gauss{i}/lambda;
    end
else
    for i = 1:21
        aber_map(:,:,i) = aberrations_avg1(i,3)/lambda;
    end    
end

aber_map(:,:,22)= psfrescale;
aber_map(:,:,23)= psfrescale;
save(p.outputfile,'aber_map','-v7'); 

cal_par =strrep(p.outputfile,'aber_map.mat','3dcal.mat');

save(cal_par,'SXY','parameters','beads_zrnikecoeff','-v7');
filefig=strrep(cal_par,'.mat','.fig');
savefig(calibrationfigure,filefig,'compact');
p.status.String='Calibration done';drawnow
```
