## Supplementary Software for "Field dependent deep learning enables high-throughput whole-cell 3D super-resolution imaging": calibrate_abr_map_GUI.m

```
%%
%
% Copyright (c) 2022 Li Lab, Southern University of Science and Technology, Shenzhen.
% 
%%
classdef calibrate_abr_map_GUI<handle
    properties
        guihandles
        zernikepar_flag
        zernikeparameters
        fitFDApos
    end
    methods
        function obj=calibrate_abr_map_GUI(varargin)  
            pathprivate=[fileparts(pwd) filesep 'fitFDaberration' filesep 'VectorPSF_Fit'];
            if exist(pathprivate,'dir')
                addpath(pathprivate)
            end
            if exist('shared','file')
                dirlist=genpath('shared');
                addpath(dirlist)
            end
            
            obj.fitFDApos.fitFDApath=[fileparts(pwd) filesep 'fitFDaberration'];  
            figureheight=550;
            
            handle=figure('Name','Field dependant aberration map calibration','MenuBar','none','ToolBar','none');
            set(handle,'NumberTitle','off')
            initPosition = handle.Position;
            handle.Position=[initPosition(1), initPosition(2)- figureheight+initPosition(4),900, figureheight];
            top=handle.Position(4)-10;
            vsep=25;
            
            if ispc
                fontsize=10;
                fieldheight=vsep;
            else 
                fontsize=14;
                fieldheight=vsep;
            end
            xpos1=10;
            xw=100;
            hatitle='left';
            obj.guihandles.handle=handle;
            obj.guihandles.title=uicontrol('style','text','String','Calibrate field dependant aberration from bead stacks. 2022 Li lab, SUSTech','Position',[xpos1,top-vsep,xw*4.5,fieldheight],'FontSize',fontsize-2,'HorizontalAlignment',hatitle,'FontWeight','bold');
         
            obj.guihandles.filelist=uicontrol('style','listbox','String','','Position',[xpos1,top-16*vsep,xw*4,vsep*15],'FontSize',fontsize);
            obj.guihandles.filelist.TooltipString='List of image files used for calibration. To change this list, use select camera files';
            
            obj.guihandles.selectfiles=uicontrol('style','pushbutton','String','Select camera files','Position',[xpos1,top-17*vsep,xw*4,vsep],'FontSize',fontsize,'Callback',@obj.selectfiles_callback);
            obj.guihandles.selectfiles.TooltipString='Select image files with bead stacks. You can select several files from different locations with the file select dialog box opend';

            obj.guihandles.outputfile=uicontrol('style','edit','String','aber_map.mat','Position',[xpos1,top-19*vsep,xw*4,fieldheight],'FontSize',fontsize);
            obj.guihandles.outputfile.TooltipString='Name of the output file';
            
            
            obj.guihandles.selectoutputfile=uicontrol('style','pushbutton','String','Select output file','Position',[xpos1,top-20*vsep,xw*4,fieldheight],'FontSize',fontsize,'Callback',@obj.selectoutputfile_callback);
            obj.guihandles.selectoutputfile.TooltipString='Select file name for output calibration file.';    
          
            %General parameters
            ha='right';
            obj.guihandles.csplinet=uicontrol('style','text','String','General parameters: ','Position',[xpos1+4.5*xw,top-2*vsep,xw*4,fieldheight],'FontSize',fontsize,'HorizontalAlignment',hatitle,'FontWeight','bold');
            obj.guihandles.modalityt=uicontrol('style','text','String','3D modality ','Position',[xpos1+4.5*xw,top-3*vsep,xw*2,fieldheight],'FontSize',fontsize,'HorizontalAlignment',ha);
            obj.guihandles.modality=uicontrol('style','popupmenu','String',{'DMO Tetrapod','Astigmatic'},'Value',1,'Position',[xpos1+6.5*xw,top-3*vsep,xw*1.3,fieldheight],'FontSize',fontsize,'Callback',@obj.modality_callback);
            obj.guihandles.modality.TooltipString='Select the kind of PSF. Astigmatic, Tatrapod';
            obj.guihandles.modalityt.TooltipString=obj.guihandles.modality.TooltipString;
            
           
            obj.guihandles.dzt=uicontrol('style','text','String','Distance between frames (nm)','Position',[xpos1+4.5*xw,top-4*vsep,xw*2,fieldheight],'FontSize',fontsize,'HorizontalAlignment',ha);
            obj.guihandles.dz=uicontrol('style','edit','String','100','Position',[xpos1+6.5*xw,top-4*vsep,xw*0.5,fieldheight],'FontSize',fontsize);
            obj.guihandles.dz.TooltipString=sprintf('Distance in nm between frames. By convention, these are objective positions (not corrected for refractive index mismatch).');
            obj.guihandles.dzt.TooltipString=obj.guihandles.dz.TooltipString;

            obj.guihandles.filtert=uicontrol('style','text','String','Filter size for peak finding','Position',[xpos1+4.5*xw,top-5*vsep,xw*2,fieldheight],'FontSize',fontsize,'HorizontalAlignment',ha);
            obj.guihandles.filter=uicontrol('style','edit','String','2','Position',[xpos1+6.5*xw,top-5*vsep,xw*0.5,fieldheight],'FontSize',fontsize);
            obj.guihandles.filter.TooltipString=sprintf('Gaussian filter for peak finding (sigma in pixels). For split PSFs (e.g. double-helix) choose larger value to segment centroid positions of the beads, not individual lobes.');
            obj.guihandles.filtert.TooltipString=obj.guihandles.filter.TooltipString;
 
            obj.guihandles.cutoffrelt=uicontrol('style','text','String','Relative cutoff','Position',[xpos1+7*xw,top-5*vsep,xw*1.,fieldheight],'FontSize',fontsize,'HorizontalAlignment',ha);
            obj.guihandles.cutoffrel=uicontrol('style','edit','String','1','Position',[xpos1+8*xw,top-5*vsep,xw*0.5,fieldheight],'FontSize',fontsize);
            obj.guihandles.cutoffrel.TooltipString=sprintf('Sometimes, the automatically determined cutoff does not work. If beads are not found, increase this value, if too many beads are found, decrease it.');
            obj.guihandles.cutoffrelt.TooltipString=obj.guihandles.cutoffrel.TooltipString;
            
            obj.guihandles.mindistancet=uicontrol('style','text','String','Minimum distance (pixels)','Position',[xpos1+4.5*xw,top-6*vsep,xw*2,fieldheight],'FontSize',fontsize,'HorizontalAlignment',ha);
            obj.guihandles.mindistance=uicontrol('style','edit','String','25','Position',[xpos1+6.5*xw,top-6*vsep,xw*.5,fieldheight],'FontSize',fontsize);
            obj.guihandles.mindistance.TooltipString=sprintf('Minimum distance between beads (in pixels). If beads are closer together, they are removed and not used for averaging. Helps eliminating background contaminations from close by beads');
            obj.guihandles.mindistancet.TooltipString=obj.guihandles.mindistance.TooltipString;           
     
            obj.guihandles.roisizet=uicontrol('style','text','String','ROI size: X,Y (pixels)','Position',[xpos1+4.5*xw,top-7*vsep,xw*2,fieldheight],'FontSize',fontsize,'HorizontalAlignment',ha);
            obj.guihandles.ROIxy=uicontrol('style','edit','String','27','Position',[xpos1+6.5*xw,top-7*vsep,xw*.5,fieldheight],'FontSize',fontsize);
            obj.guihandles.ROIxy.TooltipString=sprintf('The pixels size of the PSF ROI selected for the fitting. Usually keep 27 for astigmatism PSF and 51 for DMO Tetrapod PSF and the maximum range not to exceed 61 pixels');
            
            obj.guihandles.psfrescalet=uicontrol('style','text','String','PSF Rescale','Position',[xpos1+7*xw,top-7*vsep,xw*1,fieldheight],'FontSize',fontsize,'HorizontalAlignment',ha);
            obj.guihandles.psfrescale=uicontrol('style','edit','String','0.5','Position',[xpos1+8*xw,top-7*vsep,xw*0.5,fieldheight],'FontSize',fontsize);
            obj.guihandles.psfrescale.TooltipString=sprintf('Smoothing for the PSF model.The parameters can be adjusted according to the psf shape.\n Increasing the value of the parameter will make the shape of the PSF smoother, 0.5 to 0.8 is recommended');

            obj.guihandles.setframest=uicontrol('style','text','String','Frames steps','Position',[xpos1+4.5*xw,top-8*vsep,xw*2,fieldheight],'FontSize',fontsize,'HorizontalAlignment',ha);
            obj.guihandles.setframes=uicontrol('style','checkbox','String','set frames','Position',[xpos1+6.5*xw,top-8*vsep,xw*1,fieldheight],'FontSize',fontsize,'HorizontalAlignment',ha,'Callback',@obj.setframes_callback);
            obj.guihandles.framerange=uicontrol('style','edit','String','1:5:201','Position',[xpos1+7.5*xw,top-8*vsep,xw,fieldheight],'FontSize',fontsize,'Visible','off');
            obj.guihandles.setframes.TooltipString=sprintf('Set the frame interval of z-stacks to improve the fitting speed, but may reduce the fitting accuracy due to fewer frames.\n At the same time, you need to change the "Distance between frames" parameter to ensure that the distance between the actual fitting two frames is correct');

            %fit zernike coefficients parameters
            obj.guihandles.zernikefit=uicontrol('style','text','String','Fit zernike parameters: ','Position',[xpos1+4.5*xw,top-9*vsep,xw*2,fieldheight],'FontSize',fontsize,'HorizontalAlignment',hatitle,'FontWeight','bold');  
            obj.guihandles.NAt=uicontrol('style','text','String','Numerical aperture of obj/ NA ','Position',[xpos1+4.5*xw,top-10*vsep,xw*2,fieldheight],'FontSize',fontsize,'HorizontalAlignment',ha);
            obj.guihandles.NA=uicontrol('style','edit','String','1.35','Position',[xpos1+6.5*xw,top-10*vsep,xw*0.5,fieldheight],'FontSize',fontsize);
            obj.guihandles.NA.TooltipString=sprintf('Numerical aperture of objective. ');
            obj.guihandles.NAt.TooltipString=obj.guihandles.NA.TooltipString;
            
            obj.guihandles.refmedt=uicontrol('style','text','String','Refractive index of medium ','Position',[xpos1+4.5*xw,top-11*vsep,xw*2,fieldheight],'FontSize',fontsize,'HorizontalAlignment',ha);
            obj.guihandles.refmed=uicontrol('style','edit','String','1.406','Position',[xpos1+6.5*xw,top-11*vsep,xw*0.5,fieldheight],'FontSize',fontsize);
            obj.guihandles.refmed.TooltipString=sprintf('Refractive index of sample medium. ');
            obj.guihandles.refmedt.TooltipString=obj.guihandles.refmed.TooltipString;

            obj.guihandles.refcovt=uicontrol('style','text','String','Refractive index of converslip ','Position',[xpos1+4.5*xw,top-12*vsep,xw*2,fieldheight],'FontSize',fontsize,'HorizontalAlignment',ha);
            obj.guihandles.refcov=uicontrol('style','edit','String','1.525','Position',[xpos1+6.5*xw,top-12*vsep,xw*0.5,fieldheight],'FontSize',fontsize);
            obj.guihandles.refcov.TooltipString=sprintf('Refractive index of converslip. ');
            obj.guihandles.refcovt.TooltipString=obj.guihandles.refcov.TooltipString;
            
            obj.guihandles.refimmt=uicontrol('style','text','String','Refractive index of immersion oil ','Position',[xpos1+4.5*xw,top-13*vsep,xw*2,fieldheight],'FontSize',fontsize,'HorizontalAlignment',ha);
            obj.guihandles.refimm=uicontrol('style','edit','String','1.406','Position',[xpos1+6.5*xw,top-13*vsep,xw*0.5,fieldheight],'FontSize',fontsize);
            obj.guihandles.refimm.TooltipString=sprintf('Refractive index of immersion oil. ');
            obj.guihandles.refimmt.TooltipString=obj.guihandles.refimm.TooltipString;
            
            obj.guihandles.lambdat=uicontrol('style','text','String','wavelength of emission ','Position',[xpos1+4.5*xw,top-14*vsep,xw*2,fieldheight],'FontSize',fontsize,'HorizontalAlignment',ha);
            obj.guihandles.lambda=uicontrol('style','edit','String','680','Position',[xpos1+6.5*xw,top-14*vsep,xw*0.5,fieldheight],'FontSize',fontsize);
            obj.guihandles.lambda.TooltipString=sprintf('Wavelength of emission.  ');
            obj.guihandles.lambdat.TooltipString=obj.guihandles.lambda.TooltipString;
            
            obj.guihandles.pixelSizet=uicontrol('style','text','String','Pixel sizeX','Position',[xpos1+7*xw,top-13*vsep,xw*1.,fieldheight],'FontSize',fontsize,'HorizontalAlignment',ha);
            obj.guihandles.pixelSizeX=uicontrol('style','edit','String','110','Position',[xpos1+8*xw,top-13*vsep,xw*0.5,fieldheight],'FontSize',fontsize);
            obj.guihandles.pixelSizeX.TooltipString=sprintf('pixel size of the image. ');
            obj.guihandles.pixelSizeXt.TooltipString=obj.guihandles.pixelSizeX.TooltipString;
         
            obj.guihandles.pixelSizet=uicontrol('style','text','String','Pixel sizeY','Position',[xpos1+7*xw,top-14*vsep,xw*1.,fieldheight],'FontSize',fontsize,'HorizontalAlignment',ha);
            obj.guihandles.pixelSizeY=uicontrol('style','edit','String','110','Position',[xpos1+8*xw,top-14*vsep,xw*0.5,fieldheight],'FontSize',fontsize);
            obj.guihandles.pixelSizeY.TooltipString=sprintf(' pixel size of the image. ');
            obj.guihandles.pixelSizeYt.TooltipString=obj.guihandles.pixelSizeY.TooltipString;
            
            %Calibrate maps parameters
            obj.guihandles.mapcalibrate=uicontrol('style','text','String','Calibrate maps parameters: ','Position',[xpos1+4.5*xw,top-15*vsep,xw*2,fieldheight],'FontSize',fontsize,'HorizontalAlignment',hatitle,'FontWeight','bold'); 
            obj.guihandles.fdmapscal=uicontrol('style','checkbox','String','Generate field dependant aberration maps','Position',[xpos1+4.5*xw,top-16*vsep,xw*2.6,fieldheight],'FontSize',fontsize,'HorizontalAlignment',hatitle,'Callback',@obj.FDmaps_callback,'Value',1);   
            obj.guihandles.fdmapscal.TooltipString=sprintf('Select whether to generate field dependant aberration maps. When not select, it is the map of the average aberration coefficient generated, \n which can be applied to the case where the image area is relatively small and the field dependant aberration are not significant. ');
            
            obj.guihandles.stdofaberrationst=uicontrol('style','text','String',' Filter factor for outlier beads','Position',[xpos1+4.5*xw,top-17*vsep,xw*2,fieldheight],'FontSize',fontsize,'HorizontalAlignment',ha,'Visible','on');
            obj.guihandles.stdofaberrations=uicontrol('style','edit','String','2.5','Position',[xpos1+6.5*xw,top-17*vsep,xw*.5,fieldheight],'FontSize',fontsize,'Visible','on');
            obj.guihandles.stdofaberrations.TooltipString=sprintf('Exclude some outlier beads. The default parameter of 2.5 means to remove the beads whose Zernike coefficient standard deviation exceeds this value. Usually keep the default value.');
            
            obj.guihandles.modelrrset=uicontrol('style','text','String','Model rrse','Position',[xpos1+7*xw,top-17*vsep,xw*1,fieldheight],'FontSize',fontsize,'HorizontalAlignment',ha,'Visible','on');
            obj.guihandles.modelrrse=uicontrol('style','edit','String','0.2','Position',[xpos1+8*xw,top-17*vsep,xw*.5,fieldheight],'FontSize',fontsize,'Visible','on');
            obj.guihandles.modelrrse.TooltipString=sprintf('Discard the bad fitting precision beads. The default parameter means to remove the beads whose relative root square error (RRSE) between the raw data and fitted model was more than 0.2. Usually keep the default value.');
            
            obj.guihandles.mapsmootht=uicontrol('style','text','String','Map smooth','Position',[xpos1+4.5*xw,top-18*vsep,xw*2,fieldheight],'FontSize',fontsize,'HorizontalAlignment',ha,'Visible','on');
            obj.guihandles.mapsmooth=uicontrol('style','edit','String','200','Position',[xpos1+6.5*xw,top-18*vsep,xw*.5,fieldheight],'FontSize',fontsize,'Visible','on');
            obj.guihandles.mapsmooth.TooltipString=sprintf('Gaussian smoothing for the aberration maps. Usually keep the default value.');
           
            obj.guihandles.localroit=uicontrol('style','text','String','Local roi','Position',[xpos1+7*xw,top-18*vsep,xw*1,fieldheight],'FontSize',fontsize,'HorizontalAlignment',ha,'Visible','on');
            obj.guihandles.localroi=uicontrol('style','edit','String','256','Position',[xpos1+8*xw,top-18*vsep,xw*.5,fieldheight],'FontSize',fontsize,'Visible','on');
            obj.guihandles.localroi.TooltipString=sprintf('The default parameter means to exclude some outlier beads that 256��256 pixels around them. ');
            
            obj.guihandles.run=uicontrol('style','pushbutton','String','Calculate field dependant aberration maps','Position',[xpos1+4.5*xw,top-20*vsep,xw*4,fieldheight],'FontSize',fontsize,'Callback',@obj.run_callback,'FontWeight','bold');
                     
            obj.guihandles.status=uicontrol('style','text','String','Status','Position',[xpos1,.5*vsep,xw*4,fieldheight],'FontSize',fontsize,'HorizontalAlignment','left');
            
            modality_callback(obj,0,0)

        end

        function selectfiles_callback(obj,a,b)
            sf=selectManyFiles;
            sf.guihandles.filelist.String=(obj.guihandles.filelist.String);
            waitfor(sf.handle);
            obj.guihandles.filelist.String=sf.filelist;
            obj.guihandles.filelist.Value=1;

                ind=strfind(sf.filelist{1},';');
                if ~isempty(ind)
                    fileh=sf.filelist{1}(1:ind-1);
                else
                    fileh=sf.filelist{1};
                end
                [path,file]=fileparts(fileh);
                if length(sf.filelist)>1
                    fileh2=sf.filelist{2};
                    path2=fileparts(fileh2);
                    if ~strcmp(path,path2) %not the same: look two hierarchies down
                        if strcmp(fileparts(path),fileparts(path2))
                            path=fileparts(path);
                        elseif strcmp(fileparts(fileparts(path)),fileparts(fileparts(path2)))
                            path=fileparts(fileparts(path));
                        end
                    end
                end
                obj.guihandles.outputfile.String=[path filesep file '_aber_map.mat'];

        end
        function selectoutputfile_callback(obj,a,b)
            postfix=obj.guihandles.outputfile.String;
            if isempty(postfix)
                postfix='_aber_map.mat';
            end
            [f,p]=uiputfile(postfix);
            if f
            obj.guihandles.outputfile.String=[p,f];
            end
        end

        function modality_callback(obj,a,b)
 
            switch obj.guihandles.modality.String{obj.guihandles.modality.Value}
                case 'Astigmatic'
                    obj.guihandles.filter.String=2;
                    obj.guihandles.ROIxy.String=27;
                    obj.guihandles.mindistance.String = 25;
                case 'DMO Tetrapod'
                    obj.guihandles.filter.String=5;
                    obj.guihandles.ROIxy.String=51;
                    obj.guihandles.mindistance.String = 50;
            end

        end

        function setframes_callback(obj,a,b)
            if a.Value
                v='on';
            else
                v='off';
            end
            obj.guihandles.framerange.Visible=v;     
        end
        
        function FDmaps_callback(obj,a,b)
            if a.Value
                vis='on';
            else
                vis='off';
            end
            obj.guihandles.stdofaberrations.Visible=vis;
            obj.guihandles.stdofaberrationst.Visible=vis;
            obj.guihandles.modelrrset.Visible=vis;
            obj.guihandles.modelrrse.Visible=vis;
            obj.guihandles.mapsmootht.Visible=vis;
            obj.guihandles.mapsmooth.Visible=vis;
            obj.guihandles.localroit.Visible=vis;
            obj.guihandles.localroi .Visible=vis;     
        end

        function out=run_callback(obj,a,b)
            paraFit.NA = str2double(obj.guihandles.NA.String);
            paraFit.refmed =str2double(obj.guihandles.refmed.String);
            paraFit.refcov = str2double(obj.guihandles.refcov.String);
            paraFit.refimm =str2double(obj.guihandles.refimm.String);
            paraFit.lambda =str2double(obj.guihandles.lambda.String);
            paraFit. pixelSizeX =str2double(obj.guihandles.pixelSizeX.String);
            paraFit. pixelSizeY =str2double(obj.guihandles.pixelSizeY.String);
            paraFit.Npupil = 64;
            paraFit.sharedIB=false;
            paraFit.iterations=75;
            paraFit.zemit0 = 50;
            paraFit.objStage0 = 0; 
            obj.zernikeparameters=paraFit;
            
            p.filelist=obj.guihandles.filelist.String;
            p.outputfile=obj.guihandles.outputfile.String;
            p.dz=str2double(obj.guihandles.dz.String);
            p.modality=obj.guihandles.modality.String{obj.guihandles.modality.Value};
            p.zcorr='cross-correlation';
            p.ROIxy=str2double(obj.guihandles.ROIxy.String);

            p.smoothxy=0;

            p.smoothz=1;

            p.filtersize=str2double(obj.guihandles.filter.String);
            p.zcorrframes=1000/ p.dz;

            p.status=obj.guihandles.status;
            p.mindistance=str2double(obj.guihandles.mindistance.String);
            p.cutoffrel=str2double(obj.guihandles.cutoffrel.String);
            
            if isempty(p.filelist)
                warndlg('please select image files first')
                return
            end
            
            if obj.guihandles.setframes.Value
                p.framerange=str2num(obj.guihandles.framerange.String);
            end

            p.aberrationsstd = str2num(obj.guihandles.stdofaberrations.String);
            p.mapsmooth = str2num(obj.guihandles.mapsmooth.String);
            p.modelrrse = str2num(obj.guihandles.modelrrse.String);
            p.localroi=str2num(obj.guihandles.localroi.String);
            p.psfrescale = str2num(obj.guihandles.psfrescale.String);
%             p.lambda=obj.guihandles.lambda;
            p.zernikefit=obj.zernikeparameters;
            p.zernikefit.calculatefdmaps=obj.guihandles.fdmapscal.Value;
            
            calibrate_abr_map(p);
            
        end    
    end
end
```
