## Supplementary Software for "Field dependent deep learning enables high-throughput whole-cell 3D super-resolution imaging": bsarray.m

```
function b = bsarray(a,varargin)
% bsarray: BSpline array class constructor.
% usage: b = bsarray(a);
%    or: b = bsarray(a,'property',value,...);
%
% arguments:
%   a - N-dimensional array (vector, image, volume, etc.)
%   property/value pairs:
%       'degree': scalar or vector containing desired degree of BSpline for
%           each dimension. If 'degree' is supplied a scalar value, each
%           dimension is assumed to have that degree.  Default degree = 3.
%           Valid degree values are integers in the interval [0 7].
%       'elementSpacing': scalar or vector containing physical spacing of
%           each dimension of vector/image/volume. If 'elementSpacing' is
%           supplied a scalar value, each dimension is assumed to have that
%           spacing.  Default elementSpacing = 1.  Valid elementSpacing
%           values are positive (nonzero) numbers.
%       'lambda': scalar or vector containing coefficient for amount of
%           smoothing done in each dimension. If 'lambda' is supplied a
%           scalar value, each dimension is assumed to have that lambda.
%           Default lambda = 0 (no smoothing).  Valid lambda values are
%           nonnegative numbers.  Note: lambda values are considired to
%           apply to arrays with unit element spacing.  If elementSpacing
%           values are non-unit, appropriate scaling of lambda will be done
%           internally.
%   
%   b - BSpline array object containing BSpline coefficients of a
%

% author: Nathan D. Cahill
%
% date: 18 April 2008

switch nargin
    case 0  % no inputs supplied; create empty bsarray object
        b.coeffs = [];
        b.tensorOrder = 0;
        b.dataSize = [];
        b.coeffsSize = [];
        b.degree = [];
        b.centred = [];
        b.elementSpacing = [];
        b.lambda = [];
        b = class(b,'bsarray');
    case 1 % single input
        if isa(a,'bsarray') % if bsarray, then perform copy
            b = a;
        else % if regular array, compute BSpline coefficients
            b = constructBsarray(a);
        end
%         b = class(b,'bsarray');
    otherwise % compute BSpline coefficients
        b = constructBsarray(a,varargin{:});
%         b = class(b,'bsarray');
end

%% subfunction constructBsarray
function b = constructBsarray(a,varargin)

% first input should be array
if ~isnumeric(a)
    error([mfilename,':constructBsarray:InputNotValid'],...
        'First input argument must be numeric or of class bsarray');
end

% get tensor order of data array (i.e., 1 for vector, 2 for matrix, etc.)
n = ndims(a);
if (n==2) && (numel(a)==length(a))
    n = 1;
end

% initialize values from possible property/value pairs
deg = [];
eSp = [];
lambda = [];

% get values supplied as input in property/value pairs
k = length(varargin);
if mod(k,2)==1
    error([mfilename,':constructBsarray:PropValPairsNotValid'],...
        'Property/value inputs must come in pairs');
end
for i=1:2:(k-1)
    % test input to make sure it is a character
    if ~ischar(varargin{i})
        error([mfilename,':constructBsarray:PropertyNotString'],...
            'Property input must be a string');
    end
    % grab values
    switch lower(varargin{i})
        case 'degree'
            deg = varargin{i+1};
        case 'elementspacing'
            eSp = varargin{i+1};
        case 'lambda'
            lambda = varargin{i+1};
        otherwise
            error([mfilename,':constructBsarray:InvalidProperty'],...
                'Property must be one of {''degree'', ''elementSpacing'', or ''lambda''}');
    end
end

% set/test degree
if isempty(deg)
    deg = 3;
end
if isscalar(deg)
    deg = repmat(deg,[1 n]);
end
deg = deg(:)';
if numel(deg)~=n
    error([mfilename,':parseInputs:DegreeNotValid'],...
        'Degree must be scalar or vector of same length as the number of nonsingleton dimensions of a.');
end
if any(deg<0) || any(deg>7) || any(deg~=round(deg))
    error([mfilename,':parseInputs:DegreeNotValid'],...
        'Each element of BSpline degree must be an integer in the interval [0,7]');
end

% centred flag will be set to true, so that in each dimension, the BSpline
% basis functions are centred, not shifted
cflag = repmat(true,size(deg));

% set/test elementSpacing
if isempty(eSp)
    eSp = 1;
end
if isscalar(eSp)
    eSp = repmat(eSp,[1 n]);
end
eSp = eSp(:)';
if numel(eSp)~=n
    error([mfilename,':parseInputs:ElementSpacingNotValid'],...
        'ElementSpacing must be scalar or vector of same length as the number of nonsingleton dimensions of a.');
end
if any(eSp<=0)
    error([mfilename,':parseInputs:ElementSpacingNotValid'],...
        'Each element of ElementSpacing must be a nonzero positive number.');
end

% set/test lambda
if isempty(lambda)
    lambda = 0;
end
if isscalar(lambda)
    lambda = repmat(lambda,[1 n]);
end
lambda = lambda(:)';
if numel(lambda)~=n
    error([mfilename,':parseInputs:LambdaNotValid'],...
        'Lambda must be scalar or vector of same length as the number of nonsingleton dimensions of a.');
end
if any(lambda<0)
    error([mfilename,':parseInputs:LambdaNotValid'],...
        'Each element of lambda must be a nonzero positive number.');
end

% don't allow nonzero lambda for even degrees
if any(lambda(mod(deg,2)==0))
    error([mfilename,':parseInputs:NZLambdaEvenDegree'],...
        'Nonzero lambda values not supported for even degree BSplines.');
end

% finally, construct struture of bsarray fields/values
coeffs = directFilter(a,deg,n,lambda);
b = struct('coeffs',coeffs,...
    'tensorOrder',n,...
    'dataSize',size(a),...
    'coeffsSize',size(coeffs),...
    'degree',deg,...
    'centred',cflag,...
    'elementSpacing',eSp,...
    'lambda',lambda);
```
