## Supplementary Software for "Field dependent deep learning enables high-throughput whole-cell 3D super-resolution imaging": directFilter.m

```
function CPad = directFilter(A,D,tensorOrder,lambda)
% bsarray\directFilter: compute B-spline coefficients from data array
% usage: C = directFilter(A,D,tensorOrder,lambda);
%
% arguments: 
%   A - N-dimensional array (vector, image, volume, etc.)
%   D (1xN vector) - degree of each dimension of tensor product BSpline.
%   tensorOrder (scalar) - number of nonsingleton dimensions of A
%   lambda (1xN vector) - smoothing factor for each dimension
%
%   C - array of BSpline coefficients
%

% author: Nathan D. Cahill
%
% date: 18 April 2008

% if D = 0 or 1, and lambda = 0, return A, with reflection padding
if all(D<2) && all(lambda==0)
    padNum = floor(D/2);
    idx   = cell(1,tensorOrder);
    for k = 1:tensorOrder
        M = size(A,k);
        dimNums = [1:M (M-1):-1:2];
        p = padNum(k);
        idx{k}   = dimNums(mod(-p:M+p, 2*M-2) + 1);
    end
    CPad = A(idx{:});
    return;
end

% get coefficients for BSpline filters
F = cell(tensorOrder,1);
F0 = cell(tensorOrder,1);
for i=1:tensorOrder
    % NEED TO UPDATE THIS TO SCALE LAMBDA FOR ELEMENT SPACING
    [F{i},F0{i}] = getBSplineFiltCoeffs(D(i),true,'direct',lambda(i));
end

% initialize output
C = double(A);

% choose tolerance for K0
K0Tol = eps;

switch tensorOrder
    case 1 % X is a vector - easy case
        
        CLen = numel(C);
        
        % loop through poles of direct BSpline filter
        for i = 1:length(F{1})
            
            % compute K0 for current pole
            K0 = ceil(log(K0Tol)./log(abs(F{1}(i))));
            
            % now compute vector of indices of length K0 that runs back and
            % forth through the length of the data vector (to mimic
            % reflection of the data at each end)
            indReflect = [1:CLen (CLen-1):-1:2];
            numReps = ceil(K0/(2*CLen-2));
            KVec = repmat(indReflect,[1 numReps]);
            KVec = KVec(1:K0);
                        
            % compute scaling factor for current pole
            C0 = -F{1}(i)./(1-F{1}(i)^2);
            
            % apply symmetric exponential filter
            C(:) = symExpFilt(C(:),CLen,C0,F{1}(i),K0,KVec);
        
        end
        
        % multiply by numerator of direct BSpline filter
        C = real(C)*F0{1};
        
    case 2 % X is a matrix - a little more difficult
        
        % first perform direct filtering over each column
        [CRows,CCols] = size(C);
        
        % loop through poles of direct BSpline filter
        for i = 1:length(F{1})
            
            % compute K0 for current pole
            K0 = ceil(log(K0Tol)./log(abs(F{1}(i))));
            
            % now compute vector of indices of length K0 that runs back and
            % forth through the length of the data vector (to mimic
            % reflection of the data at each end)
            indReflect = [1:CRows (CRows-1):-1:2];
            numReps = ceil(K0/(2*CRows-2));
            KVec = repmat(indReflect,[1 numReps]);
            KVec = KVec(1:K0);
                        
            % compute scaling factor for current pole
            C0 = -F{1}(i)./(1-F{1}(i)^2);
            
            % apply symmetric exponential filter for each column
            for k = 1:CCols
                C(:,k) = symExpFilt(C(:,k),CRows,C0,F{1}(i),K0,KVec);
            end
        end
        
        % multiply by numerator of direct BSpline filter
        C = real(C)*F0{1};
        
        % now perform direct filtering across each row
        % loop through poles of direct BSpline filter
        for i = 1:length(F{2})
            
            % compute K0 for current pole
            K0 = ceil(log(K0Tol)./log(abs(F{2}(i))));
            
            % now compute vector of indices of length K0 that runs back and
            % forth through the length of the data vector (to mimic
            % reflection of the data at each end)
            indReflect = [1:CCols (CCols-1):-1:2];
            numReps = ceil(K0/(2*CCols-2));
            KVec = repmat(indReflect,[1 numReps]);
            KVec = KVec(1:K0);
                        
            % compute scaling factor for current pole
            C0 = -F{2}(i)./(1-F{2}(i)^2);
            
            % apply symmetric exponential filter for each column
            for k = 1:CRows
                C(k,:) = symExpFilt(C(k,:),CCols,C0,F{2}(i),K0,KVec);
            end
        end
        
        % multiply by numerator of direct BSpline filter
        C = real(C)*F0{2};
        
    case 3 % X is a volume - a bit more difficult
    
        % first perform direct filtering over each column
        [CRows,CCols,CSlices] = size(C);
        
        % loop through poles of direct BSpline filter
        for i = 1:length(F{1})
            
            % compute K0 for current pole
            K0 = ceil(log(K0Tol)./log(abs(F{1}(i))));
            
            % now compute vector of indices of length K0 that runs back and
            % forth through the length of the data vector (to mimic
            % reflection of the data at each end)
            indReflect = [1:CRows (CRows-1):-1:2];
            numReps = ceil(K0/(2*CRows-2));
            KVec = repmat(indReflect,[1 numReps]);
            KVec = KVec(1:K0);
                        
            % compute scaling factor for current pole
            C0 = -F{1}(i)./(1-F{1}(i)^2);
            
            % apply symmetric exponential filter for each column
            for k = 1:CSlices
                for j = 1:CCols
                    C(:,j,k) = symExpFilt(C(:,j,k),CRows,C0,F{1}(i),K0,KVec);
                end
            end
        end
        
        % multiply by numerator of direct BSpline filter
        C = real(C)*F0{1};
        
        % now perform direct filtering across each row
        % loop through poles of direct BSpline filter
        for i = 1:length(F{2})
            
            % compute K0 for current pole
            K0 = ceil(log(K0Tol)./log(abs(F{2}(i))));
            
            % now compute vector of indices of length K0 that runs back and
            % forth through the length of the data vector (to mimic
            % reflection of the data at each end)
            indReflect = [1:CCols (CCols-1):-1:2];
            numReps = ceil(K0/(2*CCols-2));
            KVec = repmat(indReflect,[1 numReps]);
            KVec = KVec(1:K0);
                        
            % compute scaling factor for current pole
            C0 = -F{2}(i)./(1-F{2}(i)^2);
            
            % apply symmetric exponential filter for each column
            for k = 1:CSlices
                for j = 1:CRows
                    C(j,:,k) = symExpFilt(C(j,:,k),CCols,C0,F{2}(i),K0,KVec);
                end
            end
        end
        
        % multiply by numerator of direct BSpline filter
        C = real(C)*F0{2};

        % now perform direct filtering across each slice
        % loop through poles of direct BSpline filter
        for i = 1:length(F{3})
            
            % compute K0 for current pole
            K0 = ceil(log(K0Tol)./log(abs(F{3}(i))));
            
            % now compute vector of indices of length K0 that runs back and
            % forth through the length of the data vector (to mimic
            % reflection of the data at each end)
            indReflect = [1:CSlices (CSlices-1):-1:2];
            numReps = ceil(K0/(2*CSlices-2));
            KVec = repmat(indReflect,[1 numReps]);
            KVec = KVec(1:K0);
                        
            % compute scaling factor for current pole
            C0 = -F{3}(i)./(1-F{3}(i)^2);
            
            % apply symmetric exponential filter for each column
            for k = 1:CCols
                for j = 1:CRows
                    C(j,k,:) = symExpFilt(C(j,k,:),CSlices,C0,F{3}(i),K0,KVec);
                end
            end
        end
        
        % multiply by numerator of direct BSpline filter
        C = real(C)*F0{3};
    
    otherwise % X is multidimensional - the most difficult
        
        % get size of coefficients array
        Csz = size(C);
        
        % set up vector of indices into each dimension
        idx = cell(1,tensorOrder);
        for k = 1:tensorOrder
            idx{k} = 1:size(C,k);
        end
        
        % loop through each dimension
        for d = 1:tensorOrder
            
            % copy original version of idx
            idxCurrent = idx;
            
            % construct vector of indices which will be used in computing
            % the initial element of the filtered data
            indReflect = [1:Csz(d) (Csz(d)-1):-1:2];
            
            % construct list of dimensions without d
            dimList = setdiff(1:tensorOrder,d);
            
            % construct index into entries of remaining dimensions
            [idxCurrent{dimList}] = ndgrid(idx{dimList});
            dimListInd = sub2ind(Csz(dimList),idxCurrent{dimList});
            dimListInd = dimListInd(:);
            
            % loop through poles of BSpline filter
            for i = 1:length(F{d})
                
                % compute K0 for current pole
                K0 = ceil(log(K0Tol)./log(abs(F{d}(i))));

                % now compute vector of indices of length K0 that runs back and
                % forth through the length of the data vector (to mimic
                % reflection of the data at each end)
                numReps = ceil(K0/(2*Csz(d)-2));
                KVec = repmat(indReflect,[1 numReps]);
                KVec = KVec(1:K0);

                % compute scaling factor for current pole
                C0 = -F{d}(i)./(1-F{d}(i)^2);
                
                % now loop through remaining dimensions, applying symmetric
                % exponential filter
                for k=1:numel(dimListInd)
                    [idxCurrent{dimList}] = ind2sub(Csz(dimList),dimListInd(k));
                    C(idxCurrent{:}) = symExpFilt(C(idxCurrent{:}),Csz(d),C0,F{d}(i),K0,KVec);
                end                
            end
            
            % multiply by numerator of direct BSpline filter
            C = real(C)*F0{d};

        end
        
end

% now pad dimensions by reflection
padNum = floor(D/2);

% Form index vectors to subsasgn input array into output array.
% Also compute the size of the output array.
idx   = cell(1,tensorOrder);
for k = 1:tensorOrder
  M = size(C,k);
  dimNums = [1:M (M-1):-1:2];
  p = padNum(k);
  idx{k}   = dimNums(mod(-p:M+p, 2*M-2) + 1);
end
CPad = C(idx{:});
```
