## Supplementary Software for "Field dependent deep learning enables high-throughput whole-cell 3D super-resolution imaging": evalBSpline.m

```
function bi = evalBSpline(xi,deg)
% evalBSpline: evaluate BSpline basis function
% usage: bi = evalBSpline(xi,deg);
%
% arguments:
%   xi (N-D arrary) - positions at which to evaluate BSpline basis function
%   deg (scalar) - degree of desired basis function
%
% Note: no error checking is performed for speed
%

% author: Nathan D. Cahill
%
% date: 18 April 2008

switch deg
    case 0
        bi = (xi>=(-1/2))&(xi<(1/2));
    case 1
        bi = zeros(size(xi));
        k = (xi>=0) & (xi<1);
        bi(k) = 1 - xi(k);
        k(:) = (xi>=-1) & (xi<0);
        bi(k) = 1 + xi(k);
    case 2
        bi = zeros(size(xi));
        x2 = xi.*xi;
        k = (xi>=1/2) & (xi<3/2);
        bi(k) = (9/8) - (3/2).*xi(k) + (1/2).*x2(k);
        k = (xi>=-1/2) & (xi<1/2);
        bi(k) = (3/4) - x2(k);
        k = (xi>=-3/2) & (xi<-1/2);
        bi(k) = (9/8) + (3/2).*xi(k) + (1/2).*x2(k);
    case 3
        bi = zeros(size(xi));
        x2 = xi.*xi; x3 = x2.*xi;
        k = (xi>=1) & (xi<2);
        bi(k) = 4/3 - 2.*xi(k) + x2(k) - (1/6).*x3(k);
        k(:) = (xi>=0) & (xi<1);
        bi(k) = 2/3 - x2(k) + (1/2).*x3(k);
        k(:) = (xi>=-1) & (xi<0);
        bi(k) = 2/3 - x2(k) - (1/2).*x3(k);
        k(:) = (xi>=-2) & (xi<-1);
        bi(k) = 4/3 + 2.*xi(k) + x2(k) + (1/6).*x3(k);
    case 4
        bi = zeros(size(xi));
        x2 = xi.*xi; x3 = x2.*xi; x4 = x3.*xi;
        k = (xi>=3/2) & (xi<5/2);
        bi(k) = 625/384 - (125/48).*xi(k) + (25/16).*x2(k) - (5/12).*x3(k) + (1/24).*x4(k);
        k = (xi>=1/2) & (xi<3/2);
        bi(k) = 55/96 + (5/24).*xi(k) - (5/4).*x2(k) + (5/6).*x3(k) - (1/6).*x4(k);
        k = (xi>=-1/2) & (xi<1/2);
        bi(k) = 115/192 - (5/8).*x2(k) + (1/4).*x4(k);
        k = (xi>=-3/2) & (xi<-1/2);
        bi(k) = 55/96 - (5/24).*xi(k) - (5/4).*x2(k) - (5/6).*x3(k) - (1/6).*x4(k);
        k = (xi>=-5/2) & (xi<-3/2);
        bi(k) = 625/384 + (125/48).*xi(k) + (25/16).*x2(k) + (5/12).*x3(k) + (1/24).*x4(k);
    case 5
        bi = zeros(size(xi));
        x2 = xi.*xi; x3 = x2.*xi; x4 = x3.*xi; x5 = x4.*xi;
        k = (xi>=2) & (xi<3);
        bi(k) = 81/40 - (27/8).*xi(k) + (9/4).*x2(k) - (3/4).*x3(k) + (1/8).*x4(k) - (1/120).*x5(k);
        k = (xi>=1) & (xi<2);
        bi(k) = 17/40 + (5/8).*xi(k) - (7/4).*x2(k) + (5/4).*x3(k) - (3/8).*x4(k) + (1/24).*x5(k);
        k = (xi>=0) & (xi<1);
        bi(k) = 11/20 - (1/2).*x2(k) + (1/4).*x4(k) - (1/12).*x5(k);
        k = (xi>=-1) & (xi<0);
        bi(k) = 11/20 - (1/2).*x2(k) + (1/4).*x4(k) + (1/12).*x5(k);
        k = (xi>=-2) & (xi<-1);
        bi(k) = 17/40 - (5/8).*xi(k) - (7/4).*x2(k) - (5/4).*x3(k) - (3/8).*x4(k) - (1/24).*x5(k);
        k = (xi>=-3) & (xi<-2);
        bi(k) = 81/40 + (27/8).*xi(k) + (9/4).*x2(k) + (3/4).*x3(k) + (1/8).*x4(k) + (1/120).*x5(k);
    case 6
        bi = zeros(size(xi));
        x2 = xi.*xi; x3 = x2.*xi; x4 = x3.*xi; x5 = x4.*xi; x6 = x5.*xi;
        k = (xi>=5/2) & (xi<7/2);
        bi(k) = 117649/46080 - (16807/3840).*xi(k) + (2401/768).*x2(k) - (343/288).*x3(k) + (49/192).*x4(k) - (7/240).*x5(k) + (1/720).*x6(k);
        k = (xi>=3/2) & (xi<5/2);
        bi(k) = 1379/7680 + (1267/960).*xi(k) - (329/128).*x2(k) + (133/72).*x3(k) - (21/32).*x4(k) + (7/60).*x5(k) - (1/120).*x6(k);
        k = (xi>=1/2) & (xi<3/2);
        bi(k) = 7861/15360 - (7/768).*xi(k) - (91/256).*x2(k) - (35/288).*x3(k) + (21/64).*x4(k) - (7/48).*x5(k) + (1/48).*x6(k);
        k = (xi>=-1/2) & (xi<1/2);
        bi(k) = 5887/11520 - (77/192).*x2(k) + (7/48).*x4(k) - (1/36).*x6(k);
        k = (xi>=-3/2) & (xi<-1/2);
        bi(k) = 7861/15360 + (7/768).*xi(k) - (91/256).*x2(k) + (35/288).*x3(k) + (21/64).*x4(k) + (7/48).*x5(k) + (1/48).*x6(k);
        k = (xi>=-5/2) & (xi<-3/2);
        bi(k) = 1379/7680 - (1267/960).*xi(k) - (329/128).*x2(k) - (133/72).*x3(k) - (21/32).*x4(k) - (7/60).*x5(k) - (1/120).*x6(k);
        k = (xi>=-7/2) & (xi<-5/2);
        bi(k) = 117649/46080 + (16807/3840).*xi(k) + (2401/768).*x2(k) + (343/288).*x3(k) + (49/192).*x4(k) + (7/240).*x5(k) + (1/720).*x6(k);
    case 7
        bi = zeros(size(xi));
        x2 = xi.*xi; x3 = x2.*xi; x4 = x3.*xi; x5 = x4.*xi; x6 = x5.*xi; x7 = x6.*xi;
        k = (xi>=3) & (xi<4);
        bi(k) = 6405119470038039/1970324836974592 - (672537544353994073/118219490218475520).*xi(k) + (64/15).*x2(k) - (16/9).*x3(k) + (4/9).*x4(k) - (1/15).*x5(k) + (1/180).*x6(k) - (1/5040).*x7(k);
        k = (xi>=2) & (xi<3);
        bi(k) = -2173612320154509/9851624184872960 + (855120979246972939/354658470655426560).*xi(k) - (23/6).*x2(k) + (49/18).*x3(k) - (19/18).*x4(k) + (7/30).*x5(k) - (1/36).*x6(k) + (1/720).*x7(k);
        k = (xi>=1) & (xi<2);
        bi(k) = 103/210 - (7/90).*xi(k) - (1/10).*x2(k) - (7/18).*x3(k) + (1/2).*x4(k) - (7/30).*x5(k) + (1/20).*x6(k) - (1/240).*x7(k);
        k = (xi>=0) & (xi<1);
        bi(k) = 151/315 - (1/3).*x2(k) + (1/9).*x4(k) - (1/36).*x6(k) + (1/144).*x7(k);
        k = (xi>=-1) & (xi<0);
        bi(k) = 151/315 - (1/3).*x2(k) + (1/9).*x4(k) - (1/36).*x6(k) - (1/144).*x7(k);
        k = (xi>=-2) & (xi<-1);
        bi(k) = 103/210 + (7/90).*xi(k) - (1/10).*x2(k) + (7/18).*x3(k) + (1/2).*x4(k) + (7/30).*x5(k) + (1/20).*x6(k) + (1/240).*x7(k);
        k = (xi>=-3) & (xi<-2);
        bi(k) = -2173612320154509/9851624184872960 - (855120979246972939/354658470655426560).*xi(k) - (23/6).*x2(k) - (49/18).*x3(k) - (19/18).*x4(k) - (7/30).*x5(k) - (1/36).*x6(k) - (1/720).*x7(k);
        k = (xi>=-4) & (xi<-3);
        bi(k) = 6405119470038039/1970324836974592 + (672537544353994073/118219490218475520).*xi(k) + (64/15).*x2(k) + (16/9).*x3(k) + (4/9).*x4(k) + (1/15).*x5(k) + (1/180).*x6(k) + (1/5040).*x7(k);
    otherwise
        bi = ((xi + ((deg+1)/2)).*evalBSpline(xi+(1/2),deg-1) + ...
            (((deg+1)/2) - xi).*evalBSpline(xi-(1/2),deg-1))./deg;
end
```
