## Supplementary Software for "Field dependent deep learning enables high-throughput whole-cell 3D super-resolution imaging": getBSplineFiltCoeffs.m

```
function [F,C0] = getBSplineFiltCoeffs(varargin)
% getBSplineFiltCoeffs: get BSpline filter coefficients
% usage: F = getBSplineFiltCoeffs(D);
%    or: F = getBSplineFiltCoeffs(D,CFlag);
%    or: F = getBSplineFiltCoeffs(D,CFlag,direction);
%    or: [F,C0] = getBSplineFiltCoeffs(D,CFlag,direction);
%    or: ... = getBSplineFiltCoeffs(D,CFlag,direction,lambda);
% 
% arguments:
%   D (scalar) - degree of BSpline 
%       D can be any integer in the range [0,7].  Default D = 3.
%   CFlag (logical scalar) - true for centred BSplines, false for shifted
%       BSplines.  Default CFlag = true.
%   direction (string) - 'direct' or 'indirect'. 
%       Default direction = 'indirect'
%   lambda (scalar) - smoothing parameter.  Default lambda = 0.
%
%   F (vector) - if direction = 'indirect', F is the convolution kernel
%       used to implement the indirect BSpline transform. If direction =
%       'direct', F is a vector of the roots of the z-transform of the
%       direct BSpline filter.
%   C0 (scalar) - if direction = 'indirect', C0 is meaningless and will be
%       set to 1.  If direction = 'direct', C0 is the numerator of the
%       direct BSpline filter.
%

% author: Nathan D. Cahill
%
% date: 18 April 2008

% parse input arguments
[D,CFlag,direction,lambda] = parseInputs(varargin{:});

switch lower(direction)
    case 'indirect' % indirect BSpline transform
        C0 = 1;
        switch D
            case 0
                switch CFlag
                    case true
                        if lambda==0
                            F = 1;
                        else
                            error([mfilename,':IndirectNonzeroLambdaNotSupported'],...
                                'Indirect BSpline transform not supported for nonzero lambda.');
                        end
                    case false
                        if lambda==0
                            F = 1;
                        else
                            error([mfilename,':IndirectNonzeroLambdaNotSupported'],...
                                'Indirect BSpline transform not supported for nonzero lambda.');
                        end
                end
            case 1
                switch CFlag
                    case true
                        if lambda==0
                            F = 1;
                        else
                            error([mfilename,':IndirectNonzeroLambdaNotSupported'],...
                                'Indirect BSpline transform not supported for nonzero lambda.');
                        end
                    case false
                        if lambda==0
                            F = [1;1]/2;
                        else
                            error([mfilename,':IndirectNonzeroLambdaNotSupported'],...
                                'Indirect BSpline transform not supported for nonzero lambda.');
                        end
                end
            case 2
                switch CFlag
                    case true
                        if lambda==0
                            F = [1;6;1]/8;
                        else
                            error([mfilename,':IndirectNonzeroLambdaNotSupported'],...
                                'Indirect BSpline transform not supported for nonzero lambda.');
                        end
                    case false
                        if lambda==0
                            F = [1;1]/2;
                        else
                            error([mfilename,':IndirectNonzeroLambdaNotSupported'],...
                                'Indirect BSpline transform not supported for nonzero lambda.');
                        end
                end
            case 3
                switch CFlag
                    case true
                        if lambda==0
                            F = [1;4;1]/6;
                        else
                            error([mfilename,':IndirectNonzeroLambdaNotSupported'],...
                                'Indirect BSpline transform not supported for nonzero lambda.');
                        end
                    case false
                        if lambda==0
                            F = [1;23;23;1]/48;
                        else
                            error([mfilename,':IndirectNonzeroLambdaNotSupported'],...
                                'Indirect BSpline transform not supported for nonzero lambda.');
                        end
                end
            case 4
                switch CFlag
                    case true
                        if lambda==0
                            F = [1;76;230;76;1]/384;
                        else
                            error([mfilename,':IndirectNonzeroLambdaNotSupported'],...
                                'Indirect BSpline transform not supported for nonzero lambda.');
                        end
                    case false
                        if lambda==0
                            F = [1;11;11;1]/24;
                        else
                            error([mfilename,':IndirectNonzeroLambdaNotSupported'],...
                                'Indirect BSpline transform not supported for nonzero lambda.');
                        end
                end
            case 5
                switch CFlag
                    case true
                        if lambda==0
                            F = [1;26;66;26;1]/120;
                        else
                            error([mfilename,':IndirectNonzeroLambdaNotSupported'],...
                                'Indirect BSpline transform not supported for nonzero lambda.');
                        end
                    case false
                        if lambda==0
                            F = [1;237;1682;1682;237;1]/3840;
                        else
                            error([mfilename,':IndirectNonzeroLambdaNotSupported'],...
                                'Indirect BSpline transform not supported for nonzero lambda.');
                        end
                end
            case 6
                switch CFlag
                    case true
                        if lambda==0
                            F = [1;722;10543;23548;10543;722;1]/46080;
                        else
                            error([mfilename,':IndirectNonzeroLambdaNotSupported'],...
                                'Indirect BSpline transform not supported for nonzero lambda.');
                        end
                    case false
                        if lambda==0
                            F = [1;57;302;302;57;1]/720;
                        else
                            error([mfilename,':IndirectNonzeroLambdaNotSupported'],...
                                'Indirect BSpline transform not supported for nonzero lambda.');
                        end
                end
            case 7
                switch CFlag
                    case true
                        if lambda==0
                            F = [1;120;1191;2416;1191;120;1]/5040;
                        else
                            error([mfilename,':IndirectNonzeroLambdaNotSupported'],...
                                'Indirect BSpline transform not supported for nonzero lambda.');
                        end
                    case false
                        if lambda==0
                            F = [1;2179;60657;259723;259723;60657;2179;1]/645120;
                        else
                            error([mfilename,':IndirectNonzeroLambdaNotSupported'],...
                                'Indirect BSpline transform not supported for nonzero lambda.');
                        end
                end
        end
    case 'direct' % direct BSpline transform
        switch D
            case 0
                switch CFlag
                    case true
                        if lambda==0
                            F = [];
                            C0 = 1;
                        else
                            error([mfilename,':DirectEvenDegreeNonzeroLambdaNotSupported'],...
                                'Direct BSpline transform of even degree not supported for nonzero lambda.');
                        end
                    case false
                        error([mfilename,':DirectShiftedNotSupported'],...
                            'Direct BSpline transform not supported for shifted basis functions.');
                end
            case 1
                switch CFlag
                    case true
                        if lambda==0
                            F = [];
                            C0 = 1;
                        else
                            F = 1 + 1/(2*lambda) - sqrt(1+4*lambda)/(2*lambda);
                            C0 = -1/lambda;
                        end
                    case false
                        error([mfilename,':DirectShiftedNotSupported'],...
                            'Direct BSpline transform not supported for shifted basis functions.');
                end
            case 2
                switch CFlag
                    case true
                        if lambda==0
                            F = 2*sqrt(2)-3;
                            C0 = 8;
                        else
                            error([mfilename,':DirectEvenDegreeNonzeroLambdaNotSupported'],...
                                'Direct BSpline transform of even degree not supported for nonzero lambda.');
                        end
                    case false
                        error([mfilename,':DirectShiftedNotSupported'],...
                            'Direct BSpline transform not supported for shifted basis functions.');
                end
            case 3
                switch CFlag
                    case true
                        if lambda==0
                            F = sqrt(3)-2;
                            C0 = 6;
                        else
                            p = [1 -4 6 -4 1];
                            p(2) = p(2) + 1/(6*lambda);
                            p(3) = p(3) + 2/(3*lambda);
                            p(4) = p(4) + 1/(6*lambda);
                            F = roots(p);
                            F = F(abs(F)<=1);
                            C0 = 1/lambda;
                        end
                    case false
                        error([mfilename,':DirectShiftedNotSupported'],...
                            'Direct BSpline transform not supported for shifted basis functions.');
                end
            case 4
                switch CFlag
                    case true
                        if lambda==0
                            F = [-0.36134122590022 -0.0137254292973391];
                            C0 = 384;
                        else
                            error([mfilename,':DirectEvenDegreeNonzeroLambdaNotSupported'],...
                                'Direct BSpline transform of even degree not supported for nonzero lambda.');
                        end
                    case false
                        error([mfilename,':DirectShiftedNotSupported'],...
                            'Direct BSpline transform not supported for shifted basis functions.');
                end
            case 5
                switch CFlag
                    case true
                        if lambda==0
                            F = [-0.430575347099973 -0.0430962882032647];
                            C0 = 120;
                        else
                            p = [1 -6 15 -20 15 -6 1];
                            p(2) = p(2) - 1/(120*lambda);
                            p(3) = p(3) - 13/(60*lambda);
                            p(4) = p(4) - 11/(20*lambda);
                            p(5) = p(5) - 13/(60*lambda);
                            p(6) = p(2) - 1/(120*lambda);
                            F = roots(p);
                            F = F(abs(F)<=1);
                            C0 = -1/lambda;
                        end
                    case false
                        error([mfilename,':DirectShiftedNotSupported'],...
                            'Direct BSpline transform not supported for shifted basis functions.');
                end
            case 6
                switch CFlag
                    case true
                        if lambda==0
                            F = [-0.488294589303046 -0.0816792710762375 -0.00141415180832582];
                            C0 = 46080;
                        else
                            error([mfilename,':DirectEvenDegreeNonzeroLambdaNotSupported'],...
                                'Direct BSpline transform of even degree not supported for nonzero lambda.');
                        end
                    case false
                        error([mfilename,':DirectShiftedNotSupported'],...
                            'Direct BSpline transform not supported for shifted basis functions.');
                end
            case 7
                switch CFlag
                    case true
                        if lambda==0
                            F = [-0.535280430796439 -0.122554615192327 -0.00914869480960828];
                            C0 = 5040;
                        else
                            p = [1 -8 28 -56 70 -56 28 -8 1];
                            p(2) = p(2) + 1/(5040*lambda);
                            p(3) = p(3) + 1/(42*lambda);
                            p(4) = p(4) + 397/(1680*lambda);
                            p(5) = p(5) + 151/(315*lambda);
                            p(6) = p(6) + 397/(1680*lambda);
                            p(7) = p(7) + 1/(42*lambda);
                            p(8) = p(8) + 1/(5040*lambda);
                            F = roots(p);
                            F = F(abs(F)<=1);
                            C0 = 1/lambda;
                        end
                    case false
                        error([mfilename,':DirectShiftedNotSupported'],...
                            'Direct BSpline transform not supported for shifted basis functions.');
                end
        end
end

%% subfunction parseInputs
function [D,CFlag,direction,lambda] = parseInputs(varargin)

% check number of inputs
nargs = length(varargin);
% error(nargchk(0,4,nargs));

% get, check D
if nargs<1
    D = [];
else
    D = varargin{1};
end
if isempty(D)
    D = 3;
end
if numel(D)>1 || D<0 || D>7 || ~isequal(D,round(D))
    error([mfilename,':parseInputs:DNotValid'],...
        'D must be an integer in the interval [0,7]');
end

% get, check CFlag
if nargs<2
    CFlag = [];
else
    CFlag = varargin{2};
end
if isempty(CFlag)
    CFlag = true;
end
if numel(CFlag)>1 || ~islogical(CFlag)
    error([mfilename,':parseInputs:CFlagNotValid'],...
        'CFlag must be a logical scalar');
end

% get, check direction
if nargs<3
    direction = '';
else
    direction = varargin{3};
end
if isempty(direction)
    direction = 'indirect';
end
if ~any(strmatch(lower(direction),{'direct','indirect'}))
    error([mfilename,':parseInputs:directionNotValid'],...
        'Direction must be either ''direct'' or ''indirect''.');
end

% get, check lambda
if nargs<4
    lambda = [];
else
    lambda = varargin{4};
end
if isempty(lambda)
    lambda = 0;
end
if lambda<0
    error([mfilename,':parseInputs:lambdaNotValid'],...
        'lambda must be a nonnegative real number.');
end
```
