## Supplementary Software for "Field dependent deep learning enables high-throughput whole-cell 3D super-resolution imaging": getstackcal_g.m

```
function [splinefit,indgood,posbeads,shift,testallrois,beadsposxy]=getstackcal_g(beads,p)
global stackcal_testfit
% isastig=contains(p.modality,'astig');%||contains(p.modality,'2D');
% alignzastig=isastig&contains(p.zcorr,'astig');
zcorr=contains(p.zcorr,'corr');
sstack=size(beads(1).stack.image);
    halfstoreframes=round((size(beads(1).stack.image,3)-1)/2);
%     if isastig    
%         for B=length(beads):-1:1
%             if  halfstoreframes<length(beads(B).stack.framerange)
%                 dframe(B)=beads(B).stack.framerange(halfstoreframes+1)-beads(B).f0;
%             else
%                 dframe(B)=NaN;
%             end
%         end
%         
%     %remove outliers:
%         badind=abs(dframe-median(dframe,'omitnan'))>10|isnan(dframe);
%         beads(badind)=[];
%     
% 
%         psfx=[beads(:).psfx0];psfy=[beads(:).psfy0];
%         dpsfx=(psfx-median(psfx(~isnan(psfx))))*10;
%         dpsfy=(psfy-median(psfy(~isnan(psfy))))*10;
%     else
        dframe=0;
        dpsfx=0;dpsfy=0;
%     end
    
    allstacks=zeros(sstack(1),sstack(2),sstack(3),length(beads))+NaN;
    allstackst=zeros(sstack(1),sstack(2),sstack(3),length(beads))+NaN;
%     allstackstm=zeros(sstack(1),sstack(2),sstack(3),length(beads))+NaN;
    goodvs=[];
%%test    Unable to perform assignment because the size of the left side is 33-by-33-by-41 and the size of the right side is 33-by-33-by-3.
%     for i=length(beads):-1:1
%         if length(beads(i).stack.framerange)~=41
%             i
%         end
%     end
    for B=length(beads):-1:1
        allstacks(:,:,:,B)=beads(B).stack.image;
        
%         if p.isglobalfit
%             if ~p.mirror
%                 allstackst(:,:,:,B)=beads(B).stack.imagetar;
%                 shiftxy(B,1:2)=beads(B).shiftxy;
%                 mirroraxis=0;
%             else
%                 if contains(p.Tmode,'up-down')
%                 %this is for up down mirror. add also for right left mirror
%                     allstackst(:,:,:,B)=beads(B).stack.imagetar(end:-1:1,:,:);
%                     shiftxy(B,1:2)=beads(B).shiftxy;
%                     shiftxy(B,2)=-shiftxy(B,2);
%                     mirroraxis=2;
%                 else %right left
%                     allstackst(:,:,:,B)=beads(B).stack.imagetar(:,end:-1:1,:);
%                     shiftxy(B,1:2)=beads(B).shiftxy;
%                     shiftxy(B,1)=-shiftxy(B,1);
%                     mirroraxis=1;
%                 end
%             end
%         else
            shiftxy(B,1:2)=[0 0];
%         end
        stackh=allstacks(:,:,:,B);
        goodvs(B)=sum(~isnan(stackh(:)))/numel(stackh);
        
    end
    
    mstack=mean(allstacks,4,'omitnan');
    mstack=mstack-min(mstack(:),[],'omitnan');
    mstack=mstack/sum(mstack(:),'omitnan');
    for k=length(beads):-1:1
    	stackh=(allstacks(:,:,:,k));
        stackh=stackh-min(stackh(:),[],'omitnan');
        stackh=stackh/sum(stackh(:),'omitnan');
        dstack(k)=sum((stackh(:)-mstack(:)).^2);
    end
    dstack=dstack/mean(dstack);  
    
%     if p.isglobalfit
%         mstack=mean(allstackst,4,'omitnan');
%         mstack=mstack-min(mstack(:),[],'omitnan');
%         mstack=mstack/sum(mstack(:),'omitnan');
%         for k=length(beads):-1:1
%             stackh=(allstackst(:,:,:,k));
%             stackh=stackh-min(stackh(:),[],'omitnan');
%             stackh=stackh/sum(stackh(:),'omitnan');
%             dstackt(k)=sum((stackh(:)-mstack(:)).^2);
%         end
%         dstackt=dstackt/mean(dstackt);  
% 
%        
%     else
        dstackt=0;
        allstackst=[];
%     end
        devs=(dpsfx.^2+dpsfy.^2+dstack+dstackt)./goodvs;


    if zcorr  
        fw2=round((p.zcorrframes-1)/2);
        
    else
        fw2=2;
    end
   

    [~,sortinddev]=sort(devs);

%     if alignzastig
%          zshift=dframe-round(median(dframe));
%     else
        zshift=[];
%     end
    
    midrange=halfstoreframes+1-round(median(dframe));
     framerange=max(1,midrange-fw2):min(midrange+fw2,size(stackh,3));
    p.status.String='calculate shift of individual PSFs';drawnow
    filenumber=[beads(:).filenumber];
%     [corrPSF,shiftedstack,shift,beadgood]=registerPSF3D_g(allrois,allroist,struct('sortind',sortinddev,'shiftxy',shiftxys,'framerange',framerange,'alignz',zcorr,'zshiftf0',zshift,'beadfilterf0',false,'status',p.status),{},filenumber(sortinddev));
%start add by shiwei 2020/08/14
%     [corrPSF,shiftedstack,shift,beadgood]=registerPSF3D_g(allstacks,allstackst,struct('sortind',sortinddev,'shiftxy',shiftxy,'framerange',framerange,'alignz',zcorr,'zshiftf0',zshift,'beadfilterf0',false,'status',p.status),{},filenumber(sortinddev));
    [corrPSF,shiftedstack,shift,beadgood,corrPSF_all]=registerPSF3D_g(allstacks,allstackst,struct('sortind',sortinddev,'shiftxy',shiftxy,'framerange',framerange,'alignz',zcorr,'zshiftf0',zshift,'beadfilterf0',false,'status',p.status),{},filenumber(sortinddev));
%     corrPSFr_all = corrPSF_all(1:size(allstacks,1),:,:,:);
    corrPSFr_all = corrPSF_all;
    beadsposxy = cell(1,length(beads));
    for k=length(beads):-1:1
        beadsposxy{k}=beads(k).pos;
    end 
    beadsposxy=beadsposxy(beadgood);
%     for k=length(beads):-1:1
%         tempx(k)=beads(k).pos(1);
%         tempy(k)=beads(k).pos(2);
%     end 
%     beadsposxy=[tempx(beadgood);tempy(beadgood)];
%end add by shiwei 2020/08/14
    corrPSFr=corrPSF(1:size(allstacks,1),:,:);
    
    indgood=beadgood;
    allrois=allstacks;
     %cut out the central part of the PSF correspoinding to the set
    %Roisize in x,y and z

    scorrPSF=size(corrPSFr);
    x=round((scorrPSF(1)+1)/2);y=round((scorrPSF(2)+1)/2);

    dRx=round((p.ROIxy-1)/2);
  
    p.ROIz=size(corrPSFr,3);
    dzroi=round((p.ROIz-1)/2);


    rangex=x-dRx:x+dRx;
    rangey=y-dRx:y+dRx;
    z=midrange;%always same reference: z=f0
    rangez=max(2,z-dzroi):min(size(corrPSFr,3)-1,z+dzroi); %first/last slice distorted by shifts...
    z0reference=find(rangez>=z,1,'first');

    %careful: like this each PSF is normalized individually. This might
    %not be the right approahc. Then normalize by one only
    %normalize PSF
    centpsfr=corrPSFr(rangex,rangey,z-1:z+1); %cut out rim from shift    
    minPSFr=min(centpsfr(:),[],'omitnan');
    corrPSFnr=corrPSFr-minPSFr;
    intglobalr=mean(sum(sum(corrPSFnr(rangex,rangey,z-1:z+1),1,'omitnan'),2,'omitnan'),'omitnan');
    corrPSFnr=corrPSFnr/intglobalr;   
    shiftedstack(1:size(allrois,1),:,:,:)=(shiftedstack(1:size(allrois,1),:,:,:)-minPSFr)/intglobalr;
    corrPSFnr(isnan(corrPSFnr))=0;
    corrPSFnr(corrPSFnr<0)=0;
    corrPSFsr=corrPSFnr(rangex,rangey,rangez);   
        
    PSFgood=true;

    %calculate effective smoothing factor. For dz=10 nm, pixelsize= 130
    %nm, a value around 1 produces visible but not too much smoothing.
%     lambdax=p.smoothxy/p.cam_pixelsize_um(1)/100000;
    if ~isfield(p,'pixelsize')
        pixelsizeh=100;
    else
        pixelsizeh=p.pixelsize{1}(1);
    end
    lambdax=p.smoothxy/pixelsizeh/100000;
    lambdaz=p.smoothz/p.dz*100;
    lambda=[lambdax lambdax lambdaz];
    %calculate smoothed bsplines
    b3_0r=bsarray(double(corrPSFsr),'lambda',lambda);
     %calculate smoothed volume
    zhd=1:1:b3_0r.dataSize(3);
    dxxhd=1;
    [XX,YY,ZZ]=meshgrid(1:dxxhd:b3_0r.dataSize(1),1:dxxhd:b3_0r.dataSize(2),zhd);
    p.status.String='calculating calibration coefficients in progress';drawnow
    corrPSFhdr = interp3_0(b3_0r,XX,YY,ZZ,0);
    %calculate cspline coefficients
    coeffr = single(Spline3D_interp(corrPSFhdr));
        
%     if p.isglobalfit
%         corrPSFt=corrPSF(size(allstacks,1)+1:end,:,:);
%         allroist=allstackst;
%         centpsft=corrPSFt(rangex,rangey,z-1:z+1);
%         minPSFt=min(centpsft(:),[],'omitnan');
%         corrPSFnt=corrPSFt-minPSFt;
%         intglobalt=mean(sum(sum(corrPSFnt(rangex,rangey,z-1:z+1),1,'omitnan'),2,'omitnan'),'omitnan');
%         %normalize also by the same as reference!
% %         intglobalt=intglobalr;
%         corrPSFnt=corrPSFnt/intglobalr;
%         shiftedstack(size(allrois,1)+1:end,:,:,:)=(shiftedstack(size(allrois,1)+1:end,:,:,:)-minPSFt)/intglobalr;        
%         corrPSFnt(isnan(corrPSFnt))=0;
%         corrPSFnt(corrPSFnt<0)=0;
%         corrPSFst=corrPSFnt(rangex,rangey,rangez);
%         b3_0t=bsarray(double(corrPSFst),'lambda',lambda);
%         corrPSFhdt = interp3_0(b3_0t,XX,YY,ZZ,0);
%         coefft = single(Spline3D_interp(corrPSFhdt));
%         
%         switch mirroraxis
%             case 0
%                 PSFtm=corrPSFhdt;
%             case 2
%                 PSFtm=corrPSFhdt(end:-1:1,:,:);
%             case 1
%                 PSFtm=corrPSFhdt(:,end:-1:1,:);
%         end
%         coefftnomirror=single(Spline3D_interp(PSFtm));
%         %assemble output structure for saving
%          b3_0r.coeffs=single(b3_0r.coeffs);
%          b3_0t.coeffs=single(b3_0t.coeffs);
%         bspline.bslpine=({b3_0r,b3_0t});
%         cspline.coeff={single(coeffr), single(coefft)};
%         splinefit.PSF={single(corrPSFr),single(corrPSFt)};
%         splinefit.PSFsmooth={single(corrPSFhdr),single(corrPSFhdt)};
%         cspline.coeffrawref=single(coeffr);
%         cspline.coeffrawtar=single(coefftnomirror);
%         cspline.normf=[intglobalr intglobalt]/intglobalr;
%         cspline.mirror=mirroraxis;
        
%     else
        b3_0r.coeffs=single(b3_0r.coeffs);
        bspline.bslpine=({b3_0r});
        cspline.coeff={single(coeffr)};    
        splinefit.PSF={single(corrPSFr)};
        splinefit.PSF_all={single(corrPSFr_all)};%add by shiwei 2020/08/14
        splinefit.PSFsmooth={single(corrPSFhdr)};   
        cspline.mirror=0;
%     end
   
        
        cspline.z0=z0reference;%round((b3_0.dataSize(3)+1)/2);
        cspline.dz=p.dz;
        cspline.x0=dRx+1;
        bspline.z0=round((b3_0r.dataSize(3)+1)/2);
        bspline.dz=p.dz;            
        splinefit.bspline=bspline;
        p.z0=cspline.z0;
        

        splinefit.cspline=cspline;

        %plot graphs
        if PSFgood      
%             dL=size(shiftedstack,1)/2;
%             ax=axes(uitab(p.tabgroup,'Title','PSFz'));
%              framerange0=max(p.fminmax(1)):min(p.fminmax(2));
%              halfroisizebig=(size(shiftedstack,2)-1)/2;         
%             ftest=z;
%             xt=x;
%             yt=y;
%             zpallr=squeeze(shiftedstack(xt,yt,:,beadgood));
%             xpall=squeeze(shiftedstack(:,yt,ftest,beadgood));
%             zprofiler=squeeze(corrPSFnr(xt,yt,:));
%             xprofiler=squeeze(corrPSFnr(:,yt,ftest));
%             dxxx=0.1;
%             xxx=1:dxxx:b3_0r.dataSize(1);            
%             zzzt=0*xxx+ftest;
%             xbsr= interp3_0(b3_0r,0*xxx+b3_0r.dataSize(1)/2+.5,xxx,zzzt);
%             zzz=1:dxxx:b3_0r.dataSize(3);xxxt=0*zzz+b3_0r.dataSize(1)/2+.5;
%             zbsr= interp3_0(b3_0r,xxxt,xxxt,zzz); 
%             hold(ax,'off')
%             h1=plot(ax,framerange0,zpallr(1:length(framerange0),:),'c');
%             hold(ax,'on')
%             dF=max(framerange0);
%             h2=plot(ax,framerange0',zprofiler(1:length(framerange0)),'k*');
%             h3=plot(ax,zzz+rangez(1)+framerange0(1)-2,zbsr,'b','LineWidth',2);
%             xlabel(ax,'frames')
%             ylabel(ax,'normalized intensity')
%             ax.XLim(1)=min(framerange0);
%             title(ax,'Profile along z for x=0, y=0');          
%             ax.XLim(2)=max(framerange0);
%              
% 
%             
%             xrange=-halfroisizebig:halfroisizebig;
%             ax=axes(uitab(p.tabgroup,'Title','PSFx'));
%             hold(ax,'off')
%            
%             axp=(uitab(p.tabgroup,'Title','PSF'));
%             
% 
%             plot(ax,xrange,xpall,'c')
%             hold(ax,'on')
%             plot(ax,xrange,vertcat(xprofiler),'k*-')
%             plot(ax,(xxx-(b3_0r.dataSize(1)+1)/2),xbsr,'b','LineWidth',2)  
%             imageslicer(corrPSFnr,'Parent',axp);
%             
%             xlabel(ax,'x (pixel)')
%             ylabel(ax,'normalized intensity')
%             title(ax,'Profile along x for y=0, z=0');
%             legend([h1(1),h2,h3],'individual PSFs','average PSF','smoothed spline')
%             
% 
%             drawnow
            %quality control: refit all beads
            if isempty(stackcal_testfit)||stackcal_testfit  %not implemented yet in fitter. Fix later
                ax=axes(uitab(p.tabgroup,'Title','validate'));
                testallrois(:,:,:,:,1)=allrois(:,:,:,beadgood); 
                corrPSFfit=corrPSF/max(corrPSF(:))*max(testallrois(:)); %bring back to some reasonable photon numbers;
                corrPSFfitf(:,:,:,1,1)=corrPSFfit(1:size(corrPSFfit,2),:,:);
                
                testallrois(isnan(testallrois))=0;
                posbeads=testfit_spline(testallrois,cspline.coeff,shiftxy(beadgood,:),p,{},ax);
                drawnow
                
                %add coordinates of rois
                
                beadshere=beads(indgood);
                posbeads.filenumber=0*posbeads.x;
                
                for k=1:size(posbeads.x,2)
                   %test if right roi!
                    posbeads.xim(:,k)=posbeads.x(:,k)+beadshere(k).pos(1); 
                    posbeads.yim(:,k)=posbeads.y(:,k)+beadshere(k).pos(2);
                    posbeads.x(:,k)=posbeads.xim(:,k)+beadshere(k).roi(1);
                    posbeads.y(:,k)=posbeads.yim(:,k)+beadshere(k).roi(2);
                    posbeads.filenumber(:,k)=beadshere(k).filenumber;
                end
                

            else
                posbeads=[];
            end
        end 
end


% function teststripes(coeff,p,ax)
% %not used, can be called to test for stripe artifacts.
% tt=tic;
% 
% zr=0:0.2:p.ROIz;
% xr=0:0.05:p.ROIxy;
% hz=zeros(1,length(zr)-1);
% hx=zeros(1,length(xr)-1);
% hy=hx;
% while toc(tt)<30
%     nn=rand(11,11,10000,'single');
% %     P=callYimingFitter(nn,single(coeff),50,5,0,1);
%     [P] =  mleFit_LM(nn,5,50,single(coeff),0,1);
%     hz=histcounts(P(:,5),zr)+hz;
%     hx=histcounts(P(:,1),xr)+hx;
%     hy=histcounts(P(:,2),xr)+hy;
%     
% end
% 
% hz(1)=[];hz(end)=[];
% hz(1)=0;hz(end)=0;
% 
% indx=(hx==0);
% hx(indx)=[];
% indy=(hy==0);
% hy(indy)=[];
% hx(1)=[];hx(end)=[];
% hy(1)=[];hy(end)=[];
% hzx=myxcorr(hz-mean(hz),hz-mean(hz));
% hxx=myxcorr(hx-mean(hx),hx-mean(hx));
% hyx=myxcorr(hy-mean(hy),hy-mean(hy));
% hzx(1)=0;hzx(end)=0;
% ax2=axes(ax.Parent);
% subplot(1,2,1,ax);
% subplot(1,2,2,ax2);
% findx=find(~indx);findy=find(~indy);
% plot(ax,zr(2:end-2),hz,zr(2:end-2),hzx/max(hzx)*(myquantile(hz,.99)));
% ax.YLim(2)=(myquantile(hz,.99))*1.1;
% ax.YLim(1)=min(myquantile(hz,.01),myquantile(hzx/max(hzx)*(myquantile(hz,.99)),.01));
% plot(ax2,xr(findx(2:end-1)),hx,xr(findx(2:end-1)),hxx/max(hxx)*max(hx),xr(findy(2:end-1)),hy,xr(findy(2:end-1)),hyx/max(hyx)*max(hy));
% end
```
