## Supplementary Software for "Field dependent deep learning enables high-throughput whole-cell 3D super-resolution imaging": images2beads_globalfit.m

```
function [b,p]=images2beads_globalfit(p)
% addpath('bfmatlab')
fs=p.filtersize;
hfilterim=fspecial('gaussian',2*round(fs*3/2)+1,fs);
fmax=0;
roisize=p.ROIxy;
roisizeh=min(round(1.5*(p.ROIxy-1)/2),(p.ROIxy-1)/2+3); %create extra space if we need to shift;
rsr=-roisizeh:roisizeh;
filelist=p.filelist;
b=[];
ht=uitab(p.tabgroup,'Title','Files');
tg=uitabgroup(ht);
p.mirror=false;
p.multifile=false;
for k=1:length(filelist)
    ax=axes(uitab(tg,'Title',num2str(k)));
%     axis(ax,'image')
    p.fileax(k)=ax;
    if contains(filelist{k},';')
        p.multifile=true;
        ind=strfind(filelist{k},';');
        filelisth=filelist{k}(1:ind-1);
        filelisth2=filelist{k}(ind+1:end);
        [imstack2, p.roi2{k}, p.pixelsize2{k},settings3D2,isem1]=readbeadimages(filelisth2,p);
        [imstack, p.roi{k}, p.pixelsize{k},settings3D,isem2]=readbeadimages(filelisth,p);
        p.emgain=[isem1,isem2];
    else
        filelisth=filelist{k};
        [imstack, p.roi{k}, p.pixelsize{k},settings3D,p.emgain]=readbeadimages(filelisth,p);
%         imstack = imstack(128:383,128:383,:); %add by shiwei 2020/09/10  Limit image size %mark
%         p.roi{k} = p.roi{k}/2;%add by shiwei 2020/09/10  Limit image size %mark
        p.roi2=p.roi;
        imstack2=imstack;
    end
    
    imstack=imstack-min(imstack(:)); %fast fix for offset;
    imstack2=imstack2-min(imstack2(:)); %fast fix for offset;
%     imageslicer(imstack)%%%%%%%%XXXXXXX
    
    mim=max(imstack,[],3);

%     mim=mean(imstack,3);
    mim=filter2(hfilterim,mim);
    imagesc(ax,mim);
    axis(ax,'image');
    axis(ax,'off')
    title(ax,'Maximum intensity projection')
    if isfield(p,'beadpos') %passed on externally
        maxima=round(p.beadpos{k});
    else
        if isfield(p,'roimask')&&~isempty(p.roimask)
            mim=double(mim).*double(p.roimask);
        end
        maxima=maximumfindcall(mim);
        indgh=maxima(:,1)>p.xrange(1) & maxima(:,1)<p.xrange(2) & maxima(:,2)>p.yrange(1) & maxima(:,2)<p.yrange(2); 
        %XXXXXXXXXXX
%         maxima(:,1)=(maxima(:,1)+2*round(rand(size(maxima,1),1)))-1; % for testing if positions match
        int=maxima(:,3);
        try
        r1=max(roisize,p.yrange(1)):min(size(mim,1)-roisize,p.yrange(2));
        r2=max(roisize,p.xrange(1)):min(size(mim,2)-roisize,p.xrange(2));
        mimc=mim(r1,r2);
        mmed=myquantile(mimc(:),0.3);
        imt=mimc(mimc<=mmed);
            sm=sort(int);
        mv=mean(sm(end-5:end));
%       cutoff=mean(imt(:))+max(2.5*std(imt(:)),(mv-mean(imt(:)))/15);
        cutoff=mean(imt(:))+max(2.5*std(imt(:)),(mv-mean(imt(:)))/15);   %mark shiwei 2022/04/07 
        catch
            cutoff=myquantile(mimc(:),.95);
        end
        cutoff=cutoff*p.cutoffrel;
        if any(int>cutoff)
            maxima=maxima(int>cutoff & indgh,:);
        else
            [~,indm]=max(int);
            maxima=maxima(indm,:);
        end
    end
    maximafound=maxima;
    indgoodb=true(size(maxima,1),1);
    %remove beads that are closer together than mindistance
    if isfield(p,'mindistance')&&~isempty(p.mindistance)
        
        for bk=1:size(maxima,1)
            for bl=bk+1:size(maxima,1)
                if  sum((maxima(bk,1:2)-maxima(bl,1:2)).^2)<p.mindistance^2
                    indgoodb(bk)=false;
                    indgoodb(bl)=false;
                end
            end
        end 
       if isfield(p,'settings_3D')
           w=p.settings_3D.width4pi;
           xm=mod(maxima(:,1),w);
           xm(xm>w/2)=xm(xm>w/2)-w;
           indgoodb(abs(xm)<p.mindistance/2)=false;
       end
       hs=size(imstack,1);
       ym=maxima(:,2);
       ym(ym>hs/2)=ym(ym>hs/2)-hs;
       indgoodb(abs(ym)<p.mindistance/2)=false;
%         maxima=maxima(indgoodb,:);
         maxima=maxima(indgoodb,:);
%          maxima=maximafound    %mark modify by shiwei 2022/04/07
    end 
    
  

    indgoodr=maxima(:,1)>p.xrange(1)&maxima(:,1)<p.xrange(end)&maxima(:,2)>p.yrange(1)&maxima(:,2)<p.yrange(end);
    maxima=maxima(indgoodr,:);
%         maxima=maxima;%mark modify by shiwei 2022/04/07
    maximaref=maxima;
    maximatar=maxima;
    dxy=zeros(size(maximatar));
%     end
   
    numframes=size(imstack,3);
    bind=length(b)+size(maximaref,1);
%     beadsdir =strcat(pwd,'\beads_imstack.mat');  %test by shiwei           
%     save(beadsdir,'imstack');
    for l=1:size(maximaref,1)
        b(bind).loc.frames=(1:numframes)';
        b(bind).loc.filenumber=zeros(numframes,1)+k;
        b(bind).filenumber=k;
%         b(bind).pos=maximaref(l,1:2);
%         b(bind).postar=maximatar(l,1:2);
        b(bind).pos=[maximaref(l,1:2),k];   %modify by shiwei to add the frame index
        b(bind).postar=[maximatar(l,1:2),k];%modify by shiwei to add the frame index
        b(bind).shiftxy=dxy(l,:);
        try
            b(bind).stack.image=imstack(b(bind).pos(2)+rsr,b(bind).pos(1)+rsr,:);
            b(bind).stack.imagetar=imstack2(b(bind).postar(2)+rsr,b(bind).postar(1)+rsr,:);
            b(bind).stack.framerange=1:numframes;
            b(bind).isstack=true;
            
        catch err
            b(bind).isstack=false;
%             err
        end
        
            b(bind).roi=p.roi{k};
            b(bind).roi2=p.roi2{k};
        bind=bind-1;
    end
    fmax=max(fmax,numframes);
    hold (ax,'on');

    plot(ax,maxima(:,1),maxima(:,2),'ko',maximafound(:,1),maximafound(:,2),'r.');
%     end
    hold (ax,'off');
    drawnow
end
indgoodbead=[b(:).isstack];
b=b(indgoodbead);

p.fminmax=[1 fmax];

p.pathhere=fileparts(filelist{1});
end
```
