## Supplementary Software for "Field dependent deep learning enables high-throughput whole-cell 3D super-resolution imaging": interp3_0.m

```
function varargout = interp3_0(varargin)
% bsarray/interp3: 3-D interpolation (table lookup)
% usage: VI = interp3(B,XI,YI,ZI);
%    or: VI = interp3(B,XI,YI,ZI,EXTRAPVAL);
%
% arguments:
%   B - bsarray object having tensorOrder 2. The positions of the
%       x-coordinates of the underlying data array are assumed to be 
%       X = s(2).*(1:Nx), where Nx is the number of data points in the X 
%       dimension (i.e., second element of get(B,'dataSize')), and s 
%       is the element spacing (i.e., get(B,'elementSpacing')). The
%       positions of the y-coordinates of the underlying data array are
%       assumed to be Y = s(1).*(1:Ny). The positions of the z-coordinates
%       of the underlying data array are assumed to be Z = s(3).*(1:Nz).
%   XI - x-coordinates of points at which to interpolate B.
%   YI - y-coordinates of points at which to interpolate B. YI must be the
%       same size as XI.
%   ZI - z-coordinates of points at which to interpolate B. ZI must be the
%       same size as XI.
%
%   EXTRAPVAL - value to return for points in XI, YI, or ZI that are 
%       outside the ranges of X, Y, and Z, respectively. 
%       Default EXTRAPVAL = NaN.
%
%   VI - the values of the underlying bsarray B evaluated at the points in
%       the array XI.
%

% author: Nathan D. Cahill
%
% date: 18 April 2008

% parse input arguments
[b,xi,yi,zi,extrapval] = parseInputs(varargin{:});

% get flag to determine if basis functions in each dimension are centred or
% shifted
% m = double(get(b,'centred'));
m = double(b.centred);
mx = m(2); my = m(1); mz = m(3);

% get number of data elements and coefficients, determine amount of padding
% that has been done to create coefficients
% nData = get(b,'dataSize'); 
nData = b.dataSize;
nDatax = nData(2); nDatay = nData(1); nDataz = nData(3);
% n = get(b,'coeffsSize'); 
n = b.coeffsSize;
nx = n(2); ny = n(1); nz = n(3);
padNum = (n-nData-1)/2; 
padNumx = padNum(2); padNumy = padNum(1); padNumz = padNum(3);

% get the spacing between elements, and then construct vectors of the
% locations of the data and of the BSpline coefficients
% h = get(b,'elementSpacing'); 
h = b.elementSpacing; 
hx = h(2); hy = h(1); hz = h(3);
xCol = hx.*((1-padNumx):(nDatax+padNumx+1))';
xDataCol = xCol((1+padNumx):(end-padNumx));
yCol = hy.*((1-padNumy):(nDatay+padNumy+1))';
yDataCol = yCol((1+padNumy):(end-padNumy));
zCol = hz.*((1-padNumz):(nDataz+padNumz+1))';
zDataCol = zCol((1+padNumz):(end-padNumz));

% turn evaluation points into a column vector, but retain original size so
% output can be returned in same size as input
siz_xi = size(xi);
siz_vi = siz_xi;

% grab the BSpline coefficients
% cMat = get(b,'coeffs');
cMat = b.coeffs;
% initialize some variables for use in interpolation
numelXi = numel(xi);
viMat = zeros(numelXi,1);
p = 1:numelXi;

% Find indices of subintervals, x(k) <= u < x(k+1),
% or u < x(1) or u >= x(m-1).
% kx = min(max(1+floor((xi(:)-xCol(1))/hx),1+padNumx),nx-padNumx) + 1-mx;
% ky = min(max(1+floor((yi(:)-yCol(1))/hy),1+padNumy),ny-padNumy) + 1-my;
% kz = min(max(1+floor((zi(:)-zCol(1))/hz),1+padNumz),nz-padNumz) + 1-mz;
%modify YL
kx = min(max(1+floor((xi(:)-xCol(1))/hx),1+padNumx),nx-padNumx-1) + 1-mx;
ky = min(max(1+floor((yi(:)-yCol(1))/hy),1+padNumy),ny-padNumy-1) + 1-my;
kz = min(max(1+floor((zi(:)-zCol(1))/hz),1+padNumz),nz-padNumz-1) + 1-mz;

sx = (xi(:) - xCol(kx))/hx;
sy = (yi(:) - yCol(ky))/hy;
sz = (zi(:) - zCol(kz))/hz;

% perform interpolation
% d = get(b,'degree'); 
d = b.degree; dx = d(2); dy = d(1); dz = d(3);
xflag = (~mx && mod(dx,2));
yflag = (~my && mod(dy,2));
zflag = (~mz && mod(dz,2));
for q = 1:ceil((dz+1)/2)
    Bz1 = evalBSpline(sz+q-(1+mz)/2,dz);
    Bz2 = evalBSpline(sz-q+(1-mz)/2,dz);
    for j=1:ceil((dx+1)/2) % loop over BSpline degree in x dimension
        Bx1 = evalBSpline(sx+j-(1+mx)/2,dx);
        Bx2 = evalBSpline(sx-j+(1-mx)/2,dx);
        for i=1:ceil((dy+1)/2) % loop over BSpline degree in y dimension
            By1 = evalBSpline(sy+i-(1+my)/2,dy);
            By2 = evalBSpline(sy-i+(1-my)/2,dy);
            for ind = 1:numelXi % loop over evaluation points, computing interpolated value
                viMat(ind) = viMat(ind) + ...
                    cMat(ky(ind)-i+my,kx(ind)-j+mx,kz(ind)-q+mz).*By1(ind).*Bx1(ind).*Bz1(ind) + ...
                    cMat(ky(ind)+i-1+my,kx(ind)-j+mx,kz(ind)-q+mz).*By2(ind).*Bx1(ind).*Bz1(ind) + ...
                    cMat(ky(ind)-i+my,kx(ind)+j-1+mx,kz(ind)-q+mz).*By1(ind).*Bx2(ind).*Bz1(ind) + ...
                    cMat(ky(ind)+i-1+my,kx(ind)+j-1+mx,kz(ind)-q+mz).*By2(ind).*Bx2(ind).*Bz1(ind) + ...
                    cMat(ky(ind)-i+my,kx(ind)-j+mx,kz(ind)+q-1+mz).*By1(ind).*Bx1(ind).*Bz2(ind) + ...
                    cMat(ky(ind)+i-1+my,kx(ind)-j+mx,kz(ind)+q-1+mz).*By2(ind).*Bx1(ind).*Bz2(ind) + ...
                    cMat(ky(ind)-i+my,kx(ind)+j-1+mx,kz(ind)+q-1+mz).*By1(ind).*Bx2(ind).*Bz2(ind) + ...
                    cMat(ky(ind)+i-1+my,kx(ind)+j-1+mx,kz(ind)+q-1+mz).*By2(ind).*Bx2(ind).*Bz2(ind);
            end
        end
        if yflag % add a correction factor if BSpline in y direction is shifted and of odd degree
            By1 = evalBSpline(sy+i+1/2,dy);
            for ind = 1:numelXi
                viMat(ind) = viMat(ind) + By1(ind).*...
                    (cMat(ky(ind)-i-1,kx(ind)-j+mx,kz(ind)-q+mz).*Bx1(ind).*Bz1(ind) + ...
                    cMat(ky(ind)-i-1,kx(ind)+j-1+mx,kz(ind)-q+mz).*Bx2(ind).*Bz1(ind) + ...
                    cMat(ky(ind)-i-1,kx(ind)-j+mx,kz(ind)+q-1+mz).*Bx1(ind).*Bz2(ind) + ...
                    cMat(ky(ind)-i-1,kx(ind)+j-1+mx,kz(ind)+q-1+mz).*Bx2(ind).*Bz2(ind));
            end
        end
    end
    if xflag % add a correction factor if BSpline in x direction is shifted and of odd degree
        Bx1 = evalBSpline(sx+j+1/2,dx);
        for i=1:ceil((dy+1)/2)
            By1 = evalBSpline(sy+i-(1+my)/2,dy);
            By2 = evalBSpline(sy-i+(1-my)/2,dy);
            for ind = 1:numelXi
                viMat(ind) = viMat(ind) + Bx1(ind).*...
                    (cMat(ky(ind)-i+my,kx(ind)-j-1,kz(ind)-q+mz).*By1(ind).*Bz1(ind) + ...
                    cMat(ky(ind)+i-1+my,kx(ind)-j-1,kz(ind)-q+mz).*By2(ind).*Bz1(ind) + ...
                    cMat(ky(ind)-i+my,kx(ind)-j-1,kz(ind)+q-1+mz).*By1(ind).*Bz2(ind) + ...
                    cMat(ky(ind)+i-1+my,kx(ind)-j-1,kz(ind)+q-1+mz).*By2(ind).*Bz2(ind));
            end
        end
    end
end
if zflag % add a correction factor if BSpline in z direction is shifted and of odd degree
    Bz1 = evalBSpline(sz+q+1/2,dz);
    for j=1:ceil((dx+1)/2)
        Bx1 = evalBSpline(sx+j-(1+mx)/2,dx);
        Bx2 = evalBSpline(sx-j+(1-mx)/2,dx);
        for i=1:ceil((dy+1)/2)
            By1 = evalBSpline(sy+i-(1+my)/2,dy);
            By2 = evalBSpline(sy-i+(1-my)/2,dy);
            for ind = 1:numelXi
                viMat(ind) = viMat(ind) + Bz1(ind).*...
                    (cMat(ky(ind)-i+my,kx(ind)-j+mx,kz(ind)-q-1).*By1(ind).*Bx1(ind) + ...
                    cMat(ky(ind)+i-1+my,kx(ind)-j+mx,kz(ind)-q-1).*By2(ind).*Bx1(ind) + ...
                    cMat(ky(ind)-i+my,kx(ind)+j-1+mx,kz(ind)-q-1).*By1(ind).*Bx2(ind) + ...
                    cMat(ky(ind)+i-1+my,kx(ind)+j-1+mx,kz(ind)-q-1).*By2(ind).*Bx2(ind));
            end
        end
    end
end

% perform extrapolation
outOfBounds = xi(:)<xDataCol(1) | xi(:)>xDataCol(nDatax) | yi(:)<yDataCol(1) | yi(:)>yDataCol(nDatay) | zi(:)<zDataCol(1) | zi(:)>zDataCol(nDataz);
viMat(p(outOfBounds)) = extrapval;

% reshape result to have same size as input xi
vi = reshape(viMat,siz_vi);
varargout{1} = vi;


%% subfunction parseInputs
function [b,xi,yi,zi,extrapval] = parseInputs(varargin)

nargs = length(varargin);
% error(nargchk(4,5,nargs));

% Process B
b = varargin{1};
if ~isequal(b.tensorOrder,3)
    error([mfilename,'parseInputs:WrongOrder'], ...
        'bsarray/interp3 can only be used with bsarray objects having tensor order 3.');
end

% Process XI
xi = varargin{2};
if ~isreal(xi)
    error([mfilename,'parseInputs:ComplexInterpPts'], ...
        'The interpolation points XI should be real.')
end

% Process YI
yi = varargin{3};
if ~isreal(yi)
    error([mfilename,'parseInputs:ComplexInterpPts'], ...
        'The interpolation points YI should be real.')
end
if ~isequal(size(xi),size(yi))
    error([mfilename,'parseInputs:YIXINotSameSize'], ...
        'YI must be the same size as XI');
end

% Process ZI
zi = varargin{4};
if ~isreal(zi)
    error([mfilename,'parseInputs:ComplexInterpPts'], ...
        'The interpolation points ZI should be real.')
end
if ~isequal(size(xi),size(zi))
    error([mfilename,'parseInputs:ZIXINotSameSize'], ...
        'ZI must be the same size as XI');
end

% Process EXTRAPVAL
if nargs > 4
    extrapval = varargin{5};
else
    extrapval = [];
end
if isempty(extrapval)
    extrapval = NaN;
end
if ~isscalar(extrapval)
    error([mfilename,':NonScalarExtrapValue'],...
        'EXTRAP option must be a scalar.')
end
```
