## Supplementary Software for "Field dependent deep learning enables high-throughput whole-cell 3D super-resolution imaging": plotCRLBcsplinePSF.m

```
function plotCRLBcsplinePSF(cspline,fit_parameters,ax)
dz=0.1;
rois=size(cspline.coeff{1},1)-3;
N=5000;bg=50;x=rois/2;y=rois/2;z=(1:dz:size(cspline.coeff{1},3))';  
z_range = abs(size(cspline.coeff{1},3)*cspline.dz /2);
v1=ones(length(z),1);
coords=[v1*x , v1*y , v1* N, v1*bg, z];
crlb=CalSplineCRLB(cspline.coeff{1}, rois, coords);
xpx=fit_parameters.pixelSizeX;
ypx=fit_parameters.pixelSizeY;
xe=sqrt(crlb(:,1))*xpx;
ye=sqrt(crlb(:,2))*ypx;
ze=sqrt(crlb(:,5))*cspline.dz;
zp=(z-cspline.z0)*cspline.dz;
indpl=abs(zp)<z_range;  
plot(ax,zp(indpl),xe(indpl),zp(indpl),ye(indpl),zp(indpl),ze(indpl));
ylim(ax,[0 quantile(ze((indpl)),0.95)*1.1]);
xlim(ax,[-z_range z_range]); 
legend(ax,'x','y','z','location','north');
title(ax,['localization precision for N= ',num2str(N), ' ,' 'bg=',num2str(bg)]);
xlabel(ax,'z (nm)');
ylabel(ax,'sqrt(CRLB) in nm');
end
```
