## Supplementary Software for "Field dependent deep learning enables high-throughput whole-cell 3D super-resolution imaging": readbeadimages.m

```
function [imstack, roi, pixelsize,settings3D,isem]=readbeadimages(file,p)
[path,f,ext]=fileparts(file);
indq=strfind(f,'_q');
settings3D=[];
multichannel_4pi=false;
if ~isempty(indq)
    allfiles=dir([path filesep f(1:indq+1)  '*' ext]);
    for k=1:length(allfiles)
        files{k}=[path filesep allfiles(k).name];
        disp(allfiles(k).name(indq:end))
    end
    file=files;
    multichannel_4pi=true;
end
pixelsize=100;
imstack=[];    

if isempty(imstack)
    disp('using simple reader')
    %comment by shiwei 20210527
    %warndlg('using simple reader, this might create problems if only part of the camera chip is used.','using simple reader','replace');
    if multichannel_4pi
        imstack=[];
        for k=1:length(file)
            imstack=horzcat(imstack,readfile_tif(file{k}));
        end
    else
         imstack=readfile_tif(file);
    end
    roi=[0 0 size(imstack,2) size(imstack,1)]; %Keep consistent with the ROI of the original tiff image
end         
%     else
%         imstack=readfile_tif(file);
%         roi=[0 0 size(imstack,1) size(imstack,2)]; %check x,y
%     end
    if multichannel_4pi
        wx=size(imstack,2)/4;wy=size(imstack,1);
        settings3D=struct('y4pi',[0 0 0 0],'x4pi',[0 wx 2*wx 3*wx], 'width4pi',wx,'height4pi',wy,'mirror4pi',[0 0 0 0],'pixelsize_nm',100,'offset',100,'conversion',0.5);
    end
    if isfield(p,'framerange')
        if length(p.framerange)~=2
            fr=p.framerange;
        else
            fr=p.framerange(1):p.framerange(2);
        end
           
        imstack=imstack(:,:,fr);
    end
    isem=false;
    if isfield(p,'emmirror') %do something with the ROI? Or already in image loader?
        switch p.emmirror
            case {0,1} %do nothing
            case 2 %from metadata
%                  if r.metadata.EMon  %modify by shiwei
                 if 1 %modify by shiwei  %ÐÞ¸Ä¾µÏñ
                     isem=true;
                     imstack=imstack(:,end:-1:1,:);
                 end                  %modify by shiwei
            case 3 %mirror
                isem=true;
                imstack=imstack(:,end:-1:1,:);
        end
        %XXXX for Andor take out?
%          if any(roi(1:2)>0) %if roi(1:2)=[0 0] it is likely that roi was not read out and set to default.
%              roi(1)=512-roi(1)-roi(3);
%          end
    end
%     if isfield(p,'roimask')&&~isempty(p.roimask)
%         imstack=imstack.*uint16(p.roimask);
%     end
end
```
