## Supplementary Software for "Field dependent deep learning enables high-throughput whole-cell 3D super-resolution imaging": registerPSF3D_g.m

```
%start add by shiwei 2020/08/14
% function [imout,shiftedstackn,shifto,indgood]=registerPSF3D_g(imin,imint,p,axs,filenumber)
function [imout,shiftedstackn,shifto,indgood,imout_all]=registerPSF3D_g(imin,imint,p,axs,filenumber)
%end add by shiwei 2020/08/14
if nargin<4
    axs={};
end
if nargin<3
    p=[];
end
if nargin<5
    filenumber=[];
end
% perform correlation on:
% p.xrange
% p.yrange
% p.framerange
% can do 2D if length(p.framerange)=1
%cutout small volumes
if ~isfield(p,'xrange')
    p.xrange=1:size(imin,1);
end
if ~isfield(p,'yrange')
    p.yrange=1:size(imin,2);
end
if ~isfield(p,'framerange')
    p.framerange=1:size(imin,3);
end
if ~isfield(p,'shiftxy')
    p.shiftxy=zeros(size(imin,4),2);
end
if ~isfield(p,'alignz')
    p.alignz='cross-correlation';
end
if ~isfield(p,'beadfilterf0')
    p.beadfilterf0=false;
end
if ~isfield(p,'status')
    p.status=[];
end

if ~isfield(p,'sortind')
    p.sortind=1:size(imin,4);
end

if ~isfield(p,'removeoutliers')
    p.removeoutliers=true;
end
if ~isfield(p,'normalize')
    p.normalize=true;
end

numbeads=size(imin,4);
if numbeads==1
    if isempty(imint)
    imout=imin;
    shiftedstackn=imin;
    else
        imout=vertcat(imin, imint);
        shiftedstackn=imout;
    end
    shifto=[0 0 0 ];
    indgood=true;
    return
end
% lx=length(p.xrange);

if isfield(p,'zshiftf0') && ~isempty(p.zshiftf0)
    zshiftf0=p.zshiftf0;
else
    zshiftf0=zeros(numbeads,1);
end

if ~isempty(imint)
    %make big image, shift target image according to shiftxy
    imina=zeros(size(imin,1)*2,size(imin,2),size(imin,3),size(imin,4));
    for k=1:size(imin,4)
        imina(1:size(imin,1),:,:,k)=imin(:,:,:,k);
        imina(size(imin,1)+1:end,:,:,k)=shiftimagexy(imint(:,:,:,k),-p.shiftxy(k,:));
    end
else 
%     imina=imin;
    for k=1:size(imin,4)
        imina(:,:,:,k)=shiftimagexy(imin(:,:,:,k),-p.shiftxy(k,:));
    end
end

numref=max(round(size(imina,4)*.5),min(5,size(imina,4)));
% numref=1;
avim=mean(imina(:,:,:,p.sortind(1:numref)),4,'omitnan');

ph=p;
lcc=ceil((min(13,length(p.yrange))-1)/2);
mp=ceil(((length(p.yrange))-1)/2)+1;
ph.yrange=p.yrange(mp-lcc:mp+lcc);

if ~isempty(imint)
    ph.xrange=[p.xrange(mp-lcc:mp+lcc) p.xrange(mp-lcc:mp+lcc)+size(imin,1)];
else
    ph.xrange=p.xrange(mp-lcc:mp+lcc);
end

%new algorithm to try:
%1. align with all frames
ph.framerange=1:size(avim,3);
[shiftedstack,shift,cc]=aligntoref(avim,imina, zshiftf0,ph);

%calculate good ones, 
shiftedstackn=normalizstack(shiftedstack,p);
indgood=true(1,size(shiftedstackn,4));
if p.removeoutliers
[indgood,res,normamp,co,cc,weightso]=getoverlap(shiftedstackn,shift,ph,indgood);
end
% meanim=nanmean(shiftedstack(:,:,:,indgood),4);meanim(isnan(meanim))=avim(isnan(meanim));   

meanim=mean(shiftedstack(:,:,:,indgood),4,'omitnan');meanim(isnan(meanim))=avim(isnan(meanim));   


%do central correlation using shiftedstack
ph.framerange=p.framerange;
[shiftedstack,shift2,cc]=aligntoref(meanim,shiftedstack, 0*zshiftf0,ph);

shifto=shift2+shift;
% cca(:,1)=cc;
% for k=1:2
% shiftedstackn=normalizstack(shiftedstack,p);
% 
% indgood=true(1,size(shiftedstackn,4));
% [indgood]=getoverlap(shiftedstackn,shift,ph,indgood);
% % sum(indgood)/length(indgood)
% meanim=mean(shiftedstack(:,:,:,indgood),4,'omitnan');
%     meanim(isnan(meanim))=avim(isnan(meanim));   
% %     refim=meanim(xrange,p.yrange,p.framerange);
% [shiftedstack,shift,cc]=aligntoref(meanim,imina, smallim, zshiftf0,ph);
% cca(:,1+k)=cc;
% figure(100);plot(cca(p.sortind,:));drawnow
% % imageslicer(shiftedstack)
% end

% xn=1:size(imina,1);yn=1:size(imina,2);zn=1:size(imina,3);
% [Xq,Yq,Zq]=meshgrid(yn,xn,zn);

% xns=1:size(smallim,1);yns=1:size(smallim,2);zns=1:size(smallim,3);
% [Xqs,Yqs,Zqs]=meshgrid(yns,xns,zns);

% meanim=[];
% % refim=avim(p.xrange,p.yrange,p.framerange);
% refim=avim;

% simin=size(imina);
% shiftedstack=zeros(simin(1),simin(2),simin(3),numbeads)+NaN;
% 
% for k=1:numbeads
%     goodframes=squeeze(sum(sum(smallim(:,:,:,k),1,'omitnan'),2,'omitnan'))>0;
%     if p.alignz
%         [shift(k,:),cc(k)]=get3Dcorrshift(refim(:,:,goodframes),smallim(:,:,goodframes,k));
%     else
%         if any(goodframes)
% %             smallimshiftf0=interp3(smallim(:,:,:,k),Xqs,Yqs,Zqs-double(zshiftf0(k)),'cubic',0);
%             [shift(k,:),cc(k)]=get2Dcorrshift(refim(:,:,goodframes),smallim(:,:,goodframes,k));
% %             [shift(k,:),cc(k)]=get2Dcorrshift(refim(:,:,goodframes),smallimshiftf0);
%         else
%             shift(k,:)=[0 0 0];cc(k)=NaN;
%         end
%     end
% %     imina(1:size(imin,1),:,:,k)=imin(:,:,:,k);
% %     imina(size(imin,1)+1:end,:,:,k)=shiftimagexy(imint(:,:,:,k),-p.shiftxy(k,:));
%     shiftedh=interp3(imina(:,:,:,k),Xq-shift(k,2),Yq-shift(k,1),Zq-shift(k,3)-double(zshiftf0(k)),'cubic',0);
% %     shiftedh=interp3(imin(:,:,:,k),Xq-shift(k,2),Yq-shift(k,1),Zq-shift(k,3)-double(zshiftf0(k)),'cubic',0);
% %     shiftedht=interp3(imin(:,:,:,k),Xq-shift(k,2)-p.shiftxy(k,1),Yq-shift(k,1)-p.shiftxy(k,2),Zq-shift(k,3)-double(zshiftf0(k)),'cubic',0);
% %     
% %     shiftedstack(1:size(imin,1),:,:,k)=shiftedh;
% %     shiftedstack(size(imin,1)+1:end,:,:,k)=shiftedht;
%     shiftedstack(:,:,:,k)=shiftedh;
%     meanim=mean(shiftedstack(:,:,:,1:k),4,'omitnan');
%     meanim(isnan(meanim))=avim(isnan(meanim));
%     
%     refim=meanim(xrange,p.yrange,p.framerange);
% end


if ph.normalize
shiftedstackn=normalizstack(shiftedstack,p);
else
    shiftedstackn=shiftedstack;
end

indgood=true(1,size(shiftedstackn,4));
if p.removeoutliers
[indgood,res,normamp,co,cc,weightso]=getoverlap(shiftedstackn,shift,ph,indgood);
[indgood,res,normamp,co,cc,weightso]=getoverlap(shiftedstackn,shift,ph,indgood,weightso);
[indgood,res,normglobal,co,cc2,weightso]=getoverlap(shiftedstackn,shift,ph,indgood,weightso);

if ph.normalize
shiftedstackn=shiftedstackn/normglobal;
end
end

imout=mean(shiftedstackn(:,:,:,indgood),4,'omitnan');
imout_all = shiftedstackn(:,:,:,indgood); %add by shiwei 2020/08/14
shiftedstackn(1,end,:,~indgood)=max(shiftedstackn(:),[],'omitnan');
shiftedstackn(1,:,1,~indgood)=max(shiftedstackn(:),[],'omitnan');


if length(axs)>0
    col=lines(max(filenumber));
    leg={};
    hold(axs{1},'off')
    for k=1:max(filenumber)
        fh=filenumber==k;
        if any((~indgood)&fh)
            plot(axs{1},(res((~indgood)&fh)),cc2((~indgood)&fh),'x','Color',col(k,:));
            hold(axs{1},'on')
            leg{end+1}=['x' num2str(k) ':' num2str(sum((~indgood)&fh))];
        end
        if any(indgood&fh) 
            plot(axs{1},(res(indgood&fh)),cc2(indgood&fh),'*','Color',col(k,:));
            hold(axs{1},'on')
            leg{end+1}=[num2str(k) ':' num2str(sum((indgood)&fh))];
        end
    end
    legend(leg);
    xlabel(axs{1},'residulas')
    ylabel(axs{1},'cross-correlation value')
    drawnow
end

if length(axs)>1
    imageslicer(vertcat(avim,imout),'Parent',axs{2}.Parent)
end

end

function [shiftedstack,shift,cc]=aligntoref(avim,imina, zshiftf0,p)
xn=1:size(imina,1);yn=1:size(imina,2);zn=1:size(imina,3);


smallim=zeros(length(p.xrange),length(p.yrange),length(p.framerange),size(imina,4));
for k=1:size(imina,4)
    frh=round(p.framerange-zshiftf0(k)); %makes sure, all beads are at same z position
    try
    smallim(:,:,:,k)=imina(p.xrange,p.yrange,frh,k);
    catch err %range out 
    end
end


[Xq,Yq,Zq]=meshgrid(yn,xn,zn);
% meanim=[];
refim=avim(p.xrange,p.yrange,p.framerange);
% refim=avim;

numbeads=size(imina,4);
simin=size(imina);
shiftedstack=zeros(simin(1),simin(2),simin(3),numbeads)+NaN;

for k=1:numbeads
        p.status.String=['calculate shift of individual PSFs: ' num2str(k) ' of ' num2str(numbeads)]; drawnow
    goodframes=squeeze(sum(sum(smallim(:,:,:,k),1,'omitnan'),2,'omitnan'))>0;
    if p.alignz
        [shift(k,:),cc(k)]=get3Dcorrshift(refim(:,:,goodframes),smallim(:,:,goodframes,k));
    else
        if any(goodframes)
%             smallimshiftf0=interp3(smallim(:,:,:,k),Xqs,Yqs,Zqs-double(zshiftf0(k)),'cubic',0);
            [shift(k,:),cc(k)]=get2Dcorrshift(refim(:,:,goodframes),smallim(:,:,goodframes,k));
%             [shift(k,:),cc(k)]=get2Dcorrshift(refim(:,:,goodframes),smallimshiftf0);
        else
            shift(k,:)=[0 0 0];cc(k)=NaN;
        end
    end
%     imina(1:size(imin,1),:,:,k)=imin(:,:,:,k);
%     imina(size(imin,1)+1:end,:,:,k)=shiftimagexy(imint(:,:,:,k),-p.shiftxy(k,:));
    
shiftedh=interp3(imina(:,:,:,k),Xq-shift(k,2),Yq-shift(k,1),Zq-shift(k,3)-double(zshiftf0(k)),'cubic',0);
%  shiftedh=interp3(imina(:,:,:,k),Xq-shift(k,2),Yq-shift(k,1),Zq+shift(k,3)-double(zshiftf0(k)),'cubic',0);

    
    %     shiftedh=interp3(imin(:,:,:,k),Xq-shift(k,2),Yq-shift(k,1),Zq-shift(k,3)-double(zshiftf0(k)),'cubic',0);
%     shiftedht=interp3(imin(:,:,:,k),Xq-shift(k,2)-p.shiftxy(k,1),Yq-shift(k,1)-p.shiftxy(k,2),Zq-shift(k,3)-double(zshiftf0(k)),'cubic',0);
%     
%     shiftedstack(1:size(imin,1),:,:,k)=shiftedh;
%     shiftedstack(size(imin,1)+1:end,:,:,k)=shiftedht;
    shiftedstack(:,:,:,k)=shiftedh;
%     meanim=mean(shiftedstack(:,:,:,1:k),4,'omitnan');
% %     meanim(isnan(meanim))=refim(isnan(r));
%     
%     refim=meanim(p.xrange,p.yrange,p.framerange);
end
end

function [indgood,res,normamp,co,cc,weightso]=getoverlap(shiftedstackn,shift,p,indgood,weights)

if nargin<5
    weights=ones(size(shiftedstackn,4),1);
end

refimn=0;
for k=size(shiftedstackn,4):-1:1
    imh=shiftedstackn(p.xrange,p.yrange,p.framerange,k);
    if indgood(k) && ~all(isnan(imh(:)))
        refimn=weights(k)*shiftedstackn(p.xrange,p.yrange,p.framerange,k)/sum(weights(indgood))+refimn;
    end
end

% refimn=mean(shiftedstackn(p.xrange,p.yrange,p.framerange,indgood),4,'omitnan');
for k=size(shiftedstackn,4):-1:1
    imh=shiftedstackn(p.xrange,p.yrange,p.framerange,k);
    badind=isnan(imh)|isnan(refimn);
    cc(k)=sum(refimn(~badind).*imh(~badind))/(sum(refimn(~badind))*sum(imh(~badind)))*sum(~badind(:));  
end

normamp=max(refimn(:),[],'omitnan');
shiftedstackn=shiftedstackn/normamp;
refimn=refimn/normamp;
for k=size(shiftedstackn,4):-1:1
     sim=shiftedstackn(p.xrange(2:end-1),p.yrange(2:end-1),p.framerange,k);
     dv=(refimn(2:end-1,2:end-1,:)-sim).^2;
    res(k)=sqrt(mean(dv(:),'omitnan'));
end
rescc=res./cc;
rescc(abs(shift(:,1))>3|abs(shift(:,2))>3)=NaN;
indtest=indgood&(cc>0);
[a,b]=robustMean(rescc(indtest));
if isnan(b)
    a=mean(rescc,'omitnan');b=std(rescc,'omitnan');
end

%hard limiter
%start add by shiwei 2020/09/10
% co=a+2.*b;
co=a+5.*b;      % shiwei ÉèÖÃÂË¹ýÌõ¼þ£¬Ôö¼ÓsigmaµÄÖµ¿ÉÒÔ¼õÉÙÂËµôµÄµã
%end add by shiwei 2020/09/10

% weightso=weights;
weightso=1./rescc;
indgood=indgood&(rescc<=co);
% norm=sum(1./rescc);
% for k=size(shiftedstackn,4):-1:1
%     shiftedstackn2(:,:,:,k)=shiftedstackn(:,:,:,k)
%      sim=shiftedstackn(p.xrange(2:end-1),p.yrange(2:end-1),p.framerange,k); 
% end

end

function out=normalizstack(in,p)
sin=size(in);
  midp=round((length(p.xrange)+1)/2);
    xr=p.xrange(midp-3:midp+3);yr=p.yrange(midp-3:midp+3);
if p.beadfilterf0 
    out=0*in+NaN;
  
    for k=1:sin(4)
        imh=in(xr,yr,p.framerange,k);
        nm=mean(imh(:),'omitnan');
        if nm>0
        out(:,:,:,k)=in(:,:,:,k)/nm;
        end
    end
else %use fitting
    inh=in;
    out=0*in+NaN;
    for iter=1:4
        meanim=mean(inh,4,'omitnan');
        for k=1:sin(4)
            imh=inh(:,:,:,k);
            if all(isnan(imh))
                continue
            end
            ims=imh(xr,yr,p.framerange);
            meanims=meanim(xr,yr,p.framerange);
            isn=isnan(ims)|isnan(meanims);
            intcutoff=meanims>myquantile(meanims(:),0.75);
            indg=~isn&intcutoff;
            ratio=ims(indg)./meanims(indg);
            
            factor=median(ratio(:),'omitnan');
            if factor>0
                out(:,:,:,k)=imh/factor;
            end
        end
        inh=out;
    end

end

end
```
