## Supplementary Software for "Field dependent deep learning enables high-throughput whole-cell 3D super-resolution imaging": shiftimagexy.m

```
function imout= shiftimagexy(imin,dx)
xn=1:size(imin,1);yn=1:size(imin,2);
[Xq,Yq]=meshgrid(yn,xn);

imout=0*imin;
for k=1:size(imin,3)
 imout(:,:,k)=interp2(imin(:,:,k),Xq-dx(1),Yq-dx(2),'cubic',0);
end
```
