## Supplementary Software for "Field dependent deep learning enables high-throughput whole-cell 3D super-resolution imaging": symExpFilt.m

```
function Y = symExpFilt(X,XLen,C0,Zi,K0,KVec)
% symExpFilt: symmetric exponential filter
% usage: Y = symExpFilt(X,XLen,C0,Zi,K0,KVec);
%
% arguments:
%   X - vector of input data
%   XLen - length(X)
%   C0 - scaling constant
%   Zi - pole of direct BSpline filter
%   K0 - parameter for computing initial condition
%   KVec - vector of indices for computing initial condition (reflects from
%       boundaries if necessary)
%
%   Y - vector of output data
%
% Note: This function implements the recursive filter in Equations (2.5)
% and (2.6) of M. Unser, A. Aldroubi, and M. Eden, "B-Spline Signal
% Processing: Part II - Efficient Design and Applications," IEEE Trans.
% Signal Processing, 41(2):834-848, February 1993.
%

% author: Nathan D. Cahill
%
% date: 18 April 2008

% initialize Y
Y = zeros(size(X));

% compute first element of Y
for k=1:K0
    Y(1) = Y(1) + (Zi^(k-1))*X(KVec(k));
end

% filter in forward direction
for k=2:XLen
    Y(k) = X(k) + Zi*Y(k-1);
end

% update last element of Y
Y(XLen) = (2*Y(XLen) - X(XLen))*C0;

% filter in reverse direction
for k=(XLen-1):-1:1
    Y(k) = (Y(k+1) - Y(k))*Zi;
end
```
