## Supplementary Software for "Field dependent deep learning enables high-throughput whole-cell 3D super-resolution imaging": testfit_spline.m

```
function posbeads=testfit_spline(teststack,coeff,shiftxy, p,linepar,ax)
if nargin<4
    linepar={};
elseif ~iscell(linepar)
    linepar={linepar};
end
% roifit = (size(teststack,1)+1)/2; %modify by shiwei 2021/12/23
roifit =size(coeff{1},1)+1;%modify by shiwei 2021/12/23
% roifit=51;   %mark tetrapod 17   61
d=round((size(teststack,1)-roifit)/2);
range=d+1:d+roifit;

numstack=size(teststack,4);
t=tic;
% f=figure(989);ax2=gca;hold off
% dx1=[];
% dx2=[];dy1=[];
% dy2=[];

% f=figure(134);ax2=gca;hold(ax2,'off')
    for k=1:size(teststack,4)
        if toc(t)>1
            p.status.String=['fitting bead stacks: ' num2str(k/numstack,'%1.2f')];drawnow
            t=tic;
        end
%         if isfield(p,'PSF2D') && p.PSF2D
%             zst=min(round(size(teststack,3)*0.4),500/p.dz)*[-1 1];
%         else
            zst=0;
%         end
        fitmode=5;
        fitstack=single(squeeze(teststack(range,range,:,k,:)));
        coeffh(:,:,:,:,1)=single(coeff{1});
        iterations=150;
        
%         if p.isglobalfit
%              shared=[1 1 1 0 0]';
%             nfits=size(fitstack,3);
%             npar=5;
%             dT=zeros(npar,2,nfits);
%             dT(1,2,:)=shiftxy(k,2);
%             dT(2,2,:)=shiftxy(k,1);           
%             coeffh(:,:,:,:,2)=single(coeff{2});
%             sharedA = repmat(shared,[1 size(fitstack,3)]);
% %             if fitmode==6
% %                 zst=500/p.dz*[-1 1];
% %             else
% %                 zst=0;
% %             end
%              [P,CRLB, LL] =mleFit_LM_global(fitstack,int32(sharedA),iterations,coeffh,single(dT),1,zst);
%             zind=3;
%             photind=4;
%         else
            if max(fitstack(:))<10
                fitstack=fitstack*1000+1;
            end
            [P,CRLB, LL] =mleFit_LM(fitstack,fitmode,iterations,coeffh,0,1,zst);
            zind=5;
            photind=3;
%         end
        

        
%         [PM,CRLBM, LLM,update, error] =  kernel_MLEfit_Spline_LM_multichannel_finalized(fitstack,coeffh, shared,dT,50);
        
%         try
%         [P,CRLB, LL] =GPUmleFit_LM_MultiChannel(fitstack,int32(sharedA),iterations,coeffh,single(dT));
%         catch err
%               [P,CRLB, LL] =CPUmleFit_LM_MultiChannel(fitstack,int32(sharedA),iterations,coeffh,single(dT));
%         end
        
        
%         [P,CRLB, LL,residuals] =CPUmleFit_LM_MultiChannel_R(fitstack,int32(sharedA),iterations,coeffh,single(dT));
 
        z=(1:size(P,1))'-1;

        znm=(P(:,zind)-p.z0)*p.dz;
        plot(ax,z,znm,linepar{:});
        hold(ax,'on');
        xlabel(ax,'frame');
        ylabel(ax,'zfit (nm)');
        zs(:,k)=P(:,zind);
        posbeads.z(:,k)=P(:,zind);
        dn=ceil((size(fitstack,1)-1)/2);
        posbeads.x(:,k)=(P(:,2)-dn);
        posbeads.y(:,k)=(P(:,1)-dn);
        posbeads.frame(:,k)=1:length(z);
        posbeads.LL(:,k)=LL;
        posbeads.phot(:,k)=P(:,photind);
        posbeads.znm(:,k)=znm;  %add by shiwei 20210623
% 
% plot(ax2,P(:,1),P(:,2),'.')
% hold(ax2,'on')


        
%         if 0% imageslicer to test
% %             coord=P1(:,[1 2 5 3 4]);
%             coord=P(:,[1 2 3 4 6]);
%             coord2=coord;
%             coord2(:,1)=coord2(:,1)+squeeze(dT(1,2,:));
%             coord2(:,2)=coord2(:,2)+squeeze(dT(2,2,:));
%             img1=renderPSF(coeff{1},coord,size(fitstack,1));
%             img2=renderPSF(coeff{2},coord2,size(fitstack,1));
%             imall=[fitstack(:,:,:,1),img1; fitstack(:,:,:,2),img2];
%             res=[fitstack(:,:,:,1)-img1, 0*img1;fitstack(:,:,:,2)-img2,0*img2];
%             ims(:,:,:,1)=imall;ims(:,:,:,2)=res;
%             f=figure(105);
%             imageslicer(ims,'Parent',f);
%         end
        
    end
    

end
```
