## Supplementary Software for "Field dependent deep learning enables high-throughput whole-cell 3D super-resolution imaging": copyfields.m

```
function [destination,missing]=copyfields(destination,source,fields)
missing={};
if nargin==2 %copy all
    if ~isempty(source)
        if isa(destination,'handle')
            fn=intersect(properties(destination),fieldnames(source));
        else
            fn=fieldnames(source);
        end
        for k=1:length(fn)
            destination.(fn{k})=source.(fn{k});
        end
    end
elseif nargin>2
    if ~iscell(fields)
        fields={fields};
    end
%     fn=intersect(fields,fnsource);
    if ~isempty(source)
    indm=1;
    
    fnsource=fieldnames(source);
%     fnc=cellfun(@makehash,fnsource);
%     tocopy=intersect(fnsource,fields);
%     missing=setdiff(fields,fnsource);

%     for k=1:length(tocopy)
%         destination.(tocopy{k})=source.(tocopy{k});
%     end
%     fn=fields;
    for k=1:length(fields)
        if any(strcmp(fnsource,fields{k}))
            destination.(fields{k})=source.(fields{k});
        else
            missing{indm}=fields{k};
            indm=indm+1;
        end
    end
    else
        missing=fields;
    end
end

function out=makehash(str)
    out=sum(char(str));
% out=1;
```
