## Supplementary Software for "Field dependent deep learning enables high-throughput whole-cell 3D super-resolution imaging": robustMean.m

```
function [finalMean, stdSample, inlierIdx, outlierIdx] = robustMean(data,dim,k,fit)
%ROBUSTMEAN calculates mean and standard deviation discarding outliers
%
% SYNOPSIS [finalMean, stdSample, inlierIdx, outlierIdx] = robustMean(data,dim,k,fit)
%
% INPUT    data : input data
%          dim  : (opt) dimension along which the mean is taken {1}
%          k    : (opt) #of sigmas at which to place cut-off {3}
%          fit  : (opt) whether or not to use fitting to robustly estimate
%                  the mean from the data that includes outliers.
%                  0 (default): mean is approximated by median(data)
%                  1 : mean is approximated by
%                      fminsearch(@(x)(median(abs(data-x))),median(data))
%                      This option is only available for scalar data
%
%
% OUTPUT   finalMean : robust mean
%          stdSample : std of the data (divide by sqrt(n) to get std of the
%                      mean)
%          inlierIdx : index into data with the inliers
%          outlierIdx: index into data with the outliers
%
% REMARKS  NaN or Inf will be counted as neither in- nor outlier
%          The code is based on (linear)LeastMedianSquares. It could be changed to
%          include weights
%
% c: jonas, 04/04
%%%%%%%%%%%%%%%%%%%%%%%%%%%%%%%%%%%%%%%%%%%%%%%%%%%%%%%%%%%%%%%%%%%%%%%%%%%%%


% test input

if isempty(data)
    error('Please supply non-empty data to robustMean')
end
if nargin<2 || isempty(dim)
    % make sure that the dimensinon is correct if there's a vector
    if any(size(data)==1) && ndims(data)==2
        dim = find(size(data)>1);
    else
        dim = 1;
    end
end
if nargin < 3 || isempty(k)
    k = 3;
end
if nargin < 5 || isempty(fit)
    fit = 0;
end
if fit == 1
    % check for vector
    if sum(size(data)>1)>1
        error('fitting is currently only supported for 1D data')
    end
end
if sum(isfinite(data(:))) < 4
    warning('ROBUSTMEAN:INSUFFICIENTDATA',...
        'Less than 4 data points!')
    if isempty(dim)
        dim=1;
    end
    finalMean = nanmean(data,dim);
%     stdSample = NaN(size(finalMean));-
    stdSample=nanstd(data,[],dim);
    inlierIdx = find(isfinite(data));
    outlierIdx = [];
    return
end


%========================
% LEAST MEDIAN SQUARES
%========================

% define magic numbers:
%k=3; %cut-off is roughly at 3 sigma, see Danuser, 1992 or Rousseeuw & Leroy, 1987
magicNumber2=1.4826^2; %see same publications

% remember data size and reduced dataSize
dataSize = size(data);
reducedDataSize = dataSize;
reducedDataSize(dim) = 1;
% need this for later repmats
blowUpDataSize = dataSize./reducedDataSize;
% count how many relevant dimensions we have besides dim
realDimensions = length(find(dataSize>1));

% calc median - reduce dimension dim to length 1
if fit
    % minimize the median deviation from the mean
    medianData = fminsearch(@(x)(median(abs(data-x))),median(data));
else
    medianData = nanmedian(data,dim);
end

% calculate statistics
res2 = (data-repmat(medianData,blowUpDataSize)).^2;
medRes2 = max(nanmedian(res2,dim),eps);

%testvalue to calculate weights
testValue=res2./repmat(magicNumber2*medRes2,blowUpDataSize);

if realDimensions == 1;
    %goodRows: weight 1, badRows: weight 0
    inlierIdx=find(testValue<=k^2);
    outlierIdx = find(testValue>k^2);
    
    % calculate std of the sample;
    if nargout > 1
        nInlier = length(inlierIdx);
        if nInlier > 4
            stdSample=sqrt(sum(res2(inlierIdx))/(nInlier-4));
        else
            stdSample = NaN;
        end
    end
    
    %====END LMS=========
    
    %======
    % MEAN
    %======
    
    finalMean = mean(data(inlierIdx));
    
else
    
    %goodRows: weight 1, badRows: weight 0
    inlierIdx=find(testValue<=k^2);
    outlierIdx=find(testValue > k^2);
    
    % mask outliers
    res2(outlierIdx) = NaN;
    % count inliers
    nInliers = sum(~isnan(res2),dim);
    
    % calculate std of the sample;
    if nargout > 1
        % put NaN wherever there are not enough data points to calculate a
        % standard deviation
        goodIdx = sum(isfinite(res2),dim) > 4;
        stdSample = NaN(size(goodIdx));
        stdSample(goodIdx)=sqrt(nansum(res2(goodIdx),dim)./(nInliers(goodIdx)-4));
    end
    
    %====END LMS=========
    
    %======
    % MEAN
    %======
    
    data(outlierIdx) = NaN;
    finalMean = nanmean(data,dim);
end
```
