## Supplementary Software for "Field dependent deep learning enables high-throughput whole-cell 3D super-resolution imaging": uigetdirs.m

```
function dirs = uigetdirs(startpath, titleString)
    % uigetdirs - Dialog for selecting multiple folders
    %
    % USAGE:
    %   dirs = uigetdirs(startpath, title)
    %
    % INPUTS:
    %   startpath:  String, path to a folder to use as the default search
    %               path. If this is an empty string, not a valid path, or
    %               not provided then it defaults to the current directory
    %
    %   title:      String, customized string to display as the figure
    %               title.
    %
    % OUTPUTS:
    %   dirs:       Cell Array, A cell array containing all of the selected
    %               directories. If no directory was selected because the
    %               dialog was cancelled or closed, then this value will be
    %               just an empty cell array

    % Copyright (c) 2016, Jonathan Suever
    % All rights reserved.
    %
    % Redistribution and use in source and binary forms, with or without
    % modification, are permitted provided that the following conditions
    % are met:
    %
    % 1. Redistributions of source code must retain the above
    %    copyright notice, this list of conditions and the following
    %    disclaimer.
    %
    % 2. Redistributions in binary form must reproduce the above copyright
    %    notice, this list of conditions and the following disclaimer in
    %    the documentation and/or other materials provided with the
    %    distribution.
    %
    % 3. Neither the name of the copyright holder nor the names of its
    %    contributors may be used to endorse or promote products derived
    %    from this software without specific prior written permission.
    %
    % THIS SOFTWARE IS PROVIDED BY THE COPYRIGHT HOLDERS AND CONTRIBUTORS
    % "AS IS" AND ANY EXPRESS OR IMPLIED WARRANTIES, INCLUDING, BUT NOT
    % LIMITED TO, THE IMPLIED WARRANTIES OF MERCHANTABILITY AND FITNESS FOR
    % A PARTICULAR PURPOSE ARE DISCLAIMED. IN NO EVENT SHALL THE COPYRIGHT
    % HOLDER OR CONTRIBUTORS BE LIABLE FOR ANY DIRECT, INDIRECT,
    % INCIDENTAL, SPECIAL, EXEMPLARY, OR CONSEQUENTIAL DAMAGES (INCLUDING,
    % BUT NOT LIMITED TO, PROCUREMENT OF SUBSTITUTE GOODS OR SERVICES; LOSS
    % OF USE, DATA, OR PROFITS; OR BUSINESS INTERRUPTION) HOWEVER CAUSED
    % AND ON ANY THEORY OF LIABILITY, WHETHER IN CONTRACT, STRICT
    % LIABILITY, OR TORT (INCLUDING NEGLIGENCE OR OTHERWISE) ARISING IN ANY
    % WAY OUT OF THE USE OF THIS SOFTWARE, EVEN IF ADVISED OF THE
    % POSSIBILITY OF SUCH DAMAGE.

    if exist('startpath', 'var') && exist(startpath, 'dir')
        startpath = java.io.File(startpath);
    else
        startpath = java.io.File(pwd);
    end

    if ~exist('titleString', 'var')
        titleString = 'Select a folder';
    elseif ~ischar(titleString)
        error(sprintf('%s:InvalidInput', mfilename), ...
            'Title must be a valid string')
    end

    fig = figure('Toolbar',             'none', ...
                 'HandleVisibility',    'off', ...
                 'windowstyle',         'modal', ...
                 'menubar',             'none', ...
                 'NumberTitle',         'off', ...
                 'Visible',             'off', ...
                 'Name',                titleString);

    [jfile, hfile] = javacomponent('javax.swing.JFileChooser', [], fig, ...
                    {'ActionPerformed', @(s,e)selectionCallback(fig,s,e)});

    % Adjust the properties of the file chooser
    jfile.setCurrentDirectory(startpath);
    jfile.setMultiSelectionEnabled(true);
    jfile.setFileSelectionMode(javax.swing.JFileChooser.DIRECTORIES_ONLY);

    % Adjust the position so that it fills the entire figure
    set(hfile, 'units',     'normalized', ...
               'position',  [0 0 1 1]);

    movegui(fig, 'center');

    % Make figure visible now that everything is ready
    set(fig, 'visible', 'on')
    set(fig, 'deletefcn', @(s,e)cleanup(hfile))

    % Wait until the callback signals a change
    waitfor(fig, 'userdata');
    if ishghandle(fig)
        dirs = get(fig, 'Userdata');
        delete(fig);
    else
        dirs = {};
    end
end

function cleanup(hfile)
    pause(0.25)
    drawnow
    delete(hfile)
end

function selectionCallback(fig, src, evnt)
    command = char(evnt.getActionCommand);
    if strcmpi(command, src.APPROVE_SELECTION)
        dirs = src.getSelectedFiles();
        dirs = arrayfun(@char, dirs, 'uni', 0);
    elseif strcmpi(command, src.CANCEL_SELECTION)
        dirs = {};
    end

    % Update userdata so caller knows that we are done here
    set(fig, 'userdata', dirs);
end
```
