## Supplementary Software for "Field dependent deep learning enables high-throughput whole-cell 3D super-resolution imaging": get3Dcorrshift.m

```
function [shift,CC]=get3Dcorrshift(refim,targetim,maxmethod)
if nargin<3
    maxmethod='interp';
end
if sum(refim(:))==0||sum(targetim(:))==0
    shift=[0 0 0];
    CC=0;
    return;
end
sim=size(targetim);
if length(sim)<3||sim(3)==1 %2D
    [shift,CC]=get2Dcorrshift(refim,targetim,maxmethod);
% G = (fftshift(real(ifft2(fft2(refim).*conj(fft2(targetim))))))/...
% 		( (mean(refim(:))*mean(targetim(:))) * size(refim,1)*size(refim,2) ) ...
% 		- 1;
else
    nanval=-1;
    refimnn=refim; refimnn(isnan(refimnn))=nanval;
    tarimnn=targetim; tarimnn(isnan(tarimnn))=nanval; 
    
    refimhd=interp3(refimnn,2,'cubic');
    targetimhd=interp3(tarimnn,2,'cubic');
    refimhd=refimhd/nanmean(refimhd(:));
    targetimhd=targetimhd/nanmean(targetimhd(:));
%         refimhd=(refim);
%     targetimhd=(targetim;);
%     targetimhd=refimhd;

    n=size(refimhd)*1;
%     reffft=fft(fft2(refimhd,n(1),n(2)),n(3),3);
%     targetfft=fft(fft2(targetimhd,n(1),n(2)),n(3),3);
    reffft=fft(fft2(refimhd),[],3);
    targetfft=fft(fft2(targetimhd),[],3);
    
    Gfft=reffft.*conj(targetfft);
    Gs = fftshift(real(ifft(ifft2(Gfft),[],3)));
    Gs(Gs<0)=NaN;
    nisn=~isnan(targetimhd);
    G=Gs/(sum(nisn(:)))-1;
    
%     px=sum(sum(G,2),3);
%      maxmethod='interp';
    
     switch maxmethod
         case 'fit'
              maxind=[getmaxFit(sum(sum(G,2),3),3),getmaxFit(sum(sum(G,3),1),3),getmaxFit(sum(sum(G,1),2),3)];
         otherwise
            [CC,ind]=max(G(:));
            if ind>1
            [x,y,z]=ind2sub(size(G),ind);
             maxind=getmaxInterp(G,[x,y,z],.05,1);
            else
                maxind=(size(refim)*2+1);
            end
     end
     shift=maxind/4-size(refim)/2+1/4;
     
%      shift=maxind/2-size(refim)/2;
%      shift=maxind-size(refim)/2;
%      shift=maxind-2*size(refim)+1;
    %maximum: quadratic fit in each dimension
    %compare with interp3 in 1 pixel 100x scaling and maximum find
end

end

function maxind=getmaxInterp(V,pos,dx,w)

%     dx=.01;w=1;
    [Xq,Yq,Zq]=meshgrid(pos(2)-w:dx:pos(2)+w,pos(1)-w:dx:pos(1)+w,pos(3)-w:dx:pos(3)+w);
%     Gsmall=V(x-2:x+2,y-2:y+2,z-2:z+2);
    Vhd=interp3(V,Xq,Yq,Zq,'cubic');
        [~,ind]=max(Vhd(:));
    [x,y,z]=ind2sub(size(Vhd),ind);
    maxind=[x*dx+pos(1)-w-dx, y*dx+pos(2)-w-dx ,z*dx+pos(3)-w-dx];
end

function maxind=getmaxFit(in,window)
if nargin<2
    window=5;
end
sw=round((window-1)/2);
[~,ind]=max(in);
range=(max(1,ind-sw):min(ind+sw,length(in)))';
warning('off')
ps=squeeze(in(range));
[psx,S] = polyfit(range,ps(:),2);
maxind=-psx(2)/2/psx(1);
warning('on')
end
```
