## Supplementary Software for "Field dependent deep learning enables high-throughput whole-cell 3D super-resolution imaging": getCameraCalibration.m

```
function [par,cam,state,error]=getCameraCalibration(imloader,l,silent,file)
error='';
if nargin<3||isempty(silent)
    silent=false;
end
if nargin<4||isempty(file)
    disp('camera calibration file not passed')
     adsf
    file='settings/cameras.mat';
end
% replacefields={'cam_pixelsize_um','pixsize'};
replacefields={};
par=[];
cam=[];
state=[];
argin{1}=imloader;argin{3}=silent;
if nargin<2||isempty(l)
    argin{2}=[];
    
    if ~exist(file,'file')
%         l=[];
        [par,cam,state]=askforcameramanager(imloader,'camera calibration file settings/camera.mat not found. ',silent,argin);
        return
%         [par,cam]=getCameraCalibration(imloader,[],silent);
    end
    l=load(file);
    argin{2}=l;
else
    argin{2}=l;
end
val=[];
for cam=1:length(l.cameras)
    valh=imloader.gettag(l.cameras(cam).ID.tag);
    if ~isempty(valh)&&strcmp(valh,l.cameras(cam).ID.value)        
        val=valh;
        break
    end
end
if isempty(val)
    cam=[];
    
    if ~silent
        answ=questdlg('Camera not recognized. Use default camera?');
    end
    if silent||strcmp(answ,'Yes')  
        camnames=getFieldAsVector(l.cameras,'ID','name');
        cam=find(strcmp(camnames,'Default'));
        state=1;
        disp('Camera not recognized. Use default');
        error='Camera not recognized. Use default camera. Make sure this is correct, e.g. by checking the parameters in the set Cam Parameters dialog.';
        if isempty(cam)&~silent
            errordlg('create Default camera with Camera Manager')
        end    
    else    
        [par,cam,state]=askforcameramanager(imloader,'Camera not recognized. ',silent,argin);
        return;
    end
end
partable=l.cameras(cam).par;
s=size(partable);
if ~isempty(strcmp(partable(:,2),'state dependent'))
    %get state
%     found=false;
    for k=length(l.cameras(cam).state):-1:1
        deftab=l.cameras(cam).state(k).defpar;
        found(k)=true;
        for k2=1:size(deftab,1)
            if strcmp('select',deftab(k2,1) )|| isempty(deftab{k2,2})
                continue
            end
            valh=imloader.gettag(deftab{k2,1});
            if ~strcmp(valh,deftab{k2,2})
                found(k)=false;
                break
            end
                
        end
    end
    state=find(found,1,'first');  
    if isempty(state)
%         askforcameramanager(imloader,'State of the camera could not be determined. Please use the CameraManager to define proper state. Create new state with Camera Manager now?',silent)
            [par,cam,state]=askforcameramanager(imloader,'State of the camera could not be determined. Please use the CameraManager to define proper state.',silent,argin);
            if ~isempty(par)
                return;
            end
    end
end
for k=1:s(1)
    switch partable{k,2}
        case 'fix'
            X=partable{k,3};
        case 'state dependent'
            if isempty(state)
                X=[];
            else
                X=l.cameras(cam).state(state).par{k,2}; 
            end
        case 'metadata'
            X=imloader.gettag(partable{k,4});

    end
    if ~isempty(partable{k,6})&&ischar(X)
        X=eval(partable{k,6});
    end
    par.(partable{k,1})=X;
    
end
if isempty(par.Width)
    w=[str2double(imloader.gettag('Width')),imloader.gettag('Width info')];
    h=[str2double(imloader.gettag('Height')),imloader.gettag('Height info')];
    par.Width=max(w);
    par.Height=max(h);
end

if isempty(par.roi)
    par.roi=[0 0 par.Width par.Height];
end
for k=1:size(replacefields,1)
    par.(replacefields{k,2})=par.(replacefields{k,1});
end
%number Of Frames needed, clean up
if isempty(par.numberOfFrames)||par.numberOfFrames==0
    f1=str2double(imloader.gettag('frames direct'));
    f2=imloader.gettag('numberOfFrames info');
    f3=str2double(imloader.gettag('Frames'));
    par.numberOfFrames=max([f1 f2 f3]);
end

if length(par.cam_pixelsize_um)==1
    par.cam_pixelsize_um(2)=par.cam_pixelsize_um(1);
end
if isfield(par,'roimode')
    par.roi=roiconverter(par.roi,par.roimode);
end

function [paro,camo,stateo]=askforcameramanager(imloader,message,silent,argin)
paro=par;camo=cam;stateo=state;
if silent
    disp(message)
    return
end
answ=warndlg(message,'Open Camera Manager now?');
if strcmp(answ,'Yes')
    disp('please open in camera manager');
    
%     camm=CameraManager;
%     camm.imloader=imloader;
%     camm.loadimages;
%     waitfor(camm.handle)
%     argin{2}=[];
%     [paro,camo,stateo]=getCameraCalibration(argin{:});
end
end
end
```
