## Supplementary Software for "Field dependent deep learning enables high-throughput whole-cell 3D super-resolution imaging": imageloaderAll.m

```
function [io]=imageloaderAll(varargin)
% imageloaderAll selects the right image loader based on the presence of
% metadata.
%file, metadata, P
file=varargin{1};
if iscell(file) %multiple channales
    file=file{1};
end
   [path,~,ext]=fileparts(file);
%    info=imfinfo(file);Tiff
%    switch ext
%        case '.tif'
%            if exist([path filesep 'metadata.txt'],'file')
% %                imloader=@imageloaderMM;s
%                if countfiles(file)>1 && ~(any(strfind(file,'MMStack'))||any(strfind(file,'.ome.')))
%                    imloader=@imageloaderMMsingle;
%                else
%                    imloader=@imageloaderMM;
%                end
%            elseif ~isempty(dir([path filesep '*metadata.txt']))
%                 imloader=@imageloaderMM;
%            elseif any(strfind(file,'MMStack'))
%                 imloader=@imageloaderMM;
%            elseif filesize(file)>4e9&&filesize(file)<4.5e9
%                imloader=@imageloaderMM;
%            elseif countfiles(file)>1000
%                imloader=@imageloaderMMsingle;
%            else
%                imloader=@imageloaderOME;
% %                imloader=@imageloaderMM;
%            end
%        case '' %directory
%            fns=dir([file filesep '*.tif']);
%            varargin{1}=[file filesep fns(1).name];
%            imloader=@imageloaderMMsingle;
%        case '.dcimg'
%            imloader=@imageloaderDCIMG;
%        otherwise
%            imloader=@imageloaderOME;
%    end    
imloader=@imageloaderOME;
   try
%        [io]=imloader(varargin{:});
       [io]=imloader(varargin);  %modify by shiwei
   catch err
       err
       disp('simple tiff loader loader')
       imloader=@imageloaderTifSimple;
       io=imloader(varargin{:});  
   end
end

function f=filesize(file)
d=dir(file);
f=d.bytes;
end
function numf=countfiles(file)
files=myfastdir(fileparts(file),'*.tif');
sstr=regexprep(files{1},'[0-9]*','[0-9]*');
isfile=regexp(files,sstr);
numf=sum(cell2mat(isfile));

end

%distinguish:
    %MM Tif single (with Metadata)
    %MM Tif stack (with Metadata)
    %Tif single wihtout metadata
    %Tif stack without metadata
    %any ome compatible file
```
