## Supplementary Software for "Field dependent deep learning enables high-throughput whole-cell 3D super-resolution imaging": mytiffreader.m

```
classdef mytiffreader<handle
    properties
        reader
        info
    end
    methods
        function obj=mytiffreader(varargin)
            warning('off','MATLAB:imagesci:tiffmexutils:libtiffWarning')
            obj.reader=Tiff(varargin{1},'r');
            obj.info.name=varargin{1};
            obj.getinfo;
        end

        function imagestack=read(obj,frames)
            obj.reader.setDirectory(frames(1));
            img=obj.reader.read;
            imagestack=zeros(obj.info.Width,obj.info.Height,length(frames),'like',img);
            imagestack(:,:,1)=img;
            
            for k=2:length(frames)
                obj.reader.setDirectory(frames(k));
                img=obj.reader.read;
                imagestack(:,:,k)=img;
            end
        end
        function imagestack=readall(obj)
             obj.reader.setDirectory(1);
            img=obj.reader.read;
            imagestack=zeros(obj.info.Width,obj.info.Height,obj.info.numberOfFrames,'like',img);
            imagestack(:,:,1)=img;
            
            for k=2:obj.info.numberOfFrames
                obj.reader.nextDirectory;
                 img=obj.reader.read;
                imagestack(:,:,k)=img;
                
            end
        end
        function getinfo(obj)
            warning('off','MATLAB:imagesci:tiffmexutils:libtiffWarning')
            while ~obj.reader.lastDirectory
                obj.reader.nextDirectory;
            end
            obj.info.numberOfFrames=obj.reader.currentDirectory;
            obj.reader.setDirectory(1);
            img=obj.reader.read;
            obj.info.Width=size(img,1);
            obj.info.Height=size(img,2);
            
            %start add by shiwei 2021/06/03
            obj.info.roi=[0 0 obj.info.Width obj.info.Height];
%             obj.info.roi=getRoiTif(obj.info.name);
%             if all(obj.info.roi(1:2)==[0 0])
%                 obj.info.roi=[0 0 obj.info.Width obj.info.Height];
%             end
             %end add by shiwei 2021/06/03
            
        end
        function close(obj)
            obj.reader.close;
            warning('on','MATLAB:imagesci:tiffmexutils:libtiffWarning')
        end
         
    end
end
```
