## Supplementary Software for "Field dependent deep learning enables high-throughput whole-cell 3D super-resolution imaging": maximumfindcall.m

```
%NMS 2D block after  Neubeck and Gool, Algorithm 4

%reprogram in c: much faster hopefully. Now 7ms for 100x100
function maximaout=maximumfindcall(imin)
%maxima=[x,y,intensity]

simg=size(imin);
maxmax=numel(imin)/9;


maxima= maximumfind2(single(imin),uint32(maxmax));
% maxima= maximumfind(single(imin),uint32(maxmax));
% sum(maxima(:)-maxima2(:))
% size(maxima)
% sum(maxima)
f=find(maxima(:,1)==0,1,'first');
if ~isempty(f)
    maximaout=maxima(1:f-1,:);
else
    maximaout=maxima;
end


% maximaout=maxima(1:maxind-1,:);
```
