## Supplementary Software for "Field dependent deep learning enables high-throughput whole-cell 3D super-resolution imaging": mleFit_LM.m

```
function [P,CRLB,LogL]=mleFit_LM(varargin)
% varargin:
%imstack, startpsf/coeff, iterations, fitmode, isemccd, hidereport
% 1. imagestack (single)
% 2. fitmode
%   1 fix PSF
%   2 free PSF
%   3 Gauss fit z
%   4 fit PSFx, PSFy elliptical
%   5 cspline
% optional:
% 3. iterations (default=30)
% 4. paramters for fitters:
%   1. fix PSF: PSFxy sigma 
%   2. free PSF: start PSFxy
%   3. Gauss fit z: parameters: PSFx, Ax, Ay, Bx, By, gamma, d, PSFy
%       (single)
%   4. fit PSFx, PSFy elliptical: start PSFx, PSFy
%   5. cspline: cspline coefficients (single)

% 5. varmap: Variance map for sCMOS. If size(varmap) ~= size(imagestack):
%       no sCMOS correction is used. Default= emCCD
% 6. silent (suppress output if 1)
% 7. z start parameter (more than one: return solution with maximum
%       LIkelihood). Units: distance of stack calibration, center based

%Output:
%P
%1. X, Y, Photons, Background, Iterations
%2. X, Y, Photons, Background, PSFxy, Iterations
%3. X, Y, Photons, Background, Z, Iterations
%4. X, Y, Photons, Background, PSFx, PSFy, Iterations
%5. X, Y, Photons, Background, Z, Iterations
%6. X, Y, Photons, Background, Z, Iterations
%CRLB: cramer-rao lower bounds, as in P
%LogL: log-likelihood.

%Only for fitmode 6: P1 etc: results with z-startparameter<0, P2 etc:
%results with z-startparameter>0

P=[];CRLB=[];LogL=[]; %in case the function exits early
narginh=nargin;
%determine of it runs on GPU, otherwise use CPU as default
persistent fitter
% fitter = 2;
allfitters={@GPUmleFit_LM_new,@CPUmleFit_LM};
allfittersnames={'GPUmleFit_LM_new','CPUmleFit_LM'};
if isempty(fitter)
    testim=single(varargin{1}(:,:,1));
    for k=1:length(allfitters)
        try
            allfitters{k}(testim,1);
            fitter=k;
            break
        catch err
            % fitter did not work
        end
    end
    disp(['using: ' char(allfitters{fitter})]);
end

%convert all parameters to the correct format
imagestack=single(varargin{1});

if narginh>1 && ~isempty(varargin{2})
    fitmode=varargin{2};
else
    fitmode=2;
end

if narginh>2 && ~isempty(varargin{3})
    iterations=varargin{3};
else
    iterations=30;
end

if narginh>3 && ~isempty(varargin{4})
    fitpar=single(varargin{4});
else
    if ~(fitmode==3) &&  ~(fitmode==5)  &&  ~(fitmode==6) 
        fitpar=1; %for Gaussian fit
    else
        disp('for z fitting (Gauss or spline) fitting parameters (e.g. spline coefficients) are required');
        return
    end
end

if narginh>4 && ~isempty(varargin{5})
    varmap=single(varargin{5});
else
    varmap=0;
end

if narginh>5 && ~isempty(varargin{6})
    silent=single(varargin{6});
else
    silent=1;
end

coeffsize=size(fitpar);
%backward compatibility
% if fitmode==6
%     fitmode=5;
%     varargin{7}=[-coeffsize(3)/6, coeffsize(3)/6];
%     narginh=max(narginh,7);
% end

if narginh>6 && ~isempty(varargin{7}) && (fitmode==5 || fitmode==6) %only for spline fitting
    z0=(varargin{7}); %in units of calibration stack distance
    zstart=single(z0+coeffsize(3)/2);
elseif (fitmode==5 || fitmode==6) 
    zstart=single(coeffsize(3)/2);
else
    zstart=0;
end
%  zstart=[-70 -30 0 30 70]+75;
if fitmode ==6
    fitmode = 5;
end
[P,CRLB,LogL]=allfitters{fitter}(imagestack,fitmode, iterations,fitpar,varmap,silent,zstart(1));

if length(zstart)>1
    for k=2:length(zstart)
        [Ph,CRLBh,LogLh]=allfitters{fitter}(imagestack,fitmode, iterations,fitpar,varmap,silent,zstart(k));
%         indbettero=LogLh<LogL;
        indbetter=LogLh-LogL>1e-8; %copy only everything if LogLh increases by more than rounding error.
        P(indbetter,:)=Ph(indbetter,:);
        CRLB(indbetter,:)=CRLBh(indbetter,:);
        LogL(indbetter)=LogLh(indbetter);
    end
end
% if varargin{2}==6 %2D fit: find proper results
%     [P1,CRLB1,LogL1,P2,CRLB2,LogL2]=allfitters{fitter}(varargin{:});
%     ind1=LogL1>=LogL2;
%     ind2=LogL1<LogL2;
%     P=zeros(size(P1),'single');CRLB=zeros(size(CRLB1),'single');LogL=zeros(size(LogL1),'single');
%     P(ind1,:)=P1(ind1,:);P(ind2,:)=P2(ind2,:);
%     CRLB(ind1,:)=CRLB1(ind1,:);CRLB(ind2,:)=CRLB2(ind2,:);
%     LogL(ind1,:)=LogL1(ind1,:);LogL(ind2,:)=LogL2(ind2,:);
% else
%     [P,CRLB,LogL]=allfitters{fitter}(varargin{:});
%     P1=[];CRLB1=[];LogL1=[];P2=[];CRLB2=[];LogL2=[];
% end
%%
% 
% <latex>
% \begin{tabular}{|c|c|} \hline
% $n$ & $n!$ \\ \hline
% 1 & 1 \\
% 2 & 2 \\
% 3 & 6 \\ \hline
% \end{tabular}
% </latex>
% 
 clear(allfittersnames{fitter})
```
