## Supplementary Software for "Field dependent deep learning enables high-throughput whole-cell 3D super-resolution imaging": CalSplineCRLB.m

```
function [CRLB] =  CalSplineCRLB(coeff, sz, Theta)
%input:
%coeff: spline coeffcients for PSF model
%sz: fit windowsize
%Theta: Nfit*5 array, collumn value [xpos,ypos,photon,bg,zpos]
%output:
%CRLB for the input Theta

spline_xsize = size(coeff, 1);
spline_ysize = size(coeff, 2);
spline_zsize = size(coeff, 3);

NV = 5; %number of parameters to optimize

off = floor(((spline_xsize+1)-sz)/2);

Nfits = size(Theta,1);

for tx = 1:Nfits
    M = zeros(NV, NV);
    Minv = zeros(NV, NV);

    xc=-1 * (Theta(tx,1) - sz / 2 + 0.5);
    yc = -1 * (Theta(tx,2) - sz / 2 + 0.5);
    zc = Theta(tx,5)-floor(Theta(tx,5));
    
    xstart=floor(xc);
    ystart=floor(yc);
    zstart=floor(Theta(tx,5));
    
    xc = xc-floor(xc);
    yc = yc-floor(yc);
    
    [delta_f,delta_dxf,delta_ddxf1,delta_dyf,delta_ddyf1,delta_dzf,delta_ddzf1] = computeDelta3Dj_v2(single(xc),single(yc),single(zc));
    
    for ii = 1:sz
        for jj = 1:sz

            newTheta = Theta(tx,:);
            
            [newDudt, model] =  kernel_DerivativeSpline(ii+xstart+off,jj+ystart+off,zstart,spline_xsize,spline_ysize,spline_zsize,delta_f,delta_dxf,delta_dyf,delta_dzf,coeff,newTheta);
            
            for l = 0:NV-1
                for m =l:NV-1
                    
                    M(l*NV+m+1) = M(l*NV+m+1)+newDudt(l+1)*newDudt(m+1)/model;
                    M(m*NV+l+1) = M(l*NV+m+1);
                end
            end
            
        end
    end
    Minv = inv(M);
    for kk = 1:NV
        CRLB(Nfits*(kk-1)+tx)=Minv(kk,kk);
    end

end

CRLB = reshape(CRLB,Nfits,NV);
```
