## Supplementary Software for "Field dependent deep learning enables high-throughput whole-cell 3D super-resolution imaging": computeDelta3Dj_v2.m

```
function [delta_f,delta_dxf,delta_ddxf,delta_dyf,delta_ddyf,delta_dzf,delta_ddzf] = computeDelta3Dj_v2(x_delta, y_delta, z_delta)

delta_f = (zeros(64,1));
delta_dxf = (zeros(64,1));
delta_dyf = (zeros(64,1));
delta_dzf = (zeros(64,1));
delta_ddxf = (zeros(64,1));
delta_ddyf = (zeros(64,1));
delta_ddzf = (zeros(64,1));
cz = (1);
for i = 0:3
    cy = (1);
    for j = 0:3
        cx = (1);
        for k = 0:3
            delta_f(i*16+j*4+k+1) = (cx*cy*cz);
            if k<3
                delta_dxf(i*16+j*4+k+1+1) = ((k+1))*cx*cy*cz;
            end
            if k<2
                delta_ddxf(i*16+j*4+k+2+1)=((k+1)*(k+2))*cx*cy*cz;
            end
            if j<3
                delta_dyf(i*16+(j+1)*4+k+1) = ((j+1))*cx*cy*cz;
            end
            if j<2
                delta_ddyf(i*16+(j+2)*4+k+1)= ((j+1)*(j+2))*cx*cy*cz;
            end
            if i<3
                delta_dzf((i+1)*16+j*4+k+1) = ((i+1))*cx*cy*cz;
            end
            if i<2
                delta_ddzf((i+2)*16+j*4+k+1)=((i+1)*(i+2))*cx*cy*cz;
            end
            cx = cx*x_delta;
        end
        cy = cy*y_delta;
    end
    cz = cz*z_delta;
end
```
