## Supplementary Software for "Field dependent deep learning enables high-throughput whole-cell 3D super-resolution imaging": copyfieldsdeep.m

```
function [destination,missing]=copyfieldsdeep(destination,source,fields)
missing={};
if nargin==2 %copy all
    if ~isempty(source)
        if isstruct(source)
        fn=fieldnames(source);
            for k=1:length(fn)
                if ~isfield(destination,fn{k})
                    destination.(fn{k})=[];
                end
                destination.(fn{k})=copyfieldsdeep(destination.(fn{k}),source.(fn{k}));
            end
        else
            destination=source;
        end
           
    end
    
elseif nargin>2

%     fn=intersect(fields,fnsource);
    if ~isempty(source)
    indm=1;
    
    fnsource=fieldnames(source);
%     fnc=cellfun(@makehash,fnsource);
%     tocopy=intersect(fnsource,fields);
%     missing=setdiff(fields,fnsource);

%     for k=1:length(tocopy)
%         destination.(tocopy{k})=source.(tocopy{k});
%     end
%     fn=fields;
    for k=1:length(fields)
        if sum(strcmp(fnsource,fields{k}))
            destination.(fields{k})=source.(fields{k});
        else
            missing{indm}=fields{k};
            indm=indm+1;
        end
    end
    else
        missing=fields;
    end
end

function out=makehash(str)
    out=sum(char(str));
% out=1;
```
