## Supplementary Software for "Field dependent deep learning enables high-throughput whole-cell 3D super-resolution imaging": imageslicer.m

```
function imageslicer(varargin)
%optional: x,y,z
%V
%optional handle

%input parser: x,y,z (optional),V,'name',value
%Parent,all gui parameters: scale,  contrastmode, contrast,
%Title
%Tags
%   cell array of length of dimensions
%   for every dimension either empty (no tag for this dimension) or cell
%   arry of tags with length corresponding to length of this dimension

%todo: rgb, more dimension, choose dim1,dim2
if isa(varargin{1},'matlab.graphics.axis.Axes')||isa(varargin{1},'matlab.ui.Figure')
    p=parseinput(varargin(2:end));
    if isempty(p.Parent)
        p.Parent=varargin{1};
    end
else
p=parseinput(varargin);
end
% extend to rgb
V=p.V;
if ~isempty(p.x)
    axl{2}=p.x;
else
    axl{2}=1:size(V,2);
end
if ~isempty(p.y)
    axl{3}=p.y;
else
    axl{3}=1:size(V,3);
end
if ~isempty(p.z)
    axl{1}=p.z;
else
    axl{1}=1:size(V,1);
end


if isempty(p.Parent)
    phandle=figure;
else
    phandle=p.Parent;
    if isa(phandle,'matlab.graphics.axis.Axes')
        phandle=phandle.Parent;
    end
    
    delete(phandle.Children)
end

maxV=nanmax(V(:));
minV=nanmin(V(:));

 dim=1:2;
 dimmenu=3;
dimrgb=[];
if isprop(phandle,'WindowKeyPressFcn')
    phandle.WindowKeyPressFcn=@keypress;
end
ax=axes('Parent',phandle,'Position',[0.05,0.18,.95,.75]);
ax.XLim=[0 Inf];

vp1=0.08;
vp2=0.02;

numf=max(1,size(V,3)-1);

hslider{1}=uicontrol('Parent',phandle,'Style','slider','Units','normalized','Position',[0.05 vp1 0.35 0.05],...
    'Min',1,'Max',size(V,3),'Value',1,'SliderStep',[1/(numf) 5/(numf)],'Callback',{@slidercallback,1});
hslider{2}=uicontrol('Parent',phandle,'Style','slider','Units','normalized','Position',[0.05 vp2 0.35 0.05],...
    'Min',1,'Max',size(V,3),'Value',1,'SliderStep',[1/(numf) 5/(numf)],'Callback',{@slidercallback,2});
hframe{1}=uicontrol('Parent',phandle,'Style','edit','Units','normalized','String','1','Position',[0.4 vp1 0.075 0.05],'Callback',{@framecallback,1});
hframe{2}=uicontrol('Parent',phandle,'Style','edit','Units','normalized','String','1','Position',[0.4 vp2 0.075 0.05],'Callback',{@framecallback,2});

hslidert{1}=uicontrol('Parent',phandle,'Style','edit','Units','normalized','Position',[0.02 vp1 0.03 0.05],'String','3', 'Callback',{@changeaxis,2});
hslidert{2}=uicontrol('Parent',phandle,'Style','edit','Units','normalized','Position',[0.02 vp2 0.03 0.05],'String','4', 'Callback',{@changeaxis,3});

hmenu{1}=uicontrol('Parent',phandle,'Style','popupmenu','Units','normalized','String',{'x','y','z'},'Position',[0.475 vp1 0.125 0.05],...
    'Callback',{@changeaxis,0});
hmenu{2}=uicontrol('Parent',phandle,'Style','popupmenu','Units','normalized','String',{'x','y','z'},'Position',[0.6 vp1 0.125 0.05],...
    'Callback',{@changeaxis,1},'Value',2);
haxscale=uicontrol('Parent',phandle,'Style','checkbox','Units','normalized','String','fill','Position',[0.475 vp2 0.1 0.05],...
    'Callback',@plotimage,'Value',p.fill);
hlut=uicontrol('Parent',phandle,'Style','popupmenu','Units','normalized','String',{'parula','gray','hot','jet'},'Position',[0.725 vp1 0.175 0.05],...
    'Callback',@plotimage);
hcontrastcheck=uicontrol('Parent',phandle,'Style','checkbox','Units','normalized','String','global contrast','Position',[0.6 vp2 0.2 0.05],...
    'Callback',@plotimage,'Value',p.globalcontrast);
hcontrast=uicontrol('Parent',phandle,'Style','edit','Units','normalized','String','1','Position',[0.8 vp2 0.1 0.05],'Callback',@plotimage);

hresetax=uicontrol('Parent',phandle,'Style','pushbutton','Units','normalized','String','reset','Position',[0.9 vp2 0.1 0.05],'Callback',@resetax);

hrgb=uicontrol('Parent',phandle,'Style','checkbox','Units','normalized','String','RGB','Position',[0.9 vp1 0.1 0.05],'Callback',@updatergb,'Value',p.rgb);
% updateall
hmenu{1}.Value=p.xdim;
hmenu{2}.Value=p.ydim;
changeaxis(0,0,0);
% plotimage


    function resetax(a,b)
        ax.XLim=[0 Inf];
        hcontrast.String='1';
        plotimage;
    end
    function slidercallback(a,b,slider)
        setslice(a.Value,slider);
    end
    function framecallback(a,b,slider)
       setslice(str2double( a.String),slider);
    end

    function changeaxis(a,b,axv)
        
        oax=~axv;
        
        if axv<2 && strcmp(hmenu{axv+1}.String{hmenu{axv+1}.Value},hmenu{oax+1}.String{hmenu{oax+1}.Value})
            if hmenu{axv+1}.Value==1
                hmenu{oax+1}.Value=2;
            else
                hmenu{oax+1}.Value=1;
            end
        end
        ax.XLim=[-inf inf];
        ax.YLim=[-inf inf];
        updateall

    end
    
    function setslice(frame,slider,plot)
        frame=min(frame,hslider{slider}.Max);
        frame=max(1,frame);
        hframe{slider}.String=num2str(round(frame));
        hslider{slider}.Value=round(frame);
        if nargin<=2 || plot
        plotimage
        end
    end
    function updatergb(a,b)
        p.rgb=a.Value;
        updateall;
    end
    function updateall(a,b)
        
        s=size(V);
            
        
        if hrgb.Value&&any(s==3)
            dimrgb=find(s==3,1,'last');
            dims=setdiff(3:length(s),dimrgb); 
        else
            dimrgb=[];
            dims=3:length(s);
        end
        str={'x','y'};
        for k=1:length(dims)
            str{end+1}=num2str(dims(k));
        end
        hmenu{1}.String=str;
        hmenu{2}.String=str;
         if hmenu{1}.Value>2 
            dim(1)=str2double(str(hmenu{1}.Value));
         else
             dim(1)=hmenu{1}.Value;
         end
         if hmenu{2}.Value>2 
            dim(2)=str2double(str(hmenu{2}.Value));
         else
             dim(2)=hmenu{2}.Value;
         end
%             dim(2)=str2double(str(hmenu{2}.Value)); 
        dimmenu=setdiff(1:length(s),[dim dimrgb]);
        
%         strm=str;strm(dimrgb)=[];
        numfh=max(1,size(V,dimmenu(1))-1);
        hslider{1}.SliderStep=[1/(numfh) 5/(numfh)];
        hslider{1}.Max=size(V,dimmenu(1));
         setslice(min(hslider{1}.Max,hslider{1}.Value),1,0);
         
         if dimmenu(1)<3
         hslidert{1}.String=str{dimmenu(1)};
         else
             hslidert{1}.String=num2str(dimmenu(1));
         end
        if length(dimmenu)>1
            if size(V,dimmenu(2))>1
            hslider{2}.SliderStep=[1/(size(V,dimmenu(2))-1) 5/(size(V,dimmenu(2))-1)];
            else
                hslider{2}.SliderStep=[1 1];
            end
            hslider{2}.Max=size(V,dimmenu(2));
             setslice(min(hslider{2}.Max,hslider{2}.Value),2,0);
            hslider{2}.Visible='on';
            hframe{2}.Visible='on';
            hslidert{2}.Visible='on';
             if dimmenu(2)<3
                hslidert{2}.String=str{dimmenu(2)};
             else
                 hslidert{2}.String=num2str(dimmenu(2));
             end
        else
            hslider{2}.Visible='off';
            hframe{2}.Visible='off';
            hslidert{2}.Visible='off';
        end
        plotimage
        
    end
    function plotimage(a,b,c)
%         disp('plot')
         s=size(V);
            
        for k=1:length(s)
%             if k<=4+double(p.rgb)
            dimall{k}=1:s(k);
%             else
%                 dimall{k}=1;
%             end
        end
        xlimold=ax.XLim;
        ylimold=ax.YLim;
        
%         slicez=round(str2double(hframe.String));
%         slicet=round(str2double(hframe2.String));
        
        if dim(1)<=3
            a1=axl{dim(1)};
        else
            a1=1:size(V,dim(1));
        end
        if dim(2)<=3
            a2=axl{dim(2)};
        else
            a2=1:size(V,dim(2));
        end
        dimall{dimmenu(1)}=round(str2double(hframe{1}.String));
        if length(dimmenu)>1
            dimall{dimmenu(2)}=round(str2double(hframe{2}.String));
        end
        for k=3:length(dimmenu)
            dimall{dimmenu(k)}=1; %XXXX later remember position
        end
        Vsl=V(dimall{:});
        dims=[dim(2) dim(1) dimrgb dimmenu];
        dimshow=(1:length(size(V)));
        dimshow(1:length(dims))=dims;
        
        Vslp=permute(Vsl,dimshow);
        
        img=(squeeze(Vslp));
       
        
        %contrast
        contrast=str2num(hcontrast.String);
        if length(contrast)==2
            imin=contrast(1);
            imax=contrast(2);
        else
            if hcontrastcheck.Value
                meanV=(minV+maxV)/2;
                dV=(maxV-minV)/2;
                imax=meanV+dV*contrast(1);
                imin=meanV-dV*contrast(1);
            else
                imaxim=nanmax(img(:));
                iminim=nanmin(img(:));
                if isnan(imaxim)
                    imax=inf;
                    imin=-inf;
                else
                    meanV=(iminim+imaxim)/2;
                    dV=(imaxim-iminim)/2;
                    imax=meanV+dV*contrast(1);
                    imin=meanV-dV*contrast(1);
                end
            end
        end
         if imax==0, imax=1;end
        
        img(img>imax)=imax;
        img(img<imin)=imin;
        if length(size(img))==3 %???
            img=(img-imin)/(imax-imin);
        end

        imagesc(ax,a1,a2,img);

        if haxscale.Value
            axis(ax,'fill')
        else
            axis(ax,'equal')
        end
        if ~isinf(xlimold(2))
            ax.XLim=xlimold;ax.YLim=ylimold;
        else
            d1=a1(2)-a1(1);
            d2=a2(2)-a2(1);
            ax.XLim=a1([1 end])+[-1 1]*d1/2;
            ax.YLim=a2([1 end])+[-1 1]*d2/2;
        end
        colormap(ax,hlut.String{hlut.Value})
%         imin=nanmin(img(:));
        ax.CLim=[imin imax];
        tagtxt='';
        titletxt='';
        if ~isempty(p.Title)
            titletxt{1}=p.Title;
        end
        if ~isempty(p.Tags)
            tags=p.Tags;
%             stags=size(tags);
            for d=3:length(dims) %1,2 of dims used for plotting
                dimp=dims(d);
                indexh=dimall{dimp};
                if dimp<=length(tags) && length(tags{dimp})>=indexh
                    tagsall=tags{dimp};
                    if ~iscell(tagsall)
                        tagh=tagsall(indexh);
                    else
                        tagh=tagsall{indexh};
                    end
                    if isnumeric(tagh)
                        tagh=num2str(tagh);
                    end
                    tagtxt=[tagtxt ' ' num2str(dimp) ': ' tagh];
                end
                    
            end
            if ~iscell(titletxt) %title defined
                titletxt{1}=tagtxt;
            else
                titletxt{end+1}=tagtxt;
            end
        end
        if iscell(titletxt)
            title(titletxt)
        end

        
        if ~p.rgb
            colorbar(ax)
        end
        
    end
    function keypress(a,b)
        if contains(b.Modifier,'shift')
            slider=2;
        else
            slider=1;
        end
        if strcmp(b.Character,'+')||strcmp(b.Key,'rightarrow')
            frame=hslider{slider}.Value;
            setslice(frame+1,slider);
        elseif strcmp(b.Character,'-')||strcmp(b.Key,'leftarrow')
            frame=hslider{slider}.Value;
            setslice(frame-1,slider);
        elseif strcmp(b.Key,'uparrow')
            hcontrast.String=num2str(str2double(hcontrast.String)*1.1,'%1.2f');
            plotimage;
        elseif strcmp(b.Key,'downarrow')
            hcontrast.String=num2str(str2double(hcontrast.String)*.9,'%1.2f');
            plotimage;
        end
            
        
    end
end


function pv=parseinput(in)
p=inputParser;
p.addOptional('x',[],@isnumeric);
p.addOptional('y',[],@isnumeric);
p.addOptional('z',[],@isnumeric);
p.addRequired('V',@isnumeric);
p.addParameter('Parent',[]);
p.addParameter('fill',false);
p.addParameter('xdim',1,@isnumeric);
p.addParameter('ydim',2,@isnumeric);
p.addParameter('rgb',false);
p.addParameter('globalcontrast',false,@islogical);
p.addParameter('Title',[]);
p.addParameter('Tags',[]);
parse(p,in{:});
pv=p.Results;

end
```
