## Supplementary Software for "Field dependent deep learning enables high-throughput whole-cell 3D super-resolution imaging": imx.m

```
classdef imx<handle
properties
    handle
    guihandles
    parameters
    V
    showframes 
    dimrgb
end
methods
        
    function obj=imx(varargin)
        %V
    %optional: x,y,z
    
    %optional handle

    %input parser:V, x,y,z (optional),'name',value
    %Parent,all gui parameters: scale,  contrastmode, contrast,
    %Title
    %Tags
    %   cell array of length of dimensions
    %   for every dimension either empty (no tag for this dimension) or cell
    %   arry of tags with length corresponding to length of this dimension

    %todo: rgb, more dimension, choose dim1,dim2
    if isa(varargin{1},'matlab.graphics.axis.Axes')||isa(varargin{1},'matlab.ui.Figure')
        p=parseinput(varargin(2:end));
        if isempty(p.Parent)
            p.Parent=varargin{1};
        end
    else
        p=parseinput(varargin);
    end
        obj.parameters=p;
        makegui(obj,p)
    end
    % extend to rgb
    
    % plotimage


        function resetax(obj,a,b)
            obj.guihandles.axis.XLim=[0 Inf];
            obj.guihandles.hcontrast.String='1';
            obj.plotimage;
        end
        function slidercallback(obj,a,b,slider)
            obj.setslice(a.Value,slider);
        end
        function framecallback(obj,a,b,slider)
           obj.setslice(str2double( a.String),slider);
        end

        function changeaxis(obj,a,b,axv)
            obj.guihandles.axis.XLim=[-inf inf];
            obj.guihandles.axis.YLim=[-inf inf];
                if obj.guihandles.hrgb.Value&&any(s==3)
                    obj.dimrgb=find(s==3,1,'last');
                    dims=setdiff(3:length(s),obj.dimrgb); 
                else
                    obj.dimrgb=[];
                    dims=3:length(size(obj.V));
                end
            str={'x','y'};
            for k=1:length(dims)
                str{end+1}=num2str(dims(k));
            end
            obj.guihandles.hmenu{1}.String=str;
            obj.guihandles.hmenu{2}.String=str;
            dim=unique(obj.getdims,'stable');
            for k=1:2
                obj.guihandles.hmenu{k}.Value=dim(k);
            end
            for k=1:length(dim)-2
                obj.guihandles.hslidert{k}.String=str(dim(k+2));
            end
                obj.updateall
        end

        function setslice(obj,frame,slider,plot)
            frame=min(frame,obj.guihandles.hslider{slider}.Max);
            frame=max(1,frame);
            obj.guihandles.hframe{slider}.String=num2str(round(frame));
            obj.guihandles.hslider{slider}.Value=round(frame);
            if nargin<=3 || plot
                obj.plotimage
            end
        end
        function updatergb(a,b)
            p.rgb=a.Value;
            updateall;
        end
        function updateall(obj,a,b)
             warning('off','MATLAB:callback:error');
            %update all gui parameters
            s=size(obj.V);
%             for k=1:length(s)
%                 obj.showframes{k}=1;
%             end
            if obj.guihandles.hrgb.Value&&any(s==3)
                obj.dimrgb=find(s==3,1,'last');
                dims=setdiff(3:length(s),obj.dimrgb); 
            else
                obj.dimrgb=[];
                dims=3:length(s);
            end
            str={'x','y'};
            for k=1:length(dims)
                str{end+1}=num2str(dims(k));
            end
            obj.guihandles.hmenu{1}.String=str;
            obj.guihandles.hmenu{2}.String=str;
            dim=obj.getdims;

    %             dim(2)=str2double(str(hmenu{2}.Value)); 
            dimmenu=setdiff(1:length(s),[dim(1:2) obj.dimrgb]);

    %         strm=str;strm(dimrgb)=[];
    
            dims1=str2double(strrep(strrep(obj.guihandles.hslidert{1}.String,'x','1'),'y','2'));
            numfh=max(1,size(obj.V,dims1));
            obj.guihandles.hslider{1}.SliderStep=[1/(numfh) 5/(numfh)];
            obj.guihandles.hslider{1}.Max=size(obj.V,dims1);
            obj.setslice(min(obj.guihandles.hslider{1}.Max,obj.guihandles.hslider{1}.Value),1,0);

            if length(dim)>3
                dims2=str2double(strrep(strrep(obj.guihandles.hslidert{2}.String,'x','1'),'y','2'));
%                 dims2=str2double(obj.guihandles.hslidert{2}.String);
                if size(obj.V,dims2)>1
                    obj.guihandles.hslider{2}.SliderStep=[1/(size(obj.V,dims2)-1) 5/(size(obj.V,dims2)-1)];
                else
                    obj.guihandles.hslider{2}.SliderStep=[1 1];
                end
                obj.guihandles.hslider{2}.Max=size(obj.V,dims2);
                obj.setslice(min(obj.guihandles.hslider{2}.Max,obj.guihandles.hslider{2}.Value),2,0);
                obj.guihandles.hslider{2}.Visible='on';
                obj.guihandles.hframe{2}.Visible='on';
                obj.guihandles.hslidert{2}.Visible='on';
            else
                obj.guihandles.hslider{2}.Visible='off';
                obj.guihandles.hframe{2}.Visible='off';
                obj.guihandles.hslidert{2}.Visible='off';
            end
            obj.plotimage
%              warning('on','MATLAB:callback:error');
        end
        function plotimage(obj,a,b,c)
            warning('off','MATLAB:callback:error');
%              s=size(obj.V);
            xlimold=obj.guihandles.axis.XLim;
            ylimold=obj.guihandles.axis.YLim;
            dim=obj.getdims;
            Vsl=obj.V(obj.showframes{:});
           
            dims=[dim(2) dim(1) obj.dimrgb dim(3:end)];
            Vslp=permute(Vsl,dims);

            img=(squeeze(Vslp));

             if dim(1)<=3
                a1=obj.parameters.axl{dim(1)};
            else
                a1=1:size(obj.V,dim(1));
            end
            if dim(2)<=3
                a2=obj.parameters.axl{dim(2)};
            else
                a2=1:size(obj.V,dim(2));
            end
            %contrast
            contrast=str2num(obj.guihandles.hcontrast.String);
            if length(contrast)==2
                imin=contrast(1);
                imax=contrast(2);
            else
                maxV=nanmax(obj.V(:));
                minV=nanmin(obj.V(:));
                if obj.guihandles.hcontrastcheck.Value
                    meanV=(minV+maxV)/2;
                    dV=(maxV-minV)/2;
                    imax=meanV+dV*contrast(1);
                    imin=meanV-dV*contrast(1);
                else
                    imaxim=nanmax(img(:));
                    iminim=nanmin(img(:));
                    if isnan(imaxim)
                        imax=inf;
                        imin=-inf;
                    else
                        meanV=(iminim+imaxim)/2;
                        dV=(imaxim-iminim)/2;
                        imax=meanV+dV*contrast(1);
                        imin=meanV-dV*contrast(1);
                    end
                end
            end
             if imax==0, imax=1;end

            img(img>imax)=imax;
            img(img<imin)=imin;
            if length(size(img))==3 %???
                img=(img-imin)/(imax-imin);
            end

            imagesc(obj.guihandles.axis,a1,a2,img);

            if obj.guihandles.haxscale.Value
                axis(obj.guihandles.axis,'fill')
            else
                axis(obj.guihandles.axis,'equal')
            end
            if ~isinf(xlimold(2))
                obj.guihandles.axis.XLim=xlimold;obj.guihandles.axis.YLim=ylimold;
            else
                d1=a1(2)-a1(1);
                d2=a2(2)-a2(1);
                obj.guihandles.axis.XLim=a1([1 end])+[-1 1]*d1/2;
                obj.guihandles.axis.YLim=a2([1 end])+[-1 1]*d2/2;
            end
            colormap(obj.guihandles.axis,obj.guihandles.hlut.String{obj.guihandles.hlut.Value})
            obj.guihandles.axis.CLim=[imin imax];
            tagtxt='';
            titletxt='';
            p=obj.parameters;
            if ~isempty(p.Title)
                titletxt{1}=p.Title;
            end
            if ~isempty(p.Tags)
                tags=p.Tags;
                for d=3:length(dims) %1,2 of dims used for plotting
                    dimp=dims(d);
                    indexh=obj.showframes{dimp};
                    if dimp<=length(tags) && length(tags{dimp})>=indexh
                        tagsall=tags{dimp};
                        if ~iscell(tagsall)
                            tagh=tagsall(indexh);
                        else
                            tagh=tagsall{indexh};
                        end
                        if isnumeric(tagh)
                            tagh=num2str(tagh);
                        end
                        tagtxt=[tagtxt ' ' num2str(dimp) ': ' tagh];
                    end

                end
                if ~iscell(titletxt) %title defined
                    titletxt{1}=tagtxt;
                else
                    titletxt{end+1}=tagtxt;
                end
            end
            if iscell(titletxt)
                title(obj.guihandles.axis,titletxt)
            end

            if ~p.rgb
                colorbar(obj.guihandles.axis)
            end
%              warning('on','MATLAB:callback:error');

        end
        function keypress(obj,a,b)
            if contains(b.Modifier,'shift')
                slider=2;
            else
                slider=1;
            end
            if strcmp(b.Character,'+')||strcmp(b.Key,'rightarrow')
                frame=obj.guihandles.hslider{slider}.Value;
                obj.setslice(frame+1,slider);
            elseif strcmp(b.Character,'-')||strcmp(b.Key,'leftarrow')
                frame=obj.guihandles.hslider{slider}.Value;
                obj.setslice(frame-1,slider);
            elseif strcmp(b.Key,'uparrow')
                obj.guihandles.hcontrast.String=num2str(str2double(obj.guihandles.hcontrast.String)*1.1,'%1.2f');
                obj.plotimage;
            elseif strcmp(b.Key,'downarrow')
                obj.guihandles.hcontrast.String=num2str(str2double(obj.guihandles.hcontrast.String)*.9,'%1.2f');
                obj.plotimage;
            end


        end
        function dim=getdims(obj)
            %dim(1:2) xy display; dim(3:4) connected to sliders
            %showframes{k}: which to show, was dimall before
             if obj.guihandles.hmenu{1}.Value>2 
                dim(1)=str2double(obj.guihandles.hmenu{1}.String{obj.guihandles.hmenu{1}.Value});
             else
                 dim(1)=obj.guihandles.hmenu{1}.Value;
             end
             if obj.guihandles.hmenu{2}.Value>2 
                dim(2)=str2double(obj.guihandles.hmenu{2}.String{obj.guihandles.hmenu{2}.Value});
             else
                 dim(2)=obj.guihandles.hmenu{2}.Value;
             end
             s=size(obj.V);
             if length(s)>3
             dim(3)=str2double(strrep(strrep(obj.guihandles.hslidert{1}.String,'x','1'),'y','2'));
             end
             if length(s)>3
             dim(4)=str2double(strrep(strrep(obj.guihandles.hslidert{2}.String,'x','1'),'y','2'));
             end
             dimmissing=setdiff(1:length(s),dim);
             dim=[dim dimmissing];
             for k=1:2
                obj.showframes{dim(k)}=1:s(dim(k));
             end
             for k=3:length(dim)
                 if dim(k)<=length(obj.showframes) && length(obj.showframes{dim(k)})>1 
                     obj.showframes{dim(k)}=round(mean(obj.showframes{dim(k)}));
                 end
             end
             if length(s)>2
            obj.showframes{dim(3)}=round(str2double(obj.guihandles.hframe{1}.String));
             end
            if length(s)>3 %&& strcmp(obj.guihandles.hframe{2}.Visible,'on')
                obj.showframes{dim(4)}=round(str2double(obj.guihandles.hframe{2}.String));
            end
        end
        function delete(obj)
             warning('on','MATLAB:callback:error');
        end
end
end

function makegui(obj,p)
V=p.V;
obj.V=V;
    if ~isempty(p.x)
        obj.parameters.axl{1}=p.x;
    else
        obj.parameters.axl{1}=1:size(V,1);
    end
    if ~isempty(p.y)
        obj.parameters.axl{2}=p.y;
    else
        obj.parameters.axl{2}=1:size(V,2);
    end
    if ~isempty(p.z)
        obj.parameters.axl{3}=p.z;
    else
        obj.parameters.axl{3}=1:size(V,3);
    end

    if isempty(p.Parent)
        obj.handle=figure;
    else
        obj.handle=p.Parent;
        if isa(obj.handle,'matlab.graphics.axis.Axes')
            obj.handle=obj.handle.Parent;
        end

        delete(obj.handle.Children)
    end

    if isprop(obj.handle,'WindowKeyPressFcn')
        obj.handle.WindowKeyPressFcn=@obj.keypress;
    end
    ax=axes('Parent',obj.handle,'Position',[0.05,0.18,.95,.75]);
    ax.XLim=[0 Inf];
    obj.guihandles.axis=ax;

    vp1=0.08;
    vp2=0.02;

    numf=max(1,size(V,3)-1);

    obj.guihandles.hslider{1}=uicontrol('Parent',obj.handle,'Style','slider','Units','normalized','Position',[0.05 vp1 0.35 0.05],...
        'Min',1,'Max',size(V,3),'Value',1,'SliderStep',[1/(numf) 5/(numf)],'Callback',{@obj.slidercallback,1});
    obj.guihandles.hslider{2}=uicontrol('Parent',obj.handle,'Style','slider','Units','normalized','Position',[0.05 vp2 0.35 0.05],...
        'Min',1,'Max',size(V,3),'Value',1,'SliderStep',[1/(numf) 5/(numf)],'Callback',{@obj.slidercallback,2});
    obj.guihandles.hframe{1}=uicontrol('Parent',obj.handle,'Style','edit','Units','normalized','String','1','Position',[0.4 vp1 0.075 0.05],'Callback',{@obj.framecallback,1});
    obj.guihandles.hframe{2}=uicontrol('Parent',obj.handle,'Style','edit','Units','normalized','String','1','Position',[0.4 vp2 0.075 0.05],'Callback',{@fobj.ramecallback,2});

    obj.guihandles.hslidert{1}=uicontrol('Parent',obj.handle,'Style','edit','Units','normalized','Position',[0.02 vp1 0.03 0.05],'String','3', 'Callback',{@obj.changeaxis,2});
    obj.guihandles.hslidert{2}=uicontrol('Parent',obj.handle,'Style','edit','Units','normalized','Position',[0.02 vp2 0.03 0.05],'String','4', 'Callback',{@obj.changeaxis,3});

    obj.guihandles.hmenu{1}=uicontrol('Parent',obj.handle,'Style','popupmenu','Units','normalized','String',{'x','y','z'},'Position',[0.475 vp1 0.125 0.05],...
        'Callback',{@obj.changeaxis,0});
    obj.guihandles.hmenu{2}=uicontrol('Parent',obj.handle,'Style','popupmenu','Units','normalized','String',{'x','y','z'},'Position',[0.6 vp1 0.125 0.05],...
        'Callback',{@obj.changeaxis,1},'Value',2);
    obj.guihandles.haxscale=uicontrol('Parent',obj.handle,'Style','checkbox','Units','normalized','String','fill','Position',[0.475 vp2 0.1 0.05],...
        'Callback',@obj.plotimage,'Value',p.fill);
    obj.guihandles.hlut=uicontrol('Parent',obj.handle,'Style','popupmenu','Units','normalized','String',{'parula','gray','hot','jet'},'Position',[0.725 vp1 0.175 0.05],...
        'Callback',@obj.plotimage);
    obj.guihandles.hcontrastcheck=uicontrol('Parent',obj.handle,'Style','checkbox','Units','normalized','String','global contrast','Position',[0.6 vp2 0.2 0.05],...
        'Callback',@obj.plotimage,'Value',p.globalcontrast);
    obj.guihandles.hcontrast=uicontrol('Parent',obj.handle,'Style','edit','Units','normalized','String','1','Position',[0.8 vp2 0.1 0.05],'Callback',@obj.plotimage);

    obj.guihandles.hresetax=uicontrol('Parent',obj.handle,'Style','pushbutton','Units','normalized','String','reset','Position',[0.9 vp2 0.1 0.05],'Callback',@obj.resetax);

    obj.guihandles.hrgb=uicontrol('Parent',obj.handle,'Style','checkbox','Units','normalized','String','RGB','Position',[0.9 vp1 0.1 0.05],'Callback',@obj.updatergb,'Value',p.rgb);
    % updateall
    obj.guihandles.hmenu{1}.Value=p.xdim;
    obj.guihandles.hmenu{2}.Value=p.ydim;
                s=size(obj.V);
            for k=1:length(s)
                obj.showframes{k}=1;
            end
    obj.changeaxis(0,0,0);
end
function pv=parseinput(in)
p=inputParser;
p.addRequired('V',@isnumeric);
p.addOptional('x',[],@isnumeric);
p.addOptional('y',[],@isnumeric);
p.addOptional('z',[],@isnumeric);

p.addParameter('Parent',[]);
p.addParameter('fill',false);
p.addParameter('xdim',1,@isnumeric);
p.addParameter('ydim',2,@isnumeric);
p.addParameter('rgb',false);
p.addParameter('globalcontrast',false,@islogical);
p.addParameter('Title',[]);
p.addParameter('Tags',[]);
parse(p,in{:});
pv=p.Results;
% pv
end
```
