## Supplementary Software for "Field dependent deep learning enables high-throughput whole-cell 3D super-resolution imaging": kernel_DerivativeSpline.m

```
function [dudt,model] =  kernel_DerivativeSpline(xc,yc,zc,xsize,ysize,zsize,delta_f,delta_dxf,delta_dyf,delta_dzf,coeff,theta)
dudt = zeros(5,1);
xc = max(xc,0);
xc = min(xc,xsize-1);

yc = max(yc,0);
yc = min(yc,ysize-1);

zc = max(zc,0);
zc = min(zc,zsize-1);

temp = coeff(xc+1,yc+1,zc+1,:);
dudt(1)=-1*theta(3)*sum(delta_dxf.*(temp(:)));
dudt(2)=-1*theta(3)*sum(delta_dyf.*(temp(:)));
dudt(5)=theta(3)*sum(delta_dzf.*(temp(:)));
dudt(3)=sum(delta_f.*(temp(:)));
dudt(4)=1;
model = theta(4)+theta(3)*dudt(3);
```
