## Supplementary Software for "Field dependent deep learning enables high-throughput whole-cell 3D super-resolution imaging": readstruct.m

```
function p=readstruct(file,replacestruct,parseascell)
if nargin<2
    replacestruct={};
end
if nargin<3
    parseascell=false;
end

if iscell(file) %convert arry of txt
    txt=file;
    p=[];
    for k=1:length(txt)
         pline=parseline(txt{k},replacestruct,parseascell);
        p=copyfieldsdeep(p,pline);
    end
else
    %read struct from text file
    fid=fopen(file);
    if fid<0
        p=[];
        return
    end
    line=fgetl(fid);
    p=[];
    while ischar(line)

        pline=parseline(line,replacestruct,parseascell);
        p=copyfieldsdeep(p,pline);
        line=fgetl(fid);

    end
    fclose(fid);
end
    

function p=parseline(txt,replacestruct,parseascell)
indp=strfind(txt,'.');
inde=strfind(txt,'=');
if isempty(inde)
    p=[];
    return
end
inde=inde(1);
indp(indp>inde)=[];
val=strtrim(txt(inde+1:end));
indend=strfind(val,';');
val(indend:end)=[];

rep=false;
for k=1:length(replacestruct)
    tr=replacestruct{k}{1};
    if strcmp(tr,val)
        rpobj=replacestruct{k}{2};
        val=rpobj;
        rep=true;
        break;
    end
end
if ~rep
    valtest=str2double(val(1:min(2,length(val))));
%     valtest=str2num(val(1:min(3,length(val))));
%     if ~isempty(valtest)&&isnumeric(valtest)
    if ~isnan(valtest)
        valn=str2num(val);
    else
        valn=[];
    end
    if ~isempty(valn)&&isnumeric(valn)
        val=valn;
    elseif parseascell %comma separated
        indcomma=strfind(val,',');
        ind=[0 indcomma length(val)+1];
        for k=1:length(ind)-1;
            valc{k}=val(ind(k)+1:ind(k+1)-1);
        end
        val=valc;
        if length(val)==1
            val=val{1};
        end
    end
end


indp(end+1)=inde;
indp=[0,indp];
% p=val;
for k=length(indp)-1:-1:1
    field=txt(indp(k)+1:indp(k+1)-1);
    p=[];
    p.(field)=val;
    val=p;
end

    
```
