## Supplementary Software for "Field dependent deep learning enables high-throughput whole-cell 3D super-resolution imaging": selectManyFiles.m

```
classdef selectManyFiles<handle
    properties %(Access=private)
        handle;
        guihandles;
        filelist;
        addpar;
        startpath;
    end

    methods
        function obj=selectManyFiles(varargin)
            obj.makegui;
            if nargin>0
                obj.startpath=varargin{1};
            end
        end
        function makegui(obj)
            if isempty(obj.handle)||~isvalid(obj.handle)
                obj.handle=figure('Units','normalized','Units','pixels','Position',[150,200,900,330],'Name','Select Files','NumberTitle','off','ToolBar','none','MenuBar','none');
                delete(obj.handle.Children);
            end
            obj.guihandles.filelist=uicontrol('Style','listbox','Parent',obj.handle,'Position',[10 10 680,300]);
            uicontrol('Style','pushbutton','String','add','Callback',@obj.addb_callback,'Parent',obj.handle,'Position',[700 270 100 30])
            uicontrol('Style','pushbutton','String','add dir','Callback',@obj.adddirb_callback,'Parent',obj.handle,'Position',[800 270 100 30])
            uicontrol('Style','pushbutton','String','remove ','Callback',@obj.removeb_callback,'Parent',obj.handle,'Position',[750 230 100 30])
%             obj.guihandles.ch2check=uicontrol('Style','checkbox','String','Ch2 in 2nd file','Callback',@obj.ch2check_callback,'Parent',obj.handle,'Position',[700 185 200 30]);
%             obj.guihandles.selectch2=uicontrol('Style','pushbutton','String','select Ch2','Callback',@obj.selectch2_callback,'Parent',obj.handle,'Position',[700 160 100 30],'Visible','off');
            obj.guihandles.findall=uicontrol('Style','pushbutton','String','find all','Callback',@obj.findall_callback,'Parent',obj.handle,'Position',[800 160 100 30],'Visible','off')       ;      
             
            uicontrol('Style','pushbutton','String','Done','Callback',@obj.done_callback,'Parent',obj.handle,'Position',[750 10 100 50])
            obj.guihandles.freepos=uicontrol('Style','text','String','','Position',[700 60 200 30]);
            c=uicontextmenu;
            obj.guihandles.filelist.UIContextMenu=c;
            
            m1=uimenu(c,'Label','move up','Callback',@obj.filelistmenu);
            m2=uimenu(c,'Label','move down','Callback',@obj.filelistmenu);
        end
        function filelistmenu(obj,in,a,b)
            select=obj.guihandles.filelist.Value;
            str=obj.guihandles.filelist.String;
            allvalues=1:length(str);
            selectnew=select;
            switch in.Text
                case 'move up'
                    if select>1
                        allvalues(select-1)=select;
                        allvalues(select)=select-1;
                        selectnew=select-1;
                    end
                case 'move down'
                    if select<length(allvalues)
                        allvalues(select+1)=select;
                        allvalues(select)=select+1;
                        selectnew=select+1;
                    end
            end
            obj.guihandles.filelist.String=str(allvalues);
            obj.guihandles.filelist.Value=selectnew;
        end
        function ch2check_callback(obj,a,b)
            if a.Value
                obj.guihandles.selectch2.Visible='on';
                obj.guihandles.findall.Visible='on';
            else
                obj.guihandles.selectch2.Visible='off';
                obj.guihandles.findall.Visible='off';
            end
        end
        function selectch2_callback(obj,a,b)
            filenumber=obj.guihandles.filelist.Value;
            filech1=obj.guihandles.filelist.String{filenumber};
            ind=strfind(filech1,';');
            if ~isempty(ind)
                filech1=filech1(1:ind-1);
            end
            [path1,file1,ext1]=fileparts(filech1);
            [f,p]=uigetfile(['*' ext1],'select Ch2 for selected file',filech1)
            if ~f
                return
            end
            filech2=[p  f];
            filecombined=[filech1 ';' filech2];
            obj.guihandles.filelist.String{filenumber}=filecombined;
        end
        function findall_callback(obj,a,b)
            filelist=obj.guihandles.filelist.String; 
            assigned=find(contains(filelist,';'));
            if isempty(assigned)
                warndlg('please select second channel file for at least one file');
                return
            end
            filenumber=assigned(1);
            filenames=filelist{filenumber};
            ind1= strfind(filenames,';');
            f1=filenames(1:ind1-1);f2=filenames(ind1+1:end);
            for k=1:length(filelist)
                fileh=filelist{k};
                ind=strfind(fileh,';');
                if ~isempty(ind)
                    fileh=fileh(1:ind-1);
                end
                fileh2=filenamereplace(fileh,f1,f2);
                obj.guihandles.filelist.String{k}=[fileh ';' fileh2];
            end

        end
        function done_callback(obj,a,b)
            obj.filelist=obj.guihandles.filelist.String;
            fn=fieldnames(obj.guihandles);
            for k=1:length(fn)
                obj.addpar.(fn{k})=copyfields([],obj.guihandles.(fn{k}),{'String','Value'});
            end
            close(obj.handle);
            
        end
        function addb_callback(obj,a,b)
            filelist=obj.guihandles.filelist.String;
            path=obj.startpath;
            if  ~isempty(filelist)
                fileselect=filelist{obj.guihandles.filelist.Value};
                if ~isempty(fileselect)
                    path=fileparts(fileselect);
                end
            end
            [f,path]=uigetfile(fullfile(path,'*.tif;*.*'),'Select image  files','MultiSelect','on');
            if ~iscell(f)
                if ~f
                return
                end
             
                f={f};
            end
               
            str=filelist;
            for k=1:length(f)
                str{end+1}=[path f{k}];
            end
            obj.guihandles.filelist.String=str;
            obj.guihandles.filelist.Value=max(min(obj.guihandles.filelist.Value,length(str)),1);
        end
        function adddirb_callback(obj,a,b)
            filelist=obj.guihandles.filelist.String;
            path=obj.startpath;
            if  ~isempty(filelist)
                fileselect=filelist{obj.guihandles.filelist.Value};
                if ~isempty(fileselect)
                    path=fileparts(fileselect);
                end
            end

            [path]=uigetdirs(path,'Select image or batch directories');
            if isempty(path)
                return
            end
            
            str=filelist;
            for k=1:length(path)
                imf=findimageindir(path{k});
                if ~isempty(imf)
                    for l=1:length(imf)
                        str{end+1}=imf{l};
                    end
                end
                    
            end
            obj.guihandles.filelist.String=str;   
            obj.guihandles.filelist.Value=max(min(obj.guihandles.filelist.Value,length(str)),1);
        end
       
        function removeb_callback(obj,a,b)
  
            vo=obj.guihandles.filelist.Value;
            str=obj.guihandles.filelist.String;
            if isempty(str)
                return
            end
            str(vo)=[];
            obj.guihandles.filelist.String=str;
            obj.guihandles.filelist.Value=max(1,min(vo,length(str)));
        end
        function processb_callback(obj,a,b)
            obj.setPar('loc_preview',false);
            p=obj.getGuiParameters;
            
            filelist=p.filelist.String;
            if obj.onlinebatch
                obj.processonline(filelist{1})
                return
            end
            for k=1:length(filelist)
                obj.locData.clear;
                filen=filelist{k};
                if ~isempty(strfind(filen,'.tif'))
                    obj.processtiff(filen);
                elseif ~isempty(strfind(filen,'.mat')) 
                    if p.useforall
                        obj.processtiffromWF(filen);
                    else
                        obj.processWF(filen);
                    end
                end
            end
        end
        
        function processonline(obj,file)
            p=obj.getGuiParameters;
            if ~isempty(obj.filesprocessed)
                answ=questdlg('already files processed. Fit everything again? (No: remaining files are fitted, cancel: abort)');
                if strcmp(answ,'Yes')
                    obj.filesprocessed={};
                elseif strcmp(answ,'Cancel')
                    return
                end
            end
            while obj.guihandles.stop.Value==0
                checksml=p.omitsml;
%                 profile on
                imf=findunprocessed(p.filelist.selection,p,obj.dirsprocessed,checksml)
%                 profile viewer
%                 imf=findimageindir(p.filelist.selection,p);
%                 unprocessed=setdiff(imf,obj.filesprocessed);
                if ~isempty(imf)
                    thisfile=imf;
                    obj.guihandles.status.String=['fitting ' thisfile];drawnow;
                    disp(['fitting ' thisfile]);
                     obj.processtiff(thisfile);
                    obj.filesprocessed{end+1}=thisfile;
                    obj.dirsprocessed{end+1}=fileparts(thisfile);
                else
                    obj.guihandles.status.String=['no file found. waiting '];drawnow; 
                    pause(1)
                end
               
                
            end 
            obj.guihandles.stop.Value=0;
            obj.guihandles.status.String=['stopped. '];drawnow;
        end
        
        function processWF(obj,wffile)
            wf=interfaces.Workflow([],obj.P);
            wf.attachLocData(obj.locData);
            wf.makeGui;
            wf.load(wffile,[],true,true);
            disp(mytextsplit(wf.description));
            %workflow: recover tiff file and start
            loadergui=wf.module(wf.startmodule);
            loader=loadergui.loaders{loadergui.getGuiParameters.loaderlist.Value};
            tiffile=loader.getGuiParameters.tiffile;
            loader.addFile(tiffile);
%             tiffile=wf.module(wf.startmodule).getGuiParameters.tiffile;
%             wf.module(wf.startmodule).addFile(tiffile);
            wf.run;  
            delete(wf)
        end
        function processtiff(obj,tiffile)
            p=obj.getGuiParameters;
            wf=interfaces.Workflow([],obj.P);
%             ld=interfaces.LocalizationData;
            ld=obj.locData;
            ld.attachPar(obj.P);
            wf.attachLocData(ld);
            wf.makeGui;
            wf.load(p.mainbatchfile,[],true,true);
            wf.module(wf.startmodule).addFile(tiffile);
            disp(mytextsplit(wf.description));
            wf.run;
            delete(wf)
%             delete(ld)
        end
        
        function processtiffromWF(obj,wffile)
            wf=interfaces.Workflow([],obj.P);
            wf.attachLocData(obj.locData);
            wf.makeGui;
            wf.load(wffile,[],true,true);
            %workflow: recover tiff file and start
            loadergui=wf.module(wf.startmodule);
            loader=loadergui.loaders{loadergui.getGuiParameters.loaderlist.Value};
            tiffile=loader.getGuiParameters.tiffile;
            obj.processtiff(tiffile);
            delete(wf)
        end

        
    end

end

function img=findunprocessed(path,p,dirlist,checksml)
img='';
searchstr=p.adddir_mask;
mintiffs=p.adddir_minimages;
files=dir([path filesep searchstr]);
matfiles=dir([path filesep '*_sml.mat']);
matfilesc={matfiles(:).name};
for k=1:length(files)
    if files(k).isdir 
        if myisstrfind(dirlist, files(k).name)% isempty(strfind(dirlist,files(k).name))
            continue
        end
        if strcmp(files(k).name,'.')||strcmp(files(k).name,'..')
            continue
        end
        if checksml&&myisstrfind(matfilesc,files(k).name)
            continue
        end
        tiffiles=(dir([path filesep files(k).name filesep '*.tif']));
        addpathh='';
        if isempty(tiffiles)
            tiffiles=(dir([path filesep files(k).name filesep 'Pos0' filesep '*.tif']));
            addpathh=['Pos0' filesep];
        end
        if length(tiffiles)>mintiffs
            img=[path filesep files(k).name filesep addpathh tiffiles(1).name];
            return
        end
    end 
%     p.hstatus.String=['directory ' num2str(k) ' of ' num2str(length(files))];
%     drawnow;
end
end
function img=findimageindir(path,p)
img={};
files=dir([path filesep '*.tif']);
if ~isempty(files)
    img={[path filesep files(1).name]};
    return
end
files=dir([path filesep 'Pos*']);
for k=1:length(files)
    if files(k).isdir
        path2=[path filesep files(k).name filesep];
        files2=dir([path2 '*.tif']);
        if ~isempty(files2)
            img={[path2 files2(1).name]};
            return
        end
    end
end
searchstr='*.tif';
mintiffs=1;
files=dir([path filesep searchstr]);
for k=1:length(files)
    if files(k).isdir 
        tiffiles=(dir([path filesep files(k).name filesep '*.tif']));
        addpathh='';
        if isempty(tiffiles)
            tiffiles=(dir([path filesep files(k).name filesep 'Pos0' filesep '*.tif']));
            addpathh=['Pos0' filesep];
        end
        if length(tiffiles)>mintiffs
            img{end+1}=[path filesep files(k).name filesep addpathh tiffiles(1).name];
        end
    end 

end
end

function pard=guidef(obj)
pard.filelist.object=struct('Style','listbox');%,'Callback',@obj.moduleselect_callback);
pard.filelist.position=[11,1];
pard.filelist.Height=9;
pard.filelist.Width=3;

pard.mainbatchfile.object=struct('Style','edit','String','x://*_batch.mat');
pard.mainbatchfile.position=[1,1];
pard.mainbatchfile.Width=3;

pard.mainbatchfile_button.object=struct('Style','pushbutton','String','load main batchfile','Callback',@obj.mainbatchfileb_callback);
pard.mainbatchfile_button.position=[1,4];
pard.mainbatchfile_button.Width=1;

pard.add_button.object=struct('Style','pushbutton','String','add','Callback',@obj.addb_callback);
pard.add_button.position=[3,4];
pard.add_button.Width=1;

pard.adddir_button.object=struct('Style','pushbutton','String','add directories','Callback',@obj.adddirb_callback);
pard.adddir_button.position=[4,4];
pard.adddir_button.Width=1;

pard.adddironline_button.object=struct('Style','pushbutton','String','add online directory','Callback',@obj.adddironlineb_callback);
pard.adddironline_button.position=[5,4];
pard.adddironline_button.Width=1;


pard.adddir_mask.object=struct('Style','edit','String','*_Localization_*');
pard.adddir_mask.position=[6,4];
pard.adddir_mask.Width=1;

pard.adddir_t.object=struct('Style','text','String','> #images');
pard.adddir_t.position=[7,4];
pard.adddir_t.Width=.5;
pard.adddir_minimages.object=struct('Style','edit','String','0');
pard.adddir_minimages.position=[7,4.5];
pard.adddir_minimages.Width=.5;


pard.omitsml.object=struct('Style','checkbox','String','skip if .sml exist','Value',1);
pard.omitsml.position=[8,4];
pard.omitsml.Width=1;


pard.remove_button.object=struct('Style','pushbutton','String','remove ','Callback',@obj.removeb_callback);
pard.remove_button.position=[9,4];
pard.remove_button.Width=1;

pard.process_button.object=struct('Style','pushbutton','String','Batch process','Callback',@obj.processb_callback);
pard.process_button.position=[11,4];
pard.process_button.Height=2;

pard.useforall.object=struct('Style','checkbox','String','use for all','Value',0);
pard.useforall.position=[2,4];
pard.useforall.Height=1;

pard.status.object=struct('Style','text','String','status','Value',0);
pard.status.position=[12.5,1];
pard.status.Width=3.5;


pard.stop.object=struct('Style','togglebutton','String','stop','Value',0);
pard.stop.position=[12.3,4.5];
pard.stop.Width=.5;

pard.plugininfo.type='ProcessorPlugin'; 
end
```
