## Supplementary Software for "Field dependent deep learning enables high-throughput whole-cell 3D super-resolution imaging": Spline3D_interp.m

```
function coeff = Spline3D_interp(PSF)
[X,Y,Z] = meshgrid(1:size(PSF,1),1:size(PSF,2),1:size(PSF,3));
[Xq,Yq,Zq] = meshgrid(1:1/3:size(PSF,1),1:1/3:size(PSF,2),1:1/3:size(PSF,3));
PSFup = interp3(X,Y,Z,PSF,Xq,Yq,Zq,'spline');

A = zeros(64,64);
for i = 1:4
    dx = (i-1)/3;
    for j = 1:4
        dy = (j-1)/3;
        for k = 1:4
            dz = (k-1)/3;
            for l = 1:4
                for m = 1:4
                    for n = 1:4
                        A((i-1)*16+(j-1)*4+k,(l-1)*16+(m-1)*4+n) = dx^(l-1)*dy^(m-1)*dz^(n-1);
                    end
                end
            end
        end
    end
end


coeff = zeros(size(PSF,1)-1,size(PSF,2)-1,size(PSF,3)-1,64);

for i = 1:size(PSF,1)-1
    for j = 1:size(PSF,2)-1
        for k = 1:size(PSF,3)-1
            temp = PSFup((i-1)*3+1:3*i+1,(j-1)*3+1:3*j+1,(k-1)*3+1:3*k+1);
            temp = reshape(temp,[64,1]);
            x = A\temp;
            coeff(i,j,k,:)=reshape(x(:),[1,1,1,64]);
        end
    end
end        
```
