## Supplementary Software for "Field dependent deep learning enables high-throughput whole-cell 3D super-resolution imaging": struct2txt.m

```
function txt=struct2txt(p,prefix)
txt={};
if isstruct(p)||isobject(p)
    
    fn=fieldnames(p);
    for k=1:length(fn)
        prefixn=[ prefix,'.'  fn{k}];
        to=struct2txt(p.(fn{k}),prefixn);
        for l=1:length(to)
            txt{end+1}=to{l};
        end
    end
else
    if iscell(p)
        to=p{1};
        if isnumeric(to)||islogical(to)
            to=num2str(to);
        end
        for k=2:length(p)
            th=p{k};
            if isnumeric(th)||islogical(th)
                th=num2str(th);
            end
            if ischar(th)
                to=[to ',' th];
            end
        end
        p=to;
    end
    if isnumeric(p)||islogical(p)
        p=num2str(p);
    end
    if ~isempty(p)
        tx=p(1,:);
        if ischar(tx)
            txt={[prefix '=' tx]};
        end
    else
        txt={[prefix '=']};
    end
end
```
