## Supplementary Software for "Field dependent deep learning enables high-throughput whole-cell 3D super-resolution imaging": writestruct.m

```
function txt=writestruct(file,p)%replacestruct,parseascell)
% if nargin<2
%     replacestruct={{}};
% end
% if nargin<3
%     parseascell=false;
% end
%read struct from text file
 txt=struct2txt(p,'');
 for k=1:length(txt)
     txt{k}(1)=[];
 end
 fid=fopen(file,'w');
 if fid>0
     for k=1:length(txt)
         fprintf(fid,'%s\n',txt{k});
     end
     fclose(fid);
 end
 
end

% line=fgetl(fid);
% p=[];
% while ischar(line)
%     
%     pline=parseline(line,replacestruct,parseascell);
%     p=copyfieldsdeep(p,pline);
%     line=fgetl(fid);
%     
% end
% fclose(fid);
    

    
% end

function p=parseline(txt,replacestruct,parseascell)
indp=strfind(txt,'.');
inde=strfind(txt,'=');
if isempty(indp)||isempty(inde)
    p=[];
    return
end
inde=inde(1);
indp(indp>inde)=[];
val=txt(inde+1:end);
indend=strfind(val,';');
val(indend:end)=[];

rep=false;
for k=1:length(replacestruct)
    tr=replacestruct{k}{1};
    if strcmp(tr,val)
        rpobj=replacestruct{k}{2};
        val=rpobj;
        rep=true;
        break;
    end
end
if ~rep
valn=str2num(val);
    if ~isempty(valn)
        val=valn;
    elseif parseascell %comma separated
        ind=[0 strfind(val,',') length(val)];
        for k=1:length(ind)-1;
            valc{k}=val(ind(k)+1:ind(k)-1);
        end
        val=valc;
    end
end


indp(end+1)=inde;
indp=[0,indp];
% p=val;
for k=length(indp)-1:-1:1
    field=txt(indp(k)+1:indp(k+1)-1);
    p=[];
    p.(field)=val;
    val=p;
end

end
```
