## Supplementary Software for "Field dependent deep learning enables high-throughput whole-cell 3D super-resolution imaging": myrmfield.m

```
function t = myrmfield(s,field)
%RMFIELD Remove fields from a structure array.
%   S = RMFIELD(S,'field') removes the specified field from the
%   m x n structure array S. The size of input S is preserved.
%
%   S = RMFIELD(S,FIELDS) removes more than one field at a time
%   when FIELDS is a character array or cell array of strings.  The
%   changed structure is returned. The size of input S is preserved.
%
%   See also SETFIELD, GETFIELD, ISFIELD, FIELDNAMES.

%   Copyright 1984-2008 The MathWorks, Inc.

%--------------------------------------------------------------------------------------------
% handle input arguments
if isa(s,'struct') 
    if ~iscell(field)
        field={field};
    end
    field=unique(field);
   field = cellstr(field); 


% get fieldnames of struct
f = fieldnames(s);

% Determine which fieldnames to delete.
idxremove = [];
for i=1:length(field)
  j = find(strcmp(field{i},f) == true);
  if isempty(j)
%     if length(field{i}) > namelengthmax
%       error(message('MATLAB:rmfield:FieldnameTooLong', field{ i }));
%     else
%       error(message('MATLAB:rmfield:InvalidFieldname', field{ i }));
%     end
  end
  idxremove = [idxremove;j];
end

% set indices of fields to keep
idxkeep = 1:length(f);
idxkeep(idxremove) = [];

% remove the specified fieldnames from the list of fieldnames.
f(idxremove,:) = [];

% convert struct to cell array
c = struct2cell(s);

% find size of cell array
sizeofarray = size(c);
newsizeofarray = sizeofarray;

% adjust size for fields to be removed
newsizeofarray(1) = sizeofarray(1) - length(idxremove);

% rebuild struct
t = cell2struct(reshape(c(idxkeep,:),newsizeofarray),f);
else
    t=[];
end

%--------------------------------------------------------------------------------------------
```
