## Supplementary Software for "Field dependent deep learning enables high-throughput whole-cell 3D super-resolution imaging": myprctile.m

```
function y = prctile(x,p,dim)
%PRCTILE Percentiles of a sample.
%   Y = PRCTILE(X,P) returns percentiles of the values in X.  P is a scalar
%   or a vector of percent values.  When X is a vector, Y is the same size
%   as P, and Y(i) contains the P(i)-th percentile.  When X is a matrix,
%   the i-th row of Y contains the P(i)-th percentiles of each column of X.
%   For N-D arrays, PRCTILE operates along the first non-singleton
%   dimension.
%
%   Y = PRCTILE(X,P,DIM) calculates percentiles along dimension DIM.  The
%   DIM'th dimension of Y has length LENGTH(P).
%
%   Percentiles are specified using percentages, from 0 to 100.  For an N
%   element vector X, PRCTILE computes percentiles as follows:
%      1) The sorted values in X are taken as the 100*(0.5/N), 100*(1.5/N),
%         ..., 100*((N-0.5)/N) percentiles.
%      2) Linear interpolation is used to compute percentiles for percent
%         values between 100*(0.5/N) and 100*((N-0.5)/N)
%      3) The minimum or maximum values in X are assigned to percentiles
%         for percent values outside that range.
%
%   PRCTILE treats NaNs as missing values, and removes them.
%
%   Examples:
%      y = prctile(x,50); % the median of x
%      y = prctile(x,[2.5 25 50 75 97.5]); % a useful summary of x
%
%   See also IQR, MEDIAN, NANMEDIAN, QUANTILE.

%   Copyright 1993-2004 The MathWorks, Inc.
%   $Revision: 1.1.8.1 $  $Date: 2010/03/16 00:16:55 $

if ~isvector(p) || numel(p) == 0
    error('stats:prctile:BadPercents', ...
          'P must be a scalar or a non-empty vector.');
elseif any(p < 0 | p > 100) || ~isreal(p)
    error('stats:prctile:BadPercents', ...
          'P must take real values between 0 and 100');
end

% Figure out which dimension prctile will work along.
sz = size(x);
if nargin < 3 
    dim = find(sz ~= 1,1);
    if isempty(dim)
        dim = 1; 
    end
    dimArgGiven = false;
else
    % Permute the array so that the requested dimension is the first dim.
    nDimsX = ndims(x);
    perm = [dim:max(nDimsX,dim) 1:dim-1];
    x = permute(x,perm);
    % Pad with ones if dim > ndims.
    if dim > nDimsX
        sz = [sz ones(1,dim-nDimsX)];
    end
    sz = sz(perm);
    dim = 1;
    dimArgGiven = true;
end

% If X is empty, return all NaNs.
if isempty(x)
    if isequal(x,[]) && ~dimArgGiven
        y = nan(size(p),class(x));
    else
        szout = sz; szout(dim) = numel(p);
        y = nan(szout,class(x));
    end

else
    % Drop X's leading singleton dims, and combine its trailing dims.  This
    % leaves a matrix, and we can work along columns.
    nrows = sz(dim);
    ncols = prod(sz) ./ nrows;
    x = reshape(x, nrows, ncols);

    x = sort(x,1);
    nonnans = ~isnan(x);

    % If there are no NaNs, do all cols at once.
    if all(nonnans(:))
        n = sz(dim);
        if isequal(p,50) % make the median fast
            if rem(n,2) % n is odd
                y = x((n+1)/2,:);
            else        % n is even
                y = (x(n/2,:) + x(n/2+1,:))/2;
            end
        else
            q = [0 100*(0.5:(n-0.5))./n 100]';
            xx = [x(1,:); x(1:n,:); x(n,:)];
            y = zeros(numel(p), ncols, class(x));
            y(:,:) = interp1q(q,xx,p(:));
        end

    % If there are NaNs, work on each column separately.
    else
        % Get percentiles of the non-NaN values in each column.
        y = nan(numel(p), ncols, class(x));
        for j = 1:ncols
            nj = find(nonnans(:,j),1,'last');
            if nj > 0
                if isequal(p,50) % make the median fast
                    if rem(nj,2) % nj is odd
                        y(:,j) = x((nj+1)/2,j);
                    else         % nj is even
                        y(:,j) = (x(nj/2,j) + x(nj/2+1,j))/2;
                    end
                else
                    q = [0 100*(0.5:(nj-0.5))./nj 100]';
                    xx = [x(1,j); x(1:nj,j); x(nj,j)];
                    y(:,j) = interp1q(q,xx,p(:));
                end
            end
        end
    end

    % Reshape Y to conform to X's original shape and size.
    szout = sz; szout(dim) = numel(p);
    y = reshape(y,szout);
end
% undo the DIM permutation
if dimArgGiven
     y = ipermute(y,perm);  
end

% If X is a vector, the shape of Y should follow that of P, unless an
% explicit DIM arg was given.
if ~dimArgGiven && isvector(x)
    y = reshape(y,size(p)); 
end
```
