## Supplementary Software for "Field dependent deep learning enables high-throughput whole-cell 3D super-resolution imaging": myquantile.m

```
function y = quantile(x,p,dim)
%QUANTILE Quantiles of a sample.
%   Y = QUANTILE(X,P) returns quantiles of the values in X.  P is a scalar
%   or a vector of cumulative probability values.  When X is a vector, Y is
%   the same size as P, and Y(i) contains the P(i)-th quantile.  When X is
%   a matrix, the i-th row of Y contains the P(i)-th quantiles of each
%   column of X.  For N-D arrays, QUANTILE operates along the first
%   non-singleton dimension.
%
%   Y = QUANTILE(X,P,DIM) calculates quantiles along dimension DIM.  The
%   DIM'th dimension of Y has length LENGTH(P).
%
%   Y = QUANTILE(X,N,...) returns quantiles at the N evenly-spaced cumulative
%   probabilities (1:N)/(N+1).  N is an integer value greater than 1.
%
%   Quantiles are specified using cumulative probabilities, from 0 to 1.
%   For an N element vector X, QUANTILE computes quantiles as follows:
%      1) The sorted values in X are taken as the (0.5/N), (1.5/n),
%         ..., ((N-0.5)/N) quantiles.
%      2) Linear interpolation is used to compute quantiles for
%         probabilities between (0.5/N) and ((N-0.5)/N)
%      3) The minimum or maximum values in X are assigned to quantiles
%         for probabilities outside that range.
%
%   QUANTILE treats NaNs as missing values, and removes them.
%
%   Examples:
%      y = quantile(x,.50); % the median of x
%      y = quantile(x,[.25 .50 .75]); % the quartiles of x
%      y = quantile(x,3); % another way to get quartiles
%      y = quantile(x,[.025 .25 .50 .75 .975]); % a useful summary of x
%
%   See also PRCTILE, IQR, MEDIAN.

%   Copyright 1993-2010 The MathWorks, Inc. 
%   $Revision: 1.1.8.2 $  $Date: 2010/04/11 20:42:29 $

if ~isvector(p) || isempty(p)
    error('stats:quantile:BadProbs', ...
          'Second input must be a scalar or a non-empty vector.');
elseif isscalar(p) && isreal(p) && (p == round(p)) && (p > 1) % 1 means 100%, not 1 quantile
    p = (1:p) / (p+1);
elseif any( p > 1) || ~isreal(p)
    error('stats:quantile:BadProbs', ...
          'Second input must be a vector of real values between 0 and 1, or a positive integer value.');
end

for k=1:length(p)
    if (p(k))<0
    p(k)=1-10^p(k);
    end
end
x=double(x);
p=double(p);

if nargin < 3
    y = myprctile(x,100.*p);
else
    y = myprctile(x,100.*p,dim);
end
```
