## Supplementary Software for "Field dependent deep learning enables high-throughput whole-cell 3D super-resolution imaging": zernikecoeffMap.m

```
function maps_gauss = zernikecoeffMap(p,beads_zrnikeparam,axmap,axdist)
pos = beads_zrnikeparam.pos;
value = beads_zrnikeparam.value_P;
RRSE1 =beads_zrnikeparam.RRSE.RRSE1;
std_ref=beads_zrnikeparam.aberrations_std;
mapsmooth_set=beads_zrnikeparam.mapsmooth;
modelrrse_set=beads_zrnikeparam.modelrrse;
localroi_set=beads_zrnikeparam.localroi;

value_all = value;
pos_all = pos;
beads_num = length(pos);
frame_size = beads_zrnikeparam.roi;
value_std = std(transpose(cell2mat(value)));
value_mean = mean(transpose(cell2mat(value)));
ZernikeModeIndx =[]; 
frame_fov = beads_zrnikeparam.roi;
value_indx=NaN(frame_size);
if min(frame_fov)>localroi_set
    xy_roi = localroi_set;
else
    xy_roi = min(frame_fov);
end

for j=1:21
%     j
    for i = beads_num:-1:1        %The flashback loop is designed to prevent skipping when successive beads have a problem
        value_indx(pos{i}(1),pos{i}(2))=value{i}(j);     
    end
    value_pos{j} = value_indx;
    value_indx=NaN(frame_size);
    
end

for i = 1:beads_num   
%     i
      pos{i} = pos{i}(1:2);
      pos_all{i} = pos_all{i}(1:2);
      for j=1:21
         value_pos_crop=  value_pos{j}(max(pos{i}(1)-xy_roi/2,0)+1:min(pos{i}(1)+xy_roi/2,frame_fov(1)),max(pos{i}(2)-xy_roi/2,0)+1:min(pos{i}(2)+xy_roi/2,frame_fov(2)));
         value_pos_crop_t = reshape(value_pos_crop, size(value_pos_crop,1)*size(value_pos_crop,2),1);  %To convert a 2D array into a 1D array
         value_pos_crop_mean(j) = mean(value_pos_crop_t,'omitnan');   
         value_pos_crop_std(j) = std(value_pos_crop_t,'omitnan');
      end
      value_roi_mean{i}=value_pos_crop_mean;
      value_roi_std{i}=value_pos_crop_std;   
end

for i = beads_num:-1:1
    pos{i} = pos{i}(1:2);
    if any(transpose(value_all{i})>value_roi_mean{i}+std_ref*value_roi_std{i}) || any(transpose(value_all{i})<value_roi_mean{i}-std_ref*value_roi_std{i}) || RRSE1(i)>modelrrse_set     % Filter outlier  beads
        pos{i} = [];
        value{i} = [];
    end

end
pos(cellfun(@isempty,pos))=[];  %Take out the null data
value(cellfun(@isempty,value))=[];  %Take out the null data

beads_num_filter = length(pos);

for j = 1:21       
     p.status.String=['calculate zernike coefficient maps of individual PSFs: ' num2str(j) ' of 21']; drawnow

    x = zeros(1,beads_num_filter);
    y = zeros(1,beads_num_filter);
    z = zeros(1,beads_num_filter);

    for i = 1:beads_num_filter
        x(i) = pos{i}(1,1);
        y(i) = pos{i}(1,2);
        z(i) = value{i}(j,1);
    end
    
   F=scatteredInterpolant(x',y',z','natural');
   [x1,y1]=meshgrid(1:1:frame_size(2),1:1:frame_size(1));
   maps{j}=F(x1,y1);
   maps_gauss{j} = imgaussfilt(maps{j},mapsmooth_set);
   maps_show(:,:,j) = maps_gauss{j};
   ZernikeModeIndx_temp = {ZernikeMode(beads_zrnikeparam.model,j)};
   ZernikeModeIndx = [ZernikeModeIndx,ZernikeModeIndx_temp];
end

BeadsCoord=cell2mat(pos_all);
X_BeadsCoord = BeadsCoord(1:2:length(BeadsCoord));
Y_BeadsCoord = BeadsCoord(2:2:length(BeadsCoord));
scatter(axdist,X_BeadsCoord,Y_BeadsCoord,30,'blue');
xlim(axdist,[0 frame_size(1)]);
ylim(axdist,[0 frame_size(2)]);
hold(axdist,'on');
scatter(axdist,x,y,30,'x','red');
set(axdist,'YDir','reverse'); 
hold(axdist,'off');
legend(axdist,'total beads','valid beads','Location','northeast');
xlabel(axdist,'X (pixel)');
ylabel(axdist,'Y (pixel)');
title(axdist,['total beads  N= ',num2str(beads_num), ' beads'  newline   'pick up beads N =',num2str(length(x)),'beads']);
box(axdist,'on');

tags={[],[],ZernikeModeIndx,[]};
imx(maps_show,'Parent',axmap,'Tags',tags);

end
```
