## Supplementary Software for "Field dependent deep learning enables high-throughput whole-cell 3D super-resolution imaging": zernikefitBeadstack.m

```
function [paraSim,PSFzernike,aberrations_avg1,RRSE,MAPE]=zernikefitBeadstack(data,p,psfrescale,axzernike,axpupil,axmode)
Npixel = size(data,1);
Nz = size(data,3);
paraFit=p;
paraFit.sizeX = size(data,1);
paraFit.sizeY = size(data,2);
paraFit.sizeZ = size(data,3);

paraFit.aberrations = [2,-2,0.0; 2,2,0.0; 3,-1,0.0; 3,1,0.0; 4,0,0.0; 3,-3,0.0; 3,3,0.0; 4,-2,0.0; 4,2,0.0; 5,-1,0.0; 5,1,0.0; 6,0,0.0; 4,-4,0.0; 4,4,0.0;  5,-3,0.0; 5,3,0.0;  6,-2,0.0; 6,2,0.0; 7,1,0.0; 7,-1,0.0; 8,0,0.0];
paraFit.aberrations(:,3) =  paraFit.aberrations(:,3)*paraFit.lambda;

numAberrations = size(paraFit.aberrations,1);
shared = [ones(1,numAberrations) 1 1 1 p.sharedIB p.sharedIB ];  % 1 is shared parameters between z slices, 0 is free parameters between z slices, only consider  [x, y, z, I, bg]

sumShared = sum(shared);
numparams = 26*Nz-sumShared*(Nz-1);


thetainit = zeros(numparams,1);

bg = zeros(1,Nz);
Nph = zeros(1,Nz);
x0 = zeros(1,Nz);
y0 = zeros(1,Nz);
z0 = zeros(1,Nz);

% center of mass with nm unit
ImageSizex = paraFit.pixelSizeX*Npixel/2;
ImageSizey = paraFit.pixelSizeY*Npixel/2;

DxImage = 2*ImageSizex/paraFit.sizeX;
DyImage = 2*ImageSizey/paraFit.sizeY;
ximagelin = -ImageSizex+DxImage/2:DxImage:ImageSizex;
yimagelin = -ImageSizey+DyImage/2:DyImage:ImageSizey;
[YImage,XImage] = meshgrid(yimagelin,ximagelin);
for i = 1:Nz
    dTemp = data(:,:,i);
    bg(i) = min(dTemp(:));
    bg(i) = max(bg(i),1);
    Nph(i) = sum(sum(dTemp-bg(i)));
    x0(i) = sum(sum(XImage.*dTemp))/Nph(i);
    y0(i) = sum(sum(YImage.*dTemp))/Nph(i);
    z0(i) = 0;
end

allTheta = zeros(numAberrations+5,Nz);
allTheta(numAberrations+1,:)=x0';
allTheta(numAberrations+2,:)=y0';
allTheta(numAberrations+3,:)=z0';
allTheta(numAberrations+4,:)=Nph';
allTheta(numAberrations+5,:)=bg';
allTheta(1:numAberrations,:) = repmat(paraFit.aberrations(:,3),[1 Nz]);

% for 
map = zeros(numparams,3);
n=1;
for i = 1:numAberrations+5
    if shared(i)==1
        map(n,1)= 1;
        map(n,2)=i;
        map(n,3)=0;
        n = n+1;
    elseif shared(i)==0
        for j = 1:Nz
            map(n,1)=0;
            map(n,2)=i;
            map(n,3)=j;
            n = n+1;
        end
    end
end

for i = 1:numparams
    if map(i,1)==1
        thetainit(i)= mean(allTheta(map(i,2),:));
    elseif map(i,1)==0
         thetainit(i) = allTheta(map(i,2),map(i,3));
    end
end

% we assume that parameters for zernike coefficiens are always linked
zernikecoefsmax = 0.25*paraFit.lambda*ones(numAberrations,1);
paraFit.maxJump = [zernikecoefsmax',paraFit.pixelSizeX*ones(1,max(Nz*double(shared(numAberrations+1)==0),1)),paraFit.pixelSizeY*ones(1,max(Nz*double(shared(numAberrations+2)==0),1)),500*ones(1,max(Nz*double(shared(numAberrations+3)==0),1)),2*max(Nph(:)).*ones(1,max(Nz*double(shared(numAberrations+4)==0),1)),100*ones(1,max(Nz*double(shared(numAberrations+5)==0),1))];


%% fit data
paraFit.numparams = numparams;
paraFit.numAberrations = numAberrations;
paraFit.zemitStack = zeros(size(data,3),1); % move emitter
zmax=(paraFit.sizeZ-1)*paraFit.dz/2;
paraFit.objStageStack=zmax:-paraFit.dz:-zmax;


paraFit.ztype = 'stage';
paraFit.map = map;
paraFit.Nitermax = paraFit.iterations;

parC.NA = paraFit.NA;
parC.refmed = paraFit.refmed;
parC.refcov = paraFit.refcov;
parC.refimm = paraFit.refimm;
parC.lambda = paraFit.lambda;
parC.zemit0 = paraFit.zemit0;
parC.objStage0 = paraFit.objStage0;
parC.pixelSizeX = paraFit.pixelSizeX;
parC.pixelSizeY = paraFit.pixelSizeY;
parC.Npupil = paraFit.Npupil;
parC.sizeX = paraFit.sizeX;
parC.sizeY = paraFit.sizeY;
parC.sizeZ = paraFit.sizeZ;
parC.aberrations = paraFit.aberrations;
parC.maxJump = paraFit.maxJump;
parC.numparams = paraFit.numparams;
parC.numAberrations = paraFit.numAberrations;
parC.zemitStack = paraFit.zemitStack;
parC.objStageStack = paraFit.objStageStack;
parC.ztype = paraFit.ztype;
parC.map = paraFit.map;
parC.Nitermax = paraFit.Nitermax;
  
paraFitCell= struct2cell(parC);
data_double = double(data.*1);
thetainit_d =thetainit.*1;
shared_d=shared.*1;


%%
% calculate the psf gauss directly using Isigma   
I_sigmax=psfrescale;
I_sigmay=psfrescale;
[x1,y1]=meshgrid( - 2 : 2 );
gauss_psf=exp(-x1.^2./2./I_sigmax^2).*exp(-y1.^2./2./I_sigmay^2);
gauss_psf = gauss_psf/sum(gauss_psf,'all');
tempGauss1=gauss_psf;
%%
   
%    tic
if gpuDeviceCount>0
    [P,model,~] = MLE_FitAbberation_Final_GPU_float(data_double,thetainit_d,paraFitCell,shared_d,0.1,tempGauss1);
else
    [P,model,~] = MLE_FitAbberation_Final_CPU(data_double,thetainit_d,paraFitCell,shared_d,0.1,tempGauss1);
end
% [P,model,err] = MLE_FitAbberation_Final_FixPara_Regu_sigma2(data,thetainit,paraFit,shared,0.1,tempGauss1);
model_all_avg_err1= model-data;
RRSE.RRSE_all_avg1= norm(model_all_avg_err1(:),2) / norm(data(:),2);
MAPE.MAPE_all_avg1 = mean(sum(abs(model_all_avg_err1),[1,2])./sum(data,[1,2]))*100;
aberrations_avg1 =  paraFit.aberrations;
aberrations_avg1(:,3)=P(1:21);
% toc 

paraSim=paraFit;
paraSim.aberrations(:,3)=P(1:21);

imx(cat(1,data,model,data-model),'Parent',axzernike);

PupilSize = 1.0;
Npupil = paraFit.Npupil;
DxyPupil = 2*PupilSize/Npupil;
XYPupil = -PupilSize+DxyPupil/2:DxyPupil:PupilSize;
[YPupil,XPupil] = meshgrid(XYPupil,XYPupil);
ApertureMask = double((XPupil.^2+YPupil.^2)<1.0);
orders = paraFit.aberrations(:,1:2);
Waberration1 = zeros(size(XPupil));
orders1 = paraFit.aberrations(:,1:2);
zernikecoefs1 = P(1:21,1);
normfac = sqrt(2*(orders(:,1)+1)./(1+double(orders(:,2)==0)));
zernikecoefs1 = normfac.*zernikecoefs1;
allzernikes1 = get_zernikefunctions(orders1,XPupil,YPupil);
for j = 1:numel(zernikecoefs1)
  Waberration1 = Waberration1+zernikecoefs1(j)*squeeze(allzernikes1(j,:,:));  
end

Waberration1 = Waberration1.*ApertureMask;
imagesc(axpupil,Waberration1);
axis(axpupil,'equal');
axis(axpupil,'tight');
xlabel(axpupil,'X (pixel)');
ylabel(axpupil,'Y (pixel)');
c =colorbar(axpupil);
c.Label.String ='nm';
c.Label.FontSize = 14;

bar(axmode,P(1:21),'FaceColor',[0.6350 0.0780 0.1840],'EdgeColor',[0.6350 0.0780 0.1840]);
xlabel(axmode,'zernike modes');
ylabel(axmode,'zernike rms value (nm)');
% legend(axmode,{'Average bead stacks'});
for k=size(orders1):-1:1
    axn{k}=[num2str(orders1(k,1)) ',' num2str(orders1(k,2))];
end
axmode.XTick=1:length(axn);
axmode.XTickLabel=axn;
axmode.XTickLabelRotation=45;

PSFzernike=model;
end
```
