## Supplementary Software for "Field dependent deep learning enables high-throughput whole-cell 3D super-resolution imaging": ZernikeMode.m

```
function [ZernikeModeIndx]=ZernikeMode(zrnikeparam_model,num)
    switch  num
        case 1   % 2, -2
            ZernikeModeIndx = ['Astigmatism y ','(',num2str(zrnikeparam_model(num,:)),')'];
        case 2   %2, 2
            
            ZernikeModeIndx = ['Astigmatism x ','(',num2str(zrnikeparam_model(num,:)),')'];

        case 3  %3, -1
         
            ZernikeModeIndx =['Coma y ','(',num2str(zrnikeparam_model(num,:)),')'];
   
        case 4  %3, 1
        
            ZernikeModeIndx = ['Coma x ','(',num2str(zrnikeparam_model(num,:)),')'];
 
        case 5  %4, 0
          
            ZernikeModeIndx = ['Spherical ','(',num2str(zrnikeparam_model(num,:)),')'];

        case 6 %3, -3
           
            ZernikeModeIndx =['Trefoil y  ','(',num2str(zrnikeparam_model(num,:)),')'];
          
         case 7   %3, 3
        
            ZernikeModeIndx =['Trefoil x ','(',num2str(zrnikeparam_model(num,:)),')'];

        case 8   %4, -2
           
            ZernikeModeIndx =['2nd Astigmatism y ','(',num2str(zrnikeparam_model(num,:)),')'];
  
        case 9   %4, 2
            
            ZernikeModeIndx =['2nd Astigmatism x ','(',num2str(zrnikeparam_model(num,:)),')'];
   
        case 10  %5, -1
           
            ZernikeModeIndx =['2nd coma y ','(',num2str(zrnikeparam_model(num,:)),')'];

        case 11  %5, 1
            
            ZernikeModeIndx =['2nd coma x ','(',num2str(zrnikeparam_model(num,:)),')'];
  
        case 12   %6, 0
 
            ZernikeModeIndx =['2nd Spherical ','(',num2str(zrnikeparam_model(num,:)),')'];
      
        case 13  %4, -4

           ZernikeModeIndx =['Tetrafoil y ','(',num2str(zrnikeparam_model(num,:)),')'];

        case 14  %4, 4
      
            ZernikeModeIndx =['Tetrafoil x ','(',num2str(zrnikeparam_model(num,:)),')'];

        case 15  %5, -3
  
            ZernikeModeIndx =['2nd Trefoil y  ','(',num2str(zrnikeparam_model(num,:)),')'];

        case 16   %5, 3
        
            ZernikeModeIndx =['2nd Trefoil x  ','(',num2str(zrnikeparam_model(num,:)),')'];

        case 17   %6, -2
        
           ZernikeModeIndx =['3nd Astigmatism y ','(',num2str(zrnikeparam_model(num,:)),')'];

        case 18   %6, 2
         
            ZernikeModeIndx =['3nd Astigmatism x ','(',num2str(zrnikeparam_model(num,:)),')'];

         case 19  %7, 1

            ZernikeModeIndx =['3nd coma x ','(',num2str(zrnikeparam_model(num,:)),')'];

        case 20   %7, -1
  
         ZernikeModeIndx = ['3nd coma y ','(',num2str(zrnikeparam_model(num,:)),')'];

        case 21 %8, 0

            ZernikeModeIndx = ['3nd Spherical ','(',num2str(zrnikeparam_model(num,:)),')'];
            
        case 22
            
            ZernikeModeIndx = 'sigam x';
            
        case 23
            
            ZernikeModeIndx = 'sigam y';
            
    end

    end
```
