## Supplementary Software for "Field dependent deep learning enables high-throughput whole-cell 3D super-resolution imaging": README.md

# FD-DeepLoc
Field dependent deep learning enables high-throughput whole-cell 3D super-resolution imaging
## Abstract
Single-molecule localization microscopy (SMLM) in a typical wide-field setup has been widely used for investigating sub-cellular structures with super resolution. However, field-dependent aberrations restrict the field of view (FOV) to only few tens of micrometers. Here, we present a deep learning method for precise localization of spatially variant point emitters (FD-DeepLoc) over a large FOV covering the full chip of a modern sCMOS camera. Using a graphic processing unit (GPU) based vectorial PSF fitter, we can fast and accurately fit the spatially variant point spread function (PSF) of a high numerical aperture (NA) objective in the entire FOV. Combined with deformable mirror based optimal PSF engineering, we demonstrate high-accuracy 3D SMLM over a volume of ~180 × 180 × 5 μm3, allowing us to image mitochondria and nuclear pore complex in the entire cells in a single imaging cycle without hardware scanning - a 100-fold increase in throughput compared to the state-of-the-art.

![traverse_large_FOV_NPC_ast3_compressed2](https://user-images.githubusercontent.com/67769465/174561057-1745a2c5-fe0c-416c-ada7-a6f2e1902207.gif)
![Supplementary Movie 5 large FOV DOF NPC(edfig 9)](https://user-images.githubusercontent.com/67769465/168954401-9c4d006f-9431-433c-9d74-d28011dc8146.gif)
![Supplementary Movie 3 large FOV DOF mito(fig5)](https://user-images.githubusercontent.com/67769465/168954626-3c10257f-6f4b-49d4-aa70-6c608f609b18.gif)
![singletom20compressed](https://user-images.githubusercontent.com/67769465/168968411-34e482a1-2241-48d4-be09-48f3d43612c9.gif)


#  Requirements
* FD-DeepLoc was tested on a workstation equipped with Windows 10 system, 128 GB of memory, an Intel(R) Core(TM) i9-11900K, 3.50GHz CPU, and an NVidia GeForce RTX 3080 GPU with 10 GB of video memory. To use FD-DeepLoc yourself, a computer with CPU memory ≥ 32GB and GPU memory ≥ 8GB is recommended since FD-DeepLoc would process large-FOV SMLM images (usually > 500GB).
* CUDA Driver (>=11.3, https://www.nvidia.com/Download/index.aspx?lang=en-us) is required for fast GPU-based PSF simulation, PSF fitting and PyTorch.
* For field-dependent aberration map calibration, we tested on Matlab 2021b with CUDA Driver 11.3 on a Windows 10 system.
* The deep learning part of FD-DeepLoc is based on Python and Pytorch. We recommend conda (https://anaconda.org) to manage the environment and provide a `fd_deeploc_env.yaml` file under the folder `FD-DeepLoc/Field Dependent PSF Learning` to build the conda environment.

# Installation
1. Download this repository (or clone it using git).
2. Open Anaconda Prompt and change the current directory to `FD-DeepLoc/Field Dependent PSF Learning`.
3. Use the command `conda env create -f fd_deeploc_env.yaml` to build the FD-DeepLoc environment, it may take several minutes.
4. Activate the environment using the command `conda activate fd_deeploc`, then check the demo using the command `jupyter notebook`.
5. The example bead stacks files for field-dependent aberration map calibration and example data for network inference can be found in the below section Demo examples.

# How to run
The notebook file `.ipynb` itself contains detailed instruction. And a general tutorial file `FD-DeepLoc tutorial.pdf` is also provided under the main directory, which illustrates the procedures for 
1. Field-dependent aberration map calibration.
2. Field-dependent deep-learning localization network (including training and inference examples)

# Demo examples
There are 4 different demo notebooks in the folder `Field Dependent PSF Learning\demo_notebooks` to illustrate the 
use of FD-DeepLoc. To run one demo, the user needs to download the corresponding test dataset using the link below and uncompress it under the 
folder `demo_datasets`. For each demo example, we provide a training notebook `train.ipynp`, an inference notebook `inference.ipynb`, 
test datasets `.tif`, aberration maps `aber_map.mat` and trained models `FD-DeepLoc.pkl`. One can run the training notebook to train a network from scratch or
 just run the inference notebook using provided trained models. For demo2 and demo3, we also provide the raw bead stacks files for field-dependent aberration map calibration. All network predictions will be saved in a `.csv` file in the format of molecule list.  We recommend to use the [SMAP](https://www.nature.com/articles/s41592-020-0938-1) to postprocess the molecule list, such as drift correction, grouping, filtering, and rendering, etc.

1. `demo1` trains 2 networks based on the simulated large-FOV field-dependent aberrated dataset (the normal aberration 
and medium SNR dataset in fig.3). One network utilized all features of FD-DeepLoc while the other one didn't use 
CoodConv and Cross Entropy. Both of them are trained without temporal context (the 3 consecutive frames input) 
for the purpose of CRLB test. This demo aims to show the superority of FD-DeepLoc over a conventional CNN in 
spatially-variant fitting case. This demo takes about 9 hours to train 2 networks and 30 minutes to do the field-dependent CRLB test. 
The test dataset can be downloaded from [![image](https://user-images.githubusercontent.com/67769465/201120098-5c305182-afdf-4dbe-be81-5d28808389ef.png)
](https://zenodo.org/record/7310921#.Y20MaGlBz30).


2. `demo2` trains a network based on our experimental large-FOV astigmatism NPC dataset (fig.4). The corresponding 
test dataset contains two cropped sub-regions of the entire FOV with different field positions. It should be noted that 
the predictions of this dataset need drift correction for better view. This demo takes about 5 hours to train
and tens of minutes to predict. The test dataset can be downloaded from [![image](https://user-images.githubusercontent.com/67769465/200457392-4b2b081c-f9a8-4839-9528-d3fed01fa74d.png)](https://zenodo.org/record/7299483#.Y2m51GlBz30). The raw bead stacks files can be downloaded from [![image](https://user-images.githubusercontent.com/67769465/200461666-9e0a7a7e-e02b-4d87-bdef-5cb240f552a5.png)
](https://zenodo.org/record/7302064).


3. `demo3` illustrates the common using pipeline of FD-DeepLoc on a large FOV and DOF with field-dependent aberrations. 
It is based on our experimental DMO-Tetrapod PSF (3μm) neuron dataset (fig.6). The dataset 
contains the first 10,000 raw frames of the entire FOV. This demo takes about 5 hours to train a network and 2 hours to 
predict. The test dataset can be downloaded from [![image](https://user-images.githubusercontent.com/67769465/200457498-a08ccde4-d3cb-4827-963b-6ff58905ae66.png)
](https://zenodo.org/record/7299064#.Y2m6CGlBz30). The raw bead stacks files can be downloaded from [![image](https://user-images.githubusercontent.com/67769465/200459917-b9eb3157-4f8a-446f-959e-c8d45e1629f4.png)](https://zenodo.org/record/7300138#.Y2m7lWlBz30).


4. `demo4` illustrates the common using pipeline of FD-DeepLoc on a FOV without field-dependent aberrations 
(aberration maps are uniform). It is based on our experimental DMO-SaddlePoint PSF (1.2μm) NPC dataset
(supplementary fig.5). The CoordConv is turned off as it will not learn any extra information from 
the spatially-invariant training data. This demo takes about 5 hours to train a network and 10 minutes to predict. The test dataset can be downloaded from
[![image](https://user-images.githubusercontent.com/67769465/200457603-cc37c13e-109e-4289-981e-fd0e172fd83f.png)](https://zenodo.org/record/7299197#.Y2m6Q2lBz30).


# Contact
For any questions / comments about this software, please contact [Li Lab](https://faculty.sustech.edu.cn/liym2019/en/).

# Copyright
Copyright (c) 2022 Li Lab, Southern University of Science and Technology, Shenzhen.
